# Concurrent loss of ciliary genes *WDR93* and *CFAP46* in phylogenetically distant birds

**DOI:** 10.1101/2023.05.22.541847

**Authors:** Salve Buddhabhushan Girish, Amia Miriam Kurian, Nagarjun Vijay

## Abstract

The respiratory system is the primary route of infection for many contagious pathogens. Mucociliary clearance of inhaled pathogens is an important innate defense mechanism sustained by the rhythmic movement of epithelial cilia. To counter host defenses, viral pathogens target epithelial cells and cilia. For instance, the influenza virus modulates the expression of *WDR93*, a central ciliary apparatus C1d projection component. Lineage-specific prevalence of such host defense genes results in differential susceptibility. Our comparative analysis of ∼500 vertebrate genomes supports the widespread conservation of *WDR93*. However, we identified loss of the *WDR93* in landfowl, geese, and other phylogenetically-independent bird species. Notably, species with *WDR93* loss have concurrently lost another C1d component, *CFAP46*, through large segmental deletions. Understanding the consequences of such gene loss may provide insight into their role in host-pathogen interactions and benefit global pathogen surveillance efforts by prioritizing species missing host defense genes and identifying putative zoonotic reservoirs.

## Introduction

The respiratory epithelium is the first line of defense that acts as a physicochemical barrier through antimicrobial host defense molecules and actively performs mucociliary clearance of inhaled pathogens [1–3]. The epithelium of the airway mostly consists of ciliated, goblet, and basal cells, along with several other rarer cell types such as club cells, deuterosomal cells, ionocytes, neuroendocrine cells, and tuft cells [4]. Adjacent epithelial cells fuse to form cell-cell junctions, which regulate the movement of molecules and physically impede viral entry into the sub-mucosa [5,6]. Additionally, pattern recognition receptors (PRRs) in the epithelium initiate the innate immune response against pathogens [7,8]. Hence, the airway epithelium provides multiple host defense factors against diverse pathogens [9–11].

Several viral pathogens, such as influenza, rhinovirus, metapneumovirus, coronavirus (including SARS-CoV-2), respiratory syncytial virus (RSV), and some bacterial pathogens, primarily infect the host through the respiratory route. Respiratory pathogens cause ciliary impairments such as ciliated epithelial cell shedding, ciliary motility disruption, loss of cilia, other types of ciliary dysfunction, and ciliogenesis suppression [12–16]. Viruses can enter host cells through various sialic acid receptors of airway epithelium cells [17,18]. Variation in sialic acid receptors has been linked to viral tropism, and a better understanding of their prevalence and tissue-specificity across diverse species may help understand zoonotic virus spillover [19–23]. For instance, a wider range of sialic acid receptor subtypes in chickens compared to ducks may cause infections from a broader range of avian influenza viruses [24]. Adaptation to differing pathogen regimes through the species-specific presence of sialic acids occurs through changes in the gene repertoire of sialic acid-converting enzymes [25,26]. Hence, gene presence/absence patterns in diverse vertebrate species could be associated with differences in the immune response [27–30]. Moreover, identifying natural knockouts of crucial genes can help uncover functional roles that could translate into biomedical insights and anticipate zoonotic events [31].

A transcriptomic study of the differential host response to high or low pathogenic H5N1 avian influenza virus in ducks has identified *WDR93* (also known as *CFAP297*) to be down-regulated in high and up-regulated in low pathogenic virus infection [32]. The *WDR93* gene has been predicted to be part of mitochondrial respiratory chain complex I due to its sequence homology with the *NDUFS4* gene. However, recent studies have conclusively established *WDR93* is part of the central apparatus projection D (C1d) along with *CFAP54*, *CFAP46* (also known as *TTC40*), *CFAP221* (also known as *PCDP1*), and *CAM1* [33–39]. Electron microscopy and gene-knockout studies have established the importance of the C1d projection in maintaining ciliary motility [34,40,41]. Single-cell transcriptomics of diverse non-model vertebrate species, including duck, has identified *WDR93* expressed in the ciliated cells of the lungs [42]. Hence, based on earlier literature, we reasoned that the *WDR93* gene expression change in response to H5N1 avian influenza in ducks might be a consequence of the virus targeting the ciliary apparatus [13]. Genome-wide RNA interference studies have also implicated *WDR93* in altering the percentage of cells infected with *M. tuberculosis* [14,43]. The interaction between the host gene *WDR93* with multiple pathogens hints at the possibility of lineage-specific gene losses that could alter host-pathogen interaction. We perform an in-depth comparative genomic analysis of the *WDR93* gene across diverse vertebrate lineages to evaluate gene presence/absence patterns. Our search of ∼500 vertebrate genomes revealed the independent loss of *WDR93* in several lineages of birds in contrast to its widespread conservation across other vertebrate species. Interestingly, *CFAP46*, another C1d gene directly interacting with *WDR93*, is concurrently lost in multiple phylogenetically independent bird lineages through segmental deletions.

## Materials and Methods

### Identification of conserved one-to-one orthologs in vertebrates

We performed the comparative genomic analysis across ∼500 vertebrate species with good-quality genome assemblies to identify one-to-one orthologs of the *WDR93* gene (see **Supplementary Table 1**). The gene annotated as *WDR93* in the human genome was used as a reference to identify one-to-one orthologs in 23 representative species from various vertebrate clades based on synteny and sequence similarity (see **Supplementary Table 2**). To overcome the heterogeneous annotation quality in different species, we re-annotated the *WDR93* gene using the gene sequence of the representative species in each clade when the NCBI annotation was incomplete or labeled as a low-quality protein. See **Supplementary Text/S1** for details of annotation curation and re-annotation. The re-annotation was done by running the TOGA (Tools to infer orthologs from genome alignment) pipeline [44] on good-quality genomes. Subsequently, the assembly sequences were verified and rectified in case of errors using Illumina short-read sequencing (see **Supplementary Table 1**).

### Genome assembly validation using long-read sequencing

We validated the genome assembly using long-read sequencing to ensure the micro-synteny of the focal gene/remnants and its flanking regions. High-quality long-read datasets (PacBio and Nanopore) are available for chicken (see **Supplementary Table 3**). We used the BWA read aligner to map the long-read data to the genome of the chicken version GRCg6a (Genome Reference Consortium Chicken Build 6a). Subsequently, we retained PacBio reads >20Kb, and Nanopore reads > 80Kb, spanning the genomic region containing the remnants of the focal gene along with flanking genes on both sides. The long-reads span the flanking genes covering the remnants of genomic features (exon, intron, and UTR) of the *WDR93* gene. However, for the *CFAP46* gene, exonic remnants were not found at the syntenic locus despite conserved gene order on both sides. Consistent lack of genomic regions homologous to the *CFAP46* locus in various versions of the chicken genome assemblies (GRCg6a (RJF), GRCg7b (maternal broiler), GCA_024206055.2 (Huxu breed)), long-read (PacBio and Nanopore), and short-read (Illumina) datasets suggests the loss of *CFAP46* due to a large segmental deletion. See **Supplementary Text/S2** for details of segmental deletion validation and estimating the size of the region lost.

### Reconstructing the history of gene loss events

The high-quality chicken genome and long-read sequencing datasets allowed for detailed validation of gene loss. However, other Galloanseriform species lack similar-quality of genomic data. Hence, reconstructing the chronology of events relies upon genome assemblies and their confirmation using short-read Illumina sequencing. First, we detected the gene-disrupting events in each species using the TOGA pipeline with Ostrich (*Struthio camelus*) as a reference species. Subsequently, we verified the events using blastn with mallard (*Anas platyrhynchos*) and white-crested guan (*Penelope pileata*) as query species (see **Supplementary Table 4**). Gene-disrupting events such as frameshift insertion/deletion, substitution leading to stop codon, exon deletion, and splice-site change were detected by TOGA. Each event was confirmed based on a blastn search of the short-read Illumina data (see **Supplementary Table 5**) and visualized using the MView utility.

### Comparison of the transcriptional level at *WDR93* gene locus

The *WDR93* gene is highly expressed in the gonads [45] and cerebellum tissues. Hence, we screened the gonadal transcriptomes of the Chinese alligator (*Alligator sinensis)*, green anole *(Anolis carolinesis*), Ostrict *(Struthio camelus*: cerebellum tissue RNA-seq data used instead of low-quality gonad data), Emu (*Dromaius novaehollandiae*), Zebra Finch (*Taenipygia guttata*), Great-tit (*Parus major*), Mallard (*Anas platyrhynchos*), Swan goose *(Anser cygnoides), Muscovy duck (Cairina moschata)*, Chicken *(Gallus gallus)*, Turkey (*Meleagris gallopava*), and common pheasant (*Phasianus colchicus*) to compare transcription at the *WDR93* gene locus. The RNA-seq reads from public datasets (see **Supplementary Table 6**) were mapped to the respective species reference genomes using STAR read mapper [46]. The flanking genes (*MINAR1* and *PEX11A*) are positive controls within the species, and the same tissue was considered between species. Furthermore, to verify the possibility of *WDR93* gene expression in other chicken tissues or sex-specific gene expression, we screened 67 RNA-seq datasets from various tissues (see **Supplementary Table 6**). The transcriptional status was assessed visually using the Integrative Genomics Viewer (IGV) browser [47]. Additionally, to overcome the restrictions of window size in IGV screenshots, the coverage of the *WDR93* gene locus and its flanking genes was calculated using the bedtools [48] genomecov command with and without-split option and saved as a bedgraph file and visualized along with the exon locations and flanking genes.

### Estimating selection strength and GC-biased gene conversion

We generated multiple sequence alignments of *WDR93* ORFs for each clade using Guidance (v2.02) with PRANK as the aligner [49]. Each clade was assessed for sequence saturation using the Index to measure substitution saturation (Iss) proposed by Xia et al. [50] and available in the DAMBE program. Time-calibrated phylogenetic species trees were downloaded for each clade from the TimeTree website [51]. We estimated selection strength in each species using RELAX program from the HyPhy package [52] and codeML from the PAML package [53] (see **Supplementary Table 7-8**). We also quantified the magnitude of gBGC using the mapNH program from testNH(v1.3.0) and phastBias from PHAST(v1.6) (see **Supplementary Table 9**). See our earlier study [54] for detailed methods.

## Results

### Conservation of micro-synteny at *WDR93* locus

Comparison of gene order in the flanking regions of the *WDR93* gene in 23 representative species of major clades identified two main types of micro-synteny (see **Fig. 1**). All therian genomes analyzed, including humans, contain *PEX11A, PLIN1*, and *KIF7* on the left flank, with *MESP1*, *MESP2*, and *ANPEP* genes on the right flank of *WDR93* (see **Supplementary Table 2**). Among monotremes, while the gene order on the left flank was the same as in other mammals, the gene order on the right flank consists of *AKAP13*, *SV2B*, and *SLCO3A1* genes in the platypus (*Ornithorhynchus anatinus*) and *NTRK3*, *AGBL1*, *AKAP13*, *SV2B*, and *SLCO3A1* genes in the short-beaked echidna (*Tachyglossus aculeatus*) (see **Supplementary Table 2**). The mammal-like gene order occurs in the Western clawed frog (*Xenopus tropicalis*) and Coelacanth (*Latimeria chalumnae*). In the case of sauropsids (including Aves), the analyzed genomes had the same left flank genes as mammals, with *MINAR1*, *MTHFS*, and *BCL2A1* being on the right flank of *WDR93*. The conserved synteny at the *WDR93* locus ensures unambiguous identification of 1-to-1 orthologs. Although *WDR93* shares the IPR006885 (NADH_UbQ_FeS_4_mit-like) domain with its only documented paralog *NDUFS4*, sequence clustering analysis identified high levels of divergence between the paralogs (see **Supplementary Figure 1**).

**Fig. 1.**
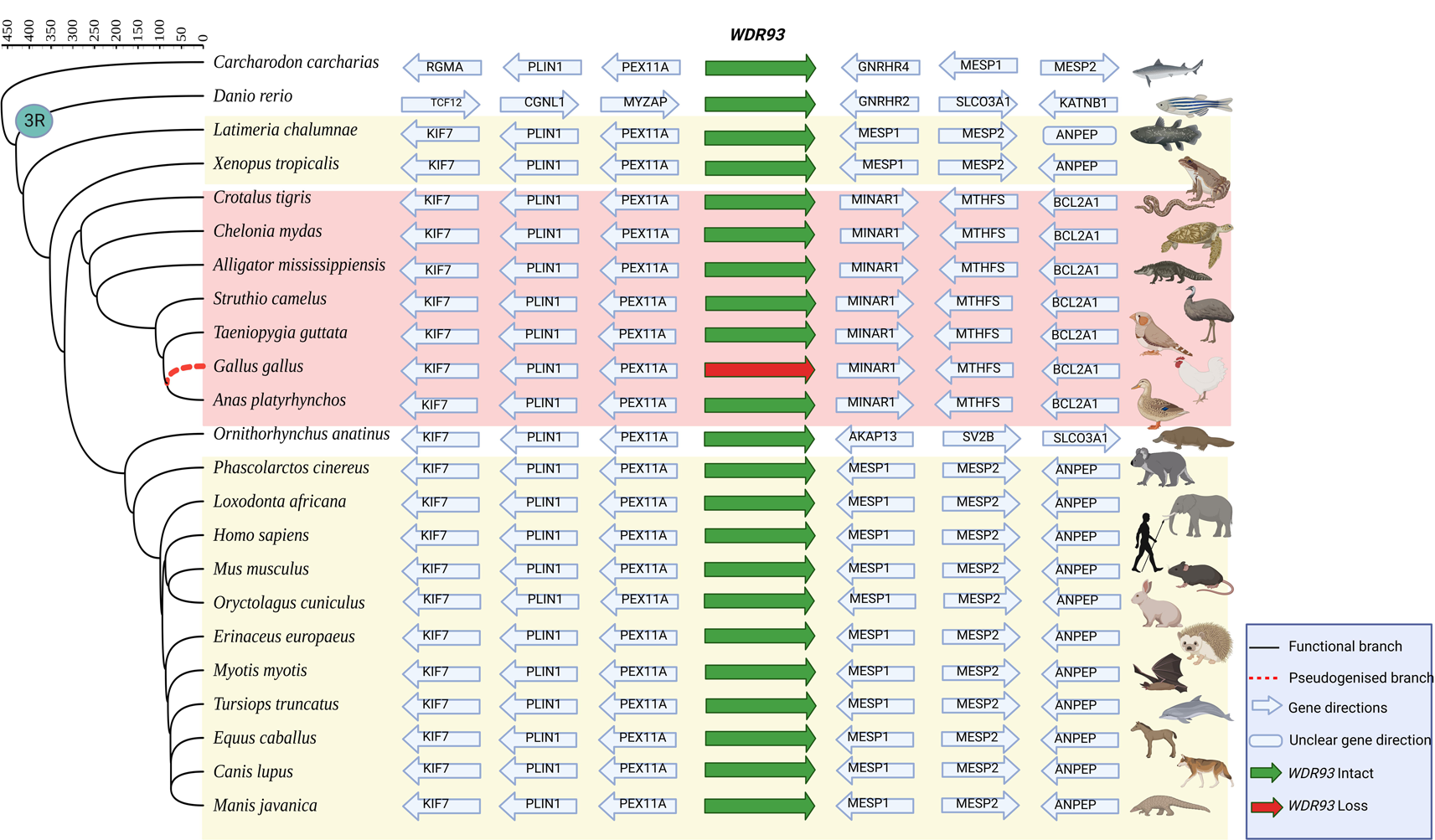
Gene order conservation in regions flanking *WDR93* in representative vertebrate species. The arrowheads indicate the transcriptional orientation of the genes relative to the *WDR93* gene for consistency across all species. A solid green arrow indicates an intact gene, and a solid red arrow indicates gene loss. In the phylogenetic tree (downloaded from TimeTree), the black-colored branches represent functional branches, whereas the red-colored dashed line indicates a pseudogenised branch. In the species highlighted in yellow, the conserved upstream genes (left flank) are *PEX11A*, *PLIN1*, and *KIF7*, and the downstream genes (right flank) are *MESP1*, *MESP2*, and *ANPEP*. In the species highlighted in red, the upstream genes are *PEX11A*, *PLIN1*, and *KIF7*, and the downstream genes are *MINAR1*, *MTHFS*, and *BCL2A1*. Timescale is millions of years. The image is generated using BioRender.com and integrative Tree of Life (iTOL).

### Chronology of *WDR93* gene inactivation in Galliformes and Geese

None of the galliform genome assemblies contain an annotation for the *WDR93* gene despite being well annotated in the closely related anseriform birds. Using long-read sequencing, we ruled out the possibility of assembly errors at the *WDR93* locus in the chicken genome. Three overlapping Nanopore reads >100 Kb (SRR15421344.63125, SRR15421342.280842, and SRR15421343.497419) span the entire region between *MINAR1* and *PEX11A* containing *WDR93* gene remnants, i.e., 5’ UTR, exons, and intronic regions (see **Fig. 2**). In galliform birds, representative species from the Numididae, Odontophoridae, and Phasianidae families share the deletion of four exons (Exon-4, 11, 13, and 14; see **Fig. 3**). Hence, the *WDR93* gene loss event shared by three of the five galliform families are estimated to have occurred between 65.4 to 46.5 MYA. Although the *WDR93* gene appears intact in the white-crested guan (*Penelope pileata*), a frame-disrupting deletion in Exon-6 of the Australian brushturkey (*Alectura lathami*) suggests an independent loss (see **Fig. 3** and **Supplementary Figure 2**). The representative species from the Odontophoridae family shared the deletion of Exon-5, 10, 12, and 15. The partial degradation of Exon-9 and Exon-12 was found to be shared across several galliform species. The high-quality chicken genome and long-read sequencing allowed the validation of the partially degraded exons.

**Fig. 2.**
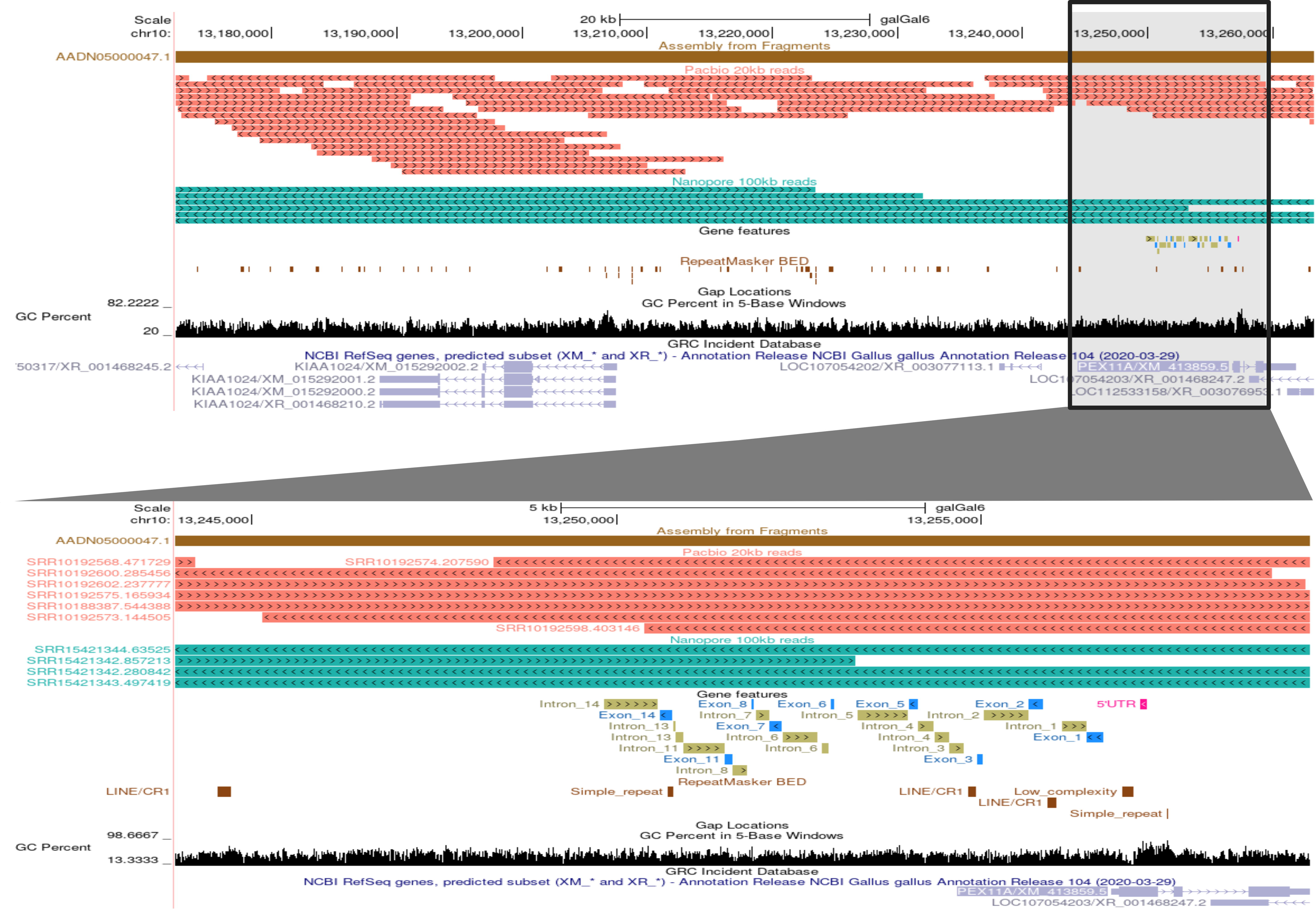
Validation of the chicken genome assembly (galGal6) using long reads. Screenshot of UCSC genome browser with the alignment of ≥20 kb long reads generated using PacBio sequencing technology (salmon colored) and ≥100 kb long reads generated using Nanopore sequencing technology (sea green colored) at the syntenic location of the *WDR93* gene remnants. The inset at the bottom is the zoomed-in view of the region highlighted (focusing on the *WDR93* remnants). Blue-colored boxes represent the *WDR93* exon remnants, olive-green-colored boxes are the *WDR93* introns, and the pink box represents the 5’ UTR (shown in the genomic features track). The RepeatMasker track (shown as dark brown boxes) depicts the repeats in that region. The accession numbers of sequencing reads are on the left beside the alignments. Overlapping reads span the entire region at the syntenic location of the *WDR93* gene, including the flanking genes (*KIAA1024/MINAR1* and *PEX11A*).

**Fig. 3.**
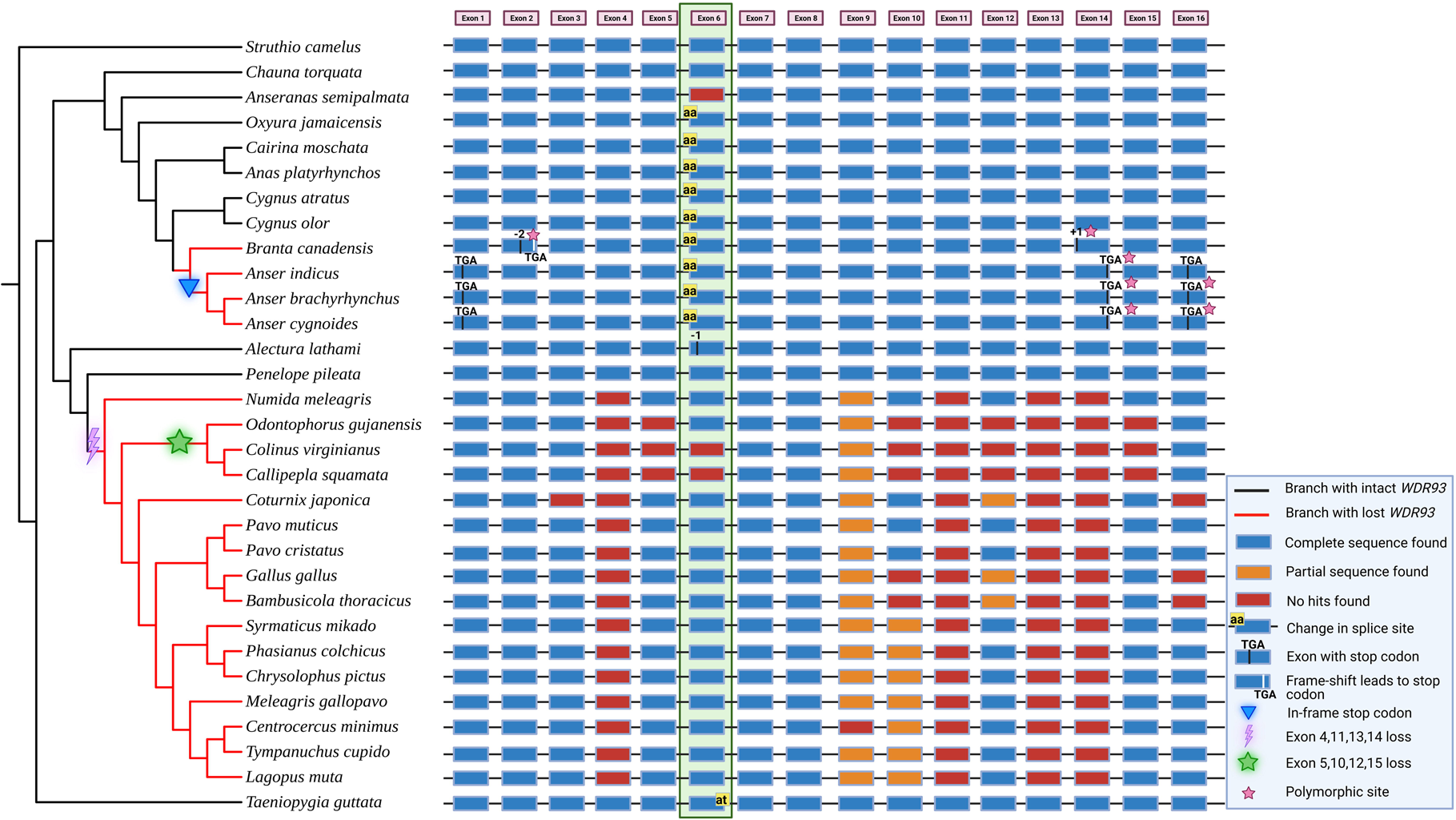
Loss of *WDR93* gene in galliform and geese species. Loss of *WDR93* in sixteen galliform species, four geese species, and their phylogenetic relationship help reconstruct the chronology of events. Black and red branches indicate species with functional and pseudogenised *WDR93*, respectively. The purple thunderbolt symbol depicts a shared loss of exons 4, 11, 13, and 14 in sixteen galliform species. The green colored star symbol represents the shared loss of exons 5, 10, 12, and 15. The blue inverted triangle represents the presence of an inframe stop codon in the Anser lineage. The filled rectangular boxes represent exons, and the lines connecting them represent the introns. The blue-colored boxes represent the intact exon sequences, the orange-colored boxes represent the exons with partial sequences, and the red-colored boxes represent the exons with no detectable remnants. The region highlighted with green represents exon 6, which has a change in the splice site (at the start of exon 6) in eight of the anseriform species. The pink star symbols indicate that the represented events are polymorphic.

Among anseriform birds, the crested screamer (*Chauna torquata*) from the Anhimidae family has an intact *WDR93* gene. In the magpie goose (*Anseranas semipalmata*), the sixth exon is missing from the genome assembly. Since short-read data supports the validity of the magpie goose genome assembly (see **Supplementary Figure 3**), it appears that the sixth exon is lost without causing any frameshift. All the representative species from the Anatidae family whose genomes were analyzed share an AG➔AA 5’ splice-site disrupting change at Exon-6 (see **Fig. 3** and **Supplementary Table 10**). Independent GT➔AT 3’ splice-site disrupting changes at Exon-6 also occur in Neoaves species such as zebra finch (*Taeniopygia guttata*). Hence, Exon-6 appears dispensable, and its loss does not result in gene loss. Duck and swan species such as the ruddy duck (*Oxyura jamaicensis*), Muscovy duck (*Cairina moschata*), mallard (*Anas platyrhynchos*), black swan (*Cygnus atratus*), and mute swan (*Cygnus olor*) have intact open reading frames after excluding Exon-6. However, geese species from the genera Anser and Branta have gene-disrupting changes. The bar-headed goose (*Anser indicus*), pink-footed goose (*Anser brachyrhynchus*), and swan goose (*Anser cygnoides*) share a C to T substitution at the 69^th^ base of Exon-1, which leads to a premature stop codon (CGA➔ TGA). Additionally, the three species from the Anser genus share a premature stop codon (CGA➔ TGA) inducing polymorphism at the 154^th^ base of Exon-14, and another polymorphic site (CAT➔TGA) occurs at the 62-64^th^ position of Exon-16 (see **Fig. 4**). Genome assembly of swan goose (*Anser cygnoides*) was verified using PacBio long-read sequencing data (see **Supplementary Figure 4**). Similarly, two gene-disrupting polymorphisms, a two-base deletion (in Exon-2) and one base insertion (in Exon-14), occur in the Canada goose (*Branta canadensis*), resulting in a premature stop codon in Exon-2 (**Supplementary Figure 5**).

**Fig. 4.**
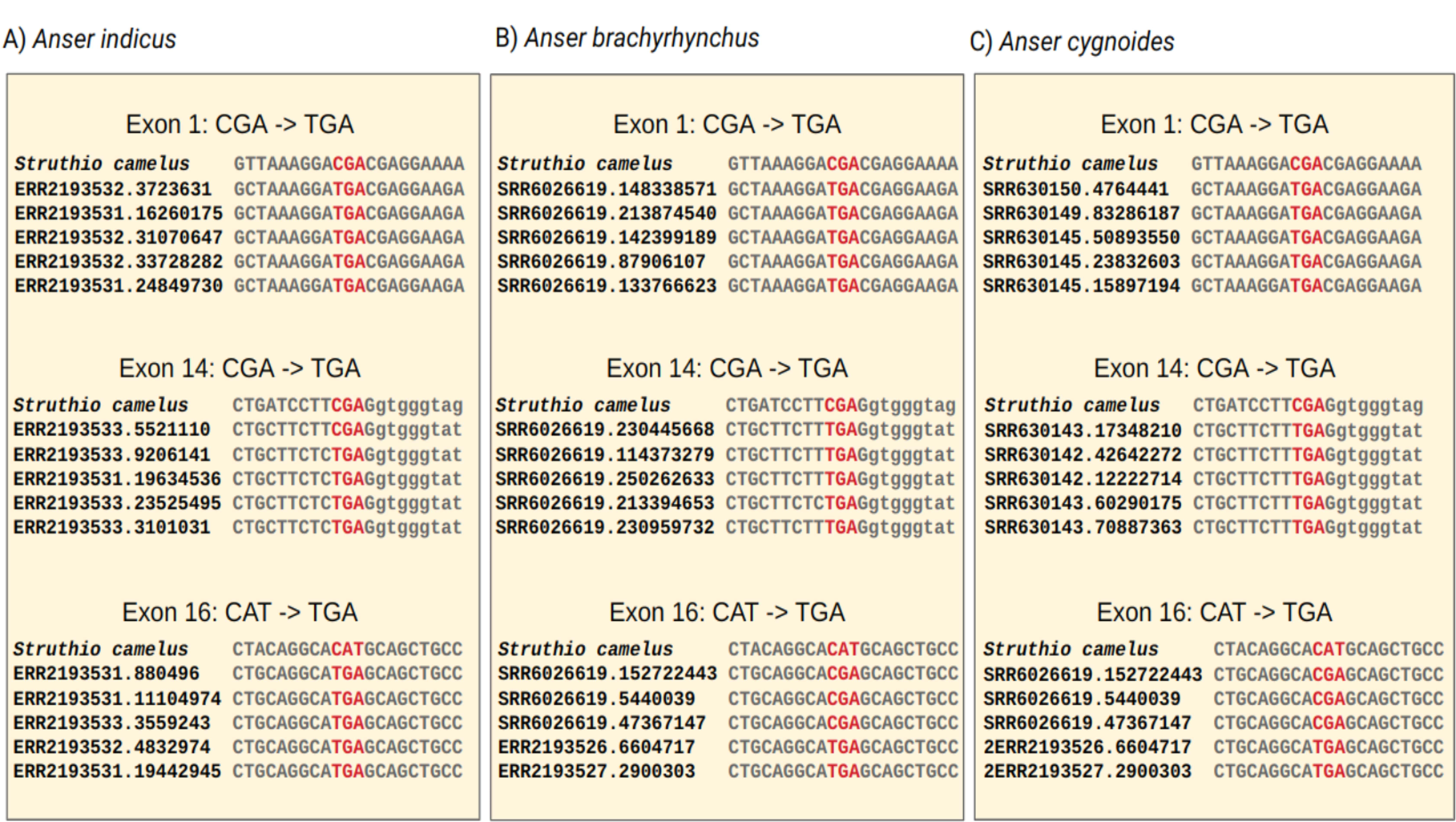
Short read support for gene disrupting changes in the *WDR93* exon sequences of geese species. In exons one and fourteen, the CGA is mutated to TGA, and in exon sixteen, CAT is mutated to TGA, leading to an inframe stop codons (shown by red text) in the three geese species: **(A)** Bar-headed goose (*Anser indicus*), **(B)** Pink-footed goose (*Anser brachyrhynchus*) and **(C)** Swan goose (*Anser cygnoides*). These results are identified based on a comparison with the sequence of the *WDR93* gene from Ostrich (*Struthio camelus*). A small case letter shows the intronic sequence beside the exon-14.

### Contrasting transcriptional patterns at the *WDR93* locus

The annotation of the *WDR93* gene consists of sixteen exons in most vertebrate species. Examination of the transcriptomes confirmed that all sixteen exons are expressed in Chinese alligator (*Alligator sinensis*), Green anole (*Anolis carolinensis*), ostrich (*Struthio camelus*), and Emu (*Dromaius novaehollandiae*) (see **Fig. 5-6** and **Supplementary Figure 6-13**). However, as noted in the previous section, some bird species have splice site-disrupting changes in Exon-6. Accordingly, only 15 exons are expressed in mallard (*Anas platyrhynchos*), Muscovy duck (*Cairina moschata*), Great tit (*Parus major*), and Zebra finch (*Taeniopygia guttata*) (**Supplementary Fig. 14-21**). Therefore, the RNA-seq data confirm the dispensability of Exon-6 and its exclusion from the transcript in species with splice site-disrupting changes. The *WDR93* gene is prominently expressed in the gonads and cerebellum (see **Fig. 5**-**6** and **Supplementary Figure 22**). Galliform species with gene-inactivating mutations, such as chicken (*Gallus gallus*), turkey (*Meleagris gallopava*), helmeted guineafowl *(Numida meleagris)*, common pheasant (*Phasianus colchicus*), Indian peafowl (*Pavo cristatus*), and Japanese quail (*Coturnix japonica*) lack transcript read support at the *WDR93* locus in the gonad (see **Supplementary Figure 23-26**). However, at the same time, *WDR93* is abundantly expressed in the gonad of anseriform species such as mallards (*Anas platyrhynchos*) and Muscovy ducks (*Cairina moschata*). Additional screening of 67 RNA-seq datasets of various chicken tissues from male and female individuals failed to identify transcripts at the *WDR93* locus. Thus, several lines of evidence support the loss of the *WDR93* gene in chickens. Although the swan goose (*Anser cygnoides*) has gene-disrupting changes, we found RNA-seq reads mapped to *WDR93* in the gonad sample (see **Fig. 5**). Upon closer inspection of the *WDR93* locus in the swan goose, the transcript support was found to be noisy and concordance of spliced reads with the exon boundaries was weak (see **Supplementary Figure 22**).

**Fig. 5.**
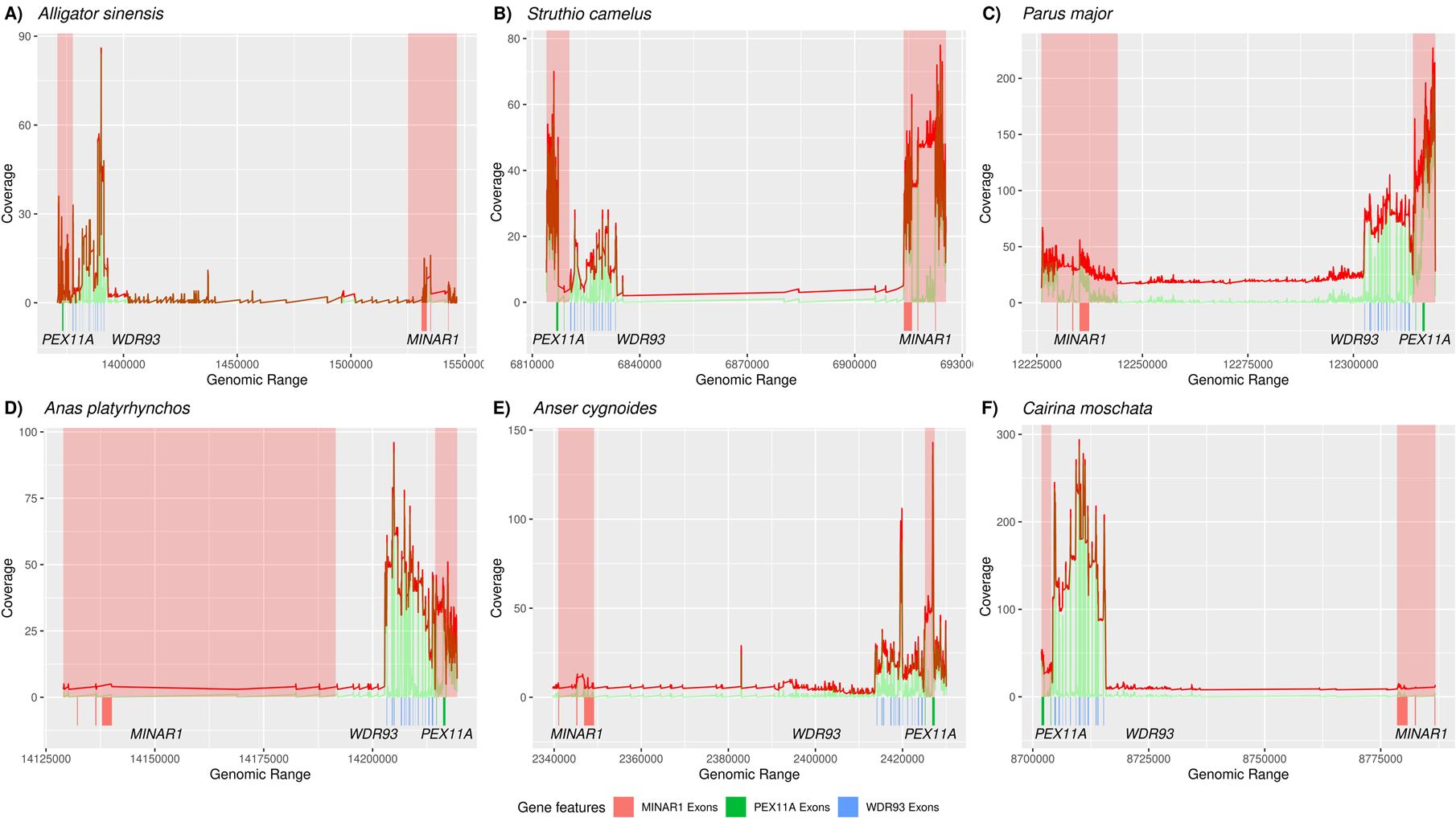
Transcriptional status of *WDR93* in Anseriformes. Read coverage of RNA-seq data from NCBI Short Read Archive (SRA) database mapped to its corresponding species genome using STAR mapper. **(A)** Chinese alligator (*Alligator sinensis*), **(B)** Common ostrich (*Struthio camelus*), **(C)** Great tit (*Parus major*), **(D)** Mallard (*Anas platyrhynchos*), **(E)** Swan goose (*Anser cygnoides*), **(F)** Muscovy duck (*Cairina moschata*). The genes adjoining *WDR93*, i.e., *PEX11A* and *MINAR1*, are highlighted with a red background as a positive control. The red line in the graph shows the total RNA-seq coverage (without the split option in bedtools), and the green line indicates the spliced alignment coverage (with split). The blast hits of *MINAR1*, *PEX11A*, and *WDR93* exons to the genome of corresponding species are shown in red, green, and blue, respectively. RNA-seq short read data is from the testis for all species except for using cerebellum in ostrich.

**Fig. 6.**
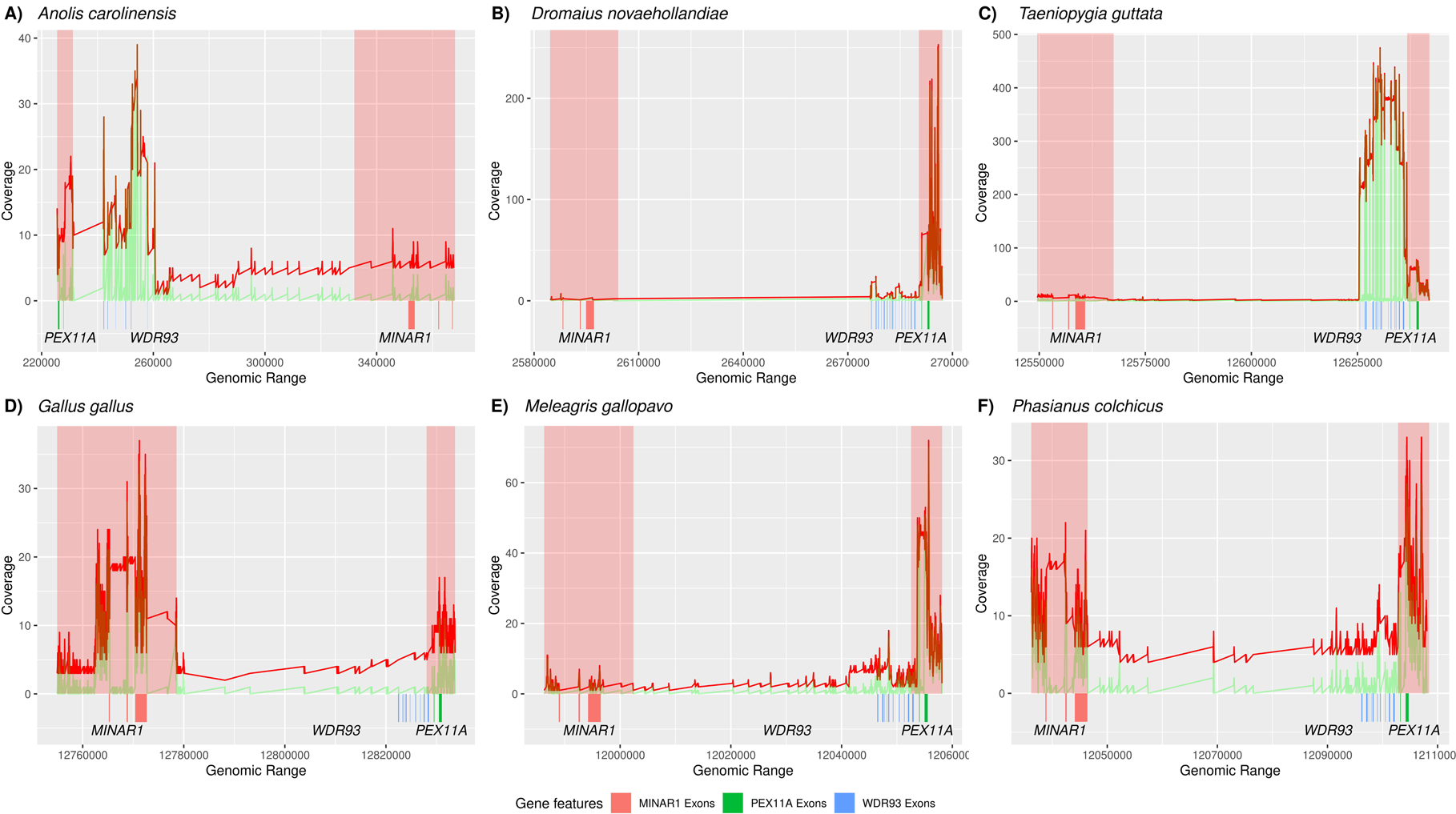
Transcriptional status of *WDR93* in Galliformes. Read coverage of RNA-seq data from NCBI Short Read Archive (SRA) database mapped to its corresponding species genome using STAR mapper. **(A)** Green anole (*Anolis carolinensis*), (**B)** Emu (*Dromaius novaehollandiae*), **(C)** Zebra finch (*Taeniopygia guttata*), **(D)** Chicken (*Gallus gallus*), **(E)** Wild turkey (*Meleagris gallopavo*), **(F)** Ring-necked Pheasant (*Phasianus colchicus*). The genes adjoining *WDR93*, i.e., *PEX11A* and *MINAR1*, are highlighted with a red background as a positive control. The red line in the graph shows the total RNA-seq coverage (without the split option in bedtools), and the green line indicates the spliced alignment coverage (with split). The blast hits of *MINAR1*, *PEX11A*, and *WDR93* exons to the genome of corresponding species are shown in red, green, and blue, respectively. RNA-seq short read data is from the testis for all species.

### Independent losses of *WDR93* in other Neognathae species

The *WDR93* locus genome assembly verification and annotation step (see **Methods**) helped identify putative gene loss events and rectify errors. We could verify four promising examples of independent losses of *WDR93* in Neognathae (see **Fig. 7**). **(1)** The fifth exon of *WDR93* in the rifleman (*Acanthisitta chloris*) bird has a G to T nucleotide substitution at its 57^th^ base, leading to an in-frame stop codon (GAG➔TAG). Short-read sequencing supports the validity of this substitution (see **Fig. 7**). Exon 7 and exons 10 to 16 are all missing from the rifleman genome assembly and short-read data. The validity of the genome assembly at the *WDR93* locus was ascertained using long-read sequencing and confirms the loss of 8 of the 16 exons (see **Supplementary Figure 27**). **(2)** In the ruff (*Calidris pugnax*), Exon-1 at the 103^rd^ base and Exon-15 at the 9^th^ base have inframe stop codons due to CGA➔TGA and AAG➔TAG substitutions, respectively. Short-read data support the validity of these stop codon-inducing substitutions (see **Fig. 7**). **(3)** Anna’s hummingbird (*Calypte anna*) has one frameshifting deletion and another polymorphic frameshifting insertion in Exon-2 (see **Supplementary Figure 28**). Exon-4 and exons 7 to 16 are deleted in Anna’s hummingbird, and the correctness of the genome assembly could be verified by a tiling path of long-read data (see **Supplementary Figure 29**). **(4)** In speckled mousebird (*Colius striatus*), the *WDR93* gene has four inframe stop codons, three single base insertions (in Exon-2, 9, and 14), and a four-base insertion in exon-10. Two exons (6 & 11) are missing from the speckled mousebird genome assembly, and short-read data suggest their loss.

**Fig. 7.**
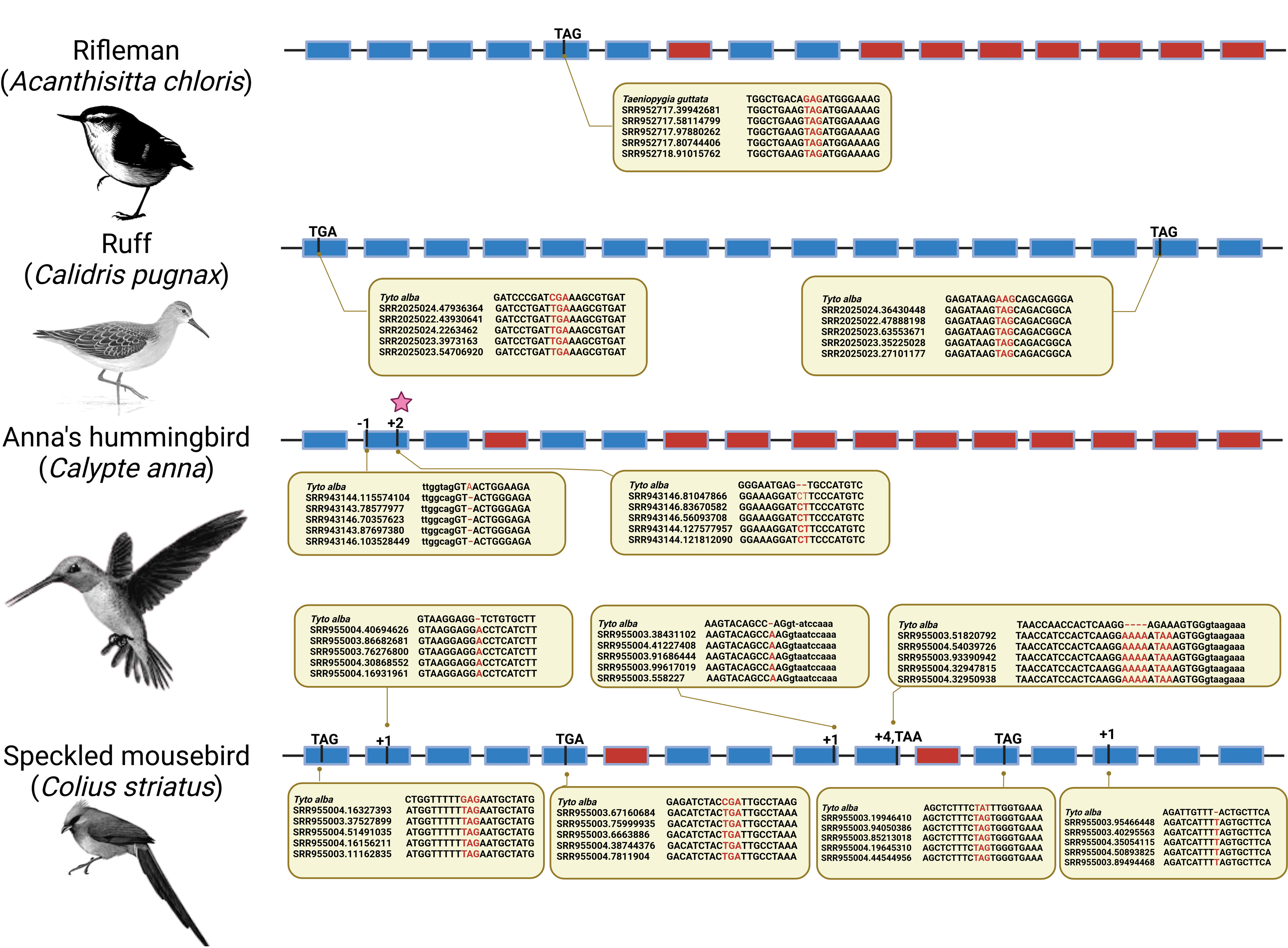
Independent loss of *WDR93* gene in Neoaves. The *WDR93* gene is lost independently in the rifleman (*Acanthisitta chloris*), ruff (*Calidris pugnax*), Anna’s hummingbird (*Calypte anna*), and speckled mousebird (*Colius striatus*). The filled rectangular boxes represent exons, and the lines connecting them represent the introns. The blue-colored ones represent the exon sequences, and the red-colored boxes represent the exons with no remnants. The pseudogenization events are shown on top of exons with black lines and events at the corresponding locations. The inset at events, i.e., in the comment box, depict the raw read support for that event visualized using Mview. The bird images are adapted from photos available in the public domain.

### Concurrent loss of *CFAP46* through large segmental deletions

The C1d projection consists of proteins encoded by *WDR93* and other genes such as *CFAP221*, *CAM1*, *CFAP54*, and *CFAP46*. A search for orthologs of these genes revealed that the *CFAP46* gene-containing region is conserved in representative bird species, such as ostrich (*Struthio camelus*), zebra finch (*Taeniopygia guttata*), tufted duck (*Aythya fuligula*), black swan (*Cygnus atratus*), mute swan (*Cygnus olor*) and ruddy duck (*Oxyura jamaicensis*) (see **Fig. 8A-D**). The *CFAP46* gene-containing region is missing in the chicken genome assembly. The micro-synteny of the *CFAP46* gene is conserved across birds, with *ALDH18A1* and *TCTN3* on the left flank and *OPNVA*, *NKX6-2*, and *INPP5A* genes on the right flank (see **Supplementary Table 11**). Based on pairwise genome alignment between mallard (*Anas platyrhynchos*) and chicken (*Gallus gallus*), we inferred a chicken-specific deletion containing the *CFAP46* gene. The *CFAP46* deletion is shared by other galliform species, such as turkey (*Numida meleagris*), common pheasant (*Phasianus colchicus*), and Japanese quail (*Coturnix japonica*) (see **Fig. 8E-H**). We approximated the size of segmental deletion in various galliform species (see **Supplementary Table 12**) to range from ∼81Kb to ∼73Kb. We confirmed the correctness of the genome assembly using PacBio long-read sequencing in chicken (*Gallus gallus*) and mallard (*Anas platyrhynchos*) (see **Supplementary Figure 30-31**). Hence, the large segmental deletion identified in chicken is not an assembly error.

**Fig. 8.**
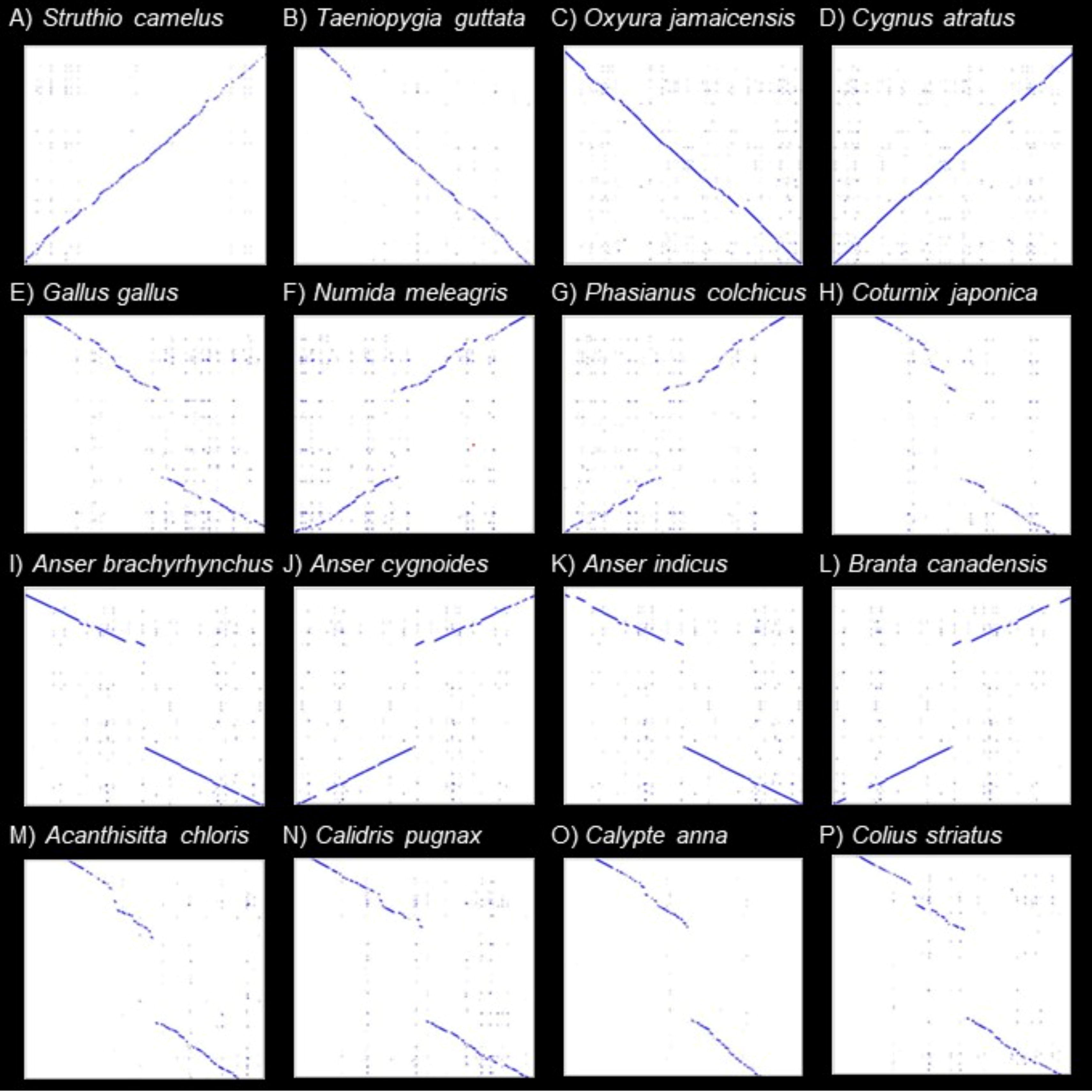
Concurrent loss of *CFAP46* gene with *WDR93*. A blast-based search of genomes of representative species (shown on the x-axis of each dot plot) using the entire *CFAP46* genic region, along with the 50 Kb flanking sequence of the mallard (*Anas platyrhynchos*) as a query (shown on the y-axis of each dotplot), was used to generate dot plots using the xsltproc utility from XSLT sandbox (https://github.com/lindenb/xslt-sandbox). Conserved orthologous regions with no segmental deletions are found in **(A)** Ostrich (*Struthio camelus*), **(B)** Zebra finch (*Taeniopygia guttata*), **(C)** ruddy duck (*Oxyura jamaicensis*), and **(D)** black swan (*Cygnus atratus*). However, alignment patterns consistent with a segmental deletion were found in the galliform lineage in species such as **(E)** chicken (*Gallus gallus*), **(F)** helmeted guineafowl (*Numida meleagris*), **(G)** common pheasant (*Phasianus colchicus*), and **(H)** Japanese quail (*Coturnix japonica*). Similarly, in geese lineage, another segmental deletion is common to species such as **(I)** pink-footed goose (*Anser brachyrhynchus*), **(J)** swan goose (*Anser cygnoides*), **(K)** bar-headed goose (*Anser indicus*), and **(L)** Canada goose (*Branta canadensis*). Other phylogenetically independent bird species, namely **(M)** rifleman (*Acanthisitta chloris*), **(N)** ruff (*Calidris pugnax*), **(O)** Anna’s hummingbird (*Calypte anna*), and **(P)** speckled mousebird (*Colius striatus*) have lost the *CFAP46* gene through segmental deletions.

We found an independent segmental deletion of *CFAP46* in the Canada goose (*Branta canadensis*), a species in which the *WDR93* gene has gene-inactivating polymorphisms. The segmental deletion found in the Canada goose (*Branta canadensis*) also occurs in the pink-footed goose (*Anser brachyrhynchus*), swan goose (*Anser cygnoides*), and bar-headed goose (*Anser indicus*) (see **Fig. 8I-L**). While the boundaries of the segmental deletion occur in the same region as chicken, they are not identical (see **Supplementary Figure 32-33**). For instance, the size of the geese-specific deletion was ∼92Kb (see **Supplementary Table 12**). To ensure this region is not prone to assembly errors, we verified the genome assembly of swan goose (*Anser cygnoides*) with PacBio long-read sequencing (see **Supplementary Figure 34**). Since we found two cases of concurrent loss of *WDR93* and *CFAP46*, the genomes of other bird species that have lost the *WDR93* gene were searched for *CFAP46* orthologs. Interestingly, the *CFAP46*-containing region of the genome appears to be lost through independent segmental deletions in the four species with independent *WDR93* gene loss, i.e., rifleman (*Acanthisitta chloris*), ruff (*Calidris pugnax*), Anna’s hummingbird (*Calypte anna*), and speckled mousebird (*Colius striatus*) (see **Fig. 8M-P**). The genome assemblies of the rifleman (*Acanthisitta chloris*) and Anna’s hummingbird (*Calypte anna*) could be verified using PacBio long-read sequencing (see **Supplementary Figure 35-36**). Consistent support for the deleted regions in long-read datasets, the prevalence in independent genome assemblies of closely related species (see **Fig. 8E-L**), and the lack of short-read support for the deleted region strongly support the recurrent deletion of this *CFAP46*-containing region without translocation within the somatic genome.

## Discussion

The central ciliary apparatus of motile cilia comprises an asymmetric pair of microtubules, C1 and C2, with interconnected distinct protein complexes known as projections [38,55]. The overall structure of the central apparatus is largely conserved despite minor species-specific differences [56,57]. The study of mutants lacking specific ciliary apparatus components suggests different roles for each projection in ensuring motility [37]. The epithelial cilia in the respiratory tract perform mucociliary clearance of pathogens as a host defense strategy [58]. For instance, mice enhance ciliary activity and mucociliary clearance when infected with the influenza A virus [59]. However, pathogens can counter almost every host defense by suppressing and evading the host immune machinery [60,61]. Tactics of the pathogens, such as downregulating ciliogenesis regulatory genes (Robinot et al., 2021), targeting ciliated epithelial cells [62], and hijacking cilia to cross the nasal epithelium barrier [63] highlight the vital role of cilia in host-pathogen interactions. The central apparatus projection D deficiency results in severe motility defects and appears to be an ideal target for pathogen-mediated disruption of cilia [34,41,64]. Changes in the expression of the *WDR93* gene upon influenza virus infection strongly indicate that respiratory pathogens may target projection D components [32]. Our discovery of the recurrent and concurrent loss of two C1D components (*WDR93* and *CFAP46*) in birds further supports the role of these proteins in host-pathogen interaction.

The concurrent loss of *WDR93* and *CFAP46* in multiple independent lineages of birds, in contrast to conservation across hundreds of other vertebrate species, suggests rapid shifts in selective forces acting on these genes. Loss of genes with host defense functions may result from relaxed selection following a change in pathogen repertoire or invasion strategy [65–67]. The ongoing arms race between host and pathogen involves constantly evolving new and complex mechanisms to counter each other and can also cause gene loss [68–70]. The disruption of *WDR93* and *CFAP46* genes could also be a casualty of this arms race. For instance, the host may have an adaptive advantage in suppressing or losing these genes if pathogens hijack the cilia to counter the immune response or overcome host defenses [12]. The sustained expression of *RIG-I*, interferon-stimulated genes, and genes from pathways involved in ciliary functions are explicitly documented in nasal epithelial cells in response to viral RNA persistence [71]. Loss of genes could also be a consequence of such persistent activation of the immune system due to increased contact with certain pathogens. Our findings indicate a hitherto unknown role for the central apparatus projection D components in host-pathogen interaction due to their involvement in ciliary motility. Hence, understanding the impact of *WDR93* and *CFAP46* gene presence/absence across diverse species in modulating the response to invading pathogens is of immunological significance. Modifying ciliary function by targeting conditionally dispensable genes such as *WDR93* and *CFAP46* could also be a promising strategy for developing new antiviral therapies.

Comparative studies of non-traditional animal models have great potential in understanding host-specific immune responses and developing new approaches to counter zoonotic pathogens [72–74]. However, restricted access to wild animals and the non-availability of molecular biology tools for non-model species have been major hurdles to experimentally studying exotic pathogens. Due to seasonal variations in microbial load, many “missing viruses” and “unknown zoonoses” are yet to be discovered and can act as potential sources of infection in the future [75,76]. Hence, novel approaches for identifying potential reservoir species, intermediate hosts, and susceptible species are required [77,78]. The availability of high-quality genomic datasets has enabled large-scale comparative genomic studies that will reveal genetic differences associated with combating various pathogens and help identify potential reservoirs and species at zoonotic risk [27,79,80]. Recent developments such as telomere-to-telomere genome assemblies and transgenics have made the chicken an amenable model to compare with the mammalian immune system [81,82]. Differences between the natural immune system vs. the transmission or spillover host could help better understand the host-pathogen interactions [83–85].

The bird species in which the *WDR93* and *CFAP46* genes are lost chiefly belong to the superorder Galloanserae. This gene loss trend can be partly explained by the fact that anseriform birds are the primary maintenance hosts for influenza A viruses and may have adapted to the pathogen load through gene loss [86,87]. However, galliform birds appear to have a more evolutionarily widespread loss than anseriform species. While this fits the general pattern of chicken-specific loss of several immune genes, the reason remains debatable [28,88,89]. Recent studies investigating the transcriptional response to viral infection in birds reveal a complex scenario of alternative pathways that can compensate for lost genes [29,90–93]. The gene loss in the ruff (*Calidris pugnax*) is consistent with the Charadriiformes order being the second most important maintenance host after anseriform birds [86,94–97]. Although the role of non-Anseriformes and non-Charadriiformes (NANC) species in acting as a reservoir for the influenza virus is difficult to evaluate using current monitoring practices, gamebirds have received considerable attention [98–101]. Nonetheless, two more species with gene loss, the Anna’s Hummingbird (*Calypte anna*) and the rifleman (*Acanthisitta chloris*), can be infected by the avian poxvirus and the zoonotic bacterium *Chlamydia psittaci*, respectively [102–105]. The rifleman is non-migratory and endemic to New Zealand but may show severity upon pathogenic infection spread by migratory birds. The spread of the virus to other wild birds can threaten endangered species and complicate pathogen containment efforts. The loss of *WDR93* and *CFAP46* in speckled mousebird (*Colius striatus*), a species reportedly missing the *MDA5* gene [106], may indicate a different host-pathogen interaction dynamic. Identifying NANC species lacking vital immune genes could help prioritize monitoring efforts and develop more sophisticated zoonotic potential evaluation strategies.

Lineage-specific loss of widely conserved genes resulting in phenotypic consequences serves as a natural experiment to understand the function of genes. Our work provides an example of how gene presence/absence patterns can identify novel candidates, even among understudied proteins involved in processes that are experimentally difficult to test. The approach outlined here also provides a framework for identifying other genes of potential relevance to host-pathogen interaction. Genome-wide extension of our study will help us understand the genomic factors involved in zoonotic spillover and find species that can act as potential reservoirs and species at zoonotic risk. We hope this approach will help in developing new strategies for disease management in the future.

## Acknowledgments

We thank the Council of Scientific & Industrial Research for a fellowship to SBG. The Department of Biotechnology, Ministry of Science and Technology, India (Grant no. BT/11/IYBA/2018/03) and Science and Engineering Research Board (Grant no. ECR/2017/001430) provided funds for procuring computational resources (i.e., Har Gobind Khorana Computational Biology cluster) used.

## Authors contributions

S.B.G. wrote the manuscript with inputs from A.M.K and N.V. S.B.G. analyzed all the data and also supervised A.M.K in validating some of the analysis. S.B.G. wrote several scripts to automate important steps of the analysis. All authors reviewed the manuscript.

## Data accessibility

The associated data is available in an easy-to-view format on the GitHub repository: https://github.com/CEGLAB-Buddhabhushan/WDR93_CFAP46.

