## Supplementary File S1 for "Concurrent loss of ciliary genes *WDR93* and *CFAP46* in phylogenetically distant birds"

```
Anas_platyrhynchos_platyrhynchos 1 TATCAGTGGGTGGGTATTAATGTTTTTCAGCAAAATCAG-----C 45
Gallus_gallus                    1 CTGTCCCACCTGGGTATTAATGCTTATCAACAAAATTAGGAAAAA 45

Anas_platyrhynchos_platyrhynchos 46 AAGTAAATATTGTACCTTCTGTTCAATTCAAAATTAAAATCTCCT 90
Gallus_gallus                    46 AAAAAAATAGTGTACACTCTGTTCAATTCAAGCTTAAGATTGT-- 90

Anas_platyrhynchos_platyrhynchos 91 GTTCTGTCTACCTGGATTACCTCCTGGCTATGCTAATTCTTAGTT 135
Gallus_gallus                    91 -TTCTCTCTGCCCAGCTTACCTCCTGGCTACGC----TCTTAGTT 135

Anas_platyrhynchos_platyrhynchos 136 ACAGAGATAAAGCATGTCTCTAATTGCCTCCCTGCCTTGCTCCCT 180
```

```
Anas_platyrhynchos_platyrhynchos 946 TGTAAATCTTCCACAACCACTTTGAGGAGTGAAGGCCATTGCTGT 990
Gallus_gallus                    946 ............................................. 990

Anas_platyrhynchos_platyrhynchos 991 CAGTTGATAGAAGCATCTTGAGATTGTCTGTCACATGCTTAAAAT
1035
Gallus_gallus                    991 .............................................
1035

Anas_platyrhynchos_platyrhynchos 1036 CAGCTGAGTGAACCTATTCAAGGGCACCACTTGAAGTATCACACT
1080
Gallus_gallus                    1036 .............................................
1080

Anas_platyrhynchos_platyrhynchos 1081 GACCTTATCGCTCATTTCCATGTCTGGGTCCTGGAATGGGAAATG
1125
Gallus_gallus                    1081 .............................................
1125

Anas_platyrhynchos_platyrhynchos 1126 CAGGACTAATGCACAAATTGCCACATACAATTTCAGACACCTGCT
1170
Gallus_gallus                    1126 .............................................
1170

Anas_platyrhynchos_platyrhynchos 1171 ATGGTCTCAGCACAGTTCTTGTCATCCTGATTTTGTCATATAGCT
1215
Gallus_gallus                    1171 .............................................
1215

Anas_platyrhynchos_platyrhynchos 1216 GGAATTACTGCAGGCACATTACACAAGTGCTATTACTTACACTTT
1260
Gallus_gallus                    1216 .............................................
1260

Anas_platyrhynchos_platyrhynchos 1261 CTCCTCACCTCACTGACTGTATGTTCCTGCTCCAATCAACTGCTG
1305
Gallus_gallus                    1261 .............................................
1305

Anas_platyrhynchos_platyrhynchos 1306 TTTATACAACAGCCTGAGGTTGTTCTGTTGTCTGCCAGCTTACCT
1350
Gallus_gallus                    1306 .............................................
1350

Anas_platyrhynchos_platyrhynchos 1351 CTTAGTAAGTGTCAGTCAGGAACACCTTTGCTAAAAAGGCGATGT
1395
Gallus_gallus                    1351 .............................................
1395

Anas_platyrhynchos_platyrhynchos 1396 GAGATATGCCAAGCTCTCTCAAGATAGGTTTGCACGTGCTCCAAG
1440
Gallus_gallus                    1396 .............................................
1440

Anas_platyrhynchos_platyrhynchos 1441 TTTATCAGTGAACTTTCAGTATATGAAAGTCCAGGAAAGGGAAGT
1485
Gallus_gallus                    1441 .............................................
1485

Anas_platyrhynchos_platyrhynchos 1486 TGTCACTAATGTAAACCAAGATCTTTATAAGAGGGACTGAAGATC
1530
Gallus_gallus                    1486 .............................................
1530
```

```
Anas_platyrhynchos_platyrhynchos 1531 TTGTGTTATAATCTTATCAACAAGTACTAATATCTGAAGGGAGGG
1575
Gallus_gallus                    1531 .............................................
1575

Anas_platyrhynchos_platyrhynchos 1576 CGTTAAGAGGATGGGGCCGGACTTTTTTCAGTAGTGCCCAGCAAC
1620
Gallus_gallus                    1576 .............................................
1620

Anas_platyrhynchos_platyrhynchos 1621 AGGACAAGAGGCAATGGGCACAGACTGAAACACAGGAAGTAGTGG
1665
Gallus_gallus                    1621 .............................................
1665

Anas_platyrhynchos_platyrhynchos 1666 TTGAATATGAGAAAACATTTATTTACTGTAAGGGTGACAGAATGC
1710
Gallus_gallus                    1666 .............................................
1710

Anas_platyrhynchos_platyrhynchos 1711 TGGCACAGGTTGCCCAGAGAGGTTATGGAGTCTCCTCCTTTGGAG
1755
Gallus_gallus                    1711 .............................................
1755

Anas_platyrhynchos_platyrhynchos 1756 TTATTCAAAACTCACCTGGACAAGATCCAGGGCCATGTGTGCTAG
1800
Gallus_gallus                    1756 .............................................
1800

Anas_platyrhynchos_platyrhynchos 1801 GTGACCCTGCTTGGCAGAAGGGTTGGACCAGGTGTTCTCCAGAGA
1845
Gallus_gallus                    1801 .............................................
1845

Anas_platyrhynchos_platyrhynchos 1846 TCCCTCCCAACCTCAAGAATTCTGTGATTCTGCTGGAGAACAGCT
1890
Gallus_gallus                    1846 .............................................
1890

Anas_platyrhynchos_platyrhynchos 1891 GTCTCTTTAAATGGAGGCAATTGACCTAGGATGATCCTGGTCTCC
1935
Gallus_gallus                    1891 .............................................
1935

Anas_platyrhynchos_platyrhynchos 1936 CTGTGTTCACAAGGCAATACTTTGTCTTTCTAAATTTCTGATTTC
1980
Gallus_gallus                    1936 .............................................
1980

Anas_platyrhynchos_platyrhynchos 1981 AGAAAGTCTCACAACAAAATTGCAAAAACTTCTGTATTATTTGGA
2025
Gallus_gallus                    1981 .............................................
2025

Anas_platyrhynchos_platyrhynchos 2026 CAAACAAAGGCCTTCAGTGCAAGAGCAGTAGCAAATTCAAACCTA
2070
Gallus_gallus                    2026 .............................................
2070

Anas_platyrhynchos_platyrhynchos 2071 TTGTGCTATAAAAAGTGGAAATGCTGTAAGTTGTAATAAGAGAGC
2115
Gallus_gallus                    2071 .............................................
```

```
2115

Anas_platyrhynchos_platyrhynchos 2116 ATGTCACAGAGAATCACTTTCAGTTGATCTGTTGAAGAAGCAAAA
2160
Gallus_gallus                    2116 .............................................
2160

Anas_platyrhynchos_platyrhynchos 2161 CAAAAGCAAACTGGGCAGCTGAGTATAAGAATATTTTTAAAGGAT
2205
Gallus_gallus                    2161 .............................................
2205

Anas_platyrhynchos_platyrhynchos 2206 TCTAAGCAGAGGACTCACAAGACAAAGACAATGCATTTAAACTGT
2250
Gallus_gallus                    2206 .............................................
2250

Anas_platyrhynchos_platyrhynchos 2251 AATCAGAATCATTACTCATGCAAGCAACTTGAACAATATCACAGC
2295
Gallus_gallus                    2251 .............................................
2295

Anas_platyrhynchos_platyrhynchos 2296 ATAAAACAACAAAATAACTGGAAACAACAGTTATCCATTCTTTAA
2340
Gallus_gallus                    2296 .............................................
2340

Anas_platyrhynchos_platyrhynchos 2341 GGGGTGTCAATGAACAGGCAATGAATTTGTGTTAGAGAAATCCAG
2385
Gallus_gallus                    2341 .............................................
2385

Anas_platyrhynchos_platyrhynchos 2386 ATAATGCCTTCTCTGTTATATTTTATTTCAGAAATCTAATTTCAT
2430
Gallus_gallus                    2386 .............................................
2430

Anas_platyrhynchos_platyrhynchos 2431 TTTAACGCCCACAAGAATTTAACGTGTCATTTTAAGCAACATAAT
2475
Gallus_gallus                    2431 .............................................
2475

Anas_platyrhynchos_platyrhynchos 2476 ATTTGAACTTGATTTTAAACTACTGGTCCAGATTTTCTGTTGTAG
2520
Gallus_gallus                    2476 .............................................
2520

Anas_platyrhynchos_platyrhynchos 2521 TATTCAGGCAATGGAGCTATGATGAATCATGCCATACAAGGATCT
2565
Gallus_gallus                    2521 .............................................
2565

Anas_platyrhynchos_platyrhynchos 2566 GCTGTCGGCATTTTAATGTCACTGTGTTTTGATCTTTCTTTCTTT
2610
Gallus_gallus                    2566 .............................................
2610

Anas_platyrhynchos_platyrhynchos 2611 TGATGCTTGGAACTTTTTAATACTTCTGTTGAATAAGCATTATTT
2655
Gallus_gallus                    2611 .............................................
2655

Anas_platyrhynchos_platyrhynchos 2656 TGTGAGTATTTTTGTTACTAGATTTGTTCTGCTTAAGCTCTGTCT
```

```
2700
Gallus_gallus                     2656 .............................................
2700

Anas_platyrhynchos_platyrhynchos 2701 TTCCCTTTCAAGTGAGTATGTTAAAGCTATTTTAAAATATCAAAA
2745
Gallus_gallus                     2701 .............................................
2745

Anas_platyrhynchos_platyrhynchos 2746 AGTTATTTTAATTATAAGCTGCTGTCATTTTTTGTTGTTTGAAAT
2790
Gallus_gallus                     2746 .............................................
2790

Anas_platyrhynchos_platyrhynchos 2791 AGATTGAAATCTTCTGATGTATAACTCTTTCTGCCTGTGCCCAGA
2835
Gallus_gallus                     2791 .............................................
2835

Anas_platyrhynchos_platyrhynchos 2836 AGCTCTTTTGATTCCATTCATGCATATCAAAAGATTTCAACCTAT
2880
Gallus_gallus                     2836 .............................................
2880

Anas_platyrhynchos_platyrhynchos 2881 GTTAAGTGTTTCACATACAAACTAAAACAAATATAAACTCAGAGG
2925
Gallus_gallus                     2881 .............................................
2925

Anas_platyrhynchos_platyrhynchos 2926 AATATATGGATAGGTAGCATTGCTCGTTCTTACTTTCAGTTCAGC
2970
Gallus_gallus                     2926 .............................................
2970

Anas_platyrhynchos_platyrhynchos 2971 TCCTGTTTTTTTCTTTGTGTGTTGCGCATGGTGCATGTCCAAATT
3015
Gallus_gallus                     2971 .............................................
3015

Anas_platyrhynchos_platyrhynchos 3016 TTGAAGACTCCAGGATGGAAACTGGATTGCTTATACATAATTGCT
3060
Gallus_gallus                     3016 .............................................
3060

Anas_platyrhynchos_platyrhynchos 3061 AACAATTATTATATATAATATAGATACATCCACTCATTTTTCTCA
3105
Gallus_gallus                     3061 .............................................
3105

Anas_platyrhynchos_platyrhynchos 3106 GAAATATCTGACTAGCAATTAATTTTTCTGTTGTCTATTGTCATG
3150
Gallus_gallus                     3106 .............................................
3150

Anas_platyrhynchos_platyrhynchos 3151 CTGTGCTGATTTTATTTGTTTCAATGTATTTGCAGGCTCTATCTT
3195
Gallus_gallus                     3151 .............................................
3195

Anas_platyrhynchos_platyrhynchos 3196 TAAAATACCACGGTGCATCTTTGAGATTTTAGCACTAAAACTGTG
3240
Gallus_gallus                     3196 .............................................
3240
```

Anas_platyrhynchos_platyrhynchos 3241 ATTATAATAATTATGCCTTCATCAGGGATAACAGCTGTACAGTCT
3285
Gallus_gallus 3241 .............................................
3285

Anas_platyrhynchos_platyrhynchos 3286 CTAGGAGAAACTAGCTGAAATTGCACTAAATAACACAGAAAACTT
3330
Gallus_gallus 3286 .........................CTAAATACCACAGAATGCTA
3330

Anas_platyrhynchos_platyrhynchos 3331 ATCTGAAGCTTCAGTTCTGGGGCCGTAAGCTAACCTGAGATGTGA
3375
Gallus_gallus 3331 ATTTGAAGCTTCATTGCTGGGGCCATGAGTTATCCTGAGACATGA
3375

Anas_platyrhynchos_platyrhynchos 3376 GCAGGGTTTTAGAATCCAGCTTCCCTGCCAAGTTTAGAGCTGCTG
3420
Gallus_gallus 3376 ACAGGGGTT-AGAATGCAGTCTCCCTGCCAAGTTTACAGCTCCTA
3420

Anas_platyrhynchos_platyrhynchos 3421 TTATAGTCCAGTGTCAGTCTTTGCAGGGCAGCCCTTCTTCAAGCA
3465
Gallus_gallus 3421
TGATAGTTCAGCATTAATCTTTTCAGGGCAGCCCTCCTTCA---- 3465

Anas_platyrhynchos_platyrhynchos 3466 AATGAGTCAGGTGGGAGCCTCCCCCTTTGCTGTGCTGGCAGCAGT
3510
Gallus_gallus 3466 -----GTCAGACAGGAGCTTCCACCTCTGCTGTGCTCCCAGCGGT
3510

Anas_platyrhynchos_platyrhynchos 3511 GCAGGGCATGGGGAAGATCACTCCACCCTCATTACTAGCAAAAGG
3555
Gallus_gallus 3511 GCAGA-CCTGGGGCAGATCACTCCACCCTCACTGCTAACAAAAGG
3555

Anas_platyrhynchos_platyrhynchos 3556 AAAGCTTCATTTTCACAGTGCTAGAGATCAGGGAAGGATTCAAGC
3600
Gallus_gallus 3556 AAATCTTTGTTTTCACCGTGCTGGAGGTCAGGAAGTGAT-CAAGC
3600

Anas_platyrhynchos_platyrhynchos 3601 AAATGTTTTGTTTTTC--CCCACAGGGTAGGGATGATCTGAGGGA
3645
Gallus_gallus 3601
AAATGTTCTGTTTTTTTTCCCACAGAATAGGAATGATTTGA---- 3645

Anas_platyrhynchos_platyrhynchos 3646 TATCTGCTCCAATCACTGCTGTTGGCCGGTACAGAAGGAGGGAAC
3690
Gallus_gallus 3646 -------TTTCAGCACTGCTGTTAGCTGCTACAGAAGGAGGGAAC
3690

Anas_platyrhynchos_platyrhynchos 3691 CTGCA---------GCTGGCTT-TATTAGAGAAGAAGACAGTATC
3735
Gallus_gallus 3691 ATGCATCTTGCATAGCTGACTTCTATTACAAAG--ACGCAGCATG
3735

Anas_platyrhynchos_platyrhynchos 3736 AGCGGTAGCTGGAGAG--GGGCAGGAGACTACCAGCAGAGAGGGG
3780
Gallus_gallus 3736 AGCTCCAGCTAGAGAGAGGGGGCAGGAGACTGCCAGCAGAGAAGGG
3780

Anas_platyrhynchos_platyrhynchos 3781 TAGAAGGCTCTCTAATAGA--TTCTCTTGCTTATCCCTGTTCCCA
3825

```
Gallus_gallus                      3781 CAGAAGGCACTCCAAGAGAGATTCCCTTGCATATTCCTCTTCCCA
3825

Anas_platyrhynchos_platyrhynchos 3826 CATGCACTGCACAGTACTAAGCCTTCCTCTGACCAAAACTTTGTT
3870
Gallus_gallus                      3826 CAT--------------------------------------TGTT
3870

Anas_platyrhynchos_platyrhynchos 3871 TCCTGGATTATTACCAGAAAGCGGAAACATGGAGAGTTTTTTTAC
3915
Gallus_gallus                      3871 TCCTGGACTATTACCAGAAAGCAGAAGCATGGAGAGTTAGTTTAT
3915

Anas_platyrhynchos_platyrhynchos 3916 ATTTCTGTTGGAGACAGACAATTTCTCCTTTTAATCCTGAACTTT
3960
Gallus_gallus                      3916 ATTT---------------GGTTTCTCCTTCTAATCTTGAATATT
3960

Anas_platyrhynchos_platyrhynchos 3961 ATTTACAGATTAGAAACTCCAATATTCTCCAATATTATTTTCACT
4005
Gallus_gallus                      3961 CTTTACAGATTTAAAACTTCATTATTATCCAACGTTACTTTTACT
4005

Anas_platyrhynchos_platyrhynchos 4006 AGTGAGAGAAAAAAACGGATGCATTCAGAGCACTTGGAAATGAGT
4050
Gallus_gallus                      4006 AGCAGGGGGGAAAAATGGATGTATTCAGAGCACTTGGAAATGAGT
4050

Anas_platyrhynchos_platyrhynchos 4051 CGCTGTTGTCTGGTTTCTCTGGTCCTGGCAGATGGGATCCCTTTC
4095
Gallus_gallus                      4051 CGCTGTTGTCGAACTCCTCTGGTCCTGCTCGATGGGATCCCTTCC
4095

Anas_platyrhynchos_platyrhynchos 4096 GTCGTCCCTTGGATTCCATCCAGCCCTGGCACTTCAGGCTTCTGG
4140
Gallus_gallus                      4096 ACCATCCCTTGGACTCCATCCAGCCCTGGCACTTCAGGCTTGTGG
4140

Anas_platyrhynchos_platyrhynchos 4141 CAGCAGTAATGCTGGTGGTGACCTCCCTATCGCTTGCTGAGAACC
4185
Gallus_gallus                      4141 CAGCAGTAATGTTCGTGGTGACCTCCCTGTCTCTTGCGGAGAACC
4185

Anas_platyrhynchos_platyrhynchos 4186 TGGCTGTGATCCTGGTAACTTTTAAGTTCAAGCAGTTGAGACAAC
4230
Gallus_gallus                      4186 TGGCTGTGATCCTGGTAACTTTTAAGTTCAAGCAGCTGAGACAAC
4230

Anas_platyrhynchos_platyrhynchos 4231 CTGTCAATTATGTTATAGTCAATTTATCTGTGGCTGATTTTCTGG
4275
Gallus_gallus                      4231 CTGTCAATTATGTTATAGTGAATTTATCTGTGGCTGATTTCCTGG
4275

Anas_platyrhynchos_platyrhynchos 4276 TCTCATTGACTGGTGGCACCATCAGCTTTTTAGCCAATCTAAAAG
4320
Gallus_gallus                      4276 TCTCTTTGACTGGTGGCACCATCAGCTTTTTAGCCAACCTAAAAG
4320

Anas_platyrhynchos_platyrhynchos 4321 GTTATTTTTATATGGGATACTGGGCTTGTGTACTGGAAGGATTTG
4365
Gallus_gallus                      4321 GCTATTTCTATATGGGGCACTGGGCTTGTGTACTGGAAGGATTTG
4365
```

```
Anas_platyrhynchos_platyrhynchos 4366 CGGTCACATTTTTTGGTAAGCTTGATTTTGTTGGTAGTTATACAT 4410
Gallus_gallus                    4366 CTGTCACGTTTTTTGGTAAGCTTGATTTTGTCTGTAGTTGTATCT 4410

Anas_platyrhynchos_platyrhynchos 4411 T-TTTTCTTTGAACTTCCTTTAATGTATTCTACTAACTCTCCTGC 4455
Gallus_gallus                    4411 TGTTTTCCTTCAGCTTCCTTCAGTGTGTTTCACTA----TCCTGC 4455

Anas_platyrhynchos_platyrhynchos 4456 AGCTTCCTGAGCCTCTGACTTTGCATGCCATTGAATGCATTGGAT 4500
Gallus_gallus                    4456 AGCTTCCTAAATCACTGACCTTGCATGCCTGTGAATGCACTGGAT 4500

Anas_platyrhynchos_platyrhynchos 4501 CTTTCCCCAGTTACGTATGTTCTGTGGTGTACCTCATGCATGAGT 4545
Gallus_gallus                    4501 CTTT-TCCAGTTGTGTGTGTTCTGTGGAGGATCTCAT--ATGAGT 4545

Anas_platyrhynchos_platyrhynchos 4546 GAATAACATGCAGCACTTTTGCTGTCATATATCAGTGGCTTGTTG 4590
Gallus_gallus                    4546 GAATAATGTGTATCACTATTGCTGTCATATGTCAGCTTCTTGCTG 4590

Anas_platyrhynchos_platyrhynchos 4591 CTTCTGTACTGTTTACTGAATATGTTTAATATAATG-AGAAAAAA 4635
Gallus_gallus                    4591 CTTCTATACTGCTCTCTGAATATAGTCAATATAATGAAAAAAAAA 4635

Anas_platyrhynchos_platyrhynchos 4636 CTTGCTTATAGAATAGCAGGA-TTGTGGGTAAAACTGGGAGAAAA 4680
Gallus_gallus                    4636 ATTGCTAATAGAATAGGAATAGTTGTGGGGAAAACAATGTGATTA 4680

Anas_platyrhynchos_platyrhynchos 4681 AAAAAATCA----------------CGATGTGCAACTATTTTCAA 4725
Gallus_gallus                    4681 AAACAAACAAACAAACAAACCACTTCACTGTGTAATTATTTTCCA 4725

Anas_platyrhynchos_platyrhynchos 4726 AGTTTGTGTCTAGGTGTGTG--TATGTGGTTAAAT---------- 4770
Gallus_gallus                    4726 GGTTTGTCTCTTGGTGTATGAATATGTGGTTAAGTGAGAGAGAGA 4770

Anas_platyrhynchos_platyrhynchos 4771 ----------GAGAGAGAGAACAAATAGCTGTTCTAAAACTGTGG 4815
Gallus_gallus                    4771 GAGAGAGAGAGAGAGAGAGAACAAATGGCTGTTCTTAAACTATGG 4815

Anas_platyrhynchos_platyrhynchos 4816 AAAATATTAGGGCACTGAAACATCTTATGTTAGCTGTCGTCCAGT 4860
Gallus_gallus                    4816 ATAATGTTATGTCACTGAAACATCTTATGTCAGCTGTCCTGCAAC 4860

Anas_platyrhynchos_platyrhynchos 4861 GGTTCTGAGGAACTATGACATGCCTGTAACTGTCACCCCGTGCAC 4905
Gallus_gallus                    4861 TGTTCTGAGGAGCCATGCCATGCCTGTGGTTGCCACCCCTCTCAT 4905

Anas_platyrhynchos_platyrhynchos 4906 CTGAATCTTTCTGTGATTGTTTATCTTGTCTTTGTTTTGCTGACT 4950
Gallus_gallus                    4906 TTGCTTCTTCCTGTGATTCTTTCTCTTGGCTTTGTTTTGC-AAGT
```

4950

Anas_platyrhynchos_platyrhynchos 4951 ACATAAAATGTATTACTTTAATTTACTTTAAATTACTTTACTAGA
4995
Gallus_gallus 4951 ATGCAAAACATAT----------------------------TGAA
4995

Anas_platyrhynchos_platyrhynchos 4996 GTACTTTACTCTAGAAGGGAA----AATTGTCACC----AGTTCA
5040
Gallus_gallus 4996 G-----TACTCTAAAAGGGAAAATCAATCATCACCAGGTAGGTCA
5040

Anas_platyrhynchos_platyrhynchos 5041 CTGTGTAAATATACCTCTGCAGAGTAAATATGAAAACAGTTTTAA
5085
Gallus_gallus 5041 CTTTGTAAATACACCTCTACAGGGTAAATAGAAAATCAG-TTTAA
5085

Anas_platyrhynchos_platyrhynchos 5086 AATTGAACATCTAATTACTTGAACATATAAATTAATTCCTGCCTC
5130
Gallus_gallus 5086 AATGGAACATCTAATTACATCGACATAGAAGTTCATTCTTACCTC
5130

Anas_platyrhynchos_platyrhynchos 5131 CTACAGAGCTTCTGTATCTCTGCCAAGTGATATTTTGGTTCTGTT
5175
Gallus_gallus 5131 CCGCGGGGCTTTTGTTTTGCTGCCAGCTAACATTTGGGTGCTGTT
5175

Anas_platyrhynchos_platyrhynchos 5176 TTAGAAGCTGCATAGTGAAGCCAGCCTTTTAACTCAGTCACTTCT
5220
Gallus_gallus 5176 TCAGAAACTGCATAGGAAAGCCATTCTTCTAACTCTGTCACTTCT
5220

Anas_platyrhynchos_platyrhynchos 5221 CTGAATCAAAACACATTGCTGGTCTTCAACCTGAAGAGGAATCCC
5265
Gallus_gallus 5221 CTGAGTC-AAACACATTGCTGATCCTCAG-CTGAAGAGGAATCCC
5265

Anas_platyrhynchos_platyrhynchos 5266 ACTGTTGCTCTCAGGTTGATTAAACTACACAGAAGGAATATGTGT
5310
Gallus_gallus 5266 GCTTTTGCTCTCAGGCTGATAAATCTGCCCAGGGAAGTTATCTGT
5310

Anas_platyrhynchos_platyrhynchos 5311 GAGGGACAATGGTTTTCTACCAGTTTGTGGCACGGAAGGCGTGTC
5355
Gallus_gallus 5311 GGGAGATGATGGTACTGTGTTAGTTTGTGGCATGGAAGGCATGTC
5355

Anas_platyrhynchos_platyrhynchos 5356 TCATGGAGTTAAGGTCAGGCTGTTTCAAGATGTTGGTGAACAACC
5400
Gallus_gallus 5356 CTATGGAG-TAAGATCAGACTATTTCAAGATTTTGGTGACCAACA
5400

Anas_platyrhynchos_platyrhynchos 5401 AGGTGGTCTTT----AATAAAGTGATACCTTAATGTGGTGGGCAG
5445
Gallus_gallus 5401 AGGTGATCTGTCCCAAATAAACTGATATTTTGATGTTGTGGGCAG
5445

Anas_platyrhynchos_platyrhynchos 5446 CTGTCTGTAGGCTAGAGTTCTACAACAGTCTCAAATATGACATTT
5490
Gallus_gallus 5446 CTGTCTGTAGGTTAGAGCT-TATGACAGTCTGAAATATGACATTT
5490

Anas_platyrhynchos_platyrhynchos 5491 CTTGCAAAGATAGTGTTTTGAGGATCTATCTTCATTTGTTAAATT

```
5535
Gallus_gallus                     5491 TTTGCAAAGATAATGTTTTGAAGATATACCTTCATTTGTTAGTTT
5535

Anas_platyrhynchos_platyrhynchos 5536 GGAAGTTATATGTAATTTGCAGCTTTTCAGAGTAGAACAGAATAA
5580
Gallus_gallus                     5536 GGAAGCTACACATAATGTGCAGTTTTTCAAAGTAAAACCTAGTGG
5580

Anas_platyrhynchos_platyrhynchos 5581 AAAAGTCATTGTGAGATAAAAAGGTCCAATTATTTGCTTGGGTTC
5625
Gallus_gallus                     5581 AAAAAGCCATGTGAGCT-AAGAGGTGCAATTAGATGCTTGGTTTT
5625

Anas_platyrhynchos_platyrhynchos 5626 GTAGTCATTATTACTGACAAACTAGACAAGTATGGAAATCTTGTG
5670
Gallus_gallus                     5626 GTAGTCATTA-TACTAACAAACTGGACAAGTATGAAAATCTTGTG
5670

Anas_platyrhynchos_platyrhynchos 5671 TTCTTAATAAAATATTAGAATAATGCACATGAATTGCACATCCTT
5715
Gallus_gallus                     5671 CTCTTAATAAAATATTAAAGCAACACATGTGATTTGCCCATCTTT
5715

Anas_platyrhynchos_platyrhynchos 5716 TGAAAACTTGGATTGATGAAAAATGTTCCTGATATGGTGGGTGGT
5760
Gallus_gallus                     5716 TGAAAACCTGTATTGCTGAAAAAGTTTCCTGATGTGGTAAGTAGG
5760

Anas_platyrhynchos_platyrhynchos 5761 CAAATGCTGCAACAGGTTACAAAGAACTGTTGGAGTCTCCACAGC
5805
Gallus_gallus                     5761 CAAACGTGGCCACTAGTTGCCCAGA-------GAAGTTCAACTGG
5805

Anas_platyrhynchos_platyrhynchos 5806 AAAGAAGCTGTGCAGTCTCCATCTGTGGAGATGTTCAAAACCTAA
5850
Gallus_gallus                     5806 AAA------GTTCAA---------ATGAAAAAGTTAAGAACCTAA
5850

Anas_platyrhynchos_platyrhynchos 5851 CTGGACGTAGTCCTGGACGACCTGTTCTATGGATTTGGCTTACGG
5895
Gallus_gallus                     5851 CTGGATGTAGTCCTGGGCACCCTGCT-GGTAGATCCTGCTTGAGC
5895

Anas_platyrhynchos_platyrhynchos 5896 TGTGTAGGGCTGGACTAGGTGACC-------------TCCAACAT
5940
Gallus_gallus                     5896 CATGTCAGACTGGACTAGATGACCTCTAGAGATGCCTTCTAACCT
5940

Anas_platyrhynchos_platyrhynchos 5941 AAATGATTCTGTGACT-CTGTGATGGTGGGATAGTGTTTGATTTT
5985
Gallus_gallus                     5941 AAATGATTCTGTGACTGCAGT----GTGGGACGAGGCTTGCTTTT
5985

Anas_platyrhynchos_platyrhynchos 5986 CAGTGAATCTATCTGATTTTTTTATGCTGACTGTGTTTTCCCACT
6030
Gallus_gallus                     5986 CAGTAAATCAATTTGATTTCGAT------GCTGTGTTTT-TTACT
6030

Anas_platyrhynchos_platyrhynchos 6031 GTAGGTATTGTTGCTCTCTGGTCTCTTGCGCTTTTGGCTTTTGAG
6075
Gallus_gallus                     6031 GTAGGCATTGTGGCTCTCTGGTCTCTTGCTCTTTTGGCTTTTGAG
6075
```

```
Anas_platyrhynchos_platyrhynchos 6076 CGGTACATAGTGATCTGCCGCCCACTGGGAAATATGCGCTTGAAG
6120
Gallus_gallus                    6076 CGGTACATTGTGATCTGCCGCCCAGTGGGAAATATGCGCTTGAGA
6120

Anas_platyrhynchos_platyrhynchos 6121 GGGAAGCATGCTGCCTTAGGTATTGCCTTTGTGTGGACCTTTTCC
6165
Gallus_gallus                    6121 GGGAAGCATGCTGCCCAAGGTATTGCTTTTGTGTGGACCTTTTCC
6165

Anas_platyrhynchos_platyrhynchos 6166 TTCATTTGGACCATCCCACCAACTATGGGTTGGAGCAGCTACACC
6210
Gallus_gallus                    6166 TTCATTTGGACCATTCCACCAACCATGGGTTGGAGCAGCTACACC
6210

Anas_platyrhynchos_platyrhynchos 6211 ACCAGTAAGATTGGAACTACTTGCGAGCCTAACTGGTAAGGCCAT
6255
Gallus_gallus                    6211 ACCAGTAAGATTGGAACTACTTGTGAGCCTAATTGGTAAGGTCAT
6255

Anas_platyrhynchos_platyrhynchos 6256 CCTTGCAGAGTATTGTGAAGAAATTACACTTACAGCTCATGACTT
6300
Gallus_gallus                    6256 CCTTGCAGAGTATGGT--AGAGTTTAAACTTGTAGCTCATGACTT
6300

Anas_platyrhynchos_platyrhynchos 6301 CTCCAAAGCAGGAGTTTAAAGGAAAGGCTTGAAGATGATGGACTT
6345
Gallus_gallus                    6301 CTTAAGCACACAGTTCCAAAGGAAAGGGTTAAGGATGATGGTCTT
6345

Anas_platyrhynchos_platyrhynchos 6346 ACAGGCAAAACTAGAGGTAGACAAAGCTATAACAAAACCAAGAAA
6390
Gallus_gallus                    6346 ---TGCCAAAAAAGAGGTAGACGAAGATATAACAA---TAAAGAA
6390

Anas_platyrhynchos_platyrhynchos 6391 AAATTGCCAAGAACAGGTTAAAAAAATGTGTGTGTGTGTGTGTAT
6435
Gallus_gallus                    6391 AAATTTCCA-----GGGTTAAAAATAAATGT-------TTTCTAA
6435

Anas_platyrhynchos_platyrhynchos 6436 AA---TTTTCTAAAAGGAGAAGCTTTTGTATCTAATGAAGATAAC
6480
Gallus_gallus                    6436 AACATTTTCCTAAAAGGAGAAGCTTACATCTCTAATGAAAGTAGC
6480

Anas_platyrhynchos_platyrhynchos 6481 TGTCCCATACCTTACGGTTTTTGTCTGGATTCAACGAGGACTAAA
6525
Gallus_gallus                    6481 TGTTCCCTATATTGGGGGTTTTGTTTG...........GACGAAT
6525

Anas_platyrhynchos_platyrhynchos 6526 ATATTTTTAGAAGTCTGGACAAGGCCCTGTGCAACCTGTTCTACC
6570
Gallus_gallus                    6526 AGATTTTCAGAAGATTGGATGAAGCCCTGAGCAATTTGTTTTAGC
6570

Anas_platyrhynchos_platyrhynchos 6571 CCCATAGTTGACATCACATTGAGCAGGAGGTTGGATTACATGTAC
6615
Gallus_gallus                    6571 AGCATAGCTGACCTCTCTTTGAGCAGGAGGTCGGACTACACGTTT
6615

Anas_platyrhynchos_platyrhynchos 6616 CTTCCCACCTGGACAGACCCACAATTCTATAGTTCTGTGTCTGTG
6660
```

```
Gallus_gallus                      6616 CTTCCCACCTAACGTACCCCACAATTCTTTGGTTCTGTGCCTATG
6660

Anas_platyrhynchos_platyrhynchos 6661 ATCCTACAAAGTAAACGTTTCCTAGCCCAGTGCTAGGGAGATGGC
6705
Gallus_gallus                      6661 ATCTTACAGAGTAAACCTTTGCTAGCCCAGAGGAAGGGAGATGGC
6705

Anas_platyrhynchos_platyrhynchos 6706 CCAGGGCTCCATCTGGTGATTTGACTAGAGCTATACAGGGGGGTT
6750
Gallus_gallus                      6706 CATGGGCTCTCTCTGGTGATTTGACTAAAGCTGTACATGGGTACT
6750

Anas_platyrhynchos_platyrhynchos 6751 TTGATTAAAAACAGTTAACAAAAATATTAGTGACACCTTTAAAGA
6795
Gallus_gallus                      6751
TTGATTAAAAACAGTTGATGAAAATATTAGTGATGCCTC------ 6795

Anas_platyrhynchos_platyrhynchos 6796 GCAAAATGAACTCTTCAAGAATATTAATCAGAACACATTAGTAAT
6840
Gallus_gallus                      6796 ------TGAACACTTTGAGAA-AATAACCAGAGCGGATTAGTAAT
6840

Anas_platyrhynchos_platyrhynchos 6841 ACATACACGACGTATTCTGTGTTTTATACAGGTATTCAGGAGCTT
6885
Gallus_gallus                      6841 ATATACACAATGTATTCTGTGTTTTATACAGGTATTCAGGAGCTT
6885

Anas_platyrhynchos_platyrhynchos 6886 ATACTGATCATACCTACATTATTGCATTTTTCACCACCTGTTTTA
6930
Gallus_gallus                      6886 ATAATGATCGTTCCTACATTATTGCATTCTTCACCACCTGTTTTA
6930

Anas_platyrhynchos_platyrhynchos 6931 TAGTGCCTTTATTGGTAATTTTGGTATCCTATGGAAAACTGATGC
6975
Gallus_gallus                      6931 TAGTGCCTTTATTGGTAATTCTGGTGTCGTATGGGAAACTGCTGC
6975

Anas_platyrhynchos_platyrhynchos 6976 AGAAGCTAAGAAAGGTAAGAAGAAGCACTAATAAG-TAGAACTGT
7020
Gallus_gallus                      6976 AGAAGCTAAGAAAGGT---AAGAAGCACTTTCAAGTTACATCTTT
7020

Anas_platyrhynchos_platyrhynchos 7021 GAAATGTGTATGTATGTAATCATTTGGAATTTTTGTGTTCAAATT
7065
Gallus_gallus                      7021 GAGATGTGTATGTATGCAATAATTT-GATTCTTCATGTTCAGATT
7065

Anas_platyrhynchos_platyrhynchos 7066 TCCATAATGATCAGATGTCAG-------ATTTCCAGACAGCAAAT
7110
Gallus_gallus                      7066 TCCTTAAGGATCAGACATAAGCAGACCTGTTTCCAGGCAGCAAAT
7110

Anas_platyrhynchos_platyrhynchos 7111 ACG-----------------CTCACTTGTCAAGCTTTGTTAAATA
7155
Gallus_gallus                      7111 ATGTACAAATATATACAAGACTCACTTAT---GTCGTGTTAAAGA
7155

Anas_platyrhynchos_platyrhynchos 7156 ATTATAATTACAAACAGAAATCTATGCTATAGTTAGAAATGATTT
7200
Gallus_gallus                      7156 ATTATTTTTGCA.................................
7200
```

```
Anas_platyrhynchos_platyrhynchos 7201 TTCAGTGTCCAAGTTTTATCACTTATGTCTCATTAGCCTAACATT
7245
Gallus_gallus                    7201 .............................................
7245

Anas_platyrhynchos_platyrhynchos 7246 GTTAGAAAAAATGCCAGCCTCATAAATAAAGAAGCTGGATGAGCC
7290
Gallus_gallus                    7246 .............................................
7290

Anas_platyrhynchos_platyrhynchos 7291 AGTATTGGTTTATTTACAATTGCAATTCAGTTCAGTGGCAGTTTT
7335
Gallus_gallus                    7291 .............................................
7335

Anas_platyrhynchos_platyrhynchos 7336 TCAGATTCAAACCAGACAATTCAGAATGTAGCTTAAAACGTCTGC
7380
Gallus_gallus                    7336 .............................................
7380

Anas_platyrhynchos_platyrhynchos 7381 ATTTGTAAGTTGGTGTTGTCAGATACCTCCAGTTGCTCATTCCGT
7425
Gallus_gallus                    7381 .............................................
7425

Anas_platyrhynchos_platyrhynchos 7426 CAGTCATTGTATTCTGTATTTTGGCATTCTAAATAGAGGCCTATT
7470
Gallus_gallus                    7426 .............................................
7470

Anas_platyrhynchos_platyrhynchos 7471 ATCTCTAATTCGACTAAAATCATTAGGATTCCCTTTCTCACAGGT
7515
Gallus_gallus                    7471 .............................................
7515

Anas_platyrhynchos_platyrhynchos 7516 GGAATTTGTCTGTATATCTGCACTGCCTTGAGGTGTGTTGCTGCA
7560
Gallus_gallus                    7516 .............................................
7560

Anas_platyrhynchos_platyrhynchos 7561 GCACCCATGAAAATATTTGAGATAAATCAACATGTGTGAGGCTGG
7605
Gallus_gallus                    7561 .............................................
7605

Anas_platyrhynchos_platyrhynchos 7606 ACAGCAAATCTGGGCCATCACTTCACTTCTCATGGTTGGCTAAAT
7650
Gallus_gallus                    7606 .............................................
7650

Anas_platyrhynchos_platyrhynchos 7651 CTAGCTTGGGTGTGCCTGTACAGGCTACAGTCATGCTGGACTGCA
7695
Gallus_gallus                    7651 .............................................
7695

Anas_platyrhynchos_platyrhynchos 7696 ACTGTGTTTTTGTACAAAGATGGTCCTAGCATTTTTTTCTGTTAT
7740
Gallus_gallus                    7696 .............................................
7740

Anas_platyrhynchos_platyrhynchos 7741 TTTGACAATAAAAATATAAAGATATGTAAACAAATGAAAATTGAA
7785
Gallus_gallus                    7741 .............................................
```

```
7785

Anas_platyrhynchos_platyrhynchos 7786 TAATCACAAGATAGAGTAATTCACATCCCTTAGCACTCATACTGA
7830
Gallus_gallus                    7786 .............................................
7830

Anas_platyrhynchos_platyrhynchos 7831 CACTCTGTGGTCCAGTAACAGCATTCTTATGTGCTATCATACTTG
7875
Gallus_gallus                    7831 .............................................
7875

Anas_platyrhynchos_platyrhynchos 7876 ACAAGAACAGATCTTCCCTTATAGCATGGCAGATTATAAAACTTC
7920
Gallus_gallus                    7876 ........................................ATTTA
7920

Anas_platyrhynchos_platyrhynchos 7921 CCCCACATTTGCTGCATGAAGGCCTTGCTTATTTTCTGGAATTTT
7965
Gallus_gallus                    7921 CACTATATTCACTCTTTGAAGGCCTTGTTTGTTTTCTGGAATTTT
7965

Anas_platyrhynchos_platyrhynchos 7966 CTGTCTTTTCTTTAAAGGTGTCAGATACTCAATGCAGGCTGGGAA
8010
Gallus_gallus                    7966 CTGTCTTTTCTTTAA-GGTGTCAAATACACAAGGCAGGTTGAGAA
8010

Anas_platyrhynchos_platyrhynchos 8011 CTACCAGGAAACCTGAAAGACAAGTGACTAGAATGGTTGTTGTTA
8055
Gallus_gallus                    8011 CTGCCAGGAAACCTGAAAGACAAGTGACTAGAATGGTGGTTGTTA
8055

Anas_platyrhynchos_platyrhynchos 8056 TGATCATTGCGTTTCTAATCTGCTGGATGCCATATGCTGCCTTTT
8100
Gallus_gallus                    8056 TGATCATTGCTTTTCTAATCTGCTGGATGCCATATGCCGTCTTTT
8100

Anas_platyrhynchos_platyrhynchos 8101 CTATCCTAGTCACTGCATATCCTTCCATTGAATTGGATCCTCGTC
8145
Gallus_gallus                    8101 CTATCCTAGCCACTGCATACCCTTCCATTGAACTGGATCCTCACC
8145

Anas_platyrhynchos_platyrhynchos 8146 TGGCAGCAATTCCAGCTTTCTTTTCCAAAACAGCTACTGTTTATA
8190
Gallus_gallus                    8146 TGGCAGCAATTCCAGCTTTCTTTTCCAAAACAGCTACTGTTTATA
8190

Anas_platyrhynchos_platyrhynchos 8191 ATCCAATTATTTATGTCTTTATGAACAAACAGGTACTGTATGCTT
8235
Gallus_gallus                    8191 ATCCAATTATTTATGTTTTTATGAACAAACAGGTACTGTATGTTC
8235

Anas_platyrhynchos_platyrhynchos 8236 TCAGAGTGTCTTTAAAACTCATATTTTCATATGTAGACTTGCTTT
8280
Gallus_gallus                    8236 TCAAAGTGTCTTCCATATCCATAGTT-------TAGAATTTTTTT
8280

Anas_platyrhynchos_platyrhynchos 8281 ATAATGAATTATT-TCTATCTACTTGTTAGTTAACCAGCAATACA
8325
Gallus_gallus                    8281 ------AATCACTGTATGTCTACTT--------------AATGCA
8325

Anas_platyrhynchos_platyrhynchos 8326 ATAACTAGCATTTAGAATTTAAAAAAATAAAATGAAAAAAATCAC
```

```
8370
Gallus_gallus                     8326 ------------TAGAGTTTTG--------------------TGC
8370

Anas_platyrhynchos_platyrhynchos 8371 TATGTCTACTTACTGGAGCATAGCATAAACCTCAGAGCATTATTA
8415
Gallus_gallus                     8371 TGTGTTT------------------------------CAGTATTA
8415

Anas_platyrhynchos_platyrhynchos 8416 CAATATGAATGTTAATAATAATTAGGTTTAACTATTAGAAAGGGA
8460
Gallus_gallus                     8416 AAATATGAATATTAGTAATCATTAGATTTAGGTATTGGAAAAGAA
8460

Anas_platyrhynchos_platyrhynchos 8461 AATCAGCCCTTAGTGAATGTCAGTTATTTCATACAGTTTTCTGTA
8505
Gallus_gallus                     8461 AATCAACTCTTAGTGAACATCAACTATTC----TAGTTTTCTGTG
8505

Anas_platyrhynchos_platyrhynchos 8506 TGAAAAGTCTATTTTCAAGACTGTCCCATGATATTTCT-----AT
8550
Gallus_gallus                     8506 AGAAAATTCTATTTCCAAGACTGTCCTGTTGTATGTTTCCATAGT
8550

Anas_platyrhynchos_platyrhynchos 8551 TCTTTAGGAATGCAATACCAGTGAATTGCTTTTGAGGTAGGGTGC
8595
Gallus_gallus                     8551 TATCTTGGAATGCAATGCCAGTGAATTGCTTCTGA.......TGC
8595

Anas_platyrhynchos_platyrhynchos 8596 TCATGGATATGAACAAAATGGAAAGAGAACATCTTTGAATAGCTG
8640
Gallus_gallus                     8596 TCAAGGCCATGAACGTGATGGAAACAAACCATCTTTGAATATCTA
8640

Anas_platyrhynchos_platyrhynchos 8641 TGTATTAGTGCATGAAAGCAACTCATGAACTCCGAGAAATGAGCA
8685
Gallus_gallus                     8641 GGTATTAGCATACAAATGCAACTCATGAGCCCCAGGAAATGACCA
8685

Anas_platyrhynchos_platyrhynchos 8686 ATTCTCTTCAGCAAGATGTGGCTAGGAACAAGCATGAGAAGAAAA
8730
Gallus_gallus                     8686 ATTCCATTCAATTAGATGCAGCTAGCAACAAGCAT-AGAGGAAAA
8730

Anas_platyrhynchos_platyrhynchos 8731 TTATTTCTTAGAATGAGGCTTACTAGTAGTTTGGGCCTTTTTTAG
8775
Gallus_gallus                     8731 TGATTTTTTAGACTGAAGCATTCCAGCAGTTTGTGTCTTTTT-AG
8775

Anas_platyrhynchos_platyrhynchos 8776 AATGTCATTGTTTCTCACTTTGAGAGTTCTTCGTGGTGTCACAAG
8820
Gallus_gallus                     8776 A----------TTCTCA--------GTCCTTTGTGGTCCCACAGA
8820

Anas_platyrhynchos_platyrhynchos 8821 CATATATATTTAATGGGGCTATTTCTGTATGTGCCCAACTGATTA
8865
Gallus_gallus                     8821
TAAATATATTTTATGGGGCTATTTC-------------------- 8865

Anas_platyrhynchos_platyrhynchos 8866 ATAAGATAATTTCCAAGAGTTCTAGTTAAGACACAAACTGCTTTA
8910
Gallus_gallus                     8866 -------------CAAGACTTTTAGTTAATATAAAAAAAA--TTA
8910
```

```
Anas_platyrhynchos_platyrhynchos 8911
TAGAAATATTTTTTAATTTCT--TGTGTTTTGATTGATCTCCT-- 8955
Gallus_gallus                    8911 AGAAAATATTTTTAAATTCCTCGTGTGTTTTTATTTTTCTTGTAT
8955

Anas_platyrhynchos_platyrhynchos 8956 -TTGTTTCAACCTTCTAGTTCAGGAGGTGTCTGGTTGAGATGTTC
9000
Gallus_gallus                    8956 GTTGTTTCAAATTTC-AGTTCAGGATGTGTCTGATTCAGATGTTC
9000

Anas_platyrhynchos_platyrhynchos 9001 AGCTGCAGTGCCATAGAGACTGCTGAGTCCAACATGAACCCGACT
9045
Gallus_gallus                    9001 AAGTGTAGTGCCATAGAGACTGCGGAGTCCAACATGAACCCAACT
9045

Anas_platyrhynchos_platyrhynchos 9046 TCAGAGAGAGCAATGCTAACCCAGGACAAAAGAGGCAGTGAGTTG
9090
Gallus_gallus                    9046 TCAGAGAGAGCAACACTCACCCAAGACAAAAGAGACAGTCAGTTG
9090

Anas_platyrhynchos_platyrhynchos 9091 TCTATCATGGCAGTATGTAGCACCGTTTCTAAGAGGAAAACTGGA
9135
Gallus_gallus                    9091 TCTGTCATGGCAGTACGAAGCACCATTTCTTAGAGGAAAAATGGA
9135

Anas_platyrhynchos_platyrhynchos 9136 GATGAACACACAAATTGCCCTTCTTTTGCTCAGTTGGCAGCTTCA
9180
Gallus_gallus                    9136 GATGAACACAAGAATTGCAATTCTTTTGCTCAGTTAACAGCTTCA
9180

Anas_platyrhynchos_platyrhynchos 9181 GAAAACAAAATCTGGCCCATGTAGGGAGTGAGATGTGATCATTAA
9225
Gallus_gallus                    9181 GGAAACAAAATCTGTCTCTTGTAGAAGGT-AGATATGATCATTAA
9225

Anas_platyrhynchos_platyrhynchos 9226 TGTAGTACTACTATATCATGTTTCCTTGTTA--CCATCACTTCAT
9270
Gallus_gallus                    9226 TGCAATACAGC---AGCGTGTTTCCTCATTACCCCATCTCTTCAT
9270

Anas_platyrhynchos_platyrhynchos 9271 ATAAAAAATGTCATGGCGGACATTATGCTCAGTTCTGTGGACACT
9315
Gallus_gallus                    9271 ACTAAAGATGTCATCGGAGCCATTGTGCTCTGTCTCATTAACACT
9315

Anas_platyrhynchos_platyrhynchos 9316 GGCTTTGAGATTCTGAGCTCCTGAGTGAATGCAAGAGCTTCTTGG
9360
Gallus_gallus                    9316 AACTTTGAGATTCTGAGCTCCCAAGCGCATATAAGAGCTGCTTGG
9360

Anas_platyrhynchos_platyrhynchos 9361 CTCTGCAGTTGCAATCAAACGCCCAGCCAAAACGTTAGTCCCTTC
9405
Gallus_gallus                    9361 CTGTGCAGTTGCAGTCAGA-GACCAGCCAGAAAGTGAGTCCCTTT
9405

Anas_platyrhynchos_platyrhynchos 9406 CTCTATTGTGTTAGTCCTCTATTATCCACTGCCTGCTTGCCCCTG
9450
Gallus_gallus                    9406 CTCTA---TGTTTGT-CTCTGCAATCCACAGCCTACTCGCCCCTCT
9450

Anas_platyrhynchos_platyrhynchos 9451 GTAATAAAACTAGAGAAGCCCAATATTTTCATGCAGGCATGTGGC
9495
```

```
Gallus_gallus                     9451 GGTATAAAACAATGGAAGCCCAACATTATTGTGCTGACACACAAT
9495

Anas_platyrhynchos_platyrhynchos 9496 ATAATAAGGATAAGAGTGTTTTACACTGACGACTGTGTAAAACAC
9540
Gallus_gallus                     9496 ACAGCAAAGAAAA----GTTTCACATTGATGACTGCTCCTTAGTC
9540

Anas_platyrhynchos_platyrhynchos 9541 GACTGTGGCAGAAACATGAAGTGGTACAGGGCTTCAACATGTTAA
9585
Gallus_gallus                     9541 AGCTG-GGCAAAAAAAATGCAGTGATGCATAGCTTCAATACATTAA
9585

Anas_platyrhynchos_platyrhynchos 9586 TGGAAAATGAATAATATTTAGCATACAACTGACTCAAGTTCAACC
9630
Gallus_gallus                     9586 TGGAAAATGAGTAATATTTAGCATACAGCTGACTCAAGTTCAACC
9630

Anas_platyrhynchos_platyrhynchos 9631 AAGTGGCGGTTCAGAGAGAGGCAGGCCTGTGTGTTTTTTGGTTTG
9675
Gallus_gallus                     9631 ACACAGAGACTAATGTAGAGACAGGCTTGAGTGCTTTTTGATTTG
9675

Anas_platyrhynchos_platyrhynchos 9676 TTTCAGAATTTTTACTTAAAACAGACAAACAAAAAGCCTAAGGCT
9720
Gallus_gallus                     9676 TTTGAAAATTTCTACTAAAAAAAG---CAGAGGAAGCGTCAGGTC
9720

Anas_platyrhynchos_platyrhynchos 9721
TCTGCACTTATGTCTGTCCGTGGTGAAAACGTCATCTG------- 9765
Gallus_gallus                     9721 TCTGCAGTTATGTCTATCCCTGATGAGAACCTCATTTGGGCCACT
9765

Anas_platyrhynchos_platyrhynchos 9766 --------CAGCAAGTCAGGTGAGCTGTTAAGACTTCCAAGGACT
9810
Gallus_gallus                     9766 AAATTCTCCAACAACATGGATGAGCCATCCATACTTTCAAGGACT
9810

Anas_platyrhynchos_platyrhynchos 9811 AACTTATTAAAAAAAATTCCCATCAAGAGTCATTCATATCCTCCT
9855
Gallus_gallus                     9811 ---TTGTTAG-------------TAAGAGCCATTCATA---CTCT
9855

Anas_platyrhynchos_platyrhynchos 9856 TCTTCATTCCCTCTTTGTCAGGAAAACTTCTGTGCCCATAGCAAC
9900
Gallus_gallus                     9856 TCTTTATGTCCTCTGTGCCAAGAAA---TCTGTACTAATGGCAAC
9900

Anas_platyrhynchos_platyrhynchos 9901 TTCTG----AAATCAACTCA------ACTTTAGCAGAGAAGGTAG
9945
Gallus_gallus                     9901 TTTTGAAATAAATCAACTCATGCCATCCTTTAGTGGGGAGG....
9945

Anas_platyrhynchos_platyrhynchos 9946 GGTATAGAAAACTGTGCAGAAATACTGAAGAAAGGGTCAGTAGCA
9990
Gallus_gallus                     9946 .............................................
9990

Anas_platyrhynchos_platyrhynchos 9991 CTACTTGAGTTCTGAAGCTGAGCTTTGTAATACAGAAGTTTTTTA
10035
Gallus_gallus                     9991 .............................................
10035
```

**Commented [Unknown A1]:** Deletion starts

```
Anas_platyrhynchos_platyrhynchos 10036 CTGTATGTCCTTGCAACGTGCAGAGCTAAGGTCTATCATATTCTT
10080
Gallus_gallus                    10036 .............................................
10080

Anas_platyrhynchos_platyrhynchos 10081 GAGATCATGTAGCTCCAGGAGACAACTATTATTTCTGTTTTTTAA
10125
Gallus_gallus                    10081 .............................................
10125

Anas_platyrhynchos_platyrhynchos 10126 TCATAGAATGGTTTGAGTTGGAAAGGACCCTTAAAGATTATCTAG
10170
Gallus_gallus                    10126 .............................................
10170

Anas_platyrhynchos_platyrhynchos 10171 TTCAACCCCCTGCCTGGGCAGGGACATCTCTCACTAGACCAGGTT
10215
Gallus_gallus                    10171 .............................................
10215

Anas_platyrhynchos_platyrhynchos 10216 GCTCAAAGCCCCATCCAACCTGAACTCAAACACTTCTAGGGATGG
10260
Gallus_gallus                    10216 .............................................
10260

Anas_platyrhynchos_platyrhynchos 10261 GGCGGGGCATCCACAGCTTCTCTTGGCAACCTGTTTCAGTACCTC
10305
Gallus_gallus                    10261 .............................................
10305

Anas_platyrhynchos_platyrhynchos 10306 ACTACCCTAGTAGTGCTCCACAGTGAGGATATTCTTCCTTGTATC
10350
Gallus_gallus                    10306 .............................................
10350

Anas_platyrhynchos_platyrhynchos 10351 TAATCTAAATCTATCCTCTTGCAGTTTAAAACCATTGCCCCTAGT
10395
Gallus_gallus                    10351 .............................................
10395

Anas_platyrhynchos_platyrhynchos 10396 CCTATCACTATAGGCACTTGTAAAATGTTTCCCTCCATTTTCTTA
10440
Gallus_gallus                    10396 .............................................
10440

Anas_platyrhynchos_platyrhynchos 10441 TAAGCCTCCTTTAAGTATTGCAAGGCTGCGATAAGGCCATCCTGG
10485
Gallus_gallus                    10441 .............................................
10485

Anas_platyrhynchos_platyrhynchos 10486 AACCTTCTCTTTTCTAGGTCTAACACTTCCATGTTTCTCAGCCTT
10530
Gallus_gallus                    10486 .............................................
10530

Anas_platyrhynchos_platyrhynchos 10531 AAAGAAAAAGCTTTGTGCACAAACAAAGCAAAAGAAGGAATTCAT
10575
Gallus_gallus                    10531 .............................................
10575

Anas_platyrhynchos_platyrhynchos 10576 TTACTACGTCCCATTGGCAGGCAGGTGTTCAGCCATCTACAGGAA
10620
Gallus_gallus                    10576 .............................................
```

```
10620

Anas_platyrhynchos_platyrhynchos 10621 AGCAATGCCCATCACATATAACAATTTATTAGGAAGACAAACACC
10665
Gallus_gallus                    10621 .............................................
10665

Anas_platyrhynchos_platyrhynchos 10666 AAAGACAAATGCACCTCACCCCCGCTAAGACCCCATTGTTCCTTC
10710
Gallus_gallus                    10666 .............................................
10710

Anas_platyrhynchos_platyrhynchos 10711 TTTACCCCAGTTTTCTTGTTCTTCATGAAGCCATGTAGTTTGGGA
10755
Gallus_gallus                    10711 .............................................
10755

Anas_platyrhynchos_platyrhynchos 10756 TATCCCTTTGGACAGTTGGGGGCAGCTGTCCTGGCTCTGTCCCCT
10800
Gallus_gallus                    10756 .............................................
10800

Anas_platyrhynchos_platyrhynchos 10801 CCCTGCTCCTTGTGCACCCCCAGCCTCCTCACTGGCAGAGCGGTG
10845
Gallus_gallus                    10801 .............................................
10845

Anas_platyrhynchos_platyrhynchos 10846 GGAGAAGCTGAAATGTCTGTGACTTACTGCTGGCACTGTTCAGGA
10890
Gallus_gallus                    10846 .............................................
10890

Anas_platyrhynchos_platyrhynchos 10891 ACAGCCAACCTATCAATGTGTTATCAACATTATTTTCACCACAAA
10935
Gallus_gallus                    10891 .............................................
10935

Anas_platyrhynchos_platyrhynchos 10936 TCCAAAACATATCATCATACAAGCCTCTACAAAACAAAAACAAAA
10980
Gallus_gallus                    10936 .............................................
10980

Anas_platyrhynchos_platyrhynchos 10981 CACTATCCCAGCCAAAACAAGGACACTTAACAATTACAATTTCAG
11025
Gallus_gallus                    10981 .............................................
11025

Anas_platyrhynchos_platyrhynchos 11026 GAAATTAGTGTCCGGAGTATACAAAAAATATGCATCTTCTGGGCC
11070
Gallus_gallus                    11026 .............................................
11070

Anas_platyrhynchos_platyrhynchos 11071 ATGTTACATCAGAGCATTGTGGATGGCTGTGTTTCCCTTCACGTT
11115
Gallus_gallus                    11071 .............................................
11115

Anas_platyrhynchos_platyrhynchos 11116 TTTGCTTTTCTCTCACTTCTGAGCTAGAGTATGTCTCAGAAAACA
11160
Gallus_gallus                    11116 .............................................
11160

Anas_platyrhynchos_platyrhynchos 11161 TAATCAGTGTTCTAGCTGAACAGGAAACTTGTAGCAGTCAATCAC
```

```
11205
Gallus_gallus                    11161 .............................................
11205

Anas_platyrhynchos_platyrhynchos 11206 ACCATGTTGCCGTTCTTCTCTTAGGAGCAGTTGCATCTGTGTCCC
11250
Gallus_gallus                    11206 .............................................
11250

Anas_platyrhynchos_platyrhynchos 11251 AGCAGTCCTTAAATTTCTGCTTCTGTACATCACTTCCTCAGCACT
11295
Gallus_gallus                    11251 .............................................
11295

Anas_platyrhynchos_platyrhynchos 11296 TTTAATTGAGCTACTGATCCTGACATAAGTTGTTAACTTGTTCTT
11340
Gallus_gallus                    11296 .............................................
11340

Anas_platyrhynchos_platyrhynchos 11341 TCACCTTTTAGGTCAAAAGCAGGGACCACAACAAAAATATACTTT
11385
Gallus_gallus                    11341 .............................................
11385

Anas_platyrhynchos_platyrhynchos 11386 CCATAAACAGTGGAAAAGAACAGAAATTGCCTGCTTTCCATGTTT
11430
Gallus_gallus                    11386 .............................................
11430

Anas_platyrhynchos_platyrhynchos 11431 GTGTGCCATCTCATATATTTAACTCATACAATGATATGGTTCAAT
11475
Gallus_gallus                    11431 .............................................
11475

Anas_platyrhynchos_platyrhynchos 11476 ACACAAAACTTTAGCTGAACATCACTTACATGCAAAAGCAAACCT
11520
Gallus_gallus                    11476 .............................................
11520

Anas_platyrhynchos_platyrhynchos 11521 GGTTATGCATGTAATAAAGACCTTTTTTGGTGAACATTGTATGAA
11565
Gallus_gallus                    11521 .............................................
11565

Anas_platyrhynchos_platyrhynchos 11566 ATGTTACTAAAATCTGGTATCTCAGATTCTGAATTGTATATAAGT
11610
Gallus_gallus                    11566 .............................................
11610

Anas_platyrhynchos_platyrhynchos 11611 TCTAAGCTTACCTGAATGCAGTTTAGAAGAAAAGGCAATGGCTGA
11655
Gallus_gallus                    11611 .............................................
11655

Anas_platyrhynchos_platyrhynchos 11656 CTACATTTAAATGCTTTACCGTGGGAAACCATTATTTCTAGGTAA
11700
Gallus_gallus                    11656 .............................................
11700

Anas_platyrhynchos_platyrhynchos 11701 GACGTTAATATACTGCCAGTAGAAGCAGCTGTTGTTTCCATGGTG
11745
Gallus_gallus                    11701 .............................................
11745
```

```
Anas_platyrhynchos_platyrhynchos 11746 ATATAGAGTCTCTACAATAGCTACACACTGTACAGGCCTTAACAA
11790
Gallus_gallus                    11746 .............................................
11790

Anas_platyrhynchos_platyrhynchos 11791 TAGAAATGTGATACAGAATCAGTCATCTGATGTTTTCATTTCAAA
11835
Gallus_gallus                    11791 .............................................
11835

Anas_platyrhynchos_platyrhynchos 11836 AGTTAAAAGTTCTTTACACATAATAAATGTGTTTTGTTTCTATTT
11880
Gallus_gallus                    11836 .............................................
11880

Anas_platyrhynchos_platyrhynchos 11881 AATTTTTTCTTCTCTTTATTTTATGAAGTTATGGTTTGAAAAACA
11925
Gallus_gallus                    11881 .............................................
11925

Anas_platyrhynchos_platyrhynchos 11926 GAAGTACCCTGAAAAGGAATGAGAGGCTACCTGAGATATGTGGGT
11970
Gallus_gallus                    11926 .............................................
11970

Anas_platyrhynchos_platyrhynchos 11971 CAGACCTAGTCTCAGAGCAGTACCCTCTTCAGTTTGAAGTCAAAA
12015
Gallus_gallus                    11971 .............................................
12015

Anas_platyrhynchos_platyrhynchos 12016 AACACTCATGAAGTAGATGGGGATGTTGGGGATCAGGGAAAGGTA
12060
Gallus_gallus                    12016 .............................................
12060

Anas_platyrhynchos_platyrhynchos 12061 AAACGGATACACAGTACAATAAATAAATAAATAAATAAATGATAA
12105
Gallus_gallus                    12061 .............................................
12105

Anas_platyrhynchos_platyrhynchos 12106 TGTCCTTAAACCCTGTTATCTTGCAAAGGCTGACCTCTTTATGAT
12150
Gallus_gallus                    12106 .............................................
12150

Anas_platyrhynchos_platyrhynchos 12151 TCCCATGCTCAAGAGGTGACGAAGCTGGAACGCTTTGTTGGGGGA
12195
Gallus_gallus                    12151 .............................................
12195

Anas_platyrhynchos_platyrhynchos 12196 AGAGGGACAGGCATAGGCAGTGTGGCCCTGCATCCACTCTGGCCC
12240
Gallus_gallus                    12196 .............................................
12240

Anas_platyrhynchos_platyrhynchos 12241 TGGCAAGCTCACATGGTTTGCTTGGCTGCTCTTTCCTGCTTGTGG
12285
Gallus_gallus                    12241 .............................................
12285

Anas_platyrhynchos_platyrhynchos 12286 CCCCTTGAAGCATTCACAATGATGGATGTGCAGAATAACCTAGCC
12330
```

```
Gallus_gallus                     12286 .............................................
12330

Anas_platyrhynchos_platyrhynchos 12331 CCCATCTCCCTGCACAGCCCAAGATTTCTGGTGCATTTTGCTTTA
12375
Gallus_gallus                     12331 .............................................
12375

Anas_platyrhynchos_platyrhynchos 12376 GCTCTGTGAAAGCAATCAATTTTAACATTGAAGCAGTAATGACAC
12420
Gallus_gallus                     12376 .............................................
12420

Anas_platyrhynchos_platyrhynchos 12421 TAGTATGTGGGAGACAGTATGAGCCAGCCTGCTCATATGGATGTG
12465
Gallus_gallus                     12421 .............................................
12465

Anas_platyrhynchos_platyrhynchos 12466 TTTCAGGAGCCCACTGTGTGCAACCTACTGCTTCTGAAGGCACAT
12510
Gallus_gallus                     12466 .............................................
12510

Anas_platyrhynchos_platyrhynchos 12511 ATTGGATTTACATGGCAAGGGTTTTGTAGCAGAGTGGCTGCAGGG
12555
Gallus_gallus                     12511 .............................................
12555

Anas_platyrhynchos_platyrhynchos 12556 GTGGGCTCTGTGAGAAGAGCCCATCAGCTGCGCCATGTCAGATAA
12600
Gallus_gallus                     12556 .............................................
12600

Anas_platyrhynchos_platyrhynchos 12601 GAGCCATCTCCAAAAGGGACCTGCTGCTGGCAGAGCCAAGCCAAT
12645
Gallus_gallus                     12601 .............................................
12645

Anas_platyrhynchos_platyrhynchos 12646 AAGTGCCATTGGATGCACCTCTGGGACAGCGTAGTTAAGAAAGGG
12690
Gallus_gallus                     12646 .............................................
12690

Anas_platyrhynchos_platyrhynchos 12691 AAAAACTTATTGCACAACAACAGCTGGGACAGAGAAAGGAAAGAA
12735
Gallus_gallus                     12691 .............................................
12735

Anas_platyrhynchos_platyrhynchos 12736 AATTATAGAGACAAACAATCCCACAGACGCCAAGGTCAATGAAGA
12780
Gallus_gallus                     12736 .............................................
12780

Anas_platyrhynchos_platyrhynchos 12781 AAGAGGTTAGGAAGAGCTCCAGGTGCTGGAGCAGATGTTACCCTT
12825
Gallus_gallus                     12781 .............................................
12825

Anas_platyrhynchos_platyrhynchos 12826 CAGCCTATGGAAAGGTCCACCGTGGAGCAGGCTGTCCACCTGTCC
12870
Gallus_gallus                     12826 .............................................
12870
CTGTATGTCCTTGCAACGTGCAGAGCTAAGGTCTATCATATTCTT
```

```
Anas_platyrhynchos_platyrhynchos 12871 ACCCATGGCTCAAGGGGTGCCACAGCAATGCAGATCTCCATGCTG
12915
Gallus_gallus                    12871 .............................................
12915

Anas_platyrhynchos_platyrhynchos 12916 CAAGCCACGGAGGAGCCCACATAGGAGCCGGGGGGGGGGGGGGGA
12960
Gallus_gallus                    12916 .............................................
12960

Anas_platyrhynchos_platyrhynchos 12961 AAAAGGGGGGGGGGGAGGGGGGGGGGAGGGTTCCACCTGTGGGGG
13005
Gallus_gallus                    12961 .............................................
13005

Anas_platyrhynchos_platyrhynchos 13006 ACCAGTGCTGGAGCAGTTTGCTGGAGATGGACTCCATGGTATGGA
13050
Gallus_gallus                    13006 .............................................
13050

Anas_platyrhynchos_platyrhynchos 13051 GCCATGTGGAAGCAGTTCTCGAAGAGCTGCAGCCTGTGGGCTTTT
13095
Gallus_gallus                    13051 .......................................GTTTTT
13095

Anas_platyrhynchos_platyrhynchos 13096 CAGATTCAGAGACCTTAACTTCATTTTATAATGGTTGGATCTAAA
13140
Gallus_gallus                    13096 CAGATTC------CCTGATTTCATTTTAGAGAGGTTAGATCTAAC
13140

Anas_platyrhynchos_platyrhynchos 13141 GAGGTGTGTTGCTCTACAAAATACATCAATTACATGAAATACATC
13185
Gallus_gallus                    13141
AACAGGAGTTACTCTACAAATAACTT------------------- 13185

Anas_platyrhynchos_platyrhynchos 13186 TACAAAATACATCAAGAGCTGCATGTATGAGAATTCAGTTCAAGG
13230
Gallus_gallus                    13186 ---------------GAGCTGCATGTGTGAGAATTCATTGCAAGC
13230

Anas_platyrhynchos_platyrhynchos 13231 CCGCTATGGGCTTTGCATTCATCTCATAAAATTTCATGAGGTGAT
13275
Gallus_gallus                    13231 CACCTGTGGTTCTGGTACTCATC-----AAATTTCATTAGGTGAT
13275

Anas_platyrhynchos_platyrhynchos 13276 TGTCTTTTACAGTATAATTTTACACTCAATGAGCTTGAACTTTCA
13320
Gallus_gallus                    13276 TGTCTTTTACAATATAATTTTAGCCTCTGTGAACTTAAACTTTCA
13320

Anas_platyrhynchos_platyrhynchos 13321 CAGTGCTCAATCTTTGAAACCAAAGTATGAACCTCATTATGTGAG
13365
Gallus_gallus                    13321 CAGTGCTCAATATT-AAACCCAAACTATAAACCTCATTATGAGAA
13365

Anas_platyrhynchos_platyrhynchos 13366 TTACATCATTT-CACAATTCCTGTTCTTTACCATTAATGCCATGT
13410
Gallus_gallus                    13366 CTACATTTTTTGCACAATTCCCGTTCTTTACCACTAACTCCATGT
13410

Anas_platyrhynchos_platyrhynchos 13411 TATCTAGATCAGAAAAATCAATATATG----AAGCTCTTCAAATG
13455
Gallus_gallus                    13411 TATCTATGTCAGAAAAGTCAATATATGTGTAAGGCTCTT--AATT
```

13455

Anas_platyrhynchos_platyrhynchos 13456 TTTTCAAAAT----------------ATTGTGAGACAGGAAGAGT
13500
Gallus_gallus 13456 TTTTTAAAATTATTATTATTATTATTATTATAAGACAGGAGGGTT
13500

Anas_platyrhynchos_platyrhynchos 13501 AAGATGTTCAATTACTATTCGTATGCATGCCTAAAAGAAAACGTT
13545
Gallus_gallus 13501 AAGATACTCAGTTATTATTTGTATGCATGACTTG-------TGTT
13545

Anas_platyrhynchos_platyrhynchos 13546 CATTAGTTTTTTAAAAAGTGAGGTATGATTTGAAAGCAAAAAATC
13590
Gallus_gallus 13546 CATTAGTTT------------------ATTTTAAA--AAAAAATC
13590

Anas_platyrhynchos_platyrhynchos 13591 TT-GCATTTAAGACTCAGATTTTCTCTTTCAAGCATATGTTGACC
13635
Gallus_gallus 13591 TTTGCTTTTAAGATTACCACTTTCTCTTTCAGGCATACTTTGACC
13635

Anas_platyrhynchos_platyrhynchos 13636 TAGGTTAATCACATGTTAATGGTATTAGCCATATTAAATCATGAA
13680
Gallus_gallus 13636 TAGGTTAATCACACATTAATGGTATTAGACATATTAAATCACAAG
13680

Anas_platyrhynchos_platyrhynchos 13681 GAATTGGCAGAAGTGATAGTCTTTAACTAAAAGCAACAGAAGTAT
13725
Gallus_gallus 13681 GAACTGGCAGCAGTGTTAGTATTTAATTAAAAGCAACAGAAGTAT
13725

Anas_platyrhynchos_platyrhynchos 13726 TTTATCTTCCAAGAATCTCAGGATATGTAGAGTATATTTCTG--T
13770
Gallus_gallus 13726 TTTATGTCCTGAGAATTTGAGGAAATGCATAGTATTTCTTTGCAC
13770

Anas_platyrhynchos_platyrhynchos 13771 AGGACAAAATGAAACATAAAAACTTAAACAATGGCAAAGAAAAAC
13815
Gallus_gallus 13771
ATATGAAAGTAAAGCATTAAAACTTAAGTTTCCATGAAG------ 13815

Anas_platyrhynchos_platyrhynchos 13816 AGTGTAGAAATTCATGGAAAACCTAAGAAAAAATGAAGTTAATAG
13860
Gallus_gallus 13816 ---------ATGCATGGAAAACCTAGGAAAA---TAATTTTAACT
13860

Anas_platyrhynchos_platyrhynchos 13861
GAATTTTGCCATCTGATAAG---------------TGTTACAT-- 13905
Gallus_gallus 13861 TATTTTTGCAATCTGATAATCATCTGTATGGGAACTGTTAGGTTG
13905

Anas_platyrhynchos_platyrhynchos 13906
CATCCT--------------------------------------- 13950
Gallus_gallus 13906 CAGCCTGAAGCACTGATTCACCACCTGGTGGCAGGACACAGCCAG
13950

Anas_platyrhynchos_platyrhynchos 13951 ------CAGTAC------AAGTCACTCAACCTGAAGTGGTATGGG
13995
Gallus_gallus 13951 CCCTGGCAGCACAGGTGGAAGCAATTCCACCAGAAGAGGTATAGG
13995

Anas_platyrhynchos_platyrhynchos 13996 TGAGTTTT------TACTTCACCAAGGAAAACTTCGGTTGCATCT

```
14040
Gallus_gallus                   13996 TGAATCTTTTAGAGTACTGCACCAAGGAAAACGTGGTTTGTATCT
14040

Anas_platyrhynchos_platyrhynchos 14041 GTAAATGAATTGCAAACACCAATTTGTAGCTGTAGTCAAAGACAA
14085
Gallus_gallus                   14041 ATACGTAAATTGTGGCCAGCAGTCTGTAATC-------AAGGCAA
14085

Anas_platyrhynchos_platyrhynchos 14086 TGTCTCCACTAGCAGAAATGGAAACACACCCACTTAATACCCTAA
14130
Gallus_gallus                   14086 TGTTTCCATT------------AACACGTTCACTTAATACATTAA
14130

Anas_platyrhynchos_platyrhynchos 14131 TAGCCATTATGAG-GTTAGCTGTAGGTCACAATACCATTGTGCAT
14175
Gallus_gallus                   14131 TAGATACTGTGAATTTTGGCTCTAGGTCACATTTCCATTGTGAGT
14175

Anas_platyrhynchos_platyrhynchos 14176 GGTGGGGTGGATGGTCTGACATCAAGCACTGAATTAAAGCAGTCA
14220
Gallus_gallus                   14176 GGTGAAGTGGATAGTCTGACATGAGGCTTTGGATTACTGCACTTA
14220

Anas_platyrhynchos_platyrhynchos 14221 AAACATTATAAGGGGATGGTATTCATATG--AGCTTTAGAGCAAT
14265
Gallus_gallus                   14221 AAACGTTGTAAGGGAA-GATATTCATAGAGCAGCTTTAGGGCAAT
14265

Anas_platyrhynchos_platyrhynchos 14266 GCTCTCATTTCAGCTATAGAGCAGACAAATAACATTGGCTACTTT
14310
Gallus_gallus                   14266 GTTCACATTTCAGCCAGAGAGCAGACGAACAACCTTGGCTACTTA
14310

Anas_platyrhynchos_platyrhynchos 14311 ATGAGGAAAAGAGAGAACTGCAGCACAACAGGAACAAAAGAGGAA
14355
Gallus_gallus                   14311 GTGAG--AAAAAAAGAACTACAATACAGTAGGAACAGAACAGGAA
14355

Anas_platyrhynchos_platyrhynchos 14356 TCAAACGAATGTATGGGAATACATTGAAAAGTCATGACATTAAGC
14400
Gallus_gallus                   14356 CTGTGAGATTATACGAGACTGAACT--AAAGTCATGACACTGAGC
14400

Anas_platyrhynchos_platyrhynchos 14401 CCTTCTTCCTTTTGCCCACTCATAAACAAGCACTAGAAAGAGAAT
14445
Gallus_gallus                   14401 CCTTCCTCCTTTTGGTTAGTCATAAACAAGTGCTAGAAAGAGAAG
14445

Anas_platyrhynchos_platyrhynchos 14446 TCAAGAATCTCTTCCCAAGGCATGTA------CCCATCCTGTCTG
14490
Gallus_gallus                   14446
GCAAGATTCTCT--CCAACTCATATAGAGGAGCCCATAATGTCT- 14490

Anas_platyrhynchos_platyrhynchos 14491 CAGAAGCCACCCAAAATACCTCCACAGTCAAAACTTCTTGCATGA
14535
Gallus_gallus                   14491 ----AACCACCT-GAATACCTC--CAGTCAGAAATCCT.......
14535

Anas_platyrhynchos_platyrhynchos 14536 CTTTTTTTCTAATGTGAAAATAATTTTCATTTATATATATATAAT
14580
Gallus_gallus                   14536 .............................................
14580
```

```
Anas_platyrhynchos_platyrhynchos 14581 GGAAAACATTATATATTTAATGGAAAACAATTTTGATATAATTTT
14625
Gallus_gallus                    14581 .............................................
14625

Anas_platyrhynchos_platyrhynchos 14626 AATGCACATTTAGAAAAAGAAAATTACTTAAATTTGTTTAGAAAT
14670
Gallus_gallus                    14626 .............................................
14670

Anas_platyrhynchos_platyrhynchos 14671 ACTAGCATTTTATGTACTTTTCGGTTATTAAAGGTGGATGATAAG
14715
Gallus_gallus                    14671 .............................................
14715

Anas_platyrhynchos_platyrhynchos 14716 GAAAAAAAAGTCACTACTTTTCTATACATTATTAGAGGCTTCAAA
14760
Gallus_gallus                    14716 .............................................
14760

Anas_platyrhynchos_platyrhynchos 14761 TTGTTGCTGTTCTCTGAATGATAATGTGCTGGCAGTATTCTTTTG
14805
Gallus_gallus                    14761 .............................................
14805

Anas_platyrhynchos_platyrhynchos 14806 GCACAGAAATATTTGTCAATGTGTTAGTTTTAAAATGAAAGGGAA
14850
Gallus_gallus                    14806 .............................................
14850

Anas_platyrhynchos_platyrhynchos 14851 TTTTAATCACATTTTCTGTTAGCCCTCAACCATTCTGCTTTTACT
14895
Gallus_gallus                    14851 .............................................
14895

Anas_platyrhynchos_platyrhynchos 14896 CAGGTCACTAACCAAAGGTGTGACAGTAGTATTTAGACATACAGA
14940
Gallus_gallus                    14896 .............................................
14940

Anas_platyrhynchos_platyrhynchos 14941 AGATAAGAAGTGTTTCCAAACATCAATTCAAATTTAGAGAAATAT
14985
Gallus_gallus                    14941 .............................................
14985

Anas_platyrhynchos_platyrhynchos 14986 TCTACAGATCCAAAATGGCACCAAAGACAATATTTATGAAAAAGC
15030
Gallus_gallus                    14986 .............................................
15030

Anas_platyrhynchos_platyrhynchos 15031 ACATTGAATTCCTAAGTGAGGTGCAGGGCATGGCCACCTCCCCCA
15075
Gallus_gallus                    15031 .............................................
15075

Anas_platyrhynchos_platyrhynchos 15076 GCTCCGGCGCAGCCCTGCTCCTGTCTGACCACTGGGGCCTAGCCC
15120
Gallus_gallus                    15076 .............................................
15120

Anas_platyrhynchos_platyrhynchos 15121 CATGTCCAGTGACCCACAGTTTATGCCCAGGCTGGAGGTCATGGA
15165
```

```
Gallus_gallus                    15121 .............................................
15165

Anas_platyrhynchos_platyrhynchos 15166 TGCCACTGTGTCCTGAGCAGGTTCAGCCCATAGCAAAGCTGAGCA
15210
Gallus_gallus                    15166 .............................................
15210

Anas_platyrhynchos_platyrhynchos 15211 AATATGCCAGAGACACATGGGGCACTAACACACCAACTGCCCTGC
15255
Gallus_gallus                    15211 .............................................
15255

Anas_platyrhynchos_platyrhynchos 15256 CTTCAGCAGCATTTACACACAGGACTCACGTAAAACAGAGATCGG
15300
Gallus_gallus                    15256 .............................................
15300

Anas_platyrhynchos_platyrhynchos 15301 AGTCCCCTTCCTCCTTTCTGCCTGATAGAGACAGAAGGTTAATAG
15345
Gallus_gallus                    15301 .............................................
15345

Anas_platyrhynchos_platyrhynchos 15346 TGTGGCCACACACACAACATTCTGGGTTCACAAGCACACACACAT
15390
Gallus_gallus                    15346 .............................................
15390

Anas_platyrhynchos_platyrhynchos 15391 TCGTAAATCCAGCACAGCCCCTCTGACTGCCAAGCATCACTGCCC
15435
Gallus_gallus                    15391 .............................................
15435

Anas_platyrhynchos_platyrhynchos 15436 AGATTCTAGGCCCCCTACCTGCTTCATGGCTCACCAGGTCACCAA
15480
Gallus_gallus                    15436 .............................................
15480

Anas_platyrhynchos_platyrhynchos 15481 CTGCTCCAGCTTAAAGTTCACAGGTCCCTCAAGAACTGGCACTGA
15525
Gallus_gallus                    15481 .............................................
15525

Anas_platyrhynchos_platyrhynchos 15526 ATTAACGCAAACGCATGCATAGGTCTCACCAGTAGCTGGCACATA
15570
Gallus_gallus                    15526 .............................................
15570

Anas_platyrhynchos_platyrhynchos 15571 GTCCTTGCAGCATGCACACAAGGGAGAGAAAGTGGGTATGCACAG
15615
Gallus_gallus                    15571 .............................................
15615

Anas_platyrhynchos_platyrhynchos 15616 AAGGATTGTTAAGAAAAATTTAATGAGGAACAGTAGGAAGAAAGA
15660
Gallus_gallus                    15616 .............................................
15660

Anas_platyrhynchos_platyrhynchos 15661 TCAGACTGTAGTGACCAGGCACAGGGCTCAGCCAGGTAAGTGTAC
15705
Gallus_gallus                    15661 .............................................
15705
```

```
Anas_platyrhynchos_platyrhynchos 15706 CGACCAGCCTCATTTTGCACATAACCAGCCTTTTTATACCCTTTT
15750
Gallus_gallus                    15706 .............................................
15750

Anas_platyrhynchos_platyrhynchos 15751 CTACCTGTCCCCCTCTTTGTTCCTCCTTCTCCCTCTTATCCCTGC
15795
Gallus_gallus                    15751 .............................................
15795

Anas_platyrhynchos_platyrhynchos 15796 AGACCCCCATGGGTCATCTGCAGTTTCGGTACCCTCCTAGTTGAG
15840
Gallus_gallus                    15796 .............................................
15840

Anas_platyrhynchos_platyrhynchos 15841 GACCACCTACTTCACAGTTAGATCAGTTGCAAAGTCCAGGTTGTT
15885
Gallus_gallus                    15841 .............................................
15885

Anas_platyrhynchos_platyrhynchos 15886 TCTCCTGGAAGCGATGCCTGTAACTTGATAGCCAACCAGCTAGTG
15930
Gallus_gallus                    15886 .............................................
15930

Anas_platyrhynchos_platyrhynchos 15931 AGGCTAGGATGTTTATATCTGGTAATTGTCTCCCAGTTTAGAAGT
15975
Gallus_gallus                    15931 .............................................
15975

Anas_platyrhynchos_platyrhynchos 15976 TCTCAATCAGGTACTCCTGCTGGAATAAATGCTTCTGCATTCTCT
16020
Gallus_gallus                    15976 .............................................
16020

Anas_platyrhynchos_platyrhynchos 16021 CCTGGTCTTACCAATTACTGTGACTGACCAGGGTTGGTTCCCATC
16065
Gallus_gallus                    16021 .............................................
16065

Anas_platyrhynchos_platyrhynchos 16066 ACTGCTTCCCCATCTTATTATCCCTTGTTAAATTGGAAAATGTCC
16110
Gallus_gallus                    16066 .............................................
16110

Anas_platyrhynchos_platyrhynchos 16111 TTGACTACATACACTTAAAGAAGCACTCTCCTTCTACATACCTTT
16155
Gallus_gallus                    16111 .............................................
16155

Anas_platyrhynchos_platyrhynchos 16156 ACATAAATATGATAAAGATGGTAACATTTTCACTTAACGAAGGCA
16200
Gallus_gallus                    16156 .............................................
16200

Anas_platyrhynchos_platyrhynchos 16201 GTGACCATGAGTCAGGAATAGTTAGGACAGAACTATGTAGACTCC
16245
Gallus_gallus                    16201 .............................................
16245

Anas_platyrhynchos_platyrhynchos 16246 TGCCTGTGAAGACAAGAAACTTCTACATACATTGTGATAGGTACA
16290
Gallus_gallus                    16246 .............................................
```

```
16290

Anas_platyrhynchos_platyrhynchos 16291 GTGGATGTCAAAGAGGTATGGCTCCATTAAAACAGGGGGTATGAT
16335
Gallus_gallus                    16291 .............................................
16335

Anas_platyrhynchos_platyrhynchos 16336 TGCATCAGGTTTAGGAACATTGTAGAAGTGGCTGGGGGGGGGGGG
16380
Gallus_gallus                    16336 .............................................
16380

Anas_platyrhynchos_platyrhynchos 16381 GGGGGGGGGGGGAGGGTTTAGAAAAAAAAATTTTAGATATTGCCA
16425
Gallus_gallus                    16381 ..........................AAAGTTTTAGCTTTTGTGA
16425

Anas_platyrhynchos_platyrhynchos 16426 GAATATCTGTGCCACAGTGCAATAAAGATGGGTCCCCAGGACCCA
16470
Gallus_gallus                    16426 GAAAATCTGTTCAAAAGTGGAATGAAGATGACTCCCCAGACCACA
16470

Anas_platyrhynchos_platyrhynchos 16471 GAAAACAGTCTTGTCAT---TGATAACAGGGTGGTGGGAGAAAGA
16515
Gallus_gallus                    16471 GGAGACAGTCTTGTTGTGGATGATTACAGGTTGATGGGAGAAAGG
16515

Anas_platyrhynchos_platyrhynchos 16516 AAGAGCAGTAGCCATGGAAAAATCTCAGGAGAGAGGATCGGAAAT
16560
Gallus_gallus                    16516 AGGAGTAGCAGCTATGGAAAAATCTCAGGAAAGATGATCAGAAAA
16560

Anas_platyrhynchos_platyrhynchos 16561 ACAGTTTTGGACGCACTACTTTGAAAATAATTGTGAAACATTCAG
16605
Gallus_gallus                    16561 ACAGTTTTGGATACACAACATTGAAAATAAGTGTGAAATATTCAG
16605

Anas_platyrhynchos_platyrhynchos 16606 AAAACAGAGGCTGGTTGAAACACAAGGCTAAGAATTCTGGAGACA
16650
Gallus_gallus                    16606 AA--CAGAGGCTGGCTGAAACAAATGGTTAGGAAATATGGAGATA
16650

Anas_platyrhynchos_platyrhynchos 16651 CATTAGAAATAAAAATCAAGATTTGAATACTACCCTTACTAACTA
16695
Gallus_gallus                    16651
CATCAGAAGTAAATGCCAAGATTTGTATACC-------------- 16695

Anas_platyrhynchos_platyrhynchos 16696 AGGGTAGTAAAGGAGTCCACATGGTTGGAGTCAGTTGCTTAAGTT
16740
Gallus_gallus                    16696 --GTTAGTATATGAACCCACACAATTGGAG------GCTTAGGTT
16740

Anas_platyrhynchos_platyrhynchos 16741 ATGCAGGAAAAAGTGGTTTGAACTAAGAGGACAGGGAGAAAGTTT
16785
Gallus_gallus                    16741 ATGCAGGAGAAAGAGG-----ACAGGGAGAACAAG-AGAAACTCT
16785

Anas_platyrhynchos_platyrhynchos 16786 CATGCCCTGAGT-GACTGGTCAAGCAGTTGTATGATAAGAAAAAC
16830
Gallus_gallus                    16786 TGTGCACTGAGCAGACAGGTCAAGCAGTTAAATAGTGAGAAAAAC
16830

Anas_platyrhynchos_platyrhynchos 16831 AGTATATAGTACTGAAAGATGCAGGACAGGATGTTAGAATGGTGG
```

16875
Gallus_gallus 16831 AG-------------------CAGGGTGTGATGTTAGGATGGAAA
16875

Anas_platyrhynchos_platyrhynchos 16876 ACATGATAAGTACCACAAAAAACAAGCAGAAAACATAATCTGGAG
16920
Gallus_gallus 16876 ACATTAACAATACCACAAAAGATAAGTGGAAAACA--ATCTGGAA
16920

Anas_platyrhynchos_platyrhynchos 16921 AGCTAGATTTATAGATGTTAGCACACATTTGCTCAGAAGCAATTT
16965
Gallus_gallus 16921 CGCTAGGTTTATAGATGTTA-CAGGTATTTGCTGAGAAGCAATTT
16965

Anas_platyrhynchos_platyrhynchos 16966 ATCAGGTTGCAAAAAAA-GAAAGCCTGAGAGAAGTGAGCAGAATC
17010
Gallus_gallus 16966 GTCAGGATGAAGAAAGGTGAAAGTCCGAGAGGAGAGAGTGAAATC
17010

Anas_platyrhynchos_platyrhynchos 17011 TGATCCATAGAAGGGAAGACATGGTAAATGAGGTGCCACTGACTT
17055
Gallus_gallus 17011 TGAGGCATAGAAGTAGAG-TATGGTTGATGAAGTGCCACTTATTT
17055

Anas_platyrhynchos_platyrhynchos 17056 TCAGGAGTTTGAAAGACACAAGATGATGGTTTGGGATGGTGTAGT
17100
Gallus_gallus 17056 CCAGGAGAGTGAAAGACATGAGATGGTGGTTAGGAGTGGTGGTGT
17100

Anas_platyrhynchos_platyrhynchos 17101 GCTCCCAGCAGCTTCCTGGAGAGACCTATTTTTGCATTGTCAAGG
17145
Gallus_gallus 17101 -CCCCCAACAGCTTGCTGAAGAGA--TATTTTTTCATTGTCAAGG
17145

Anas_platyrhynchos_platyrhynchos 17146 GAAGTAAGCAGAGAAGCAAGACAAAGTGAGGTAAAGAAGGCATAT
17190
Gallus_gallus 17146 GAAGTAAGAAGCAAAGCAGAAGAAATTGAAGCAAAGAAAGCATAT
17190

Anas_platyrhynchos_platyrhynchos 17191 AAGAGAAATAGGGATAAGCAAGTATTAATTGTCTGCTAGAAGCAG
17235
Gallus_gallus 17191
AAGAGAAATAG---------------------------------- 17235

Anas_platyrhynchos_platyrhynchos 17236 TGGGATTACATGAAGGTATCACTAATTTATTTTCAACATAGAAGT
17280
Gallus_gallus 17236 ------------AAGCTAGCACTCATTCA--------------GT
17280

Anas_platyrhynchos_platyrhynchos 17281 TCATCATGTAATTAAGTATGTTGATC--CTGAGCAGGATCTAAAT
17325
Gallus_gallus 17281 TCATCATGTAATCAAACATTAAAAACACCTGAGCAAGATCTAAAT
17325

Anas_platyrhynchos_platyrhynchos 17326 GACACA-----------CATCTGCACCTGTGTTCCTCAGGTGAAG
17370
Gallus_gallus 17326 GACATGTAAGCGCTCATCATGTACACCTGCGTTCCTTGTGTGAAG
17370

Anas_platyrhynchos_platyrhynchos 17371 CTCTATCATGTGGTCAGAATTTATGCAGAAACAGTTCATGTCTAT
17415
Gallus_gallus 17371 CTCTATCTCGTGTTCAGAGCTTATGCCAAAACAGCTCGTATCCAT
17415

Anas_platyrhynchos_platyrhynchos 17416 TTCTCCATGTAGCTCATGGCCTAATTACCTCTGTTACCATTTGAA
17460
Gallus_gallus 17416 TTCTACATGTAACTCATGGCCTAGCTACCACTGCTAATATTT-AA
17460

Anas_platyrhynchos_platyrhynchos 17461 TTGCAATCCTTTCACATTAGCTCCTCTAGAACACCTTAATAGATT
17505
Gallus_gallus 17461 TTGCACTCTTTTCACACCAGTTCCTCTAGAACACTTCAATAGTTT
17505

Anas_platyrhynchos_platyrhynchos 17506 ACATTTATGGTTGTTTTGTATTATCTTGTCACTCCTTTAGACAAA
17550
Gallus_gallus 17506 ACATTTACAGTTGTCTTATATCATCTTATCTTTCTGTTAGACAAA
17550

Anas_platyrhynchos_platyrhynchos 17551 CTCTCTGACATGCAGCAAGCATAGATAT-AAAATGTAGGCTGCAA
17595
Gallus_gallus 17551 CGCTTTGACGTGCAGCAAGCATAGACCTGAAAATGTAGGCTGCAA
17595

Anas_platyrhynchos_platyrhynchos 17596 TAGGTTGATATGAAAATTAGTGCTTCCGGGCAAATAGGAAATTAA
17640
Gallus_gallus 17596 CAGAATGATATAAAAATTAGCGCATCTAGGAAAACAGGAAATTAA
17640

Anas_platyrhynchos_platyrhynchos 17641 TATTAGAATCCAATTAATCTTCATTTCATATGGAGAAAAAAAATA
17685
Gallus_gallus 17641 TATTAGAATTTAATTAATTCTTCTTTCATGTGGAGAAGAAAT-TA
17685

Anas_platyrhynchos_platyrhynchos 17686 ATTAGTCTTAATAGGACCACAGAGGCATCACAA-TTTTTTCCTTC
17730
Gallus_gallus 17686 ATTAGACTTAATAGGACCATGGAAGCATCACAAATGTTTTCTTTC
17730

Anas_platyrhynchos_platyrhynchos 17731 CATCAAACAAAGCCAACACTGAACAACCACTAATCAATCACTCTA
17775
Gallus_gallus 17731 CATCAAATAAAGCCTACACTGAACAACCACTAATTAATCATTCTG
17775

Anas_platyrhynchos_platyrhynchos 17776 ATCAAGCACAGCATCAGCAGTATTTCTCACTGTCCATATCTCTTA
17820
Gallus_gallus 17776
ATAGAGCACAGTACCAGCAGT---TCTCATTGTCCATAACTCCT- 17820

Anas_platyrhynchos_platyrhynchos 17821 TCTTTTTACTTCGTTC--------------GGAAGTATTTTCTAA
17865
Gallus_gallus 17821 -CTTTTTATTTCCTCCATCCCCAATTCCCTGCAAGTATTTTCTAT
17865

Anas_platyrhynchos_platyrhynchos 17866 CAAGGACAAACTTTTTCTTGTTTGATTATTCTGAATACTCCCACA
17910
Gallus_gallus 17866 CAAGGGCAAGATTTTGCTTGGTTTATTATTATGAAAATTCCCATA
17910

Anas_platyrhynchos_platyrhynchos 17911 ACTGCAGGAAATGCCATTTTTTTTTTGAACAACGAATTAATTCTT
17955
Gallus_gallus 17911 ATTGTGGGAAAATCCATTTTCTT---GAGCAATGAATTAGTAATT
17955

Anas_platyrhynchos_platyrhynchos 17956 ATCTTCTGCAAGCATGAACAAATGTTTTTTTTGTAGATGTTTACC
18000

```
Gallus_gallus                    17956 ACCATCTGCACATACAAAC-----TTTTTTTTGTAGATGCCTACT
18000

Anas_platyrhynchos_platyrhynchos 18001 AATCCTCAAGCATTATCATTCCCTGTCTCAGGTAGGACACTGTCT
18045
Gallus_gallus                    18001 GATGTTCTAGCATTAATATTCTGTGTCTTAAACATGATATTTTAG
18045

Anas_platyrhynchos_platyrhynchos 18046 CTACTAGAGGTTCTGA-----------ATAAGGAATAATGGAATC
18090
Gallus_gallus                    18046 CTACTAAAGGTTCCTATCTTTGATACAACAAGGAACAATAAAAAC
18090

Anas_platyrhynchos_platyrhynchos 18091 AGTGTGAGTTTTCATGTTAAAATATCCTCGCTACTATTTATAACA
18135
Gallus_gallus                    18091
TTTGGGAGTTTTTATGTTAAAATATCCTCGGTATCATTT------ 18135

Anas_platyrhynchos_platyrhynchos 18136 ACTTTTCAGGAAGACACATTTTTT-TTTCTGCCTAT-----TGGT
18180
Gallus_gallus                    18136 ------CAGGAGGACACTTTTTTCCTTTCTGCTAGTCAGAGTGGT
18180

Anas_platyrhynchos_platyrhynchos 18181 TTAAAAAAAAAAAAAAGTTATCATCTTGATACATTGGTTGTAGTC
18225
Gallus_gallus                    18181 TTTTTAAAACAT----GTTTACATCTTGATATATAGCTCGTAGTC
18225

Anas_platyrhynchos_platyrhynchos 18226 CAACTAAACAATATTAGCCTTCACATACAGTCTTAGAGAGTTTCT
18270
Gallus_gallus                    18226 CAATAAAACAATATTAGCCTTGATAGAAAGTGTTATAAAGTTTGT
18270

Anas_platyrhynchos_platyrhynchos 18271 AGAAGTTGACTGTTTTTTAATGTAACAATAACAGAAATTTAGAAT
18315
Gallus_gallus                    18271 AGAAG------GTTTTTCAATGTAACAATAACAAAAATT-AGAAT
18315

Anas_platyrhynchos_platyrhynchos 18316
TGCAATCAA-----------GGGGGGATGCCCCTTTTTTTTT--- 18360
Gallus_gallus                    18316 TACAATAAAAATGGGTGCATGGGGTGGTGTTTTTTTGTTTTTGGT
18360

Anas_platyrhynchos_platyrhynchos 18361 ------TTTT----TTTTTTTTTTA-------------TAAAAAG
18405
Gallus_gallus                    18361 GGTTGTTTTTGGGGTTTCATTTTTAGTATGATAGTCATTGAAGAG
18405

Anas_platyrhynchos_platyrhynchos 18406 TAAGTTATGCCATGAATGGTATTGATAATCTTA-TAATTTCATGA
18450
Gallus_gallus                    18406 TAAGTTATGCCATGAATGCTGTTAATGATCTTAATGATTTCATGA
18450

Anas_platyrhynchos_platyrhynchos 18451 CTAGTGTGTATGACTGCTGCTGAAGGCAAGGTGCAAGGTGCTCGC
18495
Gallus_gallus                    18451 CTGGAATGTATGACTGCAGCTGAAAACA---------GTGTACTC
18495

Anas_platyrhynchos_platyrhynchos 18496 ACTTGATTGTTGATTAATAAACTGCTTCTCGTGTCCTTGAATTTT
18540
Gallus_gallus                    18496 AC---ATTGTT--TCAATAAACTGCTTCTCATGTCCTTGAATTTT
18540
```

```
Anas_platyrhynchos_platyrhynchos 18541 TCTTTTGCTACCAGAAGTATGCCTAGAAATTATATTATAATCCAT
18585
Gallus_gallus                    18541 CATTTTACCACCAAGAGCATGCCTAGAAATTACATCAATTTCCAT
18585

Anas_platyrhynchos_platyrhynchos 18586 CAATTTACTTGTTTATAATGTAAATAGCACTTTTTTTTCTTTTTT
18630
Gallus_gallus                    18586
CCATTGATTTGTTTACAATGGAAATAGCACTTGTATTCTTTT--- 18630

Anas_platyrhynchos_platyrhynchos 18631 TTCTGAAAATAAATTATGGTTGCAGAAAGAGCTACTGTTTTGCAT
18675
Gallus_gallus                    18631 ----GAATATAAATTACAGCTATAGTAAGAGCTACTATTCTGCAC
18675

Anas_platyrhynchos_platyrhynchos 18676 TGAAGTGGTGCTGTTTCCTTTGGGATACTACTACAAAAGGTACTA
18720
Gallus_gallus                    18676 TGAAGTGGTACTGGCCCCTTTGGGTTAT--CTGTAAGAAACACCA
18720

Anas_platyrhynchos_platyrhynchos 18721 ACACTCAAGGCTGGCTAGCTGAT--TTACTTAGGAAAGGATGGAA
18765
Gallus_gallus                    18721 GTGCTGAAGGTTGGCTAACTGCTAGTTACTGAATAAAGAATAGAA
18765

Anas_platyrhynchos_platyrhynchos 18766 ATTAATTATATAAAAGATGATAGCACTC-TTATGACCCTTCTGAT
18810
Gallus_gallus                    18766 ATTAG------AAAAAAAGAGAGCACTCATCATGCCCTTCCTGAT
18810

Anas_platyrhynchos_platyrhynchos 18811 GATGGGAACCCAGCTGTTGCAGAATGGAACAACCTCCTCCTTGTA
18855
Gallus_gallus                    18811 TCTGGGAAACTATCTGTGGCAAAACAGAAT---CTTCTCAGTGTA
18855

Anas_platyrhynchos_platyrhynchos 18856 AATCATAGTATCGTGGAATCATGCTACACTGATCTAGAATCATAG
18900
Gallus_gallus                    18856
GATCATGGCAT---------------------------------- 18900

Anas_platyrhynchos_platyrhynchos 18901 ATCATACTACACTGACCAGATCCTGCTCATGCCGAGGTTTGCGGG
18945
Gallus_gallus
18901 ----------ACTGTCCAGGTTTGGTCTGTGGTAAGATTTGC--- 18945

Anas_platyrhynchos_platyrhynchos 18946 CCTCACTGTTCTTCATATATTTTTAGTATTCCTCCAAATAAACAG
18990
Gallus_gallus                    18946 -----------------------------CCTTCTAAGAGTACAG
18990

Anas_platyrhynchos_platyrhynchos 18991 TAATCAACCTTTTATCTATGCAATGCTGAGCTTACTAGTTGACAC
19035
Gallus_gallus                    18991 TAATCAACCCTTCACTGGGACAAGGTTGAACTTTCTAGTTTGCAC
19035

Anas_platyrhynchos_platyrhynchos 19036 AACAAAGTTCACTGAGAGTGTTGCACTCTTACATATCTCCATAGG
19080
Gallus_gallus                    19036 ATGAAGATCCACTGAGACCGTTACACTCTTTAATATTTCTATTTG
19080

Anas_platyrhynchos_platyrhynchos 19081 AATGAGCAATTATAATATATATAATATATTGTATATATTATTATA
19125
Gallus_gallus                    19081 AGTGAGCTCTTAGAA---ACATTACATCCAGTAGGTATTGCTAAA
```

19125

Anas_platyrhynchos_platyrhynchos 19126
TATAATATATAATA----TATTATATAT--------TATATA--- 19170
Gallus_gallus 19126 GGCCATGTGTATTAATGTCACTGCGTATGCTCAAGCTATATAGCG
19170

Anas_platyrhynchos_platyrhynchos 19171 ------------TCGTATAT--------------------TATAA
19215
Gallus_gallus 19171 GGCCACTTGCGCTCATGTATGGGTATCTACTGCCAAGGCCTGTAC
19215

Anas_platyrhynchos_platyrhynchos 19216 ATAAATATAATAATTATAGTAATGCTAGAACTTCAGTGTGAAAAA
19260
Gallus_gallus 19216 TTAAATGTTGCATTTATAGTTAAGCCAGCATTTCAGGATGAAAAA
19260

Anas_platyrhynchos_platyrhynchos 19261 AAATCTGTTCTCCTGCCTGAGAAATAAAAATAAGTTCCTGGCTGA
19305
Gallus_gallus 19261 AT-TCTGTTCTTATGCCCAAGTAACAAAAGTAAG---CTGAGTGA
19305

Anas_platyrhynchos_platyrhynchos 19306 ATTTTGGCACACAGGAAAAAAAAAAATAAAAATCTTTCATAATCAG
19350
Gallus_gallus 19306 TTTT---AATACAGAAGAAAAAAAAAAAAAGCCTTTTTTTAATGAG
19350

Anas_platyrhynchos_platyrhynchos 19351 TTATTTCAGCATTGGGAATTTAAAATTCTGGATCAATACAACTGT
19395
Gallus_gallus 19351 --AATTTAAAATGGGCACTTTACACTTCAGGATCAGTA--ACTGT
19395

Anas_platyrhynchos_platyrhynchos 19396 AAGAATCATTTTTACTGATAAAATTCTGAAATTTAGAAGGGCACT
19440
Gallus_gallus 19396 CAGAAGCATTTTCATGGACAAAACAGTGCAGTTCAGAAGGGGACT
19440

Anas_platyrhynchos_platyrhynchos 19441 ATCCAAGCTATTTTCCATAGATTCAAATCTAGAGGGGGTTGGAAA
19485
Gallus_gallus 19441 ATCCATGCTATTTTCCATAGCCTGAAATCTAGACATGGTTGGAAA
19485

Anas_platyrhynchos_platyrhynchos 19486 AATATATTTTAAAATGTCATCACACCTCACTAGCCACTAGCAGGG
19530
Gallus_gallus 19486
AGTATGTTTTAAAGTGTCATCACACTTCACTGCCAACTAGCAGG- 19530

Anas_platyrhynchos_platyrhynchos 19531 AAAAAGGTTTTGGTGAGGGCTTAGCTACTGAATGATGTATTGGTG
19575
Gallus_gallus 19531 -------TTTTGGTGAGTACCTAGCAACTGAATG---TGTTGGTG
19575

Anas_platyrhynchos_platyrhynchos 19576 CACTTCAAAAAAGTTGTCATTGTCTTGCTCCAGCAACAGACAGCA
19620
Gallus_gallus 19576 CTCTTCAAAAAGGTTGTCATAGCTTTGCTCTGACATGACACGACA
19620

Anas_platyrhynchos_platyrhynchos 19621 CAGATTC---CCTAGAGATTTGCTGCTGCTTTTCAAAGATACAAA
19665
Gallus_gallus 19621 TAAACTCTAGCCAAGAAATATACTACTGATGTTCAAAGATACAAA
19665

Anas_platyrhynchos_platyrhynchos 19666 TATATTAATATGCAAATTTAGTTTCCTCTAAAATCACATGCAAAT

```
19710
Gallus_gallus                     19666 TGTCTTA---TACAGATTTTATTTCCTCTAAAATCACATGCAACT
19710

Anas_platyrhynchos_platyrhynchos 19711 ATGCAGAGTGCTGCTACAGGAGATACTACCTGCTCAAAGTGAAAT
19755
Gallus_gallus                     19711 GTGCACAGCAGTGCTAGAGGCGGTACT----GCTCCAAGTGAAAG
19755

Anas_platyrhynchos_platyrhynchos 19756 GGCATGGAGAAGGGCAGGTATTCTTATTAAGGACTAGCTCTACCA
19800
Gallus_gallus                     19756 GGCCTGGAGCCATTCAGGGATTCCTATCAAGGACCAGCTCTACAA
19800

Anas_platyrhynchos_platyrhynchos 19801 AACATTCATAATAAACTGGATCATTTCAGCATAGCTCTGCAAGTA
19845
Gallus_gallus                     19801 AACATTCATCAGAAGCAGGATGGTTCAAGCATAGCTCTGCAAGTA
19845

Anas_platyrhynchos_platyrhynchos 19846 G-GGAGATGGAGATTAACC-------TCATTCATCCATTTGCATT
19890
Gallus_gallus                     19846 GAGGAGATAGAAATTAACCATCTGTCTAATCCATCCTTTTGCATC
19890

Anas_platyrhynchos_platyrhynchos 19891 CTATGGTATTTAGACAGCTCCTTCTGCAAATTTTTTAACTTATCT
19935
Gallus_gallus                     19891
ATATGATAATTAGGTTGATGCTTTTGCAAAT-------------- 19935

Anas_platyrhynchos_platyrhynchos 19936 TCTTAACTTCTTAACCTTTAACTTAACTTCTGCAAATATTTAACT
19980
Gallus_gallus                     19936 -----------------------------------ATATTTACCC
19980

Anas_platyrhynchos_platyrhynchos 19981 TAATGTTTTTTGTTTGTTTGTTTGTTTTTTGTTTCTTTGACTTGA
20025
Gallus_gallus                     19981 T............................................
20025

Anas_platyrhynchos_platyrhynchos 20026 GTTGTTTTATTGGTTTTGGTTTTAGTTTTGGTAAAAATGCATTTA
20070
Gallus_gallus                     20026 .........................TTTTTCAAATAATGCATTTA
20070

Anas_platyrhynchos_platyrhynchos 20071 ATCGCATCAGTTTCCACTTTCAAGAAACCAGACCTTGTCCTTGCT
20115
Gallus_gallus                     20071 ATTACTTCAGTTTCCACTTTCAAGTAACCAGACTTTGTCCTTGCT
20115

Anas_platyrhynchos_platyrhynchos 20116 CACACTCAGTCTCCATTGTCAAGAGCTCTAAAAGGTAAAGAGAGG
20160
Gallus_gallus                     20116 GACAATC--TCTCTATTGTCAAAAG------------AAAAAAAG
20160

Anas_platyrhynchos_platyrhynchos 20161 CAACTAGCATTCAAAACAAGTTTGATAAATACATTTCTCCGTGTT
20205
Gallus_gallus                     20161 CAGGAAGTAATCAAACCAAGTTTGACAAATCCATTTCTCACTGCT
20205

Anas_platyrhynchos_platyrhynchos 20206 CCCAAATTTAGCTGCAGCATAAATTCCTTCTTTTG----TTTCCT
20250
Gallus_gallus                     20206 CCCACCTCTAGTTGCATCATTAATTCCTGCTTTTAATTTTTTCCT
20250
```

Anas_platyrhynchos_platyrhynchos 20251 GTCTTTATCTTTCATTTACTGCTTCTCCAGGATAAGATCAACCCT
20295
Gallus_gallus 20251 TTCTTC-TCTTCCATTTACTGTTTCTCCAGGCTCAGACCAAAACC
20295

Anas_platyrhynchos_platyrhynchos 20296 CCACTCCCACTCCCCTTCCCCAAACAAAACACCTCTGATGTTCCC
20340
Gallus_gallus 20296 CC---------------CCCCAAATAAAAAACTTCTGATGTTCTC
20340

Anas_platyrhynchos_platyrhynchos 20341 CTTCATCTTGTTCATACCAGCAGA-TCCATGACCTCACTCAGAGG
20385
Gallus_gallus 20341 CTTCAGCTTGTTCTTACCAGAAAAATCCATGACTCCACACAAGGG
20385

Anas_platyrhynchos_platyrhynchos 20386
AAAACTTTCCTCACAAAAAAATACACTACCAAGATA--------- 20430
Gallus_gallus 20386 AGATCTTTCTTCACCACAAA-TACACTGTTAAAGGAAGGAAGGCT
20430

Anas_platyrhynchos_platyrhynchos 20431 TTGGTGTCCAAGGATCCCT------------------------TC
20475
Gallus_gallus 20431 TTGAT-----AGGATCCTTGTCCAAGCCTAAGTAGTTTGAATCTC
20475

Anas_platyrhynchos_platyrhynchos 20476
AGTGAGATGGGCTGAT-CTGTCCAAGCCT---------------- 20520
Gallus_gallus 20476 CGTGATATTTCCTGTCGTTGTTCCAGCCTGACTGTTCCTACAGAT
20520

Anas_platyrhynchos_platyrhynchos
20521 --ATGTAGGTTCAGCA--------TTCACTACGGTTTGAA----- 20565
Gallus_gallus 20521 ATATGCAGTGCAGGCAAGTTTGTTCTCAGTCTAGCTAGAAGAATA
20565

Anas_platyrhynchos_platyrhynchos 20566 --------------------------TCTCTGTTTAATTTTGACC
20610
Gallus_gallus 20566 AAATACACACTCATCGGCTAAAACCTTTTCTTTTTAATTTTGACC
20610

Anas_platyrhynchos_platyrhynchos 20611 CCAGTGAAAATGATTGTCCAAGGAAACAGAGCCATGAATATCAAC
20655
Gallus_gallus 20611 TCAGCAAAAATGATTATCCAAGAAAACAGAACCATGAACATCAAC
20655

Anas_platyrhynchos_platyrhynchos 20656 AATAATTCTCATCTCAATTAGTAATAAATATAATTTGTAACTGAT
20700
Gallus_gallus 20656 ATTAATTCTCATCTCAATTACTAAGAGTTATAATCTGTAACTGAT
20700

Anas_platyrhynchos_platyrhynchos 20701 TAGTGAAGATTCTTATAAAATTTATGATG----------TTATAA
20745
Gallus_gallus 20701 TAGTGAAGATTCTTAGAAAATTTATGGTAGTGTCCTTTATTATAA
20745

Anas_platyrhynchos_platyrhynchos 20746 TACTTTGTATTATAAATTGTACTATTCTTT--CTCTTTACTACAT
20790
Gallus_gallus 20746 TGCTCTGTATTATAAATCGTACTATTTTTTTTCTCTTTACTGCTT
20790

Anas_platyrhynchos_platyrhynchos 20791 TTATATTGAAGTCTC-----CTCTGTCACAAAGAGTTAAAAACTA
20835

```
Gallus_gallus                    20791 TTATATTGAAGCCTCTTTTTCTTTGTCACAAAGAGTTAAAATCTG
20835

Anas_platyrhynchos_platyrhynchos 20836 TAAGAGAATTTAAGAGCCTTACAGTAATTTGGCGCATCAGTTTTA
20880
Gallus_gallus                    20836 TAAGAGAATTTAAGAGCCTTGAAGTAATTTGGCACATCAGTTTTA
20880

Anas_platyrhynchos_platyrhynchos 20881 GCTCAGCTATTATCCAGCTGCATCAAGATATTAATGTAGTCATCT
20925
Gallus_gallus                    20881 GCTCAGCTATTATCCAGCTGCATCAAGATATTAATGTAGTCATCT
20925

Anas_platyrhynchos_platyrhynchos 20926 CTCTTCAGTCTCAAAGAATTGCTATAAAAATGTCAAACAATTAGA
20970
Gallus_gallus                    20926 CTCTTTAGTCTCAATGAATTGCTATAAAAGTGTCAAACAATTAGA
20970

Anas_platyrhynchos_platyrhynchos 20971 TTATATTGGCCAACGTAGCCTTGTTTGATTGAGTACAGATGTAGG
21015
Gallus_gallus                    20971 TTATGTTGGCCAATGTAACCCTGTTTGATGGAGTACAGATGTAGG
21015

Anas_platyrhynchos_platyrhynchos 21016 CCGTGTCCCTTGAATCATACTGATGAAGAAGTAGAAGGGCTTGTA
21060
Gallus_gallus                    21016 GCATGTCCCTTGAATCACACTGACGAAAAAAATACAAAGGCTTATA
21060

Anas_platyrhynchos_platyrhynchos 21061 GAACTCCATTGCAAATTTATTTCAATATATGTTGCATTCAGGTTT
21105
Gallus_gallus                    21061 GAACTCCAGTGTAAATTTATTTGAATATACCTACTATTTAAATTT
21105

Anas_platyrhynchos_platyrhynchos 21106 CTAATGTCAGCACTTGCAATACTAATAACCC-CAAATGCTAAATA
21150
Gallus_gallus                    21106 CTAATGTCAGTGCTTACAACACCAAAAACCCAAAAATGCTAAATA
21150

Anas_platyrhynchos_platyrhynchos 21151 GGTATTTTTAGATGGAATAAAAGTCAATGTCATTCAGGACAAATA
21195
Gallus_gallus                    21151 AATGTTTTCAGATGGAGTAAAAGTAAATGTCATTTAAGACAAATA
21195

Anas_platyrhynchos_platyrhynchos 21196 CAAATGAATTTTATTGAAAGAGACAGATATCTTGTTCTAGCCTTT
21240
Gallus_gallus                    21196 TAAGTGGCCTCTATTAGAGGAGAAAAAAT................
21240

Anas_platyrhynchos_platyrhynchos 21241 TATTAATTGCTAGTCAATGCTGAAGACAGTATATTTAGGTATTTC
21285
Gallus_gallus                    21241 .............................................
21285

Anas_platyrhynchos_platyrhynchos 21286 TTGTGACTTCATCATCCTGATTCAGTGACCACATACTAGAATAGT
21330
Gallus_gallus                    21286 ................CCAATTCAGTGACCACATTCTAGAACAGT
21330

Anas_platyrhynchos_platyrhynchos 21331 TTTTCTCTATTAAATACTATGCCAGTATGTTTATTGCTATACACA
21375
Gallus_gallus                    21331 TTTTCACTATTAAGTACTAT-----CATACTTATTGTTATATGCA
21375
```

```
Anas_platyrhynchos_platyrhynchos 21376 ATTTATCTGCTACTTCCCTATCAAGATCAACCAAATGATTACAGA
21420
Gallus_gallus                    21376 GTTTAGCTGCTACTTCACTATCAAGATCAACCAAATGGTTAGAGA
21420

Anas_platyrhynchos_platyrhynchos 21421 TAGCCATCAACGTACCTCAGTTGCACCATACTTCACACTCATTTA
21465
Gallus_gallus                    21421 TAGCCAGCTACACACCTCACCTGTAGCACACTTCTTATTCATTCA
21465

Anas_platyrhynchos_platyrhynchos 21466 TGTC---CCACAAGCTACTTCAAGTGTTACAAGTCCAGAAAAATT
21510
Gallus_gallus                    21466 TCTCATTCCTTAAGCTAATTCAAAAGTTTCCAGTAAAAATAAATT
21510

Anas_platyrhynchos_platyrhynchos 21511 CTGAGGACCAGAAAGAACTGAAAATATTTT-GGAAACTCATTCTC
21555
Gallus_gallus                    21511 CTGAGGACTGAGTAGAACCGCAGATATTTTCGGAAAGCCATTCTT
21555

Anas_platyrhynchos_platyrhynchos 21556 TTCTTGAAACAGAAACTGGTAGTTAATTATCCATTGTGTATTGAC
21600
Gallus_gallus                    21556 TCTGTGAAACAGAGACTCTTAGTTAATTCTCTGCTGTGTATTGCC
21600

Anas_platyrhynchos_platyrhynchos 21601 AGGTAATTAATAAGTACCTGTCATGTTAGTAGAAGAACAGTCAGT
21645
Gallus_gallus                    21601 AGTTAGTTAATAAGGACCAATCACACCGCT---ACAACAGTCAGT
21645

Anas_platyrhynchos_platyrhynchos 21646 TCTGGTTGCTATAAGAACACTGAGAACCTCAGCAAACAAGCTTAC
21690
Gallus_gallus                    21646
TCAGGCTGCTTCAAGAACACTCAGAACTTCAGCAGACAAGCT--- 21690

Anas_platyrhynchos_platyrhynchos 21691 TAATACTAGAATTAAGGTTAATATATCTGCTCTGAACAGACTAAA
21735
Gallus_gallus                    21691 CAGTACTAGAATGAAAATCAGTATGTCTGTTCTGAACAGTCTAAG
21735

Anas_platyrhynchos_platyrhynchos 21736 TCTTTTCATCACAAACTTCTTTTTCCACATTCCTTTTCTGTTTCA
21780
Gallus_gallus                    21736 CATTTCCATCACAAACTTTTTCCTCCCCCGTTC--TTCTGTTCCA
21780

Anas_platyrhynchos_platyrhynchos 21781 GGGCGTTGCAAATTACAGCGCTACACACAATTCCATTGTTCCAGC
21825
Gallus_gallus                    21781 GTACATTGCTGATTACATCTTATCCTGTAATTCCAT-------GC
21825

Anas_platyrhynchos_platyrhynchos 21826 TCTCCCATTGAGTTTGCCTATTTTAAGGCATAATCAGAAAATTAC
21870
Gallus_gallus                    21826 TGTCCCATTGAGTTTGCCT---------TGTAAT----------T
21870

Anas_platyrhynchos_platyrhynchos 21871 ACAAATTCCTAATCAGTGCAGCCTGTTGGAGGTTTCTGGAAAAGC
21915
Gallus_gallus                    21871 ACAAGCTTGTTCTCTGTGCTACCCATAGAAGTTTTC.........
21915

Anas_platyrhynchos_platyrhynchos 21916 TGTTCATTTTTGTGGACACCTAGTTTCTACTAACAATTTACAACG
21960
Gallus_gallus                    21916 .............................................
```

```
21960

Anas_platyrhynchos_platyrhynchos 21961 GTGAATGTCAAGTACGAGAGCATCAAGAAAATTGATAGGTCATGT
22005
Gallus_gallus                    21961 .............................................
22005

Anas_platyrhynchos_platyrhynchos 22006 ACATCCAGTTATATGAAAGCACATATTACTACAGCTCAGTACTGA
22050
Gallus_gallus                    22006 .............................................
22050

Anas_platyrhynchos_platyrhynchos 22051 TCTTGCTTTCAGTTTCATCTTGCAGGTTTTCACCCTCTTCTGCCA
22095
Gallus_gallus                    22051 .............................................
22095

Anas_platyrhynchos_platyrhynchos 22096 TCATTTGTCAGATGAAGCATTATATGAAATACAGGAGAAATAAAT
22140
Gallus_gallus                    22096 .............................................
22140

Anas_platyrhynchos_platyrhynchos 22141 CACAATAGTAATTCAGTGAAAAAGAAATAGCTTACTAAATATTTC
22185
Gallus_gallus                    22141 .............................................
22185

Anas_platyrhynchos_platyrhynchos 22186 CCACGCTAATGGGCAATGCCACCAAACTGAAAACATAACTGAATA
22230
Gallus_gallus                    22186 .............................................
22230

Anas_platyrhynchos_platyrhynchos 22231 GAGGGTCATGTTATGCTTGAACATTTTCAGAGACAGCCAAATGTA
22275
Gallus_gallus                    22231 .............................................
22275

Anas_platyrhynchos_platyrhynchos 22276 TATAATTTCTATTAATGACATAATAATTTGCCAACGGTGACACAA
22320
Gallus_gallus                    22276 .............................................
22320

Anas_platyrhynchos_platyrhynchos 22321 ACATTAAGGGTTTACAATTATTGCAGATTATTTCCTTTTTCATTT
22365
Gallus_gallus                    22321 .............................................
22365

Anas_platyrhynchos_platyrhynchos 22366 CTTGTGTCTTCTTGGCCTAAGACAGAGGTTTTTAGTTTAGAAAAG
22410
Gallus_gallus                    22366 .............................................
22410

Anas_platyrhynchos_platyrhynchos 22411 GGTGTTTACTTGGTTAGCTTGAACTATGTTTGAACTTCACACTTA
22455
Gallus_gallus                    22411 .............................................
22455

Anas_platyrhynchos_platyrhynchos 22456 GTGTATATATTCTCTCTTGACAGTGGAATAATATATTCCATGGTA
22500
Gallus_gallus                    22456 .............................................
22500

Anas_platyrhynchos_platyrhynchos 22501 GATGTACTAATGTGATTATAATATTTTGATTTTTTTATAAAATCT
```

```
22545
Gallus_gallus                     22501 ............................GTTTTTAAAAAAATCA
22545

Anas_platyrhynchos_platyrhynchos 22546 ATATAATAG---ATAGCAAGATTCAGATTTTAGGGAATTACACCT
22590
Gallus_gallus                     22546 GCATAATAACATATAGCAAGATTCAGATTTCAGGG---TGTATAT
22590

Anas_platyrhynchos_platyrhynchos 22591 GATATTTTACACACTAGCAATTTTGTTAAATTATACTGCACTTTT
22635
Gallus_gallus                     22591 TACATTTTATATACCATCACTATTATTAACACCTAATGCATTTTC
22635

Anas_platyrhynchos_platyrhynchos 22636 GTAAAAATAGAATT--GTCATCTAAAATTACCCCTTGAAACACAA
22680
Gallus_gallus                     22636 TTAAGAATAGGATTAAGCCATATAAAGTTATTTCTT-----ATAA
22680

Anas_platyrhynchos_platyrhynchos 22681 GTTTTAGGCAATGTAGATTTTAAAC------TTGTTACTAATTTG
22725
Gallus_gallus                     22681 GTTTTAGACAATAAAGATTTTAAATGGCACATTATTA--AAATTA
22725

Anas_platyrhynchos_platyrhynchos 22726 TTTCCACTGTTCTTATACTTAAATGAAGACTTCTGTGACTACTGC
22770
Gallus_gallus                     22726 TTCCCTCTCTTCTTAAACTTAAATAAAAACTTTTATGACTA---C
22770

Anas_platyrhynchos_platyrhynchos 22771 TTTGTCAATATCATCTACTAGTTTCATAAAATACTACCCATGTTC
22815
Gallus_gallus                     22771 TTTGAAAATGGCATCTACTAATTTCATAAAAGGCTACCTACATTC
22815

Anas_platyrhynchos_platyrhynchos 22816 TGCTCTCACTGGCAGAGAAATACTGTATCTACCTTTTAGATGAAT
22860
Gallus_gallus                     22816 CACTCTTGCTGACAGAGAAATATTGTATCTACCTTTTATGTGAGT
22860

Anas_platyrhynchos_platyrhynchos 22861 AATGACTAAGGTCTTCCACAAACTGGTAAATATTTCAGAGGATGT
22905
Gallus_gallus                     22861 AATAAC--AGATCTTCCACAGTGTGATAAATATTTGAGAGAACAT
22905

Anas_platyrhynchos_platyrhynchos 22906 AAACATGCCTTCA-TTTTAACTTGATCACATTTTTAAAATCCATA
22950
Gallus_gallus                     22906 -AACATGCCTTCATTTTTAACTTGATCATA---TTAACATGCAGA
22950

Anas_platyrhynchos_platyrhynchos 22951 TACATGTGTGGTAGTACTAACTGT--CTGAAGATACTTCAGCTGA
22995
Gallus_gallus                     22951 TACATTCACTGGTGTACTATCTGCAGCTGAAAACA--TGAATTAA
22995

Anas_platyrhynchos_platyrhynchos 22996 TAATATCGGAGAGA---------------------------TACA
23040
Gallus_gallus                     22996 TAGTA---GAAAGACTTTTTAAAAAGTTCTTCTTATTGCTTTACA
23040

Anas_platyrhynchos_platyrhynchos 23041 GCCGGTGAGCTGTTTTTTCACCTGCAATACATAAAAGAACAGATT
23085
Gallus_gallus                     23041 TCCATTGAACTGTTCTTTTATCCACAACACATAAAGATACAGATT
23085
```

```
Anas_platyrhynchos_platyrhynchos 23086 CATATTCATGTAGGATGGTTGACTCTCCCCTGACATCGTATTCCT
23130
Gallus_gallus                    23086 CATGTTCTTGTCGGATGATTGGCTCTCCTCTGGCATCCAATTCCT
23130

Anas_platyrhynchos_platyrhynchos 23131 GGACAGCTTCCAGGTCAGACAGGGTGCCAAAGCAGTGGTCATAAA
23175
Gallus_gallus                    23131 GGGTAACGTCCAGGTCAGACAGGGTGCCAGAGTAGTAGTCATATG
23175

Anas_platyrhynchos_platyrhynchos 23176 ACTTAGCAGGCTGTAAAGTCATTTAATTAGAGCTCAAACACTGAA
23220
Gallus_gallus                    23176 GCTTAGGAAGTTGCAAAGTCGTTTAAGTATAGCTCAAACCCAGAG
23220

Anas_platyrhynchos_platyrhynchos 23221 GAATGGGCTTAACTATTCAT-GGGAACATATTTTACAACAATCTT
23265
Gallus_gallus                    23221 GATGGGATTTAACTATTCACAAGGAATATGTTTTACAGTATTCCC
23265

Anas_platyrhynchos_platyrhynchos 23266 AAACCTACTCTAATTTCCCTGGTTACAAACACCCACAGCCATCAG
23310
Gallus_gallus                    23266 AAGCCTACTGTAATTTCCCTGGTTACAAATACTCACAGCCATGAG
23310

Anas_platyrhynchos_platyrhynchos 23311 CTGCTAAGTTTTAGCATCTGCACTACCACATCTCACTCTTTGCAA
23355
Gallus_gallus                    23311 CTACTGCGTTTTGGCATCTGTTCTACCACATCCC----TCTGCAA
23355

Anas_platyrhynchos_platyrhynchos 23356 AGTAGAGAGACAGGCTTACTGAGAGCTCAAGCTATGCTAGCTTTG
23400
Gallus_gallus                    23356 AG---AGAG-----------GAGAGCACCAGCTCT-----CTTTG
23400

Anas_platyrhynchos_platyrhynchos 23401 GAGAGAGGAGCATTTGTTATGATAAATCTTTGATTGATTGCTTAT
23445
Gallus_gallus                    23401 GAGAGAGTAGCATGGGTAATGAGAGTTCTCTGATTG----CTTAA
23445

Anas_platyrhynchos_platyrhynchos 23446 GGCAGTTTAATTTTAGTGCAGTTACACCATGGACTGGTGACAGAA
23490
Gallus_gallus                    23446 GGCAGTTTAATTTTAGAACATAAACACTGTGGACTGATGTTGGAA
23490

Anas_platyrhynchos_platyrhynchos 23491 GTTCTCTTTTTTGTTTATAATTTTCTAGCAAACTTGTAGTTCTTC
23535
Gallus_gallus                    23491 CAACT------TGTTTATCAGTTTCTAGTAAACCTGTTGCTGTTT
23535

Anas_platyrhynchos_platyrhynchos 23536 CCTGCAGATCATTTAGAACTACATACTGATGTCATTCATCTTGGA
23580
Gallus_gallus                    23536 TCTGCAGGTCATTTTGAGCTATGTACTCCTGTTATTCTTCCTGGA
23580

Anas_platyrhynchos_platyrhynchos 23581 CAGTTACCTACTACTCAAATTATGTTCTTGTCCTAGAGTATTACC
23625
Gallus_gallus                    23581 CAGTTGCCTACTAACTAAATTATGTTCCTGTGCTGAAATATTATC
23625

Anas_platyrhynchos_platyrhynchos 23626 AATAAATTCAACGTGATTTTTTTCTCCTCATATTTTTT-AGATTG
23670
```

```
Gallus_gallus                    23626 AATACATTTAACGTGATTTTT--CTCCTCATATTTTTTTAGATTA
23670

Anas_platyrhynchos_platyrhynchos 23671 CATAAATTTTAGCTAACCAATTCTTTTCAATCTCAAATATATTAC
23715
Gallus_gallus                    23671 CATTAATTTTAGCTAAGCAATTCTTTTCATGCTGAAATATATTAC
23715

Anas_platyrhynchos_platyrhynchos 23716 AAAATGTCATTTCTGTTATAAAAGCTAATTGATAATGTGATTTGT
23760
Gallus_gallus                    23716 AGAATGTCATTCCTGTTATAAGAGCTAATTGATAATGTGATTTGT
23760

Anas_platyrhynchos_platyrhynchos 23761 TATGTTGCTGAGACATAATTCAGAAAGTTATTTTAAAGCCGTTGA
23805
Gallus_gallus                    23761 TATGTTGCTGAGACATAATTCAGAAAGTTATTTTAAAGCCGTTGA
23805

Anas_platyrhynchos_platyrhynchos 23806 TACATGGAATAGCACTAACCATTTAATATTTCTTGTATGAATGGA
23850
Gallus_gallus                    23806 TGCATGGAATAGCACTATCCTTTTAATATTTCTTGTGTGGATGGA
23850

Anas_platyrhynchos_platyrhynchos 23851 TACTGCATCAGGGAATGATGTGGTGCCATATAGATGAAATATTTT
23895
Gallus_gallus                    23851 TACTGCATAAGGGAATAATGTGGTGCCATGTAGATGAAATATTTT
23895

Anas_platyrhynchos_platyrhynchos 23896 CTGAAGCTTTTATTCTTGTTGTTTGGGTTTGTGCCAATGCCTTTC
23940
Gallus_gallus                    23896 CAGAAGCTTTTATTCTTGTTGTTTGGGCTTGTGCCAATGTCTTTC
23940

Anas_platyrhynchos_platyrhynchos 23941 TTGTATAGAGCTCATATACCCTCAGCTGTATATACTGTAATTTAT
23985
Gallus_gallus                    23941 TTGTCAAGAGCTCATACACCATGAGCTGTGTCTCATGTAATTTAT
23985

Anas_platyrhynchos_platyrhynchos 23986 ACCTTGGTCATCA----GTAGGAATTTTTCCTCCAAGTTGCTGAA
24030
Gallus_gallus                    23986 ACTTTGCTCATTAGGTGGCAGGAATTTTTCCTCCAAGTTGTGGAA
24030

Anas_platyrhynchos_platyrhynchos 24031 AAATAAATAATATTTTAAAAATTAAAAGCTCACAGACAAATAATT
24075
Gallus_gallus                    24031 AAAT-----GTATTTTAATTTTTAAAAGCTCACAGTCAAATAATT
24075

Anas_platyrhynchos_platyrhynchos 24076 CATGAGCATTCTCTAGAATA---CTTCAAGAAGTTGAAAAGTATT
24120
Gallus_gallus                    24076 TGTGAGCATTCTCTAGAATGTTCCTTCCAAAAGCTAAAATGTA-T
24120

Anas_platyrhynchos_platyrhynchos 24121 TTTTATTAAGATTATTTATGCAATAATTAGTACTAGGTCCCTTCT
24165
Gallus_gallus                    24121 TTTTATCACAATTATTTATATAGT-ATTAATACTAGGACTTGTCT
24165

Anas_platyrhynchos_platyrhynchos 24166 TTTTGAATGTTCTATTCGTATGTACTTAATGAATGGTTTAACAAG
24210
Gallus_gallus                    24166 TTTTAAATGTGCTATTTGT---------ATTAGCGGTTTAACAGA
24210
```

Anas_platyrhynchos_platyrhynchos 24211 ATCAAATCAGCATTTCATCATGACTATTTCCATTGCAGTTGTTAC 24255
Gallus_gallus 24211 ATTATATCAGTATTTCATCACTGCTGTTTCCACTACAGTTGTTAG 24255

Anas_platyrhynchos_platyrhynchos 24256 GGAGTAGTCTTTCTATACAATTATCATATGAAATCAACATTCCTC 24300
Gallus_gallus 24256 GGATTACTCTTTCTATACAG-TATCAGATGAAATCAACAGTCTTC 24300

Anas_platyrhynchos_platyrhynchos 24301 ATTCAACTTTTTAACCTGGAACTGGAAATTTTGTTTCATTCATTT 24345
Gallus_gallus 24301 ATGCAATTTTTGTATTTGCAGGTAGGACTTTTGGTCCCTTTATTG 24345

Anas_platyrhynchos_platyrhynchos 24346 AATATTCAGTGTTGAAGAATGAAGACCTTCAGTATGGCACCTTGT 24390
Gallus_gallus 24346 CATATTCAGTGTTAAAGAATGATAACCTTCAGAATGGCACCTCTT 24390

Anas_platyrhynchos_platyrhynchos 24391 ATGACTATGCAACAACTTAGTTCTCCAAATCATTTGAAGCCAAGG 24435
Gallus_gallus 24391 ACGACTATGCTACGGTTAAGTTTTTCGAATCAACTGAAAGCAAGG 24435

Anas_platyrhynchos_platyrhynchos 24436 AGACTCTTGGGAGTCCAC---------TCACTACAAAATAAACAT 24480
Gallus_gallus 24436 AGCCTCTTGGGAGTCCAACATCTTAGGTCTCTACAACACAAACAC 24480

Anas_platyrhynchos_platyrhynchos 24481 ATTTCTCTTTGCTCCTCTCAGGTTGCACACTCAGAAA------TT 24525
Gallus_gallus 24481 ACCTGTCTTTGTAACTATTAAGCCACACACTCAAAAGATTTATTT 24525

Anas_platyrhynchos_platyrhynchos 24526 GGCTTTCATTGCCATCACCTGTTGACTCACTCTTTGGGGG----- 24570
Gallus_gallus 24526 GGATTTAATTGCCACCATCTGCTGACTCACTCTATGGGGGGAAGG 24570

Anas_platyrhynchos_platyrhynchos 24571 --------AAAACTTGCTATTGTCTGCCCAGTTAGCTTGCTATGA 24615
Gallus_gallus 24571 AGGGAACCAAAACTTGCTATTGTCAGTCCAGTTAGTTTGCTATGA 24615

Anas_platyrhynchos_platyrhynchos 24616 GAGTGTCTCACAATTTTTTTTTCAGTCTTGAACCTTGGCTCGAAC 24660
Gallus_gallus 24616 GAGGGTCTCACAA-TTTTTGTTCAGTCTTGAGTCTTTGCTTCAAC 24660

Anas_platyrhynchos_platyrhynchos 24661 AAATATGACATTTACTCAGACCTTTGCACAGTCTGTTTGAAACCT 24705
Gallus_gallus 24661 AAAGATGACATTTACCCAGACCTCTGCATGGTCTGTTTGAAAAAT 24705

Anas_platyrhynchos_platyrhynchos 24706 GACAAGGTTGTCATTCCATGAGCACAGAAGATATCCGAGAGCCTC 24750
Gallus_gallus 24706 GACACAGTTGTAATTCCAAGAGCTCAAAACATATCTGAAAA---- 24750

Anas_platyrhynchos_platyrhynchos 24751 TGTTTCTCTCTCTCTCTCTTTTTTTTTTTTTCACTCCCACTTTCT 24795
Gallus_gallus 24751 ----------TCTCTCTATTTTTGTCTTTTTC-CTCTCCCATTCT

24795

Anas_platyrhynchos_platyrhynchos 24796 CCCCTAATTTGAGACTCCACCCAAGGTGTGGCTGCCTTGTAATTC
24840
Gallus_gallus 24796 ATCTGGATTTGAGAATCCACTTGGATAGTGGCTGAC--GTAATTC
24840

Anas_platyrhynchos_platyrhynchos 24841 TCAGCTGATGATCAAATGACTGAACAATAAATGAAAATATTTCAA
24885
Gallus_gallus 24841 TCACCTGATGCTTAAATAACTGCACTATAAATTGAAGCGTTTCCA
24885

Anas_platyrhynchos_platyrhynchos 24886 TTTGTTCTGGTGAATACACTCTAAATTATCTGTCATGTGCCATTT
24930
Gallus_gallus 24886 TCTGTTCTTGTGAATACAACCTAAACTATCCATCATGTGCCATTT
24930

Anas_platyrhynchos_platyrhynchos 24931 TTTAGAACTTGGCTGGCTGACTGTAAAATTCTTAAGTGAAAACTG
24975
Gallus_gallus 24931 CTTGGAGCTGGGTAGGTTGGCACTAAAATACTAGAGTGAAAACTA
24975

Anas_platyrhynchos_platyrhynchos 24976 TTACATCTCTGCAAAAACTTCAAAATTGCATTAAATATTCTTGTG
25020
Gallus_gallus 24976 CTGCATTTCAGCAAATGCTTCAAAATTTCTGAAAAGGTTCTTGTG
25020

Anas_platyrhynchos_platyrhynchos 25021 TGGGATCACAGAAGTAAATTT---GACTTATTTAGCTTCATATAC
25065
Gallus_gallus 25021 TATGATCACGGGAGGAAAACTTGAGACCTATTTAGCTTCATATAT
25065

Anas_platyrhynchos_platyrhynchos 25066 TCTGGGACTCTTTAGACATCTTACAATAAATTTAACTCAAAGTGG
25110
Gallus_gallus 25066 GCTGA---TCTTTAGACATCTTGCAGTAAATTTAACCGAAAGTGG
25110

Anas_platyrhynchos_platyrhynchos 25111 GTTCTTTTTATGTTTACTAATTAAAACTCATTTTGTCCTCACAAA
25155
Gallus_gallus 25111
ATTCTTTTAATGTTTACTAATTTAAACTCATTT------------ 25155

Anas_platyrhynchos_platyrhynchos 25156 GGACTCCTTCTGCTTAAATTCTCAATCACAATGTGAAGAATATAT
25200
Gallus_gallus
25156 -------TCATTCTTAAACT----ATCACAATTTGAAAAACA--- 25200

Anas_platyrhynchos_platyrhynchos 25201 ATCTAAAAGCATTATATATATATAATATAAATATATACAAAATAT
25245
Gallus_gallus 25201 --CCAAAA.......................................
25245

Anas_platyrhynchos_platyrhynchos 25246 ATAATAATATTTAAATATATAAATAATGTGTATATATTATGCTTT
25290
Gallus_gallus 25246 ..................................ATATTGAGCTGT
25290

Anas_platyrhynchos_platyrhynchos 25291 TAGATGCATACTCAGATCAGGATATGAAACAG-ATTGTTTTACAG
25335
Gallus_gallus 25291 TAGACGCATACTCAAATTAGGATCTGAAATAGAAATTTTCTAGAA
25335

Anas_platyrhynchos_platyrhynchos 25336 TGTTGTTTCTCAACAGGTTTTTGTTTGTTTGTTTTGTTTTTTTTT

```
25380
Gallus_gallus                     25336 TGTTTTTTCTCAACA-------CTGTGGTTGGTTGGTTGGCTTGT
25380

Anas_platyrhynchos_platyrhynchos 25381 TGCTTAGCTTCATTCACATTTACAGTCTTTAATAGATCAGCATTC
25425
Gallus_gallus                     25381 TGCTTAGCTCCATTCATGTCTACTGTCTTCAAGAGATTAGCACTC
25425

Anas_platyrhynchos_platyrhynchos 25426 TGAACTCTAAAATTTTATCAGCTGGACTGCTGAGTCCTTCCTTTA
25470
Gallus_gallus                     25426 TGAGCTCCAAAATTTTATTACCCAGGCTTCTAAATCTTTTGTCTA
25470

Anas_platyrhynchos_platyrhynchos 25471 TATACGATGCAGTTTCTTAGCCTTCTTTGTTAAATTGCT-TTGTA
25515
Gallus_gallus                     25471 TGTAAGATGTGGTTTCTTATCCTCCCTAGTTAAGCCGCCATTGTA
25515

Anas_platyrhynchos_platyrhynchos 25516 AGTTTACAAAGAAATACAAAATACATCAATTTGCCTATGCTCTTT
25560
Gallus_gallus                     25516 AATTTACAAAGATATATAAAACACATTAATTTATTTATGCTCTTT
25560

Anas_platyrhynchos_platyrhynchos 25561 TTCTTTCAATATCATTCCAAACTCTTCTATACATCTTTTCCTGAA
25605
Gallus_gallus                     25561 TTTTCTA-----CATTCCTAATTCTACTACATATCTTTTCCTGAA
25605

Anas_platyrhynchos_platyrhynchos 25606 GTGACTTATTACTTCT-TACCATCAGGAAATTTACTTATGTCATG
25650
Gallus_gallus                     25606 GTGACTCTCTACTTCTGTACCATCAGGAAATTTAACCACATCACT
25650

Anas_platyrhynchos_platyrhynchos 25651 GCACACTGCTAGAAACATATAATGGAAGGTAAGATATCATTTCCA
25695
Gallus_gallus                     25651 GCAAACTGCTACAAACACATAAAAGA--GAAGTTTCTCATTTCTG
25695

Anas_platyrhynchos_platyrhynchos 25696 GTACTAATTCCTGAAGCAATGAACTAGGTATCTCCTACTGTCACT
25740
Gallus_gallus                     25696 GTACTAATTCCTGAAGCAACGAATTAGATATCTTTTGGTTTCACT
25740

Anas_platyrhynchos_platyrhynchos 25741 GCATGAATATCCACTGTTACCACCTA-AAATACAGTTTATCATCT
25785
Gallus_gallus                     25741 GCATGAATATCCACTACCACCAGAGAGAAGTATAGTTTATCACTT
25785

Anas_platyrhynchos_platyrhynchos 25786 CACATCCTTAGTCTTACTACCATTCATGATTTTACTGACTGGTAT
25830
Gallus_gallus                     25786 CATGTCCTTAGTTCTACTACTATTAAGGATTTTACCGATTAGTAT
25830

Anas_platyrhynchos_platyrhynchos 25831 TTAGCAAAGCAGTACAGAAAATATTCTCCTAATAACCATGGCAAT
25875
Gallus_gallus                     25831 TTAGCAAATTAATACAGAAAGTATTCTCCTAATGCTCCAGGTCAT
25875

Anas_platyrhynchos_platyrhynchos 25876 TTGGTTATGAATACACTCCATGGTCCTAAGTTAATTATTATTTTA
25920
Gallus_gallus                     25876 TTCCTTATGACTACACTCCATAGCTCTAAGTTAATTTTTTTTCTA
25920
```

```
Anas_platyrhynchos_platyrhynchos 25921 ATACCTCAGAAATATCAATCAGATATTATGTATAAATGCATGCCA
25965
Gallus_gallus                    25921 GTACTTCAGAAATGCCAATCAGATATTATGTATATATGTATGCCA
25965

Anas_platyrhynchos_platyrhynchos 25966 CCCACTTTTACAATGCTTAATCAGCATAGATTGCAGATTTGTAGG
26010
Gallus_gallus                    25966 TCCACTTCTATCATACTTAACCAGAATGCGCTGCATATTTGTAGG
26010

Anas_platyrhynchos_platyrhynchos 26011 GATTCCA--CCAGTTTCTCTGATTAGTATTTCCAATATCTTACTC
26055
Gallus_gallus                    26011
GATTTCATCTCAGATTCTCTGATTAGCATTTCTA----------- 26055

Anas_platyrhynchos_platyrhynchos 26056 CCAACAAGTAAAGCTTACAGGTCTGAAGTTTCCTGGATAAGGCCA
26100
Gallus_gallus                    26056 --AAAAAGTAAAGCTTACAGATCCGATGTTTTATGAATAAGGCCA
26100

Anas_platyrhynchos_platyrhynchos 26101 TCACTCTTTATCAGTGCTTTCTGATAAAGTAATGCTGTCAGACTG
26145
Gallus_gallus                    26101 TTACTCTTTATCAATGCTTTCTGATAAAGTACTGCTGTCAAACTG
26145

Anas_platyrhynchos_platyrhynchos 26146 TAAAGAAATTTTTGGTCATACTTATTCATCTTTCTGACAGAGATT
26190
Gallus_gallus                    26146 CAAAGAAACTTTTGGTCATGCTCATTCATCTTTCTGACAGAGGTT
26190

Anas_platyrhynchos_platyrhynchos 26191 GTTGCAATGTAGTTAGGAATAAAGTATCAAATATAAACGAGAATA
26235
Gallus_gallus                    26191 GCTGCAATGTAGTTAGGGATTAAGTATCAAATGTAAACAGAAATA
26235

Anas_platyrhynchos_platyrhynchos 26236 TTTTTTGGAGTAAAACACTTTAATGTATTGTTATTTCCTTCAGTT
26280
Gallus_gallus                    26236 -TTTTTGGAGAAAAACACTTGAACA------TATTTCCATCAGTT
26280

Anas_platyrhynchos_platyrhynchos 26281 TCATAAATAAAGTCACAGTAGTAAACTATCAAATACGCCCATTGT
26325
Gallus_gallus                    26281 TAGTATGTAAAACCACAGTAGTGAACTACTGAATATGCCCATTGT
26325

Anas_platyrhynchos_platyrhynchos 26326 ACTTAGATATCAATGATCCTTCGAACCTTTGCAGTCTTACAGTTT
26370
Gallus_gallus                    26326 ACCAGGATATCGGTGTTCTTTCCAACCTTCACA-TCTTGTCTTTT
26370

Anas_platyrhynchos_platyrhynchos 26371 TTATCCTGGATTACTTTCAAAGAACATCACCGTCATTTCTAAAGC
26415
Gallus_gallus                    26371 TTATACTATACTACTATGAAAGAACA--ACAGTCATTTTTGAAGC
26415

Anas_platyrhynchos_platyrhynchos 26416 ATTGAAAATTTTCTGGACTTCAATATTTATGAAATGCAATATTTC
26460
Gallus_gallus                    26416 ACTGGAAGATTTCTGGACTTCACTATTT.................
26460

Anas_platyrhynchos_platyrhynchos 26461 AAGAATGTGGAAATTAATACAAGTTCATTAAAACTATCATGTAGC
26505
```

```
Gallus_gallus                    26461 .............................................
26505

Anas_platyrhynchos_platyrhynchos 26506 AAGGCAGAATAGCTTCATGATTCCTACTGAATGTAGTGCTTTTTA
26550
Gallus_gallus                    26506 .............................................
26550

Anas_platyrhynchos_platyrhynchos 26551 AAAAGCCTTTCTTATTACAGAAAAACAGTAAAAGAAAGACAGTAT
26595
Gallus_gallus                    26551 .............................................
26595

Anas_platyrhynchos_platyrhynchos 26596 TACAGGTCAAAAATGCCATTAGCAATAGGAAAACACAATTTCATA
26640
Gallus_gallus                    26596 .............................................
26640

Anas_platyrhynchos_platyrhynchos 26641 ACAAAAAATTTATTGGAAAAAGATGTGGAAAAAGTAAACTGAAAC
26685
Gallus_gallus                    26641 .............................................
26685

Anas_platyrhynchos_platyrhynchos 26686 ATTAACAGTTTAGTGTACATTTGCTATACTGAACAACAGATGCAA
26730
Gallus_gallus                    26686 .............................................
26730

Anas_platyrhynchos_platyrhynchos 26731 CCATTTGCTTCTTCAACTTCAAAAACACTTGAAATTCCCAAATGC
26775
Gallus_gallus                    26731 .............................................
26775

Anas_platyrhynchos_platyrhynchos 26776 CGTGCATGGGATGTTAAACAGGCAGCTAAGCAATGGTGACGCAAT
26820
Gallus_gallus                    26776 .............................................
26820

Anas_platyrhynchos_platyrhynchos 26821 GAAAAGGTACCTGTCGAGATTTAAAATAAATAAACAAATAAATAA
26865
Gallus_gallus                    26821 .............................................
26865

Anas_platyrhynchos_platyrhynchos 26866 ATAGTCCTAGTTTCAGCTGGGATAGAATTAATTTTCTTCATAGAG
26910
Gallus_gallus                    26866 .............................................
26910

Anas_platyrhynchos_platyrhynchos 26911 GCTGGTATGTGCCATGGTTTGGATTTAGGATGAGAATAATGCTGA
26955
Gallus_gallus                    26911 .............................................
26955

Anas_platyrhynchos_platyrhynchos 26956 TATCGTACTGATGTTTTAGTTGTTGCAGAGCAGTGCTTACACTAA
27000
Gallus_gallus                    26956 .............................................
27000

Anas_platyrhynchos_platyrhynchos 27001 GCCAAGGGCTTTTCAGCTTCTCCCACTGCCCTGCTAGTGAGGAGG
27045
Gallus_gallus                    27001 .............................................
27045
```

Anas_platyrhynchos_platyrhynchos 27046 CTGGGGGTGCACAAGGAACAGAGAGAGGACAGAGCCAGAACAGCT
27090
Gallus_gallus 27046 .............................................
27090

Anas_platyrhynchos_platyrhynchos 27091 GACCCCAAATGGCCAAAGGGATATCCCGTATCTTGTGGTGTCATT
27135
Gallus_gallus 27091 .............................................
27135

Anas_platyrhynchos_platyrhynchos 27136 GTCGTGGTTCCGCTCGAGTGGGCAGCCGAGCTCCACCACAGCCGC
27180
Gallus_gallus 27136 .............................................
27180

Anas_platyrhynchos_platyrhynchos 27181 TCTCTCACTCCCCCTCCTCAAAGAGGAATGGGGAGAAAATATGTG
27225
Gallus_gallus 27181 .............................................
27225

Anas_platyrhynchos_platyrhynchos 27226 AAAGGGGCTCAAGGACTGAGATAAGGACGAGAAAATCACGCAATA
27270
Gallus_gallus 27226 .............................................
27270

Anas_platyrhynchos_platyrhynchos 27271 ATTAGTGTAATGGGCAAACCAGACTCAGCATAAGAAGACAGTAAG
27315
Gallus_gallus 27271 .............................................
27315

Anas_platyrhynchos_platyrhynchos 27316 ATTTATTGCTCATTACTAACAAGCTAGAGAAGTGAGAAACAAAGG
27360
Gallus_gallus 27316 .............................................
27360

Anas_platyrhynchos_platyrhynchos 27361 AAAGAAACCAAAAGCACCTCCCCCCCCCCCATCCACCCTCTTCCA
27405
Gallus_gallus 27361 .............................................
27405

Anas_platyrhynchos_platyrhynchos 27406 CCTCCTCCCCCCGAGCGGCGCAGGGGAACGGGGGAATGGGGGTTA
27450
Gallus_gallus 27406 .............................................
27450

Anas_platyrhynchos_platyrhynchos 27451 TGGTCAGTCTACAGCACTTCTTCTCTGCCGCTCCTTCTCGGTCAC
27495
Gallus_gallus 27451 .............................................
27495

Anas_platyrhynchos_platyrhynchos 27496 TCTTGTCCCCTGTGCTGTGGGGTCCCTCCCACAGGATGCAGTCCT
27540
Gallus_gallus 27496 .............................................
27540

Anas_platyrhynchos_platyrhynchos 27541 TGATGAACTGATCCGGCGTGGGCTTCCCACAGGCAGCAGCTCTTC
27585
Gallus_gallus 27541 .............................................
27585

Anas_platyrhynchos_platyrhynchos 27586 CAGAACTGCTCCAGATATGGGTCCGTACCACGGGGTCCATCCCTC
27630
Gallus_gallus 27586 .............................................

27630

Anas_platyrhynchos_platyrhynchos 27631 AGGAGAAAACTGCTCCAACCTGGCTCCCCTACGGGCAGCAGCTCC
27675
Gallus_gallus 27631 .............................................
27675

Anas_platyrhynchos_platyrhynchos 27676 TGCCAGGTCACCTGCTCCTGCGTGGGCTCCTCTCCACGGGCTACA
27720
Gallus_gallus 27676 .............................................
27720

Anas_platyrhynchos_platyrhynchos 27721 GGTCCGGCCCGGAATCTGCTCCGGCAGGGGTCTTCCACAGGCGGC
27765
Gallus_gallus 27721 .............................................
27765

Anas_platyrhynchos_platyrhynchos 27766 AGCCTCCATCGGTGCAGGGCCACCTGCTCCACCGTGGTCTCCTCC
27810
Gallus_gallus 27766 .............................................
27810

Anas_platyrhynchos_platyrhynchos 27811 ACGGGCTGCAGCATGGAACCCTGCTCCACCGTGGTACTCCATGGG
27855
Gallus_gallus 27811 .............................................
27855

Anas_platyrhynchos_platyrhynchos 27856 CTGCAGGGGGACATCCTGCTTCACCATGGTCCTCACCACAGGCCG
27900
Gallus_gallus 27856 .............................................
27900

Anas_platyrhynchos_platyrhynchos 27901 CAGGGGACTTCTGCTCCGGCGCCTGGAGCACCTCTCCCCCTCCTT
27945
Gallus_gallus 27901 .............................................
27945

Anas_platyrhynchos_platyrhynchos 27946 CTACACTGACCTTGGCACCTGCAAGGCTGTTTCTCACTCCCTTGA
27990
Gallus_gallus 27946 .............................................
27990

Anas_platyrhynchos_platyrhynchos 27991 CTCTCCCGGCTGCTGTGTGGCGCAGCGTTTTTTTCCCTGTCTTAA
28035
Gallus_gallus 27991 .............................................
28035

Anas_platyrhynchos_platyrhynchos 28036 ATATGCTCTCACAGAGGCGCAAAACAACATCGCTTATTGGCTCAG
28080
Gallus_gallus 28036 .............................................
28080

Anas_platyrhynchos_platyrhynchos 28081 ATCTGGAAAACAACGGGGCCCTTCCCAAACATGGGGCGGCTTCTA
28125
Gallus_gallus 28081 .............................................
28125

Anas_platyrhynchos_platyrhynchos 28126 GATCTTTCTCACAGAAACCACCCCTATGACCCCCTGCTACCAAAA
28170
Gallus_gallus 28126 .............................................
28170

Anas_platyrhynchos_platyrhynchos 28171 CCTTGCCACGTAAACCCACTACAGTCATGTGGGACAATAAAACTG

28215
Gallus_gallus 28171 .............................................
28215

Anas_platyrhynchos_platyrhynchos 28216 GAAAGGCTGTCTGGAGGGGCTGCTGTTTCTTGGGAACTGTGAGAG
28260
Gallus_gallus 28216 .............................................
28260

Anas_platyrhynchos_platyrhynchos 28261 ATTTGGTTGGCTGATTGTGCAAAACTGCATGGTGTATTGCTTTTG
28305
Gallus_gallus 28261 .............................................
28305

Anas_platyrhynchos_platyrhynchos 28306 TTGTTTGTTTTTTGTTCCTTTCTTATTTTTTTTTCCTTTTTATTA
28350
Gallus_gallus 28306 .............................................
28350

Anas_platyrhynchos_platyrhynchos 28351 AACTGTCCTTATCTCCACCCATGAATTTCTATACTGTTCTTTGCA
28395
Gallus_gallus 28351 .............................................
28395

Anas_platyrhynchos_platyrhynchos 28396 ATTTTCTCTTCCATCTCACTGGACGGTCTGGAGTGAGTGAACAGC
28440
Gallus_gallus 28396 .............................................
28440

Anas_platyrhynchos_platyrhynchos 28441 TGTGTGGCGTTTAGCTGCCTGCCCACTTGAGCACAATTCTCTTTT
28485
Gallus_gallus 28441 .............................................
28485

Anas_platyrhynchos_platyrhynchos 28486 TTTTTTTTTTCCTCATTTTTTTTTTTTTTAGCTTGTATTGTTACC
28530
Gallus_gallus 28486 .............................................
28530

Anas_platyrhynchos_platyrhynchos 28531 AAATGCCCAGACATATGCCTCTCAAAGTATTTGTCCCATGCTTAA
28575
Gallus_gallus 28531 .............................................
28575

Anas_platyrhynchos_platyrhynchos 28576 ATAAAGAATGAAGTTACATATAGAGTGAATAACAAGGATTAGGGA
28620
Gallus_gallus 28576 .............................................
28620

Anas_platyrhynchos_platyrhynchos 28621 ACACAGTTGACAGAGAAGCTGTGATTCATATAAGGTTATCTAATT
28665
Gallus_gallus 28621 .............................................
28665

Anas_platyrhynchos_platyrhynchos 28666 AAATTGTTAGAACAATGAAGAAGGAAGTACAAAGTTGTATTATAT
28710
Gallus_gallus 28666 .............................................
28710

Anas_platyrhynchos_platyrhynchos 28711 TGTGTACATAAAAGAAGCATCTCACAAGTCATGAGTTTGTAATTA
28755
Gallus_gallus 28711 .............................................
28755

```
Anas_platyrhynchos_platyrhynchos 28756 ATGTCTATATAGCCCTGTGACTCTTATCAGGTTTTAATTAACTGT
28800
Gallus_gallus                    28756 .............................................
28800

Anas_platyrhynchos_platyrhynchos 28801 AGCACAGCAGGAATGCTAAGAACAGTATATTTTCAGTGCTATTTT
28845
Gallus_gallus                    28801 .............................................
28845

Anas_platyrhynchos_platyrhynchos 28846 CATACTCAACCAGGAACATAAAAAATCATAGGAAACAGGCATCAC
28890
Gallus_gallus                    28846 .............................................
28890

Anas_platyrhynchos_platyrhynchos 28891 CATCTTACTGGAAAAACTAAAATGTTTTTAGTTGCAATAATGATC
28935
Gallus_gallus                    28891 .............................................
28935

Anas_platyrhynchos_platyrhynchos 28936 ATAGCTTTGCATTTTCTATACATACTACATAACTTTACATATTTT
28980
Gallus_gallus                    28936 .............................................
28980

Anas_platyrhynchos_platyrhynchos 28981 TTTTTTCCTGTTATCATTACAAATAAAGTTTTAAAGTATGTATTT
29025
Gallus_gallus                    28981 .............................................
29025

Anas_platyrhynchos_platyrhynchos 29026 TAAAAACATACTGCTGATTAGCAGAACATACTACAAAATCTATGC
29070
Gallus_gallus                    29026 .............................................
29070

Anas_platyrhynchos_platyrhynchos 29071 ATATAAAATGCTTATCTGGGATTTTGGGATAATTTAATATATCTT
29115
Gallus_gallus                    29071 .............................................
29115

Anas_platyrhynchos_platyrhynchos 29116 CATAAAAGGTACTTTAATGTTATTCAGAAAGCCTATGGGAAAATA
29160
Gallus_gallus                    29116 .............................................
29160

Anas_platyrhynchos_platyrhynchos 29161 AAGAAAGCAAAACTTTAACATTGCCCTATTTTCTACTTAAACTAC
29205
Gallus_gallus                    29161 .............................................
29205

Anas_platyrhynchos_platyrhynchos 29206 AACTGCTGCTATACACAAAGTTAAAAATCAATTAACTCAGAATAA
29250
Gallus_gallus                    29206 .............................................
29250

Anas_platyrhynchos_platyrhynchos 29251 GGTGACAGTTAATTCTTCAGCTGATACATCTTCCATTTAAAAACA
29295
Gallus_gallus                    29251 .............................................
29295

Anas_platyrhynchos_platyrhynchos 29296 AAATACATCAAGGAAAAGTATTCCCAAATGGAAGCACTGACAGTA
29340
```

```
Gallus_gallus                    29296 .............................................
29340

Anas_platyrhynchos_platyrhynchos 29341 ACATTCCAATGAAATTGGCATCAACACCCCAAAACACTAGCCTGG
29385
Gallus_gallus                    29341 .............................................
29385

Anas_platyrhynchos_platyrhynchos 29386 TTTCCTTTAAGGTGTTGGTGCTAGTACTCACCAAGAATTTTTTTT
29430
Gallus_gallus                    29386 .............................................
29430

Anas_platyrhynchos_platyrhynchos 29431 TCAAATTTTCAGTGGAATCTGAAGAAAATAGAAATATTAGTAATT
29475
Gallus_gallus                    29431 .............................................
29475

Anas_platyrhynchos_platyrhynchos 29476 CTTTAGAGGGATTCGCTGAGTTCGGAAAAGCTTTGAATACCAACT
29520
Gallus_gallus                    29476 .............................................
29520

Anas_platyrhynchos_platyrhynchos 29521 GACATCTCTAAAATTAGTGTTTTTCGATAATTTCATGACACTTAT
29565
Gallus_gallus                    29521 .............................................
29565

Anas_platyrhynchos_platyrhynchos 29566 TCTAATATCAACCTGTTTATATCCAAAACCAAAATCAGTCAATCA
29610
Gallus_gallus                    29566 .............................................
29610

Anas_platyrhynchos_platyrhynchos 29611 GCATAACAGATGTTTTCTTAAGCTTACCTCTCCTTAGGCTATATA
29655
Gallus_gallus                    29611 .............................................
29655

Anas_platyrhynchos_platyrhynchos 29656 GAAGAGTATCTAGCACAATAATGTGAGATATGCCATATGCACGGT
29700
Gallus_gallus                    29656 .............................................
29700

Anas_platyrhynchos_platyrhynchos 29701 CTAGTGGTAATTCATAGTATGGAAAGTCTGACACAAATTACAGAA
29745
Gallus_gallus                    29701 .............................................
29745

Anas_platyrhynchos_platyrhynchos 29746 TAATTATGTTGGGCAAACCATATAACACACAATTAAAAAATGATG
29790
Gallus_gallus                    29746 .............................................
29790

Anas_platyrhynchos_platyrhynchos 29791 GATGAATGGCAGTTGAGGAAGTGTGAAAGTTCTGTTCTGTTTTGT
29835
Gallus_gallus                    29791 .............................................
29835

Anas_platyrhynchos_platyrhynchos 29836 CTGTAGGAGAACTGTGAGAGTCTTCATCCACTAAAAAAAAAATAA
29880
Gallus_gallus                    29836 .............................................
29880
```

```
Anas_platyrhynchos_platyrhynchos 29881 ATTTTAAATATTGTTATTTCTGAATACGACTGAATATAAGAATGA
29925
Gallus_gallus                    29881 .............................................
29925

Anas_platyrhynchos_platyrhynchos 29926 AGTGCAGTAACTTAACCACTTCTATTTGTTTAATACTTATGGTAC
29970
Gallus_gallus                    29926 .............................................
29970

Anas_platyrhynchos_platyrhynchos 29971 TTTTCCATTCCTCTCTTTAGAGGAATGTACTCAGAAATCCTCCCC
30015
Gallus_gallus                    29971 .............................................
30015

Anas_platyrhynchos_platyrhynchos 30016 TCTCCAACAAGATGCCTAGTATAATTCTTCTTGTAGCGTGCCCTT
30060
Gallus_gallus                    30016 .............................................
30060

Anas_platyrhynchos_platyrhynchos 30061 TACAATGAAGACTGTCTAATGACTGCAGGCTCAACTTTTTGCATT
30105
Gallus_gallus                    30061 .............................................
30105

Anas_platyrhynchos_platyrhynchos 30106 CTATAATCTATACCCATAAAACTAATCTACCATGCACTGTGAAGC
30150
Gallus_gallus                    30106 .............................................
30150

Anas_platyrhynchos_platyrhynchos 30151 AAAATTATGCTGTTCATGTCATGTTCATGATGTAACTTTTGAGTC
30195
Gallus_gallus                    30151 .............................................
30195

Anas_platyrhynchos_platyrhynchos 30196 ACACCATTTTTTTCATTTTATAACTGTTCAGTGGAGCATAAATTT
30240
Gallus_gallus                    30196 .............................................
30240

Anas_platyrhynchos_platyrhynchos 30241 TAGCTTTTGTTCAACTATGACAAAGTCTCCCCAAATTTTTTATTT
30285
Gallus_gallus                    30241 .............................................
30285

Anas_platyrhynchos_platyrhynchos 30286 ATTTATTTATTAATTGTTTATTTAATACATGTGAAATACAGCTAC
30330
Gallus_gallus                    30286 .............................................
30330

Anas_platyrhynchos_platyrhynchos 30331 TGCAATTGTAATATTAAATCAGCCATCTTTCCACTTTCAGTTCTT
30375
Gallus_gallus                    30331 .............................................
30375

Anas_platyrhynchos_platyrhynchos 30376 GTAAAAATATTCTCAAAACTGTCATAGCTCATTTTAAAGATTTTA
30420
Gallus_gallus                    30376 .............................................
30420

Anas_platyrhynchos_platyrhynchos 30421 TCTATATCTGTAAACTGCCTATAAGTCTTCAGGTCTACCAGGTTC
30465
Gallus_gallus                    30421 .............................................
```

30465

Anas_platyrhynchos_platyrhynchos 30466 TAATTGTGTTAATGTGCAATGCAAAGCTTTTGTACAAAGCTTTTT
30510
Gallus_gallus 30466 .............................................
30510

Anas_platyrhynchos_platyrhynchos 30511 AGTTCTTACTTTGAGCTGAATCCTTTTCCCTGTGAATCTTCCTTT
30555
Gallus_gallus 30511 .............................................
30555

Anas_platyrhynchos_platyrhynchos 30556 TTTGAAGAGTATGGATGGTCTGACCAGTTGTTCTACCAATGTGCA
30600
Gallus_gallus 30556 .............................................
30600

Anas_platyrhynchos_platyrhynchos 30601 ACAAATCTGGAAAGAAGTATGTTAGTTCCTGTCAGTTTTTACTGA
30645
Gallus_gallus 30601 .............................................
30645

Anas_platyrhynchos_platyrhynchos 30646 AAGTTTTGAAAAACAAGAAAGTATCAAACATATATGGCCTGATGT
30690
Gallus_gallus 30646 .............................................
30690

Anas_platyrhynchos_platyrhynchos 30691 CTTTACATTTAAAAAACAATAACAACAAACAAAAAGGATACTATT
30735
Gallus_gallus 30691 .............................................
30735

Anas_platyrhynchos_platyrhynchos 30736 AGAGATTTAGTTCTGTATAACTGAAGAGTTAGCTTCACTTTTTGT
30780
Gallus_gallus 30736 .............................................
30780

Anas_platyrhynchos_platyrhynchos 30781 TTTCCTCACAGTTCATGCAACTATAGATTGTCAAGCCTTAGGTCA
30825
Gallus_gallus 30781 .............................................
30825

Anas_platyrhynchos_platyrhynchos 30826 CTGCTTCTCACCACAGAAATCCTATAGAAATCCTATCTTACCATA
30870
Gallus_gallus 30826 .............................................
30870

Anas_platyrhynchos_platyrhynchos 30871 TGACCATAAATAAATCCTCTACTAAAATTGATTTAAAGAGATTAT
30915
Gallus_gallus 30871 .............................................
30915

Anas_platyrhynchos_platyrhynchos 30916 TGTTCTACTTGAAAGGTTAACCTGTCAACAACACAGGGTGTCCTG
30960
Gallus_gallus 30916 .............................................
30960

Anas_platyrhynchos_platyrhynchos 30961 CATGCTGTTTACCTGTTCTGGAAACACCAGGTAAAATTTGCCCAG
31005
Gallus_gallus 30961 .............................................
31005

Anas_platyrhynchos_platyrhynchos 31006 GAGCAAAGACAGCGGAGAAAGGGGAGAAAAATAGATACACACACC

31050
Gallus_gallus                    31006 .............................................
31050

Anas_platyrhynchos_platyrhynchos 31051 CCATGAGTAATATAAATACAAACATTTGAAATAATGCATCTGTGG
31095
Gallus_gallus                    31051 .............................................
31095

Anas_platyrhynchos_platyrhynchos 31096 ATAAAGAAGAATACAAATACTTACTCTCAAACAAAATTTCTAGCC
31140
Gallus_gallus                    31096 .............................................
31140

Anas_platyrhynchos_platyrhynchos 31141 TTTCTGCATTCTCTTGCAGGGTAGTGTACCACTGGTTGGCTATTA
31185
Gallus_gallus                    31141 .............................................
31185

Anas_platyrhynchos_platyrhynchos 31186 TAGAACGAACACCAGTAAGGCTAAGAAGCATTACTGTTTCTGTTG
31230
Gallus_gallus                    31186 .............................................
31230

Anas_platyrhynchos_platyrhynchos 31231 GTCTCTCTATAGCAAACTGTAAAGAACTAAAAAAATAAAAAGTAT
31275
Gallus_gallus                    31231 .............................................
31275

Anas_platyrhynchos_platyrhynchos 31276 GAGATCAATAAAATAAAAATGTAATTGAAATTCTGCATACTAATA
31320
Gallus_gallus                    31276 .............................................
31320

Anas_platyrhynchos_platyrhynchos 31321 TATTAAAAACATAAATATATAACACTTTTTAACAGGAGAAGAAAG
31365
Gallus_gallus                    31321 .............................................
31365

Anas_platyrhynchos_platyrhynchos 31366 AGAATTTTATACGAATCTCAGCAAACTAAGAAAAAATCCCAGAATC
31410
Gallus_gallus                    31366 .............................................
31410

Anas_platyrhynchos_platyrhynchos 31411 CATTTCTCTTGTATGTTAAGCATAAAATGCTTTCAAATATTTTGT
31455
Gallus_gallus                    31411 .............................................
31455

Anas_platyrhynchos_platyrhynchos 31456 ACCATATCTTATGTTTAAAACAAGGAATTATAATAACTTTTCTAT
31500
Gallus_gallus                    31456 .............................................
31500

Anas_platyrhynchos_platyrhynchos 31501 ACCTCTTTTCATGATGCAATTTTATGTGGATTGCTCCCTGTCATT
31545
Gallus_gallus                    31501 .............................................
31545

Anas_platyrhynchos_platyrhynchos 31546 CTACTGCTTTGGGCATCAAATGAAATACGAATGGAAGAACATCAC
31590
Gallus_gallus                    31546 .............................................
31590

```
Anas_platyrhynchos_platyrhynchos 31591 ATTAACTTTTATAAAACAATGCAGAACACCCTTTATAAGAGACTC
31635
Gallus_gallus                    31591 .............................................
31635

Anas_platyrhynchos_platyrhynchos 31636 AAAATGCTGTGAAATGGATTAAAAGACATGTTTAGTTCAGCTTCT
31680
Gallus_gallus                    31636 .............................................
31680

Anas_platyrhynchos_platyrhynchos 31681 ATTCTTAAAGGGAAAAAGAAAATATTGCAGGAACTGAAAGCTTGC
31725
Gallus_gallus                    31681 .............................................
31725

Anas_platyrhynchos_platyrhynchos 31726 TCATGTCATAAGCATGGCAGTAATATGGAGACTTAAAAAGCAATT
31770
Gallus_gallus                    31726 .............................................
31770

Anas_platyrhynchos_platyrhynchos 31771 CAGTTATTCTAAAATGACATCTCTAAAGAGGAATATTTAAATAAG
31815
Gallus_gallus                    31771 .............................................
31815

Anas_platyrhynchos_platyrhynchos 31816 TATCTTTGGCATTATTGTTTTAGTGCTTGGAGGGGAGTGATAGTC
31860
Gallus_gallus                    31816 .............................................
31860

Anas_platyrhynchos_platyrhynchos 31861 TGTAGGGCTGGAACTGAAGTAAATTAAAGGCTGAATCTTCTTTTT
31905
Gallus_gallus                    31861 .............................................
31905

Anas_platyrhynchos_platyrhynchos 31906 GATCTCTGCCTTTTATGGTAACTAAATCAGAAGATGATAACCAAA
31950
Gallus_gallus                    31906 .............................................
31950

Anas_platyrhynchos_platyrhynchos 31951 AATTTTTGTAATCTTTTTGGAAGATGCTGTCTGATTATCTATGGA
31995
Gallus_gallus                    31951 .............................................
31995

Anas_platyrhynchos_platyrhynchos 31996 GTTATGTGATTCAGTTTTAACTGAACATTATTCTAAACTTTCTGG
32040
Gallus_gallus                    31996 .............................................
32040

Anas_platyrhynchos_platyrhynchos 32041 AAACTGCTAGAATTAAACACATAAGTTCCACTGATAAATTTACCA
32085
Gallus_gallus                    32041 .............................................
32085

Anas_platyrhynchos_platyrhynchos 32086 ATCTATTAAAATAAGACTTCAGAAGTACAGAGCTAGAAAAAACAG
32130
Gallus_gallus                    32086 .............................................
32130

Anas_platyrhynchos_platyrhynchos 32131 CTTAAGAAACTGTGCCTTGTTTTGTTTTGTTTAATGGGAAAAAAG
32175
```

Gallus_gallus                    32131 .............................................
32175

Anas_platyrhynchos_platyrhynchos 32176 GATATTAAAACCAGGAGAGTTTTCAAACACTTAGAGATCTGTCTG
32220
Gallus_gallus                    32176 .............................................
32220

Anas_platyrhynchos_platyrhynchos 32221 TGTATAACATACCTTACACACAATTCATTTTAACTTACACATCAC
32265
Gallus_gallus                    32221 .............................................
32265

Anas_platyrhynchos_platyrhynchos 32266 TGGATGGATTAAACAGTATTTTAAAGGTTTTTTTTTTTTTTTTTT
32310
Gallus_gallus                    32266 .............................................
32310

Anas_platyrhynchos_platyrhynchos 32311 TTTTTTCATGAAGTGAATAATCATCAAAGTCAAGAGATATTTATG
32355
Gallus_gallus                    32311 .............................................
32355

Anas_platyrhynchos_platyrhynchos 32356 CAGCCCTAACAGGATCGGATATAGGATTTAATTGTCTACATTACA
32400
Gallus_gallus                    32356 .............................................
32400

Anas_platyrhynchos_platyrhynchos 32401 ATGGCAGTCCCAATAAAGACATGAATAAAAATACATTTTTTCATG
32445
Gallus_gallus                    32401 .............................................
32445

Anas_platyrhynchos_platyrhynchos 32446 TTCTTTTTTTGGGAGTACAAATGGGTTACCTGAAATTTAGCATGG
32490
Gallus_gallus                    32446 .............................................
32490

Anas_platyrhynchos_platyrhynchos 32491 ACTAAATAACTAATGAATAAAATCAGGACTCTAAAACTATCCTTG
32535
Gallus_gallus                    32491 .............................................
32535

Anas_platyrhynchos_platyrhynchos 32536 CTTTTAATACATTATAAGTTATTTGTTGAGCGCCATCCATTAAAA
32580
Gallus_gallus                    32536 .............................................
32580

Anas_platyrhynchos_platyrhynchos 32581 TTCATAATGTACATAAAACAAAAGTATGCGAGCCCTGTCACTTGT
32625
Gallus_gallus                    32581 .............................................
32625

Anas_platyrhynchos_platyrhynchos 32626 CAATTTAAAGGCAAGGAAAATACTGAAAATGCCTTCAAGTCTATG
32670
Gallus_gallus                    32626 .............................................
32670

Anas_platyrhynchos_platyrhynchos 32671 AAAGCCAGATGTGTAATATGAAAGATTGTGAAGAGGTCACTTAGC
32715
Gallus_gallus                    32671 .............................................
32715

```
Anas_platyrhynchos_platyrhynchos 32716 ATGCCAAAATAATTTATTACATAAAACAGTCTGATTGTTAAACTA
32760
Gallus_gallus                    32716 .............................................
32760

Anas_platyrhynchos_platyrhynchos 32761 TACAGCATAGGAAAACTTCAGAGGGACTCTTAATTATCTGTTAAA
32805
Gallus_gallus                    32761 .............................................
32805

Anas_platyrhynchos_platyrhynchos 32806 AAGAAGTTTTATATATTTAACTAGAACATCTTGAGAATCTTGAGC
32850
Gallus_gallus                    32806 .............................................
32850

Anas_platyrhynchos_platyrhynchos 32851 TGGGAGGTATGGGTAACATAACCTTTTTGTGACATTGTGTTGATT
32895
Gallus_gallus                    32851 .............................................
32895

Anas_platyrhynchos_platyrhynchos 32896 ATTGCCCTGCTGCAGAGCATTCATTCTCTTTAGTCCTACGTAAAG
32940
Gallus_gallus                    32896 .............................................
32940

Anas_platyrhynchos_platyrhynchos 32941 AATGAAGCTACAAGAGGCTAACTTCCCCCTTCAATGTAGATATGA
32985
Gallus_gallus                    32941 .............................................
32985

Anas_platyrhynchos_platyrhynchos 32986 GCCCTTCTGTTTCCAGAAGTAGATAAGAATATTTATAACTCAAAA
33030
Gallus_gallus                    32986 .............................................
33030

Anas_platyrhynchos_platyrhynchos 33031 TTTTAACTTTGCTTTGAAATAAAAATAAATCTGCATTACTGTGAA
33075
Gallus_gallus                    33031 .............................................
33075

Anas_platyrhynchos_platyrhynchos 33076 TTTTACCGAAATAAAGTTAATCTTACCTTTTGTAGATATCAGAAT
33120
Gallus_gallus                    33076 .............................................
33120

Anas_platyrhynchos_platyrhynchos 33121 TGGTTATTCTTTGATAACTTTGTTTTGATTGTACCAGATCTAAAA
33165
Gallus_gallus                    33121 .............................................
33165

Anas_platyrhynchos_platyrhynchos 33166 GTATCATCAAATGGCACTCTGGAAAAAAAAAAAACAAACCACTAA
33210
Gallus_gallus                    33166 .............................................
33210

Anas_platyrhynchos_platyrhynchos 33211 TAATAACGTCCATCGTAATAACTGTATATAAAACTGGCTTTTCTA
33255
Gallus_gallus                    33211 .............................................
33255

Anas_platyrhynchos_platyrhynchos 33256 AGTCAATGACGTACTAAGCTCAGTGTAAACAACTCATATAAAATT
33300
Gallus_gallus                    33256 .............................................
```

33300

Anas_platyrhynchos_platyrhynchos 33301 TATATTTCAGCCAAAAGAGTGTCATTAAGTAAAAGAACACCAAGA
33345
Gallus_gallus 33301 .............................................
33345

Anas_platyrhynchos_platyrhynchos 33346 AAGTCATTCTAACATTATTTAATGTAAATTTGATACACTTTTCCG
33390
Gallus_gallus 33346 .............................................
33390

Anas_platyrhynchos_platyrhynchos 33391 TAATGTTTGATGGGCTTTTGTCCTGCATCACCTTAGTATAACTAA
33435
Gallus_gallus 33391 .............................................
33435

Anas_platyrhynchos_platyrhynchos 33436 CAACAAGGTCCATGTAACCATCTTATTACAGATAATGATTTATAT
33480
Gallus_gallus 33436 .............................................
33480

Anas_platyrhynchos_platyrhynchos 33481 GACTCAAATCAGTAGTTGCAAAATCTTGTAAAGTGGAGGTATGGA
33525
Gallus_gallus 33481 .............................................
33525

Anas_platyrhynchos_platyrhynchos 33526 AATGTTATCTGACCATGTTAGAACCAAGTTACAGTTAAAACATTC
33570
Gallus_gallus 33526 .............................................
33570

Anas_platyrhynchos_platyrhynchos 33571 TTTTTAAGCTATTTTCCTTAAAGGCTCCCAATGAAATAACTGAGT
33615
Gallus_gallus 33571 .............................................
33615

Anas_platyrhynchos_platyrhynchos 33616 ACTCTTAGATTAAAAAGCTTTCCCTAAAGATTTTTGCTAAAAAGT
33660
Gallus_gallus 33616 .............................................
33660

Anas_platyrhynchos_platyrhynchos 33661 TTTTCCCTAAAGATTTTTTGCTTGGTTTGCTTGGTTTGCTGCAGC
33705
Gallus_gallus 33661 .............................................
33705

Anas_platyrhynchos_platyrhynchos 33706 GTAAACCTTTACTTAATTGTCGTATTTCACACAGATATCTGAGAA
33750
Gallus_gallus 33706 .............................................
33750

Anas_platyrhynchos_platyrhynchos 33751 GAAAATTACCTTTTTGTTTGGAAAGCCTTTGATGTACTTGAGAAC
33795
Gallus_gallus 33751 .............................................
33795

Anas_platyrhynchos_platyrhynchos 33796 TACTATCAAGCTTCTCTTCAATATTTTAACCTTTTCTCCTAAAGT
33840
Gallus_gallus 33796 .............................................
33840

Anas_platyrhynchos_platyrhynchos 33841 ATATTTATATGCATATATATGTATAAAAACACATATATATTTGTA

33885
Gallus_gallus 33841 .............................................
33885

Anas_platyrhynchos_platyrhynchos 33886 CAACTATGCACACATATCTGCACTATAGTTTTCTGACCACACTTG
33930
Gallus_gallus 33886 .............................................
33930

Anas_platyrhynchos_platyrhynchos 33931 ATCTCTACCAATAATTTATTCTAGTTTGTCAGTATTCTCCTCCAG
33975
Gallus_gallus 33931 .............................................
33975

Anas_platyrhynchos_platyrhynchos 33976 ACCTTTACAAAAGTGGAGCGTGAGATTAGTTAATTTATCTTGCCA
34020
Gallus_gallus 33976 .............................................
34020

Anas_platyrhynchos_platyrhynchos 34021 ATGACACTCCTGTATAGTTACCCCTATAACAGAACCTAGTCTTGT
34065
Gallus_gallus 34021 .............................................
34065

Anas_platyrhynchos_platyrhynchos 34066 TTGATCCCTAGCAAGATGATGTTGTTTAATTTGTGATTTATCTCC
34110
Gallus_gallus 34066 .............................................
34110

Anas_platyrhynchos_platyrhynchos 34111 TGATTGTAATCTGTAGTATCTCCCCTCAGGCAGCCATTACCTAAC
34155
Gallus_gallus 34111 .............................................
34155

Anas_platyrhynchos_platyrhynchos 34156 TTGCTAATGTTTAACTGATTATTTCTACTTAAGCACAACTCCCTC
34200
Gallus_gallus 34156 .............................................
34200

Anas_platyrhynchos_platyrhynchos 34201 TAGCCTTCCCTGAATTACATTCTGCTATTTATGCCCATAAAGCAA
34245
Gallus_gallus 34201 .............................................
34245

Anas_platyrhynchos_platyrhynchos 34246 GGTTAATTTGCATTCTAATCTTGCTGCTGAACCACCTCTTTCTCA
34290
Gallus_gallus 34246 .............................................
34290

Anas_platyrhynchos_platyrhynchos 34291 GGCTGGTCTCATATGCAATTTCAGCAAAGACATCTTCTATTTCAT
34335
Gallus_gallus 34291 .............................................
34335

Anas_platyrhynchos_platyrhynchos 34336 CACTCAAATGACCAACAAGAACATTTAATAATACTGGGTCCAAGT
34380
Gallus_gallus 34336 .............................................
34380

Anas_platyrhynchos_platyrhynchos 34381 TAGGGCCCTCTTGAACCTCAGACACACGTCATACTTAATAATGAA
34425
Gallus_gallus 34381 .............................................
34425

```
Anas_platyrhynchos_platyrhynchos 34426 AGTATTTCTTACATGAAGATCAGGATTACAATAGATTTAAATTCA
34470
Gallus_gallus                    34426 .............................................
34470

Anas_platyrhynchos_platyrhynchos 34471 GTCTGCAAGACAACAGTAATTTTATCAGCTAACAAGTTGTGTAGG
34515
Gallus_gallus                    34471 .............................................
34515

Anas_platyrhynchos_platyrhynchos 34516 TGATCAGAAAATAGAACAATTCTTTGAGGGCAACATGAATACATT
34560
Gallus_gallus                    34516 .............................................
34560

Anas_platyrhynchos_platyrhynchos 34561 TAAAGTTGGTTTATTCCCCATTAAAGGAAAATAAGATGCTATGGA
34605
Gallus_gallus                    34561 .............................................
34605

Anas_platyrhynchos_platyrhynchos 34606 GAATTCATCTAGACATATTTTAAGTGCAAGTCCCAGGGTCAGGAT
34650
Gallus_gallus                    34606 .............................................
34650

Anas_platyrhynchos_platyrhynchos 34651 TATCGAAGTGTTTACGAACTTCTCCAGAATTTCTAGTCCTATAAA
34695
Gallus_gallus                    34651 .............................................
34695

Anas_platyrhynchos_platyrhynchos 34696 TATCGGCCTACATGCATCTGAAATCTCATTTTTAAAAGTGACCAA
34740
Gallus_gallus                    34696 .............................................
34740

Anas_platyrhynchos_platyrhynchos 34741 TATATATAGGAAGATATATCTTCTTGATTTCCAGCAATATTTTCA
34785
Gallus_gallus                    34741 .............................................
34785

Anas_platyrhynchos_platyrhynchos 34786 CATACACCTTTAAATCACTTTGAAAATCAAAGCTTTTTGAATCAG
34830
Gallus_gallus                    34786 .............................................
34830

Anas_platyrhynchos_platyrhynchos 34831 CTTCCTGGTACTTTTAGAACTTCACGATCTGATATGGATTGCAGT
34875
Gallus_gallus                    34831 .............................................
34875

Anas_platyrhynchos_platyrhynchos 34876 TCTTAATTTTATATCTCGCCACAATATAGAAAGAAGCTAACATAT
34920
Gallus_gallus                    34876 .............................................
34920

Anas_platyrhynchos_platyrhynchos 34921 TCATACACCCTAATAACTTCAGCAAAACTCCAAAGCTGAATATCT
34965
Gallus_gallus                    34921 .............................................
34965

Anas_platyrhynchos_platyrhynchos 34966 TCTAGATGTACAAGGGTAATAATAAGAGAAAGAAAGACTGAATCT
35010
```

Gallus_gallus 34966 .............................................
35010

Anas_platyrhynchos_platyrhynchos 35011 CAAATAATAGTTCTGAATTTACGTCTTATGTTATATCACCAAAAA
35055
Gallus_gallus 35011 .............................................
35055

Anas_platyrhynchos_platyrhynchos 35056 GACTAATAGCTGTGAAGAAGCAGAGAAAAAATTACCTTCTCAGAG
35100
Gallus_gallus 35056 .............................................
35100

Anas_platyrhynchos_platyrhynchos 35101 TGTACAAATGCTCCATTAGAGAAGAGAAATAATTCAGTATTAACC
35145
Gallus_gallus 35101 .............................................
35145

Anas_platyrhynchos_platyrhynchos 35146 CAGTTTCTGCAGATAATTCTGCTTTGCCTGAGTGAAGACTGAGTA
35190
Gallus_gallus 35146 .............................................
35190

Anas_platyrhynchos_platyrhynchos 35191 ATTCTGAAACTGTACTACAAAGGTTGTTCTTAACAATAAAGAACA
35235
Gallus_gallus 35191 .............................................
35235

Anas_platyrhynchos_platyrhynchos 35236 CTTGTTTACCAAGTACTGCATTCCACAGAAAGGCACAGGACTTAC
35280
Gallus_gallus 35236 .............................................
35280

Anas_platyrhynchos_platyrhynchos 35281 TGGAAAAAAAACATCAGATGATGATTCAATGTTTGGAACTTAATA
35325
Gallus_gallus 35281 .............................................
35325

Anas_platyrhynchos_platyrhynchos 35326 AATGTGTTCTGATTTAAAATTCCAGCATTTGGAAGTGGAGGAGGA
35370
Gallus_gallus 35326 .............................................
35370

Anas_platyrhynchos_platyrhynchos 35371 ATAAACCCGTTTGTGAGGAATAACAAGAGATATGTGCAACAGAAT
35415
Gallus_gallus 35371 .............................................
35415

Anas_platyrhynchos_platyrhynchos 35416 AAAATATACTTAGTGCAGAGTTTTTCCAGTCAAAACCCTGGATCT
35460
Gallus_gallus 35416 .............................................
35460

Anas_platyrhynchos_platyrhynchos 35461 TGAAAAAAGGTTTTGATTACTTACGTCAATACATTGATAATACAG
35505
Gallus_gallus 35461 .............................................
35505

Anas_platyrhynchos_platyrhynchos 35506 TAAGAAGTAAACACCATTTTGTGATCATTATGAAAAAAAGGTAGT
35550
Gallus_gallus 35506 .............................................
35550

Anas_platyrhynchos_platyrhynchos 35551 GTAAATTAAAAAGTATTTTTCAGTTTGGTTATCTCACTTGGGATG
35595
Gallus_gallus 35551 .............................................
35595

Anas_platyrhynchos_platyrhynchos 35596 TTCATAGCAACCAGTCTGTTGAGTAAAATATGAGACAAGAATCTC
35640
Gallus_gallus 35596 .............................................
35640

Anas_platyrhynchos_platyrhynchos 35641 TCCATTCCATAGAACAGAAATGCACTGCAGTTTGTCAGCAGCTGC
35685
Gallus_gallus 35641 .............................................
35685

Anas_platyrhynchos_platyrhynchos 35686 TCCCATTCAGCTTGACTGCAAAGAAGGGAAATTTTTTTTGTTACT
35730
Gallus_gallus 35686 .............................................
35730

Anas_platyrhynchos_platyrhynchos 35731 GTTTAACAGGAACACTTTTAAGAAACTATCCACATCAAACACTTC
35775
Gallus_gallus 35731 .............................................
35775

Anas_platyrhynchos_platyrhynchos 35776 AGTGTTGGTCCAGAAATATTTACTAGAAAAAACAGTGTGCAAAGC
35820
Gallus_gallus 35776 .............................................
35820

Anas_platyrhynchos_platyrhynchos 35821 AAGGTGACAAACAAGAATTCTACAGAAAGTGTAATAAAATGTATG
35865
Gallus_gallus 35821 .............................................
35865

Anas_platyrhynchos_platyrhynchos 35866 GAACGTTAATAATTACTATTTTAATACGGATGTATATTTAATTTA
35910
Gallus_gallus 35866 .............................................
35910

Anas_platyrhynchos_platyrhynchos 35911 GTAACACCTTGTAAACAAAAGGGCATAGCATATTTTAAACAAATA
35955
Gallus_gallus 35911 .............................................
35955

Anas_platyrhynchos_platyrhynchos 35956 CCAGACTCTAACAAAAAAAATAAAAATCATTAGTGTACATGTATG
36000
Gallus_gallus 35956 .............................................
36000

Anas_platyrhynchos_platyrhynchos 36001 TATGCATTGTACCGCTTTGAAATTCAAAGGGAGTTTTCAAGAAGG
36045
Gallus_gallus 36001 .............................................
36045

Anas_platyrhynchos_platyrhynchos 36046 ACTTGGGGGTTCTGACGGATAAAAAGCTCAACATGAACCAGCAGT
36090
Gallus_gallus 36046 .............................................
36090

Anas_platyrhynchos_platyrhynchos 36091 GAGCACTTGCAGCCCAGAAGTCCAACTGCATCCTGGGCTGCATCA
36135
Gallus_gallus 36091 .............................................

36135

Anas_platyrhynchos_platyrhynchos 36136 ATAGAGTGGTCATCAGGCAAGGGAGGGGATTGTCCACCTCTGCTC
36180
Gallus_gallus 36136 .............................................
36180

Anas_platyrhynchos_platyrhynchos 36181 TGCCCTTGTGAGTCCCCACCTGGAGTCCTGCATCCAGGTGTGGGG
36225
Gallus_gallus 36181 .............................................
36225

Anas_platyrhynchos_platyrhynchos 36226 CCCCCAGCACCAGAAGGATGTGGGGCTGTTAGAGTGGGTCCAGAG
36270
Gallus_gallus 36226 .............................................
36270

Anas_platyrhynchos_platyrhynchos 36271 GAGGGCCACAAAGATGCTCAGAGGGCTGGAGCACCTCTCCTATGA
36315
Gallus_gallus 36271 .............................................
36315

Anas_platyrhynchos_platyrhynchos 36316 AGAAAGGCTGAGAGAGTTGGGACTCTTCATCCTGAAGAAAAGAAG
36360
Gallus_gallus 36316 .............................................
36360

Anas_platyrhynchos_platyrhynchos 36361 GCTCCGGGAAGACCTTTTTTGCATTGCAGCTTTTCAAAACTTAAA
36405
Gallus_gallus 36361 .............................................
36405

Anas_platyrhynchos_platyrhynchos 36406 AGGGGCTTGTAAAAAACGATGGAGAGCAACTTTGTGCTTGGGTAG
36450
Gallus_gallus 36406 .............................................
36450

Anas_platyrhynchos_platyrhynchos 36451 GTAATGATAGAACAAAGGGGAATGGTTTTAAACTAAAAGAAGGGA
36495
Gallus_gallus 36451 .............................................
36495

Anas_platyrhynchos_platyrhynchos 36496 GCTTTAGATTAAACGTTGGGAGGAAATTCTTCACTCAGAGGGTGG
36540
Gallus_gallus 36496 .............................................
36540

Anas_platyrhynchos_platyrhynchos 36541 TGAGGCACTGGCACAGCCTGGCCAGAGAAGCTGTGGATGCAACAT
36585
Gallus_gallus 36541 .............................................
36585

Anas_platyrhynchos_platyrhynchos 36586 CTCTGGAAGTGTTCAAAGCCAGGCTGAATGGGTCCCTGGGCAACC
36630
Gallus_gallus 36586 .............................................
36630

Anas_platyrhynchos_platyrhynchos 36631 AGATCTACCATGTGGCTTCCCTGCCCATGGCAGTGGGGTTGGGAC
36675
Gallus_gallus 36631 .............................................
36675

Anas_platyrhynchos_platyrhynchos 36676 CAGATGATCTTTAAGGTCCCTTCAACCCAAGCTGTTCTGTGATTC

36720
Gallus_gallus 36676 .............................................
36720

Anas_platyrhynchos_platyrhynchos 36721 TATGATATATGCACCCAGCAGGATATAAATAAATTAGGTTTTGAA
36765
Gallus_gallus 36721 .............................................
36765

Anas_platyrhynchos_platyrhynchos 36766 AAGAGGTTCAGTTAATCAGCCTAGCTAGCAGCAAAAACTATGCTG
36810
Gallus_gallus 36766 .............................................
36810

Anas_platyrhynchos_platyrhynchos 36811 TCTGATAACTGCTAACAATCTTTGCAATTGTTATGAAGAAACACC
36855
Gallus_gallus 36811 .............................................
36855

Anas_platyrhynchos_platyrhynchos 36856 AGAATACCTGTACAGTGATTTTACCTAAGTAGAGAAGATTTTAAA
36900
Gallus_gallus 36856 .............................................
36900

Anas_platyrhynchos_platyrhynchos 36901 GGACGGATTAGACTAGCAGTTATAAAGCAAGGTCCACAGTTCACC
36945
Gallus_gallus 36901 .............................................
36945

Anas_platyrhynchos_platyrhynchos 36946 ATGTTGCCTCACTTACAGGACTTGAATGGCTTGACACCTCAAGAT
36990
Gallus_gallus 36946 .............................................
36990

Anas_platyrhynchos_platyrhynchos 36991 CCCTTTTACGTCTCTACTTCTACAATCTTAAAATTCAAGTACTTC
37035
Gallus_gallus 36991 .............................................
37035

Anas_platyrhynchos_platyrhynchos 37036 TATAAAGAACGAACTGAACAAACTTTATAAAAATAATACGAACTT
37080
Gallus_gallus 37036 .............................................
37080

Anas_platyrhynchos_platyrhynchos 37081 CATCAACTTTCAGATGTTTGAAATTGTAAACTGAAAGATGTACCT
37125
Gallus_gallus 37081 .............................................
37125

Anas_platyrhynchos_platyrhynchos 37126 TTACTACATTGTGGCTGAAATGTGTGCTACAATCTTGCAGTTAAC
37170
Gallus_gallus 37126 .............................................
37170

Anas_platyrhynchos_platyrhynchos 37171 TCATTATGCATGGAGATAATATAGCTTTTAATACTTGGTTTTAAA
37215
Gallus_gallus 37171 .............................................
37215

Anas_platyrhynchos_platyrhynchos 37216 AAACAGTTATGAGCAAAGCCAGAAATCACACTTATATTCCCATAT
37260
Gallus_gallus 37216 .............................................
37260

```
Anas_platyrhynchos_platyrhynchos 37261 GCTCATAATAATAATTGAAGACTGCAACAGCAGCCTTTTTTTTT
37305
Gallus_gallus                    37261 .............................................
37305

Anas_platyrhynchos_platyrhynchos 37306 TCAAGATTGGATCATTTAAATATACAGCAATTTCTGGCACCGTAT
37350
Gallus_gallus                    37306 .............................................
37350

Anas_platyrhynchos_platyrhynchos 37351 TTTAGATCCTACATTTGAAACAAAGACAATGGTTACTCTGTGTTT
37395
Gallus_gallus                    37351 .............................................
37395

Anas_platyrhynchos_platyrhynchos 37396 AATAGTGCAGCGAAATGTAACCCTATTTTATGACTGAAGCCTCTG
37440
Gallus_gallus                    37396 .............................................
37440

Anas_platyrhynchos_platyrhynchos 37441 GGAATTATCACTGTTTTAAAAAAAGAAACATGGATAAAACATGAA
37485
Gallus_gallus                    37441 .............................................
37485

Anas_platyrhynchos_platyrhynchos 37486 AGCATATAGAATACTACAAACCTTGGAACATATACGTTGCCAATA
37530
Gallus_gallus                    37486 .............................................
37530

Anas_platyrhynchos_platyrhynchos 37531 ACTCCTTCCCACTGTAGTGTAAACAGGTCATGGTACTTTTCTAAG
37575
Gallus_gallus                    37531 .............................................
37575

Anas_platyrhynchos_platyrhynchos 37576 ATTTCCTTCATTTTTTGGGAGGGACTTAAAGCTTCTGTAACAATA
37620
Gallus_gallus                    37576 .............................................
37620

Anas_platyrhynchos_platyrhynchos 37621 GAATGTCATAATTGTAAGGTTAAGATGCTACAAAGTATGTTGGGC
37665
Gallus_gallus                    37621 .............................................
37665

Anas_platyrhynchos_platyrhynchos 37666 AATCTTATTTTCAAACATTTAAAAATGACAATATATATATTTAGG
37710
Gallus_gallus                    37666 .............................................
37710

Anas_platyrhynchos_platyrhynchos 37711 AATTTTTTTAGCCCGTCATTAGATCTTAAGGTTTTTAAAAACTAA
37755
Gallus_gallus                    37711 .............................................
37755

Anas_platyrhynchos_platyrhynchos 37756 GTCCAGATTTTAAAAATGCGTCAATTGCAATGTTGAAAACACCAG
37800
Gallus_gallus                    37756 .............................................
37800

Anas_platyrhynchos_platyrhynchos 37801 CACTGTTCGTGCAGTTCCATTTCAGAGACTCAGAAAGAATTTAGA
37845
```

Gallus_gallus                    37801 .............................................
37845

Anas_platyrhynchos_platyrhynchos 37846 TGGACAGAGAAAATAACTGAGTTTCTATAGCCCTGATTTTAGAAG
37890
Gallus_gallus                    37846 .............................................
37890

Anas_platyrhynchos_platyrhynchos 37891 GCAAGATTCCAAACCCACCTCTGCTCTCCAAGGACATTGAATATT
37935
Gallus_gallus                    37891 .............................................
37935

Anas_platyrhynchos_platyrhynchos 37936 TAATGACAATTTACCTTGTCTGACTATATTTTCATAATACTGTAA
37980
Gallus_gallus                    37936 .............................................
37980

Anas_platyrhynchos_platyrhynchos 37981 AATGCAGATACACTCTTACAGCCAGATATTGAGTACGTAGTAAAC
38025
Gallus_gallus                    37981 .............................................
38025

Anas_platyrhynchos_platyrhynchos 38026 CGTTTATTTTTATCTGCTGTGAAGGAGAAACTGCAAACATATGGA
38070
Gallus_gallus                    38026 .............................................
38070

Anas_platyrhynchos_platyrhynchos 38071 AGCATATTCTATCAGTATGTTCAGTACTGTTCAGTAGCTTCTTTT
38115
Gallus_gallus                    38071 .............................................
38115

Anas_platyrhynchos_platyrhynchos 38116 GTTTTGATTTTTTGTTTATTAACTATCTCAATTAAAAACCTCCAT
38160
Gallus_gallus                    38116 .............................................
38160

Anas_platyrhynchos_platyrhynchos 38161 TAAATCAATTTTTGGCATTTGATATATAAATGTGTTTGTTAACAA
38205
Gallus_gallus                    38161 .............................................
38205

Anas_platyrhynchos_platyrhynchos 38206 ATGTGAATCAATTAAACTATTACACATACTGGGAGAAAAAAAATT
38250
Gallus_gallus                    38206 .............................................
38250

Anas_platyrhynchos_platyrhynchos 38251 AAGTAAAATTGTGTTACTAAAGTATTTGTGTCAGACTGAATTGAT
38295
Gallus_gallus                    38251 .............................................
38295

Anas_platyrhynchos_platyrhynchos 38296 ATAGTAAATGCCTACTGCCTTGGTAAGTAGACTCCAAAAGAAAAC
38340
Gallus_gallus                    38296 .............................................
38340

Anas_platyrhynchos_platyrhynchos 38341 AGCAACCCTTTCAAATATCCTTACACAAGTGCCTGTAACTATCAT
38385
Gallus_gallus                    38341 .............................................
38385

```
Anas_platyrhynchos_platyrhynchos 38386 AATACCTTCCACCTTTCAAAAGACTATAACACAACATGCAAAAGC
38430
Gallus_gallus                    38386 .............................................
38430

Anas_platyrhynchos_platyrhynchos 38431 TTTGTGTGTTACACATGCTTTATATGTAGGCATTAGCTTGAGAAT
38475
Gallus_gallus                    38431 .............................................
38475

Anas_platyrhynchos_platyrhynchos 38476 CGGTATATACGTTTATATATAAGTTCCATTGTTCCTGTTCTGGTC
38520
Gallus_gallus                    38476 .............................................
38520

Anas_platyrhynchos_platyrhynchos 38521 CATTCCCATTCAGTACTGTTTGTTCCTAGTCATTCATGTATTTCA
38565
Gallus_gallus                    38521 .............................................
38565

Anas_platyrhynchos_platyrhynchos 38566 TGCAAATTTCTAGTATAATAAAGAAGATTATACAATGCATCATTT
38610
Gallus_gallus                    38566 .............................................
38610

Anas_platyrhynchos_platyrhynchos 38611 CAGTTAACCTATATCCTTCCCTGCACTACTGATGTTGTATCTTAA
38655
Gallus_gallus                    38611 .............................................
38655

Anas_platyrhynchos_platyrhynchos 38656 CGGCATATATCTGTAAAACTAAAAATGCAGTAACAAGTTCTTGAA
38700
Gallus_gallus                    38656 .............................................
38700

Anas_platyrhynchos_platyrhynchos 38701 AGAAAGCATGTATTTTTAATTGGTTTATATAGAAAGGAAGATTAC
38745
Gallus_gallus                    38701 .............................................
38745

Anas_platyrhynchos_platyrhynchos 38746 TCAAGTAACTATTTTCTTGAAGGTAAGTAGGTCATTATTTAGTGA
38790
Gallus_gallus                    38746 .............................................
38790

Anas_platyrhynchos_platyrhynchos 38791 TATGAAAAAAAAATAAGACTAATTTCTTTCTGGAACTTGAAACAG
38835
Gallus_gallus                    38791 .............................................
38835

Anas_platyrhynchos_platyrhynchos 38836 GTTATATTTTTTCATTGAGGCACCATTTGACAGGTGATAAGCAGT
38880
Gallus_gallus                    38836 .............................................
38880

Anas_platyrhynchos_platyrhynchos 38881 ACTTTTCTGACAAAAATCTAAGCTTTTTTTGTCAACACCTCAATG
38925
Gallus_gallus                    38881 .............................................
38925

Anas_platyrhynchos_platyrhynchos 38926 GTATGACTTCATTTTCTGTTAAAAAAGAAAAAAAAAATTAAATTG
38970
Gallus_gallus                    38926 .............................................
```

38970

Anas_platyrhynchos_platyrhynchos 38971 CATGGCTAGGTTGTGGAGGTAATATTATTTAATAGATATTACCTG
39015
Gallus_gallus 38971 .............................................
39015

Anas_platyrhynchos_platyrhynchos 39016 GCTCTCTTCCTTCATTATAGGGGTCCACAATATCTGATAGGCAGT
39060
Gallus_gallus 39016 .............................................
39060

Anas_platyrhynchos_platyrhynchos 39061 GAAGTGTAAACAATGCAATATAAACCAGAAAAAAAAAGTTTATCT
39105
Gallus_gallus 39061 .............................................
39105

Anas_platyrhynchos_platyrhynchos 39106 GAACTGATTCACATACGTAATCATTTGGGAAAGGATACATTTAAG
39150
Gallus_gallus 39106 .............................................
39150

Anas_platyrhynchos_platyrhynchos 39151 ATTACGAGTATCAACAGGCATGCAATTAGAGGGTAAAACACGATG
39195
Gallus_gallus 39151 .............................................
39195

Anas_platyrhynchos_platyrhynchos 39196 AACTGGTGCCTATAATTAAAAGAATGTATACAGTATTACGTGCAG
39240
Gallus_gallus 39196 .............................................
39240

Anas_platyrhynchos_platyrhynchos 39241 AAGCCTTAATCAAACATTATTATTGTATCTGCAGTGTTAAACAAC
39285
Gallus_gallus 39241 .............................................
39285

Anas_platyrhynchos_platyrhynchos 39286 TAACAAAAAGATGTCCTGAAGTTTTGAAAGTAGGCTTAAAAGCAG
39330
Gallus_gallus 39286 .............................................
39330

Anas_platyrhynchos_platyrhynchos 39331 CACAATACCCAAGTGTCAATGTAAATCAAAAGCATAAGCTATCCT
39375
Gallus_gallus 39331 .............................................
39375

Anas_platyrhynchos_platyrhynchos 39376 GCATTATACAGAATGCATATATGTACATATTGTATATATTTACAC
39420
Gallus_gallus 39376 .............................................
39420

Anas_platyrhynchos_platyrhynchos 39421 ATGTGAACATATATAGTTATATATGCATATAAAATATAAAACAGC
39465
Gallus_gallus 39421 .............................................
39465

Anas_platyrhynchos_platyrhynchos 39466 ATATAAACTATACCATATTGCTGGGGGAGGTAGTTTAGTATTTTG
39510
Gallus_gallus 39466 .............................................
39510

Anas_platyrhynchos_platyrhynchos 39511 AGACAGCTCCCTTTGTGAGTTGTGTATGTCTCGATTATGCAGAGG

```
39555
Gallus_gallus                     39511 .............................................
39555

Anas_platyrhynchos_platyrhynchos 39556 ATATGCTTTTTCAAAGAAACAACTCTATAAAATCAACAAGAGCAA
39600
Gallus_gallus                     39556 .............................................
39600

Anas_platyrhynchos_platyrhynchos 39601 GATTCTTATTATGTAATACTTATTTCCACTTCAGACTTCTAGAGT
39645
Gallus_gallus                     39601 .............................................
39645

Anas_platyrhynchos_platyrhynchos 39646 ATATAAAGAGCTTCTATGCAAGCAACTTGTATTCCTTTAACAATA
39690
Gallus_gallus                     39646 .............................................
39690

Anas_platyrhynchos_platyrhynchos 39691 TTTTTACACAACAAGAAGAGCTACCCATATTTATTTTATTTAAAA
39735
Gallus_gallus                     39691 .............................................
39735

Anas_platyrhynchos_platyrhynchos 39736 AAAATGGAAAGTTCACCATTTTGACATTCTTTTTCTGTTCGGCTT
39780
Gallus_gallus                     39736 .............................................
39780

Anas_platyrhynchos_platyrhynchos 39781 TTGGTTTTGGTTCTTTAGACCTTTGAACATCTTTAATTTCACTCT
39825
Gallus_gallus                     39781 .............................................
39825

Anas_platyrhynchos_platyrhynchos 39826 CTGTGGAATTGAAAAGAAGAGACTTTTTTTTTAAGACATGAGCAG
39870
Gallus_gallus                     39826 .............................................
39870

Anas_platyrhynchos_platyrhynchos 39871 TTTCAGAAAATACAGCTTTCTTAAAGGATTGTTTATATTCATTAG
39915
Gallus_gallus                     39871 .............................................
39915

Anas_platyrhynchos_platyrhynchos 39916 GATGTAATAATAACAACAGATTAGAAAAATACATCTTAATTTCAC
39960
Gallus_gallus                     39916 .............................................
39960

Anas_platyrhynchos_platyrhynchos 39961 ATTGCAGCAAGGCAAGGACATTCCCCAAACCCTCTTGCCTAAAAA
40005
Gallus_gallus                     39961 .............................................
40005

Anas_platyrhynchos_platyrhynchos 40006 TAGTAGCTTCAAAGTGTAAAACAACCAACACACACAATCTCTTAG
40050
Gallus_gallus                     40006 .............................................
40050

Anas_platyrhynchos_platyrhynchos 40051 ATGTGCAAATGCTTCACTACGTTACGCGTTCAGGAGAAAAGTGTT
40095
Gallus_gallus                     40051 .............................................
40095
```

Anas_platyrhynchos_platyrhynchos 40096 GAATAGTGACTACTGGAGCACTTTTTAAGCCTATTGCTATATCTA
40140
Gallus_gallus 40096 .............................................
40140

Anas_platyrhynchos_platyrhynchos 40141 TTCAACATTGATGTTACTGGAATAACTGAAGCTTCTGTGAAATAC
40185
Gallus_gallus 40141 .............................................
40185

Anas_platyrhynchos_platyrhynchos 40186 ACAGAAATCAAAAGAATAAAGGAAGTTTTATATTTGAATCTGATT
40230
Gallus_gallus 40186 .............................................
40230

Anas_platyrhynchos_platyrhynchos 40231 TTTTACTTCAAGAGGCTTCAAAAAAGCAATGGGAACTGTGTTATA
40275
Gallus_gallus 40231 .............................................
40275

Anas_platyrhynchos_platyrhynchos 40276 TTTCTCACTGCACAAATGACTATAACCCAAAGTATAAGTCTCAGA
40320
Gallus_gallus 40276 .............................................
40320

Anas_platyrhynchos_platyrhynchos 40321 TAGTTATCGCTGGAATGCACTGCAAATTAGGGGAAGTTTGTGATA
40365
Gallus_gallus 40321 .............................................
40365

Anas_platyrhynchos_platyrhynchos 40366 TGCATTTGAGTGCATTTGGCCATTGATTCTCTTTTTATTCAGTGG
40410
Gallus_gallus 40366 .............................................
40410

Anas_platyrhynchos_platyrhynchos 40411 CTACTTTCATAATAGCATAGGTAGCTTTTCAATTATGCTTCCTCT
40455
Gallus_gallus 40411 .............................................
40455

Anas_platyrhynchos_platyrhynchos 40456 CATTAATTAACTTCTAAAATTATGCAACCATAAGAATTTTCACTT
40500
Gallus_gallus 40456 .............................................
40500

Anas_platyrhynchos_platyrhynchos 40501 TAAAAGGTGGAAAAACAAATTAAAAAAAAAAAAAAAAGTTCTGAA
40545
Gallus_gallus 40501 .............................................
40545

Anas_platyrhynchos_platyrhynchos 40546 AGTCTTATTTATAACAGATATTTAAGAGTATATGAACGTGTAGAC
40590
Gallus_gallus 40546 .............................................
40590

Anas_platyrhynchos_platyrhynchos 40591 TAAAAATATTCTCATCTCAGTTTGTTTACAGTTCTTTCCATGCAG
40635
Gallus_gallus 40591 .............................................
40635

Anas_platyrhynchos_platyrhynchos 40636 AAATGGCCAAACAGATGACAGTTACACTAAAGTTGCTAAATATTG
40680

```
Gallus_gallus                    40636 .............................................
40680

Anas_platyrhynchos_platyrhynchos 40681 TAGATAAGAATGCAATACAAGAACCTGTATGGAACAGAAGACTGT
40725
Gallus_gallus                    40681 .............................................
40725

Anas_platyrhynchos_platyrhynchos 40726 CATTTTTCACAGTGAGACTTTTCTAAATTTAACCTATTTATTGCC
40770
Gallus_gallus                    40726 .............................................
40770

Anas_platyrhynchos_platyrhynchos 40771 TGAGGCCCACTAACTTTTAAGTGAGTTAAAATGTGTTATCTATAT
40815
Gallus_gallus                    40771 .............................................
40815

Anas_platyrhynchos_platyrhynchos 40816 CTGAGTAAAATGTGGCTTCAAATTTTTATAGGTTCAGTAATAGGA
40860
Gallus_gallus                    40816 .............................................
40860

Anas_platyrhynchos_platyrhynchos 40861 GGTATCTTTGATCTATAAGAAAAAAAAGCTTAGCAGTTCTATAAG
40905
Gallus_gallus                    40861 .............................................
40905

Anas_platyrhynchos_platyrhynchos 40906 TAAAACTGCTTAGCAGTTTTTCTCAGCAGAAAAATGCTATGTGCC
40950
Gallus_gallus                    40906 .............................................
40950

Anas_platyrhynchos_platyrhynchos 40951 TGCATTTCTTACTGATTTTTAGAATAAATTTAAAAAACAAACAAA
40995
Gallus_gallus                    40951 .............................................
40995

Anas_platyrhynchos_platyrhynchos 40996 CAAGCAAAAAACAACAACAACAAAAAAAAGAAGATGCCATAGGGA
41040
Gallus_gallus                    40996 .............................................
41040

Anas_platyrhynchos_platyrhynchos 41041 CAAGGTTATAAAACCATTTCAAAATCTACCCAACCATCTCACTTA
41085
Gallus_gallus                    41041 .............................................
41085

Anas_platyrhynchos_platyrhynchos 41086 ACAAACTCATTTAATTGAAGAGTTTATACAGTGTACAAGTGCCTG
41130
Gallus_gallus                    41086 .............................................
41130

Anas_platyrhynchos_platyrhynchos 41131 TGGTAGTGATAAAAGTATCTACAGCCTGTGGAGTCTGACTTTGCT
41175
Gallus_gallus                    41131 .............................................
41175

Anas_platyrhynchos_platyrhynchos 41176 CATGCGTAGAACAAGAATCACTTAACTGACCTGAGGACTTAACGT
41220
Gallus_gallus                    41176 .............................................
41220
```

```
Anas_platyrhynchos_platyrhynchos 41221 AATCTTAGCTATACATAGAGGAAGTAGCTCAATATATCAATATTC
41265
Gallus_gallus                    41221 .............................................
41265

Anas_platyrhynchos_platyrhynchos 41266 TAAGCAGTCTTACATTAATGTCAAAAGAAAACTAACAGAATTCTA
41310
Gallus_gallus                    41266 .............................................
41310

Anas_platyrhynchos_platyrhynchos 41311 AATTCTTATTTGATTGTTTTCAATGTAGATGAAAATAGTCATAAA
41355
Gallus_gallus                    41311 .............................................
41355

Anas_platyrhynchos_platyrhynchos 41356 GTTTGATGAAAAATCAAAAAAAAAGGTATACATGAAAACAGTTGT
41400
Gallus_gallus                    41356 .............................................
41400

Anas_platyrhynchos_platyrhynchos 41401 AGTTTGAGACCATATGTGTTCAGTTGAAACAAGTCATTTTAGAAT
41445
Gallus_gallus                    41401 .............................................
41445

Anas_platyrhynchos_platyrhynchos 41446 CTACTTTTTAGCATATTCTTTAATATCTAGAGGTTATTTTGGATT
41490
Gallus_gallus                    41446 .............................................
41490

Anas_platyrhynchos_platyrhynchos 41491 AGCTAACATTTCAACTATAATCAGCCTGTATATTTCTAAATAAGT
41535
Gallus_gallus                    41491 .............................................
41535

Anas_platyrhynchos_platyrhynchos 41536 CTATTATACTAGCTATCCCATCTAAAATAATGGAAACATTATATG
41580
Gallus_gallus                    41536 .............................................
41580

Anas_platyrhynchos_platyrhynchos 41581 ACAGGCTATGAAACAATGCTATGACATTATATGACAGGCTATGAA
41625
Gallus_gallus                    41581 .............................................
41625

Anas_platyrhynchos_platyrhynchos 41626 GCAGGCTATGACAGGATATCAAACAATATATGACAGGCTTCCAGT
41670
Gallus_gallus                    41626 .............................................
41670

Anas_platyrhynchos_platyrhynchos 41671 AAATTACATGACACCATGCATATTTACATTTTAACTATCTTCCTT
41715
Gallus_gallus                    41671 .............................................
41715

Anas_platyrhynchos_platyrhynchos 41716 CCTATATCAACATACAAACACAAAAAATGTTTATCTAATAGTACC
41760
Gallus_gallus                    41716 .............................................
41760

Anas_platyrhynchos_platyrhynchos 41761 TGATTCATCCACTTGCAGTCGATTGTAGAGAATCTGGAGAGAAAA
41805
Gallus_gallus                    41761 .............................................
```

41805

Anas_platyrhynchos_platyrhynchos 41806 ATCTCTGGATACAGAACATATTCCTTCCTCCTGAAAGACACTCAG 41850
Gallus_gallus 41806 ............................................. 41850

Anas_platyrhynchos_platyrhynchos 41851 AGTTTCCAAAGGCAATTCCATAAATGACTTATCTGCTAGTAATAC 41895
Gallus_gallus 41851 ............................................. 41895

Anas_platyrhynchos_platyrhynchos 41896 CACACACAATCCTAACTCCATAGGTGTTCCTGAATAAATGTGACA 41940
Gallus_gallus 41896 ............................................. 41940

Anas_platyrhynchos_platyrhynchos 41941 AACCACAATTAGGATTTTATATTTCTCAGTATGTATTTAAATAAA 41985
Gallus_gallus 41941 ............................................. 41985

Anas_platyrhynchos_platyrhynchos 41986 TGAAGTTAGTTATAAAAACTAGGTTTCCTCTGTTGTCTGACAATT 42030
Gallus_gallus 41986 ............................................. 42030

Anas_platyrhynchos_platyrhynchos 42031 GAACTTCATTTTACTGTGAAAAGCATTATTCAAAAATATGTTTTA 42075
Gallus_gallus 42031 ............................................. 42075

Anas_platyrhynchos_platyrhynchos 42076 ATACAGTTAATTCCTAAGATGTTATTAGACTTCTAAGCATATACA 42120
Gallus_gallus 42076 ............................................. 42120

Anas_platyrhynchos_platyrhynchos 42121 TTTGAGAGTACTAATTAATCAGAGGGAAAAATATTCCTGTATTAA 42165
Gallus_gallus 42121 ............................................. 42165

Anas_platyrhynchos_platyrhynchos 42166 TAATCAAAACAATCCAGAACAAAATCTGGGCAAATGCTATGCATT 42210
Gallus_gallus 42166 ............................................. 42210

Anas_platyrhynchos_platyrhynchos 42211 TTAAGAAAATCATTTTGGAAATGAAAACTTGTACTAAAAATGTTT 42255
Gallus_gallus 42211 ............................................. 42255

Anas_platyrhynchos_platyrhynchos 42256 GGCTAAAGAGTACAGACTCGTAAAAGAGCACATTTCCAATAATAA 42300
Gallus_gallus 42256 ............................................. 42300

Anas_platyrhynchos_platyrhynchos 42301 CCGCGTATTTCTGGAGGAAACTTTTTGTTTCTGTTATTTGGGAAA 42345
Gallus_gallus 42301 ............................................. 42345

Anas_platyrhynchos_platyrhynchos 42346 AACAAAATGTATCATTTTATCAACATTTTCCTGTTTTGTTACAGA

```
42390
Gallus_gallus                    42346 .............................................
42390

Anas_platyrhynchos_platyrhynchos 42391 GAAGTAACAAATGCAAATGACAAGACAGCTGCTGGATTGCAGAAA
42435
Gallus_gallus                    42391 .............................................
42435

Anas_platyrhynchos_platyrhynchos 42436 GAAACTCCTGCAAATTTCAAAAAGCAAATGCATTTTTTTCTAGTA
42480
Gallus_gallus                    42436 .............................................
42480

Anas_platyrhynchos_platyrhynchos 42481 ACTTTTAGCTGAGAAATTTGATTGCTCTTCTCCATAAAGAAAAAA
42525
Gallus_gallus                    42481 .............................................
42525

Anas_platyrhynchos_platyrhynchos 42526 AAAAAATCAATCCCCAAAGCAAGCCATGGACTTGGAATGTCTTCT
42570
Gallus_gallus                    42526 .............................................
42570

Anas_platyrhynchos_platyrhynchos 42571 CTCAGAACATGCTTATACTCCCACATTAAGTTTGTTGCATTAAAG
42615
Gallus_gallus                    42571 .............................................
42615

Anas_platyrhynchos_platyrhynchos 42616 CAAACAAACTGTGAGAGTCCACAAAGTAAAAAGCAGAAAGCAGGA
42660
Gallus_gallus                    42616 .............................................
42660

Anas_platyrhynchos_platyrhynchos 42661 AAAGACAAAACTAGACCAACAGACTTATGTTCTCTTTTGGTATTT
42705
Gallus_gallus                    42661 .............................................
42705

Anas_platyrhynchos_platyrhynchos 42706 CTCCTCAAAAGACTGGCTTGTAATAAAGACATTTTATTAACCTGC
42750
Gallus_gallus                    42706 .............................................
42750

Anas_platyrhynchos_platyrhynchos 42751 CTGAAGCCTGTATACTATAGAAGACTTTTAATACATGCTGAAGGA
42795
Gallus_gallus                    42751 .............................................
42795

Anas_platyrhynchos_platyrhynchos 42796 TAGGCCCAGTGTACAAATAATCTTTGGAAGTATACTGAAATATTT
42840
Gallus_gallus                    42796 .............................................
42840

Anas_platyrhynchos_platyrhynchos 42841 AACTTGGTATAGAAGATAGGCAACACTGTTAAAGTGCACATATAT
42885
Gallus_gallus                    42841 .............................................
42885

Anas_platyrhynchos_platyrhynchos 42886 TATACTCCAGAGAGCACATACCCTGCATTCTAGTGTGTTTGGTGT
42930
Gallus_gallus                    42886 .............................................
42930
```

```
Anas_platyrhynchos_platyrhynchos 42931 CCTTATCTTTCACTTTTATTTTTCCTGATTCAGCTGCTGAAGCTG
42975
Gallus_gallus                    42931 .............................................
42975

Anas_platyrhynchos_platyrhynchos 42976 ATGTGCTGTGTTCTCTGGGAAGAAAAATTGCATTGAAAATATTTT
43020
Gallus_gallus                    42976 .............................................
43020

Anas_platyrhynchos_platyrhynchos 43021 TGGTATTAATACTCATGCAAGTAAAGGTTTTCATGGGATACACAG
43065
Gallus_gallus                    43021 .............................................
43065

Anas_platyrhynchos_platyrhynchos 43066 AGACAGAGATATTTCAATCTCAATGAAATTTTACACAAAGTACTC
43110
Gallus_gallus                    43066 .............................................
43110

Anas_platyrhynchos_platyrhynchos 43111 CTTTTGGAATTCCATCTTTAATTAGTTAAATTATAATGTTTCATT
43155
Gallus_gallus                    43111 .............................................
43155

Anas_platyrhynchos_platyrhynchos 43156 CTGAACTGTTGGAGGACAGTGCTTGTTAGGCATAGGCATTAGTCT
43200
Gallus_gallus                    43156 .............................................
43200

Anas_platyrhynchos_platyrhynchos 43201 TCCATGTGGTTCTATCCTCTTTGTCTTGCATATACTGCATATATT
43245
Gallus_gallus                    43201 .............................................
43245

Anas_platyrhynchos_platyrhynchos 43246 TATATATACCTTTAAAGATGCAGAAGTCCATTCAATGGAATCTAA
43290
Gallus_gallus                    43246 .............................................
43290

Anas_platyrhynchos_platyrhynchos 43291 TCTTGCTTTTCATTTACAACCTTTCCTGTATTTGTACGAGCTGAG
43335
Gallus_gallus                    43291 .............................................
43335

Anas_platyrhynchos_platyrhynchos 43336 ACCATTCTATTTTCACAAATGATAAAAACAACTCTAAATGTCTTA
43380
Gallus_gallus                    43336 .............................................
43380

Anas_platyrhynchos_platyrhynchos 43381 CCTAATGTTAGGGAAATCAAACTGTGATAGCAGTGGTTTCAGATA
43425
Gallus_gallus                    43381 .............................................
43425

Anas_platyrhynchos_platyrhynchos 43426 TTCTTCCATAATTTCTAATATTTCAGAGAAATCTGTTATCAAATT
43470
Gallus_gallus                    43426 .............................................
43470

Anas_platyrhynchos_platyrhynchos 43471 GTGTCCATTCTTGTCTATTTTCTCTGTTGGAGTCTATCACACACA
43515
```

Gallus_gallus                    43471 .............................................
43515

Anas_platyrhynchos_platyrhynchos 43516 TTGCAACAAAAAACAAAATACATACAAAATTCTATTTAACAAGGA
43560
Gallus_gallus                    43516 .............................................
43560

Anas_platyrhynchos_platyrhynchos 43561 TGGTGGGTGTAATTTGGTAAATCAGAGGCTAAATCAGATCATTTG
43605
Gallus_gallus                    43561 .............................................
43605

Anas_platyrhynchos_platyrhynchos 43606 AAACATTATAAAGTGGAAAATATAAATAGAGCAAATCAGTTTCAA
43650
Gallus_gallus                    43606 .............................................
43650

Anas_platyrhynchos_platyrhynchos 43651 AACTACTTCACACCATCATTGCCACCACCATTTTGCAAAATATTC
43695
Gallus_gallus                    43651 .............................................
43695

Anas_platyrhynchos_platyrhynchos 43696 TAACATAAGAATCAGTATAAAATGTCAATATTTATTATCAAAGCA
43740
Gallus_gallus                    43696 .............................................
43740

Anas_platyrhynchos_platyrhynchos 43741 AATAATATATTTCTGAAACTTAAATGTGGAACAGGAAGGGAAGGA
43785
Gallus_gallus                    43741 .............................................
43785

Anas_platyrhynchos_platyrhynchos 43786 AAGTCCAGGAAAAAAAAAAAAAAAAAAAAAGAATAAAAACTGGAA
43830
Gallus_gallus                    43786 .............................................
43830

Anas_platyrhynchos_platyrhynchos 43831 AGGAACTGTGAAAGGACAGCATTGAGAAGTAAATAGAAGAAGATG
43875
Gallus_gallus                    43831 .............................................
43875

Anas_platyrhynchos_platyrhynchos 43876 AGTGTTGAAAAAGCAAATTATATTGTATGGAGATCAGTGGAACCA
43920
Gallus_gallus                    43876 .............................................
43920

Anas_platyrhynchos_platyrhynchos 43921 ACTGAGGAAAACAAAATAAACAGAATGAAGAAAACAGATAAGGAG
43965
Gallus_gallus                    43921 .............................................
43965

Anas_platyrhynchos_platyrhynchos 43966 AGAATTGAGATGACACTATAAAACTGAATCTTCTAAAAGCCAATA
44010
Gallus_gallus                    43966 .............................................
44010

Anas_platyrhynchos_platyrhynchos 44011 TAGATGGACACAAAAAAATAAATGGACAAATGAGCTCAACTAAAT
44055
Gallus_gallus                    44011 .............................................
44055

```
Anas_platyrhynchos_platyrhynchos 44056 GTCAAAGAGTTTTAGTACTAATGGATGGACTTTCTATCCAAAGCC
44100
Gallus_gallus                    44056 .............................................
44100

Anas_platyrhynchos_platyrhynchos 44101 CTGTTTGCATGGCTTTGTGCTGGTTATATGCAAGATTAATGAAAG
44145
Gallus_gallus                    44101 .............................................
44145

Anas_platyrhynchos_platyrhynchos 44146 TTTTAATGTACATGTGTTTTAAAAATCTAAGCCACTTTTTTTTTT
44190
Gallus_gallus                    44146 .............................................
44190

Anas_platyrhynchos_platyrhynchos 44191 CTGTCTGTTCTACCTGACATTATCACCTCTGTACTGATGGAACTA
44235
Gallus_gallus                    44191 .............................................
44235

Anas_platyrhynchos_platyrhynchos 44236 CATCAATCTTTACATAGCTTGCCATTACAACAAAGGCTGGCCATG
44280
Gallus_gallus                    44236 .............................................
44280

Anas_platyrhynchos_platyrhynchos 44281 GACAAGAGAAAAAGTGTTCAGCACTTTAACCATCAGGAATAATCA
44325
Gallus_gallus                    44281 .............................................
44325

Anas_platyrhynchos_platyrhynchos 44326 TTATTTTAGCTGAAAGATGGTAACACAATGAATAAAACAGAATTA
44370
Gallus_gallus                    44326 .............................................
44370

Anas_platyrhynchos_platyrhynchos 44371 TATTTCAAGAACCAGATGTGTCAGAGTCATGTGATTAAAAGAAGA
44415
Gallus_gallus                    44371 .............................................
44415

Anas_platyrhynchos_platyrhynchos 44416 AAACAAATGCAGAAAATTCAGTATGATCTACACTTTTCATATCAG
44460
Gallus_gallus                    44416 .............................................
44460

Anas_platyrhynchos_platyrhynchos 44461 CTACATTTTTATCATAGTTCAAGTTTTTCATATCAATGATTAAAA
44505
Gallus_gallus                    44461 .............................................
44505

Anas_platyrhynchos_platyrhynchos 44506 AAGGTCTTTACCAGTTTTTGTTAATACATAACATTTAACCAGGGA
44550
Gallus_gallus                    44506 .............................................
44550

Anas_platyrhynchos_platyrhynchos 44551 GTATAGAACGTAGCGGAGACAAAATTAAAGTAAATGAGAATAGGA
44595
Gallus_gallus                    44551 .............................................
44595

Anas_platyrhynchos_platyrhynchos 44596 GTTTATAGACTTACAGAGTGCAGAAAACCTGTCGAAAAGCATAGG
44640
Gallus_gallus                    44596 .............................................
```

```
44640

Anas_platyrhynchos_platyrhynchos 44641 CTTTTCCAGATGTTACACTTGGAATAAATGGAAAATTCTATTATG
44685
Gallus_gallus                    44641 .............................................
44685

Anas_platyrhynchos_platyrhynchos 44686 CTATACAATAACTGTTCTCTGAAAAAGTATTTCCTTACAGAACTA
44730
Gallus_gallus                    44686 .............................................
44730

Anas_platyrhynchos_platyrhynchos 44731 AGAATTCTTACAATAATAGAACTAAACCCTCTTACAGTAACAGAA
44775
Gallus_gallus                    44731 .............................................
44775

Anas_platyrhynchos_platyrhynchos 44776 AACTTTTTTCATCTAGAATTTAATGTTCAGAGCCACAGTTGTGAA
44820
Gallus_gallus                    44776 .............................................
44820

Anas_platyrhynchos_platyrhynchos 44821 TTTGGAATTCCATTGAGATGTGTACTTAAAGAAAAATTTTGTGAC
44865
Gallus_gallus                    44821 .............................................
44865

Anas_platyrhynchos_platyrhynchos 44866 TAACATACTTGGATCAACTCCACCCAAACAGCCAAAGTGCCAAAG
44910
Gallus_gallus                    44866 .............................................
44910

Anas_platyrhynchos_platyrhynchos 44911 GACAGTGTGCCCTAGAAAGGAACTTTTCTGTAAAAATCAGTAATT
44955
Gallus_gallus                    44911 .............................................
44955

Anas_platyrhynchos_platyrhynchos 44956 TTCTGGAAATCAGTAATAATTCTCTGGTTAAGACTGATTTATTTT
45000
Gallus_gallus                    44956 .............................................
45000

Anas_platyrhynchos_platyrhynchos 45001 GAATAAAGGGTTGAGATCTATGATCCCATTTTGGGGGGGGGACAT
45045
Gallus_gallus                    45001 .............................................
45045

Anas_platyrhynchos_platyrhynchos 45046 CTGGGGCCAATGTACACAATGTAGATGTTAGACACAAAGCACCTT
45090
Gallus_gallus                    45046 .............................................
45090

Anas_platyrhynchos_platyrhynchos 45091 TTCAAGTAAATTGTTCTCAGTTGTATCAACTTTCTATCACAATAA
45135
Gallus_gallus                    45091 .............................................
45135

Anas_platyrhynchos_platyrhynchos 45136 ACTACAGAACATCTCAAAGTTGTCGAATGACATTGGAAACAGTAA
45180
Gallus_gallus                    45136 .............................................
45180

Anas_platyrhynchos_platyrhynchos 45181 CAGAACAATATAGTAACATTAAATGTTGACGCAGAATACACAAAA
```

```
45225
Gallus_gallus                     45181 .............................................
45225

Anas_platyrhynchos_platyrhynchos 45226 CATTTAGTAGTTGCGAAGATTAGGGATGTTACACTTGGAATAAAT
45270
Gallus_gallus                     45226 .............................................
45270

Anas_platyrhynchos_platyrhynchos 45271 GGAAAATTCTACATAAAGAGTGTTCATGATAGTTTTTTAGTACAT
45315
Gallus_gallus                     45271 .............................................
45315

Anas_platyrhynchos_platyrhynchos 45316 ATTTTTCAGTCTTGTTTGAATATAAGGTAATCAAATATATTACCG
45360
Gallus_gallus                     45316 .............................................
45360

Anas_platyrhynchos_platyrhynchos 45361 TATCAGTTTTAGTGATGCACTCTTGCAGGCTTTGAATGACAGTTT
45405
Gallus_gallus                     45361 .............................................
45405

Anas_platyrhynchos_platyrhynchos 45406 CTTTAAGGTGACTCTTCATATTTTGTTGCTTGTAAAGGTACACTT
45450
Gallus_gallus                     45406 .............................................
45450

Anas_platyrhynchos_platyrhynchos 45451 TTTCCAACAAAGTACAAAATGTATTGGGATTCACAGGAAATCGAG
45495
Gallus_gallus                     45451 .............................................
45495

Anas_platyrhynchos_platyrhynchos 45496 AAACTTTAGCTTGCACTAATAGTGAAGGAGAATAACATCATCATG
45540
Gallus_gallus                     45496 .............................................
45540

Anas_platyrhynchos_platyrhynchos 45541 AGACACATTCTTTTTGAGACATCATATTTGTACTTGAAGAAATTA
45585
Gallus_gallus                     45541 .............................................
45585

Anas_platyrhynchos_platyrhynchos 45586 CAAGAGAGACCATCAGTATCTAAAACAATATATTCTGTTGTTCTT
45630
Gallus_gallus                     45586 .............................................
45630

Anas_platyrhynchos_platyrhynchos 45631 TGAAAAACTGTTTACATATATGTTGCATAACAGATATGTACAATG
45675
Gallus_gallus                     45631 .............................................
45675

Anas_platyrhynchos_platyrhynchos 45676 AAAAAACAATCTCCTGACATGTATTTAACAGCAACTTTAGCCTTA
45720
Gallus_gallus                     45676 .............................................
45720

Anas_platyrhynchos_platyrhynchos 45721 AAATATACACTAAAATTCTGTAAATTCTGTCTGGATACACTGTAT
45765
Gallus_gallus                     45721 .............................................
45765
```

```
Anas_platyrhynchos_platyrhynchos 45766 CTGATAATGAAACATGCATTTAGGTATCTACTTTTCTCAGGTACT
45810
Gallus_gallus                    45766 .............................................
45810

Anas_platyrhynchos_platyrhynchos 45811 AATGAGTTACCACTATTTAAACAATTATTTAAAAATTTCATTGAA
45855
Gallus_gallus                    45811 .............................................
45855

Anas_platyrhynchos_platyrhynchos 45856 GGAAACAGTGAAACATATAAGCAATCAGGGTAATGATTTTTATCT
45900
Gallus_gallus                    45856 .............................................
45900

Anas_platyrhynchos_platyrhynchos 45901 CTAACTGTAAGAGGCACTTACCCATTGGCTGAGTGGATTTTCCCT
45945
Gallus_gallus                    45901 .............................................
45945

Anas_platyrhynchos_platyrhynchos 45946 TCTGCTGTGCAGTACTTACTTTTGGTTTCTCAATCAATGAGCTGT
45990
Gallus_gallus                    45946 .............................................
45990

Anas_platyrhynchos_platyrhynchos 45991 ACAAGTTTGATCTATAAAAAGCAGAAAACTGCTGAGTAGATTATT
46035
Gallus_gallus                    45991 .............................................
46035

Anas_platyrhynchos_platyrhynchos 46036 TAACTGTTCAGGCTTATTGATGTGAATGAGTTTAGTTGCATGAGT
46080
Gallus_gallus                    46036 .............................................
46080

Anas_platyrhynchos_platyrhynchos 46081 AATTACTACCCTACAATTTAATGTGCTAGTACTCTCATGATGCAT
46125
Gallus_gallus                    46081 .............................................
46125

Anas_platyrhynchos_platyrhynchos 46126 TAATTACCCCCCACCTTGGGGTGCAAAAACCTGCAAAATCCAATT
46170
Gallus_gallus                    46126 .............................................
46170

Anas_platyrhynchos_platyrhynchos 46171 GGAACAAACTGACAGCTTTAAGAAAACATCTTTCATTGCTATATA
46215
Gallus_gallus                    46171 .............................................
46215

Anas_platyrhynchos_platyrhynchos 46216 TCCGGTAAGGGAGATTTTTGTTTGTTCTATGACAGTTGTTTAAGA
46260
Gallus_gallus                    46216 .............................................
46260

Anas_platyrhynchos_platyrhynchos 46261 CTTTTCACAGTTGGTGAAACTGTACATAACCAGTTGCTACAGTTG
46305
Gallus_gallus                    46261 .............................................
46305

Anas_platyrhynchos_platyrhynchos 46306 TTTACGAGTTCTCACTCATGAGGCTGCAATAACCAGGCTATGCGA
46350
```

```
Gallus_gallus                      46306 .............................................
46350

Anas_platyrhynchos_platyrhynchos 46351 GGTTTTATAAAGCTAAATGGGACTCCTCCTGACATTTTCAGAAGC
46395
Gallus_gallus                      46351 .............................................
46395

Anas_platyrhynchos_platyrhynchos 46396 ATAACCAGTGAGAGATGAAAAATATAAACAGGATTTAGGAGGATA
46440
Gallus_gallus                      46396 .............................................
46440

Anas_platyrhynchos_platyrhynchos 46441 ACAAGAATGGTCTAAAACCTTTCAAATAAACTCTGGAGATTAAAC
46485
Gallus_gallus                      46441 .............................................
46485

Anas_platyrhynchos_platyrhynchos 46486 ATATAACTACAGAAAAAAAAAAAAAAAAGGCCCAGAAATAGTTAAA
46530
Gallus_gallus                      46486 .............................................
46530

Anas_platyrhynchos_platyrhynchos 46531 AGTACAGGTTTGAGCTGCAGGTGGTTTTATTTGTTTAATAATTTG
46575
Gallus_gallus                      46531 .............................................
46575

Anas_platyrhynchos_platyrhynchos 46576 CACACTAACCTAAAATTCAAATCCAGCAAAATTAGAAGTAAAAGA
46620
Gallus_gallus                      46576 .............................................
46620

Anas_platyrhynchos_platyrhynchos 46621 CAATAAGCAAACAAAATAGTGCTTATTCACAGGTATAGAAGGATG
46665
Gallus_gallus                      46621 .............................................
46665

Anas_platyrhynchos_platyrhynchos 46666 ACATCTGTAACAAACAGAATAACAAATGAAAATGATGACATTTTA
46710
Gallus_gallus                      46666 .............................................
46710

Anas_platyrhynchos_platyrhynchos 46711 TACCTGTCTTCTGAATGCTGGAGAATAATTATGTAGAAATTAGAT
46755
Gallus_gallus                      46711 .............................................
46755

Anas_platyrhynchos_platyrhynchos 46756 GGCAATTCATTCACTATATTAAAATGTTCTACAGGAATACAGAGA
46800
Gallus_gallus                      46756 .............................................
46800

Anas_platyrhynchos_platyrhynchos 46801 TTTCTCCATGCCTGCATAATAATTCAAAAATAAAATTATTTTAAT
46845
Gallus_gallus                      46801 .............................................
46845

Anas_platyrhynchos_platyrhynchos 46846 ACATTTTATTAACAACAATTTCAGTTCTTCCATGAGAAAATCGAC
46890
Gallus_gallus                      46846 .............................................
46890
```

```
Anas_platyrhynchos_platyrhynchos 46891 AATTCTAGTAAATAAGGTTTTAATAATTTAACAGGACTAGAGGTA
46935
Gallus_gallus                    46891 .............................................
46935

Anas_platyrhynchos_platyrhynchos 46936 ATGGACACATGCTGAAACATAGGAGGCTCCATCCAGACATAAGGA
46980
Gallus_gallus                    46936 .............................................
46980

Anas_platyrhynchos_platyrhynchos 46981 AACATTTACATGTATTCATTGTGGGGTGACAGAGCACTGGCACAG
47025
Gallus_gallus                    46981 .............................................
47025

Anas_platyrhynchos_platyrhynchos 47026 CTTATCCAGAGAGGCTGTGAAGTCTCCCTCCTTGGAGATGCTCAA
47070
Gallus_gallus                    47026 .............................................
47070

Anas_platyrhynchos_platyrhynchos 47071 AAGCCATCTGGACATGATCCTGGGTAACCTGCTCAGAGTAGCCCT
47115
Gallus_gallus                    47071 .............................................
47115

Anas_platyrhynchos_platyrhynchos 47116 GCTTGAGCAGGACAGATTGGATCAGATGACTTCCACGGGTCTCTT
47160
Gallus_gallus                    47116 .............................................
47160

Anas_platyrhynchos_platyrhynchos 47161 CCAACCTCAACCTTTCTGTGATTCTGTGATTTTTATGCAGTGCAG
47205
Gallus_gallus                    47161 .............................................
47205

Anas_platyrhynchos_platyrhynchos 47206 AACTTATAAATAATATTTAAAACAAATTTGAATTACTGTAACTTA
47250
Gallus_gallus                    47206 .............................................
47250

Anas_platyrhynchos_platyrhynchos 47251 GCCCACTTTTCATAGCAACATGACTTTTTTGGCCACTACACTATA
47295
Gallus_gallus                    47251 .............................................
47295

Anas_platyrhynchos_platyrhynchos 47296 TGTGAAGTTAAAATGTTATCAAATCCTAGCATAATATTATCAAAG
47340
Gallus_gallus                    47296 .............................................
47340

Anas_platyrhynchos_platyrhynchos 47341 GCAGGTAGCAGAATAATTCAATGTAACACACACTTGTAAAAGTGT
47385
Gallus_gallus                    47341 .............................................
47385

Anas_platyrhynchos_platyrhynchos 47386 TTCACGAATTTGTTAAAAACAGACACTTCACCAAAAGAAACTATA
47430
Gallus_gallus                    47386 .............................................
47430

Anas_platyrhynchos_platyrhynchos 47431 TAAAATTTTGCAATTATTGCTAACAAATATAGAACTGTTTTCAAG
47475
Gallus_gallus                    47431 .............................................
```

47475

Anas_platyrhynchos_platyrhynchos 47476 AGAACACGGAAGAAAAGTGATTCAACAGTCAGTCACCAGATGAAA 47520
Gallus_gallus 47476 ............................................. 47520

Anas_platyrhynchos_platyrhynchos 47521 TATTTGTTCTAATACAGAAAGCCCCATAAATGTCTTTGACTCTGT 47565
Gallus_gallus 47521 ............................................. 47565

Anas_platyrhynchos_platyrhynchos 47566 TACATTTAAGCATGTGCATGTATGTGTTCTTCAGATTGCAGCAAC 47610
Gallus_gallus 47566 ............................................. 47610

Anas_platyrhynchos_platyrhynchos 47611 CCCAAAGGTGTCTTCAGAACAAAATATTGTCCTATTCCTTGTGCT 47655
Gallus_gallus 47611 ............................................. 47655

Anas_platyrhynchos_platyrhynchos 47656 AAATAAGATAATACAGCACATACAGAGAAGACCCAAAATAATGGC 47700
Gallus_gallus 47656 ............................................. 47700

Anas_platyrhynchos_platyrhynchos 47701 CTACAAATACTTTTTTTTATTCTGTACTGTACTTCTGCTTGCCAC 47745
Gallus_gallus 47701 ............................................. 47745

Anas_platyrhynchos_platyrhynchos 47746 CTTTTTTAAATGTTTAATTCCTGCCAGTTACAGTTAAATCTTTGT 47790
Gallus_gallus 47746 ............................................. 47790

Anas_platyrhynchos_platyrhynchos 47791 ACCTAACTTTCTGGCAGTTTTAAATTTCATTTAATTTTTATTCTT 47835
Gallus_gallus 47791 ............................................. 47835

Anas_platyrhynchos_platyrhynchos 47836 TTCTAACTGAGGGTCTTCAAATTTTAGTTTTGGTATCCACTTTTA 47880
Gallus_gallus 47836 ............................................. 47880

Anas_platyrhynchos_platyrhynchos 47881 GATGTCAGATGTTTGAAGAACACATATTTCTCAGATGGTTTAGAA 47925
Gallus_gallus 47881 ............................................. 47925

Anas_platyrhynchos_platyrhynchos 47926 CAAATGACAGATTACTTTTGTCCCAGAAAAGTTTATCTGTCATCT 47970
Gallus_gallus 47926 ............................................. 47970

Anas_platyrhynchos_platyrhynchos 47971 GCTCCAGAACTATGAATTACATGAAATATTTTTTAACTGGTTTTT 48015
Gallus_gallus 47971 ............................................. 48015

Anas_platyrhynchos_platyrhynchos 48016 TTTTTTTTTTTTTAATATCATAGTTTAGATGATGTCTAGAGGCTA

48060
Gallus_gallus 48016 .............................................
48060

Anas_platyrhynchos_platyrhynchos 48061 ACCAACAGAGATTTTAATAAAAAGATCAATTATATATATACGTAT
48105
Gallus_gallus 48061 .............................................
48105

Anas_platyrhynchos_platyrhynchos 48106 ATATATGTAAATGAACCTTGTCATAGGGTAGCACTTGTGTGGGGG
48150
Gallus_gallus 48106 .............................................
48150

Anas_platyrhynchos_platyrhynchos 48151 CTTAAGTATCTCAAACAGCACGACTGGTTGCTGTCTAGATGGTGA
48195
Gallus_gallus 48151 .............................................
48195

Anas_platyrhynchos_platyrhynchos 48196 CTAGAGTCTCTGCTGCCAGGGTCCTTTACAACAGCATCGTCAGCT
48240
Gallus_gallus 48196 .............................................
48240

Anas_platyrhynchos_platyrhynchos 48241 GGATTGGATGAGACTCACATGGAACTAAGGATTTAAGTGTACTGC
48285
Gallus_gallus 48241 .............................................
48285

Anas_platyrhynchos_platyrhynchos 48286 ATTCAGAGAGCACAGCTCTGACAACCAGATACACCAAATTGTAAC
48330
Gallus_gallus 48286 .............................................
48330

Anas_platyrhynchos_platyrhynchos 48331 TACAGGACAAATTTGGATTTCATCTTTGTATCTCAATTCTAATTT
48375
Gallus_gallus 48331 .............................................
48375

Anas_platyrhynchos_platyrhynchos 48376 AAATTACATAGCTCATGCCACACAGATAACCTAGCATAAACAAAA
48420
Gallus_gallus 48376 .............................................
48420

Anas_platyrhynchos_platyrhynchos 48421 CCACATGACAATACGGACCATACTAACAAACAGTGAGTTTCAGTC
48465
Gallus_gallus 48421 .............................................
48465

Anas_platyrhynchos_platyrhynchos 48466 ATGACCAAGTCAACCCATACCTGAAATTAAGAAAATAAAAATGTT
48510
Gallus_gallus 48466 .............................................
48510

Anas_platyrhynchos_platyrhynchos 48511 CAACAGAGTTCAAAGGAAACAGATATGTATGTATATGTCTATATG
48555
Gallus_gallus 48511 .............................................
48555

Anas_platyrhynchos_platyrhynchos 48556 TGTGTTTGTATAACATACATATAAACACCTCATAAGTAACGCATA
48600
Gallus_gallus 48556 .............................................
48600

```
Anas_platyrhynchos_platyrhynchos 48601 TCAAAAAGTTACCATTGAAGTAGCAGTCAGTTGCTGTTCCACATT
48645
Gallus_gallus                    48601 .............................................
48645

Anas_platyrhynchos_platyrhynchos 48646 TTTCAGAAGATCACTTGTGTTCCTGTTTTGTTTCAATTGATTTTG
48690
Gallus_gallus                    48646 .............................................
48690

Anas_platyrhynchos_platyrhynchos 48691 AAGATGCAGTAATGCTGCCAGCTGAGAGCTGCTGGTGTCAGAATT
48735
Gallus_gallus                    48691 .............................................
48735

Anas_platyrhynchos_platyrhynchos 48736 AGCTGTTAGAAGGATATGTTTCATCATCATAGATGCTGAACAGCT
48780
Gallus_gallus                    48736 .............................................
48780

Anas_platyrhynchos_platyrhynchos 48781 CTGTAAAACGTTTAGAGGTGCTTAGTTTCTCTCCCCAAAATCATA
48825
Gallus_gallus                    48781 .............................................
48825

Anas_platyrhynchos_platyrhynchos 48826 TAAATACATATATATATATTATATGTTAATTATAACATGGATATA
48870
Gallus_gallus                    48826 .............................................
48870

Anas_platyrhynchos_platyrhynchos 48871 TATGTATGTTAAAGAAATAGTCTCTCACTTAAAAATCTGATTTAT
48915
Gallus_gallus                    48871 .............................................
48915

Anas_platyrhynchos_platyrhynchos 48916 TGCATTCCACTCACTGATGGAGCTATTTGGAAATCTAAGTGTTGA
48960
Gallus_gallus                    48916 .............................................
48960

Anas_platyrhynchos_platyrhynchos 48961 CACAGTGACATATGCTTAATGTCTGCAATAATTTCCTGTGGTTGC
49005
Gallus_gallus                    48961 .............................................
49005

Anas_platyrhynchos_platyrhynchos 49006 CTTGATGTGGGAGGAAGAAAATTAGTATAAGGTGCATTTCTCTAT
49050
Gallus_gallus                    49006 .............................................
49050

Anas_platyrhynchos_platyrhynchos 49051 TTCTGCAGTGGAGAAGACGGTGTAGGAAGTACGTATTATTTGTTC
49095
Gallus_gallus                    49051 .............................................
49095

Anas_platyrhynchos_platyrhynchos 49096 CTGATGTAGGGAATATGAGACTTACAGCCATTCCTTATCTTCGTT
49140
Gallus_gallus                    49096 .............................................
49140

Anas_platyrhynchos_platyrhynchos 49141 GTGGTGAAGGCCCATAGTCACAATAGTCACACTAGTGATTCATAT
49185
```

```
Gallus_gallus                    49141 .............................................
49185

Anas_platyrhynchos_platyrhynchos 49186 ACCCTTGACTCACCTACAGAATCATGCTAAGTAGGTGCTGATTTG
49230
Gallus_gallus                    49186 .............................................
49230

Anas_platyrhynchos_platyrhynchos 49231 ACAAATTGAACATAAACCTTTCCTATCTACATGAAAATAATGAAT
49275
Gallus_gallus                    49231 .............................................
49275

Anas_platyrhynchos_platyrhynchos 49276 GAATAAATGCACAGAAACATTTTTTAAATTATTTGTCAGGGAAGG
49320
Gallus_gallus                    49276 .............................................
49320

Anas_platyrhynchos_platyrhynchos 49321 ATTAAAATGTTTTGTGTTAATGTCTTATCTTACCTAATCACAATA
49365
Gallus_gallus                    49321 .............................................
49365

Anas_platyrhynchos_platyrhynchos 49366 TAATAAATTATTCATAATAAAATATTTGCATGTATGCAATCAACA
49410
Gallus_gallus                    49366 .............................................
49410

Anas_platyrhynchos_platyrhynchos 49411 ATGAAAACTTCATGTTAAGCACAACAGATACTGGAATTGTCAAAA
49455
Gallus_gallus                    49411 .............................................
49455

Anas_platyrhynchos_platyrhynchos 49456 AGCCATGGAACAAACATTTTGACCTCAGTTAACATACTGTATATT
49500
Gallus_gallus                    49456 .............................................
49500

Anas_platyrhynchos_platyrhynchos 49501 TAGTTACCCATATTACCAATTAGTGCCTATATGTATCTGTACGAT
49545
Gallus_gallus                    49501 .............................................
49545

Anas_platyrhynchos_platyrhynchos 49546 CAATTATCAGGCAGGATCTGTACGATGAATGATCAGGCAATGTCT
49590
Gallus_gallus                    49546 .............................................
49590

Anas_platyrhynchos_platyrhynchos 49591 ATAAACAAACATTCTTGCTTATGTAACTCAACACAACCTGGAAAA
49635
Gallus_gallus                    49591 .............................................
49635

Anas_platyrhynchos_platyrhynchos 49636 ATGAATAATACATTAGCTGCATTACCTGATGAAGGGCTAAAAACT
49680
Gallus_gallus                    49636 .............................................
49680

Anas_platyrhynchos_platyrhynchos 49681 GGCTAGTCGAAACTGGATCAAATTGTCCAAAGCATTCAACCATTT
49725
Gallus_gallus                    49681 .............................................
49725
```

Anas_platyrhynchos_platyrhynchos 49726 CCAAACTAGCAGGACCAAGTACATTGGTAATATTATAGCTGAATG
49770
Gallus_gallus 49726 .............................................
49770

Anas_platyrhynchos_platyrhynchos 49771 CAATACCCATACACTGTAGTAAAGCTTCACTGCACTGAGAAAGAT
49815
Gallus_gallus 49771 .............................................
49815

Anas_platyrhynchos_platyrhynchos 49816 ATTTCTGAGCCAACCTAAGTTCCTTCTAGATGATAAAAATAGAAA
49860
Gallus_gallus 49816 .............................................
49860

Anas_platyrhynchos_platyrhynchos 49861 GGTTTATATTAAAATATGCCTTAATATTAAATATTTATTAAATGT
49905
Gallus_gallus 49861 .............................................
49905

Anas_platyrhynchos_platyrhynchos 49906 ATACAGATTATTTTTGTGTTAGAAAAATAAATATTATTTTTCACG
49950
Gallus_gallus 49906 .............................................
49950

Anas_platyrhynchos_platyrhynchos 49951 GTCAATAATGTTTTTCTCCTTTTTTAGACATTTATTATGTTCAAA
49995
Gallus_gallus 49951 .............................................
49995

Anas_platyrhynchos_platyrhynchos 49996 ATATGCAAAATAGATAAATTATTCAGCAGATGCCACATTTGTTAT
50040
Gallus_gallus 49996 .............................................
50040

Anas_platyrhynchos_platyrhynchos 50041 GCAGTATTAAAACATAAATGTTTTAAGAAAATACCTTCCTTAATA
50085
Gallus_gallus 50041 .............................................
50085

Anas_platyrhynchos_platyrhynchos 50086 TAAAACATGTAGTACATTTTAGCAAAACTATACTTAAATTCCAGA
50130
Gallus_gallus 50086 .............................................
50130

Anas_platyrhynchos_platyrhynchos 50131 GAATAGGAGCTGCTCAGAACAAAGGAATTTTCAGTCAATTTACAA
50175
Gallus_gallus 50131 .............................................
50175

Anas_platyrhynchos_platyrhynchos 50176 AACAGTTGTTTCCATTTACAGTAAATTATATTTAATACAAATAAA
50220
Gallus_gallus 50176 .............................................
50220

Anas_platyrhynchos_platyrhynchos 50221 ACCACAATTTTCTTAATTTAATTATTACATCAAGTTATCAATTAA
50265
Gallus_gallus 50221 .............................................
50265

Anas_platyrhynchos_platyrhynchos 50266 CAATATAAACTGTATATCTCCTCCTTCTAGTGTGCTTTGAGTTGC
50310
Gallus_gallus 50266 .............................................

50310

Anas_platyrhynchos_platyrhynchos 50311 AGTAGTTAACACAGATCAGCTCAGCATACAGCTTAAATTTTGTAA
50355
Gallus_gallus 50311 .............................................
50355

Anas_platyrhynchos_platyrhynchos 50356 GAATTTTACAGATGTTTTATGACAGAAGTTATCTACTTGTTGCCA
50400
Gallus_gallus 50356 .............................................
50400

Anas_platyrhynchos_platyrhynchos 50401 ATCTGCAAAATGTTACATTCCTTTAAAAAGGCATGTCTGTGCAGC
50445
Gallus_gallus 50401 .............................................
50445

Anas_platyrhynchos_platyrhynchos 50446 AAAAAGCTGCTACACGCTGGATCAATGAAAATCTTTAAAGGGGAA
50490
Gallus_gallus 50446 .............................................
50490

Anas_platyrhynchos_platyrhynchos 50491 ATTACTTCTACCTGGGAGCTCAATCAGAATCATAGAATCATAGAA
50535
Gallus_gallus 50491 .............................................
50535

Anas_platyrhynchos_platyrhynchos 50536 TCTCCCAAGTTGGACGGGACTCATAAGGATCATCGAGTCCAACTC
50580
Gallus_gallus 50536 .............................................
50580

Anas_platyrhynchos_platyrhynchos 50581 CTGGCACCACACAGGTCTACCCAAAATTTTAGACCATGTGACTAA
50625
Gallus_gallus 50581 .............................................
50625

Anas_platyrhynchos_platyrhynchos 50626 GTGCACAGTCCAAATGCTTCTTAAACTCCAACAGACTTGGTGCAG
50670
Gallus_gallus 50626 .............................................
50670

Anas_platyrhynchos_platyrhynchos 50671 TGACTGCTTCCCTGGGGAGCCTGTTCCAGTGTGCAACCACCCTCT
50715
Gallus_gallus 50671 .............................................
50715

Anas_platyrhynchos_platyrhynchos 50716 CAGTGAAGAACCTCTTCCTGACGTCTAGCCTAAACCTCCCCTGCC
50760
Gallus_gallus 50716 .............................................
50760

Anas_platyrhynchos_platyrhynchos 50761 TTCAGATTAACACCATTCCCGTGGGTCCTATCACTGGTGACTAAG
50805
Gallus_gallus 50761 .............................................
50805

Anas_platyrhynchos_platyrhynchos 50806 GAGAATAGATTGACGCCTTGCTCTCCACTCCCTCTCGTGAGGAAG
50850
Gallus_gallus 50806 .............................................
50850

Anas_platyrhynchos_platyrhynchos 50851 TTGTAGACCGCAATGAGCTCTCCCCTCAGCCTCCTCTTTTCCAGG

```
50895
Gallus_gallus                      50851 .............................................
50895

Anas_platyrhynchos_platyrhynchos 50896 CTGAACAGGCCAAGTGACCTCAGCCGCTCCTCATACATCTTCGCC
50940
Gallus_gallus                      50896 .............................................
50940

Anas_platyrhynchos_platyrhynchos 50941 ATCTTTGTAGCCCACCTCTGGACACTCTCCAACAGTTTCATGTCT
50985
Gallus_gallus                      50941 .............................................
50985

Anas_platyrhynchos_platyrhynchos 50986 TTTTTGTACCATGGTGCCCAGAACTGCATCCAGTACTCAAGATGA
51030
Gallus_gallus                      50986 .............................................
51030

Anas_platyrhynchos_platyrhynchos 51031 GGCTCCTGCAGAAGCTCCCACAGCATGTATTGAGATGCAGGACAG
51075
Gallus_gallus                      51031 .............................................
51075

Anas_platyrhynchos_platyrhynchos 51076 AGACACAAGAGAGGTAAAAAGATGGGATTTTCTGGGCGTTTTGGA
51120
Gallus_gallus                      51076 .............................................
51120

Anas_platyrhynchos_platyrhynchos 51121 GATTCTCATTTGTAAGATCAAATTTATGTTACAAACATTTTCATT
51165
Gallus_gallus                      51121 .............................................
51165

Anas_platyrhynchos_platyrhynchos 51166 ACCATTTCAAACAGAACTGATAAGGTAAGAAAATTATTTTTTTAA
51210
Gallus_gallus                      51166 .............................................
51210

Anas_platyrhynchos_platyrhynchos 51211 ACTACTAAAACATGCCCAAATACCTTTAATTCCTGTACTTTCCTC
51255
Gallus_gallus                      51211 .............................................
51255

Anas_platyrhynchos_platyrhynchos 51256 CTATACTTATCAAACCGACACTTATTAGACAGTAAACTGATGTCT
51300
Gallus_gallus                      51256 .............................................
51300

Anas_platyrhynchos_platyrhynchos 51301 GTCACTTTCTGTTCATTGCATTCTCCTGAAATCAGGAAAGGTTTA
51345
Gallus_gallus                      51301 .............................................
51345

Anas_platyrhynchos_platyrhynchos 51346 GTGATGTCCAATTTAGCTTCTTCCTACTCATCACGGCAGGAAAAA
51390
Gallus_gallus                      51346 .............................................
51390

Anas_platyrhynchos_platyrhynchos 51391 AAAAAAAAAAGTTAAACATTACATCATTACATCAGGTACAGGTG
51435
Gallus_gallus                      51391 .............................................
51435
```

```
Anas_platyrhynchos_platyrhynchos 51436 GCTTCATTTCTTTTGTGAATGTTCAGGTGAAAATGAAACCCATGT
51480
Gallus_gallus                    51436 .............................................
51480

Anas_platyrhynchos_platyrhynchos 51481 TAAAAGATATCATATCACTACATGAATTCCTCGATGGTGATTTAG
51525
Gallus_gallus                    51481 .............................................
51525

Anas_platyrhynchos_platyrhynchos 51526 CAATACTAACAGACATCAAAATTTCTTAATATGTTATGATGACAA
51570
Gallus_gallus                    51526 .............................................
51570

Anas_platyrhynchos_platyrhynchos 51571 TACGTGTAATAGAAGCTAAAAAGGATGCCCATACAATCTCTTAGA
51615
Gallus_gallus                    51571 .............................................
51615

Anas_platyrhynchos_platyrhynchos 51616 TTTTTGATTTTATGGTCTTTCTTCCTCCGTTTATTTCTTATGAAC
51660
Gallus_gallus                    51616 .............................................
51660

Anas_platyrhynchos_platyrhynchos 51661 ACTTCAATCCATCACTAAAATTGATTGACTCAATCAACATTCATT
51705
Gallus_gallus                    51661 .............................................
51705

Anas_platyrhynchos_platyrhynchos 51706 TTCTTACATGGCGTTCTTACTGCAACACAAAAGGAAGTTATAAAC
51750
Gallus_gallus                    51706 .............................................
51750

Anas_platyrhynchos_platyrhynchos 51751 CTGAGCATTTCAAATACAAGTCCTTACTTTTCTTCCAAGGTTACA
51795
Gallus_gallus                    51751 .............................................
51795

Anas_platyrhynchos_platyrhynchos 51796 AAGACAAAGCTAACAAGCCCTTTAAGCCCTTCAAGAATTTTTTAA
51840
Gallus_gallus                    51796 .............................................
51840

Anas_platyrhynchos_platyrhynchos 51841 TCTCTCCTGAAAATTTTGGAATTATCATATATCAGGAGTAAGTAA
51885
Gallus_gallus                    51841 .............................................
51885

Anas_platyrhynchos_platyrhynchos 51886 AAAAGTAATCTACATAACAATGGCATGTTGACAGAAGCATTTTTT
51930
Gallus_gallus                    51886 .............................................
51930

Anas_platyrhynchos_platyrhynchos 51931 TCTGAAATTCATTGGTTCAGCAAACATTTTTCTTACGGCTTGTAT
51975
Gallus_gallus                    51931 .............................................
51975

Anas_platyrhynchos_platyrhynchos 51976 GCAATATGTAAAATCCAGAATAAAGAGTATTCCTATACAAGTGCA
52020
```

```
Gallus_gallus                      51976 .............................................
52020

Anas_platyrhynchos_platyrhynchos 52021 TTTAGAAAAAAATACTGTGGTAGACAAACTTTACACACAAAAACT
52065
Gallus_gallus                      52021 .............................................
52065

Anas_platyrhynchos_platyrhynchos 52066 AATTTCTATGTAGAGAAGAATGCATACCATAATGTTTGGCTTCCA
52110
Gallus_gallus                      52066 .............................................
52110

Anas_platyrhynchos_platyrhynchos 52111 GTAGATTTCTGAATGTACAGGATCTATGTTAATGGCAAGTAAATG
52155
Gallus_gallus                      52111 .............................................
52155

Anas_platyrhynchos_platyrhynchos 52156 AAGACATTTTCCAGTGAGGTACAGGCACTTGGATTTGATGTCAGG
52200
Gallus_gallus                      52156 .............................................
52200

Anas_platyrhynchos_platyrhynchos 52201 GCAGCCTTTACAAAGATTCTGTATGTTTGCTAACTGAGCCAGTGT
52245
Gallus_gallus                      52201 .............................................
52245

Anas_platyrhynchos_platyrhynchos 52246 AATATGACCTATAGTGTACTCCATATGCTTCCAATCCTGGTAAAA
52290
Gallus_gallus                      52246 .............................................
52290

Anas_platyrhynchos_platyrhynchos 52291 TATGTAGTCATGTTTATTTGCTCATATATACCCACACATACAGAA
52335
Gallus_gallus                      52291 .............................................
52335

Anas_platyrhynchos_platyrhynchos 52336 ACATACTTTTAAAATATATATATATATATACATGTGTGTACACAC
52380
Gallus_gallus                      52336 .............................................
52380

Anas_platyrhynchos_platyrhynchos 52381 AAGAACATTTGTTAAATCCAATTACTTTTAACAACAAAATAGCTG
52425
Gallus_gallus                      52381 .............................................
52425

Anas_platyrhynchos_platyrhynchos 52426 CATCTAACAGATACACAAATTCATTACCAATTGATAAAAAAAAAA
52470
Gallus_gallus                      52426 .............................................
52470

Anas_platyrhynchos_platyrhynchos 52471 AAAGTACACTAACGTATATAAGCAAATGACAAACAAATCTGAGAA
52515
Gallus_gallus                      52471 .............................................
52515

Anas_platyrhynchos_platyrhynchos 52516 AGAGTAAGGGGTAAAGGGGGTACCAGAGAGAGTGCTGGGGCTGCT
52560
Gallus_gallus                      52516 .............................................
52560
```

```
Anas_platyrhynchos_platyrhynchos 52561 ACGCAACAGTAGCATTTCCTACTGTTGCTGTCAGAGGTAGAATAA
52605
Gallus_gallus                    52561 .............................................
52605

Anas_platyrhynchos_platyrhynchos 52606 TGGGCAAGAAATACTGCTCTGTTCCAGCGGAGCACGTCTTGTTTT
52650
Gallus_gallus                    52606 .............................................
52650

Anas_platyrhynchos_platyrhynchos 52651 CTAAATCACCAGCTCAGTTAAGCAGTTCTGCTTAAAACCATTTAA
52695
Gallus_gallus                    52651 .............................................
52695

Anas_platyrhynchos_platyrhynchos 52696 AAAAGACAGTATTTTTAAAAAGCTCAATTATGAAGCCTCAAGTAA
52740
Gallus_gallus                    52696 .............................................
52740

Anas_platyrhynchos_platyrhynchos 52741 TTTCCCTGTAACAATTAGTTGTTTTTAAGTCTTTGCATTTAAGAA
52785
Gallus_gallus                    52741 .............................................
52785

Anas_platyrhynchos_platyrhynchos 52786 ACAGAATTTTTCTTAAAAATCTTAACAAATAACAACAACAACAAA
52830
Gallus_gallus                    52786 .............................................
52830

Anas_platyrhynchos_platyrhynchos 52831 TGCACCAAAAAAAAAAACACTAGAACTTTCTTTGCTGCAAAACTA
52875
Gallus_gallus                    52831 .............................................
52875

Anas_platyrhynchos_platyrhynchos 52876 AATGAATGGCCTGAGAAGCCCTTTCCTCAGTAGTGGAAGTTCTAC
52920
Gallus_gallus                    52876 .............................................
52920

Anas_platyrhynchos_platyrhynchos 52921 TACAGAAAACAGCCCCGGTAAAGTTTGTTATACCTTTATATGCAA
52965
Gallus_gallus                    52921 .............................................
52965

Anas_platyrhynchos_platyrhynchos 52966 ATCACAAGAATAATATGTTGTGTTACCTGTTCTGTTGCATTATCT
53010
Gallus_gallus                    52966 .............................................
53010

Anas_platyrhynchos_platyrhynchos 53011 TTAGTAGTTTCTCCACAGAATGATTCTGCAATTCTATGGAATGTT
53055
Gallus_gallus                    53011 .............................................
53055

Anas_platyrhynchos_platyrhynchos 53056 TTTCCCACTTTTTTAAGCTTTTTCTCGGTAGTAACAACTTGAAAA
53100
Gallus_gallus                    53056 .............................................
53100

Anas_platyrhynchos_platyrhynchos 53101 ATATCCAGAAGAATTTCCACAAGGTTTAGCTTCCTGTTACCTAGT
53145
Gallus_gallus                    53101 .............................................
```

```
53145

Anas_platyrhynchos_platyrhynchos 53146 TGCCTCATTAATGGTATGCTAGTATTTGTATTCTGAAATACAGAA
53190
Gallus_gallus                    53146 .............................................
53190

Anas_platyrhynchos_platyrhynchos 53191 GAAAGGAAAGTACAGTTTTACTTCCAAATATATTTGCAAAGTTAG
53235
Gallus_gallus                    53191 .............................................
53235

Anas_platyrhynchos_platyrhynchos 53236 CTGCTAAGAAGTTAGAATGTTGTATTTCTTTTAACTAAAGAAGCA
53280
Gallus_gallus                    53236 .............................................
53280

Anas_platyrhynchos_platyrhynchos 53281 AAAGTTTTAACATTATTGCCTCGATAGATCATAGATATGGCTATG
53325
Gallus_gallus                    53281 .............................................
53325

Anas_platyrhynchos_platyrhynchos 53326 TAACTTAAAATAATTTAAGAGTAATTTAATTTTCAAAGATAAATC
53370
Gallus_gallus                    53326 .............................................
53370

Anas_platyrhynchos_platyrhynchos 53371 TTAAATACAATACAAGACAATGAAAGTAGCTGCTTAGAACAAGAG
53415
Gallus_gallus                    53371 .............................................
53415

Anas_platyrhynchos_platyrhynchos 53416 TGGCTGGATTTCTGCTTGCCATGTTTTATGTACCCAGTACAAATA
53460
Gallus_gallus                    53416 .............................................
53460

Anas_platyrhynchos_platyrhynchos 53461 TCTAATTATAAAGGCATTAAAAATGTGTACATAAAAACAGAATTT
53505
Gallus_gallus                    53461 .............................................
53505

Anas_platyrhynchos_platyrhynchos 53506 TATAATTGCTCATTTTTGCAACAGTCTTTAGAAGCAACACAAGAA
53550
Gallus_gallus                    53506 .............................................
53550

Anas_platyrhynchos_platyrhynchos 53551 TACTTTAAAAAAGGCAAAGCAGAAAAAAAGCTAGATGCTAGAGTA
53595
Gallus_gallus                    53551 .............................................
53595

Anas_platyrhynchos_platyrhynchos 53596 TAATAACAGTACACACTAAGCAAAGTAGACTGCAGGCTGTTATTA
53640
Gallus_gallus                    53596 .............................................
53640

Anas_platyrhynchos_platyrhynchos 53641 CCCAAAATACAGTATTCTAAAGACAAGTTTTCTTTATTTACATAA
53685
Gallus_gallus                    53641 .............................................
53685

Anas_platyrhynchos_platyrhynchos 53686 CAGTCAATGTAAATGTTTTTCTCCTCAGGAATACATTTTATTGTG
```

```
53730
Gallus_gallus                     53686 .............................................
53730

Anas_platyrhynchos_platyrhynchos 53731 AGTAGTTTAAACCTGTCAGGTCTTTAGAGGATAATTTCATCAAAA
53775
Gallus_gallus                     53731 .............................................
53775

Anas_platyrhynchos_platyrhynchos 53776 TATCTCCAAGTTTATATTTTTTCCTTTTGTAGCTTTTAGATAAAT
53820
Gallus_gallus                     53776 .............................................
53820

Anas_platyrhynchos_platyrhynchos 53821 TTTAGAATTATAGACAATTGACAAAATAAATCTTCTCATTTCTGT
53865
Gallus_gallus                     53821 .............................................
53865

Anas_platyrhynchos_platyrhynchos 53866 CTTTCTTCTTCCTCTCTTAAACTGTAAATGTTTTGGGCATCAGAA
53910
Gallus_gallus                     53866 .............................................
53910

Anas_platyrhynchos_platyrhynchos 53911 TGTTCCCTTCCGTGTTCATACAATGCCTATCGTTACAGGAAGATA
53955
Gallus_gallus                     53911 .............................................
53955

Anas_platyrhynchos_platyrhynchos 53956 ATCTCATTTGGGGCTTCCACAAATCATCCAAAAGCAAACAAATAG
54000
Gallus_gallus                     53956 .............................................
54000

Anas_platyrhynchos_platyrhynchos 54001 TAGAACTAAATTAGATTTATTTTCTTAAGTAATAAACATATATAT
54045
Gallus_gallus                     54001 .............................................
54045

Anas_platyrhynchos_platyrhynchos 54046 ATATATCCATCCATCCATACCTCATTAACTGCAATTAAGCTGCAA
54090
Gallus_gallus                     54046 .............................................
54090

Anas_platyrhynchos_platyrhynchos 54091 ACATTATGAAACACTTCTTCTGCTTTGGATACTGCCATCTGAGCT
54135
Gallus_gallus                     54091 .............................................
54135

Anas_platyrhynchos_platyrhynchos 54136 AAGTTATAAGCATCTAGACAGTGACTATATTTTTCTTCTTCATTT
54180
Gallus_gallus                     54136 .............................................
54180

Anas_platyrhynchos_platyrhynchos 54181 CCTTCCTGCTTGGCAATCATCCTGAAAAAAACATAGAAACAAAAA
54225
Gallus_gallus                     54181 .............................................
54225

Anas_platyrhynchos_platyrhynchos 54226 AATAATGTAAAAAAGTATTTATTTTTACAGTATCAACAATATTGA
54270
Gallus_gallus                     54226 .............................................
54270
```

```
Anas_platyrhynchos_platyrhynchos 54271 ACATACATGATAGATAAAACACACTTTCGTGTGTTTGTTTTTTAT
54315
Gallus_gallus                    54271 .............................................
54315

Anas_platyrhynchos_platyrhynchos 54316 ATATTCAAATAGTAAAAGTAAATGTTCACACATTATTGTTTTTGA
54360
Gallus_gallus                    54316 .............................................
54360

Anas_platyrhynchos_platyrhynchos 54361 GGGGTTATTCCTGGAACTGTACTACTGCAATGCAAAGAGATTTTA
54405
Gallus_gallus                    54361 .............................................
54405

Anas_platyrhynchos_platyrhynchos 54406 ATAAATAAAATAACAGCAGATAATACTCCAGTGTATCTATTCACC
54450
Gallus_gallus                    54406 .............................................
54450

Anas_platyrhynchos_platyrhynchos 54451 AAGAACAAAATTCCAGAACAGCTAGTTATAATACGTATCTCAGTC
54495
Gallus_gallus                    54451 .............................................
54495

Anas_platyrhynchos_platyrhynchos 54496 ATTATAATATATACTGTATATTCTGGAAACAGCTTAGAGTGAAGG
54540
Gallus_gallus                    54496 .............................................
54540

Anas_platyrhynchos_platyrhynchos 54541 CATACTTCGTTTTGCACCAGAATTTCTAGTAAAACATTTCAACAG
54585
Gallus_gallus                    54541 .............................................
54585

Anas_platyrhynchos_platyrhynchos 54586 CAATAATTGTAAAAGCCTATAGATACTTGAGAATGATTGGCATGG
54630
Gallus_gallus                    54586 .............................................
54630

Anas_platyrhynchos_platyrhynchos 54631 TTGCCCAAATGCCTTTACAAATGGAAAGACAAAAAACTGGCTTAG
54675
Gallus_gallus                    54631 .............................................
54675

Anas_platyrhynchos_platyrhynchos 54676 AGACCTTGATACTTTGTTTTGCTCAAAGACAGTACAAAAAACAGT
54720
Gallus_gallus                    54676 .............................................
54720

Anas_platyrhynchos_platyrhynchos 54721 AGACTAGGTAAAAATAACTCTGCCTTCAGTGGAAAAGATTTTTGA
54765
Gallus_gallus                    54721 .............................................
54765

Anas_platyrhynchos_platyrhynchos 54766 TTCTGAAGTATTTTCTGAATATAGAACACATTCGGAATACAAATA
54810
Gallus_gallus                    54766 .............................................
54810

Anas_platyrhynchos_platyrhynchos 54811 TACACCTAATATCCAGATATAAAAGCTATAAATATTATCTGCTCT
54855
```

```
Gallus_gallus                    54811 .............................................
54855

Anas_platyrhynchos_platyrhynchos 54856 TTTCTATCTTAGACTACAATAATGTAAATAATACACTTTCTAGAT
54900
Gallus_gallus                    54856 .............................................
54900

Anas_platyrhynchos_platyrhynchos 54901 TAGTTAAACTTAAATACAATTTTTCAGCGTAGGTATAATCTTACC
54945
Gallus_gallus                    54901 .............................................
54945

Anas_platyrhynchos_platyrhynchos 54946 TGTGAATATTTGCACGTTCCAGCAAAATCTCAGCACTGCAGTTTT
54990
Gallus_gallus                    54946 .............................................
54990

Anas_platyrhynchos_platyrhynchos 54991 TATGCCCATAATTAAGGAAGTCTTCCTCAATCTGAATTAATTTAC
55035
Gallus_gallus                    54991 .............................................
55035

Anas_platyrhynchos_platyrhynchos 55036 TGTAAGATCCCTGTAGCATATGTGGCGACTGACAAGGCTCAGTAT
55080
Gallus_gallus                    55036 .............................................
55080

Anas_platyrhynchos_platyrhynchos 55081 TAACAGTCAAACAATTTTGAGCAAAATGTATCTCAATAAGTGCTT
55125
Gallus_gallus                    55081 .............................................
55125

Anas_platyrhynchos_platyrhynchos 55126 TCCTAGGGATAAAAATTATGTACAGGATGGCAGTCAGAAAGCACA
55170
Gallus_gallus                    55126 .............................................
55170

Anas_platyrhynchos_platyrhynchos 55171 GGCTGTATGTATTTCTCTCTGTCTTCCTTTAAATTATAATAAGCA
55215
Gallus_gallus                    55171 .............................................
55215

Anas_platyrhynchos_platyrhynchos 55216 TATAAGAAGCCTCTGAATTCTTTAAATTAATATTCTACAAATGAT
55260
Gallus_gallus                    55216 .............................................
55260

Anas_platyrhynchos_platyrhynchos 55261 AGTATGTCCTGAAAAATATCATAATTTCAGATTAGAACAGAGGTG
55305
Gallus_gallus                    55261 .............................................
55305

Anas_platyrhynchos_platyrhynchos 55306 TCAGTTGTTCCATGTATCTAATTTTTCAAACTACCTAGGCCTAGT
55350
Gallus_gallus                    55306 .............................................
55350

Anas_platyrhynchos_platyrhynchos 55351 CAGTTTTCTCAGCATAGCTGAAATGAGGAAACTATTAGTCTTCCC
55395
Gallus_gallus                    55351 .............................................
55395
```

```
Anas_platyrhynchos_platyrhynchos 55396 TTTCTAATCCAGAAGTTTAGTATTTCTGGAAAGAAAAGGAAAAAG
55440
Gallus_gallus                    55396 .............................................
55440

Anas_platyrhynchos_platyrhynchos 55441 TTACGTCTTTCTCTCTTTTTACCTACACATTTTAAGCAATCTGTA
55485
Gallus_gallus                    55441 .............................................
55485

Anas_platyrhynchos_platyrhynchos 55486 CTTCGATTTCAATATCACAAAAGTCATTGTTTAGTTTCACCATAA
55530
Gallus_gallus                    55486 .............................................
55530

Anas_platyrhynchos_platyrhynchos 55531 AAAGACCTCTTGGATAATGGTGAAGAAAAAAGTAGGTCAATACTG
55575
Gallus_gallus                    55531 .............................................
55575

Anas_platyrhynchos_platyrhynchos 55576 TGGTTTATGAGAGTTGTGGAAATATATGAATGTTGTGGAAATATC
55620
Gallus_gallus                    55576 .............................................
55620

Anas_platyrhynchos_platyrhynchos 55621 TGCAACAGAGACCTCTGCACAGGCAAATGTATCTCAGTCCTTTGG
55665
Gallus_gallus                    55621 .............................................
55665

Anas_platyrhynchos_platyrhynchos 55666 GTAGCCTACACCTCAAACTATGCCTAATAACATTGTGGTATCTGA
55710
Gallus_gallus                    55666 .............................................
55710

Anas_platyrhynchos_platyrhynchos 55711 ACTAGCTTACTAACTTACCAGGTAATAGGACTCTGTACAAGAATA
55755
Gallus_gallus                    55711 .............................................
55755

Anas_platyrhynchos_platyrhynchos 55756 AAGAAATTGGAAAGGAAATTTAAACAAACAAACAAACAAACAGAA
55800
Gallus_gallus                    55756 .............................................
55800

Anas_platyrhynchos_platyrhynchos 55801 TAAGCAGAATTACTTGGGAGAAAAATATTTGAGAATCTTCCGAAT
55845
Gallus_gallus                    55801 .............................................
55845

Anas_platyrhynchos_platyrhynchos 55846 GCACTATACTAAAATGAAGGTGAGTGACAAAACTAGTATTTACCT
55890
Gallus_gallus                    55846 .............................................
55890

Anas_platyrhynchos_platyrhynchos 55891 GCAGCTGGAGGCATATAGTGGAGCATAGTGGAGGCTGGATATATG
55935
Gallus_gallus                    55891 .............................................
55935

Anas_platyrhynchos_platyrhynchos 55936 TTAATAAAAAGACAATTACCTGGCTTCAAGGGATGCAATTATAAA
55980
Gallus_gallus                    55936 .............................................
```

```
55980

Anas_platyrhynchos_platyrhynchos 55981 ACCCAACTCAGATTCTCTGTTAGGTCTCTTCAGTAAGGTAGATCT
56025
Gallus_gallus                    55981 .............................................
56025

Anas_platyrhynchos_platyrhynchos 56026 AAGTACATATACAGTGTGTTCCAGTATCTTACAAGCCTGCAGGTA
56070
Gallus_gallus                    56026 .............................................
56070

Anas_platyrhynchos_platyrhynchos 56071 ACATCAGAATTTCACTTTAAATGTTTGCTGTTAATATTAAACATT
56115
Gallus_gallus                    56071 .............................................
56115

Anas_platyrhynchos_platyrhynchos 56116 AATTTAGAAAGCTTATATTTTGTACAGAAAATGTTAAATATCTTT
56160
Gallus_gallus                    56116 .............................................
56160

Anas_platyrhynchos_platyrhynchos 56161 CCCTAATATTCACAGCGACAGCTGATTTTTGGCAACTGTAGGCAA
56205
Gallus_gallus                    56161 .............................................
56205

Anas_platyrhynchos_platyrhynchos 56206 TTCTAGAAAGTGTAAGTTGTAATTGTATTTAGGTATTTCGCTATG
56250
Gallus_gallus                    56206 .............................................
56250

Anas_platyrhynchos_platyrhynchos 56251 GTTAAAAGATTGAAGCACATACACACAAGGAAAGGATAAAAGAGT
56295
Gallus_gallus                    56251 .............................................
56295

Anas_platyrhynchos_platyrhynchos 56296 TTGGCCTGAAGACAAAGTTAAGAGGTGCGTACTACTGCTGCCCTC
56340
Gallus_gallus                    56296 .............................................
56340

Anas_platyrhynchos_platyrhynchos 56341 TTTTCTCTAAGCACACCATGCACAGAAGATGGAGCCAGACTCATC
56385
Gallus_gallus                    56341 .............................................
56385

Anas_platyrhynchos_platyrhynchos 56386 TTGAAGTCCCTAGCTGGTAAGTGAGAATGCCCCTTTGATGACGCT
56430
Gallus_gallus                    56386 .............................................
56430

Anas_platyrhynchos_platyrhynchos 56431 CAAAACCTCATATATAGATTTGTCCTTATTAAAGGTTTTTGTTAC
56475
Gallus_gallus                    56431 .............................................
56475

Anas_platyrhynchos_platyrhynchos 56476 TACAGTCATGTTGGTTAGCCATGTAGACAAAAACAAAACAAAACA
56520
Gallus_gallus                    56476 .............................................
56520

Anas_platyrhynchos_platyrhynchos 56521 AAAAAAACCAACGCATACTACATTAACAGCCATACTGGTAAACCC
```

```
56565
Gallus_gallus                    56521 .............................................
56565

Anas_platyrhynchos_platyrhynchos 56566 ACAGTAAATGCCTGCCCTCACTGGTAACTAGGTATTTTGTCTTTC
56610
Gallus_gallus                    56566 .............................................
56610

Anas_platyrhynchos_platyrhynchos 56611 AACGATTGTAGATGTGAATTCACCCAAAGCCAAAAAAATCTCTCT
56655
Gallus_gallus                    56611 .............................................
56655

Anas_platyrhynchos_platyrhynchos 56656 GTACAGGAAAGACTTGCATGTAATATCAATAAAGGACAGAGCATC
56700
Gallus_gallus                    56656 .............................................
56700

Anas_platyrhynchos_platyrhynchos 56701 ACAGACACAACAGAACTATATCTGTGCTGAAGCTCATGGAAGTTA
56745
Gallus_gallus                    56701 .............................................
56745

Anas_platyrhynchos_platyrhynchos 56746 AGGGCTGAACAGTAAATTTTAATGTTAAGGGTAAAATAAGAAAAT
56790
Gallus_gallus                    56746 .............................................
56790

Anas_platyrhynchos_platyrhynchos 56791 GTTTTTAATAAATATATGTATTACATTAGTATCTTTCAGGCTCTA
56835
Gallus_gallus                    56791 .............................................
56835

Anas_platyrhynchos_platyrhynchos 56836 AAAATCAGTATTTGAAATGTAACTACATACTGGGAAACAATTCTA
56880
Gallus_gallus                    56836 .............................................
56880

Anas_platyrhynchos_platyrhynchos 56881 ACTTCTAACTAAGATTCTAAAGTTAGAATCTTTATTAAATATGGA
56925
Gallus_gallus                    56881 .............................................
56925

Anas_platyrhynchos_platyrhynchos 56926 AAACAAACACATGTCAACTGTTTACAGTTTACAAAGGATCTTATC
56970
Gallus_gallus                    56926 .............................................
56970

Anas_platyrhynchos_platyrhynchos 56971 CAAGAGAATTGCATCCCTGACATTTAGTTACACATACTAGAGTAC
57015
Gallus_gallus                    56971 .............................................
57015

Anas_platyrhynchos_platyrhynchos 57016 TGTTGTTCTTATTTCAATAGAAGCATATACAACCTAACACTAAAA
57060
Gallus_gallus                    57016 .............................................
57060

Anas_platyrhynchos_platyrhynchos 57061 AAAATTATACTTCTGTGGGGACAGTCTCATTCCTACTTGCATGCA
57105
Gallus_gallus                    57061 .............................................
57105
```

```
Anas_platyrhynchos_platyrhynchos 57106 TCTGTTTGGAATTCATAATTAATGTTTGTTGAACGCACAGCCAAG
57150
Gallus_gallus                    57106 .............................................
57150

Anas_platyrhynchos_platyrhynchos 57151 CAAACAACATTAACTGCATTTAACACCTAATTTTGTACTGAAATG
57195
Gallus_gallus                    57151 .............................................
57195

Anas_platyrhynchos_platyrhynchos 57196 CCAAGGGCATAAAATTTCTCTTCATTGAAAGTGAGGTACAGCTGC
57240
Gallus_gallus                    57196 .............................................
57240

Anas_platyrhynchos_platyrhynchos 57241 AGAAAAGTAAACAATAGTTCCATACCGATCTCCGTTTTTCTTCCC
57285
Gallus_gallus                    57241 .............................................
57285

Anas_platyrhynchos_platyrhynchos 57286 CCTCTTCCTCTAGAATTGCCTCTATTAGACTAACAGTGCTGTTGT
57330
Gallus_gallus                    57286 .............................................
57330

Anas_platyrhynchos_platyrhynchos 57331 ACCAAAACTGTTCATTTCCTCCTAGAAGCTGCGCTTTCTCAAGAA
57375
Gallus_gallus                    57331 .............................................
57375

Anas_platyrhynchos_platyrhynchos 57376 GTGCTTTAGCTTGTCTGTGTTTTTTTTCCAGGTTAGCCAAAACAG
57420
Gallus_gallus                    57376 .............................................
57420

Anas_platyrhynchos_platyrhynchos 57421 CCAATAAATACAAGCACCTTGACGCATCATATGGAGTCCTCAATT
57465
Gallus_gallus                    57421 .............................................
57465

Anas_platyrhynchos_platyrhynchos 57466 CCTAGGAAAAAAGATACATTACCTGTATATGAAAGAACCACTACC
57510
Gallus_gallus                    57466 .............................................
57510

Anas_platyrhynchos_platyrhynchos 57511 ACTATTCATATTTATTTATATTCAGATTTTAGATAAAACTTATGA
57555
Gallus_gallus                    57511 .............................................
57555

Anas_platyrhynchos_platyrhynchos 57556 ATAGGGTCAGAAATAAGAATTGAATAGTATTTCTTCATTTACTTG
57600
Gallus_gallus                    57556 .............................................
57600

Anas_platyrhynchos_platyrhynchos 57601 TAGAATTTGTTAGTTACTTTTTATATTACTTGTCCTTTTTCAAAA
57645
Gallus_gallus                    57601 .............................................
57645

Anas_platyrhynchos_platyrhynchos 57646 GCACAAAGTGTCTACTATGAACCAAAAACTTCAAATCTTCAAATA
57690
```

```
Gallus_gallus                    57646 .............................................
57690

Anas_platyrhynchos_platyrhynchos 57691 TAGTTTGTATTTCTTTGTTTCCACACCTTCTACCCCACAGAAGTC
57735
Gallus_gallus                    57691 .............................................
57735

Anas_platyrhynchos_platyrhynchos 57736 AAATAAGCAATTTTTCTTCTATTATTTGATACCATAGAATCATCT
57780
Gallus_gallus                    57736 .............................................
57780

Anas_platyrhynchos_platyrhynchos 57781 AGGCTGGAAAAGGCCTTCAAGATCATCAAGTTCAACCATCAACCT
57825
Gallus_gallus                    57781 .............................................
57825

Anas_platyrhynchos_platyrhynchos 57826 TACCTACCAAGTCCCACCACTTAAACATGTCACCTAGTGCCATAT
57870
Gallus_gallus                    57826 .............................................
57870

Anas_platyrhynchos_platyrhynchos 57871 ATGCATACACCTACTGTCATTTGTCTGTATATGATGGAGATGGAG
57915
Gallus_gallus                    57871 .............................................
57915

Anas_platyrhynchos_platyrhynchos 57916 GTATTAAATTTTTTTTTAGTGAACTGGGATCTAGATACTGGAGC
57960
Gallus_gallus                    57916 ............................................
57960

Anas_platyrhynchos_platyrhynchos 57961 TCCAGTAAATTTACAACACTATTTTATGCTACTTAAAAAGTATAT
58005
Gallus_gallus                    57961 .............................................
58005

Anas_platyrhynchos_platyrhynchos 58006 ACTAGTAGTATACATGATCCTAACATACAGCATACTAAAGATAGT
58050
Gallus_gallus                    58006 .............................................
58050

Anas_platyrhynchos_platyrhynchos 58051 ATACTGGCATGCAGACATATAGCATAAAAAACGCTATAGAAAAAA
58095
Gallus_gallus                    58051 .............................................
58095

Anas_platyrhynchos_platyrhynchos 58096 AATGTATCATTAAAATACGTTGTCCTCCCTGATGTCATCTGAACT
58140
Gallus_gallus                    58096 .............................................
58140

Anas_platyrhynchos_platyrhynchos 58141 TTTCTCAATGTACAATACTAAATTGATGCTTCAATAAAAGAAATG
58185
Gallus_gallus                    58141 .............................................
58185

Anas_platyrhynchos_platyrhynchos 58186 TATTATTTGTTAATATATGATTCCATATATTAAAATCATATAAAT
58230
Gallus_gallus                    58186 .............................................
58230
```

```
Anas_platyrhynchos_platyrhynchos 58231 TATTGAAATCACATTATATATTTAAAATATTAATCTTTTAAATAT
58275
Gallus_gallus                    58231 .............................................
58275

Anas_platyrhynchos_platyrhynchos 58276 AAAAATATTAATAATATTATATAATACAATGTGTATAGGATATAT
58320
Gallus_gallus                    58276 .............................................
58320

Anas_platyrhynchos_platyrhynchos 58321 CATATATAATATGTATAATGTATATAATATGTAATTAGTCATAAG
58365
Gallus_gallus                    58321 .............................................
58365

Anas_platyrhynchos_platyrhynchos 58366 TATTAAATTGAAATATTAAAATATTAAAGTTATATTTACATTATT
58410
Gallus_gallus                    58366 .............................................
58410

Anas_platyrhynchos_platyrhynchos 58411 AAAATTTTAGCTTTCTTCATGCTTAACGTACTTCAACAGTTTTTA
58455
Gallus_gallus                    58411 .............................................
58455

Anas_platyrhynchos_platyrhynchos 58456 CTTACATCAGCAAAATTAGTTAGTGGAATAACAGTATATTTATAT
58500
Gallus_gallus                    58456 .............................................
58500

Anas_platyrhynchos_platyrhynchos 58501 AATCTTATTTTTTTTTAATTAAAGCAATTACAATAATTATGCAAC
58545
Gallus_gallus                    58501 .............................................
58545

Anas_platyrhynchos_platyrhynchos 58546 AGTAACCTGCATAAGCAGGAAGAGAGCTGTAGTACAGAATACAAG
58590
Gallus_gallus                    58546 .............................................
58590

Anas_platyrhynchos_platyrhynchos 58591 TTGTAGGAAGAATATTTTTAATATAAAAACTTCACATTAAAGAGA
58635
Gallus_gallus                    58591 .............................................
58635

Anas_platyrhynchos_platyrhynchos 58636 TTAAATGGCTTAGTTTGGTGGAACACAGGGACACTTTAAATCAGA
58680
Gallus_gallus                    58636 .............................................
58680

Anas_platyrhynchos_platyrhynchos 58681 TCTGCCAGAGTTCTAAGTCATAATCCCTGCATAACTGGCTATCAC
58725
Gallus_gallus                    58681 .............................................
58725

Anas_platyrhynchos_platyrhynchos 58726 TAAACATAATAACATTTAATAAAACATGTTATAAAACGTGTTTAT
58770
Gallus_gallus                    58726 .............................................
58770

Anas_platyrhynchos_platyrhynchos 58771 AAAAATGTAGCACCAAAATCACAATAAAATACTAAGTGATATAAT
58815
Gallus_gallus                    58771 .............................................
```

```
58815

Anas_platyrhynchos_platyrhynchos 58816 TAGATTAAATGGTGAGGGGATGGAGATAAATGTCATATCTCTCTT
58860
Gallus_gallus                    58816 .............................................
58860

Anas_platyrhynchos_platyrhynchos 58861 ATTAACAAATCTTGAACATACATAAACTATCTACACAAAGAAAGT
58905
Gallus_gallus                    58861 .............................................
58905

Anas_platyrhynchos_platyrhynchos 58906 TTGAACATAACTAGATGTTCCAAGGGTCACTGCAAAGCAGAGAGA
58950
Gallus_gallus                    58906 .............................................
58950

Anas_platyrhynchos_platyrhynchos 58951 GATGCTAGCACTATTTCACAAGTTGAAATAATCTTCAATAATCAG
58995
Gallus_gallus                    58951 .............................................
58995

Anas_platyrhynchos_platyrhynchos 58996 AAAGTTGAACTTAAGTTTCATATATTTAAATGCATTTATATTTTC
59040
Gallus_gallus                    58996 .............................................
59040

Anas_platyrhynchos_platyrhynchos 59041 AACTTCAATTGGTTTTTGCTTGGTCAGTTTTCTTTATTAGCACTA
59085
Gallus_gallus                    59041 .............................................
59085

Anas_platyrhynchos_platyrhynchos 59086 TGAATGTTCTGGTTAGTTTCCCTTACTTTAATGAATAACAATGCA
59130
Gallus_gallus                    59086 .............................................
59130

Anas_platyrhynchos_platyrhynchos 59131 TTCTCAGTTTAACAGTACTTTTATTTCTGCTTTAAAAAATATATC
59175
Gallus_gallus                    59131 .............................................
59175

Anas_platyrhynchos_platyrhynchos 59176 CCAGCAGTTACTTGTTTTTCTATTTGATTATTATTATGAGAAATG
59220
Gallus_gallus                    59176 .............................................
59220

Anas_platyrhynchos_platyrhynchos 59221 ACTATATCAAAATTAACATTTTTCCTATATGTTATATTTTTTTTT
59265
Gallus_gallus                    59221 .............................................
59265

Anas_platyrhynchos_platyrhynchos 59266 TTTTCTGTTTTATTTCTGATGTACCTGAGTTGCAGCATGGGCCTC
59310
Gallus_gallus                    59266 .............................................
59310

Anas_platyrhynchos_platyrhynchos 59311 TGAAAGCAGTAGTCTTGCCGGCTGATACAAACCTAGCTGAATTAG
59355
Gallus_gallus                    59311 .............................................
59355

Anas_platyrhynchos_platyrhynchos 59356 CACTTCAGCTTTTTCTATCCATAAGTAGGGGAAAGAAAGTGCACT
```

```
59400
Gallus_gallus                    59356 .............................................
59400

Anas_platyrhynchos_platyrhynchos 59401 CAGTCCTTTGCCTGTTATTGCATTTAACTCAAAAATCTGAAAAAT
59445
Gallus_gallus                    59401 .............................................
59445

Anas_platyrhynchos_platyrhynchos 59446 AATAAAAATATTAGACTGTTATAAATTTTAAAACTAAGCTATATGA
59490
Gallus_gallus                    59446 .............................................
59490

Anas_platyrhynchos_platyrhynchos 59491 AATACAAACTGTTATTAACTATTATATCCAGTCCCTGATATATCT
59535
Gallus_gallus                    59491 .............................................
59535

Anas_platyrhynchos_platyrhynchos 59536 CACAGTCCAGCTCTATTTTGTATTTTCTTAAAATTTTGAAGTTGT
59580
Gallus_gallus                    59536 .............................................
59580

Anas_platyrhynchos_platyrhynchos 59581 TTTATAAGACAAACAGGTTTACTGTGAGTTTTGGACCTTTGTAAT
59625
Gallus_gallus                    59581 .............................................
59625

Anas_platyrhynchos_platyrhynchos 59626 ATGCAAAAGGTAGTTAACTGATAAATTTATACTATTACAATTAAT
59670
Gallus_gallus                    59626 .............................................
59670

Anas_platyrhynchos_platyrhynchos 59671 TTTAGGACTGATTAACTACTTCAGATCTGCTACTGTAAGGAAGTA
59715
Gallus_gallus                    59671 .............................................
59715

Anas_platyrhynchos_platyrhynchos 59716 ATTGACTCTTACTAGTAACTTTAGTAACTAGCAACTTGCTTTTGC
59760
Gallus_gallus                    59716 .............................................
59760

Anas_platyrhynchos_platyrhynchos 59761 TCCTCAGCCATTTTCATTGATTTGTTTAGTCCCAAATTCCATTTG
59805
Gallus_gallus                    59761 .............................................
59805

Anas_platyrhynchos_platyrhynchos 59806 TAGAAAAGTGGCTTTTCTTTGTATGAATACTGATTCTATTCTTGT
59850
Gallus_gallus                    59806 .............................................
59850

Anas_platyrhynchos_platyrhynchos 59851 TCCACAATATAGATAATATGCCAGGCAAACCCAGCTCACATTCAG
59895
Gallus_gallus                    59851 .............................................
59895

Anas_platyrhynchos_platyrhynchos 59896 CTGATGACTGCCAGCAGAATTTAAGTAGTATGCACTTTTAATTAA
59940
Gallus_gallus                    59896 .............................................
59940
```

```
Anas_platyrhynchos_platyrhynchos 59941 CTTTTTGTGTGTGTTGAATTACTTCTCAGGTATAAAATTATTTTA
59985
Gallus_gallus                    59941 .............................................
59985

Anas_platyrhynchos_platyrhynchos 59986 AAACAATGTAGGAAGTACATTAAAATCAAAAGGAAAGAGTAATTT
60030
Gallus_gallus                    59986 .............................................
60030

Anas_platyrhynchos_platyrhynchos 60031 ATTAAGGAAATGGGAGTAGAAATAAAAACCTTTTTCCCTGAAGTG
60075
Gallus_gallus                    60031 .............................................
60075

Anas_platyrhynchos_platyrhynchos 60076 AGGATATTTTCATCCTTACTACCATTCAGCTGAGCACTCTCCAAG
60120
Gallus_gallus                    60076 .............................................
60120

Anas_platyrhynchos_platyrhynchos 60121 AATGTTTTTCTTTCTTCTGTCTCAGTATGTTTTTTTTTTTCCTTT
60165
Gallus_gallus                    60121 .............................................
60165

Anas_platyrhynchos_platyrhynchos 60166 TTCAGTGCAATCTCTTGCCTACACCTAAATTGTAACAAAGAAAAA
60210
Gallus_gallus                    60166 .............................................
60210

Anas_platyrhynchos_platyrhynchos 60211 ATTAATTTGCGTATGTTTTAGAAGTATTTCATAATGAAATATTAG
60255
Gallus_gallus                    60211 .............................................
60255

Anas_platyrhynchos_platyrhynchos 60256 TGTATTAGGCAGCTTTACCACACACTCACTCTGTATATAGAGTCC
60300
Gallus_gallus                    60256 .............................................
60300

Anas_platyrhynchos_platyrhynchos 60301 CCTATATAAATTTGTATTAAGAGATCATTTTTTAAATACCTTTCT
60345
Gallus_gallus                    60301 .............................................
60345

Anas_platyrhynchos_platyrhynchos 60346 TTTCCTGTCAACACTTAGTAACTGTAGGGAAATAAATATGTACGT
60390
Gallus_gallus                    60346 .............................................
60390

Anas_platyrhynchos_platyrhynchos 60391 ACATATATACATTAAGCCTATGTATGTAGAAATACATACATTACT
60435
Gallus_gallus                    60391 .............................................
60435

Anas_platyrhynchos_platyrhynchos 60436 TGTTCTAATTCACTGATGTATGCTTCTCCAGCAGCTTTCTCATGA
60480
Gallus_gallus                    60436 .............................................
60480

Anas_platyrhynchos_platyrhynchos 60481 TATGAAGATGCCTGACTAAGTTTTAGGTCTGAACATATCAGGGCA
60525
```

Gallus_gallus 60481 .............................................
60525

Anas_platyrhynchos_platyrhynchos 60526 ATTCTAAGGGAAAAAAAAAGAAAAGAAAAAAAAAAGGAAGAAATGT
60570
Gallus_gallus 60526 .............................................
60570

Anas_platyrhynchos_platyrhynchos 60571 TTAAGAAACAAGAATCTTACAAAAGTTGTTTCATATAAACACATT
60615
Gallus_gallus 60571 .............................................
60615

Anas_platyrhynchos_platyrhynchos 60616 AAAAACATATAAACACATTAAAAAGAAGAGCATATATTCCATATT
60660
Gallus_gallus 60616 .............................................
60660

Anas_platyrhynchos_platyrhynchos 60661 TTTCAACCTTGCTTTCATATGTCTTAACTAATATGGGTAGTAACA
60705
Gallus_gallus 60661 .............................................
60705

Anas_platyrhynchos_platyrhynchos 60706 AACATTTTTTAAAGAAATATAGGTATATTTGAAATTACCTATTGA
60750
Gallus_gallus 60706 .............................................
60750

Anas_platyrhynchos_platyrhynchos 60751 AGAGGTAAAAATATGAAAATAGCATTTTTAATATGGTCTTATTAC
60795
Gallus_gallus 60751 .............................................
60795

Anas_platyrhynchos_platyrhynchos 60796 AGCATGCAAAAACCACAGGCATATGAGCTAACCGGCTACACTACA
60840
Gallus_gallus 60796 .............................................
60840

Anas_platyrhynchos_platyrhynchos 60841 TTAGTAAAGAATCACCTATTCATGTATATCTATCAGATCATATAT
60885
Gallus_gallus 60841 .............................................
60885

Anas_platyrhynchos_platyrhynchos 60886 GAAAAACAGGTCACGAATAAAATAAATAAATAACGAAATAAAAAC
60930
Gallus_gallus 60886 .............................................
60930

Anas_platyrhynchos_platyrhynchos 60931 AACTCTTGAGGCATTAAGAAAAAAATTTATCTGAAAAAGCTTCAT
60975
Gallus_gallus 60931 .............................................
60975

Anas_platyrhynchos_platyrhynchos 60976 TGCAAATCAATTTCGTATTAGTAAACCACTTCAGTCTTGTGAGAC
61020
Gallus_gallus 60976 .............................................
61020

Anas_platyrhynchos_platyrhynchos 61021 AGTAACTCAGTAATCAATTATCCAGATGAGCACACATCCATCTGT
61065
Gallus_gallus 61021 .............................................
61065

```
Anas_platyrhynchos_platyrhynchos 61066 ATACAGAAATGGATGTTTCTACATTTGTCAACTCTATTATGTCAA
61110
Gallus_gallus                    61066 .............................................
61110

Anas_platyrhynchos_platyrhynchos 61111 ATCAGTTTCTAGCATTTTAAAAATAGATTCTATGGCATTTCAGAA
61155
Gallus_gallus                    61111 .............................................
61155

Anas_platyrhynchos_platyrhynchos 61156 ACTATTGGGAATAAAGCTAGTATTAATTATTGCTGATAATAACTT
61200
Gallus_gallus                    61156 .............................................
61200

Anas_platyrhynchos_platyrhynchos 61201 AATTATTAAGCCATTATTATGCAGTACACTTCTGAAGTGTGATAA
61245
Gallus_gallus                    61201 .............................................
61245

Anas_platyrhynchos_platyrhynchos 61246 AATGTTTTAACTGATAAGCAATAATTCACACCTGCATGCATATTA
61290
Gallus_gallus                    61246 .............................................
61290

Anas_platyrhynchos_platyrhynchos 61291 AATATTTTTGTTTTAAAAAAATCAGGGCTTAACATCCATAACGAT
61335
Gallus_gallus                    61291 .............................................
61335

Anas_platyrhynchos_platyrhynchos 61336 CCAGAATTTTGGTCACAAAAGCAACTCTGGAAAGAGAATGACAGA
61380
Gallus_gallus                    61336 .............................................
61380

Anas_platyrhynchos_platyrhynchos 61381 GCTAACTGAGGATACCTATCTGGGTTCTGAAGTAATATATTTTTG
61425
Gallus_gallus                    61381 .............................................
61425

Anas_platyrhynchos_platyrhynchos 61426 ATGCAAGATGGGAGCCTTCCTTTCGAATATTTAAAACATAATATT
61470
Gallus_gallus                    61426 .............................................
61470

Anas_platyrhynchos_platyrhynchos 61471 TTAATTTAATTTTGTTTGGACTCCTTTCAGTTAGATGAGAATAAG
61515
Gallus_gallus                    61471 .............................................
61515

Anas_platyrhynchos_platyrhynchos 61516 ACATTACAAAAATTATTCACATGAAAGTGGCTTGAGACCCCAAAT
61560
Gallus_gallus                    61516 .............................................
61560

Anas_platyrhynchos_platyrhynchos 61561 CTTGTATATCTTTCCTCAAGGAGATGATATTTAGTAACTGATTTT
61605
Gallus_gallus                    61561 .............................................
61605

Anas_platyrhynchos_platyrhynchos 61606 CTGGCTTTAATGCCACAAGCAAAACAATTCATCAAAACACCCTAT
61650
Gallus_gallus                    61606 .............................................
```

61650

Anas_platyrhynchos_platyrhynchos 61651 TAAAATGAAATGAATGAATGAAATGAAGTGAATGAAAAAAAAATA
61695
Gallus_gallus 61651 .............................................
61695

Anas_platyrhynchos_platyrhynchos 61696 ATTTAGAAAATTACAAATATAAAACAAATCACAAAAAGCATTGTT
61740
Gallus_gallus 61696 .............................................
61740

Anas_platyrhynchos_platyrhynchos 61741 TACTATTTAGTGAAACACTGATGGACAGTCCACTTCATTCTGGTT
61785
Gallus_gallus 61741 .............................................
61785

Anas_platyrhynchos_platyrhynchos 61786 CTCTTTTATTTAAAACAAATAAAGAATACAAACCGAAGGTGATAG
61830
Gallus_gallus 61786 .............................................
61830

Anas_platyrhynchos_platyrhynchos 61831 AGATCGGACAGACTTCTACATTCCACAACTTCATTAGCAATAACT
61875
Gallus_gallus 61831 .............................................
61875

Anas_platyrhynchos_platyrhynchos 61876 TCAGCTAGCTGAAAGATGGGAAGTATCAAGTGGGGGCAAAATATT
61920
Gallus_gallus 61876 .............................................
61920

Anas_platyrhynchos_platyrhynchos 61921 TGCTGTAGTTCTTGAGCTAAAATGTCTATAAAATGCAAAGTGCAT
61965
Gallus_gallus 61921 .............................................
61965

Anas_platyrhynchos_platyrhynchos 61966 GTCTGAAAAAAAAATAATTTTGTAGAAAATATATTTAAAATTACT
62010
Gallus_gallus 61966 .............................................
62010

Anas_platyrhynchos_platyrhynchos 62011 GATTAAATATTAAGTTCTGAATGCTACTAAAACAAATATAAACAA
62055
Gallus_gallus 62011 .............................................
62055

Anas_platyrhynchos_platyrhynchos 62056 ACTTTCAAGATGAAAAAGCAGCTACATACACACGAATACAGCAAA
62100
Gallus_gallus 62056 .............................................
62100

Anas_platyrhynchos_platyrhynchos 62101 TTTACTCTGTAGGTTAAGTTCATTTCAAAAATGCAGAGATAGACA
62145
Gallus_gallus 62101 .............................................
62145

Anas_platyrhynchos_platyrhynchos 62146 TTTGAAATATTAAGGAACGGAATGACACAAAAGTATTTCTGATAG
62190
Gallus_gallus 62146 .............................................
62190

Anas_platyrhynchos_platyrhynchos 62191 CTGTCTGAACATACTCTTTATCACAAATACAAGAAAACTAAAATG

```
62235
Gallus_gallus                     62191 .............................................
62235

Anas_platyrhynchos_platyrhynchos 62236 GTACATAGGTCTCCACTATTCAATTTTTCCCTTAAATAAGTGCTC
62280
Gallus_gallus                     62236 .............................................
62280

Anas_platyrhynchos_platyrhynchos 62281 TAGAATTTTGAGGAGTAGACTCACAGGTTTTGTGAAATTATTCCA
62325
Gallus_gallus                     62281 .............................................
62325

Anas_platyrhynchos_platyrhynchos 62326 GTTAATACCATAACTATTTGTTTTCTGTTTAAAAGTATTTCTAAC
62370
Gallus_gallus                     62326 .............................................
62370

Anas_platyrhynchos_platyrhynchos 62371 TTCATTGGGACAGTCATAATGAGCCCATTCTTCCACATTTATTGG
62415
Gallus_gallus                     62371 .............................................
62415

Anas_platyrhynchos_platyrhynchos 62416 TAAGGTATCAACTGACTCCTTCCTTTTAGGCTGCTCTTTCACAAA
62460
Gallus_gallus                     62416 .............................................
62460

Anas_platyrhynchos_platyrhynchos 62461 AGATTCTTCAACCTGTAATTTAAAATACAGCCATTCATGAGGCCC
62505
Gallus_gallus                     62461 .............................................
62505

Anas_platyrhynchos_platyrhynchos 62506 AAGTTGATATTTTGTTTGGTATTAAATTTTTAGCTAACATAAAAA
62550
Gallus_gallus                     62506 .............................................
62550

Anas_platyrhynchos_platyrhynchos 62551 ACTTACTTGTTTAGTGTCTTTTGAAGTTTTGTCTCTTTTTTGACC
62595
Gallus_gallus                     62551 .............................................
62595

Anas_platyrhynchos_platyrhynchos 62596 TCTAGACTGGGGGTTCTGCAAAAAAGCAGTATTGTAACCAATGCA
62640
Gallus_gallus                     62596 .............................................
62640

Anas_platyrhynchos_platyrhynchos 62641 GCAAACAAAAGAAAAGTATCTGTACATTTATGACCTGCCAAGACC
62685
Gallus_gallus                     62641 .............................................
62685

Anas_platyrhynchos_platyrhynchos 62686 ATGCAATCAGAAGTGAAAATATGATAAGAGTCAGAGAAAAGTTCT
62730
Gallus_gallus                     62686 .............................................
62730

Anas_platyrhynchos_platyrhynchos 62731 TCACAAAATACAAGAGTTTTTATAGGAAAGCACTTAAGTTATTTA
62775
Gallus_gallus                     62731 .............................................
62775
```

```
Anas_platyrhynchos_platyrhynchos 62776 ACAGACAAGCACAGGGAAATATCTGTTATGAAGAGACTATTGATT
62820
Gallus_gallus                    62776 .............................................
62820

Anas_platyrhynchos_platyrhynchos 62821 TTATTTCATAGGAACAGGGTTCTTACTTGTTTCTCTTCCTTTTTG
62865
Gallus_gallus                    62821 .............................................
62865

Anas_platyrhynchos_platyrhynchos 62866 TCCTCTTTCTTACGAGCAATTGTTGGCTGTGCATCCTGAGGAGCT
62910
Gallus_gallus                    62866 .............................................
62910

Anas_platyrhynchos_platyrhynchos 62911 GTAAATTGTGAAGACTTTGATAATGCTCCTATAATATGTCCTGAT
62955
Gallus_gallus                    62911 .............................................
62955

Anas_platyrhynchos_platyrhynchos 62956 GCTGACAATGATACCTAGATGAGAAAATAATTACGGTGTTGTTTT
63000
Gallus_gallus                    62956 .............................................
63000

Anas_platyrhynchos_platyrhynchos 63001 TTTTGTTTGTTTGTTTGTTTTTCCTGGAAATTTCTAGTAGTAATA
63045
Gallus_gallus                    63001 .............................................
63045

Anas_platyrhynchos_platyrhynchos 63046 ATTATTCATTGTAAATCACTACCTACTAAAAATTGTCATACACCA
63090
Gallus_gallus                    63046 .............................................
63090

Anas_platyrhynchos_platyrhynchos 63091 TGCTTTTTTACATGCGAGCTTTCTTCACAAAACAAGAAAATTCAA
63135
Gallus_gallus                    63091 .............................................
63135

Anas_platyrhynchos_platyrhynchos 63136 AGTATCCTAGATGCTGGCATTATTTGTACAAGCATTTTGGAAGAT
63180
Gallus_gallus                    63136 .............................................
63180

Anas_platyrhynchos_platyrhynchos 63181 CATGATAATGCACGTAAGCAGCTGAACTGTCAGATGACATCCTAT
63225
Gallus_gallus                    63181 .............................................
63225

Anas_platyrhynchos_platyrhynchos 63226 CCTCCCTGGATCTATAATAAAGCTACTTGGGTGGTTGGGCACTGG
63270
Gallus_gallus                    63226 .............................................
63270

Anas_platyrhynchos_platyrhynchos 63271 AACAGGCTCCCCAAGGAAGTGGTCAAGGAACCGAGCCTGCTGGAG
63315
Gallus_gallus                    63271 .............................................
63315

Anas_platyrhynchos_platyrhynchos 63316 TTCAAGAAGCATTTGGAGAATGCTCTCAGACACGTGATCTTATTT
63360
```

Gallus_gallus                    63316 .............................................
63360

Anas_platyrhynchos_platyrhynchos 63361 TTATGTTTATCTATTTATTTATTGTGTGGTCCTTTGAGGACCCAG
63405
Gallus_gallus                    63361 .............................................
63405

Anas_platyrhynchos_platyrhynchos 63406 GAGTTGGACTCAATGATCCTAGTTGGTCCCTTCCAACTCAGATTT
63450
Gallus_gallus                    63406 .............................................
63450

Anas_platyrhynchos_platyrhynchos 63451 TCTATGATTCTGTGATTTTACTTTTTTAATCTATATGGTTGGAAA
63495
Gallus_gallus                    63451 .............................................
63495

Anas_platyrhynchos_platyrhynchos 63496 TTAAATTCTTACTTTAAACTACTAGTCTAGATTGCTTAGTCCTCT
63540
Gallus_gallus                    63496 .............................................
63540

Anas_platyrhynchos_platyrhynchos 63541 ATAAATAGAGACAGAGGATCAGGGGTTCAAATGAATCTGGAAGAC
63585
Gallus_gallus                    63541 .............................................
63585

Anas_platyrhynchos_platyrhynchos 63586 TGAAAATATGAAAGAGCCACTTTATTGTACTTTTGGAGTGGTCTT
63630
Gallus_gallus                    63586 .............................................
63630

Anas_platyrhynchos_platyrhynchos 63631 CTATATTGCGTCTAGTGAATGTGATATAACAGAAATTTCTAAATA
63675
Gallus_gallus                    63631 .............................................
63675

Anas_platyrhynchos_platyrhynchos 63676 TTTAAACATATGGAAGCATCCAGAGAAAGTTCCTCGCATGTAAAA
63720
Gallus_gallus                    63676 .............................................
63720

Anas_platyrhynchos_platyrhynchos 63721 TAACAACACTTCTGATCTTGCCTAAAGCATAGAAACACGTATTAA
63765
Gallus_gallus                    63721 .............................................
63765

Anas_platyrhynchos_platyrhynchos 63766 TCTGAGAAGATGAACGTATTTCATTAGAGTTATCTAGAGTAGAAG
63810
Gallus_gallus                    63766 .............................................
63810

Anas_platyrhynchos_platyrhynchos 63811 AAAAAATATCACCAAAAAAAAAATAATGCTGGATGCTGCACAACT
63855
Gallus_gallus                    63811 .............................................
63855

Anas_platyrhynchos_platyrhynchos 63856 TCTGCAATGCCACCCGTAGCATTAAGACTGTTTATATGAAGATTT
63900
Gallus_gallus                    63856 .............................................
63900

```
Anas_platyrhynchos_platyrhynchos 63901 GAGGCATTACTATTATCAAGTATAGATTTAGTGCTTTCGAAGATT
63945
Gallus_gallus                    63901 .............................................
63945

Anas_platyrhynchos_platyrhynchos 63946 TACTCATATTTAACATATAATAATAATCCTATGATTTTTGCTATA
63990
Gallus_gallus                    63946 .............................................
63990

Anas_platyrhynchos_platyrhynchos 63991 ATGCTCAATTAACCATTTAAATTGTGCATTCATATCAACGAGAAA
64035
Gallus_gallus                    63991 .............................................
64035

Anas_platyrhynchos_platyrhynchos 64036 TCAAATACACTTGTTTGATTTGAAATGCTGTACCAATGAAAAAGC
64080
Gallus_gallus                    64036 .............................................
64080

Anas_platyrhynchos_platyrhynchos 64081 GATTGAAGAATAATATTTAATGAAAACTGCCCGTCTATACTTTTT
64125
Gallus_gallus                    64081 .............................................
64125

Anas_platyrhynchos_platyrhynchos 64126 TTTCTCAACATGTAACAAACAAATCAACACTTTAATTTATTTTAA
64170
Gallus_gallus                    64126 .............................................
64170

Anas_platyrhynchos_platyrhynchos 64171 CTACATCAGATGATTCCATTTAGTTGAATCACAGCCACTTTAAAT
64215
Gallus_gallus                    64171 .............................................
64215

Anas_platyrhynchos_platyrhynchos 64216 CCATCTAAGTGACAGTCTAAGTCCCCTCCCTCAGAACTGTCAGCA
64260
Gallus_gallus                    64216 .............................................
64260

Anas_platyrhynchos_platyrhynchos 64261 ATGTGTTCATTGTGTATTCTGATGCTAGTGAGATTCTTTAACTAA
64305
Gallus_gallus                    64261 .............................................
64305

Anas_platyrhynchos_platyrhynchos 64306 TTAAAAATACAAAATTTTAGAAACACAAAAAAATCACAAGAAAAG
64350
Gallus_gallus                    64306 .............................................
64350

Anas_platyrhynchos_platyrhynchos 64351 AATTATTGCATTCCAAGGAAAAAAATGGGCCACTGTTTTTATCCTT
64395
Gallus_gallus                    64351 .............................................
64395

Anas_platyrhynchos_platyrhynchos 64396 AAATGTCTGTTTCCCTATTTACCTGAGAGAAATAATTTTCTTGTT
64440
Gallus_gallus                    64396 .............................................
64440

Anas_platyrhynchos_platyrhynchos 64441 TAATCCTAAACCTGAGAAAATATACTAACTACAGCATAAAATTGG
64485
Gallus_gallus                    64441 .............................................
```

```
64485

Anas_platyrhynchos_platyrhynchos 64486 AAACAGTAAAGACATAGCTAAACTGTCAAATGAGTAGTTGATTCC
64530
Gallus_gallus                    64486 .............................................
64530

Anas_platyrhynchos_platyrhynchos 64531 GCAGTAACACTGACAGTTGAAAATATAATGGCCACATTATAACGT
64575
Gallus_gallus                    64531 .............................................
64575

Anas_platyrhynchos_platyrhynchos 64576 AATATCATATAATCTCACCTGCCAGATACGGATAACACAAGCATA
64620
Gallus_gallus                    64576 .............................................
64620

Anas_platyrhynchos_platyrhynchos 64621 TGCCATCAAACAGTGTTGCTGATGAAAAGGAGAACTATGACCAGA
64665
Gallus_gallus                    64621 .............................................
64665

Anas_platyrhynchos_platyrhynchos 64666 AACTACAGCCATTAAAGTCTGTGCTCTAAACAAAGCTTCCAATTG
64710
Gallus_gallus                    64666 .............................................
64710

Anas_platyrhynchos_platyrhynchos 64711 TTTAATATTAGTCAAATTTTCCAAATTGATTCGGGGCATGACATC
64755
Gallus_gallus                    64711 .............................................
64755

Anas_platyrhynchos_platyrhynchos 64756 ATTTCTTCCAACATCTATCTTCAGATTTTCCATAGGCAATAATTT
64800
Gallus_gallus                    64756 .............................................
64800

Anas_platyrhynchos_platyrhynchos 64801 TTTAATAGATTTTTCTTCTGTTAAAAACGAAAAAATAACATGCCT
64845
Gallus_gallus                    64801 .............................................
64845

Anas_platyrhynchos_platyrhynchos 64846 ATATTTGTAGCTATAATAGTATTTTTAAAATTACTCAAAAGTTAA
64890
Gallus_gallus                    64846 .............................................
64890

Anas_platyrhynchos_platyrhynchos 64891 ATTTCTAAATTACAAATAAAATCAGAAAAAAAATCATCTACTGAA
64935
Gallus_gallus                    64891 .............................................
64935

Anas_platyrhynchos_platyrhynchos 64936 TATGTTCTTCTTATGGCAACGTAATGCTACTTGTACTGTGTTTTA
64980
Gallus_gallus                    64936 .............................................
64980

Anas_platyrhynchos_platyrhynchos 64981 AATTAATAGTGTAACAATTATGCATATTAAATGACAGAAGTTCAA
65025
Gallus_gallus                    64981 .............................................
65025

Anas_platyrhynchos_platyrhynchos 65026 CACAGAAATGAAATACACAACAGACACTTTAACTATGTCTTATAT
```

65070
Gallus_gallus                     65026 .............................................
65070

Anas_platyrhynchos_platyrhynchos 65071 GTCTTATTTAACAGGCCACAGTTCCTGCCAGATAACCTATGTGGC
65115
Gallus_gallus                     65071 .............................................
65115

Anas_platyrhynchos_platyrhynchos 65116 TCCTCTCCCTGGGGTCACAGCTCCTGCCAGGAGTCTGCTCCTGTG
65160
Gallus_gallus                     65116 .............................................
65160

Anas_platyrhynchos_platyrhynchos 65161 CAGGCTTTCCACAGACTGCAGCTGCCTGCAGGGCACTTCCACCTC
65205
Gallus_gallus                     65161 .............................................
65205

Anas_platyrhynchos_platyrhynchos 65206 CTCTGGAAGCTGCAGAGTAGATATCTGTTCCAGTGTGGTCCTCCA
65250
Gallus_gallus                     65206 .............................................
65250

Anas_platyrhynchos_platyrhynchos 65251 TAGGCTACAGGGGCAAAACCTGCTTCACTACTGGCCTGCAGGGGA
65295
Gallus_gallus                     65251 .............................................
65295

Anas_platyrhynchos_platyrhynchos 65296 ATCTCTGCTCCAGTATGTGGAGCATCTCCTCCTCCTCCTTTTCTG
65340
Gallus_gallus                     65296 .............................................
65340

Anas_platyrhynchos_platyrhynchos 65341 ACTTTCTGACCTTTTAGCATCTACAAAGTTTTTTTTCTCTCACAT
65385
Gallus_gallus                     65341 .............................................
65385

Anas_platyrhynchos_platyrhynchos 65386 TTTTCTTACTGCAGTCTCTCACAGCTGCCACACAGAGGTGTGGTT
65430
Gallus_gallus                     65386 .............................................
65430

Anas_platyrhynchos_platyrhynchos 65431 TTTTTTTTTTTAACCCTCTCCTATCTTATGCAGAGGAGCAAACAG
65475
Gallus_gallus                     65431 .............................................
65475

Anas_platyrhynchos_platyrhynchos 65476 CATTCTTCATTGTCTCAGCTCTGGACAGTAGTGGATACCTTTTGG
65520
Gallus_gallus                     65476 .............................................
65520

Anas_platyrhynchos_platyrhynchos 65521 AGCCAGCTGGAACTGGCTCTATCCGACACAGGGCAGCACTCGGTC
65565
Gallus_gallus                     65521 .............................................
65565

Anas_platyrhynchos_platyrhynchos 65566 TCTTCTCTAACAGGCCGCCCCTGCAACAACCTCCCTCCCTCCCTG
65610
Gallus_gallus                     65566 .............................................
65610

```
Anas_platyrhynchos_platyrhynchos 65611 CCCCAACCATAAGCATTTTGTGATTTGGTAATGAGCGTAGATCTT
65655
Gallus_gallus                    65611 .............................................
65655

Anas_platyrhynchos_platyrhynchos 65656 TATGACACAACACAGAATTCACAGGTGCTTTTAAAATATCCTTGA
65700
Gallus_gallus                    65656 .............................................
65700

Anas_platyrhynchos_platyrhynchos 65701 ACAAATGACAGTGCATTCTGGAGAAAGTACTGCCATGCTGCTGCC
65745
Gallus_gallus                    65701 .............................................
65745

Anas_platyrhynchos_platyrhynchos 65746 ATTCTTTGTTTGCTAACTAAGTTATAAATTTAAATTGTCCCTTGC
65790
Gallus_gallus                    65746 .............................................
65790

Anas_platyrhynchos_platyrhynchos 65791 ATTTGTTACTGGGAAATTTATTTGAATAATATACACATTTACATA
65835
Gallus_gallus                    65791 .............................................
65835

Anas_platyrhynchos_platyrhynchos 65836 CCTTCTTCTTGAGAAGATTGCATAGAAAATTTCATATGTAGCAAG
65880
Gallus_gallus                    65836 .............................................
65880

Anas_platyrhynchos_platyrhynchos 65881 AGACCTACTGCCCAATCCAGAGGTTTAATAACATCATTAAGGGGA
65925
Gallus_gallus                    65881 .............................................
65925

Anas_platyrhynchos_platyrhynchos 65926 AACTGGTTACAGTATAACCATTCAGCAAATTCCATCAGGTAATCA
65970
Gallus_gallus                    65926 .............................................
65970

Anas_platyrhynchos_platyrhynchos 65971 ACTTTCTGCCATTCATTTTCTTGTTTCTATGAAAGTGATAAAGGA
66015
Gallus_gallus                    65971 .............................................
66015

Anas_platyrhynchos_platyrhynchos 66016 AGAGTTAGTTTATATCTGGTGGTGTAAACCACTGAACTGATATTC
66060
Gallus_gallus                    66016 .............................................
66060

Anas_platyrhynchos_platyrhynchos 66061 AAAAGTTCAATATAGTCTTAATGGAAGCAAAAAAATAATTTGTCA
66105
Gallus_gallus                    66061 .............................................
66105

Anas_platyrhynchos_platyrhynchos 66106 CTGAGCAGTTTCATTTGAATCACATAATCAGAGACATATGTAAAA
66150
Gallus_gallus                    66106 .............................................
66150

Anas_platyrhynchos_platyrhynchos 66151 CAGAAAGCAACAGAAAACTCTATTTAGTTTATTCTTTGCTTTTTC
66195
```

```
Gallus_gallus                    66151 .............................................
66195

Anas_platyrhynchos_platyrhynchos 66196 AAAATTTATTGCAATTAGCTATTCTTAGACTTTTTCTTTGTCAGG
66240
Gallus_gallus                    66196 .............................................
66240

Anas_platyrhynchos_platyrhynchos 66241 ACTCAGCTGAACTCTTATGGCCCTCAAAAGGTTCCTCATTTGAGA
66285
Gallus_gallus                    66241 .............................................
66285

Anas_platyrhynchos_platyrhynchos 66286 AGGTGCTTGTGCACTGAAGTCAAGGCACTGAGCCTGAGAGAGTTC
66330
Gallus_gallus                    66286 .............................................
66330

Anas_platyrhynchos_platyrhynchos 66331 AAGAGGTGTTCGGAAAATGTTGTCAAATATATAATTACTAGTCTA
66375
Gallus_gallus                    66331 .............................................
66375

Anas_platyrhynchos_platyrhynchos 66376 ATAGTTAGCCCTGCATGGAGCCAGGAGTGGGACTCAATGGTCCTT
66420
Gallus_gallus                    66376 .............................................
66420

Anas_platyrhynchos_platyrhynchos 66421 GTAGATTCCCTTCCCACTAAGGATATTCTATGATTCTATGATTTT
66465
Gallus_gallus                    66421 .............................................
66465

Anas_platyrhynchos_platyrhynchos 66466 CCAACTGAAAAGCAATTTCAAATTTACAAGAAACAATGATCATAT
66510
Gallus_gallus                    66466 .............................................
66510

Anas_platyrhynchos_platyrhynchos 66511 ATATGGCCTATCTACTCCTTAAATTCTATACTTCTCAATACTTGC
66555
Gallus_gallus                    66511 .............................................
66555

Anas_platyrhynchos_platyrhynchos 66556 TATCTCATTAAACACTTTAAAAACATATTTTTATCAGTATTGGAA
66600
Gallus_gallus                    66556 .............................................
66600

Anas_platyrhynchos_platyrhynchos 66601 CTGATAGTAAAAATTCACAAACCTGCAAAGCATTGATTGCATTTT
66645
Gallus_gallus                    66601 .............................................
66645

Anas_platyrhynchos_platyrhynchos 66646 TAAAACAGTTTAACTGTTCAACAATACTTCTGGAAACTGTTACCA
66690
Gallus_gallus                    66646 .............................................
66690

Anas_platyrhynchos_platyrhynchos 66691 TATGACGCCAAATACGGGACACATAATCTTCACTTTCACCTGCAA
66735
Gallus_gallus                    66691 .............................................
66735
```

```
Anas_platyrhynchos_platyrhynchos 66736 ATTTCTGAATTTCCATCATGAAATTACATCCCAATCTTGCCTTTA
66780
Gallus_gallus                    66736 .............................................
66780

Anas_platyrhynchos_platyrhynchos 66781 CTGTAGCCAAATGTTTAAAAATCAGACTAGAAAAGGACAAAAACA
66825
Gallus_gallus                    66781 .............................................
66825

Anas_platyrhynchos_platyrhynchos 66826 ATTGCACATTAATTGCATCTGTATCTGTAAAGTAACTCGAAGTGG
66870
Gallus_gallus                    66826 .............................................
66870

Anas_platyrhynchos_platyrhynchos 66871 AATAGATATAGTAGTTCAGTGTAGTGCAGAGATGTGTAAGTTCAT
66915
Gallus_gallus                    66871 .............................................
66915

Anas_platyrhynchos_platyrhynchos 66916 AAACTAATATAAAAGGGCATAAATCTTTGAACTCATGTAAGACTA
66960
Gallus_gallus                    66916 .............................................
66960

Anas_platyrhynchos_platyrhynchos 66961 TATATAAACAGTAGCCTAGATAATTTAGTTTCCATTTTACATTAC
67005
Gallus_gallus                    66961 .............................................
67005

Anas_platyrhynchos_platyrhynchos 67006 GTGTGTTTACAGCTAGAAAAATTATAACTTCAGCTACATTTCGTA
67050
Gallus_gallus                    67006 .............................................
67050

Anas_platyrhynchos_platyrhynchos 67051 CTCCTATTTTCTTGGAAAAATTAGATAGTACCTGTATACAAGTTT
67095
Gallus_gallus                    67051 .............................................
67095

Anas_platyrhynchos_platyrhynchos 67096 TAGAAAATGACATGTACAAATGTTATGAAAAGATTAATGATAAGA
67140
Gallus_gallus                    67096 .............................................
67140

Anas_platyrhynchos_platyrhynchos 67141 AACACTGCAAACATAAGTCTTTTCACTGAAAAGACACACATGTGT
67185
Gallus_gallus                    67141 .............................................
67185

Anas_platyrhynchos_platyrhynchos 67186 TTTCTGTTATAAGAATTGCCCAGCTGACCAAAACGTAGTCATGAT
67230
Gallus_gallus                    67186 .............................................
67230

Anas_platyrhynchos_platyrhynchos 67231 TTTATTCTGCCAATGCAAAATAGAAAAATAGTAAAAACAGAACTC
67275
Gallus_gallus                    67231 .............................................
67275

Anas_platyrhynchos_platyrhynchos 67276 AGAGTTTTAGAGTTCTCAAGAACCCAAATGCTTATTTTAATAAAA
67320
Gallus_gallus                    67276 .............................................
```

```
67320

Anas_platyrhynchos_platyrhynchos 67321 ATAAATAAATAACTTACAGCCTGTGTCTTGTCTGAGGCAAAGCCT
67365
Gallus_gallus                    67321 .............................................
67365

Anas_platyrhynchos_platyrhynchos 67366 GAATAGCTTCATTCAGGACTTCTAATGCTTTTTCCCAATCAGATT
67410
Gallus_gallus                    67366 .............................................
67410

Anas_platyrhynchos_platyrhynchos 67411 TATCAGCATATATGTGGAATAATAACCCATATAAAGAAGTCCTTA
67455
Gallus_gallus                    67411 .............................................
67455

Anas_platyrhynchos_platyrhynchos 67456 ATGTCAGATCTTCATCAGCACCTTTGAAGTATATTCAAGTCATTT
67500
Gallus_gallus                    67456 .............................................
67500

Anas_platyrhynchos_platyrhynchos 67501 ATTACTATACTGCAGAATTAGAACAATGCTATTACAAAATACAGG
67545
Gallus_gallus                    67501 .............................................
67545

Anas_platyrhynchos_platyrhynchos 67546 TTTTTGTTAGTTTGTTTTTAATTATCATATCAAAACAAGAAAGAA
67590
Gallus_gallus                    67546 .............................................
67590

Anas_platyrhynchos_platyrhynchos 67591 CGTGCCTGGCAGAAAGCATCCATCTGATAATCCACTACTTTGAAA
67635
Gallus_gallus                    67591 .............................................
67635

Anas_platyrhynchos_platyrhynchos 67636 ATCCTTTGTGAGCCACTGATGCAAAGGCAATGTATCATTTTTCTC
67680
Gallus_gallus                    67636 .............................................
67680

Anas_platyrhynchos_platyrhynchos 67681 CTAAAAAGACAGTAGGAACAATGCCCTTTATATTTTGGTACATAT
67725
Gallus_gallus                    67681 .............................................
67725

Anas_platyrhynchos_platyrhynchos 67726 CTCAAGTAAGCAAACAACCACTGCCTTGCTGGGCTAAACCAAAAG
67770
Gallus_gallus                    67726 .............................................
67770

Anas_platyrhynchos_platyrhynchos 67771 TTATCTTCAGGTATACATGTATGCTTCTCTAGATATGTGAACAAA
67815
Gallus_gallus                    67771 .............................................
67815

Anas_platyrhynchos_platyrhynchos 67816 TTTAAAAACTGCTTAGAGTTAGCCTGATCAAGGATCACTGGATAC
67860
Gallus_gallus                    67816 .............................................
67860

Anas_platyrhynchos_platyrhynchos 67861 CAGTTCCTGCCTCTTACAGCAGTTACTATAACATATAAAGAATTA
```

67905
Gallus_gallus 67861 .............................................
67905

Anas_platyrhynchos_platyrhynchos 67906 GTAACTCTGGAAATACTCAAATACAGAACTTATTTGAAATATTTT
67950
Gallus_gallus 67906 .............................................
67950

Anas_platyrhynchos_platyrhynchos 67951 TGTCTAAATAATATCAGAAAATCCTATATAGTCTAATTCACAAGA
67995
Gallus_gallus 67951 .............................................
67995

Anas_platyrhynchos_platyrhynchos 67996 ATTACTATTCTTTACAGATATAAAAAGTCTGGGCTCTCTGTGACT
68040
Gallus_gallus 67996 .............................................
68040

Anas_platyrhynchos_platyrhynchos 68041 GTACACTTCCATTATCTAAATAAAGGTCTTGACTTTGCATATATC
68085
Gallus_gallus 68041 .............................................
68085

Anas_platyrhynchos_platyrhynchos 68086 TTATTAAGAGTTTGTTCTGAAAATATCCTGTAGCAGCAAGTTATT
68130
Gallus_gallus 68086 .............................................
68130

Anas_platyrhynchos_platyrhynchos 68131 TATATATACATAATGAAGTATTCCTTCTTACTGGTTGGTTTTTTT
68175
Gallus_gallus 68131 .............................................
68175

Anas_platyrhynchos_platyrhynchos 68176 TTTTTTTTACTCTGCAAATAAGTAGACCTTTAGAAATGTCCCTAT
68220
Gallus_gallus 68176 .............................................
68220

Anas_platyrhynchos_platyrhynchos 68221 GGGAAATCACAGTGTTAGATATCTTAAAGTTCCTTTAAGTTTACG
68265
Gallus_gallus 68221 .............................................
68265

Anas_platyrhynchos_platyrhynchos 68266 ATCTTCATGTCACTGTATTTCCTCGCACACAGAGTCCTCACAAAT
68310
Gallus_gallus 68266 .............................................
68310

Anas_platyrhynchos_platyrhynchos 68311 TCTGTCTCCTACACACTCAAGCAGACCAGTCACCATCTCTGATTA
68355
Gallus_gallus 68311 .............................................
68355

Anas_platyrhynchos_platyrhynchos 68356 TTAGTGATTTATACATGGCTTGAGGAGAAAGGAAAAATAAGTTTT
68400
Gallus_gallus 68356 .............................................
68400

Anas_platyrhynchos_platyrhynchos 68401 CTATTATTATACTACAGTACTAGATGATCTGAGATGCAGACTGAG
68445
Gallus_gallus 68401 .............................................
68445

```
Anas_platyrhynchos_platyrhynchos 68446 ATGTCACATTGGTACCACTGAACTTGAATGCAAAAGTTATTAATA
68490
Gallus_gallus                    68446 .............................................
68490

Anas_platyrhynchos_platyrhynchos 68491 AAATCTTCAAGGTCTAATTCTGCAACCACTGAAGTGCATGGGGAA
68535
Gallus_gallus                    68491 .............................................
68535

Anas_platyrhynchos_platyrhynchos 68536 ACTCACATTAGCTTCATTGACATTTAAATCAAATCTAGAAAGAGA
68580
Gallus_gallus                    68536 .............................................
68580

Anas_platyrhynchos_platyrhynchos 68581 GAAATGGATCAGCTCTAGGAATTTTGAAGTACGAACAGTTGTAGA
68625
Gallus_gallus                    68581 .............................................
68625

Anas_platyrhynchos_platyrhynchos 68626 TGTGTAGGGATGTTTCAGCAGGAAAAGAAATTTCATGTGATGAGA
68670
Gallus_gallus                    68626 .............................................
68670

Anas_platyrhynchos_platyrhynchos 68671 TTTGTTAAAAAATAATTTTGCAGGTAATTTGAATCTGCAGGTGTG
68715
Gallus_gallus                    68671 .............................................
68715

Anas_platyrhynchos_platyrhynchos 68716 AGGAACCTTAATTTAATTCCAACTTTTATGAGTTTGTTTTTTAAG
68760
Gallus_gallus                    68716 .............................................
68760

Anas_platyrhynchos_platyrhynchos 68761 GCCTATTATCATTCTTACTGGAAGTGACAGATTTTTTCTGTCACT
68805
Gallus_gallus                    68761 .............................................
68805

Anas_platyrhynchos_platyrhynchos 68806 TTATCTTTTTTTGATTATTTTCTAAGCATTCCCTTTTGTGAGCTT
68850
Gallus_gallus                    68806 .............................................
68850

Anas_platyrhynchos_platyrhynchos 68851 CACAAAGTGCAACACTACACACATCCAGACCGTAAATTATTTTGA
68895
Gallus_gallus                    68851 .............................................
68895

Anas_platyrhynchos_platyrhynchos 68896 AGAAAACAGCTGCAAGTTTGTCAATTGCCTAGAACCAGAAGAATC
68940
Gallus_gallus                    68896 .............................................
68940

Anas_platyrhynchos_platyrhynchos 68941 ACCAAATGGAAAACAAGTATGTGATTCCTTGTTGTGTCTGAATGT
68985
Gallus_gallus                    68941 .............................................
68985

Anas_platyrhynchos_platyrhynchos 68986 GTGCTATGCAAATGAAGATTAGAAAGCATTTACGTGTTTATGATA
69030
```

```
Gallus_gallus                     68986 .............................................
69030

Anas_platyrhynchos_platyrhynchos 69031 AGACCTTTTCCTTTTAGGAAAGATATCTTGAAGTATGCACACAGC
69075
Gallus_gallus                     69031 .............................................
69075

Anas_platyrhynchos_platyrhynchos 69076 TTCAGAACAAAGCAAAACTTTTCTTTATCATCAGTACCAATCACT
69120
Gallus_gallus                     69076 .............................................
69120

Anas_platyrhynchos_platyrhynchos 69121 AAACGCATGCATAGTTAACAATATTTATTAGCTATTCTTACCTGT
69165
Gallus_gallus                     69121 .............................................
69165

Anas_platyrhynchos_platyrhynchos 69166 TTATTTTCAGATTCTGCTTTCATTATATTCTTAAGAATTATTTCA
69210
Gallus_gallus                     69166 .............................................
69210

Anas_platyrhynchos_platyrhynchos 69211 GTAGGTTCTTTAAACTGTTCTCTGTCATGTGGAAAACCTAGCAAA
69255
Gallus_gallus                     69211 .............................................
69255

Anas_platyrhynchos_platyrhynchos 69256 GGTAAACATGCGTTCCAAAAGTGTCTCGCTGCAAACATTACAAGG
69300
Gallus_gallus                     69256 .............................................
69300

Anas_platyrhynchos_platyrhynchos 69301 GAATAATTTCCAGCTTCACCTGCATATCTGAAGAAAACAAATTAC
69345
Gallus_gallus                     69301 .............................................
69345

Anas_platyrhynchos_platyrhynchos 69346 AGCTTAATGAAGAAAAAAACCTCACTGTAACAAAGATACAGGATT
69390
Gallus_gallus                     69346 .............................................
69390

Anas_platyrhynchos_platyrhynchos 69391 TTACTTCTGCAAACATCATCTGTGGTTTACCTTATCTAGACAACA
69435
Gallus_gallus                     69391 .............................................
69435

Anas_platyrhynchos_platyrhynchos 69436 CTGTATCTCAGCAATGAGGGGTTAGCATATCAGCTTAATACAAAA
69480
Gallus_gallus                     69436 .............................................
69480

Anas_platyrhynchos_platyrhynchos 69481 TATATTTTCCATTTTTATATACAAAACACCTCCATACATCCAACA
69525
Gallus_gallus                     69481 .............................................
69525

Anas_platyrhynchos_platyrhynchos 69526 GAAGGAATATTGAACAAGTGTTTGTACTGTATTCCTGCATAACTT
69570
Gallus_gallus                     69526 .............................................
69570
```

```
Anas_platyrhynchos_platyrhynchos 69571 TAACTGAAGTGAGACCTGATTTTAAAAGTTAGTAGAAACCTACTT
69615
Gallus_gallus                    69571 .............................................
69615

Anas_platyrhynchos_platyrhynchos 69616 GAGAGGTTCTTTGGAATAACCTACTTTTTTTTTTCTTTTCTTCCT
69660
Gallus_gallus                    69616 .............................................
69660

Anas_platyrhynchos_platyrhynchos 69661 ATTATGTTAATTCCTCGTTATAAAAGCATCACACACACAAAAAAA
69705
Gallus_gallus                    69661 .............................................
69705

Anas_platyrhynchos_platyrhynchos 69706 ATAAACAAACAAACAAACACAACTCTTCAGTCCTATGGATTTTGC
69750
Gallus_gallus                    69706 .............................................
69750

Anas_platyrhynchos_platyrhynchos 69751 AAAGTACAGAATCACTGTAAAATAAAAATATACTTCAAATGCCTC
69795
Gallus_gallus                    69751 .............................................
69795

Anas_platyrhynchos_platyrhynchos 69796 TACTTGCATGCAAATTGTTTTAAGCACATTCCATGCAAATGTTAC
69840
Gallus_gallus                    69796 .............................................
69840

Anas_platyrhynchos_platyrhynchos 69841 TTTTAATTAATAAAATATAATAATTTTCCAGACTGTAATACAACA
69885
Gallus_gallus                    69841 .............................................
69885

Anas_platyrhynchos_platyrhynchos 69886 ATAACACACATGAGCGAGCAAAATCTTTAGAATTTAGCTAAAAAT
69930
Gallus_gallus                    69886 .............................................
69930

Anas_platyrhynchos_platyrhynchos 69931 TATGGATTGGTATTAAAAAATCAACTAATTAATACTCTTACTATT
69975
Gallus_gallus                    69931 .............................................
69975

Anas_platyrhynchos_platyrhynchos 69976 TTCATAGAATGATAAAATATCCTGAGCTGGAAAGGACCCACAAGG
70020
Gallus_gallus                    69976 .............................................
70020

Anas_platyrhynchos_platyrhynchos 70021 ATCATCAAGTCCAACTCCTGGCACCACACGGGAGCACCCAAAAAT
70065
Gallus_gallus                    70021 .............................................
70065

Anas_platyrhynchos_platyrhynchos 70066 TCAGACCGTATGTCTAAGAGCACAGTCCAAACACTTCTTAAACCC
70110
Gallus_gallus                    70066 .............................................
70110

Anas_platyrhynchos_platyrhynchos 70111 TGACAGGCTTGGTGCCGTGACTACGTCCCTGGTGAGCCTGTTCCA
70155
Gallus_gallus                    70111 .............................................
```

70155

Anas_platyrhynchos_platyrhynchos 70156 GTGCACGACCATCCTCTTGGTGAAGAACCATTTTCTGATATCCAG
70200
Gallus_gallus 70156 .............................................
70200

Anas_platyrhynchos_platyrhynchos 70201 CCTGAACCTCCCCTGTCACAGCTTAATCATTTTTGAATCAGGCAA
70245
Gallus_gallus 70201 .............................................
70245

Anas_platyrhynchos_platyrhynchos 70246 TCTGTCATTTTCTCCCAAAATTTCACAATGTTTAGACTCTATCTG
70290
Gallus_gallus 70246 .............................................
70290

Anas_platyrhynchos_platyrhynchos 70291 CTCACATGCTGTATATATTCACCCTAGACTTACTAAGTTTTTGCT
70335
Gallus_gallus 70291 .............................................
70335

Anas_platyrhynchos_platyrhynchos 70336 CACCACTTTGTGTCCCTTTTGACATTTGTTGAGTTATTTGTATAT
70380
Gallus_gallus 70336 .............................................
70380

Anas_platyrhynchos_platyrhynchos 70381 GTCCACTCCCAAAAAAAAGTGCTTGTCATGGTATATAAATTGAAA
70425
Gallus_gallus 70381 .............................................
70425

Anas_platyrhynchos_platyrhynchos 70426 AAATATGGATTTGTCTGTTCACAACACAACATAACTGTAGAGTTC
70470
Gallus_gallus 70426 .............................................
70470

Anas_platyrhynchos_platyrhynchos 70471 TGTTACGCTGGCCTTTCCAACTTCATTCATCACTCTCTTTTTAAT
70515
Gallus_gallus 70471 .............................................
70515

Anas_platyrhynchos_platyrhynchos 70516 TATTAAGGAAAAAAAATGATACTGCTAAGGGCTTCACATCAGCCA
70560
Gallus_gallus 70516 .............................................
70560

Anas_platyrhynchos_platyrhynchos 70561 TCACATTTTCATCCTCTAACAGTTTCAATTGCCTACTTTTTTGCT
70605
Gallus_gallus 70561 .............................................
70605

Anas_platyrhynchos_platyrhynchos 70606 CTTTTTTTTTATTTTTTTTTTTCCCATAAAGCAAAGAATTTAAA
70650
Gallus_gallus 70606 .............................................
70650

Anas_platyrhynchos_platyrhynchos 70651 AATCAGGCATTATCCATCCATGTATCACCTTCTTCCTATTTATTA
70695
Gallus_gallus 70651 .............................................
70695

Anas_platyrhynchos_platyrhynchos 70696 CTTAAAGCAATCTTTCCACTTCGGTTTCTAAACTGATAAAATTTA

```
70740
Gallus_gallus                     70696 .............................................
70740

Anas_platyrhynchos_platyrhynchos 70741 TTTTAAATTTACCAAACTGCTAAATTCTGTCAAATCTTTTGTTAA
70785
Gallus_gallus                     70741 .............................................
70785

Anas_platyrhynchos_platyrhynchos 70786 GCATTTAAACTCTCAACTATTTTGATTTTTTATCAGTCTTGGGAA
70830
Gallus_gallus                     70786 .............................................
70830

Anas_platyrhynchos_platyrhynchos 70831 AGAATGATGATCCCCAAATCTTTAAATCAATTTTTTTTGTCTGCT
70875
Gallus_gallus                     70831 .............................................
70875

Anas_platyrhynchos_platyrhynchos 70876 TTGTATTAGTCCCCTCTTAAAACAAAACCTACCTGGCACTCTGAG
70920
Gallus_gallus                     70876 .............................................
70920

Anas_platyrhynchos_platyrhynchos 70921 CAAAGGCTTTTGATGCAGCTATACGCAAATGTTTTTGTCCAGCCA
70965
Gallus_gallus                     70921 .............................................
70965

Anas_platyrhynchos_platyrhynchos 70966 AATTTTCCATTATACTCTTTCCTTGGATACAGGCAGCAGTACTCA
71010
Gallus_gallus                     70966 .............................................
71010

Anas_platyrhynchos_platyrhynchos 71011 ACATTTCTTGTCTCGCTGTGTTATCAGGTCTGAATAAAGACAAGT
71055
Gallus_gallus                     71011 .............................................
71055

Anas_platyrhynchos_platyrhynchos 71056 AACTCAAAATGTGCTAGTCTTACAGTCTAAGAACACACCAGTTCC
71100
Gallus_gallus                     71056 .............................................
71100

Anas_platyrhynchos_platyrhynchos 71101 CATTAATAATGCTTTCAAGTTGTTACTAGAGAGATGTTAGTAGCT
71145
Gallus_gallus                     71101 .............................................
71145

Anas_platyrhynchos_platyrhynchos 71146 TTGAAAGTTAGAAAGGCTTCTAATGACTTTTTTCCTACAATCAGA
71190
Gallus_gallus                     71146 .............................................
71190

Anas_platyrhynchos_platyrhynchos 71191 AAATATTTGAAAAATCTGAACACTCTGTGAACAGGTATCAGAAAT
71235
Gallus_gallus                     71191 .............................................
71235

Anas_platyrhynchos_platyrhynchos 71236 AAAGTATCTTTTTTACAGTTCTAATGAAAATAAATGTGAACTAAG
71280
Gallus_gallus                     71236 .............................................
71280
```

```
Anas_platyrhynchos_platyrhynchos 71281 ATTTGAAAAATATTTTACCTACTTCTAGTGACCAAAGAAAATAGA
71325
Gallus_gallus                    71281 .............................................
71325

Anas_platyrhynchos_platyrhynchos 71326 TTTGTGATTCTCTCCCCATTATTCACCATTGTACATCAAACAACA
71370
Gallus_gallus                    71326 .............................................
71370

Anas_platyrhynchos_platyrhynchos 71371 TAAAAATATTAGATCAGAATGAAATTTACATAGCAAGTAATTAAA
71415
Gallus_gallus                    71371 .............................................
71415

Anas_platyrhynchos_platyrhynchos 71416 AACAGAAATTGGGAAAATCAGACAATTTCAAATACTTTTTTTTTT
71460
Gallus_gallus                    71416 .............................................
71460

Anas_platyrhynchos_platyrhynchos 71461 TTCTTGGAAGTGGTCTCTCTTATGTCTGGTAAAGTACGGACAGGG
71505
Gallus_gallus                    71461 .............................................
71505

Anas_platyrhynchos_platyrhynchos 71506 TTCTAGAGAAATTGACTTGGATTATACTCTTCTACATTTAAAAGT
71550
Gallus_gallus                    71506 .............................................
71550

Anas_platyrhynchos_platyrhynchos 71551 AAGGAGCAGAGATTTCAATCCCATCAGTTAACTAGTTAAAAACAA
71595
Gallus_gallus                    71551 .............................................
71595

Anas_platyrhynchos_platyrhynchos 71596 ACAAACAAACAAACAAAAACAATAACAAAGCAATAATAAAAAATA
71640
Gallus_gallus                    71596 .............................................
71640

Anas_platyrhynchos_platyrhynchos 71641 TCTGATATACTTGAATAGAAGTGGTTTTGAAGGAAAATAATACAA
71685
Gallus_gallus                    71641 .............................................
71685

Anas_platyrhynchos_platyrhynchos 71686 TAAAGTAATACAATATGGAAGATTTTTAAAAGAAAAGTAAAAAAA
71730
Gallus_gallus                    71686 .............................................
71730

Anas_platyrhynchos_platyrhynchos 71731 AAAAAGAAAAGAGAAAAAAGAACAAAAAAAAAAAAGAAAGAAAGA
71775
Gallus_gallus                    71731 .............................................
71775

Anas_platyrhynchos_platyrhynchos 71776 AAGGGGAGAATTCACCTTTACTAAACTGGCATTTCTCGTTTCTAG
71820
Gallus_gallus                    71776 .............................................
71820

Anas_platyrhynchos_platyrhynchos 71821 TTGGCTTAGTTCCTACCGCGCAGTGAGGGCTATTATGCAGTTATG
71865
```

```
Gallus_gallus                     71821 .............................................
71865

Anas_platyrhynchos_platyrhynchos 71866 ATTTGTGTCCACAAGAGGACACCAGTGTTCTACAGTACAAGTTAA
71910
Gallus_gallus                     71866 .............................................
71910

Anas_platyrhynchos_platyrhynchos 71911 CAACAACAACAACAACAACAACAACAAAGTAGGTATCTAAGCCTT
71955
Gallus_gallus                     71911 .............................................
71955

Anas_platyrhynchos_platyrhynchos 71956 CTTTGTCCTTAAGTTGTGCTTTAACGTGAAAAGAATGGAGACGTT
72000
Gallus_gallus                     71956 .............................................
72000

Anas_platyrhynchos_platyrhynchos 72001 CATCTTCCATTTTTCTGGAGATAATATTGACAACCCACAACTCAA
72045
Gallus_gallus                     72001 .............................................
72045

Anas_platyrhynchos_platyrhynchos 72046 CTTCTTGGCCAAAGAAGGAAACATTTTACATTTTAACATTCACCA
72090
Gallus_gallus                     72046 .............................................
72090

Anas_platyrhynchos_platyrhynchos 72091 GATTGCTAAATATTCATTATTTTCTCTCTATAATATTTTTAAATT
72135
Gallus_gallus                     72091 .............................................
72135

Anas_platyrhynchos_platyrhynchos 72136 ATATGTTCCATATTTTAAGATGTTTTGGAGAGAAAAACTGGTTTT
72180
Gallus_gallus                     72136 .............................................
72180

Anas_platyrhynchos_platyrhynchos 72181 AAATGTTCTTCTCCCATGAAATAAACAACCATTAATTTCAAAGAG
72225
Gallus_gallus                     72181 .............................................
72225

Anas_platyrhynchos_platyrhynchos 72226 CAGCATTGTTTCCCAGACATCTTCACAGAGAGAAAAATGAACACA
72270
Gallus_gallus                     72226 .............................................
72270

Anas_platyrhynchos_platyrhynchos 72271 TAAGTATATTCAAAGTATTTATCATATTCATTTCAAACTTCCTGC
72315
Gallus_gallus                     72271 .............................................
72315

Anas_platyrhynchos_platyrhynchos 72316 ATTCTGATTTTTTTTTTTTTTAAATAATTGCTGTCAATTCCCTCT
72360
Gallus_gallus                     72316 .............................................
72360

Anas_platyrhynchos_platyrhynchos 72361 TGTTTTGATGAAATTACATTCTGTTTTTGGGGGGGTAAAATCGTA
72405
Gallus_gallus                     72361 .............................................
72405
```

```
Anas_platyrhynchos_platyrhynchos 72406 TATAAGCAGAAAGATCTAAGGACTCCACCAACAGATCCTCCTATA
72450
Gallus_gallus                    72406 .............................................
72450

Anas_platyrhynchos_platyrhynchos 72451 TAGGAAACCTTTTTTCTTTTTTCCTTCTATTCCCCACAAGATCAA
72495
Gallus_gallus                    72451 .............................................
72495

Anas_platyrhynchos_platyrhynchos 72496 AGCTTTAAATTAACAGTCCAGTAAGACTCTGGGCTTTTTAAGAAA
72540
Gallus_gallus                    72496 .............................................
72540

Anas_platyrhynchos_platyrhynchos 72541 TGAATTACACTCCTATTGCTCCCACATACAGAATTTATAAGGGAA
72585
Gallus_gallus                    72541 .............................................
72585

Anas_platyrhynchos_platyrhynchos 72586 GCCTGCTACCTGGGGATACCTGACACCAGAAATTAGAAAACAGTA
72630
Gallus_gallus                    72586 .............................................
72630

Anas_platyrhynchos_platyrhynchos 72631 GCTGTTATCACCTGTTATCACTCACAGCATCTTCTTACATGAACT
72675
Gallus_gallus                    72631 .............................................
72675

Anas_platyrhynchos_platyrhynchos 72676 GTGTAGACCCTTCATGTATTATCAGAAGCAATTTTCATCAATGTA
72720
Gallus_gallus                    72676 .............................................
72720

Anas_platyrhynchos_platyrhynchos 72721 TTAGAATAAGTTACTTTCTGCTTGGAAACTACAAAAGCACGAAAA
72765
Gallus_gallus                    72721 .............................................
72765

Anas_platyrhynchos_platyrhynchos 72766 GACTGTGACTGCCACAAACCAATTTCAGAAAAGCACTGCTCATCA
72810
Gallus_gallus                    72766 .............................................
72810

Anas_platyrhynchos_platyrhynchos 72811 GGAAGACACTCAAACACAAGTATATAACCTGTTGAGTCACTTGGA
72855
Gallus_gallus                    72811 .............................................
72855

Anas_platyrhynchos_platyrhynchos 72856 TTACAAAATTCACTTATCTTCCTGAACTGAGTCCAGAGACACAAT
72900
Gallus_gallus                    72856 .............................................
72900

Anas_platyrhynchos_platyrhynchos 72901 CTGGATCTCACACCATCTTCTACTCAGGCTACACTGTAGAGTTAA
72945
Gallus_gallus                    72901 .............................................
72945

Anas_platyrhynchos_platyrhynchos 72946 AACATTATCTGTAGTACTTAAGTAATTGCTAATGACAAAATCAGG
72990
Gallus_gallus                    72946 .............................................
```

72990

Anas_platyrhynchos_platyrhynchos 72991 AGTGGTAGGAAAATTTCACTTTCTGTACAGTTTCTTTGTAAATAA
73035
Gallus_gallus 72991 .............................................
73035

Anas_platyrhynchos_platyrhynchos 73036 ATAAATAAATAAATAAATTTCTCTCAATTTTGATTAAGTAGTCAG
73080
Gallus_gallus 73036 .............................................
73080

Anas_platyrhynchos_platyrhynchos 73081 ATGCATTTTCCCTATATATTCATTCTTCAAAATTTATATCAAGTA
73125
Gallus_gallus 73081 .............................................
73125

Anas_platyrhynchos_platyrhynchos 73126 TACCACATTATAATGCAAAATTCAATGTACTTACATTTCATATTT
73170
Gallus_gallus 73126 .............................................
73170

Anas_platyrhynchos_platyrhynchos 73171 CTTGGTACCGCTATTGTGAAAAAAATTATTAACTGACTGCAATAT
73215
Gallus_gallus 73171 .............................................
73215

Anas_platyrhynchos_platyrhynchos 73216 CCTTTGGGAACAAGTCATCACTATTTCGTAGTCATGTAATACATA
73260
Gallus_gallus 73216 .............................................
73260

Anas_platyrhynchos_platyrhynchos 73261 CGCAAAATATGTAAGTCGTGTCAGTGTCTGCAGTTCTACTAATGA
73305
Gallus_gallus 73261 .............................................
73305

Anas_platyrhynchos_platyrhynchos 73306 ATCTGACCAGCTGCATTCTAAAGCCAATTTCACTAACTGGAAAAA
73350
Gallus_gallus 73306 .............................................
73350

Anas_platyrhynchos_platyrhynchos 73351 AAAAAAAAAAAAAAAAGTAAAACAAAAATAAATCTTGTGTTTCTG
73395
Gallus_gallus 73351 .............................................
73395

Anas_platyrhynchos_platyrhynchos 73396 CCATGCCCTTTTCATACATAATACTTACTTATCCTCACAACAGCT
73440
Gallus_gallus 73396 .............................................
73440

Anas_platyrhynchos_platyrhynchos 73441 TACTCATATACAATCAGATTACATTCTTCCAGATGTACTTTTCAT
73485
Gallus_gallus 73441 .............................................
73485

Anas_platyrhynchos_platyrhynchos 73486 ACAACCCACCTTGGCATGGAACAAGTAAACAATAAAACAATCTAG
73530
Gallus_gallus 73486 .............................................
73530

Anas_platyrhynchos_platyrhynchos 73531 AAAACCAAAGATTTACAAGAAAAATCAAAATAGGCAATAAACATA

73575
Gallus_gallus                    73531 .............................................
73575

Anas_platyrhynchos_platyrhynchos 73576 TAATCAACATTTACGCTGCAGTAAATATTGTAAAAGCAAATGTAA
73620
Gallus_gallus                    73576 .............................................
73620

Anas_platyrhynchos_platyrhynchos 73621 ACCCAGCTATAAATATGCTGAATACCACTGAAATAATAGCTAAAG
73665
Gallus_gallus                    73621 .............................................
73665

Anas_platyrhynchos_platyrhynchos 73666 CTAGCTAATAGTATTTGGCAGAAATCTGCAGACCAATTGGAATTG
73710
Gallus_gallus                    73666 .............................................
73710

Anas_platyrhynchos_platyrhynchos 73711 CATACCTATATCTTTATAAGGACTGATATCCTGGTCTTTAGAGAG
73755
Gallus_gallus                    73711 .............................................
73755

Anas_platyrhynchos_platyrhynchos 73756 AAGCATTTCTACAGAAGTGTTGGACAATGTCATCAGATTCCTAGC
73800
Gallus_gallus                    73756 .............................................
73800

Anas_platyrhynchos_platyrhynchos 73801 TATCGAAGGAAATATGCCTTAGTCTTTTAAGAAACCCATTTATAA
73845
Gallus_gallus                    73801 .............................................
73845

Anas_platyrhynchos_platyrhynchos 73846 CAGGAAGTATAAATGATTAGGTGTAGTAATCAGACAGGTAAGACA
73890
Gallus_gallus                    73846 .............................................
73890

Anas_platyrhynchos_platyrhynchos 73891 TCAGCTCATAGTTTTGAAAATGAATGATTTAAACATTTTTATACT
73935
Gallus_gallus                    73891 .............................................
73935

Anas_platyrhynchos_platyrhynchos 73936 ACCTTCATTCCTCCAGACTGGTAATCAACAGAAGCAAGTGTTTTA
73980
Gallus_gallus                    73936 .............................................
73980

Anas_platyrhynchos_platyrhynchos 73981 ATGAACTATGCTAGCAATGTAGCTCTGTCTCATATAAGTCTTACA
74025
Gallus_gallus                    73981 .............................................
74025

Anas_platyrhynchos_platyrhynchos 74026 ATAAAGCCACCTAATTCTTTAAATAACAGGTAGTTCTACCCCATA
74070
Gallus_gallus                    74026 .............................................
74070

Anas_platyrhynchos_platyrhynchos 74071 ACTGCCAAAAGGAAATTCACAATAACATGAAGTTCTCTGTTGAAA
74115
Gallus_gallus                    74071 .............................................
74115

```
Anas_platyrhynchos_platyrhynchos 74116 TTTGGAATGCTCCCAATACAAACCCTAAGATAAGACTTTATCTTC
74160
Gallus_gallus                    74116 .............................................
74160

Anas_platyrhynchos_platyrhynchos 74161 TCAATAAAATAAAATAAAAGTCCCAAGTATGGTTCTGTACAGAAT
74205
Gallus_gallus                    74161 .............................................
74205

Anas_platyrhynchos_platyrhynchos 74206 GCATCCGAACCCATTATGTTCATCAGATTGTTATGATAATAACCT
74250
Gallus_gallus                    74206 .............................................
74250

Anas_platyrhynchos_platyrhynchos 74251 GTTAGAGCAACGTAGGAATAATTGTAATACTGGTTTTTAATGCTT
74295
Gallus_gallus                    74251 .............................................
74295

Anas_platyrhynchos_platyrhynchos 74296 TTTCTTTTTTGTATAAATGAAAATAGTTACACAATTTTTCTACCT
74340
Gallus_gallus                    74296 .............................................
74340

Anas_platyrhynchos_platyrhynchos 74341 GTTGTAATTACTCAATCTCTTGAAAACTTTGAAAGTAAATAATAA
74385
Gallus_gallus                    74341 .............................................
74385

Anas_platyrhynchos_platyrhynchos 74386 AAAGTTTCAACATAATATGTATAAGTAATATATATTATGTATGTA
74430
Gallus_gallus                    74386 .............................................
74430

Anas_platyrhynchos_platyrhynchos 74431 CTTCATATATATAACAATATAATAGAAGTTTCAAATTAAATTTTG
74475
Gallus_gallus                    74431 .............................................
74475

Anas_platyrhynchos_platyrhynchos 74476 TCTGTATTCAAAACCAAAACTGAAATCACATGCTCTTTCTCAAAA
74520
Gallus_gallus                    74476 .............................................
74520

Anas_platyrhynchos_platyrhynchos 74521 AACAGAGCTAATAAAATAAAAGGAAATACCTGAGATAAAGCAGGA
74565
Gallus_gallus                    74521 .............................................
74565

Anas_platyrhynchos_platyrhynchos 74566 ACTGTGAAATCCATGAGACCCAAGCCAGTACATGAATACATTTCT
74610
Gallus_gallus                    74566 .............................................
74610

Anas_platyrhynchos_platyrhynchos 74611 AGACCAACAAGAATTCTTGCAATAGGATTTTGTGCTTCATCACTA
74655
Gallus_gallus                    74611 .............................................
74655

Anas_platyrhynchos_platyrhynchos 74656 CTCTGAGCAGAAATTAAAATTTATTTATTTATTTATTTATTTTTA
74700
```

```
Gallus_gallus                       74656 .............................................
74700

Anas_platyrhynchos_platyrhynchos 74701 CTACCATTGTTAATGCTGAACTACTACAGTATCACCAAAAATACA
74745
Gallus_gallus                       74701 .............................................
74745

Anas_platyrhynchos_platyrhynchos 74746 GCAACAATATAATTAATTCACATTCCATACTCTGGTATTATAGAC
74790
Gallus_gallus                       74746 .............................................
74790

Anas_platyrhynchos_platyrhynchos 74791 ATATTTAAAATTAAAAGATTGGAATAAAATATTATTTTTCAAGTT
74835
Gallus_gallus                       74791 .............................................
74835

Anas_platyrhynchos_platyrhynchos 74836 ACTGAATCGACAGTCTCCATATAAAGAGTTCATACTAAATGTTAA
74880
Gallus_gallus                       74836 .............................................
74880

Anas_platyrhynchos_platyrhynchos 74881 TATAAACATAGCTGATATTTCAGCTACAGGCAAAAATTTTCTGTC
74925
Gallus_gallus                       74881 .............................................
74925

Anas_platyrhynchos_platyrhynchos 74926 ACATCAAATTAAAATATATACAAGAAATACACACTGATAATTTAA
74970
Gallus_gallus                       74926 .............................................
74970

Anas_platyrhynchos_platyrhynchos 74971 TCATTATTAAATAAATTAAGCATTTAATATTAATTTTTCAAAGAA
75015
Gallus_gallus                       74971 .............................................
75015

Anas_platyrhynchos_platyrhynchos 75016 AAAATTAACTTATTGTAGGATTTCAGAGAAATAATACCTCCTCCT
75060
Gallus_gallus                       75016 .............................................
75060

Anas_platyrhynchos_platyrhynchos 75061 CAGTCCAAAGTTTATGTCCTATCTGTTGCTGATTTAACTGTTTTG
75105
Gallus_gallus                       75061 .............................................
75105

Anas_platyrhynchos_platyrhynchos 75106 CTTTCACCCATGTAGCAATGATCTGTTGTTTTACAGCAACAGGTA
75150
Gallus_gallus                       75106 .............................................
75150

Anas_platyrhynchos_platyrhynchos 75151 CTATTTCATTAGGTATGGTTCCACTTGTCACATTTAGAGCAAATT
75195
Gallus_gallus                       75151 .............................................
75195

Anas_platyrhynchos_platyrhynchos 75196 CACAAATCTAAACAGCAAGAAGAAAAAAAGACTGAAACTTAAGGC
75240
Gallus_gallus                       75196 .............................................
75240
```

```
Anas_platyrhynchos_platyrhynchos 75241 AAATTTGAACAGCTCTTTTTATGGAAAATGTTATCCATTAGTTAT
75285
Gallus_gallus                    75241 .............................................
75285

Anas_platyrhynchos_platyrhynchos 75286 TCTACAATTTCTGTGCATGCATCTCTAATACTAATAACATCTCCA
75330
Gallus_gallus                    75286 .............................................
75330

Anas_platyrhynchos_platyrhynchos 75331 TAAATAAATCCATACAGCGAATAAAAGCTGCAGTCTTCGGATTCA
75375
Gallus_gallus                    75331 .............................................
75375

Anas_platyrhynchos_platyrhynchos 75376 ATTGACAAGGAACTTCAGTGCATATTTGGGGAGAACCTGTGATGG
75420
Gallus_gallus                    75376 .............................................
75420

Anas_platyrhynchos_platyrhynchos 75421 CAGGAACCATAAAAGATGGTAGTTTTTCTGCAAGAATCTCTGGAA
75465
Gallus_gallus                    75421 .............................................
75465

Anas_platyrhynchos_platyrhynchos 75466 TTTAAATTTCTATGAATCATAATACATGTCAGCATATTTAAAAAA
75510
Gallus_gallus                    75466 .............................................
75510

Anas_platyrhynchos_platyrhynchos 75511 AAAAAAAAAAAGACAAAGTGGATTTTGTACCTCTAAGGCTGCTTT
75555
Gallus_gallus                    75511 .............................................
75555

Anas_platyrhynchos_platyrhynchos 75556 GATGTCAGGAAGTCCACTGGGATCTACTGACAAAGAAGGGATAGC
75600
Gallus_gallus                    75556 .............................................
75600

Anas_platyrhynchos_platyrhynchos 75601 ATTTGCTTTCTCATGTCCCTTTGCTGAAGCCTTTCTACTTTTGTC
75645
Gallus_gallus                    75601 .............................................
75645

Anas_platyrhynchos_platyrhynchos 75646 TGGTGATGCACATAACTTTTTCTCCTCAGTACGTTTTGGTTTTGG
75690
Gallus_gallus                    75646 .............................................
75690

Anas_platyrhynchos_platyrhynchos 75691 AATCCAACAGAGAATTAAGCCCCGAGCTACAGCATTACATAACAT
75735
Gallus_gallus                    75691 .............................................
75735

Anas_platyrhynchos_platyrhynchos 75736 CAACAAAGTGACAGTGCTTCTGAAGAAGAAAAAAGAAAACTTAAA
75780
Gallus_gallus                    75736 .............................................
75780

Anas_platyrhynchos_platyrhynchos 75781 TGCAAACTGCATCAGACCAAAATAAGTGAAACAAATAAAGTCATG
75825
Gallus_gallus                    75781 .............................................
```

75825

Anas_platyrhynchos_platyrhynchos 75826 AGCACTATTCAAAAATCTAGATTTTTCTTAGAAAACAAAATTATG
75870
Gallus_gallus 75826 .............................................
75870

Anas_platyrhynchos_platyrhynchos 75871 GATTTTTCAAAATTTCCCTGTTGGAAATGCAGTTAAAATGGTTAA
75915
Gallus_gallus 75871 .............................................
75915

Anas_platyrhynchos_platyrhynchos 75916 ATCTACTCTCCCTCCTCCCCCTCTCTCCAATAAAAACTTTCATAT
75960
Gallus_gallus 75916 .............................................
75960

Anas_platyrhynchos_platyrhynchos 75961 TTTATATTGCATTTTTTCAGGTCTTTCTGTGATTGAAGCTTCTAT
76005
Gallus_gallus 75961 .............................................
76005

Anas_platyrhynchos_platyrhynchos 76006 TTCTTCACAGAAAAAGGTATCTTTTCAAATTTATTTCAGAGGTGA
76050
Gallus_gallus 76006 .............................................
76050

Anas_platyrhynchos_platyrhynchos 76051 ACAAATTTAAAATCTCACTTTTTAGTAGTGAACTGTTACGTGTGA
76095
Gallus_gallus 76051 .............................................
76095

Anas_platyrhynchos_platyrhynchos 76096 AAGTGGAAAAGGTTTCCTAATAACCTACACCAAACTCTTTACATA
76140
Gallus_gallus 76096 .............................................
76140

Anas_platyrhynchos_platyrhynchos 76141 AAATCATTTGTTACTCTGTATTTTTTTATATGATAATTATTTTTT
76185
Gallus_gallus 76141 .............................................
76185

Anas_platyrhynchos_platyrhynchos 76186 TTGATTGATTTTTTTTTTTTTTTTTTACAGACAACACAAACTGAAT
76230
Gallus_gallus 76186 .............................................
76230

Anas_platyrhynchos_platyrhynchos 76231 TATGATACATATTAATATAAGTGCCCGGATTAATAAGTAGGTATT
76275
Gallus_gallus 76231 .............................................
76275

Anas_platyrhynchos_platyrhynchos 76276 ACCTGTTTTCATCTCTGTACAAACTGAACTTTAGGCCTTGACATG
76320
Gallus_gallus 76276 .............................................
76320

Anas_platyrhynchos_platyrhynchos 76321 CATGTTAAAAAAATCATTCTTTGCTGTTGGTCCTCTCAGTTTGTC
76365
Gallus_gallus 76321 .............................................
76365

Anas_platyrhynchos_platyrhynchos 76366 TAGTTCTGAAGTCACTGCCAGTAAAAAGTGCTAAAAGTTCTAGGG

76410
Gallus_gallus 76366 .............................................
76410

Anas_platyrhynchos_platyrhynchos 76411 TTTTGTCCGACAGAACTTTGACTTGAGTATTCCTCTGGAATAGTT
76455
Gallus_gallus 76411 .............................................
76455

Anas_platyrhynchos_platyrhynchos 76456 AATTATTCTGCCATTGCATTTGCTACAGAACAGAATTCCTCTGTG
76500
Gallus_gallus 76456 .............................................
76500

Anas_platyrhynchos_platyrhynchos 76501 AAGTCTATTTTTGTGGTCAATTATTGAACACATTCCCTTTACTTT
76545
Gallus_gallus 76501 .............................................
76545

Anas_platyrhynchos_platyrhynchos 76546 CTTTTTTCCTCCTTTGTTTTGTATTTCAGAACTTACTGTGATTAG
76590
Gallus_gallus 76546 .............................................
76590

Anas_platyrhynchos_platyrhynchos 76591 AAATATTTACAATAAACTATATTTACTATATAATTCAACTTCAGA
76635
Gallus_gallus 76591 .............................................
76635

Anas_platyrhynchos_platyrhynchos 76636 GTAACTGAGAGCAAGTAAACACATTTCAAGCTCCTGCCAACAGAA
76680
Gallus_gallus 76636 .............................................
76680

Anas_platyrhynchos_platyrhynchos 76681 AGGCTACAAAAATTTTCTTGATTCTTTCTGAGCAGGTCAACAAAA
76725
Gallus_gallus 76681 .............................................
76725

Anas_platyrhynchos_platyrhynchos 76726 TTGACACATGCACTATTTCACATGCACATGTTGCTATTCCATAAT
76770
Gallus_gallus 76726 .............................................
76770

Anas_platyrhynchos_platyrhynchos 76771 TAAGAGGAATTAACATAATTTTAACTCAGGTTAATATGGATTATA
76815
Gallus_gallus 76771 .............................................
76815

Anas_platyrhynchos_platyrhynchos 76816 ATGAACAGAAATAAATAGGATTAAAACCTCACTTTCCAACCACAA
76860
Gallus_gallus 76816 .............................................
76860

Anas_platyrhynchos_platyrhynchos 76861 AATTTAATATAACTTCTAAAGTATTCTGTATATGCTGACGTCCTT
76905
Gallus_gallus 76861 .............................................
76905

Anas_platyrhynchos_platyrhynchos 76906 CTACTTCTAATAAAAGCCAGCAATAACATTCTTTCATGACATAAA
76950
Gallus_gallus 76906 .............................................
76950

```
Anas_platyrhynchos_platyrhynchos 76951 CTAATATTGTGAAAATATTTTTTCTGTTATAGAAACAAGACCATG
76995
Gallus_gallus                    76951 .............................................
76995

Anas_platyrhynchos_platyrhynchos 76996 TTTGTAACCAAAACTCTAAATTTACCCAGTGTGCCCAACAGTCTT
77040
Gallus_gallus                    76996 .............................................
77040

Anas_platyrhynchos_platyrhynchos 77041 GACTGCTCCAAGAAGAGTCTGCAGATACGCAATAATTTCCCTCTG
77085
Gallus_gallus                    77041 .............................................
77085

Anas_platyrhynchos_platyrhynchos 77086 TCTTCCTGATGTGATAAGATGCCTATTGTAGTTTAAGATATACAC
77130
Gallus_gallus                    77086 .............................................
77130

Anas_platyrhynchos_platyrhynchos 77131 CACAGCATTATGAACAATCCAGGCTTCATTCAGTTCTTCTCCTAT
77175
Gallus_gallus                    77131 .............................................
77175

Anas_platyrhynchos_platyrhynchos 77176 TTCAGCTGCACGCAGGAAATTTTCCATAGCATACCGAGATAAGTA
77220
Gallus_gallus                    77176 .............................................
77220

Anas_platyrhynchos_platyrhynchos 77221 GTCTATCCATGAACTATAAAATTAGTATTAAGAAACTGTTATTCC
77265
Gallus_gallus                    77221 .............................................
77265

Anas_platyrhynchos_platyrhynchos 77266 AAAAGTACAACACTAGAAGGAATTATATCTTAAGGCTAAATATGT
77310
Gallus_gallus                    77266 .............................................
77310

Anas_platyrhynchos_platyrhynchos 77311 AAAAGTAGAACTGAATCAACAAAAAATATATATTTTTGGAAATTA
77355
Gallus_gallus                    77311 .............................................
77355

Anas_platyrhynchos_platyrhynchos 77356 CTTTTCAGTTGCATTTATTACATCTCTATCCATTAACTTATAATT
77400
Gallus_gallus                    77356 .............................................
77400

Anas_platyrhynchos_platyrhynchos 77401 GGATGACAAGATGTCTCAGGAAAGTTATTAGTATCAACATCGACC
77445
Gallus_gallus                    77401 .............................................
77445

Anas_platyrhynchos_platyrhynchos 77446 AGTCTCTGATACTCAAATGCTATTAGCTTAAAAATATTGGCTCTA
77490
Gallus_gallus                    77446 .............................................
77490

Anas_platyrhynchos_platyrhynchos 77491 ACGTAAATTTCACAGTTCATCCACACTTTTTAAACTGTGTGTGAA
77535
```

```
Gallus_gallus                     77491 .............................................
77535

Anas_platyrhynchos_platyrhynchos 77536 TCACTGCTAGAAATCATTAAATATCTATGGCAAAGCTCTACTTGA
77580
Gallus_gallus                     77536 .............................................
77580

Anas_platyrhynchos_platyrhynchos 77581 ATAAAAAGTAAGACATTCCACTCTAGTGTTGAATGAGATATCTTT
77625
Gallus_gallus                     77581 .............................................
77625

Anas_platyrhynchos_platyrhynchos 77626 TTTACAACCTCATGTCTTCTTGCACTCAAAGACATCTCCAGATGT
77670
Gallus_gallus                     77626 .............................................
77670

Anas_platyrhynchos_platyrhynchos 77671 TCTGAAATGCTAAGTCCATTTTTTATAAAATAACATTCCATAGTC
77715
Gallus_gallus                     77671 .............................................
77715

Anas_platyrhynchos_platyrhynchos 77716 AACACAATAACATTCTTTTATGTACAAATTAATTATCAAGATATA
77760
Gallus_gallus                     77716 .............................................
77760

Anas_platyrhynchos_platyrhynchos 77761 TATCACACTAGATAAAGCAATACTGAAAACATCATCTTGTATGAT
77805
Gallus_gallus                     77761 .............................................
77805

Anas_platyrhynchos_platyrhynchos 77806 CAGACAGAATGACCTTGGTTAAGTCACAAATTCAAAAAAACATAA
77850
Gallus_gallus                     77806 .............................................
77850

Anas_platyrhynchos_platyrhynchos 77851 ACTGACATAGGGTATTACTATGTCACCAACAGTAGACAACATAGT
77895
Gallus_gallus                     77851 .............................................
77895

Anas_platyrhynchos_platyrhynchos 77896 AGACCAAAGTGATTTTTTTCCCTATGTGTTCTTAGCCTAATTGGA
77940
Gallus_gallus                     77896 .............................................
77940

Anas_platyrhynchos_platyrhynchos 77941 TGAAAGAGACACTGGTACAAGTAAATTGGTTAAACTCAAAAACAG
77985
Gallus_gallus                     77941 .............................................
77985

Anas_platyrhynchos_platyrhynchos 77986 CTTCACAGCGAATTTTAGAACAGAATCTTCATCTTCTTTGTCCCT
78030
Gallus_gallus                     77986 .............................................
78030

Anas_platyrhynchos_platyrhynchos 78031 TGTTTCTGATTAGATTTTACTTTTTCATATTTTACTGAGCATTTG
78075
Gallus_gallus                     78031 .............................................
78075
```

```
Anas_platyrhynchos_platyrhynchos 78076 AAGCTGACTGCGAGAAACTTCACTTCATCTTCAGACTTAATGAAG
78120
Gallus_gallus                    78076 .............................................
78120

Anas_platyrhynchos_platyrhynchos 78121 AGCCTTAGGGACACAACTCTGATAACCTGCTCTTACAGGACATGA
78165
Gallus_gallus                    78121 .............................................
78165

Anas_platyrhynchos_platyrhynchos 78166 TGACCATTGTGACAACAGTTAATACTAGTACAAATGTAGTAGAGT
78210
Gallus_gallus                    78166 .............................................
78210

Anas_platyrhynchos_platyrhynchos 78211 TTAAATATCATATTTAAAATTCTCTCTCCCTCTTCCTTCACAATA
78255
Gallus_gallus                    78211 .............................................
78255

Anas_platyrhynchos_platyrhynchos 78256 AATAATACTGTGAGACCAAAAGAAAGAAACAGAAACATAAAAAGA
78300
Gallus_gallus                    78256 .............................................
78300

Anas_platyrhynchos_platyrhynchos 78301 AGGAGAAGGAGGAGAAGGAGAAGAAGGGTACACTAGCTTCTGATT
78345
Gallus_gallus                    78301 .............................................
78345

Anas_platyrhynchos_platyrhynchos 78346 TCAGAGAAATAAACATTAAATGAAAAACAAAACAAACATTGCAGA
78390
Gallus_gallus                    78346 .............................................
78390

Anas_platyrhynchos_platyrhynchos 78391 ATGAAAGCAAATTAGAGAAAAATTATTTCAAAATATTTTTTAGAT
78435
Gallus_gallus                    78391 .............................................
78435

Anas_platyrhynchos_platyrhynchos 78436 TCTTCTTTTCCCACTACCATTCCTAGCTAAACATATGAACTATCT
78480
Gallus_gallus                    78436 .............................................
78480

Anas_platyrhynchos_platyrhynchos 78481 GATAACAATCAGAGAATCATTAAAGTTGGAAAAGACCTTCGAGAT
78525
Gallus_gallus                    78481 .............................................
78525

Anas_platyrhynchos_platyrhynchos 78526 CATCTGGTCCAACCATCACCCTACTGCCAATGTCCCCAAGCACCA
78570
Gallus_gallus                    78526 .............................................
78570

Anas_platyrhynchos_platyrhynchos 78571 GGTCCAACCTTTCCTAGAACACCCCCAGGAACGATGATGTCGCCA
78615
Gallus_gallus                    78571 .............................................
78615

Anas_platyrhynchos_platyrhynchos 78616 CTTCCCTGGGTAACCTGTTTCAATGTCTAACTACTCTTTCTGAGA
78660
Gallus_gallus                    78616 .............................................
```

78660

Anas_platyrhynchos_platyrhynchos 78661 AGAAAAATGATTTGTACTAAAAAGAAATAACTTAGTAAATTATCA
78705
Gallus_gallus 78661 .............................................
78705

Anas_platyrhynchos_platyrhynchos 78706 TCTGTGTTTGATAATAAAACAATTTGTTAGAGTTGGTAAGTACCT
78750
Gallus_gallus 78706 .............................................
78750

Anas_platyrhynchos_platyrhynchos 78751 GTATGCTTTCCATTCTAGATCACTTTCAGGAAAATCTAAAACATA
78795
Gallus_gallus 78751 .............................................
78795

Anas_platyrhynchos_platyrhynchos 78796 TTCTGAAGGATGTTGGCTTTTATCTTCTGTAGGTACAGTGTGGTC
78840
Gallus_gallus 78796 .............................................
78840

Anas_platyrhynchos_platyrhynchos 78841 ATTCAGCTCAAGTCCCTGCAATCTTAATAAATGAATCGTTGCCTG
78885
Gallus_gallus 78841 .............................................
78885

Anas_platyrhynchos_platyrhynchos 78886 GAAAGATACAAAGAAATACATTTTCTACCTGTGTTATCTACGTGA
78930
Gallus_gallus 78886 .............................................
78930

Anas_platyrhynchos_platyrhynchos 78931 AAATGGACTCTGACTTCTTATTTGTGTCTACACCTCTTGAAAAAA
78975
Gallus_gallus 78931 .............................................
78975

Anas_platyrhynchos_platyrhynchos 78976 GAAAAAAAAAAAAAGTTTTTGTTGTACAACATCTGAAGATGTACA
79020
Gallus_gallus 78976 .............................................
79020

Anas_platyrhynchos_platyrhynchos 79021 GGCATACAAAACCAATTATATGAAATAATCTTCAAATTACTTATG
79065
Gallus_gallus 79021 .............................................
79065

Anas_platyrhynchos_platyrhynchos 79066 CTTAATTTTATTAAATAAATAACTCAAATGAGAAAAAGAAAATAT
79110
Gallus_gallus 79066 .............................................
79110

Anas_platyrhynchos_platyrhynchos 79111 GCCATATATATACACACAACTATATTTATCATGCTGGAACTTTAA
79155
Gallus_gallus 79111 .............................................
79155

Anas_platyrhynchos_platyrhynchos 79156 GGATCCTGGAACATAATGTCACATGTCAAATATTTACAACAACAT
79200
Gallus_gallus 79156 .............................................
79200

Anas_platyrhynchos_platyrhynchos 79201 CCCTTGTAAAAAAGTGTTAAAACCATATGCAGTGATACTATAGTT

```
79245
Gallus_gallus                     79201 .............................................
79245

Anas_platyrhynchos_platyrhynchos  79246 ACTTTCACACAGTATTTTTATACAGATGCTTCACAGAACTACTTC
79290
Gallus_gallus                     79246 .............................................
79290

Anas_platyrhynchos_platyrhynchos  79291 CTCTAATGTCTGTGGAGACATAATGAATCTAACTCAAAATTCTAC
79335
Gallus_gallus                     79291 .............................................
79335

Anas_platyrhynchos_platyrhynchos  79336 TGCTGTAAGATGAATATTCCTAGAATTAGACATACAACCAAAGCT
79380
Gallus_gallus                     79336 .............................................
79380

Anas_platyrhynchos_platyrhynchos  79381 AAATGAAACAAATATGAGTAACATCTGTGTAGGTTGATTACCTCT
79425
Gallus_gallus                     79381 .............................................
79425

Anas_platyrhynchos_platyrhynchos  79426 GCATTAATAAATCTTATTTCTGCTAATATTCTGAGTAAATCTCTT
79470
Gallus_gallus                     79426 .............................................
79470

Anas_platyrhynchos_platyrhynchos  79471 TCAAAAGAAAAGGACTTAGCAGGTAGTAATAACTCTCCTGGTAAG
79515
Gallus_gallus                     79471 .............................................
79515

Anas_platyrhynchos_platyrhynchos  79516 CCATCATCTTGGTCATCATCCATCATAGTTATACTAGCTTTCTTC
79560
Gallus_gallus                     79516 .............................................
79560

Anas_platyrhynchos_platyrhynchos  79561 TTCTGTGCTAAAAAAAAAAAAAGTTACATTATTATTAACACGTAA
79605
Gallus_gallus                     79561 .............................................
79605

Anas_platyrhynchos_platyrhynchos  79606 ATCTGACAAAAATTACCTTAATTTTAGAATTTAAATATCCCTATA
79650
Gallus_gallus                     79606 .............................................
79650

Anas_platyrhynchos_platyrhynchos  79651 TAATCCGAAATCCACAATAAAGCATTCACAAGCACAGATTTGAAG
79695
Gallus_gallus                     79651 .............................................
79695

Anas_platyrhynchos_platyrhynchos  79696 TTACTTCACTCAGACTACATCTCAAGTTTAATATTCAGTGATAGA
79740
Gallus_gallus                     79696 .............................................
79740

Anas_platyrhynchos_platyrhynchos  79741 TCTAATTTATTAGTCTTTGTCCAATAATTTTAATCATATTTCAAT
79785
Gallus_gallus                     79741 .............................................
79785
```

```
Anas_platyrhynchos_platyrhynchos 79786 ATTCCAAAGTAACATTTTAGTTTAAGTTGAAAATTACATGTTTTT
79830
Gallus_gallus                    79786 .............................................
79830

Anas_platyrhynchos_platyrhynchos 79831 GTTTTCTTAAAACCTACAGACTGTTAATTTGAATGAAGAGCTCCT
79875
Gallus_gallus                    79831 .............................................
79875

Anas_platyrhynchos_platyrhynchos 79876 CATTCCTTGAGCAATCATACATATTCACTTTCCCCAGACTAGCCG
79920
Gallus_gallus                    79876 .............................................
79920

Anas_platyrhynchos_platyrhynchos 79921 TGTTTCATCTCTCATCTCTTTTTCAAACTAAAGGAAAAAAAAAAA
79965
Gallus_gallus                    79921 .............................................
79965

Anas_platyrhynchos_platyrhynchos 79966 GTACTTATCAGGAACTCAGGGCAACCATGCACCAAAAGCATCATT
80010
Gallus_gallus                    79966 .............................................
80010

Anas_platyrhynchos_platyrhynchos 80011 AATAGGACCTGGCATATATCTGTAACACTATAGCACTTTGAATTA
80055
Gallus_gallus                    80011 .............................................
80055

Anas_platyrhynchos_platyrhynchos 80056 TTTGCTTTTATTCAGGCCTGATAATACAGCAAGTCTTTTTCCAGG
80100
Gallus_gallus                    80056 .............................................
80100

Anas_platyrhynchos_platyrhynchos 80101 ATCTTCATATTGCCCAAATTATCTTTGCCCTATCTTCAGTATATG
80145
Gallus_gallus                    80101 .............................................
80145

Anas_platyrhynchos_platyrhynchos 80146 TCACATACAGAGAAGACCTTCAATTTTTCTCACCCACTATTCCTA
80190
Gallus_gallus                    80146 .............................................
80190

Anas_platyrhynchos_platyrhynchos 80191 GGATTAGAAGTAGTAAAGGATGAAACATAACACAAAAATGAATGT
80235
Gallus_gallus                    80191 .............................................
80235

Anas_platyrhynchos_platyrhynchos 80236 TGCCAACAGTCCAAGTGCAAGGTGCTGCACCTGGGTCAGGGCAAT
80280
Gallus_gallus                    80236 .............................................
80280

Anas_platyrhynchos_platyrhynchos 80281 CCCAGATATGAGGGCAGACTGGGAGAAGAACTCATTGAGAGCAGC
80325
Gallus_gallus                    80281 .............................................
80325

Anas_platyrhynchos_platyrhynchos 80326 CCTGCAGAGGACTTGGGAGTTCTGGTGAAGAAAAGGTTCAACATG
80370
```

```
Gallus_gallus                    80326 .............................................
80370

Anas_platyrhynchos_platyrhynchos 80371 AGCCAGCAGTGTGCACTTGCAGCCCAGAAGGCTGTGTGTGACCAG
80415
Gallus_gallus                    80371 .............................................
80415

Anas_platyrhynchos_platyrhynchos 80416 CAGGTCGATGGAGGTGATTGCCTCCCTCTACTCTGCCCTTGTGAA
80460
Gallus_gallus                    80416 .............................................
80460

Anas_platyrhynchos_platyrhynchos 80461 GCCCCACTTGAAGTACTATCTCCAGACCTGGAGCCCCAAGCACAA
80505
Gallus_gallus                    80461 .............................................
80505

Anas_platyrhynchos_platyrhynchos 80506 GAAGGATGTGGACCTGTTAGAGAAGGTCTGGAAAAGGGCCACAAA
80550
Gallus_gallus                    80506 .............................................
80550

Anas_platyrhynchos_platyrhynchos 80551 GTTGAACGGAGGACTGGAGCACCTCTCCAATGAAAAAAAGGCTGG
80595
Gallus_gallus                    80551 .............................................
80595

Anas_platyrhynchos_platyrhynchos 80596 AAGACCTGGGGCTCTTCATCCTGAAGAGGAGAAGGCTCCGTGGAG
80640
Gallus_gallus                    80596 .............................................
80640

Anas_platyrhynchos_platyrhynchos 80641 ACTTCATTGCGACCTTTCTATACTTAAAGAGGTCTTATAAAAAAG
80685
Gallus_gallus                    80641 .............................................
80685

Anas_platyrhynchos_platyrhynchos 80686 ATGGAGAAGGACTCTATTTGGGTAGACAATCCCCTCCCTCACCTC
80730
Gallus_gallus                    80686 .............................................
80730

Anas_platyrhynchos_platyrhynchos 80731 TTTTGATGCAGCCCAGGATGCAGTTGGAATTCTGGGCTGCAAGCA
80775
Gallus_gallus                    80731 .............................................
80775

Anas_platyrhynchos_platyrhynchos 80776 CAAACTGCTGGCTCATGTCAAGCTTCATCCACCAGAACACCCAAG
80820
Gallus_gallus                    80776 .............................................
80820

Anas_platyrhynchos_platyrhynchos 80821 TCTTTTCTCTGCTAGGCTGCTCTCAATGAGTTCATCCAGTCTGTA
80865
Gallus_gallus                    80821 .............................................
80865

Anas_platyrhynchos_platyrhynchos 80866 CTCCTGTCCGGGATTGCCCCAACCCGGGTGCAGCACCTTGCACTC
80910
Gallus_gallus                    80866 .............................................
80910
```

```
Anas_platyrhynchos_platyrhynchos 80911 AGACTTGTTGAACCTCCTCAGGTTCACATGGGCCCACTGTTCTAG
80955
Gallus_gallus                    80911 .............................................
80955

Anas_platyrhynchos_platyrhynchos 80956 CTTGTCCAGCTCCCTTTAGCTGGCATCCCTTCCTTTTGTTGTATT
81000
Gallus_gallus                    80956 .............................................
81000

Anas_platyrhynchos_platyrhynchos 81001 AACTGCACCACTCCACTTGCATCTGCAAACTTGCTGAGGAACAAC
81045
Gallus_gallus                    81001 .............................................
81045

Anas_platyrhynchos_platyrhynchos 81046 TAAGAACACTGATAATGATCTCAAGAACACTAAATAATGATCTTA
81090
Gallus_gallus                    81046 .............................................
81090

Anas_platyrhynchos_platyrhynchos 81091 AAGGTCTTTTCCAACCTAAACGATTCCACGATTCTATGCTTCTAA
81135
Gallus_gallus                    81091 .............................................
81135

Anas_platyrhynchos_platyrhynchos 81136 ATATACATACTTTAACTAGCTCAGGCCCAAAGCTTATATTTAATA
81180
Gallus_gallus                    81136 .............................................
81180

Anas_platyrhynchos_platyrhynchos 81181 GCATCACCTCATAGCACTGTCATTGAATTATCATGAAAAGTGAAA
81225
Gallus_gallus                    81181 .............................................
81225

Anas_platyrhynchos_platyrhynchos 81226 ATAATTTCCAAACCTCTCCTTCCTAAGCTTTTCTGCAGTTGTTAA
81270
Gallus_gallus                    81226 .............................................
81270

Anas_platyrhynchos_platyrhynchos 81271 TTTCTGAGAGCACACATTGCCTTCTACATTTCTAAACTTTTTGAA
81315
Gallus_gallus                    81271 .............................................
81315

Anas_platyrhynchos_platyrhynchos 81316 AACAATGTGCACAATATTCATTGGAACACCATACCTTTACACTGG
81360
Gallus_gallus                    81316 .............................................
81360

Anas_platyrhynchos_platyrhynchos 81361 CTTACTCCTTCAAGTGACTCTAATAAAGGAGGAGAAAGTTCACTT
81405
Gallus_gallus                    81361 .............................................
81405

Anas_platyrhynchos_platyrhynchos 81406 TATGACACTCAAAATAATGTTTTCACTTACTTTTTTTAGGTTTTG
81450
Gallus_gallus                    81406 .............................................
81450

Anas_platyrhynchos_platyrhynchos 81451 TTGCTCTCCTAAACAAAGTGTCATCATACAGGAGACAAAATCTGC
81495
Gallus_gallus                    81451 .............................................
```

```
81495

Anas_platyrhynchos_platyrhynchos 81496 ATGCTGTACGGCAGACATCCCACACTTCTTGTGTACGTGCAACTT
81540
Gallus_gallus                    81496 .............................................
81540

Anas_platyrhynchos_platyrhynchos 81541 TTGCCAAATTTGCCCAAAGTCTAATTCTATCAAAGAAAAAGCATA
81585
Gallus_gallus                    81541 .............................................
81585

Anas_platyrhynchos_platyrhynchos 81586 TGTAAGATTACTTTATATCCAGAATTATCTTCCACACTTTAGAAA
81630
Gallus_gallus                    81586 .............................................
81630

Anas_platyrhynchos_platyrhynchos 81631 CCACAAAAATTATGTCAATTTTGATGGCCACCACAAATAAGATTT
81675
Gallus_gallus                    81631 .............................................
81675

Anas_platyrhynchos_platyrhynchos 81676 TTAATCTAGTAAACATTACTACGTGGAAGAGTTTTTATCAGTAAC
81720
Gallus_gallus                    81676 .............................................
81720

Anas_platyrhynchos_platyrhynchos 81721 AAAGATAACACTATTACTTTACTCACTTAAAAAAAATACCCTCAG
81765
Gallus_gallus                    81721 .............................................
81765

Anas_platyrhynchos_platyrhynchos 81766 CTTATCAAGATTGCAAGAAAAAAATATTTTGGAGAATAGCTGTACT
81810
Gallus_gallus                    81766 .............................................
81810

Anas_platyrhynchos_platyrhynchos 81811 GAAACTTTTAAAATGTGTTTGTGACATATCCCTAATTATCTATCT
81855
Gallus_gallus                    81811 .............................................
81855

Anas_platyrhynchos_platyrhynchos 81856 TATAAAAAATAAACCAAATTAAACCAAGTATTTCAAAAGCAATGC
81900
Gallus_gallus                    81856 .............................................
81900

Anas_platyrhynchos_platyrhynchos 81901 TGAGAATTTCAAAATGAGTCTTTAATACAAGTTTGGATTTTGAAT
81945
Gallus_gallus                    81901 .............................................
81945

Anas_platyrhynchos_platyrhynchos 81946 TGTTTAGAAAAACACTGTTTAGATTTCATTCCAGGATTTTTTTTA
81990
Gallus_gallus                    81946 .............................................
81990

Anas_platyrhynchos_platyrhynchos 81991 ATGCTGCAAACAAAACAAAACAAATAAAGAAAAACAAACAACAAC
82035
Gallus_gallus                    81991 .............................................
82035

Anas_platyrhynchos_platyrhynchos 82036 AAAATTGTTTATAAAATAAACAATAAAATTATATAAACATGCTGA
```

```
82080
Gallus_gallus                     82036 .............................................
82080

Anas_platyrhynchos_platyrhynchos 82081 AAACAAAGTCACACAGAATCACAGAAAGGTTTAAGTTGGAAGAGA
82125
Gallus_gallus                     82081 .............................................
82125

Anas_platyrhynchos_platyrhynchos 82126 CCTCTGGAAGTTACCTGGTCCAATCCTCATGCTCTAGCAGGGACA
82170
Gallus_gallus                     82126 .............................................
82170

Anas_platyrhynchos_platyrhynchos 82171 AAGCTATGATTTTCCTACAGATCAGTATTTCTGACACTCAGAAAG
82215
Gallus_gallus                     82171 .............................................
82215

Anas_platyrhynchos_platyrhynchos 82216 GACTGTGTGGTTTATCAGAAAATATGTAGCATGCAAAGTGGTACA
82260
Gallus_gallus                     82216 .............................................
82260

Anas_platyrhynchos_platyrhynchos 82261 AGAACCAAAAAAGACGCTTCAGCAAATTTTCAGCTATGCATCAAG
82305
Gallus_gallus                     82261 .............................................
82305

Anas_platyrhynchos_platyrhynchos 82306 TTATTCAGCCTTAATCCACCCATTCTTCTGTGTGTCCAGTTTCTA
82350
Gallus_gallus                     82306 .............................................
82350

Anas_platyrhynchos_platyrhynchos 82351 TCAACAGAATAGGTGGTAGGGGGAAGGTTGCACCAGATGATATTG
82395
Gallus_gallus                     82351 .............................................
82395

Anas_platyrhynchos_platyrhynchos 82396 GAGGTCTTTTCCAACCTTAATGATTCTATGATAAGAAATTCCCTT
82440
Gallus_gallus                     82396 .............................................
82440

Anas_platyrhynchos_platyrhynchos 82441 ATTAGACACTTCTACGCAAAAAGTGGTAATTTCTGTGGTAATTAC
82485
Gallus_gallus                     82441 .............................................
82485

Anas_platyrhynchos_platyrhynchos 82486 CTCTCTTTGTCATTTTCCTTTTTTAGGTGTTTCAAATGTCCATCA
82530
Gallus_gallus                     82486 .............................................
82530

Anas_platyrhynchos_platyrhynchos 82531 ACTTTGTCCACACTTTTAACATGATGTTGTGCTTTTGCACAGAGG
82575
Gallus_gallus                     82531 .............................................
82575

Anas_platyrhynchos_platyrhynchos 82576 TAACTTACTTGACTCTGACTATAACCTGAGGACACTTACAAAGAA
82620
Gallus_gallus                     82576 .............................................
82620
```

```
Anas_platyrhynchos_platyrhynchos 82621 ATAAGAATTTAATGTAATTTTTTAAGTACCAGAGTTGAAACAAAT
82665
Gallus_gallus                    82621 .............................................
82665

Anas_platyrhynchos_platyrhynchos 82666 CTAAATGAAAGGTATGCATTTTGATTCACAACTACTTTCTGAAAC
82710
Gallus_gallus                    82666 .............................................
82710

Anas_platyrhynchos_platyrhynchos 82711 TCCTGAAATCCAGTCCTTTTTTCAATCCCTCAATCTTTCCCAACG
82755
Gallus_gallus                    82711 .............................................
82755

Anas_platyrhynchos_platyrhynchos 82756 CCAATCAAAATGAAAAAAACACACCCGACTAGCATCTATTCTGTT
82800
Gallus_gallus                    82756 .............................................
82800

Anas_platyrhynchos_platyrhynchos 82801 TTTAATTTATCTGCATAGATAGTCTCCCCATATAAAAATGGATTT
82845
Gallus_gallus                    82801 .............................................
82845

Anas_platyrhynchos_platyrhynchos 82846 ATGTTTTTTTAGCCCCAAGGTCTCTTTTCTGACAGGACATGCATT
82890
Gallus_gallus                    82846 .............................................
82890

Anas_platyrhynchos_platyrhynchos 82891 GCTCTAATTAAGCTTCAGTCAAAAGGAAAGTGATGTAATATATGG
82935
Gallus_gallus                    82891 .............................................
82935

Anas_platyrhynchos_platyrhynchos 82936 AATGTTGCATTCTGGCTGCATTTTAAGAACACTAATCCAATGAAA
82980
Gallus_gallus                    82936 .............................................
82980

Anas_platyrhynchos_platyrhynchos 82981 TACGTTTTTGCTCAGTAGCAAAATTTGTTCTTTGTGTTTTTGTTT
83025
Gallus_gallus                    82981 .............................................
83025

Anas_platyrhynchos_platyrhynchos 83026 TTTTTTGTTTAAATTTGAGAATAATGACTTTTTTTCACGTAGTCA
83070
Gallus_gallus                    83026 .............................................
83070

Anas_platyrhynchos_platyrhynchos 83071 ACCAGATATAGTTATCTAGGAAAAAGTATTAAACAGAAATAACCA
83115
Gallus_gallus                    83071 .............................................
83115

Anas_platyrhynchos_platyrhynchos 83116 ACATAGAAAAATGACACAAGGGAGGAATGAGCTGCCCTCCTGAGG
83160
Gallus_gallus                    83116 .............................................
83160

Anas_platyrhynchos_platyrhynchos 83161 TAGAAAATTACTCCAGTGATACAGAAAAATCAAAAAGAGATTTTT
83205
```

```
Gallus_gallus                     83161 .............................................
83205

Anas_platyrhynchos_platyrhynchos 83206 TTTTACTGTAATGACATAGAGGAGATGCTATTTATTAGAATACAA
83250
Gallus_gallus                     83206 .............................................
83250

Anas_platyrhynchos_platyrhynchos 83251 ATAACTTGATATGATTATTTGGGAAATCAATATTCACTTGAATCT
83295
Gallus_gallus                     83251 .............................................
83295

Anas_platyrhynchos_platyrhynchos 83296 AAGCCATGGTTGCTAATATTACTTAGACTAAGTTTTCTGTTCCAA
83340
Gallus_gallus                     83296 .............................................
83340

Anas_platyrhynchos_platyrhynchos 83341 ATGAATACTGGCTGGACTGCTGAGATCAAAGGGTATTTGCAAATC
83385
Gallus_gallus                     83341 .............................................
83385

Anas_platyrhynchos_platyrhynchos 83386 GAGTTAAATCCAGCTGGCAGCCAAGTGGCCATAGTGGCGTTCCCC
83430
Gallus_gallus                     83386 .............................................
83430

Anas_platyrhynchos_platyrhynchos 83431 AGGGGTCCGCATTAGGGCCAGTTTTGTTTAATCAATGATCTTGAT
83475
Gallus_gallus                     83431 .............................................
83475

Anas_platyrhynchos_platyrhynchos 83476 GAGGGAATCATTTAATCTTGATGAGGGATAAAATTGTAGACAACA
83520
Gallus_gallus                     83476 .............................................
83520

Anas_platyrhynchos_platyrhynchos 83521 CCAAGTTGGGCAGGAATGATGATCTGTGAGAAGGTAGGCAGGCTC
83565
Gallus_gallus                     83521 .............................................
83565

Anas_platyrhynchos_platyrhynchos 83566 TGCAGATGGGTTTGGGTAAGCTAGATTTATGGACCACATCCATTT
83610
Gallus_gallus                     83566 .............................................
83610

Anas_platyrhynchos_platyrhynchos 83611 GTATGGCATGCAACAAGGCTAAATGCCAGGTGATGCACTTGGGTC
83655
Gallus_gallus                     83611 .............................................
83655

Anas_platyrhynchos_platyrhynchos 83656 ATTACAACCCCATGCAGTGGTATAGGCCTGGGGAAGAGTAGCTGG
83700
Gallus_gallus                     83656 .............................................
83700

Anas_platyrhynchos_platyrhynchos 83701 AAAGCTGCCTGGAAGAAAAGGAGTTGGGGATGTTGTGTTGATCAA
83745
Gallus_gallus                     83701 .............................................
83745
```

```
Anas_platyrhynchos_platyrhynchos 83746 TAGCCAGTTGCACATGAACCAGCTGTGTGCTCAGGTGGCCAAGAA
83790
Gallus_gallus                    83746 .............................................
83790

Anas_platyrhynchos_platyrhynchos 83791 GACCAACAGCATCCTGGCTTGTATCAGAAATAGCATGGCCAGAGG
83835
Gallus_gallus                    83791 .............................................
83835

Anas_platyrhynchos_platyrhynchos 83836 TACTATGGAAGTGACTGTCCCTCTGTACTCTGTACTTGACACACC
83880
Gallus_gallus                    83836 .............................................
83880

Anas_platyrhynchos_platyrhynchos 83881 TTGGGCTGGGCTCCTCACTACATAAAGATATCGAGTTTCTGGAAT
83925
Gallus_gallus                    83881 .............................................
83925

Anas_platyrhynchos_platyrhynchos 83926 ACATTCATAGAAGAACAGCAAAGCTGCTGAAGGGACTAGAAAATA
83970
Gallus_gallus                    83926 .............................................
83970

Anas_platyrhynchos_platyrhynchos 83971 TTCCTGGCTTTAAAACGTGAATATAACTGAAACTGCTTAGACATA
84015
Gallus_gallus                    83971 .............................................
84015

Anas_platyrhynchos_platyrhynchos 84016 TTTACGTTGAAATGAGGAAGGTATAGTTTCTTATAATTTTTGTTT
84060
Gallus_gallus                    84016 .............................................
84060

Anas_platyrhynchos_platyrhynchos 84061 AAAATACTTTGCAATGCATATGACAGAACTGTTACTTAACTACAC
84105
Gallus_gallus                    84061 .............................................
84105

Anas_platyrhynchos_platyrhynchos 84106 ACCCAGATTGACAGACAGAAATTTTTCAGCATTAATACATGACAG
84150
Gallus_gallus                    84106 .............................................
84150

Anas_platyrhynchos_platyrhynchos 84151 CAATATCATAGAATTATATTCATTTTTCATTCTCTAATTTTTGCC
84195
Gallus_gallus                    84151 .............................................
84195

Anas_platyrhynchos_platyrhynchos 84196 ACAGAATTCAGTAAACTTGTCACTCTATTTGCGCTGTAATATTCC
84240
Gallus_gallus                    84196 .............................................
84240

Anas_platyrhynchos_platyrhynchos 84241 CCACATATTGCCACTATATGTTATGTCCGTTCTTTGAGGTTGGGC
84285
Gallus_gallus                    84241 .............................................
84285

Anas_platyrhynchos_platyrhynchos 84286 AAAGCAGTAAAAGTTAAGTGTCGTAATTTTAGCCACACACATACA
84330
Gallus_gallus                    84286 .............................................
```

```
84330

Anas_platyrhynchos_platyrhynchos 84331 TTACCTTTAGCTTCATTTTCACTATCAAGAACCACCTGAAATAAA
84375
Gallus_gallus                    84331 .............................................
84375

Anas_platyrhynchos_platyrhynchos 84376 TCAGGAGCTAGTGCAAGACCAGCATTTACCAGAAGGGATCTCTTT
84420
Gallus_gallus                    84376 .............................................
84420

Anas_platyrhynchos_platyrhynchos 84421 TGCCTCACATTATTCTTTTCCTTTCCCCTTTTAGCCTATGTAAAT
84465
Gallus_gallus                    84421 .............................................
84465

Anas_platyrhynchos_platyrhynchos 84466 AAAAGAACTAAATGTTACAAGGTTCATTTAGACCTATTAAAATGC
84510
Gallus_gallus                    84466 .............................................
84510

Anas_platyrhynchos_platyrhynchos 84511 TATTTACATTTTTATTACATTAATATATCTTACTTACTAAACATG
84555
Gallus_gallus                    84511 .............................................
84555

Anas_platyrhynchos_platyrhynchos 84556 TAAGAATGTAGATCATAGACTTTGGTATCTTGAGCCAATACAATA
84600
Gallus_gallus                    84556 .............................................
84600

Anas_platyrhynchos_platyrhynchos 84601 CAGGAATTGTTTATCTTAGCATATCTTAAACAACATCCTGCAGTA
84645
Gallus_gallus                    84601 .............................................
84645

Anas_platyrhynchos_platyrhynchos 84646 TTATTACTCTATATACCTTATTGTATGACCTGTCTTAATGTTGTA
84690
Gallus_gallus                    84646 .............................................
84690

Anas_platyrhynchos_platyrhynchos 84691 TTTCCTTAAATAAAGAATAGTATTTATTTAGAAACCAAAAATAAC
84735
Gallus_gallus                    84691 .............................................
84735

Anas_platyrhynchos_platyrhynchos 84736 ATACCTGTCGTGCAATAATTAGGACTCACTATGAAATATAAAAAA
84780
Gallus_gallus                    84736 .............................................
84780

Anas_platyrhynchos_platyrhynchos 84781 TCCCTAGTGTACCCAGAATTAGCTTAAGTCAGAGGAAAATTAAGT
84825
Gallus_gallus                    84781 .............................................
84825

Anas_platyrhynchos_platyrhynchos 84826 GCTTTCTTTAAAAAATATTTAAATGGAATAAGCTTGTGTGAAAAA
84870
Gallus_gallus                    84826 .............................................
84870

Anas_platyrhynchos_platyrhynchos 84871 AAAAATAATAAACTAGAATAAACTACACCAAAACTCTGTCATAAC
```

```
84915
Gallus_gallus                    84871 .............................................
84915

Anas_platyrhynchos_platyrhynchos 84916 AATAATATAACAATCATACCAGCAGCTTTCCAATACTGCTTTTTA
84960
Gallus_gallus                    84916 .............................................
84960

Anas_platyrhynchos_platyrhynchos 84961 GGTTCAATGAGCATATCTCAATTTCTGTGAATATCCCCAAATGGT
85005
Gallus_gallus                    84961 .............................................
85005

Anas_platyrhynchos_platyrhynchos 85006 GTATAGGTGAAATTTCTTTTAGCAAACTACCAAATCAGTTGCCAA
85050
Gallus_gallus                    85006 .............................................
85050

Anas_platyrhynchos_platyrhynchos 85051 TATGAATTTTGTTCATGTGCTGAAGGCATCACAACAATTATCTCA
85095
Gallus_gallus                    85051 .............................................
85095

Anas_platyrhynchos_platyrhynchos 85096 ATTTCTGTGAATATCCCCAAATGGTGTATAGGTGAAATTTCTTTT
85140
Gallus_gallus                    85096 .............................................
85140

Anas_platyrhynchos_platyrhynchos 85141 AGCAAACTACCAAATCAGCTGCCAATATGAATTTTGTTCATGTGC
85185
Gallus_gallus                    85141 .............................................
85185

Anas_platyrhynchos_platyrhynchos 85186 TGAAGGCATTGCAACAATTAGATGTAATAACTCTTGAAAATGACT
85230
Gallus_gallus                    85186 .............................................
85230

Anas_platyrhynchos_platyrhynchos 85231 ACACTGTGACTTTAGATTGTATTACTGGATTTATTATCTGTATTT
85275
Gallus_gallus                    85231 .............................................
85275

Anas_platyrhynchos_platyrhynchos 85276 TCTCACATTTTAAAGTGTGATATAATACTGATTTATCTTCAGTCC
85320
Gallus_gallus                    85276 .............................................
85320

Anas_platyrhynchos_platyrhynchos 85321 AGATGGGATATTCAGCTATTTGGCCACTGCTCCTCTCATTATTTA
85365
Gallus_gallus                    85321 .............................................
85365

Anas_platyrhynchos_platyrhynchos 85366 TAAAAATGTTCCACACAATGAAATTTTAATTGTTGACAATTCTGA
85410
Gallus_gallus                    85366 .............................................
85410

Anas_platyrhynchos_platyrhynchos 85411 ACATGCAAAACAGTAACACTGAGTTTCTTTTTTGCAGATGAGTAT
85455
Gallus_gallus                    85411 .............................................
85455
```

```
Anas_platyrhynchos_platyrhynchos 85456 TTCTGCCAAAGGATAATGCTTTTTCATTTTTTTCATAAACAGGTG
85500
Gallus_gallus                    85456 .............................................
85500

Anas_platyrhynchos_platyrhynchos 85501 GTTTGAACATAATTGGAAAAAATCCTCCATCATAACCATTAATCT
85545
Gallus_gallus                    85501 .............................................
85545

Anas_platyrhynchos_platyrhynchos 85546 TCAGAGTATGAAGCAGTGTGATGACCAGAGGGAAAATCCAGAGAG
85590
Gallus_gallus                    85546 .............................................
85590

Anas_platyrhynchos_platyrhynchos 85591 CTTTCAAGGCAACAAACTAATGTTGCTGAAAGCAAGCCCGTGTTA
85635
Gallus_gallus                    85591 .............................................
85635

Anas_platyrhynchos_platyrhynchos 85636 GCTCTACTAATATGCAGTTCTACTAAATGGCTAACTTTTCTCTGC
85680
Gallus_gallus                    85636 .............................................
85680

Anas_platyrhynchos_platyrhynchos 85681 CAGAATTTAATAAGCAAACCTATTTCAGCTACTAATCTTATTCTG
85725
Gallus_gallus                    85681 .............................................
85725

Anas_platyrhynchos_platyrhynchos 85726 TACGATTACCTGTTCAATCATCATTAGCGCCTGATCCTCTAGACG
85770
Gallus_gallus                    85726 .............................................
85770

Anas_platyrhynchos_platyrhynchos 85771 CTCAGGAGACTTATACAGCATAGCGTACAAAGAAAGACGGTTAAA
85815
Gallus_gallus                    85771 .............................................
85815

Anas_platyrhynchos_platyrhynchos 85816 ATTCAATCTTAGATGTTCTTGATACTGCCCACTGTTATCTAGATG
85860
Gallus_gallus                    85816 .............................................
85860

Anas_platyrhynchos_platyrhynchos 85861 TATAGCTTTCTGTAAATGTTGCACAGCAATCTCAATTCTATCCTC
85905
Gallus_gallus                    85861 .............................................
85905

Anas_platyrhynchos_platyrhynchos 85906 ATCTTCTTCAATACGAGCAATTTCCATGTGAACTTGGCAACGTAA
85950
Gallus_gallus                    85906 .............................................
85950

Anas_platyrhynchos_platyrhynchos 85951 AAAAATCAGCATGCTATGGAATAAAAATTCAAATGTAATCTGTTA
85995
Gallus_gallus                    85951 .............................................
85995

Anas_platyrhynchos_platyrhynchos 85996 AATGGAGCAAACTAAAATTGTTAGACCTAAAAATTAACCAAAAGC
86040
```

```
Gallus_gallus                      85996 .............................................
86040

Anas_platyrhynchos_platyrhynchos 86041 ACAACCCAGTAATAAAAAGTGACAATAGTATGAAAATATACTCCA
86085
Gallus_gallus                      86041 .............................................
86085

Anas_platyrhynchos_platyrhynchos 86086 TTTTTGACTTAATGTTTCTTTCATTTTCATTATCATGTCTTTGTA
86130
Gallus_gallus                      86086 .............................................
86130

Anas_platyrhynchos_platyrhynchos 86131 ACGAAATGATTATGAAAACTTGTTGTGGTTTAACCTGGCTGGCAG
86175
Gallus_gallus                      86131 .............................................
86175

Anas_platyrhynchos_platyrhynchos 86176 CTAAGCACTACATAGACATTCACTCACCCTCCCCTGTCCCTCTGG
86220
Gallus_gallus                      86176 .............................................
86220

Anas_platyrhynchos_platyrhynchos 86221 GATGGGAGAAGAGAATCAGAAAGAAATGAAAGCCTGTGGGTTGAG
86265
Gallus_gallus                      86221 .............................................
86265

Anas_platyrhynchos_platyrhynchos 86266 ATAAAGACAGTTTATGGGGACAGAAAAGAAAGGAAAATAATAATA
86310
Gallus_gallus                      86266 .............................................
86310

Anas_platyrhynchos_platyrhynchos 86311 ACAGTGATAACAGTACTACTACTAATAGTGTGTACAGAACAAGTG
86355
Gallus_gallus                      86311 .............................................
86355

Anas_platyrhynchos_platyrhynchos 86356 ATGCACAATGCAATTGCTCACCACCCGCTGACCAATGCCCAGCCT
86400
Gallus_gallus                      86356 .............................................
86400

Anas_platyrhynchos_platyrhynchos 86401 ATCCCCAAGCAGCCGGTCCCCCCAACCCAGCCAGCCACCCCATAT
86445
Gallus_gallus                      86401 .............................................
86445

Anas_platyrhynchos_platyrhynchos 86446 TAATTGTTCAGCATGATGTGATATCATATGGGATATCCCTTTGGC
86490
Gallus_gallus                      86446 .............................................
86490

Anas_platyrhynchos_platyrhynchos 86491 GAGTTTGGGTCAGCTGTCCTGGGTCTGTCCTCCCCCAGCTCCTGC
86535
Gallus_gallus                      86491 .............................................
86535

Anas_platyrhynchos_platyrhynchos 86536 TGTACACCCAATCTCCTCACTGGCAGGGCACTGAGAGAAGCTGAA
86580
Gallus_gallus                      86536 .............................................
86580
```

```
Anas_platyrhynchos_platyrhynchos 86581 AAGCCCTTGGCTTAGTGTAAGCACTTCTCTGCAACAATTAAAACA
86625
Gallus_gallus                    86581 .............................................
86625

Anas_platyrhynchos_platyrhynchos 86626 TCACTGTGGTATCAATATTATTCTCATCCTAAATCCCAAGCACAG
86670
Gallus_gallus                    86626 .............................................
86670

Anas_platyrhynchos_platyrhynchos 86671 CAACATACCAGCTACAAGGAGGAAAATTAACTCTATCCTAGCCAA
86715
Gallus_gallus                    86671 .............................................
86715

Anas_platyrhynchos_platyrhynchos 86716 CACTAGGACAAAACTCAACTGCATTTTACTTGTGAACGCCGCATG
86760
Gallus_gallus                    86716 .............................................
86760

Anas_platyrhynchos_platyrhynchos 86761 AGAAAACAGCCAAAAGGAAGCAACACATCTTGTAAACTGACTGAT
86805
Gallus_gallus                    86761 .............................................
86805

Anas_platyrhynchos_platyrhynchos 86806 CTTTGTGTTTTAGCAGAATTATTCCTTAATACCACACTTCCCATT
86850
Gallus_gallus                    86806 .............................................
86850

Anas_platyrhynchos_platyrhynchos 86851 CAGAAATGCCCACTTTGGTTGACAGATGAAGTGACTGGATTTTTA
86895
Gallus_gallus                    86851 .............................................
86895

Anas_platyrhynchos_platyrhynchos 86896 ACCTGCTTAACTTCTAGCAAGGGCATGATTGACACTTTTGATAAT
86940
Gallus_gallus                    86896 .............................................
86940

Anas_platyrhynchos_platyrhynchos 86941 ACTTCACACTCTAATGATAACATATAAGTAGAAGCTTCTAAAAAT
86985
Gallus_gallus                    86941 .............................................
86985

Anas_platyrhynchos_platyrhynchos 86986 CTAGTCCTGATTGCAGTTTCTTACCACTTTTATAAATGCAGCTCA
87030
Gallus_gallus                    86986 .............................................
87030

Anas_platyrhynchos_platyrhynchos 87031 TGACAGGTGTACATACTCACTGCAGTCACTGCCTTTGCCCAGACT
87075
Gallus_gallus                    87031 .............................................
87075

Anas_platyrhynchos_platyrhynchos 87076 GCTGAAAACCACTCCATGAAGTCAATACATTTTCACTGAAAACTG
87120
Gallus_gallus                    87076 .............................................
87120

Anas_platyrhynchos_platyrhynchos 87121 CATGCTCCCCAGACTTGGGTACATTACACTTCCAATTTACCACCT
87165
Gallus_gallus                    87121 .............................................
```

```
87165

Anas_platyrhynchos_platyrhynchos 87166 GCCTGAGGAAGCTGTTCATAGCAGCTGTAAGCTATAAATTATTTT
87210
Gallus_gallus                    87166 .............................................
87210

Anas_platyrhynchos_platyrhynchos 87211 AGATTACTTCCAGGCAAAAATAGATAGATAGATAGATAGATAGAT
87255
Gallus_gallus                    87211 .............................................
87255

Anas_platyrhynchos_platyrhynchos 87256 AGATAGATAGATAGATAGATAGATAGATGAAAGCAATGTTAGGAA
87300
Gallus_gallus                    87256 .............................................
87300

Anas_platyrhynchos_platyrhynchos 87301 AAACAGAACAGAAGGAGGGCTTTCAGTGCCTCTAGGATATCAAAT
87345
Gallus_gallus                    87301 .............................................
87345

Anas_platyrhynchos_platyrhynchos 87346 GTCTTGCAAAGAACTAATGTGTGCCTACATTAGAGTAAAGCTGAC
87390
Gallus_gallus                    87346 .............................................
87390

Anas_platyrhynchos_platyrhynchos 87391 TGAGCAGCAAGTATTGCCCTTATGCTGCTGACCAAAGTAGTTAGA
87435
Gallus_gallus                    87391 .............................................
87435

Anas_platyrhynchos_platyrhynchos 87436 ATTAAAAAAGTGGTGTTAGTACAAGAACCTATATTCCAACACATT
87480
Gallus_gallus                    87436 .............................................
87480

Anas_platyrhynchos_platyrhynchos 87481 TACATGAATGGGACCTATACTTAAACAACTAATGAGGAAATGTAC
87525
Gallus_gallus                    87481 .............................................
87525

Anas_platyrhynchos_platyrhynchos 87526 ACAGTATATGCAGATGTTAACTTGTTACAAGCTAGAGCAAAAATT
87570
Gallus_gallus                    87526 .............................................
87570

Anas_platyrhynchos_platyrhynchos 87571 TTCAGACTCTTCACATCATACAATTTGAGTTTTTATGAGTTTGAT
87615
Gallus_gallus                    87571 .............................................
87615

Anas_platyrhynchos_platyrhynchos 87616 GCCATGTTTAAATTGCTTCTTAGTTCTTTATTATGAGCTGTATTT
87660
Gallus_gallus                    87616 .............................................
87660

Anas_platyrhynchos_platyrhynchos 87661 TCTTTTCAGTCATTAGAGATTTCAAAGTAACTGATAAGAATGTAA
87705
Gallus_gallus                    87661 .............................................
87705

Anas_platyrhynchos_platyrhynchos 87706 ACAGCATAAAAAGCTACCTGTCAATCTCTTCAAGGACATCAGCAA
```

```
87750
Gallus_gallus                    87706 .............................................
87750

Anas_platyrhynchos_platyrhynchos 87751 CTGCTATCAAAGGTTTTCGCACATGTTGATGCAAATTTTGTTGTA
87795
Gallus_gallus                    87751 .............................................
87795

Anas_platyrhynchos_platyrhynchos 87796 GTAGTGGTAAGCACAGATTCCACTGGGTTGTGCAAACAACCTGAA
87840
Gallus_gallus                    87796 .............................................
87840

Anas_platyrhynchos_platyrhynchos 87841 TCACATAAGGATCTCCCAGCTGAACCGCATGTTGCAAGGTTAAAT
87885
Gallus_gallus                    87841 .............................................
87885

Anas_platyrhynchos_platyrhynchos 87886 CAAGATTTTTAATCAATTTCAGCTGAGCCTACACAAAAGAAAAAT
87930
Gallus_gallus                    87886 .............................................
87930

Anas_platyrhynchos_platyrhynchos 87931 TGCTTAGCATATACTGCATTTAATGAATACATCTTCTTACATACT
87975
Gallus_gallus                    87931 .............................................
87975

Anas_platyrhynchos_platyrhynchos 87976 CCTCTAAATTCTAAATTTAGGCTCCTACTTTTGATGCTCTAAATT
88020
Gallus_gallus                    87976 .............................................
88020

Anas_platyrhynchos_platyrhynchos 88021 ATAACCAAGTCCACTGACTATAGAAATCTAAAAAAAATCCTTCCC
88065
Gallus_gallus                    88021 .............................................
88065

Anas_platyrhynchos_platyrhynchos 88066 AGTGTGACAATTTGTCAGGAATAACCCAAATTTTTCAGAAATAAT
88110
Gallus_gallus                    88066 .............................................
88110

Anas_platyrhynchos_platyrhynchos 88111 AAGATTGCCTTTAATATGCAATAATACATCTTCCATAAATGTAAA
88155
Gallus_gallus                    88111 .............................................
88155

Anas_platyrhynchos_platyrhynchos 88156 ATATGAAAAATTATAGTATTGCCTTTGCATGTTTCAATGCTACCA
88200
Gallus_gallus                    88156 .............................................
88200

Anas_platyrhynchos_platyrhynchos 88201 TCTTTACTCTTAGAAATTAATATATTATGAATCTTCATTATCAGA
88245
Gallus_gallus                    88201 .............................................
88245

Anas_platyrhynchos_platyrhynchos 88246 CTGAAGACTTAAGAAAGGAATATATGACTATTGTTTATTTCTCTG
88290
Gallus_gallus                    88246 .............................................
88290
```

```
Anas_platyrhynchos_platyrhynchos 88291 TTTTATTCTTGCCAGCCCAACTTCCTGCTTTCTTCTTGGAAGACC
88335
Gallus_gallus                    88291 .............................................
88335

Anas_platyrhynchos_platyrhynchos 88336 AATCAGAAAGAATATAATGCTTAAAGCAATTTCTGATCAGTGTAA
88380
Gallus_gallus                    88336 .............................................
88380

Anas_platyrhynchos_platyrhynchos 88381 TGGAATTAAGACACACAAGTCCAGTGCAATTTGCATATTTCACAA
88425
Gallus_gallus                    88381 .............................................
88425

Anas_platyrhynchos_platyrhynchos 88426 ACTAAGCCTTTTATTTTTCTCTCAAATCTAAAGAACATACCTCAA
88470
Gallus_gallus                    88426 .............................................
88470

Anas_platyrhynchos_platyrhynchos 88471 CAACTCCTTTGGTATAAGTCACTATTTTAGGGCCAAATTTTTGAA
88515
Gallus_gallus                    88471 .............................................
88515

Anas_platyrhynchos_platyrhynchos 88516 TTTCATATTCACATTCCAGGCATTCTACTTCAATAAGTTTCCCTT
88560
Gallus_gallus                    88516 .............................................
88560

Anas_platyrhynchos_platyrhynchos 88561 GTTCCTATGAAGGATTGATACAAAATGCTGATTTTAATAATTAAA
88605
Gallus_gallus                    88561 .............................................
88605

Anas_platyrhynchos_platyrhynchos 88606 AAGGAATATGATTCAGACTAAAACAATGCAATGATGCTAATACTG
88650
Gallus_gallus                    88606 .............................................
88650

Anas_platyrhynchos_platyrhynchos 88651 CATTGTAATACCTTTTTGAAAATCCAACTGTACTAGCTAATTAAT
88695
Gallus_gallus                    88651 .............................................
88695

Anas_platyrhynchos_platyrhynchos 88696 TGAAACACTAAAAGAGGGATTAGAATTTGTTATAGCATAGTGTTA
88740
Gallus_gallus                    88696 .............................................
88740

Anas_platyrhynchos_platyrhynchos 88741 TTAAGATGGTATATTATTGGAATAAACAAGAGATAGTTGAGACAG
88785
Gallus_gallus                    88741 .............................................
88785

Anas_platyrhynchos_platyrhynchos 88786 TAATGAATAATCTTAATGGGTTGTGCAACTTGTAAGTGAAAGTCA
88830
Gallus_gallus                    88786 .............................................
88830

Anas_platyrhynchos_platyrhynchos 88831 CTTTAACACCCACTACTTCTAATGGAGATGAGAAGGTTGGTATAA
88875
```

```
Gallus_gallus                    88831 .............................................
88875

Anas_platyrhynchos_platyrhynchos 88876 AAGCACGCCTTCTCTACTGCTGCCACTGAAGGGTTACTTGCTGTT
88920
Gallus_gallus                    88876 .............................................
88920

Anas_platyrhynchos_platyrhynchos 88921 ACTGTCTATAAAAATATTGAAACTAAATCCTAATGAACCCTTGAT
88965
Gallus_gallus                    88921 .............................................
88965

Anas_platyrhynchos_platyrhynchos 88966 CTAGAGAATTTAAACACATTCTAGGTGAATGCAATTTAAACACGT
89010
Gallus_gallus                    88966 .............................................
89010

Anas_platyrhynchos_platyrhynchos 89011 TCTAGGTGAATGCTATGTAACTGTGTACAACCTCAGACTTGTTGG
89055
Gallus_gallus                    89011 .............................................
89055

Anas_platyrhynchos_platyrhynchos 89056 AATTTTTCAGAAAAAAAAAAAAAAAAGTGAGGGGCCTGAGGCAGAC
89100
Gallus_gallus                    89056 .............................................
89100

Anas_platyrhynchos_platyrhynchos 89101 ATCTGGAATGAAAAATTGCAGTTAGGATAAGTAAAATCTACATTA
89145
Gallus_gallus                    89101 .............................................
89145

Anas_platyrhynchos_platyrhynchos 89146 CAAGTAACTCACAATAGGATCATTACAACAGAAATATGGACTGGT
89190
Gallus_gallus                    89146 .............................................
89190

Anas_platyrhynchos_platyrhynchos 89191 ATGTTTGGCAGTATTTGCCATTACAATTTCATTACATACACTTGA
89235
Gallus_gallus                    89191 .............................................
89235

Anas_platyrhynchos_platyrhynchos 89236 ATATTTCAGGTAAGTAAGGACTTACATGGGCATCACAACACTATT
89280
Gallus_gallus                    89236 .............................................
89280

Anas_platyrhynchos_platyrhynchos 89281 GGGCACTTCTACATACTATTTTTTTTTTTTCCTGAAAAACTTACT
89325
Gallus_gallus                    89281 .............................................
89325

Anas_platyrhynchos_platyrhynchos 89326 TTTTATCTGAATACAGGGGTGAACAGTTCTGCTTATCAACCAAGA
89370
Gallus_gallus                    89326 .............................................
89370

Anas_platyrhynchos_platyrhynchos 89371 AGGAAGTGAAAAAATTGCAGAGCTAGGTGGACATAAGAATCAAAA
89415
Gallus_gallus                    89371 .............................................
89415
```

```
Anas_platyrhynchos_platyrhynchos 89416 AAGGATATACACATTGAAAGACCAGTTATTACAGAGACCACACAA
89460
Gallus_gallus                    89416 .............................................
89460

Anas_platyrhynchos_platyrhynchos 89461 AATTTCCATCTAGGAATGCATCTAGTAATAATGATATTAATCAGT
89505
Gallus_gallus                    89461 .............................................
89505

Anas_platyrhynchos_platyrhynchos 89506 ATATAAGCAATGCAGAGAAGAAGACAGAAGAACCCAGAACCATCA
89550
Gallus_gallus                    89506 .............................................
89550

Anas_platyrhynchos_platyrhynchos 89551 GCATTATTTTTAAAGTAGAATCATTGTCAAAAAAAAATCATTTCT
89595
Gallus_gallus                    89551 .............................................
89595

Anas_platyrhynchos_platyrhynchos 89596 AATTAATACTTACAGTAATGCCAGCTTTCTTCAATTCAAGTATAC
89640
Gallus_gallus                    89596 .............................................
89640

Anas_platyrhynchos_platyrhynchos 89641 AATCATGTGTAAGCTTAATGCAGTTTACTTTCACAGAAAAACGAG
89685
Gallus_gallus                    89641 .............................................
89685

Anas_platyrhynchos_platyrhynchos 89686 CCAATTCCAGGATCAAAGGAATTCTTCAAAAACAGAAAAACATCA
89730
Gallus_gallus                    89686 .............................................
89730

Anas_platyrhynchos_platyrhynchos 89731 CTTCAGAGAATCACACACTCCTCTTTAATGAAATCAGAAAAACAT
89775
Gallus_gallus                    89731 .............................................
89775

Anas_platyrhynchos_platyrhynchos 89776 TATATGAAACTAATTTTGTGCAATATATCTGAAACTCTTGAAGAT
89820
Gallus_gallus                    89776 .............................................
89820

Anas_platyrhynchos_platyrhynchos 89821 ATCTGAATATTAGAAGATGTGATTTTACCAGTAATGCTCGCAGTC
89865
Gallus_gallus                    89821 .............................................
89865

Anas_platyrhynchos_platyrhynchos 89866 AGTAAGGAAAAAGATTGAAGTTGGTTGGATGAAAAATACAAAAAT
89910
Gallus_gallus                    89866 .............................................
89910

Anas_platyrhynchos_platyrhynchos 89911 TCAGCTGTTTATGGAAAATAAATAGTATAATTCTAATGTAAAAAA
89955
Gallus_gallus                    89911 .............................................
89955

Anas_platyrhynchos_platyrhynchos 89956 TTGTTTCTGAATAACATCACACAATGAAGGATTAAACGCATGCAG
90000
Gallus_gallus                    89956 .............................................
```

90000

Anas_platyrhynchos_platyrhynchos 90001 CACAAAGATGGTGAAATAGGAAATGGGTTAAATTGCAAGTAATTC
90045
Gallus_gallus 90001 .............................................
90045

Anas_platyrhynchos_platyrhynchos 90046 TGCAAACATAATGCAGTCAACTGGCACACATCTTGAAACAGTCAA
90090
Gallus_gallus 90046 .............................................
90090

Anas_platyrhynchos_platyrhynchos 90091 ATACAATGGTCAATTCCTTAAAAAAATAGGAACACACTATTCTCA
90135
Gallus_gallus 90091 .............................................
90135

Anas_platyrhynchos_platyrhynchos 90136 TGGTAGGACACTTCTGAATATTTTGGATCTCAGTAAATAACCAAA
90180
Gallus_gallus 90136 .............................................
90180

Anas_platyrhynchos_platyrhynchos 90181 TGAGTCCCAATGTCTGCACTAGAGGTAATAAGAAAACTGAGTAGA
90225
Gallus_gallus 90181 .............................................
90225

Anas_platyrhynchos_platyrhynchos 90226 GTCAGAAAAGATTGAACTTCATTTGCAAAAATGTAGTATTACTTC
90270
Gallus_gallus 90226 .............................................
90270

Anas_platyrhynchos_platyrhynchos 90271 CATATTTAACAGCTGTGATCTAACTACTAGAATAGAGTTTGGTTT
90315
Gallus_gallus 90271 .............................................
90315

Anas_platyrhynchos_platyrhynchos 90316 TGTCTGCATTTAAAATAACACTGAAGAATTAGGAACGATTCAAAG
90360
Gallus_gallus 90316 .............................................
90360

Anas_platyrhynchos_platyrhynchos 90361 AAAAGCTAAAAAATGCATTAAAAACTATTGTGAGATCATAAAAAA
90405
Gallus_gallus 90361 .............................................
90405

Anas_platyrhynchos_platyrhynchos 90406 AATCTCTTCAAGATTTTGTCACACTTTGCTGGAAGCAGCATCAGC
90450
Gallus_gallus 90406 .............................................
90450

Anas_platyrhynchos_platyrhynchos 90451 TTCAGCATCAATTTTCAGGGACAAGTAAACTCACTTCATACTGGC
90495
Gallus_gallus 90451 .............................................
90495

Anas_platyrhynchos_platyrhynchos 90496 AACAGATGCTGGAGGTACATCATACTGTTTCAGAAGTTCATATAG
90540
Gallus_gallus 90496 .............................................
90540

Anas_platyrhynchos_platyrhynchos 90541 AGAATACAGAGTTGCGGTGGCATCACTTGGAAATTCATTTGTGTC

```
90585
Gallus_gallus                    90541 .............................................
90585

Anas_platyrhynchos_platyrhynchos 90586 CCACTGCCTGCAATATGTTACGAATTGATCACTTTTTATCACAGG
90630
Gallus_gallus                    90586 .............................................
90630

Anas_platyrhynchos_platyrhynchos 90631 AAATATTTCAATTTTTTAATTCAATAATAACTCTCCTAACTTCAG
90675
Gallus_gallus                    90631 .............................................
90675

Anas_platyrhynchos_platyrhynchos 90676 AGCAGATGGATTTTAGTCCAATATATTAGTGGATTATCTAGACTA
90720
Gallus_gallus                    90676 .............................................
90720

Anas_platyrhynchos_platyrhynchos 90721 CAAACTATTACAAAAANNNNNNNNNNNNNNNNNNNNNNNNNNNNN
90765
Gallus_gallus                    90721 .............................................
90765

Anas_platyrhynchos_platyrhynchos 90766 NNNNNNNNNNNNNNNNNNNNNNNNNNNNNNNNNNNNNNNNNNNNN
90810
Gallus_gallus                    90766 .............................................
90810

Anas_platyrhynchos_platyrhynchos 90811 NNNNNNNNNNNNNNNNNNNNNNNNNNNAAAATAAAATACAACAAAA
90855
Gallus_gallus                    90811 .............................................
90855

Anas_platyrhynchos_platyrhynchos 90856 CTCACTTCAGTATACCAAGTAAGGTTCTCGGAGAACCAGATACAA
90900
Gallus_gallus                    90856 .............................................
90900

Anas_platyrhynchos_platyrhynchos 90901 AGCAGAAAGAGCAAAATATTCCAGAATAAGAATATTTGAGGCAGT
90945
Gallus_gallus                    90901 .............................................
90945

Anas_platyrhynchos_platyrhynchos 90946 ACATTTTATTTGAAAGGCAAAGAAATATGGAGTGCAAAACTCCTT
90990
Gallus_gallus                    90946 .............................................
90990

Anas_platyrhynchos_platyrhynchos 90991 TCCCTTAATGTCAATATAAATTATTCCACGATGAAAAGAATTAGT
91035
Gallus_gallus                    90991 .............................................
91035

Anas_platyrhynchos_platyrhynchos 91036 ACGTTAAAAAGTGGAACTGTTGAGACACTGAAAATCATGTGGAAT
91080
Gallus_gallus                    91036 .............................................
91080

Anas_platyrhynchos_platyrhynchos 91081 TCTGTTACAGTCTAATTTTCAAAATTGCTTTCATGTAAAATTTAA
91125
Gallus_gallus                    91081 .............................................
91125
```

```
Anas_platyrhynchos_platyrhynchos 91126 AGTTTTAGAAAATACCATGCTTCCCATCCTTGTGATTCATCCATT
91170
Gallus_gallus                    91126 .............................................
91170

Anas_platyrhynchos_platyrhynchos 91171 ATATTTCAGATGTGATTCCTCTTCTCTATCTTTACTTTCTTATAT
91215
Gallus_gallus                    91171 .............................................
91215

Anas_platyrhynchos_platyrhynchos 91216 ATGACTTATGTACCAAGCCCCAATCTGCTAGATACAAAAACGTCT
91260
Gallus_gallus                    91216 .............................................
91260

Anas_platyrhynchos_platyrhynchos 91261 TTCACTCTCTTGTAAATTCTTTCACTTGAGACTGTCAGTACAACA
91305
Gallus_gallus                    91261 .............................................
91305

Anas_platyrhynchos_platyrhynchos 91306 CACTATTCTACACATTCCCCAAGAATTCTCTCACCTTTTAGTTTC
91350
Gallus_gallus                    91306 .............................................
91350

Anas_platyrhynchos_platyrhynchos 91351 GGCAGTTTCATGGACAGGACTGAGCTTCACTGCTCAAAACTATTT
91395
Gallus_gallus                    91351 .............................................
91395

Anas_platyrhynchos_platyrhynchos 91396 CCATTAGAGAAACTAATCTTTGTAGTATGGTCTTCTACTGGCATA
91440
Gallus_gallus                    91396 .............................................
91440

Anas_platyrhynchos_platyrhynchos 91441 TGAGATATCAGAACCCGACTCTACTTCTGTAATATCTAGTATCAA
91485
Gallus_gallus                    91441 .............................................
91485

Anas_platyrhynchos_platyrhynchos 91486 AATCCAAGTGAAAGGATACAGTTTAGTACTCACGATTTTATCATT
91530
Gallus_gallus                    91486 .............................................
91530

Anas_platyrhynchos_platyrhynchos 91531 TGCATTTTATAAATCACTGACAAATGGATGGAATTTTGTATTTCC
91575
Gallus_gallus                    91531 .............................................
91575

Anas_platyrhynchos_platyrhynchos 91576 TTCTCTATCTGGGAAATATCCATTAGTTTGTGGTATACCTATAAA
91620
Gallus_gallus                    91576 .............................................
91620

Anas_platyrhynchos_platyrhynchos 91621 ATAATAATTTCATTATTAATCCACTAAATAAGCCAGTGAAGGACA
91665
Gallus_gallus                    91621 .............................................
91665

Anas_platyrhynchos_platyrhynchos 91666 GACACTCAACAGGGCTAAAATAAATCATGTATGTTGATTGTAGGC
91710
```

Gallus_gallus 91666 .............................................
91710

Anas_platyrhynchos_platyrhynchos 91711 TTTAAGGGCAATGGAATTTAATGGACATTAAGGGCAATGAAATTA
91755
Gallus_gallus 91711 .............................................
91755

Anas_platyrhynchos_platyrhynchos 91756 CTATGGTAATCTATGTTGTTTTGCATACATACCACTTAGTAAAGC
91800
Gallus_gallus 91756 .............................................
91800

Anas_platyrhynchos_platyrhynchos 91801 TGTATTAATTTTCATATGCTGAGAATCATAAGTATTAGCAGAAAT
91845
Gallus_gallus 91801 .............................................
91845

Anas_platyrhynchos_platyrhynchos 91846 CTAAATCTGGAACTATTATCTGAGTTAGATTTCTAATTAATCAAT
91890
Gallus_gallus 91846 .............................................
91890

Anas_platyrhynchos_platyrhynchos 91891 TTTTTAGAACATTCTGATAACACTCATTCTGAAATCATCTCATCC
91935
Gallus_gallus 91891 .............................................
91935

Anas_platyrhynchos_platyrhynchos 91936 AGATAAGGGAAATTAGACATTCAAATTTAACTGCAACTATAAATT
91980
Gallus_gallus 91936 .............................................
91980

Anas_platyrhynchos_platyrhynchos 91981 AAATATAACAAAAATATTACAAAGAAATTACAAGGTACTGTAGTT
92025
Gallus_gallus 91981 .............................................
92025

Anas_platyrhynchos_platyrhynchos 92026 ACTTCAATTTATCTGTAGAGTCACATATTAGACAAATAACATCAT
92070
Gallus_gallus 92026 .............................................
92070

Anas_platyrhynchos_platyrhynchos 92071 TTCTTTTGACTCTCTGACACTGACTGTCCACAGCAATCAATAATG
92115
Gallus_gallus 92071 .............................................
92115

Anas_platyrhynchos_platyrhynchos 92116 ATTACTTAAGTTGTAATGGAACACTGCATTTTGTAGGACTTGATT
92160
Gallus_gallus 92116 .............................................
92160

Anas_platyrhynchos_platyrhynchos 92161 ACTCTGTCATTAAAGCATCCAATAGTGTAGGTCATACTTCTGTAG
92205
Gallus_gallus 92161 .............................................
92205

Anas_platyrhynchos_platyrhynchos 92206 CTCTGCTGATAAACACAGCCCAAGGAAACTTTTAGAACTACTTTA
92250
Gallus_gallus 92206 .............................................
92250

```
Anas_platyrhynchos_platyrhynchos 92251 TTCTTTTAAGACCAATGTTAAGTAGTTTTGTTCAAAATTGCCTCT
92295
Gallus_gallus                    92251 .............................................
92295

Anas_platyrhynchos_platyrhynchos 92296 AAAAAAAAAATACATGTACAAAAATAATTTGGATCCCTTGCCTAA
92340
Gallus_gallus                    92296 .............................................
92340

Anas_platyrhynchos_platyrhynchos 92341 ATGTCATTGCTTCACTGATTTCATACCTCTGAAAGGTCATTAAGG
92385
Gallus_gallus                    92341 .............................................
92385

Anas_platyrhynchos_platyrhynchos 92386 TTCTCTTAATTCAAATGCTAAAGAACATATAACATAAACATAATC
92430
Gallus_gallus                    92386 .............................................
92430

Anas_platyrhynchos_platyrhynchos 92431 TTTTGAAATGCAGTGAAAACGCGTAAGTAAGACAAGTAAGAGTGT
92475
Gallus_gallus                    92431 .............................................
92475

Anas_platyrhynchos_platyrhynchos 92476 AGCAGGGAACATAGAAGGAATGCTTAAGAACATGACTGCATGCAA
92520
Gallus_gallus                    92476 .............................................
92520

Anas_platyrhynchos_platyrhynchos 92521 CTCCAAAAGTTTTATTTGCCTATTTCTGCTTTTGTCACATCTATC
92565
Gallus_gallus                    92521 .............................................
92565

Anas_platyrhynchos_platyrhynchos 92566 TTCACCCTCTTATTTCCCAAATGAAGGTCATGCAAAAAAATGTGA
92610
Gallus_gallus                    92566 .............................................
92610

Anas_platyrhynchos_platyrhynchos 92611 ACTAATATCATATACTTTAATTCCCAGAACTCTATAGGGTTCACA
92655
Gallus_gallus                    92611 .............................................
92655

Anas_platyrhynchos_platyrhynchos 92656 TTGGGTCATTTAGGTATATTTCAGCATAAGAATAGTCAACAAAAC
92700
Gallus_gallus                    92656 .............................................
92700

Anas_platyrhynchos_platyrhynchos 92701 CTAAATAGAGCCTAGATTATTTTTTCCCAGTGGCATAAAACAGGT
92745
Gallus_gallus                    92701 .............................................
92745

Anas_platyrhynchos_platyrhynchos 92746 AAAGTTACACCCTTTACACCATCTGTGATAATAGCAGTTATTCCC
92790
Gallus_gallus                    92746 .............................................
92790

Anas_platyrhynchos_platyrhynchos 92791 ACACAGATTTTTTAGGGGTAGCAAAAGTTTTCCCTATAATATTAT
92835
Gallus_gallus                    92791 .............................................
```

```
92835

Anas_platyrhynchos_platyrhynchos 92836 GGTGTACTGAAATAACATGGGGCACTAAGGAGCTGTTTAGCTTGG
92880
Gallus_gallus                    92836 .............................................
92880

Anas_platyrhynchos_platyrhynchos 92881 AGTATGAAAAATACAAACAATAAAAATATCTGCTTATTTGGTGTA
92925
Gallus_gallus                    92881 .............................................
92925

Anas_platyrhynchos_platyrhynchos 92926 TCATTTGTTAGTTCACAATATTTAAGTACAGGAAATAAAATACTT
92970
Gallus_gallus                    92926 .............................................
92970

Anas_platyrhynchos_platyrhynchos 92971 AAATAAAACAAGGATGTTAATAAATGTAGAGAAAGCTCCTGTCTT
93015
Gallus_gallus                    92971 .............................................
93015

Anas_platyrhynchos_platyrhynchos 93016 ACTGGGAATTTTAAATATTTTACAGCATTACCATTAAAGAAAATA
93060
Gallus_gallus                    93016 .............................................
93060

Anas_platyrhynchos_platyrhynchos 93061 TTTGAGAGTATTTATCTGGAGCATTTTCCTTAATAAAGACTGCTG
93105
Gallus_gallus                    93061 .............................................
93105

Anas_platyrhynchos_platyrhynchos 93106 CAGTCGATGAAAACTCTTTTGCTTCTTTCAGTTTTGAAGCATCGA
93150
Gallus_gallus                    93106 .............................................
93150

Anas_platyrhynchos_platyrhynchos 93151 GGAGGCACTCAAGTAAATTTCTGCAGATTAAATAAATATTTATAT
93195
Gallus_gallus                    93151 .............................................
93195

Anas_platyrhynchos_platyrhynchos 93196 TAAACATTTATCAATGTTTTTGAAACAATTCTTATTCATATTTTT
93240
Gallus_gallus                    93196 .............................................
93240

Anas_platyrhynchos_platyrhynchos 93241 TCAAGAGACTCTTTATATTCAAGGTACCACTCTTAAATGCAGTAC
93285
Gallus_gallus                    93241 .............................................
93285

Anas_platyrhynchos_platyrhynchos 93286 AATCAATTAATGCTATAAAGCATTGACTCAGCAAAGAGAACTTCT
93330
Gallus_gallus                    93286 .............................................
93330

Anas_platyrhynchos_platyrhynchos 93331 AACTTTTATGCATTTGTTTCTTCCAGAAAATACACAGCTCTTAAT
93375
Gallus_gallus                    93331 .............................................
93375

Anas_platyrhynchos_platyrhynchos 93376 ATCCATTTATTACAGATCACAAAAATAATCTTGGAACAGGAGATT
```

```
93420
Gallus_gallus                    93376 .............................................
93420

Anas_platyrhynchos_platyrhynchos 93421 TTTCTGAACACTCTGCACAACAGGACACCTGGAATTAAAATGATC
93465
Gallus_gallus                    93421 .............................................
93465

Anas_platyrhynchos_platyrhynchos 93466 AATGATTTAACACTTCATGAAGAAAAGAAAAAAAAAGGTGAGCTG
93510
Gallus_gallus                    93466 .............................................
93510

Anas_platyrhynchos_platyrhynchos 93511 CATTCTGTGGTTCTCCATTAACTACTGGAAAGGCTAGTATTTCAA
93555
Gallus_gallus                    93511 .............................................
93555

Anas_platyrhynchos_platyrhynchos 93556 AGTCATCTAGTAAAAGAGAACCCATACATGCTGGAAACTTCAATG
93600
Gallus_gallus                    93556 .............................................
93600

Anas_platyrhynchos_platyrhynchos 93601 TTTTTCAATCCTTCAACTGTTTTCACTGAAAAAGAGCCTGTGGCT
93645
Gallus_gallus                    93601 .............................................
93645

Anas_platyrhynchos_platyrhynchos 93646 TCATAAAATAAATAATAAAAATACCATAGAAATAATAGTAAATGA
93690
Gallus_gallus                    93646 .............................................
93690

Anas_platyrhynchos_platyrhynchos 93691 TTCCATAAAAATATAAAAATACAAATATAAGAAAAAATAAAGGTT
93735
Gallus_gallus                    93691 .............................................
93735

Anas_platyrhynchos_platyrhynchos 93736 ACCATTAGTTTCATACCACAAATTCATTGTGCACTTGCTAATTCT
93780
Gallus_gallus                    93736 .............................................
93780

Anas_platyrhynchos_platyrhynchos 93781 CATCAATACCTCAGCAAGATTATGTTTGTTATTAAAAAACTGTAT
93825
Gallus_gallus                    93781 .............................................
93825

Anas_platyrhynchos_platyrhynchos 93826 TGTATTCTATATAACCTTTAGTGTATAACATGAAATGTTTTGATT
93870
Gallus_gallus                    93826 .............................................
93870

Anas_platyrhynchos_platyrhynchos 93871 TTTTATTTCCCTCCAGTTTCAACTTAGAAAGAAAACACAAAAAAG
93915
Gallus_gallus                    93871 .............................................
93915

Anas_platyrhynchos_platyrhynchos 93916 GCTGTACAGCAATAAACTATCAAGTCCATAGACCAAAGAAATAGA
93960
Gallus_gallus                    93916 .............................................
93960
```

```
Anas_platyrhynchos_platyrhynchos 93961 AAAGTTCACAAAGAGAAAGTTAAAAACTGTGGTAGAGATGTATCA
94005
Gallus_gallus                    93961 .............................................
94005

Anas_platyrhynchos_platyrhynchos 94006 AGAACTGGAAATGGTCTGTAGAATAGCAGACCCAAATACCTCTAA
94050
Gallus_gallus                    94006 .............................................
94050

Anas_platyrhynchos_platyrhynchos 94051 AGAAAGGTTGTCTGATCTTGAAAAATTATATAGACGCTCTTTATT
94095
Gallus_gallus                    94051 .............................................
94095

Anas_platyrhynchos_platyrhynchos 94096 TTGCACATAAAACCGGTTCTCTCAGTTATTTGTAAAGGCATCTCA
94140
Gallus_gallus                    94096 .............................................
94140

Anas_platyrhynchos_platyrhynchos 94141 GTGTTTATGATCCAGCTTTACTCAGAGAATCCAAGACAATGATGT
94185
Gallus_gallus                    94141 .............................................
94185

Anas_platyrhynchos_platyrhynchos 94186 AACAATTCATCATTGTTAATTGATATTGTTACATCATTAGTAATA
94230
Gallus_gallus                    94186 .............................................
94230

Anas_platyrhynchos_platyrhynchos 94231 ATGGGTAATAATGTGAATTGTTATCAGAGTACCAATAGCGCACAG
94275
Gallus_gallus                    94231 .............................................
94275

Anas_platyrhynchos_platyrhynchos 94276 TTTTTGGAATGTTTGTTGGTTTTTTTTTGTTTGGTTGTTTGTTTG
94320
Gallus_gallus                    94276 .............................................
94320

Anas_platyrhynchos_platyrhynchos 94321 AAACACTTCCATTTATATCTTGTTTAGAGACAAAAAACAACCTGA
94365
Gallus_gallus                    94321 .............................................
94365

Anas_platyrhynchos_platyrhynchos 94366 TTCTTTGTTCAGTTCTAGAGCCAAAAAACATCAATGGCAAATTAA
94410
Gallus_gallus                    94366 .............................................
94410

Anas_platyrhynchos_platyrhynchos 94411 TTTAATATCAGAATCAGGCATTATATAAATATCAATTTCTAACTT
94455
Gallus_gallus                    94411 .............................................
94455

Anas_platyrhynchos_platyrhynchos 94456 TCTGTTTTCCTTATAGTCTGTAAACTAGAAGACTTAGGCTGACAC
94500
Gallus_gallus                    94456 .............................................
94500

Anas_platyrhynchos_platyrhynchos 94501 TCTGCCTGCTTTGCTTCAGTCTTCACCAAAAAAAACTCTCATGCC
94545
```

```
Gallus_gallus                   94501 .............................................
94545

Anas_platyrhynchos_platyrhynchos 94546 TCAGTACTCTCAGTTTCTGCTCAACCAACATTATTTTTTTTAATT
94590
Gallus_gallus                   94546 .............................................
94590

Anas_platyrhynchos_platyrhynchos 94591 TATTTTTATGAGGGTTTTGCATATAATTTGCCATTTCTTTTGTTG
94635
Gallus_gallus                   94591 .............................................
94635

Anas_platyrhynchos_platyrhynchos 94636 AAATTGAAACCAAAGACTCAATAGTATGACTAAGCTTCTAATATT
94680
Gallus_gallus                   94636 .............................................
94680

Anas_platyrhynchos_platyrhynchos 94681 GCTTTTCGTTAGTGATCAGAGACTTAATGAATTTCTTAGCCAACA
94725
Gallus_gallus                   94681 .............................................
94725

Anas_platyrhynchos_platyrhynchos 94726 TGTGCTGCTCAGTCCATAAATATCTGCTCTTATCTGCAAAAACTT
94770
Gallus_gallus                   94726 .............................................
94770

Anas_platyrhynchos_platyrhynchos 94771 CAAATTATCTAACAGATAGCTCTTGTTCTTAAAATGTGTATCTCA
94815
Gallus_gallus                   94771 .............................................
94815

Anas_platyrhynchos_platyrhynchos 94816 GTAAAATCTGAATTGCAATCAGAATGCAAATCAGATGCTATCAGT
94860
Gallus_gallus                   94816 .............................................
94860

Anas_platyrhynchos_platyrhynchos 94861 TCAGAGAGGCTGAGGGGAGACCACATCGCTCTATGCAACTACCTC
94905
Gallus_gallus                   94861 .............................................
94905

Anas_platyrhynchos_platyrhynchos 94906 AAAGGAGAGTGTCGGTCTTATTTCTCAGATGACAAGTGACAAGAC
94950
Gallus_gallus                   94906 .............................................
94950

Anas_platyrhynchos_platyrhynchos 94951 GTGAGGAAACAGCTTCAAGTTGCACCAGGGGAGGTTTAGATTTGG
94995
Gallus_gallus                   94951 .............................................
94995

Anas_platyrhynchos_platyrhynchos 94996 TATTGGGAAGCACCTCTTCATGGAGAGGGTGCTTAGGAATTGAAG
95040
Gallus_gallus                   94996 .............................................
95040

Anas_platyrhynchos_platyrhynchos 95041 TGGACTACCAAGGGAAGTAGTAGATTCCCCATCCCTGGAGGCATT
95085
Gallus_gallus                   95041 .............................................
95085
```

```
Anas_platyrhynchos_platyrhynchos 95086 TGAGAGAGATGTGGACATGGCAGTTAGCGACACAGTTCAGTGATG
95130
Gallus_gallus                    95086 .............................................
95130

Anas_platyrhynchos_platyrhynchos 95131 AAACTCAGTACGTTGGGTTGACAGTTGGACTTGATCATGAGGGTC
95175
Gallus_gallus                    95131 .............................................
95175

Anas_platyrhynchos_platyrhynchos 95176 TTTTCCAACCTGAATGATTCCAGGATTCTGTAAGTGTGTGGTATA
95220
Gallus_gallus                    95176 .............................................
95220

Anas_platyrhynchos_platyrhynchos 95221 TACACTGTTCCATACACAAAGGGAAGTCAGATTGAGACCACTCAC
95265
Gallus_gallus                    95221 .............................................
95265

Anas_platyrhynchos_platyrhynchos 95266 AGCTCAAACAAAAAAAGAAAGAACTTTCCAACAGATAGTGATAAT
95310
Gallus_gallus                    95266 .............................................
95310

Anas_platyrhynchos_platyrhynchos 95311 TATTTATAACGTACCTTTCTGTAATTCTGGAAAAAAATATTTGAA
95355
Gallus_gallus                    95311 .............................................
95355

Anas_platyrhynchos_platyrhynchos 95356 CGTGATGTATCAAAAAGTAGCAATTTCTTCAGTCTCTTAAAAATA
95400
Gallus_gallus                    95356 .............................................
95400

Anas_platyrhynchos_platyrhynchos 95401 ACAGCTATAGCAGAAAGTAACAACATACTATTTAAAAACCCTGTG
95445
Gallus_gallus                    95401 .............................................
95445

Anas_platyrhynchos_platyrhynchos 95446 ATTCTATCTGTCTTTTAAAATGACATTTAATCAAATTGTATTTTT
95490
Gallus_gallus                    95446 .............................................
95490

Anas_platyrhynchos_platyrhynchos 95491 CACTGTCTTCTACAAGCCATAAGGACTTTTCAGTCATGCTTAGTA
95535
Gallus_gallus                    95491 .............................................
95535

Anas_platyrhynchos_platyrhynchos 95536 AGACTGTCCTTGAAGTGAAACTACCTTTGATTGTCCTTGAAACTA
95580
Gallus_gallus                    95536 .............................................
95580

Anas_platyrhynchos_platyrhynchos 95581 CCATGTACTTCTTCAGCCACAAACTGTCTTCAGTGAAAACCAATT
95625
Gallus_gallus                    95581 .............................................
95625

Anas_platyrhynchos_platyrhynchos 95626 GCTAAAGATTCTGATTAGAGCTAATTATTTAATTCAGTGTAAATT
95670
Gallus_gallus                    95626 .............................................
```

95670

Anas_platyrhynchos_platyrhynchos 95671 ACACTATACTCACATCTTAACACTACAAATGCAGAAAAAGCTAGG
95715
Gallus_gallus 95671 .............................................
95715

Anas_platyrhynchos_platyrhynchos 95716 TATTTCCTGAATAGGACAGAAATCAGCTAACAAATTTCAGTTATG
95760
Gallus_gallus 95716 .............................................
95760

Anas_platyrhynchos_platyrhynchos 95761 ACATAGGTGTAAACGAGTCAGTAACACATTTTTATCCAACCCTGG
95805
Gallus_gallus 95761 .............................................
95805

Anas_platyrhynchos_platyrhynchos 95806 ATTACTAATCTCCTCCCTAAAAATTATTAGACTTCTAATTTTATT
95850
Gallus_gallus 95806 .............................................
95850

Anas_platyrhynchos_platyrhynchos 95851 TATTTTTTTTTTTTTGCATAAAGCTTAAAGATATTCTCTGAATTC
95895
Gallus_gallus 95851 .............................................
95895

Anas_platyrhynchos_platyrhynchos 95896 ACCCTGTTGGGAGGTTTAGTTCTCAAAAATGCACTAATTACCAGA
95940
Gallus_gallus 95896 .............................................
95940

Anas_platyrhynchos_platyrhynchos 95941 AAGAAAACCTGCCAGTTTAAAACTTAGCAGCCTTGTTTATGCAAG
95985
Gallus_gallus 95941 .............................................
95985

Anas_platyrhynchos_platyrhynchos 95986 TTTCTCTTTCACATAACTTTTTGACTTCTGTTTGTCATAAAACTG
96030
Gallus_gallus 95986 .............................................
96030

Anas_platyrhynchos_platyrhynchos 96031 GTTAAAAAAAGGACCTATTTCATTTATACTTTGAAACATTAGGCC
96075
Gallus_gallus 96031 .............................................
96075

Anas_platyrhynchos_platyrhynchos 96076 AAGCAGTTTATTCAAATTAGTACGAGTAGCCTTTAATTTAATATT
96120
Gallus_gallus 96076 .............................................
96120

Anas_platyrhynchos_platyrhynchos 96121 GAAGAAAACTACATATAAAACTAATTAGGAAGAAAAAGGCTTGAT
96165
Gallus_gallus 96121 .............................................
96165

Anas_platyrhynchos_platyrhynchos 96166 CTGAACTAGTTACATTAGTAAGAATCATACAGAACCAGATAGTAC
96210
Gallus_gallus 96166 .............................................
96210

Anas_platyrhynchos_platyrhynchos 96211 TTAAAACATGTGACTAACAACATGAAAAAAAAAACACACAACAAC

```
96255
Gallus_gallus                     96211 .............................................
96255

Anas_platyrhynchos_platyrhynchos 96256 ACAAAAACATCACTACTGGTGATGATTCATCTCTGTTTTCTTGAT
96300
Gallus_gallus                     96256 .............................................
96300

Anas_platyrhynchos_platyrhynchos 96301 GTTGCACTTCCAATATATTTATTCTGCTCACTGAACTTATTTTTC
96345
Gallus_gallus                     96301 .............................................
96345

Anas_platyrhynchos_platyrhynchos 96346 TCAAAGTTTCATTTCAACGTACAGCTACAGACTACCAAGAGAAAA
96390
Gallus_gallus                     96346 .............................................
96390

Anas_platyrhynchos_platyrhynchos 96391 TACAGTAAGAAAGATAACTAAGATAAGGATATATATAAAATCTTC
96435
Gallus_gallus                     96391 .............................................
96435

Anas_platyrhynchos_platyrhynchos 96436 AGAATGAAGAAGTAAATTAGAAGATACACTCTCACAGCCTTACCA
96480
Gallus_gallus                     96436 .............................................
96480

Anas_platyrhynchos_platyrhynchos 96481 GGCTGTTTCTGTCTCATCAGGTATCAGAGGATAGAAGAGGACAGA
96525
Gallus_gallus                     96481 .............................................
96525

Anas_platyrhynchos_platyrhynchos 96526 AGATATTTGATTATCTATTTAAACATTATGTTTCCTTACCTGAAG
96570
Gallus_gallus                     96526 .............................................
96570

Anas_platyrhynchos_platyrhynchos 96571 AAAAAGGTTCTTAAAGAAAACTGGAGTTTGTATTTCTCCTCACTT
96615
Gallus_gallus                     96571 .............................................
96615

Anas_platyrhynchos_platyrhynchos 96616 CAAAGGACTACACATAAGCAATGTGTTGAACACTATATCTGAAAT
96660
Gallus_gallus                     96616 .............................................
96660

Anas_platyrhynchos_platyrhynchos 96661 GACTGTGACAGTTCAGATAGGACTTGGGGTCTGACTGCATCTGCT
96705
Gallus_gallus                     96661 .............................................
96705

Anas_platyrhynchos_platyrhynchos 96706 ACTAACATTTTAACCTCAAAGTCAATATTAAAAGGAAGAATTATG
96750
Gallus_gallus                     96706 .............................................
96750

Anas_platyrhynchos_platyrhynchos 96751 TGTATATTTATATATTGTTAAAAAGTTATGAATTATGACTCCCCA
96795
Gallus_gallus                     96751 .............................................
96795
```

```
Anas_platyrhynchos_platyrhynchos 96796 GAGTGGTGAAGCATTGGAATGGGCTGCCCAGGGAAGTGGTGGAGT
96840
Gallus_gallus                    96796 .............................................
96840

Anas_platyrhynchos_platyrhynchos 96841 TCCCTGCCCCTAGAGGTATTTAAAAGATGTGTGGATGTGGCACTA
96885
Gallus_gallus                    96841 .............................................
96885

Anas_platyrhynchos_platyrhynchos 96886 AGGGACATATTATAGTGATGGAACTCAGTAGGCTGGGCTGACAGT
96930
Gallus_gallus                    96886 .............................................
96930

Anas_platyrhynchos_platyrhynchos 96931 TGGATTGATGATTTCAAAGGTCTTTTCCAACCTAAATGATTCTAT
96975
Gallus_gallus                    96931 .............................................
96975

Anas_platyrhynchos_platyrhynchos 96976 AATAAAATAAACATTATTAAAAACTACCTCAGGTGGCAGCCACAT
97020
Gallus_gallus                    96976 .............................................
97020

Anas_platyrhynchos_platyrhynchos 97021 GCACAAAAGGACTACAGAATACCCATATGGAAATTAAAATTAAAC
97065
Gallus_gallus                    97021 .............................................
97065

Anas_platyrhynchos_platyrhynchos 97066 TTACATCATGAGTTCTGCTCTCCATTCACCGTCTGGCTCATCAGT
97110
Gallus_gallus                    97066 .............................................
97110

Anas_platyrhynchos_platyrhynchos 97111 TTGGTTTAATGCTTTAACAATTTGAGACAGGCTGGGAACAAGGTA
97155
Gallus_gallus                    97111 .............................................
97155

Anas_platyrhynchos_platyrhynchos 97156 ACAGCGGAATCCAGGCTTAAGATAGGGTCTCACAAATTGCCAATA
97200
Gallus_gallus                    97156 .............................................
97200

Anas_platyrhynchos_platyrhynchos 97201 AAGAACTGAGGCATTATATATCAAAAAATAGTATCTAAAAAAAAA
97245
Gallus_gallus                    97201 .............................................
97245

Anas_platyrhynchos_platyrhynchos 97246 AAAGAGAGAGAAGAATCAAGAATGGAGGATGTACTTAGATTAACT
97290
Gallus_gallus                    97246 .............................................
97290

Anas_platyrhynchos_platyrhynchos 97291 TGAAACTACAGAGTAAGAATTTTAAGACACTCCTAGGAAATAAAA
97335
Gallus_gallus                    97291 .............................................
97335

Anas_platyrhynchos_platyrhynchos 97336 ATTTTAACTTTTCCTACTGAAGTTCCTGAATTAACCTTGACTCAG
97380
```

```
Gallus_gallus                     97336 .............................................
97380

Anas_platyrhynchos_platyrhynchos 97381 AACTTTGAGCACACGTACAGCATAGGAAGAGTAAGGTAACTCTTA
97425
Gallus_gallus                     97381 .............................................
97425

Anas_platyrhynchos_platyrhynchos 97426 AGACTACACTTTACCCAGTAATTAACTACTTAAAATAAGCTAAAA
97470
Gallus_gallus                     97426 .............................................
97470

Anas_platyrhynchos_platyrhynchos 97471 TGCTGCTCTCCTCCTCCAGAGGATAAATATAAAAGAGGAAGAAAA
97515
Gallus_gallus                     97471 .............................................
97515

Anas_platyrhynchos_platyrhynchos 97516 GCCCTGAGATACCTACCAGGTCTAAACAGTCTGTTTCATCCTGTT
97560
Gallus_gallus                     97516 .............................................
97560

Anas_platyrhynchos_platyrhynchos 97561 TATTTAATACTATTTACAGTATTCTATACTGTCACAGCATGCACA
97605
Gallus_gallus                     97561 .............................................
97605

Anas_platyrhynchos_platyrhynchos 97606 GTAGTTAAAGAAATAAAGTAAGAAAACGGAAGGGAAATCCATCAA
97650
Gallus_gallus                     97606 .............................................
97650

Anas_platyrhynchos_platyrhynchos 97651 GAAAAAAAAAAAAGACTATCTAGTGGAAAACTGAGACAAACACCT
97695
Gallus_gallus                     97651 .............................................
97695

Anas_platyrhynchos_platyrhynchos 97696 ATGAAAACCTAATTCACTAACATCTCCTGAGATTTAAAGAGTAAA
97740
Gallus_gallus                     97696 .............................................
97740

Anas_platyrhynchos_platyrhynchos 97741 TACTATCAGGAGGTCCTACCTTTGATCATGTGTTGCAAAATCTAT
97785
Gallus_gallus                     97741 .............................................
97785

Anas_platyrhynchos_platyrhynchos 97786 TGCTTTCATAAAAAACAGCACAAATTTTTCAAATGCTTCCTAAAA
97830
Gallus_gallus                     97786 .............................................
97830

Anas_platyrhynchos_platyrhynchos 97831 TAGAGCAAACATGATTTAGGAAATTTGTACATAGCCTTTATAAAT
97875
Gallus_gallus                     97831 .............................................
97875

Anas_platyrhynchos_platyrhynchos 97876 TACCAATACATTATTTTCATATTTCAATATGTTCATGTCTCATTA
97920
Gallus_gallus                     97876 .............................................
97920
```

Anas_platyrhynchos_platyrhynchos 97921 TGTTTATTTTCAGACCATTCCCAAATAAAACATGTTGTCATAATA
97965
Gallus_gallus 97921 .............................................
97965

Anas_platyrhynchos_platyrhynchos 97966 ACGTGTTGCCATACACCAGAACAACTGAAAAACAGTAGCATTGCC
98010
Gallus_gallus 97966 .............................................
98010

Anas_platyrhynchos_platyrhynchos 98011 TGTTACCATGCTTGCCAGATTTCAAAATTTCAGACAATTTTTTCT
98055
Gallus_gallus 98011 .............................................
98055

Anas_platyrhynchos_platyrhynchos 98056 TTAATTTCTGCCAAAGTTTAATCATTAGGACTGAAATTGTCCAAA
98100
Gallus_gallus 98056 .............................................
98100

Anas_platyrhynchos_platyrhynchos 98101 GCTGTAAAGTTGGGTCCTTTTAAATTTTTCAGCAAAGAAAGCTCA
98145
Gallus_gallus 98101 .............................................
98145

Anas_platyrhynchos_platyrhynchos 98146 GAAATTTCCAAGATGAAAGCTAGGAAAAAAAATATTGGTTTGTTAC
98190
Gallus_gallus 98146 .............................................
98190

Anas_platyrhynchos_platyrhynchos 98191 ATTAAAAAAATGTAGGTATCATTTTCTTCGGAAAAGTTCCAACAC
98235
Gallus_gallus 98191 .............................................
98235

Anas_platyrhynchos_platyrhynchos 98236 GCCATACTCGTAATCATTTGTCAACTGAATGCTGTCAGGCATGTT
98280
Gallus_gallus 98236 .............................................
98280

Anas_platyrhynchos_platyrhynchos 98281 CTTTTAATCGTCAATGCTGCTGAAAAGCAGCTTTTCAGGAAAAAC
98325
Gallus_gallus 98281 .............................................
98325

Anas_platyrhynchos_platyrhynchos 98326 AAACAAACAGAAACACAAAAATCAGGGGGCCTGGTGGATACCAGC
98370
Gallus_gallus 98326 .............................................
98370

Anas_platyrhynchos_platyrhynchos 98371 TTGAACATAAGCCAGTAACTTATGTATGTGAATTTGCAGCAAAGG
98415
Gallus_gallus 98371 .............................................
98415

Anas_platyrhynchos_platyrhynchos 98416 AGGTGAGCATTATCCTGGACTGCATAAGGACAAGTGTCACCAGCA
98460
Gallus_gallus 98416 .............................................
98460

Anas_platyrhynchos_platyrhynchos 98461 GACTGAGGGAGGTGACCCTTCCCCTCTGTTTTGCAGTGGTGAGAC
98505
Gallus_gallus 98461 .............................................

98505

Anas_platyrhynchos_platyrhynchos 98506 CACACCTGGAGAGCTGGGTCCAGCTGTGGGCTTCCACAGGACAAG 98550
Gallus_gallus 98506 ............................................. 98550

Anas_platyrhynchos_platyrhynchos 98551 AGAGACATGGACATACTGGAAAGAGTCCAGGGAAGCACCACAAAT 98595
Gallus_gallus 98551 ............................................. 98595

Anas_platyrhynchos_platyrhynchos 98596 AGGGTCAATAAAGGTCTAGAGCATCTCACCTATAAGGAAAGACTA 98640
Gallus_gallus 98596 ............................................. 98640

Anas_platyrhynchos_platyrhynchos 98641 AGAGAGCTGGGACTGCTAAGCGTGGAGAAAAGAAGTCTCAGAGGA 98685
Gallus_gallus 98641 ............................................. 98685

Anas_platyrhynchos_platyrhynchos 98686 TCTTATCAGTGCATACAAATTACTGAAGGGAGGGTGCAAAAAAGA 98730
Gallus_gallus 98686 ............................................. 98730

Anas_platyrhynchos_platyrhynchos 98731 TGGATCCAGGCTTTTTCAGTAGTGTCCAGCAACAGGACAGGAGGC 98775
Gallus_gallus 98731 ............................................. 98775

Anas_platyrhynchos_platyrhynchos 98776 AATGGGCACAAACTGAAATACAGGAGGTTTCCTCCGAACACAAGA 98820
Gallus_gallus 98776 ............................................. 98820

Anas_platyrhynchos_platyrhynchos 98821 CAACACTTTTCTATTGTGAGTGTGACTGAGCACTAGCAGAGGTTG 98865
Gallus_gallus 98821 ............................................. 98865

Anas_platyrhynchos_platyrhynchos 98866 CCCAGAAGCTGTAGAGTCACGATCTCTGTGAGATATTCAAAAGTT 98910
Gallus_gallus 98866 ............................................. 98910

Anas_platyrhynchos_platyrhynchos 98911 GTCTGGACTTGGTCCTGGAAAACCACCTCTGGGTGGCCCTGCTTG 98955
Gallus_gallus 98911 ............................................. 98955

Anas_platyrhynchos_platyrhynchos 98956 GAGTAGATGACCTCCACAGGTCCCCTCCAGCCTTTACCATTCCAT 99000
Gallus_gallus 98956 ............................................. 99000

Anas_platyrhynchos_platyrhynchos 99001 ACTTTCTTTGAAACAGCAGTCAAATCCAGTAAGCTGCAAATCCAT 99045
Gallus_gallus 99001 ............................................. 99045

Anas_platyrhynchos_platyrhynchos 99046 TTTCAATTTATATATGTGTTCAATTATGTTTTACATTTTATGTAT

```
99090
Gallus_gallus                    99046 .............................................
99090

Anas_platyrhynchos_platyrhynchos 99091 AACATTCCTCTAAGCTTCCACATAGAGTGAGCTTGCTTCCACATG
99135
Gallus_gallus                    99091 .............................................
99135

Anas_platyrhynchos_platyrhynchos 99136 GACTGAGCAAGACTTTCCCAAGAACTGCAAAAAGCTATGCTGAAC
99180
Gallus_gallus                    99136 .............................................
99180

Anas_platyrhynchos_platyrhynchos 99181 AGCTTTCTACCGACCTAAGAGGTCCTCTCAGTGGTCTTTATCTTG
99225
Gallus_gallus                    99181 .............................................
99225

Anas_platyrhynchos_platyrhynchos 99226 TAGCCATTCAGCACCAAAGTGATCAGGCTAGGACAGGGAAAAAGC
99270
Gallus_gallus                    99226 .............................................
99270

Anas_platyrhynchos_platyrhynchos 99271 GAAGAAAGTAAAGATAAGGAACCTAAAGAGCTGGATAACAGGATG
99315
Gallus_gallus                    99271 .............................................
99315

Anas_platyrhynchos_platyrhynchos 99316 GGATACTTGTAAAATGAAATGTATATTGAGAGCACAAATGATGAC
99360
Gallus_gallus                    99316 .............................................
99360

Anas_platyrhynchos_platyrhynchos 99361 AGAAAGGAATACAGGGTTTAGAGAAGACTAGAACTAACTGAAGAG
99405
Gallus_gallus                    99361 .............................................
99405

Anas_platyrhynchos_platyrhynchos 99406 ACTGTGGCTGAGAAACAGATAGGAGAAAAAGCATATAATGAAAAG
99450
Gallus_gallus                    99406 .............................................
99450

Anas_platyrhynchos_platyrhynchos 99451 CAAAGGTACAAGAGGAATAGAAATATGGTGAAGAATAAAAAAATT
99495
Gallus_gallus                    99451 .............................................
99495

Anas_platyrhynchos_platyrhynchos 99496 GGGAGAAACTAGAATTTGGTCACATAAGGAAGGGATTTTTGAAAA
99540
Gallus_gallus                    99496 .............................................
99540

Anas_platyrhynchos_platyrhynchos 99541 TCTGTGAGAAAGACACAGTGGAAGCAGAAAAGCAGGAGTGGAACT
99585
Gallus_gallus                    99541 .............................................
99585

Anas_platyrhynchos_platyrhynchos 99586 GGGTGGGCAAGAAGGGGATGCTGGCAAAGGGCTCTGTATAGTGTG
99630
Gallus_gallus                    99586 .............................................
99630
```

```
Anas_platyrhynchos_platyrhynchos 99631 ACAAAACATAATAAAACTCTAGGAGAGATGAGACTGATTTACTGG
99675
Gallus_gallus                    99631 .............................................
99675

Anas_platyrhynchos_platyrhynchos 99676 ATTCTGGGATTGGTGTTAAAAGGTTACTGAGCAATGAGTGGGAAC
99720
Gallus_gallus                    99676 .............................................
99720

Anas_platyrhynchos_platyrhynchos 99721 TGCCCAGATAAAGGGAAAAACAGGATCAAGAGCCTGAGAGGTGAC
99765
Gallus_gallus                    99721 .............................................
99765

Anas_platyrhynchos_platyrhynchos 99766 AGATGTGAAAGTAGAATAGTCACAACTGGTGCATAAGAATGTAAG
99810
Gallus_gallus                    99766 .............................................
99810

Anas_platyrhynchos_platyrhynchos 99811 AGTCCTAGTTGACTAACAAGGTTTTCCCAGGATGTAAACCAAAAG
99855
Gallus_gallus                    99811 .............................................
99855

Anas_platyrhynchos_platyrhynchos 99856 TAAGGACAATTTTATCTGGTTTCTAAACGAAAGATGATTACTAGT
99900
Gallus_gallus                    99856 .............................................
99900

Anas_platyrhynchos_platyrhynchos 99901 GGTAGACCATGTGTAGTTGTGCCGCTCACTGTTGCATAAATGACC
99945
Gallus_gallus                    99901 .............................................
99945

Anas_platyrhynchos_platyrhynchos 99946 TAGGAACACAGCAGTGAACAATCCTCACAACTTACAAAAAGTGTA
99990
Gallus_gallus                    99946 .............................................
99990

Anas_platyrhynchos_platyrhynchos 99991 CCTTAAAAGCACATAAGAACACATGTCACAAGGGAGCCACAGCTG
100035
Gallus_gallus                    99991 .............................................
100035

Anas_platyrhynchos_platyrhynchos 100036 GGATTGCACATGCATGAATGGTTCTGAACTTCTATCAAATTGACA
100080
Gallus_gallus                    100036 .............................................
100080

Anas_platyrhynchos_platyrhynchos 100081 AGAGCAAAGAAGAAAACAGCAAAATTTGGCTGACATTTATAGAAC
100125
Gallus_gallus                    100081 .............................................
100125

Anas_platyrhynchos_platyrhynchos 100126 ATCACAGACCAGTTTGATTAATTTAGAAAGATAACACATGCTCTG
100170
Gallus_gallus                    100126 .............................................
100170

Anas_platyrhynchos_platyrhynchos 100171 AGAAGGCAAAGAAAGCATGGGTACCATGTTAGGGAAAAGTTTCTA
100215
```

```
Gallus_gallus                    100171 .............................................
100215

Anas_platyrhynchos_platyrhynchos 100216 GCTTGAAGGGTAATTCAAAAACATTATTAAAATAAACTATGGGAC
100260
Gallus_gallus                    100216 .............................................
100260

Anas_platyrhynchos_platyrhynchos 100261 GACTATCCATCAGAGAGATTTGTGAACACAAATTACACAAATGCT
100305
Gallus_gallus                    100261 .............................................
100305

Anas_platyrhynchos_platyrhynchos 100306 TAACACCTGCTAAAGCTATTTCAGTTCAGAGCTTAGGTTAGAAAT
100350
Gallus_gallus                    100306 .............................................
100350

Anas_platyrhynchos_platyrhynchos 100351 TGTCAAAGCTAAGTGATGACAGTATGAGGAATAGTGAAATACTCA
100395
Gallus_gallus                    100351 .............................................
100395

Anas_platyrhynchos_platyrhynchos 100396 AGACAGAAAATAGGTCCAACTCCATCTGAAATTCAGAAGGAGGTA
100440
Gallus_gallus                    100396 .............................................
100440

Anas_platyrhynchos_platyrhynchos 100441 AGACCTGCTCATCACGTACAGAGTATAACCAAATACTAACACAAA
100485
Gallus_gallus                    100441 .............................................
100485

Anas_platyrhynchos_platyrhynchos 100486 GGAGCTTTTAGAAGCAGAAGTATACTACTGAGTTGCTACCAGAAA
100530
Gallus_gallus                    100486 .............................................
100530

Anas_platyrhynchos_platyrhynchos 100531 GTTTCATCTCTCATTTCAAAAAGCCACATCCAAAGCACTACATTG
100575
Gallus_gallus                    100531 .............................................
100575

Anas_platyrhynchos_platyrhynchos 100576 AAAGGCAACTGAAGAGAGGAATGAACTGGACGACCTCTCTGAGGT
100620
Gallus_gallus                    100576 .............................................
100620

Anas_platyrhynchos_platyrhynchos 100621 ACCTTTAGTGTATTTTTGAATGAAACATTTTGCAACTGTTTTTCT
100665
Gallus_gallus                    100621 .............................................
100665

Anas_platyrhynchos_platyrhynchos 100666 GCTTACAACTAAATTTGGTCTGAAAATATCATAATGATTTAATAG
100710
Gallus_gallus                    100666 .............................................
100710

Anas_platyrhynchos_platyrhynchos 100711 TGAAAAAGTTAACTTTGAAATTATTGACCTTCTGAAAAACCTTCC
100755
Gallus_gallus                    100711 .............................................
100755
```

```
Anas_platyrhynchos_platyrhynchos 100756 AAGTGTTCATGTTTTTTCCCGAACATATTTGCTATATTAGTTCAA
100800
Gallus_gallus                    100756 .............................................
100800

Anas_platyrhynchos_platyrhynchos 100801 TTCAGCGGATAGCTTTGTTTACTCCCAAACACGTTAATGTACTTC
100845
Gallus_gallus                    100801 .............................................
100845

Anas_platyrhynchos_platyrhynchos 100846 TATTCGATTTTTCCCAGAGTTCTGATGACTGACCTTTGTATGTTT
100890
Gallus_gallus                    100846 .............................................
100890

Anas_platyrhynchos_platyrhynchos 100891 TAAATGACTGAAAGTTTTGCATTAAAAGCAATACATAAAAACCTA
100935
Gallus_gallus                    100891 .............................................
100935

Anas_platyrhynchos_platyrhynchos 100936 GAATGAATACTCTTAAGACGTAACTTAATATAATACAACCTTCAT
100980
Gallus_gallus                    100936 .............................................
100980

Anas_platyrhynchos_platyrhynchos 100981 CATTTCTCACCAAATTATCTGTAGAGAGTGGAGTGTGCAACTGTC
101025
Gallus_gallus                    100981 .............................................
101025

Anas_platyrhynchos_platyrhynchos 101026 CTTGGCACAAATATGCTCTACCAATAAACTGGTTTACAGGAAATC
101070
Gallus_gallus                    101026 .............................................
101070

Anas_platyrhynchos_platyrhynchos 101071 TTCCTTTAAAATACATCTGTAGACAGTCACTGCTAATTTGCAGGT
101115
Gallus_gallus                    101071 .............................................
101115

Anas_platyrhynchos_platyrhynchos 101116 ATCCCAGCTAAGAGAAAAAAAAAAGCATTTTATTTTTTTTTAAATG
101160
Gallus_gallus                    101116 .............................................
101160

Anas_platyrhynchos_platyrhynchos 101161 AATGCTTATTCTAACTGTGTAGCACATAACAGCATTATCATTATG
101205
Gallus_gallus                    101161 .............................................
101205

Anas_platyrhynchos_platyrhynchos 101206 AAAACAATATATCATAAAATTGCATTCCACTAGGCCATTAAGTTT
101250
Gallus_gallus                    101206 .............................................
101250

Anas_platyrhynchos_platyrhynchos 101251 TCTTAAAATTTAAAGGTAGAAATTTTGTCATTCCAAACACTTAAA
101295
Gallus_gallus                    101251 .............................................
101295

Anas_platyrhynchos_platyrhynchos 101296 ATGTTTCCTTTATTTAATACAGCTCCTGAATCACTTAGTTATAGA
101340
Gallus_gallus                    101296 .............................................
```

```
101340

Anas_platyrhynchos_platyrhynchos 101341 ACCCCTTACAGTATACAAAAAAGTATAATTTTTTAGCAGCCATTA
101385
Gallus_gallus                    101341 .............................................
101385

Anas_platyrhynchos_platyrhynchos 101386 ATAGAATTAAGATAAATGCAAAGTTTCTTGTATGATTAAAAAAAC
101430
Gallus_gallus                    101386 .............................................
101430

Anas_platyrhynchos_platyrhynchos 101431 AAAAAGTATGTATATTAGCCTGCAGGATTTTGCATGGATGTCAGC
101475
Gallus_gallus                    101431 .............................................
101475

Anas_platyrhynchos_platyrhynchos 101476 ACTGCACCTCTATTTGCAGACAACAGAAAACACTTGGGTGGCTTG
101520
Gallus_gallus                    101476 .............................................
101520

Anas_platyrhynchos_platyrhynchos 101521 CTGCTTGGAAAAAAATAAGTTACTTTATTAGAAACTATGTCTGCT
101565
Gallus_gallus                    101521 .............................................
101565

Anas_platyrhynchos_platyrhynchos 101566 TCAATTCCAATCACTTATACCCATGGATTGAGAATTATAAAATAC
101610
Gallus_gallus                    101566 .............................................
101610

Anas_platyrhynchos_platyrhynchos 101611 TTCAAAATTAAGAACCTCTACCATATGTTATGGAAGTCAGCTCAA
101655
Gallus_gallus                    101611 .............................................
101655

Anas_platyrhynchos_platyrhynchos 101656 AAAGCGACAGTTTTTCATAATTAGAATGCAATTCTTCTTTTGGTC
101700
Gallus_gallus                    101656 .............................................
101700

Anas_platyrhynchos_platyrhynchos 101701 CACTAGGGACGCTTCATAAACAGTTTATATTGTTGACAACAAGGG
101745
Gallus_gallus                    101701 .............................................
101745

Anas_platyrhynchos_platyrhynchos 101746 AAAAAAAAAAAAGACATAGAAAACTATTTCATATTACAATGGTAT
101790
Gallus_gallus                    101746 .............................................
101790

Anas_platyrhynchos_platyrhynchos 101791 GCTTTATATATTTTGGACTAAATATAAACTTTATAAACCACAGGT
101835
Gallus_gallus                    101791 .............................................
101835

Anas_platyrhynchos_platyrhynchos 101836 TACAGATTATAAAAAAGTGATGATTCAGGGAAGAGGCTGAGTTTT
101880
Gallus_gallus                    101836 .............................................
101880

Anas_platyrhynchos_platyrhynchos 101881 TAACAAGGAATGCATGCTAAAACAATTGCTCTGCAAAAAAAAGGA
```

```
101925
Gallus_gallus                    101881 .............................................
101925

Anas_platyrhynchos_platyrhynchos 101926 AAGGAAAGTAATATTTAAGTAATAACTTTAACACAGGTAAACTAA
101970
Gallus_gallus                    101926 .............................................
101970

Anas_platyrhynchos_platyrhynchos 101971 CTAAATTACAAACCTGAAATGCCTGTTCAGCACACAGAACGTATA
102015
Gallus_gallus                    101971 .............................................
102015

Anas_platyrhynchos_platyrhynchos 102016 AGTCTGTGCTGAAGTTATCTGATGAGTCAAGTGCAGATCTTCCTT
102060
Gallus_gallus                    102016 .............................................
102060

Anas_platyrhynchos_platyrhynchos 102061 GACAGGCTGATTTGATCAACTCATAAGCATCCTTCAAAGCCTGAA
102105
Gallus_gallus                    102061 .............................................
102105

Anas_platyrhynchos_platyrhynchos 102106 CATCTAAAGTAATAGATAATTTATGATTCAAGATGGACCAATAGA
102150
Gallus_gallus                    102106 .............................................
102150

Anas_platyrhynchos_platyrhynchos 102151 CATGTAATTGCATTATTCAGATCAACTGTGTAACACCGTCTTCCT
102195
Gallus_gallus                    102151 .............................................
102195

Anas_platyrhynchos_platyrhynchos 102196 CCTTTAAAGAAAGAAATTACCACAGAATCAAAAATGGATGGGTTG
102240
Gallus_gallus                    102196 .............................................
102240

Anas_platyrhynchos_platyrhynchos 102241 GCGGGGACCCTACAGCCCACCCGGGTCCCCCCATGCCATGGGCAG
102285
Gallus_gallus                    102241 .............................................
102285

Anas_platyrhynchos_platyrhynchos 102286 GACCCCTGCCCCCAGCGCAGGCTGCCCAAAGCCCCCCAGCCCGGC
102330
Gallus_gallus                    102286 .............................................
102330

Anas_platyrhynchos_platyrhynchos 102331 CTTGGGCACTGCCAGGGGAGGGGCACCCACGGCTCCTCTGGGCAA
102375
Gallus_gallus                    102331 .............................................
102375

Anas_platyrhynchos_platyrhynchos 102376 CTGCCCAGAGGTTTTAATTAGGAATTGCCCCAAAATTATGAATTT
102420
Gallus_gallus                    102376 .............................................
102420

Anas_platyrhynchos_platyrhynchos 102421 TTAGGGAAAAAAAGGTCAAGCAGATGTTGTTCAAAGTCAGCAGCA
102465
Gallus_gallus                    102421 .............................................
102465
```

```
Anas_platyrhynchos_platyrhynchos 102466 GTTGTAAAATGAAGCATTAACAACTCTGGTGCAGTTTAAAATCAC
102510
Gallus_gallus                    102466 .............................................
102510

Anas_platyrhynchos_platyrhynchos 102511 TGATGTAGGCAAACAATAAAGTAAATCACTTTGAAGGTCTCAATT
102555
Gallus_gallus                    102511 .............................................
102555

Anas_platyrhynchos_platyrhynchos 102556 GCAACCCGCAGGCCCCAAGAATACGTCCTGCTGAAAAAGCAACTT
102600
Gallus_gallus                    102556 .............................................
102600

Anas_platyrhynchos_platyrhynchos 102601 GGATTTGGGCTGCTTGATGAAGTAAGTTTGACTGGCAGAAAATCA
102645
Gallus_gallus                    102601 .............................................
102645

Anas_platyrhynchos_platyrhynchos 102646 GGCTAAGAAGCACAAATAAAGCGTTTGGGAAAAAAAAAAAAAAAA
102690
Gallus_gallus                    102646 .............................................
102690

Anas_platyrhynchos_platyrhynchos 102691 AGGCCGTACGATTTTTTCGGGACAGGCGCACATCCCGCGGTGCCC
102735
Gallus_gallus                    102691 .............................................
102735

Anas_platyrhynchos_platyrhynchos 102736 GCGGTGCGGGGCCGCAGCCTGACACCCCCTCACCTCGGCGGCTCT
102780
Gallus_gallus                    102736 .............................................
102780

Anas_platyrhynchos_platyrhynchos 102781 GCGCGGCCAGGAGCCTCTCGCGGACGGCCACGTCCATGTCGCCCC
102825
Gallus_gallus                    102781 .............................................
102825

Anas_platyrhynchos_platyrhynchos 102826 CCCCGCCGCTCTGCCAAGCTCGCCGGAGCCGTTTGCCGACGCGTT
102870
Gallus_gallus                    102826 .............................................
102870

Anas_platyrhynchos_platyrhynchos 102871 CTTCCGAGAGCTTTGCCGCACACCCGGCAGAGCAGCCGGGGCACG
102915
Gallus_gallus                    102871 .............................................
102915

Anas_platyrhynchos_platyrhynchos 102916 TCCTCCGTCCGGCGGCCCCGCCGCGCTGCCTCCCTGTGCTCGGCC
102960
Gallus_gallus                    102916 .............................................
102960

Anas_platyrhynchos_platyrhynchos 102961 GCCGTTGCCCGCGCGCGTCCCAGCAACGCTCCCGCCGCGCGCGTC
103005
Gallus_gallus                    102961 .............................................
103005

Anas_platyrhynchos_platyrhynchos 103006 CTAGCAACTAGGTCCTGTCCCCGCCCCGCGGGCGGTGCCGTTCCT
103050
```

```
Gallus_gallus                    103006 .............................................
103050

Anas_platyrhynchos_platyrhynchos 103051 CGGAAAGAGCCCGCCGCGGCTGGTCGGCAGCGCTATTGCTGCGGC
103095
Gallus_gallus                    103051 .............................................
103095

Anas_platyrhynchos_platyrhynchos 103096 GTGCGCCGTGGCAGAAATCCCGCACACTCCGTGAAAGCAAAGGCT
103140
Gallus_gallus                    103096 .............................................
103140

Anas_platyrhynchos_platyrhynchos 103141 CTCGAGCTGTTAATGCGTGCTTACCGCCGCGCACACGCTGTACGA
103185
Gallus_gallus                    103141 .............................................
103185

Anas_platyrhynchos_platyrhynchos 103186 AATCACCGCACAAGCGGCCTCAGGTTAACAAGGGCACCGACCGAT
103230
Gallus_gallus                    103186 .............................................
103230

Anas_platyrhynchos_platyrhynchos 103231 AGCTTCCTGTAGGTAGAACTACTGCTGAGAAGCACTACCAGCAGT
103275
Gallus_gallus                    103231 .............................................
103275

Anas_platyrhynchos_platyrhynchos 103276 AATTTTGGTTGCTGTTTATAGAGCTTCCAACGTGAAACCTTTTTA
103320
Gallus_gallus                    103276 .............................................
103320

Anas_platyrhynchos_platyrhynchos 103321 AATAGGCCACCAGTGCCTGCAGCTTCGGGCTGGAGCAGAATGAAG
103365
Gallus_gallus                    103321 .............................................
103365

Anas_platyrhynchos_platyrhynchos 103366 TGCCGAGCGATGTTTGGGTCATATCAGCCCACACAGAAGGAAGCT
103410
Gallus_gallus                    103366 .............................................
103410

Anas_platyrhynchos_platyrhynchos 103411 GCAGGCTACAAACAGTGATCTTCCAAAACTCGAGGCTTTTGCATC
103455
Gallus_gallus                    103411 .............................................
103455

Anas_platyrhynchos_platyrhynchos 103456 TTTTTAGTCATTTCACAGTATATCGGCCAAGAACAATTTTATGAT
103500
Gallus_gallus                    103456 .............................................
103500

Anas_platyrhynchos_platyrhynchos 103501 AAAAAAATAACCTTCAGAGAACAACATTGCAGAAAAGATACAGTC
103545
Gallus_gallus                    103501 .............................................
103545

Anas_platyrhynchos_platyrhynchos 103546 AAAACAACACAAAACAAACCAAAAAAAAACACAGAAACAAAATAA
103590
Gallus_gallus                    103546 .............................................
103590
```

```
Anas_platyrhynchos_platyrhynchos 103591 ATCTGTAACAAAATAAATCTGTAAAAATAAATCTGGACATGCTGC
103635
Gallus_gallus                    103591 .............................................
103635

Anas_platyrhynchos_platyrhynchos 103636 CATTTGACAGACCAAATGAAGCACTGAAGCTGTCGTGCTCTCTCT
103680
Gallus_gallus                    103636 .............................................
103680

Anas_platyrhynchos_platyrhynchos 103681 CCCTCCAGCTGCTGAGACGTGGAACGCTGTTTTGCATAACACCGT
103725
Gallus_gallus                    103681 .............................................
103725

Anas_platyrhynchos_platyrhynchos 103726 TGGCATGAAGACTATCTATGGACTTAAACGTGTAAATCCAACATT
103770
Gallus_gallus                    103726 .............................................
103770

Anas_platyrhynchos_platyrhynchos 103771 TAAACGTAAACCCTTTGCACTTTAAGCTAAATTGAAATTCTGCAC
103815
Gallus_gallus                    103771 .............................................
103815

Anas_platyrhynchos_platyrhynchos 103816 TGGTAGGATGTTGCAGCCCTGCGAAGACTGAACGCATCGTGCAGA
103860
Gallus_gallus                    103816 .............................................
103860

Anas_platyrhynchos_platyrhynchos 103861 AGGACAGCATTGCTATTAAGTTTCCGAAGACACTTCTTGCAGTCA
103905
Gallus_gallus                    103861 .............................................
103905

Anas_platyrhynchos_platyrhynchos 103906 TCTATATCTGAATTATGAGAGGCCAGTAGGGTGAAAGCCGATAAA
103950
Gallus_gallus                    103906 .............................................
103950

Anas_platyrhynchos_platyrhynchos 103951 AACGGTAAGTGCATCAAACTCTTATGCTTCAGGCCCAGCTACTGA
103995
Gallus_gallus                    103951 .............................................
103995

Anas_platyrhynchos_platyrhynchos 103996 GTAAGTGACACATACAGTACATTTCAAGACCTTTCCATTTGTGAA
104040
Gallus_gallus                    103996 .............................................
104040

Anas_platyrhynchos_platyrhynchos 104041 GTGCTGCTTCATATGAAACATTACAAATGTATTCAAGAACCTGGA
104085
Gallus_gallus                    104041 .............................................
104085

Anas_platyrhynchos_platyrhynchos 104086 AAGAATTCAGCTTTAGACACCTTCCTATTCTCTAGCTGTTCTCCA
104130
Gallus_gallus                    104086 .............................................
104130

Anas_platyrhynchos_platyrhynchos 104131 GCCTACTCCTGACCACACAGCCCATCAAAAAGAACAAACTGCCTT
104175
Gallus_gallus                    104131 .............................................
```

104175

Anas_platyrhynchos_platyrhynchos 104176 GCTTGCATGAACGTTAGTAAACAAGAACAAAACTCGTGAGACGAA
104220
Gallus_gallus 104176 .............................................
104220

Anas_platyrhynchos_platyrhynchos 104221 CAAAACCTCTGTGCCCTTGTGCATTACAAAATCCTTCAAACTCAC
104265
Gallus_gallus 104221 .............................................
104265

Anas_platyrhynchos_platyrhynchos 104266 TGCTGCTGATCCCTACACACCTGCGTGCAGTGTAGCCCTCCTCTT
104310
Gallus_gallus 104266 .............................................
104310

Anas_platyrhynchos_platyrhynchos 104311 TTGGATTGTGGCACACGGCAGGTCACAGCAAGCCTAAGTACCCTA
104355
Gallus_gallus 104311 .............................................
104355

Anas_platyrhynchos_platyrhynchos 104356 AGTACCTAACGTAACTACACAGACCAATACATGGTCTCATAGAAT
104400
Gallus_gallus 104356 .............................................
104400

Anas_platyrhynchos_platyrhynchos 104401 CACGAGACAGCTTTGGGAGGGAGAGGAAAGAAAACTCTGTAGCGA
104445
Gallus_gallus 104401 .............................................
104445

Anas_platyrhynchos_platyrhynchos 104446 TTTTGTGGCAATTTCTGAAACAGAAATGCATGCCTACGTAGATAT
104490
Gallus_gallus 104446 .............................................
104490

Anas_platyrhynchos_platyrhynchos 104491 TTGCCCTATGCCGTGGCGAATTCTCATCCCAGTGAAATTGACAGC
104535
Gallus_gallus 104491 .............................................
104535

Anas_platyrhynchos_platyrhynchos 104536 AACATTAATATCCTCCGCTTGTTTTGAATTGTTTTTCCATTGTCA
104580
Gallus_gallus 104536 .............................................
104580

Anas_platyrhynchos_platyrhynchos 104581 ACCCTAACGTAACGAGAAAAAAAAAAAAAAAAAGAGGACGTGCTTC
104625
Gallus_gallus 104581 .............................................
104625

Anas_platyrhynchos_platyrhynchos 104626 TGGGGCCACGCCCGTCCCNNNNNNNNNNNNNNNNNNNNNNNNNNN
104670
Gallus_gallus 104626 .............................................
104670

Anas_platyrhynchos_platyrhynchos 104671 NNNNNNNNNNNNNNNNNNNNNNNNNNNNNNNNNNNNNNNNNNNNN
104715
Gallus_gallus 104671 .............................................
104715

Anas_platyrhynchos_platyrhynchos 104716 NNNNNNNNNNNNNNNNNNNNNNNNNNNNNNNNNNNNNNNNNNNNN

```
104760
Gallus_gallus                    104716 .............................................
104760

Anas_platyrhynchos_platyrhynchos 104761 NNNNNNNNNNNNNNNNNNNNNNNNNNNNNNNNNNNNNNNNNNNNN
104805
Gallus_gallus                    104761 .............................................
104805

Anas_platyrhynchos_platyrhynchos 104806 NNNNNNNNNNNNNNNNNNNNNNNNNNNNNNNNNNNNNNNNNNNNN
104850
Gallus_gallus                    104806 .............................................
104850

Anas_platyrhynchos_platyrhynchos 104851 NNNNNNNNNNNNNNNNNNNNNNNNNNNNNNNNNNNNNNNNNNNNN
104895
Gallus_gallus                    104851 .............................................
104895

Anas_platyrhynchos_platyrhynchos 104896 NNNNNNNNNNNNNNNNNNNNNNNNNNNNNNNNNNNNNNNNNNNNN
104940
Gallus_gallus                    104896 .............................................
104940

Anas_platyrhynchos_platyrhynchos 104941 NNNNNNNNNNNNNNNNNNNNNNNNNNNNNNNNNNNNNNNNNNNNN
104985
Gallus_gallus                    104941 .............................................
104985

Anas_platyrhynchos_platyrhynchos 104986 NNNNNNNNNNNNNNNNNNNNNNNNNNNNNNNNNNNNNNNNNNNNN
105030
Gallus_gallus                    104986 .............................................
105030

Anas_platyrhynchos_platyrhynchos 105031 NNNNNNNNNNNNNNNNNNNNNNNNNNNNNNNNNNNNNNNNNNNNN
105075
Gallus_gallus                    105031 .............................................
105075

Anas_platyrhynchos_platyrhynchos 105076 NNNNNNNNNNNNNNNNNNNNNNNNNNNNNNNNNNNNNNNNNNNNN
105120
Gallus_gallus                    105076 .............................................
105120

Anas_platyrhynchos_platyrhynchos 105121 NNNNNNNNNNNNNNNNNNNNNNNNNNNNNNNNNNNNNNNNNNNNN
105165
Gallus_gallus                    105121 .............................................
105165

Anas_platyrhynchos_platyrhynchos 105166 NNNNNNNNNNNNNNNNNNNNNNNNNNNNNNNNNNNNNNNNNNNNN
105210
Gallus_gallus                    105166 .............................................
105210

Anas_platyrhynchos_platyrhynchos 105211 NNNNNNNNNNNNNNNNNNNNNNNNNNNNNNNNNNNNNNNNNNNNN
105255
Gallus_gallus                    105211 .............................................
105255

Anas_platyrhynchos_platyrhynchos 105256 NNNNNNNNNNNNNNNNNNNNNNNNNNNNNNNNNNNNNNNNNNNNN
105300
Gallus_gallus                    105256 .............................................
105300
```

```
Anas_platyrhynchos_platyrhynchos 105301 NNNNNNNNNNNNNNNNNNNNNNNNNNNNNNNNNNNNNNNNNNNNN
105345
Gallus_gallus                    105301 .............................................
105345

Anas_platyrhynchos_platyrhynchos 105346 NNNNNNNNNNNNNNNNNNNNNNNNNNNNNNNNNNNNNNNNNNNNN
105390
Gallus_gallus                    105346 .............................................
105390

Anas_platyrhynchos_platyrhynchos 105391 NNNNNNNNNNNNNNNNNNNNNNNNNNNNNNNNNNNNNNNNNNNNN
105435
Gallus_gallus                    105391 .............................................
105435

Anas_platyrhynchos_platyrhynchos 105436 NNNNNNNNNNNNNNNNNNNNNNNNNNNNNNNNNNNNNNNNNNNNN
105480
Gallus_gallus                    105436 .............................................
105480

Anas_platyrhynchos_platyrhynchos 105481 NNNNNNNNNNNNNNNNNNNNNNNNNNNNNNNNNNNNNNNNNNNNN
105525
Gallus_gallus                    105481 .............................................
105525

Anas_platyrhynchos_platyrhynchos 105526 NNNNNNNNNNNNNNNNNNNNNNNNNNNNNNNNNNNNNNNNNNNNN
105570
Gallus_gallus                    105526 .............................................
105570

Anas_platyrhynchos_platyrhynchos 105571 NNNNNNNNNNNNNNNNNNNNNNNNNNNNNNNNNNNNNNNNNNNNN
105615
Gallus_gallus                    105571 .............................................
105615

Anas_platyrhynchos_platyrhynchos 105616 NNNNNNNNNNNNNNNNNNNNNNNNNNNNNNNNNNNNNNNNNNNNN
105660
Gallus_gallus                    105616 .............................................
105660

Anas_platyrhynchos_platyrhynchos 105661 NNNNNNNNNNNNNNNNNNNNNNNNNNNNNNNNNNNNNNNNNNNNN
105705
Gallus_gallus                    105661 .............................................
105705

Anas_platyrhynchos_platyrhynchos 105706 NNNNNNNNNNNNNNNNNNNNNNNNNNNNNNNNNNNNNNNNNNNNN
105750
Gallus_gallus                    105706 .............................................
105750

Anas_platyrhynchos_platyrhynchos 105751 NNNNNNNNNNNNNNNNNNNNNNNNNNNNNNNNNNNNNNNNNNNNN
105795
Gallus_gallus                    105751 .............................................
105795

Anas_platyrhynchos_platyrhynchos 105796 NNNNNNNNNNNNNNNNNNNNNNNNNNNNNNNNNNNNNNNNNNNNN
105840
Gallus_gallus                    105796 .............................................
105840

Anas_platyrhynchos_platyrhynchos 105841 NNNNNNNNNNNNNNNNNNNNNNNNNNNNNNNNNNNNNNNNNNNNN
105885
```

```
Gallus_gallus                    105841 .............................................
105885

Anas_platyrhynchos_platyrhynchos 105886 NNNNNNNNNNNNNNNNNNNNNNNNNNNNNNNNNNNNNNNNNNNNN
105930
Gallus_gallus                    105886 .............................................
105930

Anas_platyrhynchos_platyrhynchos 105931 NNNNNNNNNNNNNNNNNNNNNNNNNNNNNNNNNNNNNNNNNNNNN
105975
Gallus_gallus                    105931 .............................................
105975

Anas_platyrhynchos_platyrhynchos 105976 NNNNNNNNNNNNNNNNNNNNNNNNNNNNNNNNNNNNNNNNNNNNN
106020
Gallus_gallus                    105976 .............................................
106020

Anas_platyrhynchos_platyrhynchos 106021 NNNNNNNNNNNNNNNNNNNNNNNNNNNNNNNNNNNNNNNNNNNNN
106065
Gallus_gallus                    106021 .............................................
106065

Anas_platyrhynchos_platyrhynchos 106066 NNNNNNNNNNNNNNNNNNNNNNNNNNNNNNNNNNNNNNNNNNNNN
106110
Gallus_gallus                    106066 .............................................
106110

Anas_platyrhynchos_platyrhynchos 106111 NNNNNNNNNNNNNNNNNNNNNNNNNNNNNNNNNNNNNNNNNNNNN
106155
Gallus_gallus                    106111 .............................................
106155

Anas_platyrhynchos_platyrhynchos 106156 NNNNNNNNNNNNNNNNNNNNNNNNNNNNNNNNNNNNNNNNNNNNN
106200
Gallus_gallus                    106156 .............................................
106200

Anas_platyrhynchos_platyrhynchos 106201 NNNNNNNNNNNNNNNNNNNNNNNNNNNNNNNNNNNNNNNNNNNNN
106245
Gallus_gallus                    106201 .............................................
106245

Anas_platyrhynchos_platyrhynchos 106246 NNNNNNNNNNNNNNNNNNNNNNNNNNNNNNNNNNNNNNNNNNNNN
106290
Gallus_gallus                    106246 .............................................
106290

Anas_platyrhynchos_platyrhynchos 106291 NNNNNNNNNNNNNNNNNNNNNNNNNNNNNNNNNNNNNNNNNNNNN
106335
Gallus_gallus                    106291 .............................................
106335

Anas_platyrhynchos_platyrhynchos 106336 NNNNNNNNNNNNNNNNNNNNNNNNNNNNNNNNNNNNNNNNNNNNN
106380
Gallus_gallus                    106336 .............................................
106380

Anas_platyrhynchos_platyrhynchos 106381 NNNNNNNNNNNNNNNNNNNNNNNNNNNNNNNNNNNNNNNNNNNNN
106425
Gallus_gallus                    106381 .............................................
106425
```

```
Anas_platyrhynchos_platyrhynchos 106426 NNNNNNNNNNNNNNNNNNNNNNNNNNNNNNNNNNNNNNNNNNNNN
106470
Gallus_gallus                    106426 .............................................
106470

Anas_platyrhynchos_platyrhynchos 106471 NNNNNNNNNNNNNNNNNNNNNNNNNNNNNNNNNNNNNNNNNNNNN
106515
Gallus_gallus                    106471 .............................................
106515

Anas_platyrhynchos_platyrhynchos 106516 NNNNNNNNNNNNNNNNNNNNNNNNNNNNNNNNNNNNNNNNNNNNN
106560
Gallus_gallus                    106516 .............................................
106560

Anas_platyrhynchos_platyrhynchos 106561 NNNNNNNNNNNNNNNNNNNNNNNNNNNNNNNNNNNNNNNNNNNNN
106605
Gallus_gallus                    106561 .............................................
106605

Anas_platyrhynchos_platyrhynchos 106606 NNNNNNNNNNNNNNNNNNNNNNNNNNNNNNNNNNNNNNNNNNNNN
106650
Gallus_gallus                    106606 .............................................
106650

Anas_platyrhynchos_platyrhynchos 106651 NNNNNNNNNNNNNNNNNNNNNNNNNNNNNNNNNNNNNNNNNNNNN
106695
Gallus_gallus                    106651 .............................................
106695

Anas_platyrhynchos_platyrhynchos 106696 NNNNNNNNNNNNNNNNNNNNNNNNNNNNNNNNNNNNNNNNNNNNN
106740
Gallus_gallus                    106696 .............................................
106740

Anas_platyrhynchos_platyrhynchos 106741 NNNNNNNNNNNNNNNNNNNNNNNNNNNNNNNNNNNNNNNNNNNNN
106785
Gallus_gallus                    106741 .............................................
106785

Anas_platyrhynchos_platyrhynchos 106786 NNNNNNNNNNNNNNNNNNNNNNNNNNNNNNNNNNNNNNNNNNNNN
106830
Gallus_gallus                    106786 .............................................
106830

Anas_platyrhynchos_platyrhynchos 106831 NNNNNNNNNNNNNNNNNNNNNNNNNNNNNNNNNNNNNNNNNNNNN
106875
Gallus_gallus                    106831 .............................................
106875

Anas_platyrhynchos_platyrhynchos 106876 NNNNNNNNNNNNNNNNNNNNNNNNNNNNNNNNNNNNNNNNNNNNN
106920
Gallus_gallus                    106876 .............................................
106920

Anas_platyrhynchos_platyrhynchos 106921 NNNNNNNNNNNNNNNNNNNNNNNNNNNNNNNNNNNNNNNNNNNNN
106965
Gallus_gallus                    106921 .............................................
106965

Anas_platyrhynchos_platyrhynchos 106966 NNNNNNNNNNNNNNNNNNNNNNNNNNNNNNNNNNNNNNNNNNNNN
107010
Gallus_gallus                    106966 .............................................
```

107010

Anas_platyrhynchos_platyrhynchos 107011 NNNNNNNNNNNNNNNNNNNNNNNNNNNNNNNNNNNNNNNNNNNNN
107055
Gallus_gallus 107011 .............................................
107055

Anas_platyrhynchos_platyrhynchos 107056 NNNNNNNNNNNNNNNNNNNNNNNNNNNNNNNNNNNNNNNNNNNNN
107100
Gallus_gallus 107056 .............................................
107100

Anas_platyrhynchos_platyrhynchos 107101 NNNNNNNNNNNNNNNNNNNNNNNNNNNNNNNNNNNNNNNNNNNNN
107145
Gallus_gallus 107101 .............................................
107145

Anas_platyrhynchos_platyrhynchos 107146 NNNNNNNNNNNNNNNNNNNNNNNNNNNNNNNNNNNNNNNNNNNNN
107190
Gallus_gallus 107146 .............................................
107190

Anas_platyrhynchos_platyrhynchos 107191 NNNNNNNNNNNNNNNNNNNNNNNNNNNNNNNNNNNNNNNNNNNNN
107235
Gallus_gallus 107191 .............................................
107235

Anas_platyrhynchos_platyrhynchos 107236 NNNNNNNNNNNNNNNNNNNNNNNNNNNNNNNNNNNNNNNNNNNNN
107280
Gallus_gallus 107236 .............................................
107280

Anas_platyrhynchos_platyrhynchos 107281 NNNNNNNNNNNNNNNNNNNNNNNNNNNNNNNNNNNNNNNNNNNNN
107325
Gallus_gallus 107281 .............................................
107325

Anas_platyrhynchos_platyrhynchos 107326 NNNNNNNNNNNNNNNNNNNNNNNNNNNNNNNNNNNNNNNNNNNNN
107370
Gallus_gallus 107326 .............................................
107370

Anas_platyrhynchos_platyrhynchos 107371 NNNNNNNNNNNNNNNNNNNNNNNNNNNNNNNNNNNNNNNNNNNNN
107415
Gallus_gallus 107371 .............................................
107415

Anas_platyrhynchos_platyrhynchos 107416 NNNNNNNNNNNNNNNNNNNNNNNNNNNNNNNNNNNNNNNNNNNNN
107460
Gallus_gallus 107416 .............................................
107460

Anas_platyrhynchos_platyrhynchos 107461 NNNNNNNNNNNNNNNNNNNNNNNNNNNNNNNNNNNNNNNNNNNNN
107505
Gallus_gallus 107461 .............................................
107505

Anas_platyrhynchos_platyrhynchos 107506 NNNNNNNNNNNNNNNNNNNNNNNNNNNNNNNNNNNNNNNNNNNNN
107550
Gallus_gallus 107506 .............................................
107550

Anas_platyrhynchos_platyrhynchos 107551 NNNNNNNNNNNNNNNNNNNNNNNNNNNNNNNNNNNNNNNNNNNNN

```
107595
Gallus_gallus                      107551 .............................................
107595

Anas_platyrhynchos_platyrhynchos 107596 NNNNNNNNNNNNNNNNNNNNNNNNNNNNNNNNNNNNNNNNNNNNN
107640
Gallus_gallus                      107596 .............................................
107640

Anas_platyrhynchos_platyrhynchos 107641 NNNTGCCGTGCCGTGCCGTGCCGTGCCGTGCCGTGCCGTGCCGGC
107685
Gallus_gallus                      107641 .............................................
107685

Anas_platyrhynchos_platyrhynchos 107686 GGAAGCCGTCCCGGGGCTACCAGGCGCGGACAGGGGCGGGCGGCG
107730
Gallus_gallus                      107686 .............................................
107730

Anas_platyrhynchos_platyrhynchos 107731 GGGCCCCGGCTCGGGCTGACTGCAGGAGCTGCGGCGCGGCCGCCA
107775
Gallus_gallus                      107731 .............................................
107775

Anas_platyrhynchos_platyrhynchos 107776 GTCCCGCCGCCCTGCTTGCCCGCACCGGGGGAACGGCTCGGTTCT
107820
Gallus_gallus                      107776 .............................................
107820

Anas_platyrhynchos_platyrhynchos 107821 GAGCCGGGGCTCGGAAAGGCTGCCGATGGCAGGTGTTCTAGCGGC
107865
Gallus_gallus                      107821 .............................................
107865

Anas_platyrhynchos_platyrhynchos 107866 CCGGGTAAAGCTGGTCTCGACTGCTCTGTGACATGGCTTGTGGTG
107910
Gallus_gallus                      107866 .............................................
107910

Anas_platyrhynchos_platyrhynchos 107911 TGGAAACTAGGGAGTCTTACCTGAGGAAAAAAGAACTCGTTTCCT
107955
Gallus_gallus                      107911 .............................................
107955

Anas_platyrhynchos_platyrhynchos 107956 ACCGGGAGGGAGCCTGCAGAGCAATTACAGACCGAGAGATTCTGT
108000
Gallus_gallus                      107956 .............................................
108000

Anas_platyrhynchos_platyrhynchos 108001 CAGAGAGGAGGGTATTTTCAGCTGGTTGGAGAGTGAAATCTGTGG
108045
Gallus_gallus                      108001 .............................................
108045

Anas_platyrhynchos_platyrhynchos 108046 CAGAGCCCTGAATCTCCAGGTTGTGGTTTTGGAAAGAACCCAGAG
108090
Gallus_gallus                      108046 ...........................TTAGGAAAGAACCAGGAG
108090

Anas_platyrhynchos_platyrhynchos 108091 GAGCTGTAAGACACAGACGCTGAATTTCTTTACCGCTTCTTGTTC
108135
Gallus_gallus                      108091 GAGCTGTAATATGTGGATGCCGAATTCCTTAACCATTTCTTGTTC
108135
```

**Commented [Unknown A2]:** Deletion Ends

```
Anas_platyrhynchos_platyrhynchos 108136 CAAGGAATCTCAGAACGATTTCTAAATAAACAAATAGCGAGTATC
108180
Gallus_gallus                    108136 TGAG---TGTCGGAATGATTTCTAAATATACAGATCGAGTGTATC
108180

Anas_platyrhynchos_platyrhynchos 108181 CACAGAGTTACTACAGTGGGAAAGGAGGCTGTCTATAATTGATTA
108225
Gallus_gallus                    108181 CACAGAGTTACTGCTGTGGGGGAGGTGGCCTTCTGAAATTGATTG
108225

Anas_platyrhynchos_platyrhynchos 108226 TGAACACAAACTGTACAGTAACTTACTTAAGCAGATGAAAAGTAA
108270
Gallus_gallus                    108226 CCAATGCGAATTGTACAGTAACTCCTGTAAGCAGGTTAATGATAA
108270

Anas_platyrhynchos_platyrhynchos 108271 TACTGTGTTTTATTGTTGGCATTTAGTAATGCAAGAAGTGAAATT
108315
Gallus_gallus                    108271 CACTGT----------TGGCCTTTAGTAAGGTAAAAAGTGAAATA
108315

Anas_platyrhynchos_platyrhynchos 108316
CTACAGGAAAAGTCTATTTAACATTCTTTTG-------------- 108360
Gallus_gallus                    108316 CTGCAAGAAAATACTACTTAATTGACTTTTGTGGTGGGGTGGGGG
108360

Anas_platyrhynchos_platyrhynchos 108361 GGGGTGTGTCTGTTTGCAGGTCTGTGAATCAAGTGTGTGTGGAAA
108405
Gallus_gallus                    108361 GGGGTG-GTTTGTTTGCAGGTCTGTGAATCAAGTGTGTGTGGAAA
108405

Anas_platyrhynchos_platyrhynchos 108406 AATCGCTTGTGTTCAGAACTAATACTCCTTATCATACTGATACCA
108450
Gallus_gallus                    108406 AATCACTTATATTCAGAACTAATATTCCTTATCATACTGATACCA
108450

Anas_platyrhynchos_platyrhynchos 108451 TTTCTGTCCCTGGTGAACATGATCTGCTGTTCTGTGTGCAGCTGG
108495
Gallus_gallus                    108451 TCGCTGTCTCTGATGGACCAGAT---CTGTTCTGTGTGCAGCTGG
108495

Anas_platyrhynchos_platyrhynchos 108496 ATGACTGTAAGTAAC---TTTTCTTTCCATTAACTTAATTCATAG
108540
Gallus_gallus                    108496 ATGACCGTAAGTAACTTTTTTTTTTTCCATTAACCTGATTCTAAT
108540

Anas_platyrhynchos_platyrhynchos 108541 ATAGCCTTTTCTTAGGGAATCTCTTTTTAAACTTACATGCTATAA
108585
Gallus_gallus                    108541 ATAGCCTCTGCATAGGGAATCTCTTCTGAAACTT--GTGTTATAG
108585

Anas_platyrhynchos_platyrhynchos 108586 TTGCTAGAAATTATTTCGTTTGTTATAGTCTCTGAGAGACCATAG
108630
Gallus_gallus                    108586 TTGTTACTAATTATTTTGGCTACTGTAGTTCC----ATACCATGT
108630

Anas_platyrhynchos_platyrhynchos 108631 GGAATCACCATCCCATCACTGAAGACAAAAAATTATCATTGCCCT
108675
Gallus_gallus                    108631 TGCATCA.....................................
108675

Anas_platyrhynchos_platyrhynchos 108676 CATTTTGAGGGTAGTGCTCCCTTGGCTAAACACGAATCACTCTTT
108720
```

**Commented [Unknown A3]:** Chicken and Duck TCTN3 Exon2

```
Gallus_gallus                      108676 .............................................
108720

Anas_platyrhynchos_platyrhynchos 108721 AGCAATTGTTTCTGTATCTAAGACACCTTAATTTGCTACACGTAG
108765
Gallus_gallus                      108721 .............................................
108765

Anas_platyrhynchos_platyrhynchos 108766 AGCAGTAAAGAAAGTTTGACATCTGTCATATTTTTCCTGTTAGGA
108810
Gallus_gallus                      108766 .............................................
108810

Anas_platyrhynchos_platyrhynchos 108811 AAGGAAGGATTCCTTTTATCTTCCTATTATCCTTTTCAGAAAGCG
108855
Gallus_gallus                      108811 .............................................
108855

Anas_platyrhynchos_platyrhynchos 108856 GTTTATTATCTTATCTTGGATTACTCATTTGATATACTTATTTTC
108900
Gallus_gallus                      108856 .............................................
108900

Anas_platyrhynchos_platyrhynchos 108901 ATACTTCCTTCTTTGCTGTCACTGATTTGCAGTGATCACATACCC
108945
Gallus_gallus                      108901 .............................................
108945

Anas_platyrhynchos_platyrhynchos 108946 TAGTAGGGCATGTCAATAATGTTTTTTCTCCTTCTGCTTTCCTCA
108990
Gallus_gallus                      108946 .............................................
108990

Anas_platyrhynchos_platyrhynchos 108991 ATGCGTTTTGTTTGTCTGATTTGGTGTCCAGCTCACTGGTATTCA
109035
Gallus_gallus                      108991 .............................................
109035

Anas_platyrhynchos_platyrhynchos 109036 TTGAGATTGCTTAGGAGCTCATGTTTGAACACATACTATTTTCTC
109080
Gallus_gallus                      109036 .............................................
109080

Anas_platyrhynchos_platyrhynchos 109081 TTTTGACACTCGTGTCTGGGAGTTTTGATCAACAGTCCCATCTAC
109125
Gallus_gallus                      109081 .............................................
109125

Anas_platyrhynchos_platyrhynchos 109126 TTCGGCCTTGTGTCATAAAATCAGAGAACATTCTAGGCTGGGAGA
109170
Gallus_gallus                      109126 .............................................
109170

Anas_platyrhynchos_platyrhynchos 109171 GGCTTAGGAGGTTACCTGGTCCAGGTACTAGTCAAAGAAGGGTCC
109215
Gallus_gallus                      109171 .............................................
109215

Anas_platyrhynchos_platyrhynchos 109216 TGTAATTTAGTCCACCTTTCAAATCCTTGCAGATGTTACCTGAAT
109260
Gallus_gallus                      109216 ................TCTTAAGTGCTTATGAAAGGTGTCT---T
109260
```

```
Anas_platyrhynchos_platyrhynchos 109261 GTTTGTATGTGTTAAAAAGTGT---GTTCAAACAAATCCATCAGG
109305
Gallus_gallus                    109261
ATTTGCATGCATTTAAAAGCTTACTGTTTTAACAAGCCTAT---- 109305

Anas_platyrhynchos_platyrhynchos 109306 ATGAATCTCTTTAGTATGTCCTTAATCGTTTGACTGAATTATATG
109350
Gallus_gallus                    109306 ----------GTGTTATGTCATTAATCATTTGTC---ATTTTATG
109350

Anas_platyrhynchos_platyrhynchos 109351 ACATAGTGCCTTAGCAGTAATAATTTAAAACATTGTAATACTGTA
109395
Gallus_gallus                    109351 ACTTAGTGTTTTAGCAGTAATAATTTAATACATTATAGCATTGTG
109395

Anas_platyrhynchos_platyrhynchos 109396 ACACTACTGAATGGTGCTGGGACAAGAAGACATCACTGTGAGATA
109440
Gallus_gallus                    109396 ACAGTTCTGAATATCAACTGGCAAGGAAGACATTACTCTGAGATA
109440

Anas_platyrhynchos_platyrhynchos 109441 CCAGTAGAATTTTGTCTTTTTAGGATGTCTATTCACAATTCATAC
109485
Gallus_gallus                    109441 CGAGTGGGAGTTTGTCTTTTAAGGATATGTATTCACAGTTCATAC
109485

Anas_platyrhynchos_platyrhynchos 109486 TTACTTTAAAGTTTATCAGCCTTAATTACTTTTTAATCCATTTAA
109530
Gallus_gallus                    109486 TTAAGTTAAAGTTTATTTGGC---ATTA-TTTTTAATCAATTTAA
109530

Anas_platyrhynchos_platyrhynchos 109531 TGTTGCTTTTACTAAAT--GTATACTGCTGGTGTTTTGTCATTAA
109575
Gallus_gallus                    109531 TGTTGTTTTTACTGATTTCACATGCTAATAGTGTTTTGACATAAA
109575

Anas_platyrhynchos_platyrhynchos 109576 AACATCATTAAAACATCGTATGGGTCCAACTTGGGCATCTTACAA
109620
Gallus_gallus                    109576 AATGTCA-----------GATGACTGTAACTTGGACACCTCACAA
109620

Anas_platyrhynchos_platyrhynchos 109621 AAATGAGAGCACATGTTAAACTAGTTGCCTTAAGCAGTTTTCACA
109665
Gallus_gallus                    109621 ATG-GAGAGCACAT--TAAGATAGTTGCCTTAAATAACTTTCATT
109665

Anas_platyrhynchos_platyrhynchos 109666 ATTTCAGAAAATCACATATGTGTATCAACACGTTTCTCAAGAAAC
109710
Gallus_gallus                    109666 ATTTCTCAAGATCAGATTTGTATATGAACACGTTTTTCATGCAAC
109710

Anas_platyrhynchos_platyrhynchos 109711 AGTACAAAGTAGTTCCCATGATATCATCCTCCTGTTCATAGCTCG
109755
Gallus_gallus                    109711 ATTACGAATTACTACTCATTATGTCA---TCTTGTTCAAAGCTGG
109755

Anas_platyrhynchos_platyrhynchos 109756 TATTAGTCCGTTAGCAATCTTACTAAGGATTAATTACTGTAAGGC
109800
Gallus_gallus                    109756 TATTACTGTGT.......CCTGTTAGCAATTAATTAATATATGGC
109800

Anas_platyrhynchos_platyrhynchos 109801 ACTATCTAGCTTATAATTACTGAAGTTGTCCCTCTTAAAAATATT
109845
Gallus_gallus                    109801 ACTATATAGCTTATAATTAATAAAATTGTCACTTTTAAAA-TATT
```

109845

Anas_platyrhynchos_platyrhynchos 109846 AGCACAATGTTTACTTTTTCATAGAACTATCTTTTTTAGTAATTT
109890
Gallus_gallus 109846 GGCA---TGTTTACTTTTT---AAAAAT-TCTTTTCTAGTAATTT
109890

Anas_platyrhynchos_platyrhynchos 109891 TA-CGTTGGTAGATT---------TTTTTTTTTTCACCTTTTAGC
109935
Gallus_gallus 109891 GAATGTTGGTAGATTATTATATATTTTTTTTTCCCAACCTTTAGT
109935

Anas_platyrhynchos_platyrhynchos 109936 AAAGCTGAACTATTTCCAGCAGCCACAGCCTATCAAAGAATCTGA
109980
Gallus_gallus 109936 GAAGTTGAACTATTTCCGACAGCCAAAGCGTATCAAAGAATCTGA
109980

Anas_platyrhynchos_platyrhynchos 109981 TTTTCTGGAGTTCTCAGAGAAGTACGGCAGACTCTCTTTCTTAGC
110025
Gallus_gallus 109981 TTTTCTGAAGCTCTT---GAAGTATAGCAAGCACTCTTTCTTAGC
110025

Anas_platyrhynchos_platyrhynchos 110026 AGTATCACAAGTCCAGCCATCGTTTTCTGCCTTTTACCGGGTGAG
110070
Gallus_gallus 110026 ACAATCACAAGTCCAGCCATCATTCTCTGCCTTTTACCGAGTGAG
110070

Anas_platyrhynchos_platyrhynchos 110071 TTGAAAAAAAGATAGCAACCAGTTTGCAGATATGCCTTTGCTATT
110115
Gallus_gallus 110071 TTGAAAGAAA-ATACCAACTGGCTTGCAGGTATACCTTTGCTGTA
110115

Anas_platyrhynchos_platyrhynchos 110116 GATGTCTGTAATGAGACCTAACGAGAACAGAATTAAATATTCCTC
110160
Gallus_gallus 110116 GCTGTCTGTAATGAGACCTAATGAGAACAAATTCAAAGATACTTT
110160

Anas_platyrhynchos_platyrhynchos 110161 ATGTATTTTTCCCCTATAAACCACGGTATTATCAGTTCTTTTCAT
110205
Gallus_gallus 110161 GTATTTTTCTATCCTTTGAACCA---TATTATTTTTCATTTCTTT
110205

Anas_platyrhynchos_platyrhynchos 110206
TCCTTGTTGTGTAAAAAGTTTCTAATTAGTTTTCTTGTATGTA-- 110250
Gallus_gallus 110206 TCCTTACTGTGTAAAATGTTCCTAATTAGTTTTCTTGTATATAAA
110250

Anas_platyrhynchos_platyrhynchos 110251 -GTTTGGACA---TTTTAGATAAAACTGGCTGTGCACTAAGGAGA
110295
Gallus_gallus 110251 AGTTTGTACAGCCCTTTAAACAAAACTGACTGTATGCAAAGGAGA
110295

Anas_platyrhynchos_platyrhynchos 110296 AGTTTCAGTTTGTCTCTCTGCCCTGAGGATGTCGTTACCTTGCAC
110340
Gallus_gallus 110296 AGTTAAAGTTTGCCTCTTTGCCCTGA---TGTAATCACCCTGAAC
110340

Anas_platyrhynchos_platyrhynchos 110341 TTGGACTTACTATTCCACAGTGATGTTGTTTTTAGTTTCTCCGAA
110385
Gallus_gallus 110341 TTAGTCTTACAATTCCACAGTAATGTTGTCTGTAAGTTTTCCAAA
110385

Anas_platyrhynchos_platyrhynchos 110386 TTGTCTTTTCAGGCTGGTGATCCAATCCTAATCTATTTTGATTCA

```
110430
Gallus_gallus                     110386 TTGTCTTTTCAGGCTGGCGATCCAATCCTAATCTACTTTGATTCA
110430

Anas_platyrhynchos_platyrhynchos 110431 TCATCTACACTGAGTATGCTTAGACAACCAGTGAAAATGGGAGTC
110475
Gallus_gallus                     110431 TTATCTGTACTGAGTGTGCTTAGACAGCCAGTAAAAATGGGAGTC
110475

Anas_platyrhynchos_platyrhynchos 110476 AGTGGACTCTGTGTTGATGGAAATCCTGCTGGTTAGTAAAGGGGA
110520
Gallus_gallus                     110476 AGTGGACTTTGTGTAGATGGAAATCCAGCTGGTTAGTAAAAGGAA
110520

Anas_platyrhynchos_platyrhynchos 110521 AAGCGTTTCTAAGGGCTGTGTGCTTCTGTTGTGTCTTAGCTGACA
110565
Gallus_gallus                     110521 AGGCATTTCTGAGGTTTATGTGCTTCTGTTTTCTCTTATTTGATG
110565

Anas_platyrhynchos_platyrhynchos 110566 ATGTCTCTTAACTCAGGCTTCCTGGAAAGTAAAAGCACAAGCTGT
110610
Gallus_gallus                     110566 TCATCTCTTCACTCAGGCTTTCTGGAAAGTAAAAGCACAAGCTGT
110610

Anas_platyrhynchos_platyrhynchos 110611 ACTCGGATTTTCACCAACTTGAGCAAGAGCTGCATTACTGACCCT
110655
Gallus_gallus                     110611 ACTCGCATTTTCACAAACCTGAGTGGGAGCTGTGTCAGTGACCCT
110655

Anas_platyrhynchos_platyrhynchos 110656 GCCCTAGATGCTGCCTCATACTACCGTGACTTCACAGTGCTAAAG
110700
Gallus_gallus                     110656 GCCCTAAATGCTGCCTCCTACTACCGTGACTTCTCAGTACTAAAG
110700

Anas_platyrhynchos_platyrhynchos 110701
GTGCATGTCGTATTTCTTCTTTTGCTAAGCAT------------- 110745
Gallus_gallus                     110701 GTGTGTCT----TTTTTCCTTTTGTTCAGTGCCATGTGTGCTTGA
110745

Anas_platyrhynchos_platyrhynchos 110746 CTTTCCATTGCCTTCCATTGATACTTCAGATACC--AAATCACTC
110790
Gallus_gallus                     110746 CTTAGCATTGTCATCCATTTCAACATCACGTAACTTGGAAAAGGT
110790

Anas_platyrhynchos_platyrhynchos 110791 TGTTTTCAAGTTATGTAGTACCCCTTGATCTTCTGATTATTATAC
110835
Gallus_gallus                     110791
TATTT----GGTGTGTAGTATCCCTCGATCTTCTGATCATG---- 110835

Anas_platyrhynchos_platyrhynchos 110836 CCCTCTTGCTAGAAATCCTGTTAAAAATGTCCCAGTAATCAGTAA
110880
Gallus_gallus                     110836 --CCCTTGCTAGAAATCCTGTTGAAAATGTCCC--TGATCAGTGA
110880

Anas_platyrhynchos_platyrhynchos 110881 TAATCCATCACTTTTCTTTTTTTTCTTTTACACCCAAGTACTTTT
110925
Gallus_gallus                     110881 TAAACCATCACTT--------------------CAATGTGCTTTC
110925

Anas_platyrhynchos_platyrhynchos 110926 CAATAAACTTCAGTTATTATCAAGTTATTAATGCATAGAACAGGT
110970
Gallus_gallus                     110926 CCATAAACTTCAGTTATTAGCAAGTACTTAGTGCATATAACAGCT
110970
```

Anas_platyrhynchos_platyrhynchos 110971 ACTTCATCAATATTTCATAATTTGAACAGTTTAACCACAACTCTC 111015
Gallus_gallus 110971
ACTTCATGGATATTTCATATTCT--ATGGTTTAAGTACAAATCT- 111015

Anas_platyrhynchos_platyrhynchos 111016 TTACATTTTAAACAGACTGTTTAT-GAAAGTATTTTGATC---TC 111060
Gallus_gallus 111016 TACCTTTTCAAATAGACCATTTATCAAAAATATTTTCTCCCTGTC 111060

Anas_platyrhynchos_platyrhynchos 111061 TTCCGTCAACTTGATCAACCAAGCAAAGCATATTCTTAGGAAAAA 111105
Gallus_gallus 111061 TTCCATCAACTTGATCAATGAAGTAAAGCATATTCTCAGGAAATA 111105

Anas_platyrhynchos_platyrhynchos 111106 GTAAGGTGTTTTTTTTTTTGATACAACATGTTCTCTCTAAAATCA 111150
Gallus_gallus 111106 ATTAGGGTTTTTTTTTTTTGTTACAATCTGTTCTCTGTAAAATCC 111150

Anas_platyrhynchos_platyrhynchos 111151 CTTTGGAAGCCAGTT---ATATTATCTTTATTTTGCACAATCACA 111195
Gallus_gallus 111151 TATCTAATGCCAGTTGTCATATAAGCTGTATTTCACAGAATCACA 111195

Anas_platyrhynchos_platyrhynchos 111196 AAATGGTTAAGGTAGGAAGGGACTCCTGGAGGTCATCTAGTCTAA 111240
Gallus_gallus 111196 GAATGGTTGAAGTTGGAAGGGACCTCTATAGGTCATCTCTTCTAA 111240

Anas_platyrhynchos_platyrhynchos 111241 CCAACCCCCACTCAAGCAGGGACACCAAGAGCAGGTTACCCAGGA 111285
Gallus_gallus 111241 CCTG.TCATGCTCCAGCAGAGGCACCTAGAGCAGGGCGCCCAGGA 111285

Anas_platyrhynchos_platyrhynchos 111286 CCACATCCAAGCAGCTTTTGA------------------GTCTCC 111330
Gallus_gallus 111286 CCATGTCCTGGTGGCTTTTGAAGACCACCAAGGCGGAGAGTCCAC 111330

Anas_platyrhynchos_platyrhynchos 111331 TTCCT------------------------------AAGGAGGAAGC 111375
Gallus_gallus 111331 CACCTCCGTGGGCAACTTTTGCCAGTGCTCCATCAACAAGACAGC 111375

Anas_platyrhynchos_platyrhynchos 111376 GTTTAGATG----------------------------TGTTCCAG 111420
Gallus_gallus 111376 ATGCAAATGCTTCCTCATGATCAGATGGAACCTCTTGTGTTCCAG 111420

Anas_platyrhynchos_platyrhynchos 111421 TTTGTGCCCATTGCCTCTTCTCCTGGCACCGGGCACCACTGATAG 111465
Gallus_gallus 111421 TTTGTGTCCACTGCCTCTTGCCCTGGTATTGAGCACCTCTGAAAA 111465

Anas_platyrhynchos_platyrhynchos 111466 GTGCATGTCTCTGTCCCCTTTGCAACCTACCCTCTGGGTATTTAT 111510
Gallus_gallus 111466 GAGCACAGCTACATCCTCTTTGCAGCCT-CCCTTCAGGTATTTAT 111510

Anas_platyrhynchos_platyrhynchos 111511 ATACACTGATGAGATTTCCCCCAAGCTTCCTCTTCTCCGGACTGG 111555

```
Gallus_gallus                     111511 ACACATTGTTCAGACCTGCCCTGAACCTCGTCTTCTCCAGTCTGA
111555

Anas_platyrhynchos_platyrhynchos 111556 ACAGTCCCAGCTCTCCCAGCCCCTCCTCATAGGAAAGGCGATCCA
111600
Gallus_gallus                     111556 ACAGCCCCAGCTCTCCCAGCTTCTCTTCACAGGAGAGGAGCTCCA
111600

Anas_platyrhynchos_platyrhynchos 111601 GTTCTTTGATAATCTCGATGCCTTTCCACTGGGCTCTTTCCAGCA
111645
Gallus_gallus                     111601 GTTCCTTCACCATCTTGGTGCCCTTCCATTGGCCTCTTTTCAGTA
111645

Anas_platyrhynchos_platyrhynchos 111646 TATCTGG-TCTCTTGTGTTCCGGGGGGCCCAGAACTGGACCCAGC
111690
Gallus_gallus                     111646
TGCCCAGATCTCTGTTGTTCTGGGA-------------------- 111690

Anas_platyrhynchos_platyrhynchos 111691 CCTTCCGTTGTGGCCTCACCAATGCTGAGTAGAGGGGAACGGATC
111735
Gallus_gallus                     111691
CCCTCTAATGTAGATGCA--------------------------- 111735

Anas_platyrhynchos_platyrhynchos 111736 ACTTCTCTTGACCTGCTGGCAAGACTTTGCTTAATGCAGCTGAGA
111780
Gallus_gallus                     111736 ACCTCTCTTTACTTGCTGGTGATACTTTGCTTAATGCTGCTGTGA
111780

Anas_platyrhynchos_platyrhynchos 111781 ATACCGTTAGCCTTCTTA-GCCAAGGGTGCATTTGCTGATTCATG
111825
Gallus_gallus                     111781 ATAACATTAGCATTCTTGTGAGAAGGGCACATT-GCTGACTCATG
111825

Anas_platyrhynchos_platyrhynchos 111826 TTTGACTTGGTGGCCATGAGGA-CTGCTAGATCTCTTTCTGCAGA
111870
Gallus_gallus                     111826 TTCTACTTGGTGTCTATTAGGGTCTGCTATATCCTTTTCTGTAGT
111870

Anas_platyrhynchos_platyrhynchos 111871 GCTGCTTTTGTACATTCATAGATCGAATTTCATATCAATTTTCCC
111915
Gallus_gallus                     111871 GCTGCTATCATATATTCTAAGATTGAATTCCATATCAATTTCTAC
111915

Anas_platyrhynchos_platyrhynchos 111916 ATTATTTATGCTGAGAATAATGTCAAAATAACCAGCTTCTTATTC
111960
Gallus_gallus                     111916 TTTATTAATTCTGAAAAAAATGCCAAATTG-CTAGTTTATT--TT
111960

Anas_platyrhynchos_platyrhynchos 111961
TGGGTTCATCATGCTGGTCTTGCTGAATGTTAGCATAAGAACGT- 112005
Gallus_gallus                     111961 TGTGTTCATTGTGTTAGCTTTACTGAGGGATAGCTTTATCTCATT
112005

Anas_platyrhynchos_platyrhynchos 112006 --------CTCACCACTTTCTCTGGTATTCCAAGATAAAAGAGCA
112050
Gallus_gallus                     112006 CATTTCTTCTTAACAGTTTCTCTGTTATGCTTAGACAATACAATG
112050

Anas_platyrhynchos_platyrhynchos 112051 ATATAAATATTAACACCTTATTAAGACTAATGTCCCAGTTTACCC
112095
Gallus_gallus                     112051 -TGTCAGTATTAGTATCTTATTAAAGTTAATGGCTCTGCTTACGC
112095
```

Anas_platyrhynchos_platyrhynchos 112096 TTCAGTTGGAAATACTCTTCTAAATCTATATAATTGTAGTA-GAG 112140
Gallus_gallus 112096 TTTGATTGGAAATATTCTTCTAAACCTGCATGACTGTAGTAAGAG 112140

Anas_platyrhynchos_platyrhynchos 112141 AGTATTGCATTGACCAAATTTAACTCATGTTTTCAGGGCTTGCTT 112185
Gallus_gallus 112141 AGTATTCCACTGACCAAATGTAATTCATGTTT-CTGTGCTTACTT 112185

Anas_platyrhynchos_platyrhynchos 112186 TGTTACTGAG-----------------CCCCAAATAAAAA----- 112230
Gallus_gallus 112186 GGTTACTGATGGCAAAACTTAATTTTTCCTTAAGTGGAAACTAGA 112230

Anas_platyrhynchos_platyrhynchos 112231 -------------------------------------------AA 112275
Gallus_gallus 112231 GTAGGTACTTCATTCCACAAGGGGGAGCTCCAGGAATGGATTCAA 112275

Anas_platyrhynchos_platyrhynchos 112276 AAT--------------------------AAGGCTAATAATAAAA 112320
Gallus_gallus 112276 GATGCTAGTAATATCTTCAGTTTGTTGACAGGGCCAGGAATAAAT 112320

Anas_platyrhynchos_platyrhynchos 112321 GTTTTCTTTTGGTCT----------AAAAATATATTCAG---TAC 112365
Gallus_gallus 112321 GCTGTTTTCTGTTCTTCTAAAAAAAAAAAAAAAAATCCCACTCTAT 112365

Anas_platyrhynchos_platyrhynchos 112366 AGTATGTCTGTGATTTCGAGAACACAAATGGATGACAAACTTTAC 112410
Gallus_gallus 112366 ATTATGTCTGTAATTTC-----TATCTATGGATGACAAATTTTAC 112410

Anas_platyrhynchos_platyrhynchos 112411 ATCTGTGGAATAATC--AGTCATTCTTGTCAATGTAATATCTAAG 112455
Gallus_gallus 112411 ATACCTTAAGTAAGCCAAGTGACTTTTGTCACTGAAATACCTAGG 112455

Anas_platyrhynchos_platyrhynchos 112456 GTCTTTTTAATGGTGGCTTCATGACTTTAGGCAGTTGTAAACCCA 112500
Gallus_gallus 112456 ATGTATCT--TGTTTGTCTTTTGATTTTAC----TTGCTAACTCA 112500

Anas_platyrhynchos_platyrhynchos 112501 GCA----AAACTGTTGTTTTTTGGTGGTGTTTGTTGGTTTGTTTG 112545
Gallus_gallus 112501 GCACCAAAAACAACTAGATTTTGTT-----TTTTTGGTTTGTTTG 112545

Anas_platyrhynchos_platyrhynchos 112546 TTTTCCCCAGGTTCCAATCGATGGCACTCTTGTGGAGTCCATGAA 112590
Gallus_gallus 112546 TTTTCCCTAGGTTCCGATCAATGACACCATTGCAGAGTCCATGAA 112590

Anas_platyrhynchos_platyrhynchos 112591 GGTCAGTCTCTATTGCTAATGAAAGTAATTTCTGCTTCTGAAATA 112635
Gallus_gallus 112591 GGTCAGTCTCTGTTGCCAGTGAAAGTAACTTCTGCTTCTGAAATT 112635

Anas_platyrhynchos_platyrhynchos 112636 AAACAT---------------ATCTGGAATATTAGGCATTCATAG 112680
Gallus_gallus 112636 AAGTATTGGGAACAGTAGAGTATCGGGAATAGTAAACATTGGTAA

112680

Anas_platyrhynchos_platyrhynchos 112681 TAAGGCCCATGAGTGAGAGTGGAAAGCCAACTTGCGAGAGTGGAA
112725
Gallus_gallus 112681 TAATGCTCATGGGTTCAG----------AACTTGAAAGAGCGGGA
112725

Anas_platyrhynchos_platyrhynchos 112726 ATATTTTGTGCCTGAAACAAGACTTTTTGTAAG----TGTGACAT
112770
Gallus_gallus 112726 ACATTCGGTGTCTGAAATAAGACTCCTTGTAAGGAAGTGTCACAT
112770

Anas_platyrhynchos_platyrhynchos 112771 ---AGACAAAA-TACAGGGTTATAAGGCAAGGTGTACTTTTAAAT
112815
Gallus_gallus 112771 TACAGACAAAAATACAGGGGTATAAGGCAGCATGTGCTTTTAAAT
112815

Anas_platyrhynchos_platyrhynchos 112816 GATTGAGCATGGGTAGGGTTGGTGTTAGGGAAATAGCTTGTTCTC
112860
Gallus_gallus 112816 GATTGAGCACAGATGGCATTGGAGTCAGGGAGGTAGCTTTCTTTT
112860

Anas_platyrhynchos_platyrhynchos 112861 TT-------------AGGAAAAACTG-------GGCTTCTGGAAC
112905
Gallus_gallus 112861 TTTAAAAACAACAACAACAAAAACAGTCAAAATGGCATCTGGAAC
112905

Anas_platyrhynchos_platyrhynchos 112906 AGAGTGACATATCCCCAGATTTCAGCTACAGCAGGAAAGTAAATC
112950
Gallus_gallus 112906 AGTGTGATGTTTCTCCAGATCACAGTTACAGCAGGAATGTAAATC
112950

Anas_platyrhynchos_platyrhynchos 112951 CAGCTTTGACAGAAAAGCAGTGGAAAAGAGTAAAAAAGGCACACT
112995
Gallus_gallus 112951 ----TTGGTTGGAAAAGCAGTGGAAAAGAGTAA----GATATGCT
112995

Anas_platyrhynchos_platyrhynchos 112996 CTGTTTAAACAGTTGCTGTTTGTTAATATGCTTAATATGTTTCCA
113040
Gallus_gallus 112996 CTTTTTAAACAACTGCTATTCATTAATATCTTTACT----CTCCA
113040

Anas_platyrhynchos_platyrhynchos 113041 GGTGAAGATCATCCCAGTCACACCACCTAGAGCTCCCTACATGAA
113085
Gallus_gallus 113041 GGTAAGGATCAGTCCAATTGGACCACCTGGAGCTCCCTACATGGA
113085

Anas_platyrhynchos_platyrhynchos 113086 AGACAACACATGTAACAATGCTGTTTCTGAGGTATGTCAACCTGT
113130
Gallus_gallus 113086 AGATAACGTGTGTAACAATGCTGTTTCTGAGGTATCTCGACCTTT
113130

Anas_platyrhynchos_platyrhynchos 113131 TTTCCTTAGGGCAGCTCAACACCAGCTTGAAGTCATTACCCGTAT
113175
Gallus_gallus 113131
TTTTCTTAGG----TTCAATATGAACTTCAAGTTGTTACCC---- 113175

Anas_platyrhynchos_platyrhynchos 113176 AAAGTCATTACCTATATAAAGTCATTACCTGACTATGTCTCTGAT
113220
Gallus_gallus 113176 ----------------------TATTAC-----TGTGTCTCTGAT
113220

Anas_platyrhynchos_platyrhynchos 113221 TCTTCTCTGAATTGGACTCTCAGGTGACCTATGAAATAGAATTCA

113265
Gallus_gallus 113221 TCTTCTCTGGATTGGACTTTCAGGTGATCTATGAGATAGAGTTTA
113265

Anas_platyrhynchos_platyrhynchos 113266 ATGGCACCCATGGGATTCAGAGTGTGTCTGTCCGGTTCAAAGTGA
113310
Gallus_gallus 113266 ATGGTACCTATGGGATTCAGAGTGTGTCTGTCCAGTTCAAAGTGA
113310

Anas_platyrhynchos_platyrhynchos 113311 CCAACATCTCTGGGAACTCAGGAGTCTCTCTGCAGCAGCA---CT
113355
Gallus_gallus 113311 CCAGCATCTCTGGGAGTTCTGGATTCTCACTGCAGCAGCAGCACT
113355

Anas_platyrhynchos_platyrhynchos 113356 TCACTTTGCACTTCTGGGTCAGTGCCCCCATCCACCCTGAGCTCT
113400
Gallus_gallus 113356 TCACTTTGCGTTTCTGGGTCAGTATCCCAGTCT---TTGGGCTGT
113400

Anas_platyrhynchos_platyrhynchos 113401 TGCCAAGTTTGTGTTCTGTAGCATGTGAACTAGTTTTGTAT-GGA
113445
Gallus_gallus 113401 TGTCAAGTTTGTGTTCTGTAACTTTTGACTCAATTCTGTACAGGT
113445

Anas_platyrhynchos_platyrhynchos 113446 GAAATGTGAAAAGATAAGAACAGCATCACAGTCGTATGATAGCAT
113490
Gallus_gallus 113446 GAGATGCAAATAGA-----GCCGTGCCATAGTACTATGATAACAC
113490

Anas_platyrhynchos_platyrhynchos 113491 GAGAACCCTGGTACATAGCTGGTACTTGAAGGAAGTGGG-GACAT
113535
Gallus_gallus 113491 ---------------CAGCTG----TCAAAGAAAGTAGGAGATGC
113535

Anas_platyrhynchos_platyrhynchos 113536 AGCTTTGGTCTCAAGTAAACTGTGTTAGTAGAACAGGCTCTGTGC
113580
Gallus_gallus 113536 AGCTCTGCTCTTAAGTAAAGCATGCTAGTAGAACAAACTCAGTTT
113580

Anas_platyrhynchos_platyrhynchos 113581 AAGCATGCACACTTGAGCAAATTTGAAGGGGCTAGTAATTTTGAT
113625
Gallus_gallus 113581 AAGTAAGCAACTTTGTGCAACTTTGAAGGGACTGGTTATTTTGAT
113625

Anas_platyrhynchos_platyrhynchos 113626 CTGGCTTTTTGTTGGCGTTTTATTTGCTTTGTATTTGTGTGTGTG
113670
Gallus_gallus 113626 CCGGCT--------GGGTTGGATTAGTTTGGTTTTGGTTGGTTTA
113670

Anas_platyrhynchos_platyrhynchos 113671 TGTTTTGTT--------TGTTTTTTTCTGGGAG--AGCAGGGCAG
113715
Gallus_gallus 113671 TATCAAGTTACTTGCACTGGTTGTTTTTGGTGGGCAGAAGGGGAG
113715

Anas_platyrhynchos_platyrhynchos 113716 AGGG-------------------ATGATGAATGAAAGGGTTGGAG
113760
Gallus_gallus 113716 AGGGTCAGGGTCATCAGCACAGTATCAGAAAGGAAAAGGTTGGAG
113760

Anas_platyrhynchos_platyrhynchos 113761 CTGATACTGTATCCAAGATGTTCAGAGTTTGTTCACAAGTTCATG
113805
Gallus_gallus 113761
CTGATACTGTATCTGTGATGTTCAGTGTTTGTGAACAG------- 113805

Anas_platyrhynchos_platyrhynchos 113806 TTCTTGTTGGTCCCTTCCAACCTGGGATATTCTATGAAAGTTGCT
113850
Gallus_gallus 113806 --------------------------------------AGTTGGT
113850

Anas_platyrhynchos_platyrhynchos 113851 AAATCTAATCCAGGTATTTTTTTTTAATTTTGCCTGTGCAGTGAG
113895
Gallus_gallus 113851 AGGTTTATTCCAGGTACTG------AACTTTG----TGTAGTAGG
113895

Anas_platyrhynchos_platyrhynchos 113896 AATAAGAGTTATTCTGGAGGGGG-----------TGGACATCTGG
113940
Gallus_gallus 113896 AATGAGATTTACTTTGGAGGGGGGGTGATGGGAGGTGGACAGCTGG
113940

Anas_platyrhynchos_platyrhynchos 113941 GAAATGTTAATATGTGTTAGAGTTCGTATAAGCCTTCTCCCATGG
113985
Gallus_gallus 113941 GAGACACTAATGTGTTTTGAAGATTATATAAACTTTCTCCTATAG
113985

Anas_platyrhynchos_platyrhynchos 113986 AGG-------AATTTGGATAAATGGTTACCT-CAGGAGTTAGTGT
114030
Gallus_gallus 113986
AGACAATGCAAACTTGGATAAGTGGTGACCCGCAGGAGTTAGT-- 114030

Anas_platyrhynchos_platyrhynchos 114031 GACCTCAGGAGTTAAGTGTGCAGTGCAACCTTGCTAAAATTATCC
114075
Gallus_gallus 114031 -------------AAGTGTGCATTGCAACCTTGCTGAATTCATTC
114075

Anas_platyrhynchos_platyrhynchos 114076 CTAGAGGCTACCTCCATAAAATAATGTCAAGTGGAATCTGCCACT
114120
Gallus_gallus 114076 TCAAAGTCTGTCTCCATAAAACAACTTAGAGGTGG---TGTCTCT
114120

Anas_platyrhynchos_platyrhynchos 114121 CAGAATCAAGGTTAAGGCAGAGCTTGTTCTGCAGATAATCTGTAG
114165
Gallus_gallus 114121 CAGAATCAAGGT----------CTTGTTCTGCAAATCATCTTTAG
114165

Anas_platyrhynchos_platyrhynchos 114166 AATTGCCAAAACTGATACATCTGCTTCCTTAGAGAGAAATACTGA
114210
Gallus_gallus 114166 AATTGCCCAGACTGATTTATTTGCTTCCTTAGACAGAAATGCTGA
114210

Anas_platyrhynchos_platyrhynchos 114211 GCAATTCAATGTCTCAAAAAAACATCCTGGTAGAATGCACATGTC
114255
Gallus_gallus 114211 GCAACTGACTCTC---AAAAACTGCCCTGGTAGAGTCCGCATTTC
114255

Anas_platyrhynchos_platyrhynchos 114256
AGGATAAAGACATCTAGT---------TTGTTTGTTCG------- 114300
Gallus_gallus 114256 AGGATAATGTCATCTAGTCAGTTTAGATTGTTTGCTTGGGTTTCT
114300

Anas_platyrhynchos_platyrhynchos 114301 -----------------------TTTTGCTTCCTGGACTGTTGAT
114345
Gallus_gallus 114301 TTGTTTTTGTTGTTTGTTTTTTTTTTTACCTCCTAGACTAGATAT
114345

Anas_platyrhynchos_platyrhynchos 114346 GGACTTGTCAGGTCAAAAGCACTATTTGGAAACAAATAGCGGAGG
114390

```
Gallus_gallus                    114346 TAACTCATCAGGTGAAAAAGACTACTTGAAATCAAACAGTGGAGG
114390

Anas_platyrhynchos_platyrhynchos 114391
CTTTG--------------GGAGTGTATCATATTTAG-------- 114435
Gallus_gallus                    114391 TTTTAATAGTATGTAGTAAGTAGTATGTAATATTCAGGTATTCAC
114435

Anas_platyrhynchos_platyrhynchos 114436 TCACTCAAAAAATGCATTGCATCTATATTTCTGCAAATACAAATT
114480
Gallus_gallus                    114436 TCATCCAAAAAATGGGTTAAGTCTACTTTTCTGAGAATGCAAGTT
114480

Anas_platyrhynchos_platyrhynchos 114481 TGGTCTCCTCAGAAAAATGAATCTCATTGGTGATAACATTTTCCT
114525
Gallus_gallus                    114481 TACTCTCCTGGGATGAAGAGATCTAACTAAT-ATTTCTCTTTTTT
114525

Anas_platyrhynchos_platyrhynchos 114526 TTTTTCCTTTAACATTCTCTTTTGCTTTGTTTCTTCCCTTTGCCT
114570
Gallus_gallus                    114526 TTTTTTTTGTAATGTTCTCTCTTGCTTT--CTCTTTTCTTTGCCT
114570

Anas_platyrhynchos_platyrhynchos 114571 GTTGATAGACCAGGACTCCATCTCATACGTTGCCTAGAAGCGGAA
114615
Gallus_gallus                    114571 CTTTGTAGACCAGGATGCTTCCTCATACACTGCCTAGAAGTGGAA
114615

Anas_platyrhynchos_platyrhynchos 114616 ACCCTGGCTTTATCATTGGAGCACCACTGCTGATTGCAAACACTG
114660
Gallus_gallus                    114616 ATCCTGGCTATATCACTGGAGCACCACTGCTGATTGCAAATGATA
114660

Anas_platyrhynchos_platyrhynchos 114661 GTGCCACGCAGCATGTATCCTTTCTGCTGTCAAGTTTTATTTATC
114705
Gallus_gallus                    114661 GTGCAGTGCAACACGTATCCTTTCTGCTGTCATGTTTTATCTGTC
114705

Anas_platyrhynchos_platyrhynchos 114706 TCTGGGTTTCTTGACGCCTGGTACTATTCTGTTCGGACTCTTGAG
114750
Gallus_gallus                    114706 ACTGGGTTTCTTAACTTCTGTTACTATGTTGTGTGAATTTCTGAG
114750

Anas_platyrhynchos_platyrhynchos 114751 ATTCTACCTTTACCTGTGAGGGCAGACTTCAATATTACTTTGCAG
114795
Gallus_gallus                    114751 ATTGTACCTTTGCCTGTGAGGGCATACTTCAGTATAACTTTGCGG
114795

Anas_platyrhynchos_platyrhynchos 114796 CTATCAGTAGGCCTTTTGGTCAGG-----------AAACCCTG-T
114840
Gallus_gallus                    114796 CCAGTAGTATGCTGTTTGGTCAGGGCTGAATTGTAAAGCTGTGAT
114840

Anas_platyrhynchos_platyrhynchos 114841 AGGGTTAAGGAGGAATGCGGTTTACGG--AGAATGTTAAATAAAA
114885
Gallus_gallus                    114841 AGTGCAAAAGAGGAATG-----TACAGTAAGGTTGTTAAATACAA
114885

Anas_platyrhynchos_platyrhynchos 114886 GTTGTGTGTCATTTATTATCTGGTAGTTTTTTGTTTGGCTTCTTG
114930
Gallus_gallus                    114886 GTTGAGTGTCCTTTGTGATTTG----------------CTTCTTG
114930
```

```
Anas_platyrhynchos_platyrhynchos 114931 ACATGCAGTCTATTAGATGAGCATTTTAAGGAGTGAGGGTGATGG
114975
Gallus_gallus                    114931 ACATGCAGATTGTTAGATGAGCATTCTGCAGAGCGAGGGTGATGG
114975

Anas_platyrhynchos_platyrhynchos 114976 AAGTTGTTCACAGTTCCTCAGACACACGGTACAATTTGGAAAAAA
115020
Gallus_gallus                    114976 AAGTTGCTCACAGTTTCTGAGACATACAGTACGATTTGGAAGCAA
115020

Anas_platyrhynchos_platyrhynchos 115021 TATGAGGACTGGCTGCAAGCTCAGGTAAGAACACGCT--GCAGTC
115065
Gallus_gallus                    115021 TACGAGGACTGGCTGCAAACTCAGGTAGGAACATGCTCTGTGATC
115065

Anas_platyrhynchos_platyrhynchos 115066 ACCCTTACTGACATACTTGGAAGACTC----TCACTCAGCACCTG
115110
Gallus_gallus                    115066 ACCCTTCCTGAAACATCTGGAAGGCATGGAGTCATTTAGCACCTT
115110

Anas_platyrhynchos_platyrhynchos 115111 TTTCCATAGCTTCTCCCTCACTTCTGTGGAGCCAAGCGCTTTCTC
115155
Gallus_gallus                    115111 TTTTTGTAGCTTTTCACTCATTTCTGTGGAGCCAAGGACCCTCTT
115155

Anas_platyrhynchos_platyrhynchos 115156 TAAGGTCACCAATGGTTCTC-TACAGATTTTTCAT-CTACTCTAA
115200
Gallus_gallus                    115156 TAAGGTCACCCATGGTTTTAACACAGGTCTTTCACGTTAGCCCTA
115200

Anas_platyrhynchos_platyrhynchos 115201 AGAGGACTAGACTGTCAGAGGGATGGTGGACATCTAGAGCAGTTT
115245
Gallus_gallus                    115201 AGAAAACTAGACCATCAGAGGGATGGTAAGCCTACAAAACAGTTT
115245

Anas_platyrhynchos_platyrhynchos 115246 GCACAT-AGTGATGGCCCTCTGGGAAGGTCTGAGTCTTTGGTCTG
115290
Gallus_gallus                    115246 GAATGTGAGAGATAATCCTCTAGGAAGTCTTCAGGTTTTGGTCTG
115290

Anas_platyrhynchos_platyrhynchos 115291 GCAACATGTCTTGGCCAAGAGTAGCGTTTGTATTTGAGAAGTGAT
115335
Gallus_gallus                    115291 GTGACATGTCCCAGACAAGAGTAAAGTTCATGTTTGAGAGGTGAA
115335

Anas_platyrhynchos_platyrhynchos 115336 TTATAATGGAACCTTAAAAACTGATT--GATTTGA----TGATGT
115380
Gallus_gallus                    115336 TTATAATAAAACTTTAAATGATTATTCTAATTTGAATTATTATTT
115380

Anas_platyrhynchos_platyrhynchos 115381 GGGTGA-----------------CACTGAGTTGATATGATAGCTA
115425
Gallus_gallus                    115381 TGTTGAAAGGTCAGGTATACCATCACTGAGGTGGTAAGATGGGTG
115425

Anas_platyrhynchos_platyrhynchos 115426 AAAAGCTGTCAGTTTTCTGCATCATGTTTTATCCTGCCATGTTCT
115470
Gallus_gallus                    115426 AAAAGCTGTCGGTATTC---ATCATGTT--ATCTTGCCATGTTCT
115470

Anas_platyrhynchos_platyrhynchos 115471 TCTCGCTTCTCATTGCAGAAAGGATTACAACTTTGCCTGCTTAAT
115515
Gallus_gallus                    115471 TCACCCTTGTCCTTGCAGAAAGGATTGGAAGTTGACTC.......
```

115515

Anas_platyrhynchos_platyrhynchos 115516 ACTGGTACTTCTGTAATGATTCTATTGACTTCATATTCTATTACC
115560
Gallus_gallus 115516 .....................................TGTTGCT
115560

Anas_platyrhynchos_platyrhynchos 115561 AGTGAAGGGTTTCGATATCAGCTGCTTCTGCCCAGGAAAGCGCTT
115605
Gallus_gallus 115561 GAGGAAAGTTGTTGATGCCATCTGCTTCTGTCTTGGAAATCCCTT
115605

Anas_platyrhynchos_platyrhynchos 115606 TCTG-AAAGCTGTAATTCTATTTCATTTCCAGCCTACCCCAAACA
115650
Gallus_gallus 115606 TAGATAAAGCTGGTACTTTTTTTC----CTAGGCTAACTCAAATG
115650

Anas_platyrhynchos_platyrhynchos 115651 GAAGAAAGTAACTGCAGTTACATCCAGATGAAGTTATATGAGGCT
115695
Gallus_gallus 115651 GAAGAAAGTAATTGCAGTTATATCCAGCTGAAGTTATACAAGGCT
115695

Anas_platyrhynchos_platyrhynchos 115696 CTTCAGGGGATGAACACAGCAGAGGACCTTGCTATAACTAGCAGT
115740
Gallus_gallus 115696 TTTCAGGGGATGAACCAAGCAGAGGACCTTGCTATAATTGGTAGT
115740

Anas_platyrhynchos_platyrhynchos 115741 GCTCATTCAACCCAGGTAGAAGAGCGGACAACCATTCTGATTCAG
115785
Gallus_gallus 115741 GCTCGTTCAACCCAAGAAGAAGAGTGGGTGACCATTCTGATCCAG
115785

Anas_platyrhynchos_platyrhynchos 115786 AACTGCAGTGTACAGGTAATGTAAATGTAAAAAGACCATCTGAAT
115830
Gallus_gallus 115786 AACTGCAGTGTACAGGTAATACAAATGCAAAAAGACAATCTGAAT
115830

Anas_platyrhynchos_platyrhynchos 115831 ATTTATTTTTTATATCTTTGTGTTAGTTTGAGGAGC-----ATGA
115875
Gallus_gallus 115831 AC---CTTTTCATATCCTTGTTTTAGTTTGAGAAGTGAGTAATAT
115875

Anas_platyrhynchos_platyrhynchos 115876 GTGAGGGGGGAGGAAAAAAAAAAAGAACCGTCTTGTTCATCAGGG
115920
Gallus_gallus 115876 TTAAAAAAAAAAAAAAAAAAAAAAGAACCAGTTTGTTCATCAGGG
115920

Anas_platyrhynchos_platyrhynchos 115921
CTTGGTTT-CTTTGTGTGATGATCCTCTGAATATCAGGTGCCTT- 115965
Gallus_gallus 115921 CTAGTTTTAATTTGTGTGATAATCCACTGGCTTTCAGGAGTCTTA
115965

Anas_platyrhynchos_platyrhynchos 115966 ------GTAAACAGCCATCCTTATTAAGGTTCTTACTATTAGAAT
116010
Gallus_gallus 115966 ATATTCATAAACATTCATGTTTATTAGTGTTCATATTGTTAGGAT
116010

Anas_platyrhynchos_platyrhynchos 116011 ATTTATTAAACA--AAAGAAGAACTGATAGATTTAATCTCTCTTG
116055
Gallus_gallus 116011 ATTTAGTAAGCAACAACAAAAAACTGATAGGTTTA--TTCTCTTA
116055

Anas_platyrhynchos_platyrhynchos 116056

```
AATCCCGTTCCTGATATTCAAAGAGTCAAAAAAAAGTTTAA---- 116100
Gallus_gallus                   116056 ATTCCTTGTTGTTGTATTTAAATGGCC-AAGAAGAGTTTAATGCT
116100

Anas_platyrhynchos_platyrhynchos 116101 TAATCTAGCGTTGCACTACCTAGTTTTCTGACAGCTTTCTAGAAC
116145
Gallus_gallus                   116101 TAGTCTAGTGCTGTACTACCTAGTTTTCTGACTGCATTCTAAGAC
116145

Anas_platyrhynchos_platyrhynchos 116146 T------ACCTGGTTACTTCCATTAATTTGATGCTTATATTATTA
116190
Gallus_gallus                   116146 TAATACAACCTGTTTACTCCCATTAATTTGATACT-------TTA
116190

Anas_platyrhynchos_platyrhynchos 116191 TTGTCCTTTCTGA-TTTTCTAGTTTTTAG-GATATTTATGCATTA
116235
Gallus_gallus                   116191 TCATCTTTTCTGATTTTTTTGGATTCTAGTGATATTTGTATATTG
116235

Anas_platyrhynchos_platyrhynchos 116236 CCTAACTAAAGTTAAGAGCTTGCCATCCAAGTATTTATAGAAGAG
116280
Gallus_gallus                   116236 CTTAACTAAAGCTAAGAGCTTGCAATCCAAGTACTTGCTGGTGAG
116280

Anas_platyrhynchos_platyrhynchos 116281 GGAAGTTCTGTTTTGATCATT-TTAATCTATCTGGAGATGGCCTA
116325
Gallus_gallus                   116281 GTAGATTCTGTTTTAATAATTACTCATCTATTAGAAGGTGTCCTC
116325

Anas_platyrhynchos_platyrhynchos 116326 ATGGAATGTAAAATCCAAGTGATAGAACTGAGACAAATTCTATTC
116370
Gallus_gallus                   116326 ATAGAAGGCAAAATCCAGGTGATAGAGGTGAGATAAAT--TGTCT
116370

Anas_platyrhynchos_platyrhynchos 116371 AGAAATGAGATGACAGCGAGATGAGTATAGCTCTAAAACGTGCAG
116415
Gallus_gallus                   116371 GGAAATGAGACGACAGTGAGATAGATACAGCTCTAAAATGGGTAG
116415

Anas_platyrhynchos_platyrhynchos 116416 CTGACCTGACCAGAAGTTACGGGATTAGTTCTGAAGTAACTCATT
116460
Gallus_gallus                   116416 CTGACCTGACAAGAAGTTATGGGGTTAGTCCTCAAGTAAGTGATT
116460

Anas_platyrhynchos_platyrhynchos 116461 GAAGT--TGTGACCTCTGCTACACAAAGTGTTAGACTTTGACTAT
116505
Gallus_gallus                   116461 GAGGTTCTGAGACCTTTGCTACTCGAAGTATTGAGCTTTGACCGT
116505

Anas_platyrhynchos_platyrhynchos 116506 GATGGTCCTTTTCAACTTTTCAATCTTTAAAATACTATCTTGAGA
116550
Gallus_gallus                   116506 GATGGTCCTTTTTAAC--------ATGTAAATTAATGCATTGAGA
116550

Anas_platyrhynchos_platyrhynchos 116551 TGCTTGGGTTGCCTTTTATTTTCAGGCTATGAACTGTACTTCCTG
116595
Gallus_gallus                   116551 TGTTTGAGTTGCATTTTA...........................
116595

Anas_platyrhynchos_platyrhynchos 116596 TTGCATGGTNNNNNNNNNNNNNNNNNNNNNNNNNNNNNNNNNNNN
116640
Gallus_gallus                   116596 .............................................
116640
```

```
Anas_platyrhynchos_platyrhynchos 116641 NNNNNNNNNNNNNNNNNNNNNNNNNNNNNNNNNNNNNNNNNNNNN
116685
Gallus_gallus                    116641 .............................................
116685

Anas_platyrhynchos_platyrhynchos 116686 NNNNNNNNNNNNNNNNNNNNNNNNNNNNNNNNNNNNNNNNNNNNN
116730
Gallus_gallus                    116686 .............................................
116730

Anas_platyrhynchos_platyrhynchos 116731 NNNNNNNNNNNNNNNNNNNNNNNNNNNNNNNNNNNNNNNNNNNNN
116775
Gallus_gallus                    116731 .............................................
116775

Anas_platyrhynchos_platyrhynchos 116776 NNNNNNNNNNNNNNNNNNNNNNNNNNNNNNNNNNNNNNNNNNNNN
116820
Gallus_gallus                    116776 .............................................
116820

Anas_platyrhynchos_platyrhynchos 116821 NNNNNNNNNNNNNNNNNNNNNNNNNNNNNNNNNNNNNNNNNNNNN
116865
Gallus_gallus                    116821 .............................................
116865

Anas_platyrhynchos_platyrhynchos 116866 NNNNNNNNNNNNNNNNNNNNNNNNNNNNNNNNNNNNNNNNNNNNN
116910
Gallus_gallus                    116866 .............................................
116910

Anas_platyrhynchos_platyrhynchos 116911 NNNNNNNNNNNNNNNNNNNNNNNNNNNNNNNNNNNNNNNNNNNNN
116955
Gallus_gallus                    116911 .............................................
116955

Anas_platyrhynchos_platyrhynchos 116956 NNNNNNNNNNNNNNNNNNNNNNNNNNNNNNNNNNNNNNNNNNNNN
117000
Gallus_gallus                    116956 .............................................
117000

Anas_platyrhynchos_platyrhynchos 117001 NNNNNNNNNNNNNNNNNNNNNNNNNNNNNNNNNNNNNNNNNNNNN
117045
Gallus_gallus                    117001 .............................................
117045

Anas_platyrhynchos_platyrhynchos 117046 NNNNNNNNNNNNNNNNNNNNNNNNNNNNNNNNNNNNNNNNNNNNN
117090
Gallus_gallus                    117046 .............................................
117090

Anas_platyrhynchos_platyrhynchos 117091 NNNNNNNNNNNNNNNNNNNNNNNNNNNNNNNNNNNNNNNNNNNNN
117135
Gallus_gallus                    117091 .............................................
117135

Anas_platyrhynchos_platyrhynchos 117136 NNNNNNNNNNNNNNNNNNNNNNNNNNNNNNNNNNNNNNNNNNNNN
117180
Gallus_gallus                    117136 .............................................
117180

Anas_platyrhynchos_platyrhynchos 117181 NNNNNNNNNNNNNNNNNNNNNNNNNNNNNNNNNNNNNNNNNNNNN
117225
```

```
Gallus_gallus                   117181 .............................................
117225

Anas_platyrhynchos_platyrhynchos 117226 NNNNNNNNNNNNNNNNNNNNNNNNNNNNNNNNNNNNNNNNNNNNN
117270
Gallus_gallus                   117226 .............................................
117270

Anas_platyrhynchos_platyrhynchos 117271 NNNNNNNNNNNNNNNNNNNNNNNNNNNNNNNNNNNNNNNNNNNNN
117315
Gallus_gallus                   117271 .............................................
117315

Anas_platyrhynchos_platyrhynchos 117316 NNNNNNNNNNNNNNNNNNNNNNNNNNNNNNNNNNNNNNNNNNNNN
117360
Gallus_gallus                   117316 .............................................
117360

Anas_platyrhynchos_platyrhynchos 117361 NNNNNNNNNNNNNNNNNNNNNNNNNNNNNNNNNNNNNNNNNNNNN
117405
Gallus_gallus                   117361 .............................................
117405

Anas_platyrhynchos_platyrhynchos 117406 NNNNNNNNNNNNNNNNNNNNNNNNNNNNNNNNNNNNNNNNNNNNN
117450
Gallus_gallus                   117406 .............................................
117450

Anas_platyrhynchos_platyrhynchos 117451 NNNNNNNNNNNNNNNNNNNNNNNNNNNNNNNNNNNNNNNNNNNNN
117495
Gallus_gallus                   117451 .............................................
117495

Anas_platyrhynchos_platyrhynchos 117496 NNNNNNNNNNNNNNNNNNNNNNNNNNNNNNNNNNNNNNNNNNNNN
117540
Gallus_gallus                   117496 .............................................
117540

Anas_platyrhynchos_platyrhynchos 117541 NNNNNNNNNNNNNNNNNNNNNNNNNNNNNNNNNNNNNNNNNNNNN
117585
Gallus_gallus                   117541 .............................................
117585

Anas_platyrhynchos_platyrhynchos 117586 NNNNNNNNNNNNNNNNNNNNNNNNNNNNNNNNNNNNNNNNNNNNN
117630
Gallus_gallus                   117586 .............................................
117630

Anas_platyrhynchos_platyrhynchos 117631 NNNNNNNNNNNNNNNNNNNNNNNNNNNNNNNNNNNNNNNNNNNNN
117675
Gallus_gallus                   117631 .............................................
117675

Anas_platyrhynchos_platyrhynchos 117676 NNNNNNNNNNNNNNNNNNNNNNNNNNNNNNNNNNNNNNNNNNNNN
117720
Gallus_gallus                   117676 .............................................
117720

Anas_platyrhynchos_platyrhynchos 117721 NNNNNNNNNNNNNNNNNNNNNNNNNNNNNNNNNNNNNNNNNNNNN
117765
Gallus_gallus                   117721 .............................................
117765
```

```
Anas_platyrhynchos_platyrhynchos 117766 NNNNNNNNNNNNNNNNNNNNNNNNNNNNNNNNNNNNNNNNNNNNN
117810
Gallus_gallus                    117766 .............................................
117810

Anas_platyrhynchos_platyrhynchos 117811 NNNNNNNNNNNNNNNNNNNNNNNNNNNNNNNNNNNNNNNNNNNNN
117855
Gallus_gallus                    117811 .............................................
117855

Anas_platyrhynchos_platyrhynchos 117856 NNNNNNNNNNNNNNNNNNNNNNNNNNNNNNNNNNNNNNNNNNNNN
117900
Gallus_gallus                    117856 .............................................
117900

Anas_platyrhynchos_platyrhynchos 117901 NNNNNNNNNNNNNNNNNNNNNNNNNNNNNNNNNNNNNNNNNNNNN
117945
Gallus_gallus                    117901 .............................................
117945

Anas_platyrhynchos_platyrhynchos 117946 NNNNNNNNNNNNNNNNNNNNNNNNNNNNNNNNNNNNNNNNNNNNN
117990
Gallus_gallus                    117946 .............................................
117990

Anas_platyrhynchos_platyrhynchos 117991 NNNNNNNNNNNNNNNNNNNNNNNNNNNNNNNNNNNNNNNNNNNNN
118035
Gallus_gallus                    117991 .............................................
118035

Anas_platyrhynchos_platyrhynchos 118036 NNNNNNNNNNNNNNNNNNNNNNNNNNNNNNNNNNNNNNNNNNNNN
118080
Gallus_gallus                    118036 .............................................
118080

Anas_platyrhynchos_platyrhynchos 118081 NNNNNNNNNNNNNNNNNNNNNNNNNNNNNNNNNNNNNNNNNNNNN
118125
Gallus_gallus                    118081 .............................................
118125

Anas_platyrhynchos_platyrhynchos 118126 NNNNNNNNNNNNNNNNNNNNNNNNNNNNNNNNNNNNNNNNNNNNN
118170
Gallus_gallus                    118126 .............................................
118170

Anas_platyrhynchos_platyrhynchos 118171 NNNNNNNNNNNNNNNNNNNNNNNNNNNNNNNNNNNNNNNNNNNNN
118215
Gallus_gallus                    118171 .............................................
118215

Anas_platyrhynchos_platyrhynchos 118216 NNNNNNNNNNNNNNNNNNNNNNNNNNNNNNNNNNNNNNNNNNNNN
118260
Gallus_gallus                    118216 .............................................
118260

Anas_platyrhynchos_platyrhynchos 118261 NNNNNNNNNNNNNNNNNNNNNNNNNNNNNNNNNNNNNNNNNNNNN
118305
Gallus_gallus                    118261 .............................................
118305

Anas_platyrhynchos_platyrhynchos 118306 NNNNNNNNNNNNNNNNNNNNNNNNNNNNNNNNNNNNNNNNNNNNN
118350
Gallus_gallus                    118306 .............................................
```

118350

Anas_platyrhynchos_platyrhynchos 118351 NNNNNNNNNNNNNNNNNNNNNNNNNNNNNNNNNNNNNNNNNNNNN
118395
Gallus_gallus 118351 .............................................
118395

Anas_platyrhynchos_platyrhynchos 118396 NNNNNNNNNNNNNNNNNNNNNNNNNNNNNNNNNNNNNNNNNNNNN
118440
Gallus_gallus 118396 .............................................
118440

Anas_platyrhynchos_platyrhynchos 118441 NNNNNNNNNNNNNNNNNNNNNNNNNNNNNNNNNNNNNNNNNNNNN
118485
Gallus_gallus 118441 .............................................
118485

Anas_platyrhynchos_platyrhynchos 118486 NNNNNNNNNNNNNNNNNNNNNNNNNNNNNNNNNNNNNNNNNNNNN
118530
Gallus_gallus 118486 .............................................
118530

Anas_platyrhynchos_platyrhynchos 118531 NNNNNNNNNNNNNNNNNNNNNNNNNNNNNNNNNNNNNNNNNNNNN
118575
Gallus_gallus 118531 .............................................
118575

Anas_platyrhynchos_platyrhynchos 118576 NNNNNNNNNNNNNNNNNNNNNNNNNNNNNNNNNNNNNNNNNNNNN
118620
Gallus_gallus 118576 .............................................
118620

Anas_platyrhynchos_platyrhynchos 118621 NNNNNNNNNNNNNNNNNNNNNNNNNNNNNNNNNNNNNNNNNNNNN
118665
Gallus_gallus 118621 .............................................
118665

Anas_platyrhynchos_platyrhynchos 118666 NNNNNNNNNNNNNNNNNNNNNNNNNNNNNNNNNNNNNNNNNNNNN
118710
Gallus_gallus 118666 .............................................
118710

Anas_platyrhynchos_platyrhynchos 118711 NNNNNNNNNNNNNNNNNNNNNNNNNNNNNNNNNNNNNNNNNNNNN
118755
Gallus_gallus 118711 .............................................
118755

Anas_platyrhynchos_platyrhynchos 118756 NNNNNNNNNNNNNNNNNNNNNNNNNNNNNNNNNNNNNNNNNNNNN
118800
Gallus_gallus 118756 .............................................
118800

Anas_platyrhynchos_platyrhynchos 118801 NNNNNNNNNNNNNNNNNNNNNNNNNNNNNNNNNNNNNNNNNNNNN
118845
Gallus_gallus 118801 .............................................
118845

Anas_platyrhynchos_platyrhynchos 118846 NNNNNNNNNNNNNNNNNNNNNNNNNNNNNNNNNNNNNNNNNNNNN
118890
Gallus_gallus 118846 .............................................
118890

Anas_platyrhynchos_platyrhynchos 118891 NNNNNNNNNNNNNNNNNNNNNNNNNNNNNNNNNNNNNNNNNNNNN

```
118935
Gallus_gallus                  118891 .............................................
118935

Anas_platyrhynchos_platyrhynchos 118936 NNNNNNNNNNNNNNNNNNNNNNNNNNNNNNNNNNNNNNNNNNNNN
118980
Gallus_gallus                  118936 .............................................
118980

Anas_platyrhynchos_platyrhynchos 118981 NNNNNNNNNNNNNNNNNNNNNNNNNNNNNNNNNNNNNNNNNNNNN
119025
Gallus_gallus                  118981 .............................................
119025

Anas_platyrhynchos_platyrhynchos 119026 NNNNNNNNNNNNNNNNNNNNNNNNNNNNNNNNNNNNNNNNNNNNN
119070
Gallus_gallus                  119026 .............................................
119070

Anas_platyrhynchos_platyrhynchos 119071 NNNNNNNNNNNNNNNNNNNNNNNNNNNNNNNNNNNNNNNNNNNNN
119115
Gallus_gallus                  119071 .............................................
119115

Anas_platyrhynchos_platyrhynchos 119116 NNNNNNNNNNNNNNNNNNNNNNNNNNNNNNNNNNNNNNNNNNNNN
119160
Gallus_gallus                  119116 .............................................
119160

Anas_platyrhynchos_platyrhynchos 119161 NNNNNNNNNNNNNNNNNNNNNNNNNNNNNNNNNNNNNNNNNNNNN
119205
Gallus_gallus                  119161 .............................................
119205

Anas_platyrhynchos_platyrhynchos 119206 NNNNNNNNNNNNNNNNNNNNNNNNNNNNNNNNNNNNNNNNNNNNN
119250
Gallus_gallus                  119206 .............................................
119250

Anas_platyrhynchos_platyrhynchos 119251 NNNNNNNNNNNNNNNNNNNNNNNNNNNNNNNNNNNNNNNNNNNNN
119295
Gallus_gallus                  119251 .............................................
119295

Anas_platyrhynchos_platyrhynchos 119296 NNNNNNNNNNNNNNNNNNNNNNNNNNNNNNNNNNNNNNNNNNNNN
119340
Gallus_gallus                  119296 .............................................
119340

Anas_platyrhynchos_platyrhynchos 119341 NNNNNNNNNNNNNNNNNNNNNNNNNNNNNNNNNNNNNNNNNNNNN
119385
Gallus_gallus                  119341 .............................................
119385

Anas_platyrhynchos_platyrhynchos 119386 NNNNNNNNNNNNNNNNNNNNNNNNNNNNNNNNNNNNNNNNNNNNN
119430
Gallus_gallus                  119386 .............................................
119430

Anas_platyrhynchos_platyrhynchos 119431 NNNNNNNNNNNNNNNNNNNNNNNNNNNNNNNNNNNNNNNNNNNNN
119475
Gallus_gallus                  119431 .............................................
119475
```

```
Anas_platyrhynchos_platyrhynchos 119476 NNNNNNNNNNNNNNNNNNNNNNNNNNNNNNNNNNNNNNNNNNNNN
119520
Gallus_gallus                    119476 .............................................
119520

Anas_platyrhynchos_platyrhynchos 119521 NNNNNNNNNNNNNNNNNNNNNNNNNNNNNNNNNNNNNNNNNNNNN
119565
Gallus_gallus                    119521 .............................................
119565

Anas_platyrhynchos_platyrhynchos 119566 NNNNNNNNNNNNNNNNNNNNNNNNNNNNNNNNNNNNNNNAGCAGG
119610
Gallus_gallus                    119566 .............................................
119610

Anas_platyrhynchos_platyrhynchos 119611 CAGATGAGGAAGAGTCTAATTCTGATGGTGGTAGGTTTCAAGAAT
119655
Gallus_gallus                    119611 .............................................
119655

Anas_platyrhynchos_platyrhynchos 119656 TAGGGGGAAAGATCATGAGCCAGAAAGCAAGTTTTTAAGAGAAGA
119700
Gallus_gallus                    119656 .............................................
119700

Anas_platyrhynchos_platyrhynchos 119701 TTACTTAATGTCAGCAGGAAGGTAGAGCTAACAGGACTGTGAAGG
119745
Gallus_gallus                    119701 .............................................
119745

Anas_platyrhynchos_platyrhynchos 119746 AAAGTGAATAGTTTTTTTTTGAGTATGTGTGGATCCCATTTTGGT
119790
Gallus_gallus                    119746 .............................................
119790

Anas_platyrhynchos_platyrhynchos 119791 GCAAGGCTGCATTCCTGCGATGACAAAATCTTTTGGGCAGCATGT
119835
Gallus_gallus                    119791 .............................................
119835

Anas_platyrhynchos_platyrhynchos 119836 GTATTTCAAGGCTTATACTTCTCTGCAAGAGCTTTAATGGGGAGC
119880
Gallus_gallus                    119836 .............................................
119880

Anas_platyrhynchos_platyrhynchos 119881 AGCTGACACCTGTCCATCATGCCTTCATGGCAATGAGGTATGACA
119925
Gallus_gallus                    119881 .............................................
119925

Anas_platyrhynchos_platyrhynchos 119926 TTGCAAATGTCTGATATACTACTATTCTTTGAACTTCGGATTCAG
119970
Gallus_gallus                    119926 .............................................
119970

Anas_platyrhynchos_platyrhynchos 119971 CCCAAATGGGACTTCACATCACTTGTTTGGCCGCGTCTCATGGTT
120015
Gallus_gallus                    119971 .............................................
120015

Anas_platyrhynchos_platyrhynchos 120016 CCTCTTTGTCTCCCTTCCCTGCAACGTAATTTAAAAGAACATGAC
120060
```

```
Gallus_gallus                    120016 .............................................
120060

Anas_platyrhynchos_platyrhynchos 120061 AGCAAAATGTTTACCCTGTAAGACATACTTTACCTTGAGGCAAGC
120105
Gallus_gallus                    120061 .............................................
120105

Anas_platyrhynchos_platyrhynchos 120106 TGACACCACTCCCCATGTGCTTGCCTGCTATTGGGCGGATAATTT
120150
Gallus_gallus                    120106 .............................................
120150

Anas_platyrhynchos_platyrhynchos 120151 CTAAATCTGAACTATTATATGGCTCTATTGATATAGCTTCTGTCT
120195
Gallus_gallus                    120151 .............................................
120195

Anas_platyrhynchos_platyrhynchos 120196 TCATTCCTAGTAAAAGTGCAGGTTGGACATCTCTCAATATGGATT
120240
Gallus_gallus                    120196 .............................................
120240

Anas_platyrhynchos_platyrhynchos 120241 AGCGAGTCAGGAATAATGAGATTCAGATTTCATCTGCTTGAGCAA
120285
Gallus_gallus                    120241 .............................................
120285

Anas_platyrhynchos_platyrhynchos 120286 CTGATGCTAATTACTTTAGAATTATAGAATATCCTGAGTTGAAAG
120330
Gallus_gallus                    120286 .............................................
120330

Anas_platyrhynchos_platyrhynchos 120331 GGTCCCACAAGGATCACCGAGTCCAACTCCTGAGTCCCCAAAGGA
120375
Gallus_gallus                    120331 .............................................
120375

Anas_platyrhynchos_platyrhynchos 120376 CTACCCAAAAAATCAGACCATGTGTCTGAGAGCGTTGTCCAAATG
120420
Gallus_gallus                    120376 .............................................
120420

Anas_platyrhynchos_platyrhynchos 120421 CTTCCTGAACTCTGGCAGGCTCAGTGCTGTGACTACTGCCCTAGG
120465
Gallus_gallus                    120421 .............................................
120465

Anas_platyrhynchos_platyrhynchos 120466 GAGCCTGTTTCAGTGCCTGACCACTCTCTCGGTAAAGAACCTGTT
120510
Gallus_gallus                    120466 .............................................
120510

Anas_platyrhynchos_platyrhynchos 120511 CCTGATATTCAGCCTGAACTGCCCCTTTAAGCTGTTTCCTTGGGT
120555
Gallus_gallus                    120511 .............................................
120555

Anas_platyrhynchos_platyrhynchos 120556 TCTATAGCTGGTCATCGGAGAGAGAAGCTCAGCAACTTTAGGGCT
120600
Gallus_gallus                    120556 .............................................
120600
```

```
Anas_platyrhynchos_platyrhynchos 120601 AGAAGATGTGTTGCAAAGCAAATGATAAGGAAGTTTCTGGCAGCA
120645
Gallus_gallus                    120601 .............................................
120645

Anas_platyrhynchos_platyrhynchos 120646 GAAAACACCATTAAAATCAGTAATGACTTCATGAGTGGCAATCCA
120690
Gallus_gallus                    120646 .............................................
120690

Anas_platyrhynchos_platyrhynchos 120691 CAAATCAGGGTTGTTAGTAAAGACATTTTAGCAATGACACGAAGA
120735
Gallus_gallus                    120691 .............................................
120735

Anas_platyrhynchos_platyrhynchos 120736 GCCTTAGAAATCCTTTTAGGATGTACCTGTCTTAGGAACAGTTTA
120780
Gallus_gallus                    120736 .............................................
120780

Anas_platyrhynchos_platyrhynchos 120781 CATTGGCACCGTAAGCCAGTAGTAGTGCCAGTGTCAAACCCTGAT
120825
Gallus_gallus                    120781 .............................................
120825

Anas_platyrhynchos_platyrhynchos 120826 GTGTGAAATTGATAAAAAGCTGAATCATTCAGCCAGATCTAGGAC
120870
Gallus_gallus                    120826 .............................................
120870

Anas_platyrhynchos_platyrhynchos 120871 AAAACTGGAACAAGGCAGGAGAAGAAAATTAATTCCCCTAAGTGC
120915
Gallus_gallus                    120871 .............................................
120915

Anas_platyrhynchos_platyrhynchos 120916 CATCTGAACTGCATTATACCAGTTAGACAGCTGGATCATAAATCT
120960
Gallus_gallus                    120916 .............................................
120960

Anas_platyrhynchos_platyrhynchos 120961 GCTGGAAGTATTTGTATCAATGGATGAGAATGTATGTTTCTAGTT
121005
Gallus_gallus                    120961 .............................................
121005

Anas_platyrhynchos_platyrhynchos 121006 GTGACTAATTTCAGGGCCATCTAGGATATTCTGATATACATCACA
121050
Gallus_gallus                    121006 .............................................
121050

Anas_platyrhynchos_platyrhynchos 121051 GAGACAGCGCCTGCTTACTCGTTGATACAAGAACTTACATCCATC
121095
Gallus_gallus                    121051 .............................................
121095

Anas_platyrhynchos_platyrhynchos 121096 AAGAGTCTTGGTATTTTTCTCTTGAGACTCGTGAATTCCATCCTG
121140
Gallus_gallus                    121096 .............................................
121140

Anas_platyrhynchos_platyrhynchos 121141 GAGCTTCTTTGAATGCATTCTGCACCACTTTGCTCTAAAGGCAAT
121185
Gallus_gallus                    121141 .............................................
```

121185

Anas_platyrhynchos_platyrhynchos 121186 CTTGTTGAGGTTTTCAGAAAAGTGAAATGAGATAGTGATTTCACA 121230
Gallus_gallus 121186 ............................................. 121230

Anas_platyrhynchos_platyrhynchos 121231 GATTGGCTTAAATCAGTACTAGCACAAAATGCATACTGCCAGTAG 121275
Gallus_gallus 121231 ............................................. 121275

Anas_platyrhynchos_platyrhynchos 121276 GTTCCATCTCAAGGCTAGCAATGCGGAGCTGCGAAATGGCTGGGA 121320
Gallus_gallus 121276 ............................................. 121320

Anas_platyrhynchos_platyrhynchos 121321 GGTGGTGACAGGTTGTGAGACTGCTCCTGCCAAAAGAACGTGGTC 121365
Gallus_gallus 121321 ............................................. 121365

Anas_platyrhynchos_platyrhynchos 121366 TCATCTCAAACCAAGCTAGCCATGCAACTATCCTCAGCTCGGGGT 121410
Gallus_gallus 121366 ............................................. 121410

Anas_platyrhynchos_platyrhynchos 121411 GCGTGTGTGTTCTCTTACAGCATTCTGTTGAAACAGAAAGATGCT 121455
Gallus_gallus 121411 ............................................. 121455

Anas_platyrhynchos_platyrhynchos 121456 GAGGGCTTAGGATAATTTCAGTCCCCTGTTTCGTCTACCAACTGA 121500
Gallus_gallus 121456 ............................................. 121500

Anas_platyrhynchos_platyrhynchos 121501 CCTACCAATCTATCTTCTTTCTGAAACAATATCATAAAATAATGA 121545
Gallus_gallus 121501 ............................................. 121545

Anas_platyrhynchos_platyrhynchos 121546 GAAAGAAACTCATATTGGGTCCCTGGCTTATTTTATGGAAAGGAT 121590
Gallus_gallus 121546 ............................................. 121590

Anas_platyrhynchos_platyrhynchos 121591 CAAACTCTGTGGCAATCCTGTCCTTTTCTGGCAGCAATGAGCTGC 121635
Gallus_gallus 121591 ............................................. 121635

Anas_platyrhynchos_platyrhynchos 121636 TCAAAGGTGATGTTATCTCAAATGTTATTTAGAAAAATAAAGCTT 121680
Gallus_gallus 121636 ............................................. 121680

Anas_platyrhynchos_platyrhynchos 121681 GCATTGCTTCTGTTCTTATTACCAAGTAGCAGATGTGCCTTTCCT 121725
Gallus_gallus 121681 ............................................. 121725

Anas_platyrhynchos_platyrhynchos 121726 CTGAATAGAAGGCTTACTGTAGCAGAGCATGGAGACTGCATCCAT

121770
Gallus_gallus 121726 .............................................
121770

Anas_platyrhynchos_platyrhynchos 121771 TCTGGAGATCTGTAACACAATGCACAGTAGCTACTGATCTTTCTT
121815
Gallus_gallus 121771 .............................................
121815

Anas_platyrhynchos_platyrhynchos 121816 GAGACCTGTGCTGTTAATTTACCTGCTAAGTATCAATTTAGAGAA
121860
Gallus_gallus 121816 .............................................
121860

Anas_platyrhynchos_platyrhynchos 121861 AGGAGTTACTCCACTAATTATGTGTGTATCTTTCTATACATATAT
121905
Gallus_gallus 121861 .............................................
121905

Anas_platyrhynchos_platyrhynchos 121906 ATCTGGATTTCCACTGGCAGATGTCTTAAGAGGAAAGATGTGTAC
121950
Gallus_gallus 121906 .............................................
121950

Anas_platyrhynchos_platyrhynchos 121951 CTATGGTGTAATAGATGTATCCGTCAAGGTGTTCTCATGGTCTGC
121995
Gallus_gallus 121951 .............................................
121995

Anas_platyrhynchos_platyrhynchos 121996 ATTTCATCTGGCAGCTCAGAGTCTACAGCTCTTAACAGATGTTGA
122040
Gallus_gallus 121996 .............................................
122040

Anas_platyrhynchos_platyrhynchos 122041 GTTGTATGGAGGAAAAGAAAAATCCATCCAGGTTTACACGTAAAG
122085
Gallus_gallus 122041 .............................................
122085

Anas_platyrhynchos_platyrhynchos 122086 CTATGACATTACATCCTGACTCTTCTGACCTAAATATTGTGAAGA
122130
Gallus_gallus 122086 .............................................
122130

Anas_platyrhynchos_platyrhynchos 122131 CCCAAGGTCAACTACAGGATCACAACCAGTCCTTAATCTTTCATG
122175
Gallus_gallus 122131 .............................................
122175

Anas_platyrhynchos_platyrhynchos 122176 TAGCCTTTTATTGAAAGGTGTCAGTTTTCATCTTAAAAATGACAA
122220
Gallus_gallus 122176 .............................................
122220

Anas_platyrhynchos_platyrhynchos 122221 AGGGCAGCACTGTGAAGATACTGAGTTTTCTTTTTCCCCACTAAA
122265
Gallus_gallus 122221 .............................................
122265

Anas_platyrhynchos_platyrhynchos 122266 ATACTTCTCTGTACCGTTTCATGAAAACTACATGACGAAGAGGAT
122310
Gallus_gallus 122266 .............................................
122310

```
Anas_platyrhynchos_platyrhynchos 122311 ATGAAGAACTACTGATGGAACTTGCAAGCTATTTTGAGTGACCAT
122355
Gallus_gallus                    122311 ...........CTTGTGGAACACGCAAGCTGTTTCAAGAGTCCAT
122355

Anas_platyrhynchos_platyrhynchos 122356 TAGGTAATTTATCTGGGAAACGTATCTTTTGAGTGAGGTATGCTA
122400
Gallus_gallus                    122356 TGCAGAACTTAGCTGGGAGATGCATCTTTTGCATTAGACATGCTG
122400

Anas_platyrhynchos_platyrhynchos 122401 AACTCTGT--ACTTTACTTTTTGTTACAGCTCCTAGACGTGAGCC
122445
Gallus_gallus                    122401 AATTCTTTCTGCTTTCCATTTTGTTACAGTCCCTAAGCACAGGCA
122445

Anas_platyrhynchos_platyrhynchos 122446 TAGTGCCTTTGACAACTGTCGTTACCTTCGCTGACATGACAGAAT
122490
Gallus_gallus                    122446 TGGTGCCTTTGACAACAGTTGTTACCTTCACTGACATGACGGAAT
122490

Anas_platyrhynchos_platyrhynchos 122491 GGCCAGAACCTCCACATGGCCAGCCCAAAATACAGTGGAAACTCC
122535
Gallus_gallus                    122491 GGCCAGAACCTCCACGTGGTCTGCCCAAAGTACACTGGAAACTCC
122535

Anas_platyrhynchos_platyrhynchos 122536 CATTTGACTTCTTTTTCCCCTTCAAGGTGGTACTGAATGGAGAAA
122580
Gallus_gallus                    122536 CGTTTGACTTTTTTTTTCCATTCAAGATGGTCCTGAGTATTGAAA
122580

Anas_platyrhynchos_platyrhynchos 122581 GAAGCTACAGAGGTGATCTTGCTGCCTATTTTTTGTTGATTCTAA
122625
Gallus_gallus                    122581 GAAGTTACAGAGATGATTTGACTGCCTACTTTGTTGTGATTCTAA
122625

Anas_platyrhynchos_platyrhynchos 122626 TATTGTCTAGTATTCCTTGCCTTTGTTGAGAGTCAAACAGAAGAG
122670
Gallus_gallus                    122626 TACTGTCTAGTATTCTCTGCTTTTGAAGAGAGTCAAATAGATGGG
122670

Anas_platyrhynchos_platyrhynchos 122671 TTCAG----------TAAGTGAGTTAACACATTCATCTTATGATT
122715
Gallus_gallus                    122671
ACCTGTACTTAAAGTTACGTGAATTTATACATTTGTATACTTTT- 122715

Anas_platyrhynchos_platyrhynchos 122716 CAGATCATAAACTTTTCCTCTCAGGTTCTTATACTGTGCCTATGG
122760
Gallus_gallus                    122716 ----TTGTAAACTTTTTTCCTCAGATTCTTATACTGT--CTATAA
122760

Anas_platyrhynchos_platyrhynchos 122761 TATGTATGTGCCTTCTGTTCTTGGATTAAGCCACATGACTAGCTT
122805
Gallus_gallus                    122761 TAT--------CTTCTGTTGATGGATTAATTCACGTGACTACTTT
122805

Anas_platyrhynchos_platyrhynchos 122806 CTGTCCTGTGTCCTCTCACACTCCTGTTCTCCATACGCATGTTCT
122850
Gallus_gallus                    122806 CTGTCCTGTGTCCTCTCATTCTCCTGTTCTCCATATATACAATCT
122850

Anas_platyrhynchos_platyrhynchos 122851
GTAAA-----------CCTCACTGAAGTGAAATTCTGGAGGGA-- 122895
```

```
Gallus_gallus                      122851 TTAAAGTGAAATTTACACTCTTTGAAGTGAAATTCTTGAGGGATG
122895

Anas_platyrhynchos_platyrhynchos 122896 CGGGAGTTGTTGAGAGATGTTGCAGGTATGAGGAGCAGGAAGGAG
122940
Gallus_gallus                      122896 CGGGGGTAGTTGAAAGATGTTGCAAGAATGGGGAGCAGGAAGGAG
122940

Anas_platyrhynchos_platyrhynchos 122941 GAATTTAAAATGAACTTATTCTCAGAAACCAAGAGTGCTTTCTAA
122985
Gallus_gallus                      122941 GAAGTTAAAATGAACTTGTTCCCAGAAATC--GAGTTTTTTCTAA
122985

Anas_platyrhynchos_platyrhynchos 122986 ACCAATGAATGTGGAGAGAGGAGCGAAGTTCAGTTCAAGCATTGT
123030
Gallus_gallus                      122986 ACCAGGGAACATGGAGAGAG.........................
123030

Anas_platyrhynchos_platyrhynchos 123031 GGAGCTACATTCACTCATCTTGTTAGAGAAGCACCTGATTTTTCT
123075
Gallus_gallus                      123031 .............................................
123075

Anas_platyrhynchos_platyrhynchos 123076 ATGTGGAGCTAACAGAACACTTAAAATATGTGATGTCTCCCAGGA
123120
Gallus_gallus                      123076 .............................................
123120

Anas_platyrhynchos_platyrhynchos 123121 GACCCAAATTGTTATTTTGTGAGTGGGCGTGTCTGCCAGTCACCT
123165
Gallus_gallus                      123121 .............................................
123165

Anas_platyrhynchos_platyrhynchos 123166 TTGAGCTGTCCTTATTTTGTGACAGCCATGACTGTGTGACAATTT
123210
Gallus_gallus                      123166 .............................................
123210

Anas_platyrhynchos_platyrhynchos 123211 TTTTTCTATGTATTTTCAAGTATAGGTATTCAATTGAAATTTTCT
123255
Gallus_gallus                      123211 .............................................
123255

Anas_platyrhynchos_platyrhynchos 123256 TGATTTAATCACTGTTTGCATTAGCAATAGTGTTACGTGAAGTGA
123300
Gallus_gallus                      123256 .............................................
123300

Anas_platyrhynchos_platyrhynchos 123301 TGTAATTTGAGGTATTGCACGGACTGAAGTGTAACTGAATTTTAC
123345
Gallus_gallus                      123301 .............................................
123345

Anas_platyrhynchos_platyrhynchos 123346 AGTTGTTTGTTGAGACAGCTTCCTTCCTGCTCATGTGTATATGTG
123390
Gallus_gallus                      123346 .............................................
123390

Anas_platyrhynchos_platyrhynchos 123391 AGTGAACAGTTTTTCCTACTCAAGAAACATTCACCTGTGGCTCCA
123435
Gallus_gallus                      123391 .............................................
123435
```

```
Anas_platyrhynchos_platyrhynchos 123436 GATTGGAACAGGACTGATAATCTTAGAATCGTTCAGGTTGGAAAA
123480
Gallus_gallus                    123436 .............................................
123480

Anas_platyrhynchos_platyrhynchos 123481 GACCTCTACGATTGTCTAGTCTAACCTTTAAACTAGTACTGACTA
123525
Gallus_gallus                    123481 .............................................
123525

Anas_platyrhynchos_platyrhynchos 123526 GTCCACCACAAAACCACACTACTTAGCTCTACATGTAATGACAGC
123570
Gallus_gallus                    123526 .............................................
123570

Anas_platyrhynchos_platyrhynchos 123571 CACTGAAGTTCTAGGTGGCACATCAGAGAGCTGTGAACAGTCTTA
123615
Gallus_gallus                    123571 .............................................
123615

Anas_platyrhynchos_platyrhynchos 123616 AAGGACAACATCAATTATTTTAGGAGGAGTTGATGGGACTAAAGA
123660
Gallus_gallus                    123616 .............................................
123660

Anas_platyrhynchos_platyrhynchos 123661 TCACTGCAACACCTTACTCACCTGTGTGTGAGTACCCTACTCATC
123705
Gallus_gallus                    123661 .............................................
123705

Anas_platyrhynchos_platyrhynchos 123706 TGGGACCAGATTGTAAGTTGCTGTTTTTAAAGTCAAAGGTGATCA
123750
Gallus_gallus                    123706 .............................................
123750

Anas_platyrhynchos_platyrhynchos 123751 TTTTGATATCTAGTCTGACCCACTGAATCACGAAGACTAAGGAAA
123795
Gallus_gallus                    123751 .............................................
123795

Anas_platyrhynchos_platyrhynchos 123796 TCTGCTCAATCATTCCATTAAGTTAACTACTGTATATCTT-AGCA
123840
Gallus_gallus                    123796 ......CAGTTGTTTAAGTTGATTAACAACAGTAAATTTTTAGCT
123840

Anas_platyrhynchos_platyrhynchos 123841 AAGAATTTATTACTTATCCTAAATTCTTCA---AATGGAAAAATC
123885
Gallus_gallus                    123841 AAGGATTTATTATTTATCTTAGATTCCTCATATGATGGAAAAATC
123885

Anas_platyrhynchos_platyrhynchos 123886 TGCCACATCTAATGTTGAGTAATCCAGTGGAATCAATAATAGGCC
123930
Gallus_gallus                    123886 TGCCCCCTGTGAGGTTCAGTAATCCATTGGAATCA----CGGGTC
123930

Anas_platyrhynchos_platyrhynchos 123931 AGAAAA----------------AGAGAAATCTGG---------CC
123975
Gallus_gallus                    123931 AGAAAACCTGAGGACAAACAACAGAGAA---TGGGTCAGAAAACT
123975

Anas_platyrhynchos_platyrhynchos 123976 TGCCTTTGACAAAATGAAGTTGTTGCTTTTTTACTTAATACTGCA
124020
Gallus_gallus                    123976 TACCCTTGACAAAATG----TATAGATTTT--ACTTAATACTGCA
```

```
124020

Anas_platyrhynchos_platyrhynchos 124021 TTTAGCATAACTCTGTTGCTGTTCTTCATTATAACTTAAACAGAG
124065
Gallus_gallus                    124021 TTTATCGTAACTTCATTGTTGTTCCTCACTATTACTTCTGCACTA
124065

Anas_platyrhynchos_platyrhynchos 124066 CAGTGTACAAATAAACTGAAAAATTCTTATTTATAAGTCAGGACA
124110
Gallus_gallus                    124066 CTCTGT-----TTAACTGAAAAACTTGTATTTA----GTAGGATG
124110

Anas_platyrhynchos_platyrhynchos 124111 TCAAAGAAGATAACCTGGAT-------------AAAACTAAACAT
124155
Gallus_gallus                    124111 TCA---AAGATAAGCTTGATTTTAAAATAAAAAAAAAAAATACAT
124155

Anas_platyrhynchos_platyrhynchos 124156 GACCAGTATAAAACATGCATATTAGAGAGTAACTTCACAGTAACT
124200
Gallus_gallus                    124156 GGCCAGTATGAATCATGCTGATTA.....................
124200

Anas_platyrhynchos_platyrhynchos 124201 GATTGGAAAATCAGTTCTGATTGTTTGCATGGATTATTAGCTTGG
124245
Gallus_gallus                    124201 .............................................
124245

Anas_platyrhynchos_platyrhynchos 124246 ACAGAATCCTGTTCAAGTGACTAGTGCTTCCATTGCTCCTGATTC
124290
Gallus_gallus                    124246 .............................................
124290

Anas_platyrhynchos_platyrhynchos 124291 TGAAGCAGGCTACGTATGTTCTAGTGTTAGATGGGGTTGGTGATA
124335
Gallus_gallus                    124291 .............................................
124335

Anas_platyrhynchos_platyrhynchos 124336 GTACCAGTACAGCAATGAGCTTTTATATTAGATAGGTACGGTTGA
124380
Gallus_gallus                    124336 .............................................
124380

Anas_platyrhynchos_platyrhynchos 124381 GAAGCCTCTTCTAGGTCTTTGGAAAGCTTTTAGCCGCACACATTT
124425
Gallus_gallus                    124381 .............................................
124425

Anas_platyrhynchos_platyrhynchos 124426 GAAGAGTAGATCATGTCCTCTTATGCCCCTAGTATTTTTCAATCC
124470
Gallus_gallus                    124426 .............................................
124470

Anas_platyrhynchos_platyrhynchos 124471 ACAAAGTTAATTTATGGGTGAGAGTAACTTAATTGAAGAAAGAGT
124515
Gallus_gallus                    124471 .............................................
124515

Anas_platyrhynchos_platyrhynchos 124516 CAGTTGGAGAAACTTGCGAGTCATCACGCTGCTGTGTCAGGACAC
124560
Gallus_gallus                    124516 .............................................
124560

Anas_platyrhynchos_platyrhynchos 124561 AGCTTTAATTAAAAGAACATGGTGAATATGGCAGCTTTTTTTTTC
```

124605
Gallus_gallus 124561 .............................................
124605

Anas_platyrhynchos_platyrhynchos 124606 TGGCTTACTGCAATAATCCAATTAAGTGATTACAGGGACTTTTTT
124650
Gallus_gallus 124606 .............................................
124650

Anas_platyrhynchos_platyrhynchos 124651 TTATTCTACAAGCTGCAAAGACAGCTAAGTGCTGTAATAATGTGT
124695
Gallus_gallus 124651 .............................................
124695

Anas_platyrhynchos_platyrhynchos 124696 CACCTTACTATTACAATTCTTTCATATTCCTAGCTTCTAATATGT
124740
Gallus_gallus 124696 .............................................
124740

Anas_platyrhynchos_platyrhynchos 124741 CACTTTCTCTAGCACATTTTTGCTTCCTCTGAAATACCCTTATGA
124785
Gallus_gallus 124741 .............................................
124785

Anas_platyrhynchos_platyrhynchos 124786 CAGAAAATAGAGCTGGGAATAGCATTGTTCATAACTTGGTCAAGC
124830
Gallus_gallus 124786 .............................................
124830

Anas_platyrhynchos_platyrhynchos 124831 CTGCTTTAAGGTACGCTCAATTGTAGATGTTCCATACCACATAGT
124875
Gallus_gallus 124831 .............................................
124875

Anas_platyrhynchos_platyrhynchos 124876 TGTGGGGTTTAGAGTCCACAATTTCCTATGATAGTTTATGTTGGT
124920
Gallus_gallus 124876 .............................................
124920

Anas_platyrhynchos_platyrhynchos 124921 GTCTACCTTGTTGTGTTTGTATTTTCCCCTACTGTCTGAACTTAA
124965
Gallus_gallus 124921 .............................................
124965

Anas_platyrhynchos_platyrhynchos 124966 ATGACCTCTACCAGTCGAGTATCAGGGAGCAGTATGACAGTGTCA
125010
Gallus_gallus 124966 .............................................
125010

Anas_platyrhynchos_platyrhynchos 125011 CGCAAGGCAACGTTCATAAAATAATCAGGAGACTCTTATGCCAGA
125055
Gallus_gallus 125011 .............................................
125055

Anas_platyrhynchos_platyrhynchos 125056 TATGTAGAATAGGGTTTCTTGTTTGGCATATTTTTTGTAAAGCCC
125100
Gallus_gallus 125056 .............................................
125100

Anas_platyrhynchos_platyrhynchos 125101 CAGCTGAAACTGCTGCAGAAATCGCTCAGTGGGGGATGCCAAAGA
125145
Gallus_gallus 125101 .............................................
125145

```
Anas_platyrhynchos_platyrhynchos 125146 GGCATGTAAACATGCTGTTTTCCANNNNNNNNNNNNNNNNNNNNN
125190
Gallus_gallus                    125146 .............................................
125190

Anas_platyrhynchos_platyrhynchos 125191 NNNNNNNNNNNNNNNNNNNNNNNNNNNNNNNNNNNNNNNNNNNNN
125235
Gallus_gallus                    125191 .............................................
125235

Anas_platyrhynchos_platyrhynchos 125236 NNNNNNNNNNNNNNNNNNNNNNNNNNNNNNNNNNNNNNNNNNNNN
125280
Gallus_gallus                    125236 .............................................
125280

Anas_platyrhynchos_platyrhynchos 125281 NNNNNNNNNNNNNNNNNNNNNNNNNNNNNNNNNNNNNNNNNNNNN
125325
Gallus_gallus                    125281 .............................................
125325

Anas_platyrhynchos_platyrhynchos 125326 NNNNNNNNNNNNNNNNNNNNNNNNNNNNNNNNNNNNNNNNNNNNN
125370
Gallus_gallus                    125326 .............................................
125370

Anas_platyrhynchos_platyrhynchos 125371 NNNNNNNNNNNNNNNNNNNNNNNNNNNNNNNNNNNNNNNNNNNNN
125415
Gallus_gallus                    125371 .............................................
125415

Anas_platyrhynchos_platyrhynchos 125416 NNNNNNNNNNNNNNNNNNNNNNNNNNNNNNNNNNNNNNNNNNNNN
125460
Gallus_gallus                    125416 .............................................
125460

Anas_platyrhynchos_platyrhynchos 125461 NNNNNNNNNNNNNNNNNNNNNNNNNNNNNNNNNNNNNNNNNNNNN
125505
Gallus_gallus                    125461 .............................................
125505

Anas_platyrhynchos_platyrhynchos 125506 NNNNNNNNNNNNNNNNNNNNNNNNNNNNNNNNNNNNNNNNNNNNN
125550
Gallus_gallus                    125506 .............................................
125550

Anas_platyrhynchos_platyrhynchos 125551 NNNNNNNNNNNNNNNNNNNNNNNNNNNNNNNNNNNNNNNNNNNNN
125595
Gallus_gallus                    125551 .............................................
125595

Anas_platyrhynchos_platyrhynchos 125596 NNNNNNNNNNNNNNNNNNNNNNNNNNNNNNNNNNNNNNNNNNNNN
125640
Gallus_gallus                    125596 .............................................
125640

Anas_platyrhynchos_platyrhynchos 125641 NNNNNNNNNNNNNNNNNNNNNNNNNNNNNNNNNNNNNNNNNNNNN
125685
Gallus_gallus                    125641 .............................................
125685

Anas_platyrhynchos_platyrhynchos 125686 NNNNNNNNNNNNNNNNNNNNNNNNNNNNNNNNNNNNNNNNNNNNN
125730
```

```
Gallus_gallus                    125686 .............................................
125730

Anas_platyrhynchos_platyrhynchos 125731 NNNNNNNNNNNNNNNNNNNNNNNNNNNNNNNNNNNNNNNNNNNNN
125775
Gallus_gallus                    125731 .............................................
125775

Anas_platyrhynchos_platyrhynchos 125776 NNNNNNNNNNNNNNNNNNNNNNNNNNNNNNNNNNNNNNNNNNNNN
125820
Gallus_gallus                    125776 .............................................
125820

Anas_platyrhynchos_platyrhynchos 125821 NNNNNNNNNNNNNNNNNNNNNNNNNNNNNNNNNNNNNNNNNNNNN
125865
Gallus_gallus                    125821 .............................................
125865

Anas_platyrhynchos_platyrhynchos 125866 NNNNNNNNNNNNNNNNNNNNNNNNNNNNNNNNNNNNNNNNNNNNN
125910
Gallus_gallus                    125866 .............................................
125910

Anas_platyrhynchos_platyrhynchos 125911 NNNNNNNNNNNNNNNNNNNNNNNNNNNNNNNNNNNNNNNNNNNNN
125955
Gallus_gallus                    125911 .............................................
125955

Anas_platyrhynchos_platyrhynchos 125956 NNNNNNNNNNNNNNNNNNNNNNNNNNNNNNNNNNNNNNNNNNNNN
126000
Gallus_gallus                    125956 .............................................
126000

Anas_platyrhynchos_platyrhynchos 126001 NNNNNNNNNNNNNNNNNNNNNNNNNNNNNNNNNNNNNNNNNNNNN
126045
Gallus_gallus                    126001 .............................................
126045

Anas_platyrhynchos_platyrhynchos 126046 NNNNNNNNNNNNNNNNNNNNNNNNNNNNNNNNNNNNNNNNNNNNN
126090
Gallus_gallus                    126046 .............................................
126090

Anas_platyrhynchos_platyrhynchos 126091 NNNNNNNNNNNNNNNNNNNNNNNNNNNNNNNNNNNNNNNNNNNNN
126135
Gallus_gallus                    126091 .............................................
126135

Anas_platyrhynchos_platyrhynchos 126136 NNNNNNNNNNNNNNNNNNNNNNNNNNNNNNNNNNNNNNNNNNNNN
126180
Gallus_gallus                    126136 .............................................
126180

Anas_platyrhynchos_platyrhynchos 126181 NNNNNNNNNNNNNNNNNNNNNNNNNNNNNNNNNNNNNNNNNNNNN
126225
Gallus_gallus                    126181 .............................................
126225

Anas_platyrhynchos_platyrhynchos 126226 NNNNNNNNNNNNNNNNNNNNNNNNNNNNNNNNNNNNNNNNNNNNN
126270
Gallus_gallus                    126226 .............................................
126270
```

```
Anas_platyrhynchos_platyrhynchos 126271 NNNNNNNNNNNNNNNNNNNNNNNNNNNNNNNNNNNNNNNNNNNNN
126315
Gallus_gallus                    126271 .............................................
126315

Anas_platyrhynchos_platyrhynchos 126316 NNNNNNNNNNNNNNNNNNNNNNNNNNNNNNNNNNNNNNNNNNNNN
126360
Gallus_gallus                    126316 .............................................
126360

Anas_platyrhynchos_platyrhynchos 126361 NNNNNNNNNNNNNNNNNNNNNNNNNNNNNNNNNNNNNNNNNNNNN
126405
Gallus_gallus                    126361 .............................................
126405

Anas_platyrhynchos_platyrhynchos 126406 NNNNNNNNNNNNNNNNNNNNNNNNNNNNNNNNNNNNNNNNNNNNN
126450
Gallus_gallus                    126406 .............................................
126450

Anas_platyrhynchos_platyrhynchos 126451 NNNNNNNNNNNNNNNNNNNNNNNNNNNNNNNNNNNNNNNNNNNNN
126495
Gallus_gallus                    126451 .............................................
126495

Anas_platyrhynchos_platyrhynchos 126496 NNNNNNNNNNNNNNNNNNNNNNNNNNNNNNNNNNNNNNNNNNNNN
126540
Gallus_gallus                    126496 .............................................
126540

Anas_platyrhynchos_platyrhynchos 126541 NNNNNNNNNNNNNNNNNNNNNNNNNNNNNNNNNNNNNNNNNNNNN
126585
Gallus_gallus                    126541 .............................................
126585

Anas_platyrhynchos_platyrhynchos 126586 NNNNNNNNNNNNNNNNNNNNNNNNNNNNNNNNNNNNNNNNNNNNN
126630
Gallus_gallus                    126586 .............................................
126630

Anas_platyrhynchos_platyrhynchos 126631 NNNNNNNNNNNNNNNNNNNNNNNNNNNNNNNNNNNNNNNNNNNNN
126675
Gallus_gallus                    126631 .............................................
126675

Anas_platyrhynchos_platyrhynchos 126676 NNNNNNNNNNNNNNNNNNNNNNNNNNNNNNNNNNNNNNNNNNNNN
126720
Gallus_gallus                    126676 .............................................
126720

Anas_platyrhynchos_platyrhynchos 126721 NNNNNNNNNNNNNNNNNNNNNNNNNNNNNNNNNNNNNNNNNNNNN
126765
Gallus_gallus                    126721 .............................................
126765

Anas_platyrhynchos_platyrhynchos 126766 NNNNNNNNNNNNNNNNNNNNNNNNNNNNNNNNNNNNNNNNNNNNN
126810
Gallus_gallus                    126766 .............................................
126810

Anas_platyrhynchos_platyrhynchos 126811 NNNNNNNNNNNNNNNNNNNNNNNNNNNNNNNNNNNNNNNNNNNNN
126855
Gallus_gallus                    126811 .............................................
```

126855

Anas_platyrhynchos_platyrhynchos 126856 NNNNNNNNNNNNNNNNNNNNNNNNNNNNNNNNNNNNNNNNNNNNN
126900
Gallus_gallus                    126856 .............................................
126900

Anas_platyrhynchos_platyrhynchos 126901 NNNNNNNNNNNNNNNNNNNNNNNNNNNNNNNNNNNNNNNNNNNNN
126945
Gallus_gallus                    126901 .............................................
126945

Anas_platyrhynchos_platyrhynchos 126946 NNNNNNNNNNNNNNNNNNNNNNNNNNNNNNNNNNNNNNNNNNNNN
126990
Gallus_gallus                    126946 .............................................
126990

Anas_platyrhynchos_platyrhynchos 126991 NNNNNNNNNNNNNNNNNNNNNNNNNNNNNNNNNNNNNNNNNNNNN
127035
Gallus_gallus                    126991 .............................................
127035

Anas_platyrhynchos_platyrhynchos 127036 NNNNNNNNNNNNNNNNNNNNNNNNNNNNNNNNNNNNNNNNNNNNN
127080
Gallus_gallus                    127036 .............................................
127080

Anas_platyrhynchos_platyrhynchos 127081 NNNNNNNNNNNNNNNNNNNNNNNNNNNNNNNNNNNNNNNNNNNNN
127125
Gallus_gallus                    127081 .............................................
127125

Anas_platyrhynchos_platyrhynchos 127126 NNNNNNNNNNNNNNNNNNNNNNNNNNNNNNNNNNNNNNNNNNNNN
127170
Gallus_gallus                    127126 .............................................
127170

Anas_platyrhynchos_platyrhynchos 127171 NNNNNNNNNNNNNNNNNNNNNNNNNNNNNNNNNNNNNNNNNNNNN
127215
Gallus_gallus                    127171 .............................................
127215

Anas_platyrhynchos_platyrhynchos 127216 NNNNNNNNNNNNNNNNNNNNNNNNNNNNNNNNNNNNNNNNNNNNN
127260
Gallus_gallus                    127216 .............................................
127260

Anas_platyrhynchos_platyrhynchos 127261 NNNNNNNNNNNNNNNNNNNNNNNNNNNNNNNNNNNNNNNNNNNNN
127305
Gallus_gallus                    127261 .............................................
127305

Anas_platyrhynchos_platyrhynchos 127306 NNNNNNNNNNNNNNNNNNNNNNNNNNNNNNNNNNNNNNNNNNNNN
127350
Gallus_gallus                    127306 .............................................
127350

Anas_platyrhynchos_platyrhynchos 127351 NNNNNNNNNNNNNNNNNNNNNNNNNNNNNNNNNNNNNNNNNNNNN
127395
Gallus_gallus                    127351 .............................................
127395

Anas_platyrhynchos_platyrhynchos 127396 NNNNNNNNNNNNNNNNNNNNNNNNNNNNNNNNNNNNNNNNNNNNN

127440
Gallus_gallus 127396 .............................................
127440

Anas_platyrhynchos_platyrhynchos 127441 NNNNNNNNNNNNNNNNNNNNNNNNNNNNNNNNNNNNNNNNNNNNN
127485
Gallus_gallus 127441 .............................................
127485

Anas_platyrhynchos_platyrhynchos 127486 NNNNNNNNNNNNNNNNNNNNNNNNNNNNNNNNNNNNNNNNNNNNN
127530
Gallus_gallus 127486 .............................................
127530

Anas_platyrhynchos_platyrhynchos 127531 NNNNNNNNNNNNNNNNNNNNNNNNNNNNNNNNNNNNNNNNNNNNN
127575
Gallus_gallus 127531 .............................................
127575

Anas_platyrhynchos_platyrhynchos 127576 NNNNNNNNNNNNNNNNNNNNNNNNNNNNNNNNNNNNNNNNNNNNN
127620
Gallus_gallus 127576 .............................................
127620

Anas_platyrhynchos_platyrhynchos 127621 NNNNNNNNNNNNNNNNNNNNNNNNNNNNNNNNNNNNNNNNNNNNN
127665
Gallus_gallus 127621 .............................................
127665

Anas_platyrhynchos_platyrhynchos 127666 NNNNNNNNNNNNNNNNNNNNNNNNNNNNNNNNNNNNNNNNNNNNN
127710
Gallus_gallus 127666 .............................................
127710

Anas_platyrhynchos_platyrhynchos 127711 NNNNNNNNNNNNNNNNNNNNNNNNNNNNNNNNNNNNNNNNNNNNN
127755
Gallus_gallus 127711 .............................................
127755

Anas_platyrhynchos_platyrhynchos 127756 NNNNNNNNNNNNNNNNNNNNNNNNNNNNNNNNNNNNNNNNNNNNN
127800
Gallus_gallus 127756 .............................................
127800

Anas_platyrhynchos_platyrhynchos 127801 NNNNNNNNNNNNNNNNNNNNNNNNNNNNNNNNNNNNNNNNNNNNN
127845
Gallus_gallus 127801 .............................................
127845

Anas_platyrhynchos_platyrhynchos 127846 NNNNNNNNNNNNNNNNNNNNNNNNNNNNNNNNNNNNNNNNNNNNN
127890
Gallus_gallus 127846 .............................................
127890

Anas_platyrhynchos_platyrhynchos 127891 NNNNNNNNNNNNNNNNNNNNNNNNNNNNNNNNNNNNNNNNNNNNN
127935
Gallus_gallus 127891 .............................................
127935

Anas_platyrhynchos_platyrhynchos 127936 NNNNNNNNNNNNNNNNNNNNNNNNNNNNNNNNNNNNNNNNNNNNN
127980
Gallus_gallus 127936 .............................................
127980

```
Anas_platyrhynchos_platyrhynchos 127981 NNNNNNNNNNNNNNNNNNNNNNNNNNNNNNNNNNNNNNNNNNNNN
128025
Gallus_gallus                    127981 .............................................
128025

Anas_platyrhynchos_platyrhynchos 128026 NNNNNNNNNNNNNNNNNNNNNNNNNNNNNNNNNNNNNNNNNNNNN
128070
Gallus_gallus                    128026 .............................................
128070

Anas_platyrhynchos_platyrhynchos 128071 NNNNNNNNNNNNNNNNNNNNNNNNNNNNNNNNNNNNNNNNNNNNN
128115
Gallus_gallus                    128071 .............................................
128115

Anas_platyrhynchos_platyrhynchos 128116 NNNNNNNNNNNNNNNNNNNNNNNNNNNNNNNNNNNNNNNNNNNNN
128160
Gallus_gallus                    128116 .............................................
128160

Anas_platyrhynchos_platyrhynchos 128161 NNNNNNNNNCGGTGGCCGCGCTGCCGGCGGGAAGCCGCGGTCGCG
128205
Gallus_gallus                    128161 ............TGGCCGCGCCGC-GGCGGGAGATGGCGTTGGCG
128205

Anas_platyrhynchos_platyrhynchos 128206 AGG---TGCGGCCGGTGCCG-GAGCCGTAGGGGG-GCGG-GGCGG
128250
Gallus_gallus                    128206 CGGGAGTTTGGTCGTGGCTGCGTGTCGCCGAGCGCGCGGCGGCTG
128250

Anas_platyrhynchos_platyrhynchos 128251 GGCGGGGCGTGTGGCCG-CGCCGCCTCCGCTTGAGCGCCGCGGGG
128295
Gallus_gallus                    128251 TGCGAAG-GTGCGGCCGGCACTGCGTTATGTTGACCGCCG--GTG
128295

Anas_platyrhynchos_platyrhynchos 128296
ACCAGCTCTGCGGCTTGGGGTTAGGAGCCGCTGCC---------- 128340
Gallus_gallus                    128296 CTCAGCCGCGGGTCCCGGCG------GCCGCTGTCCGTTTCCCCT
128340

Anas_platyrhynchos_platyrhynchos 128341 ---GTGGCTCG--CTGCTCTCCCAGGAGCTCTCGTTGCTCCGCAG
128385
Gallus_gallus                    128341 CTTGTGACTCGGTCTG-GCCCCGACGAGGTTCCCTTGTAAACGAG
128385

Anas_platyrhynchos_platyrhynchos 128386 CGAGCCAAGCCTTGCGGCGGTTGAGG-TACGCCCCGTGGGGTGTG
128430
Gallus_gallus                    128386 TGAGCTAAGCCTTGCGACGACAGAGGGTACCGTACACACAGTGTG
128430

Anas_platyrhynchos_platyrhynchos 128431 CGATGTGCTGCCTGCGTCTGGAGGCGCCCCATCCTCCCGG---TC
128475
Gallus_gallus                    128431 CGCCGTGCTGCAGGTATCTGGGCAC---TCATCCTTCTGGCCCTC
128475

Anas_platyrhynchos_platyrhynchos 128476 TCAGGTTTTCTTGAGACAAGCGTGTAGGCTTTGTTTTACG-TTTT
128520
Gallus_gallus                    128476 TGGAGTTGCCTTGAG----TTGCATAGACTTTATTTCAGGCTTTT
128520

Anas_platyrhynchos_platyrhynchos 128521 CTTGATGGCCACAGAGCATTGCTGCTACGTGGAGTCATTGGCAGA
128565
```

```
Gallus_gallus                   128521 TTTGGTGGCTGTAGAGCATTACTGTTATACCGAGCCAACGACAGA
128565

Anas_platyrhynchos_platyrhynchos 128566 GTGAAGCTGCTGTTTGGATCTCTGCAAGCAGTAGAACCAATAAGT
128610
Gallus_gallus                   128566 ATGGAATTGCTCCTTGGGCCTTTGCGAGCAGTAGA--CAGAGAAT
128610

Anas_platyrhynchos_platyrhynchos 128611 AGCATCAATAAGAACTGTTCTGACTTGGTTCTTCTGGAATTTTGC
128655
Gallus_gallus                   128611 TCTCTGAA--------GCATTGGCTTCATTCATTGGGAGATTTGG
128655

Anas_platyrhynchos_platyrhynchos 128656 AAAAGCAGAGCTGTGGCAC-TGTGAGGCTTTCTTT---CTTTGCC
128700
Gallus_gallus                   128656 AAAAACAGGGCTGCTGCTCATCTTTGGCCTTCCTTTTGCTCTGTA
128700

Anas_platyrhynchos_platyrhynchos 128701 AGTCTCTTATGCCTACCGAGAAGATTTCTCCATGCCCAAAGAAAA
128745
Gallus_gallus                   128701 TGTCTCTTGGGCCTACCATGGAAGTTTCTCTGAGCCCCAAAAAAC
128745

Anas_platyrhynchos_platyrhynchos 128746 CATTTAAACCCCCCTGACACTTGCGTGAGACATAGGAGTTCTTAC
128790
Gallus_gallus                   128746 CATGTAAATCCTCCTGACACCTGCGTGAGACGCGCAGGTTATAAC
128790

Anas_platyrhynchos_platyrhynchos 128791 CTATGTTTTGTCTTAACACTTGGTGATAAGCCTGTGTTGGTTGCA
128835
Gallus_gallus                   128791 CCAGGCTTTGGCTTTACTCCTGGTGATAAGTCTGTGTCTGCTGCA
128835

Anas_platyrhynchos_platyrhynchos 128836
TAGCATTTCTTC------------ACAGGATCAGTTTTT------ 128880
Gallus_gallus                   128836 TGCTGTTTCTTCACGCAAGGTACAACAGGGCCTGTTTTTTTTTAT
128880

Anas_platyrhynchos_platyrhynchos 128881 -----GTAAGAAGAAACCTGATAGCTCTCTGAAAGAAGTAATAAA
128925
Gallus_gallus                   128881 TTTTAATGAGAAGGGATCTGATAGGTCTCAAAA............
128925

Anas_platyrhynchos_platyrhynchos 128926 ACTGCAGATAAAGGAGACTTGATGACCAAAACTTACACAGATTTC
128970
Gallus_gallus                   128926 .............................................
128970

Anas_platyrhynchos_platyrhynchos 128971 CCAAAACCCTCAGCAACTGTCTCTGTACTATGAAGTCAAGGACAG
129015
Gallus_gallus                   128971 .............................................
129015

Anas_platyrhynchos_platyrhynchos 129016 TTTTATTTTCTGAAATGACAAATAACTAAAATGCTTCCATAAATG
129060
Gallus_gallus                   129016 .............................................
129060

Anas_platyrhynchos_platyrhynchos 129061 TTTGTACCTGTGTTGCCTCTGTACCCTTAAAAACACTGGGTACTT
129105
Gallus_gallus                   129061 .............................................
129105
```

```
Anas_platyrhynchos_platyrhynchos 129106 CAAAGTCTCTGCCTTCTTTCTCTTGTCAGATTTGCTCTTGATAAC
129150
Gallus_gallus                    129106 .............................................
129150

Anas_platyrhynchos_platyrhynchos 129151 ACTGAAAAGCACCTTATAAAGGGTTAACATTTTCTTCTTTCTATT
129195
Gallus_gallus                    129151 .............................................
129195

Anas_platyrhynchos_platyrhynchos 129196 TTGTAGTAAAGTATTTATATTTAGGCACTTTAGAAAGGGTATTAC
129240
Gallus_gallus                    129196 .............................................
129240

Anas_platyrhynchos_platyrhynchos 129241 CTTTTCTGTGTTTGTGTTGGTGAAAGCTTTTTCCTTCAAGTTCTG
129285
Gallus_gallus                    129241 .............................................
129285

Anas_platyrhynchos_platyrhynchos 129286 TATGGATTTATTAGTTTGTGTGCTGTACTGCTTTCCTCTTCTAAA
129330
Gallus_gallus                    129286 .............................................
129330

Anas_platyrhynchos_platyrhynchos 129331 CTACTCTTTGCCAGGGTTGTTGAAAGTAGCAGAGTGATCGTCTCC
129375
Gallus_gallus                    129331 .............................................
129375

Anas_platyrhynchos_platyrhynchos 129376 CAGGCTTTTTCAGGTAATTTTTCAGGTGCTCTTCTTGCTGTTGTA
129420
Gallus_gallus                    129376 .............................................
129420

Anas_platyrhynchos_platyrhynchos 129421 TCGGTAGGAATATGAAAGCGATGGTATTTTCACAGTTTTCTACAG
129465
Gallus_gallus                    129421 .............................................
129465

Anas_platyrhynchos_platyrhynchos 129466 TATCCAGGGAAGAAAACCGCTGAGAGTCACTGTAGATGATACTCT
129510
Gallus_gallus                    129466 .............................................
129510

Anas_platyrhynchos_platyrhynchos 129511 GAAACATCAGTTTGAGGTGTTAGTGGCAGTCAAAAAGCAAATAGA
129555
Gallus_gallus                    129511 .............................................
129555

Anas_platyrhynchos_platyrhynchos 129556 ATGTTAGAAATTACCAGGCAAGGAATGAAGTGCAAAATGAGTATA
129600
Gallus_gallus                    129556 .............................................
129600

Anas_platyrhynchos_platyrhynchos 129601 TCATTTGATGTATTCATTTATATCGTTTGATATATCATTTGATGT
129645
Gallus_gallus                    129601 .............................................
129645

Anas_platyrhynchos_platyrhynchos 129646 ATCATGACCATATGTATCATGCCACTATAAATACATGCTCCAACC
129690
Gallus_gallus                    129646 .............................................
```

```
129690

Anas_platyrhynchos_platyrhynchos 129691 AACTCTGAGCATCGTGTGCTGTTCTGGGCTCGTTTTTCTGGCTAA
129735
Gallus_gallus                    129691 .............................................
129735

Anas_platyrhynchos_platyrhynchos 129736 CATGCTGTACTCTTTTGAGGTTAGTGGGGGAGGTCAAGGAAAAGC
129780
Gallus_gallus                    129736 .............................................
129780

Anas_platyrhynchos_platyrhynchos 129781 AGTGCACATGATCATAGATATGAAATAGCTTTTGTTTAAGAACAG
129825
Gallus_gallus                    129781 .............................................
129825

Anas_platyrhynchos_platyrhynchos 129826 CAAGAGCATCGTAAACAGCTTTATTAGTGGCAGTACAGTCTCAGG
129870
Gallus_gallus                    129826 .............................................
129870

Anas_platyrhynchos_platyrhynchos 129871 AATTGCTGTGTTTCTCTTCTGCTGGTGCTACTGCTCATGTGATTG
129915
Gallus_gallus                    129871 .............................................
129915

Anas_platyrhynchos_platyrhynchos 129916 TGTAGTCAGGTGGGTTGGTTGGTCTCTGACCAGAACATTAGTGAT
129960
Gallus_gallus                    129916 .............................................
129960

Anas_platyrhynchos_platyrhynchos 129961 AAAGCTGAACACTGGTTTAAGGAGTGGGATGGGAGCTGTAGCTTT
130005
Gallus_gallus                    129961 .............................................
130005

Anas_platyrhynchos_platyrhynchos 130006 GAGAAGGAGGGGTGGCAAAACTGACAATTGTGGTGACAGAACTGT
130050
Gallus_gallus                    130006 ....AGTAGGGGTGGCAAAACTGATGCTT------------CTGC
130050

Anas_platyrhynchos_platyrhynchos 130051 TTAGGAGCAACTTGAGCACTGCAGCCACAATTCACGTGCAGGAAT
130095
Gallus_gallus                    130051 TTAGGTGCAGCTTGGGCTCCACAGCCAT-------GGGGAGGAAT
130095

Anas_platyrhynchos_platyrhynchos 130096 GAGAAGGTGCGTGTTAC---AAGTCTGTAAGATACTAAGTGGTGT
130140
Gallus_gallus                    130096 GAGAAGGTATGTGC-ACGGGAAGTCCGCAGAATTACAGGTGCTGT
130140

Anas_platyrhynchos_platyrhynchos 130141 GGAGAAGGTGGATAGCCCCA--GTCACACAAAGATAGAGACCAGG
130185
Gallus_gallus                    130141 GGAGAAGGTGCATAGCACCATAGTCACACAAAGCTGGAGACCAGA
130185

Anas_platyrhynchos_platyrhynchos 130186 GGAGTGTTTCGGTAGTTTTAATCGTTTTCTTGTCACACCCATATA
130230
Gallus_gallus                    130186 GGAGTGCTTTAGGAATTTTAATTGTGTTCTTATCATAGCCATGTA
130230

Anas_platyrhynchos_platyrhynchos 130231 CTTTATTTGGGCATCTGATGGAAACTGCTGTCATACCAATTTTTG
```

```
130275
Gallus_gallus                     130231
CTGTATTTAGGAATTGGACACAAATTGCTGTCTTACTGATTTTT- 130275

Anas_platyrhynchos_platyrhynchos 130276 GGGGGTAAGTT------------------TTTCCCCATTAGGATT
130320
Gallus_gallus                     130276 -----TATTTTATTTTATTTTATTTTAAATTTCTCTATTAAGATT
130320

Anas_platyrhynchos_platyrhynchos 130321 GGGTGTTGGCTTATCCTTGGTAGAAACCAAACACCTTCTTTATTC
130365
Gallus_gallus                     130321 GG-TATCAGCTTATCTTTGATAAAAACAGAACGCCTTCTTTAATC
130365

Anas_platyrhynchos_platyrhynchos 130366 TCCCCAACCACAAAAGAATAAGTAAACACATCATTGTGAGGACAC
130410
Gallus_gallus                     130366
TC---AGCAATAAAAGAAGAGAAAAGCACATCATTGTGAGGACA- 130410

Anas_platyrhynchos_platyrhynchos 130411 AGAATAAAAAAAGAGAGGTAGGTAAAGGC-------TTAAAAATA
130455
Gallus_gallus                     130411 -AAACAAAAAAGGAGAGGTTGGTAAAGGCAAATGTCTTAAAAACA
130455

Anas_platyrhynchos_platyrhynchos 130456 AGGAACTGGAG-GGAATGCCA-------------------TCAGG
130500
Gallus_gallus                     130456 GGGAACTGAAATGGAATACAAAGCTGAGTACTGCATGGAGTTGAG
130500

Anas_platyrhynchos_platyrhynchos 130501 ATACTGATAAGTGCTGTTTTAAGTGATACTCTTCATTGAGAAGTG
130545
Gallus_gallus                     130501 ATTATAATAAATGCTATTTTGAGTGATGCTCTTCTTGGAGAAATG
130545

Anas_platyrhynchos_platyrhynchos 130546 ACAGGACAAGGACCTATAAATAAATCATACAGCTGGTGATTTTTC
130590
Gallus_gallus                     130546 ACAGGG--------TATGAACAAATCTTACTGTTGGTGAATTTCC
130590

Anas_platyrhynchos_platyrhynchos 130591 CCTGTCTTTCTTCTCAGGATGCTATGGTTTTCTGGCTTAAAGACC
130635
Gallus_gallus                     130591 CCTGTCTTTCTTAGCAAGAGGCTGCAGCCAGCTGGCTTGCAGGTT
130635

Anas_platyrhynchos_platyrhynchos 130636 GCTGTAAGGCAGTTGTTGAAGAGGTTC----CACAATGGGGCCTT
130680
Gallus_gallus                     130636 ACTATAAGGCAGTAG---AAGAGATTCATTCCATAGTGGGGGCTT
130680

Anas_platyrhynchos_platyrhynchos 130681 GGATACAGAGGGTATGAATAAATAATTTAGAAGGGTGCAAAACTG
130725
Gallus_gallus                     130681 GGGTATGGAGGGTTTGAAAAAATAATTTGGAAGGGTGAAGAAGTG
130725

Anas_platyrhynchos_platyrhynchos 130726
CATCTAAAGCTTCTATAATACATGCTGGAGGAGCTG--------- 130770
Gallus_gallus                     130726 CATCTAAAGCTTCAATAAGATGTACTGGAGGAGCTGTAAGAAGCT
130770

Anas_platyrhynchos_platyrhynchos 130771 -------ATAGGGTACTAAATGTAGATATCAGTCTTTGTCCGGGA
130815
Gallus_gallus                     130771 GACTAACTGAGGGTACTAACTGTAAATGTTCATCTTTCTCCTGAA
130815
```

```
Anas_platyrhynchos_platyrhynchos 130816 ATATTGAACAAGACTGTAGACTTCAGTTTATATTTTTCGAATTTC
130860
Gallus_gallus                    130816 ATATTGAACAAAACTGCTGACTTCAGTTTACATTTTTGAAA---C
130860

Anas_platyrhynchos_platyrhynchos 130861 TTTTCTTTCTATCCTTCTGTTTCTCTGGCTACATTTACTCGTGAA
130905
Gallus_gallus                    130861 TTTTCTTTCTATCCTTACATTTCTGTGGCTGCATTTACTCCTGAA
130905

Anas_platyrhynchos_platyrhynchos 130906
TATGAATGTGTGTCCAGGTACTGAATTTTCATCTTGTCATTCAG- 130950
Gallus_gallus                    130906 CAAGAGCATGTGTCCAGGTACTCGATTTGCATCTTATCTCTCAGT
130950

Anas_platyrhynchos_platyrhynchos 130951 GACACTAGTGTCAGTATATTAGCCAGAGCTATTATAAAATCTTTT
130995
Gallus_gallus                    130951 GATACTAGTGTTACTGTTTTAGCTGTAGCTTTTATAAAAACTTTA
130995

Anas_platyrhynchos_platyrhynchos 130996 TTCCGTCTCCGTTGCTTTCCC-GAACGACTTTTGATGCTTGTACA
131040
Gallus_gallus                    130996 TTCAGTCT--GTTTCTTTCCCTGAATGACTTGTGATGGTTGTACA
131040

Anas_platyrhynchos_platyrhynchos 131041 GCTTTGATTTGATGATTTGATGTCTGTAGGGAAACTGATACAGGA
131085
Gallus_gallus                    131041 ACCTTGATTTGATGG--------CCATAGAAAAACTGATCCAGAA
131085

Anas_platyrhynchos_platyrhynchos 131086 CTGAGATTTTAGCTCTGTTGTCTTTATCTTTAACAGCAGGTGGAA
131130
Gallus_gallus                    131086
CTGAGATGTTTTCTATCATCTCTTTATCCTTAATGGCAGATG--- 131130

Anas_platyrhynchos_platyrhynchos 131131 TCTGATGTTTCACTGTTTGGTGTTAAGAACAGGTTTTAGAGGAAA
131175
Gallus_gallus                    131131 ---GATGTTTTACTGCTTGGTG-------AAGGTTTTAGAGGAAA
131175

Anas_platyrhynchos_platyrhynchos 131176 GTAGCCAGATTTGATCCAGGCAAG--AGTAGAGTGGTGATGCTAA
131220
Gallus_gallus                    131176 TT------ATTTGATCCAGGCCAGAAAAGAGAGCGATGATTTTTG
131220

Anas_platyrhynchos_platyrhynchos 131221 GACGAGGGAAATAAATGAAAGATGCTAACTGACCCTGTCTCTAGG
131265
Gallus_gallus                    131221
GAAGAAGGAAATACTGGAAAAATGCTAACTGAACC---------- 131265

Anas_platyrhynchos_platyrhynchos 131266 TTTCATAGTGATGACCAGCACTTGATTAAATTTACCTGTGTCCTC
131310
Gallus_gallus                    131266 ------ACTGATGACT-GTGCTTAATTAAA-----CTGTGTCCTC
131310

Anas_platyrhynchos_platyrhynchos 131311 ATGTAGTAATCTGGATTAAAACACCTACTCTTTCTTGGTTTTGCT
131355
Gallus_gallus                    131311 ACAT........................TTTTTGGTGGTGGTGGC
131355

Anas_platyrhynchos_platyrhynchos 131356 TGTTTTTGTTCAATTAGAGAAAAATGCAAACCTTTTGTTTAGAGA
131400
```

```
Gallus_gallus                   131356 AATTTTTTTTTTATTGGAGAAAAAAGTGAAGTCTTTGAATAGAGA
131400

Anas_platyrhynchos_platyrhynchos 131401 TATACCTACCTATAATTGTAGCAATTGTACCTAGGACTGAAGGTG
131445
Gallus_gallus                   131401 TAAGCCT-------------------GTGATCAGGACTGCAGGTG
131445

Anas_platyrhynchos_platyrhynchos 131446 GACTGTAGCATGCTGGCCTACGCATATTGTTACTGCCACCAGGAT
131490
Gallus_gallus                   131446 GGATGTAGCATGTTCACCTCCACGTGTTGTTAATGTTATCGGTAT
131490

Anas_platyrhynchos_platyrhynchos 131491 GGTTCTTTGGCCTTTTCT-TTGAAGATGAGGCTCTGCTTGAGATT
131535
Gallus_gallus                   131491 GGTTCCTGTGCTTTTTTTCTTGAAGATGGAACTCTTCTTGAGATA
131535

Anas_platyrhynchos_platyrhynchos 131536 TCTCAG--TGTTCACTACCCTTAACTATCTGAAACCA---GTCTT
131580
Gallus_gallus                   131536 TATCAGAGTATTCACTACCTTTAACAATCTGAAACCACAGTTTTT
131580

Anas_platyrhynchos_platyrhynchos 131581 TCATTGCTGACTTCCTTATGTGTTTGTTGTACACTGAAGTAAATT
131625
Gallus_gallus                   131581 CCACTGCTGACTTCCTAGCATACTTGCTGTACATTGAGGTAAACT
131625

Anas_platyrhynchos_platyrhynchos 131626 ACTGCTCATCAGGCAGGAAATCTAGGATTCTTTGCTGTTAACAAG
131670
Gallus_gallus                   131626 GCTACTCTTCAGGGAAGAAATCTTGGACTCCTTGATGCTAACAAG
131670

Anas_platyrhynchos_platyrhynchos 131671
CATCCTTCTGTAAAAGCACTCAGTTTTGTTTGGTAAAGGAAAG-- 131715
Gallus_gallus                   131671 CATCCTTGTGTAAAAGCTAAAAGTTTTATATGGGAAACAGAAGTG
131715

Anas_platyrhynchos_platyrhynchos 131716 --GTTTTTGTT---TATTTTCTCATACCAATTCTTCCTTTATTTC
131760
Gallus_gallus                   131716 CTGATTTAGCTCGTTAGTTTCTCTTACCAATTCCTCTCTTATTTC
131760

Anas_platyrhynchos_platyrhynchos 131761 AGTGTGTGAAGAGCAAAAGCAGCTCTTCAGTAGCTGAAGGTGTTG
131805
Gallus_gallus                   131761 AGCATG--AGGAGCCAAAGCATCTCATCAGTGGCTGCAGGGGTTG
131805

Anas_platyrhynchos_platyrhynchos 131806 AGGGGAACCTTTTGCTTTTTATTTAGAAGCACGGTTCAAAGGAGT
131850
Gallus_gallus                   131806 AGGGGAACCTGTTCCTTTTTATTTGCAAGCATAGATCAGAGGAGT
131850

Anas_platyrhynchos_platyrhynchos 131851 TGCATGTGCTAAAAAGTGGACAGATCAATCTAAAATAAGGGCATT
131895
Gallus_gallus                   131851
TGCATGTTCAAAAAATTGTCTAGCTCAATCTAAAATAGGG----- 131895

Anas_platyrhynchos_platyrhynchos 131896 CCTGTTTTTTGTTAGGTGTTTAAAAATAAAATTGAAGGCTTAGCA
131940
Gallus_gallus                   131896 CACGTTTCTTGTTATGTGTTTAAAAATCAAAAGGAAGAGTTAGCA
131940
```

```
Anas_platyrhynchos_platyrhynchos 131941 GCTTTTT
131947
Gallus_gallus                    131941 GCTCTTT
131947
```
