## Supplementary Text for "Concurrent loss of ciliary genes *WDR93* and *CFAP46* in phylogenetically distant birds"

**Section S1: Gene curation and re-annotation**

We used NCBI datasets to gather the data of all species with annotated *WDR93* orthologs (Human GeneID: 56964). The *WDR93* gene annotation occurs in ~464 vertebrate species; 33 are annotated as low-quality proteins. The low-quality remark in NCBI was due to corrections such as substitution at genomic stop codon, deletion of bases, or insertion of bases to obtain an open reading frame. Careful verification of such changes showed that these corrections were often false and caused by errors in genomic assemblies, such as gapped assemblies, assemblies with low coverage, or assemblies made using error-prone sequencing technologies such as PacBio. For further verification, we searched other genome assembly versions of the same species to determine whether the corrections suggested by NCBI were correct. If the genome assemblies lacked annotation, we used TOGA (Tools to infer orthologs from genome alignment) for gene annotation to find the gene intactness or gene disruption events. Subsequently, we performed a blast search of the raw read data to confirm the correctness of the genome assemblies. For details regarding the curation approaches used, see **Supplementary Table 1**.

Prominent examples of assembly or annotation correction that recovered an intact *WDR93* gene include the flier cichlid (*Archocentrus centrarchus*), white killdeer (*Charadrius vociferus*), nine-banded armadillo (*Dasypus novemcinctus*), colocolo opossum (*Dromiciops gliroides*), ballan wrasse (*Labrus bergylta*), Wuchang bream (*Megalobrama amblycephala*), brown mesite (*Mesitornis unicolor*), Rio pearlfish (*Nematolebias whitei*), spotty (*Notolabrus celidotus*), Dalmatian pelican (*Pelecanus crispus*), leopard coral grouper (*Plectropomus leopardus*), Gibel carp (*Carassius gibelio*), copperband butterflyfish (*Chelmon rostratus*), Etruscan shrew (*Suncus etruscus*), green swordtail (*Xiphophorus hellerii*), downy woodpecker (*Dryobates pubescens*), bar-tailed Trogon (*Apaloderma vittatum*), rabbit (*Oryctolagus cuniculus*), elephant (*Loxodonta africana*), leopard (*Panthera pardus*), California sea lion (*Zalophus californianus*), Chinese river dolphin (*Lipotes vexillifer*) and Velvety free-tailed bat (*Molossus molossus*). Although genome assemblies are available, gene annotation was missing for many vertebrates (such as *Heterohyrax brucei*, *Microgale talazaci*, *Procavia capensis*, *Axis porcinus*, *Catagonus wagneri*, *Muntiacus muntjac*, *Rangifer tarandus*, *Acinonyx jubatus*, *Lynx pardinus*, *Ursus americanus*, *Ursus arctos*, *Miniopterus natalensis*, *Molossus molossus*, *Chiloscyllium punctatum*, *Hemiscyllium ocellatum*, *Scyliorhinus torazame*, *Galemys pyrenaicus*, *Scalopus aquaticus*, *Solenodon paradoxus, Chiloscyllium punctatum, Hemiscyllium ocellatum, Scyliorhinus torazame, Galemys pyrenaicus, Scalopus aquaticus,* and *Solenodon paradoxus*) and was re-annotated using TOGA. In the order, Perissodactyla, species of the Rhinocerotidae family (*Ceratotherium simum*, *Diceros bicornis*, *Dicerorhinus sumatrensis*, and *Rhinoceros unicornis*) and Tapiridae family (*Tapirus indicus* and *Tapirus terrestris*) were found to have frame-shift changes in the 16^th^ exon leading to a premature stop codon resulting in proteins of 656/673/680 amino acids compared to 771 amino acids in the Equidae family (*Equus asinus*, *Equus caballus*, *Equus przewalskii*, and *Equus quagga*) (see **Supplementary Table 1**). We had to inspect the RNA-seq data for the central bearded dragon (*Pogona vitticeps*) and Townsend's dwarf sphaero (*Sphaerodactylus townsendi*) to verify and re-annotate the *WDR93* gene using the expressed transcript. In the case of a few species, such as *Eptesicus fuscus* and *Myotis lucifugus,* gaps in the genome prevented us from recovering the entire gene. Few other species, such as the burrowing owl (*Athene cunicularia*), had unclear data.

The European rabbit (*Oryctolagus cuniculus*) genome version GCA_000003625 was found to have gaps in the *WDR93* gene region, and Ensembl annotated the gene as a pseudogene. However, a more recent version (GCF_009806435.1) of the genome assembly was complete in the region, ruling out gene loss. RNA-seq data (SRR12227967 and SRR12228022) supports the expression of the exonic regions of the *WDR93* gene mapped to the recent genome. In the swan goose (*Anser cygnoides*), the *WDR93* region is incomplete in the genome assemblies GCF_002166845.1 and GCA_000971095.1, whereas the GCA_013030995.1 genome is gapless at the *WDR93* gene locus.

Gene annotation was missing for several species of the superorder Galloanserae. Hence, we evaluated the coding status of the *WDR93* gene in ~44 species of Galloanserae. The NCBI annotation of *WDR93* in *Struthio camelus* (Ostrich) and *Anas platyrhynchos* (Mallard) was used to screen the genomes of Galloanserae species.

**Section S2: Validation of segmental deletions and size estimation**

The *CFAP46* gene-containing region flanked by the *TCTN3* and *OPNVA* genes is missing from the genome assembly in chicken. However, this region is intact in the mallard (*Anas platyrhynchos*) genome. The lack of this region in the chicken genome could result from **(a)** an assembly error or **(b)** a translocation event, or **(c)** a segmental deletion event. We validated the correctness of the genome assembly using PacBio long-read sequencing data in chicken (*Gallus gallus*), mallard (*Anas platyrhynchos*), and swan goose (*Anser cygnoides*) (see **Supplementary Figure 31**, **30,** and **34**) to rule out the possibility of an assembly error. In the case of chicken (*Gallus gallus*), we also verified the assembly using Nanopore reads ≥ 80 Kb. Our search of the short-read and long-read blast databases using the *CFAP46* gene from the mallard genome as a query failed to recover the gene. If the *CFAP46* gene were translocated to another location in the genome, we would have expected to recover the reads corresponding to this translocated copy. The lack of such reads matching the *CFAP46* query supports the scenario of a segmental deletion in the chicken genome. We downloaded the pairwise genome alignment between mallard (*Anas platyrhynchos*) and chicken (*Gallus gallus*) from the Ensembl genome browser in the *CFAP46* gene-containing region. The pairwise alignment was consistent with a segmental deletion scenario (see **Supplementary File S1**). Unfortunately, the mallard genome version (GCA_002743455.1) available on the Ensembl genome browser at the time of the analysis had gaps in the assembly at the *CFAP46* gene-containing region. Hence, we used a newer version (GCF_015476345.1) of the genome without gaps in the assembly at the *CFAP46* gene-containing region for assembly validation using long-read data and generation of the dot plot (**Fig. 8** and **Supplementary Figure 30-31**, **34-36**).

The *CFAP46* gene region is deleted in the rifleman (*Acanthisitta chloris*), ruff (*Calidris pugnax*), Anna's hummingbird (*Calypte anna*), and speckled mousebird (*Colius striatus*). We searched short-read datasets of all four species with query sequence of *CFAP46* gene from mallard (*Anas platyrhynchos;* XM_038181248.1), common swift (*Apus apus*; XM_051616636.1), kākāpō (*Strigops habroptilus*; XM_030486345.1), the zebra finch (*Taeniopygia guttata*; XM_030276648.3) and ostrich (*Struthio camelus*; XM_009670554.1). However, we failed to recover the *CFAP46* gene from these four species. We found hits for *CFAP46* exon-61 of mallard in the rifleman (*Acanthisitta chloris*) genome. Short-read data supported the genomic sequence corresponding to the exon-61 in the rifleman bird.

The exact boundaries of genomic rearrangements are challenging to identify as the orthologous regions diverge over time and tend to be repeat-rich, making accurate alignments error-prone. Hence, we first performed a blast-based search of the focal genomes using the mallard genomic sequence from the *TCTN3* start and *OPNVA* end (1 kb each) as a query. In addition to the regions identified by the blast search in the focal genome, we considered 50 Kb flanking regions beside each hit as the region of interest. We extracted the nucleotide sequence of this entire region slightly longer than 100 Kb from each focal genome. These extracted sequences served as the query for a blastn search of the mallard (*Anas platyrhynchos*) repeat masked chromosome 6 (*CFAP46* containing region). The resultant blast hits were discontinuous but located nearby except for a large region of ~90 Kb that lacked hits (see **Supplementary Figure 33**). We used the bedtools merge command with the -d 30000 option to merge the nearby hits to extend the alignments in regions of poor sequence identity. We found one large region orthologous to the *CFAP46* gene region lacking blast hits, corresponding to the segmental deletion (see **Supplementary Figure 32**). The size of the deletion was estimated as the length of this region lacking blast hits. Interestingly, the deletion size was considerably different in each independent event but somewhat comparable among species sharing a common deletion event.

**Section S3: Raw read-based verification of gene loss events**

Gene loss candidates were initially identified based on the results of TOGA and blastn search of the genome assemblies. Genome assemblies can have base pair-level errors artefactually introduced during genome assembly. Hence, the nucleotide sequence at the genomic positions corresponding to the gene-disrupting events requires verification in the raw-read datasets. The short-read datasets were searched using blastn with the gene sequences as queries to asses raw-read support. The nucleotide sequence of reads identified by blast search was extracted and reformatted for inspection in the Mview utility for easy visualization. Gene-disrupting changes at the exon boundaries are harder to reformat using Mview. Hence, the exonic sequence is extended in these cases by including the flanking intronic sequence. For instance, the premature stop codon in Exon-14 occurs just one base before the splice site in the Anser genus. Using intronic flank sequence as a query helped recover the raw reads supporting the genomic sequence (see **Fig. 4**). The reformatting done by Mview is error-prone when two or more events occur in close proximity. Hence, we generated multiple sequence alignments for the region of interest using the clustalW aligner in MegaX.

The rifleman (*Acanthisitta chloris*) and Anna's hummingbird (*Calypte anna*) are missing several consecutive *WDR93* exons towards the 3’ end. While the lack of consecutive exons could result from gaps or errors in the genome assembly, searching raw read datasets can provide a more definitive answer. Our search of short-read databases for the missing exons using the *WDR93* query sequences from the mallard (*Anas platyrhynchos*), the zebra finch (*Taeniopygia guttata*), the barn owl (*Tyto alba*), common swift (*Apus apus*) and ostrich (*Struthio camelus*) failed to recover these exons at the 3’ end. To ensure the correctness of genome assembly, we used the long-read sequencing data from PacBio. The reads were mapped to the genome of these species using BWA and filtered for reads ≥ 10 Kb. The presence of overlapping reads that span the gene remnants at the syntenic region establishes the validity of the genome assembly. The PacBio long-read assembly verification of the *WDR93* gene was done for the swan goose (*Anser cygnoides*), mallard (*Anas platyrhynchos*), rifleman (*Acanthisitta chloris*), Anna's hummingbird (*Calypte anna*), and chicken (*Gallus gallus*). In the case of chicken (*Gallus gallus*), we also verified assembly correctness using Nanopore reads ≥ 100 Kb spanning the entire region of the *WDR93* gene and its flanking genes.

GC-biased gene conversion can lead to strong sequence divergence between closely related species contributing to rapid change in the gene sequence. For instance, in the sand rat (*Psammomys obesus*), highly divergent regions have resulted from GC-biased changes (Hargreaves et al., 2017). Identification of orthologous sequences in such divergent regions is challenging due to uneven and rapid changes. Moreover, genomic regions with extreme GC content are hard to recover using short-read technologies like Illumina sequencing (Chen et al., 2013). Hence, to ensure that such artifacts don’t affect the inference of gene loss, we quantified the magnitude of gBGC (see **Methods**). In the case of *WDR93*, certain species were found to have weak to intermediate levels of GC-biased gene conversion (see **Supplementary Table 9**). However, all cases of *WDR93* gene loss involve gene-disrupting mutations in addition to exonic loss. Moreover, the exon loss events have been verified using long-read sequencing and are unlikely to be an artifact of GC-biased gene conversion. In contrast to *WDR93*, we found evidence of strong gBGC in the *CFAP46* gene (see **Supplementary Table 13-14**). The loss of *CFAP46* through a large segmental deletion could be verified using long-read sequencing. However, the possibility of translocation to another genomic locus and strong gBGC complicates verifying *CFAP46* gene loss. Nonetheless, our search of short-read and long-read sequencing datasets failed to find any evidence of translocation. We found sequence saturation in the Actinopterygii clade potentially due to exon structure changes/ annotation issues. Hence, the molecular evolutionary analysis within this clade is not reliable.

**Section S4: Other functions of *WDR93* and *CFAP46***

We discuss the host-pathogen interaction changes based on the species that have experienced *WDR93* and *CFAP46* gene loss. However, several other important biological roles of these genes are reported in the literature. For instance, the assembly and disassembly process of cilia is closely linked to cell division, and the impairment or misregulation of these processes can lead to oncogenesis (Fliegauf et al., 2007). Changes in the expression of *WDR93* have been noted in lung squamous cell carcinoma (Tian et al., 2017), esophageal carcinoma (Sheng et al., 2021), and hepatocellular carcinoma (Wang et al., 2017). Prevalence of *CFAP46* fusion transcripts and aberrant methylation-associated gene expression changes are reported in hepatocellular carcinoma (Tsuge et al., 2019) and nasopharyngeal carcinoma (Ayadi et al., 2014), respectively. The expression of *WDR93* is downregulated in *NDP* gene knockout and has been associated with Norrie disease and other neuronal disorders (Alazami et al., 2015; Hayashi et al., 2021). Both *WDR93* and *CFAP46* are highly expressed in the testis (McKenzie et al., 2015), and *WDR93* has also been identified at the protein level using multiple assays (Vandenbrouck et al., 2016). Hence, the loss of these genes may be related to a phenotype associated with the sperm.
