## Supplementary Figures for "Concurrent loss of ciliary genes *WDR93* and *CFAP46* in phylogenetically distant birds"

**Figure S1:*WDR93* and *NDUSF4* are highly diverged from each other**

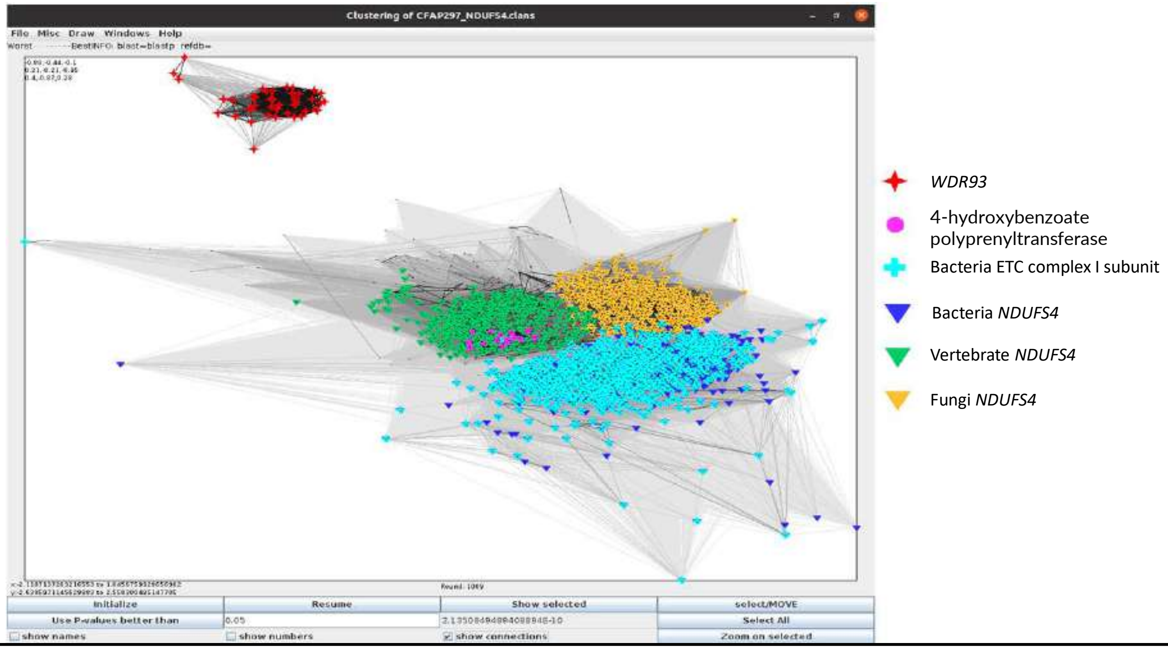


**Supplementary Figure 1: CLANS (CLuster ANalysis of Sequences) for IPR006885 (NADH_UbQ_FeS_4_mit-like) domain**. The IPR006885 domain which is shared by *WDR93* and *NDUFS4*, shows the high levels of divergence between the paralogs. The E-value was <1e-4 and p-value < 0.05 used while doing CLANS, and screenshot taken after 1009 rounds of iteration.

**Figure S2: Frame-disrupting deletion in Exon-6 of the Australian brush turkey (*Alectura lathami*)**

```
Struthio camelus Exon6      ggcagAAAAGTCTGGAGCTG
SRR10019968.19894788        ggcagAAAA-TCTGGAGCAG
SRR10019966.36801233        ggcagAAAA-TCTGGAGCAG
SRR10019966.30444977        ggcagAAAA-TCTGGAGCAG
SRR10019966.25524677        ggcagAAAA-TCTGGAGCAG
SRR10019966.38768001        ggcagAAAA-TCTGGAGCAG
SRR10019966.14067359        ggcagAAAA-TCTGGAGCAG
SRR10019968.8712523         ggcagAAAA-TCTGGAGCAG
SRR10019968.35890367        ggcagAAAA-TCTGGAGCAG
```

**Supplementary Figure 2:** Illumina short read data (SRA) supports the deletion of G base from Exon-6 of Australian brush turkey (*Alectura lathami*) in comparison to Ostrich (*Struthio camelus*).

**Figure S3: Short-read data supports the validity of the magpie goose (*Anseranas semipalmata*) genome assembly**

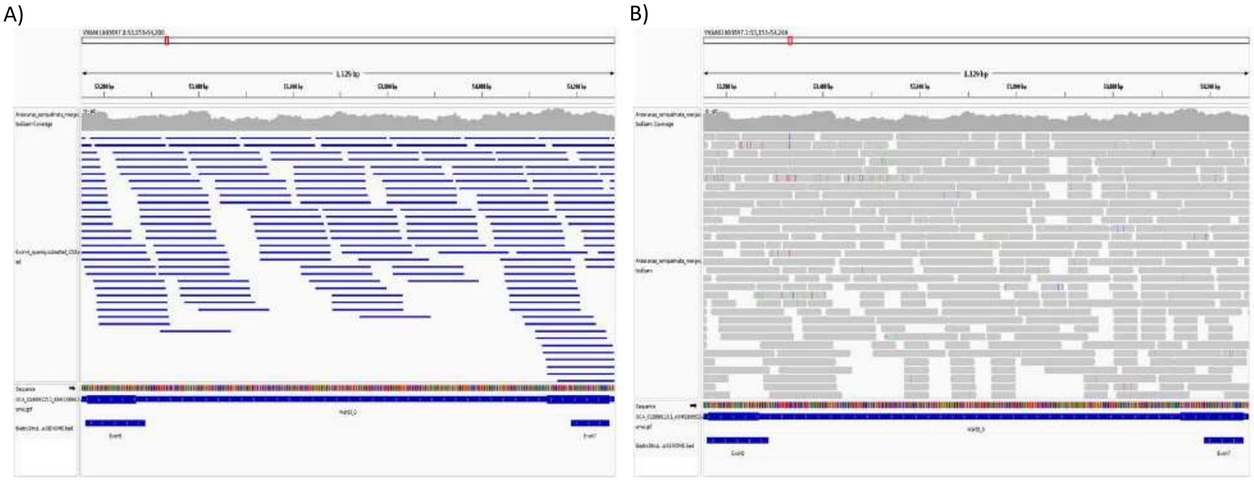


**Supplementary Figure 3: A)** IGV screenshot showing the continuous overlapping of 150 bp reads (in bed format) spanning the region between *WDR93* Exon-5 and Exon-7 of magpie goose (*Anseranas semipalmata*). **B)** IGV screenshot showing the continuous overlapping of Illumina reads (in bam format) spanning the region between *WDR93* Exon-5 and Exon-7 of magpie goose (*Anseranas semipalmata*). The SRR10019943, SRR10019944, SRR10019946, and SRR9946951 short read datasets mapped to GCA_013399115.1 genome of magpie goose (*Anseranas semipalmata*) for verification.

**Figure S4: Swan goose (*Anser cygnoides*) assembly verification at *WDR93* gene locus**

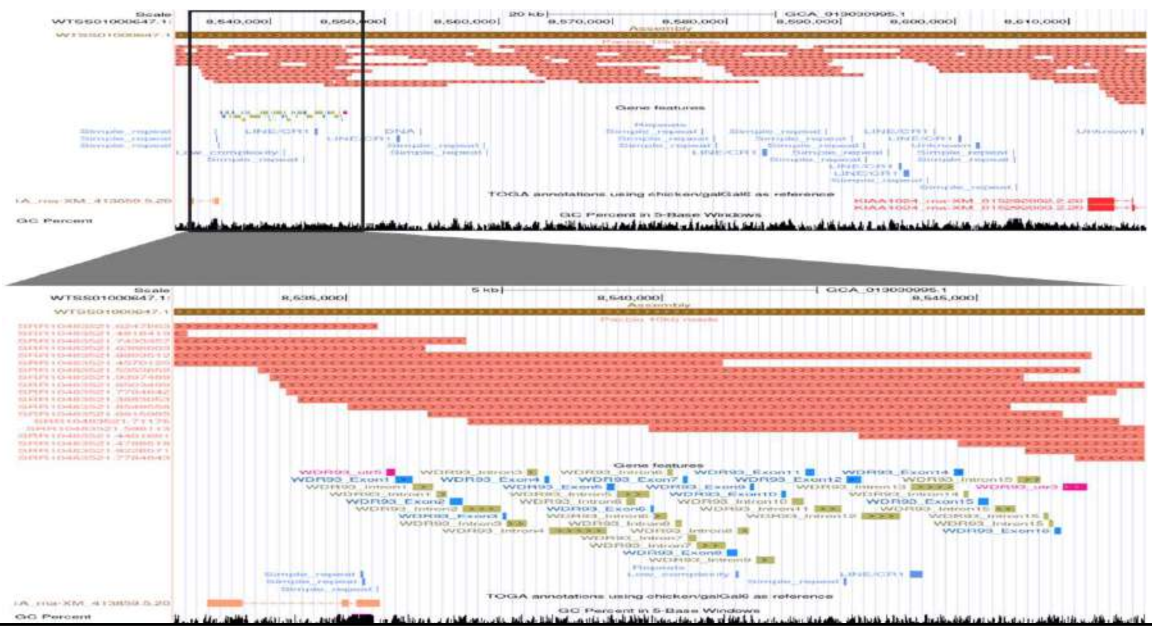


**Supplementary Figure 4: Verification of the Swan goose (*Anser cygnoides*) genome assembly using long reads.** The UCSC genome browser image showing the alignment of ≥10 kb long reads generated using PacBio (SRR12418405, SRR10483521) sequencing (salmon colored), aligned to the swan goose (GCA_013030995.1) at the syntenic location of the *WDR93* gene remnants. The image at the bottom is the zoomed-in view of the region highlighted (focusing on the *WDR93* remnants). Blue-colored boxes represent the *WDR93* exon remnants, olive-green-colored boxes show the *WDR93* introns, and the pink box represents the 5' and 3' UTR (shown in the genomic features track). The RepeatMasker BED track shows the repeats in that region (indicated by blue boxes). The accession IDs of the reads are specified on the left side of the reads beside them. Overlapping reads are found to span the entire region at the syntenic location of the *WDR93* gene, including the flanking genes (*KIAA1024/MINAR1* and *PEX11A*).

**Figure S5: SRA data supports the polymorphic frame-disrupting events in Canada goose (*Branta canadensis*)**

Exon 2: GC -> GT/--

```
Struthio camelus Exon2     AAGACAGAGGTTTGTGCCATCCATGCCTCGGACTT
SRR8867566.192711190       AAGACAGAAATTTGTGTCATCCGCTCCTCGGAGTT
SRR8867566.40549898        AAGACAGAAATTTGTGTCATCCGCTCCTCGGAGTT
SRR8867566.306892560       AAGACAGAAATTTGTGTCATCCGCTCCTCGGAGTT
SRR8867566.402489482       AAGACAGAAATTTGTGTCATCCGATCCTCGGAGTT
SRR8867566.102040110       AAGACAGAAATTTGTGTCATCCGCTCCTCGGAGTT
SRR8867566.97091013        AAGACAGAGATTTGT--CATCCGCTCCTCGGACTT
SRR8867566.289880847       AAGACAGAGATTTGT--CATCCGCTCCTCGGACTT
SRR8867566.321493811       AAGACAGAGATTTGT--CATCCGCTCCTCGGACTT
SRR8867566.7502768         AAGACAGAGATTTGT--CATCCGCTCCTCGGACTT
SRR8867566.190699042       AAGACAGAGATTTGT--CATCCGCTCCTCGGACTT
```

Exon 14: insertion of T

```
Struthio camelus Exon14    GCCCTGATCTT-CTCCTGGGATGGCACAGTGT
SRR8867566.382733177       GCCCTGTTTTTTCTCCTGGGATGGCACGGTGT
SRR8867566.264008827       GCCCTGTTTTTTCTCCTGGGATGGCACGGTGT
SRR8867566.344767627       GCCCTGTTTTTTCTCCTGGGATGGCACGGTGT
SRR8867566.138652956       GCCCTGTTTTTTCTCCTGGGATGGCACGGTGT
SRR8867566.193304061       GCCCTGTTTTTTCTCCTGGGATGGCACGGTGT
SRR8867566.363217162       GCCCTGGTTTT-CTCCTGGGATGGCACAGTGT
SRR8867566.380610938       GCCCTGGTTTT-CTCCTGGGATGGCACAGTGT
SRR8867566.121491133       GCCCTGGTTTT-CTCCTGGGATGGCACAGTGT
SRR8867566.252462180       GCCCTGGTTTT-CTCCTGGGATGGCACAGTGT
SRR8867566.65061994        GCCCTGGTTTT-CTCCTGGGATGGCACAGTGT
```

**Supplementary Figure 5:** Short read data supports the polymorphic deletion of GT base in Exon-2 and Insertion of T base in Exon-14 of Canada goose (*Branta canadensis*) compared to Ostrich (*Struthio camelus*). The red and blue colors show the two different haplotypes.

**Figure S6: Chinese alligator (*Alligator sinensis*) *WDR93* Expressed: IGV screenshot**

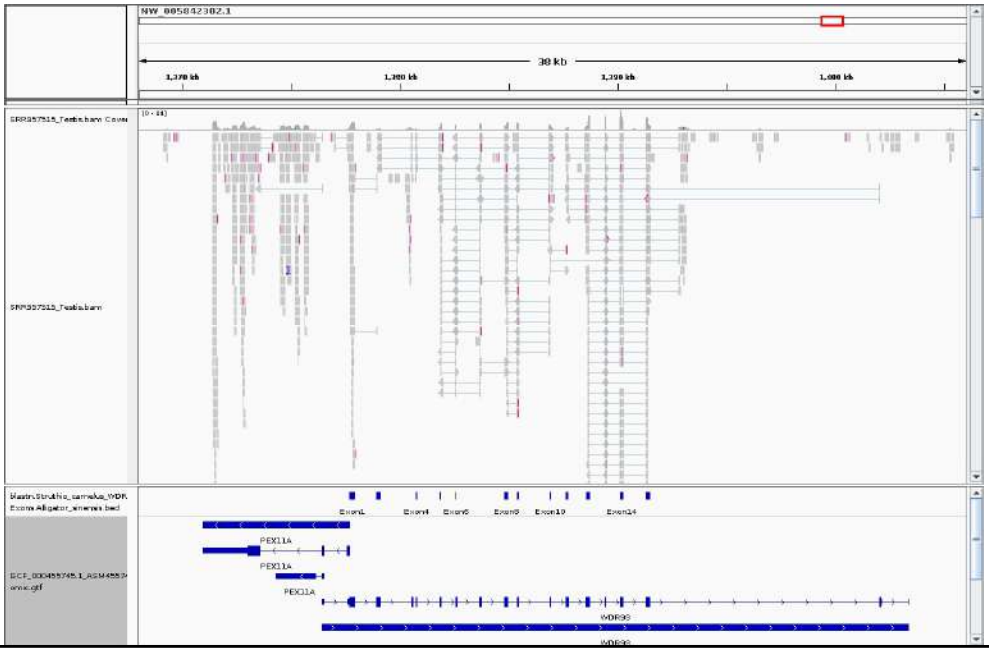


**Supplementary Figure 6: The *WDR93* gene is expressed in the gonads of the Chinese alligator (*Alligator sinensis*).** IGV screenshot for *WDR93* expression for testis (SRR957515) mapped to GCF_000455745.1 genome assembly of Chinese alligator (*Alligator sinensis*), using STAR mapper.

**Figure S7: Chinese alligator (*Alligator sinensis*) *WDR93* Expressed: Sashimi plot**

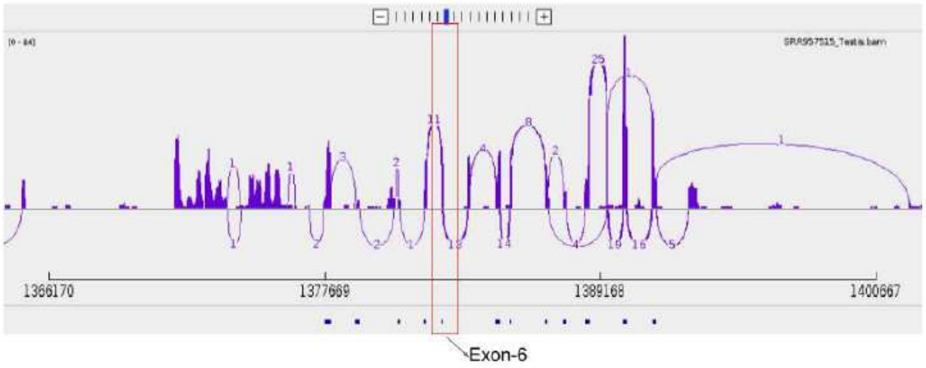


**Supplementary Figure 7:** The sashimi plot showing the splice junction at the Exon boundaries of the Chinese alligator (*Alligator sinensis*). All sixteen exons are expressed, including Exon6 (shown in a red rectangular box).

**Figure S8: Green anole (*Anolis carolinensis*) *WDR93* Expressed: IGV screenshot**

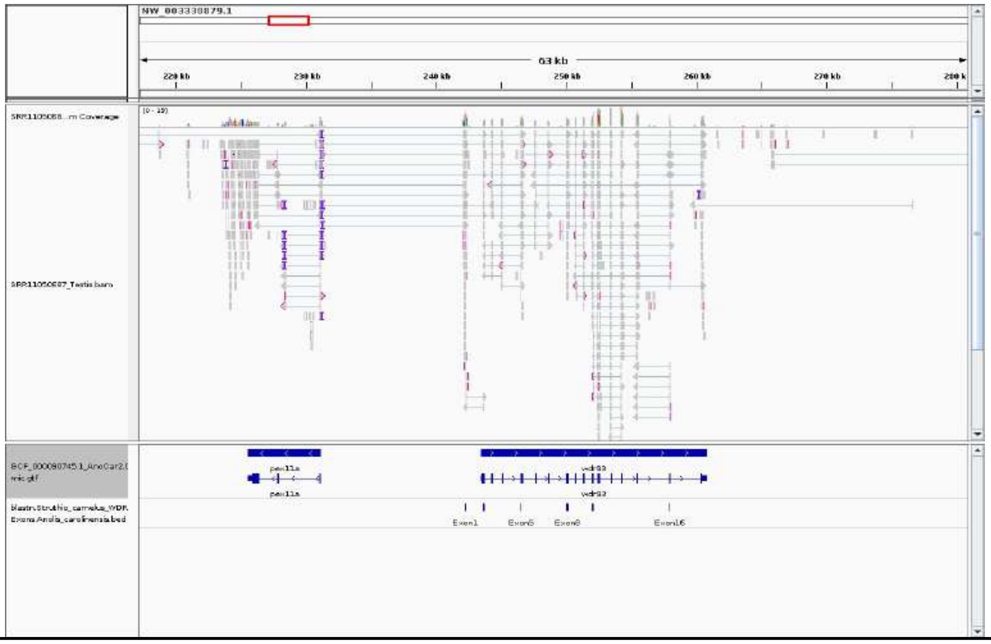


**Supplementary Figure 8: The *WDR93* gene is expressed in the gonads of green anole (*Anolis carolinensis*).** IGV screenshot for *WDR93* expression for Testis (SRR11050687) mapped to GCF_000090745.1 genome assembly of Green anole (*Anolis carolinensis*), using STAR mapper.

**Figure S9: Green anole (*Anolis carolinensis*) *WDR93* Expressed: Sashimi plot**

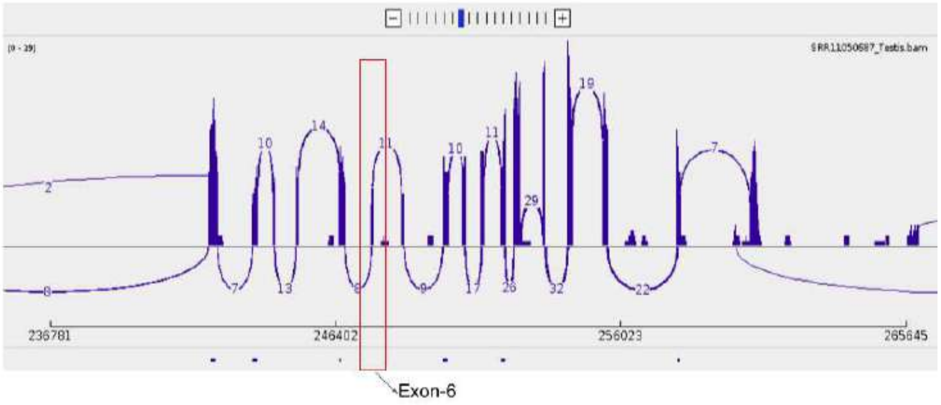


**Supplementary Figure 9:** The sashimi plot showing the splice junction at Exon boundaries of green anole (*Anolis carolinensis*). All sixteen exons are expressed, including Exon6 (shown in a red rectangular box).

**Figure S10: Ostrich (*Struthio camelus*) *WDR93* Expressed: IGV screenshot**

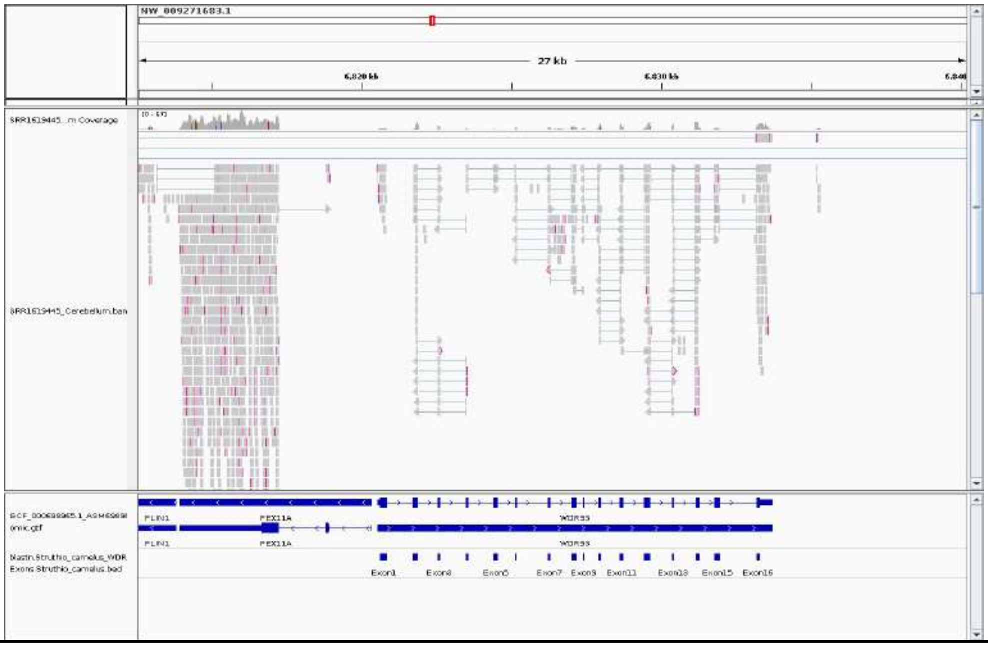


**Supplementary Figure 10: The *WDR93* gene is expressed in the cerebellum of ostrich (*Struthio camelus*).** IGV screenshot for *WDR93* expression for Cerebellum (SRR1619445) mapped to GCF_000698965.1 genome assembly of ostrich (*Struthio camelus*), using STAR mapper.

**Figure S11: Ostrich (*Struthio camelus*) *WDR93* Expressed: Sashimi plot**

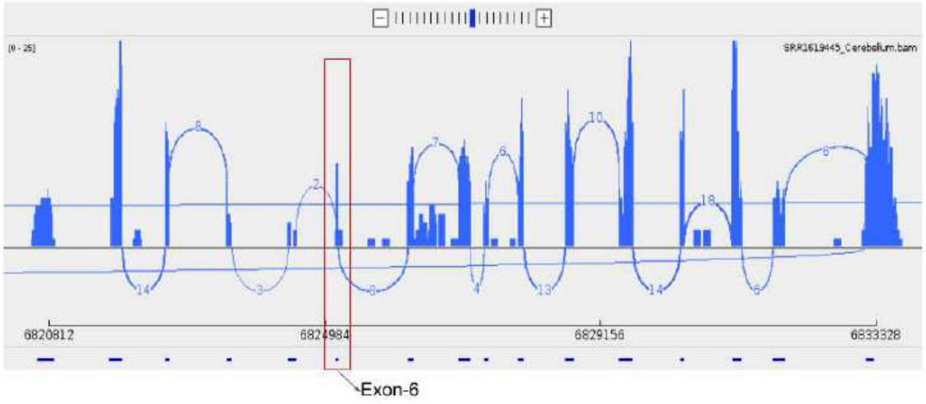


**Supplementary Figure 11:** The sashimi plot showing the splice junction at the Exon boundaries of Ostrich (*Struthio camelus*). All sixteen exons are expressed, including Exon6 (shown in a red rectangular box).

**Figure S12: Emu (*Dromaius novaehollandiae*) *WDR93* Expressed: IGV screenshot**

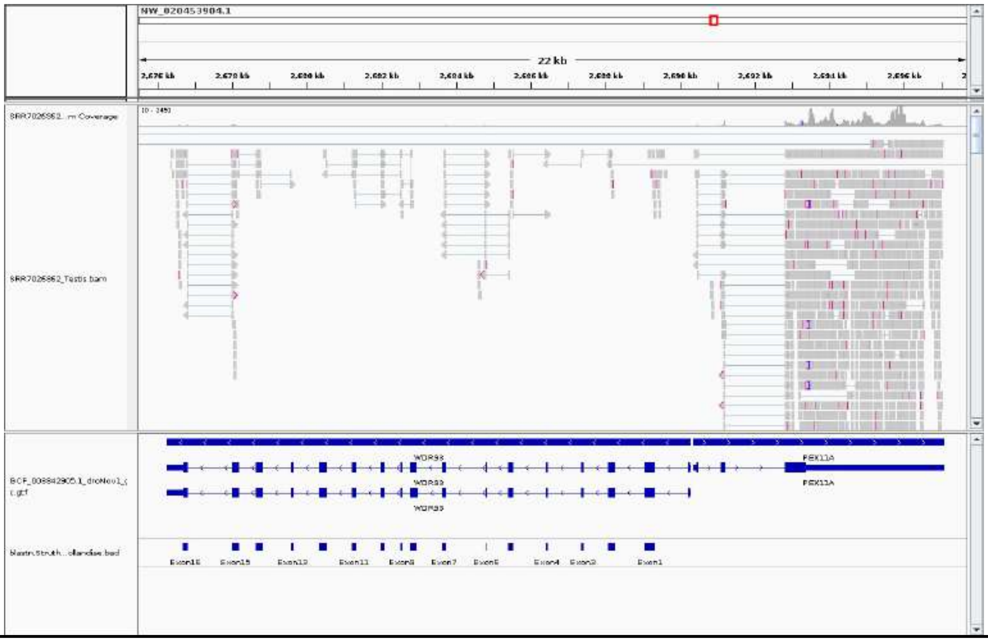


**Supplementary Figure 12: The *WDR93* gene is expressed in the testis of emu (*Dromaius novaehollandiae*).** IGV screenshot for *WDR93* expression for Testis (SRR7026362) mapped to GCF_003342905.1 genome assembly of emu (*Dromaius novaehollandiae*), using STAR mapper.

**Figure S13: Emu (*Dromaius novaehollandiae*) *WDR93* Expressed: Sashimi plot**

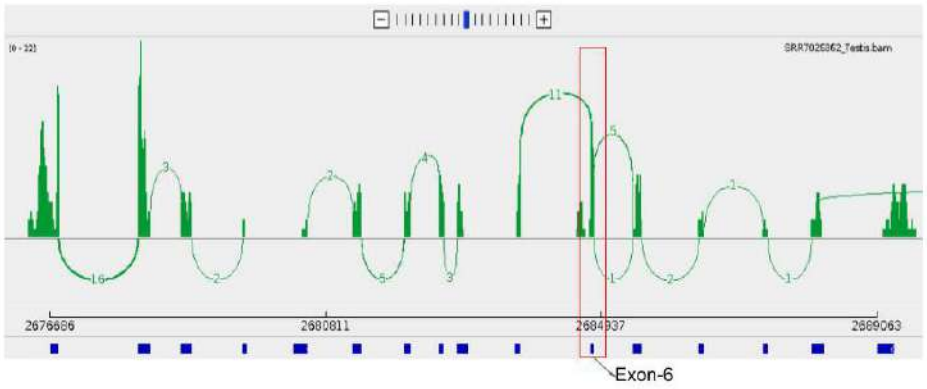


**Supplementary Figure 13:** The sashimi plot showing the splice junction at the Exon boundaries of emu (*Dromaius novaehollandiae*). All sixteen exons are expressed, including Exon6 (shown in a red rectangular box).

**Figure S14: Mallard (*Anas platyrhynchos*) *WDR93* Expressed: IGV screenshot**

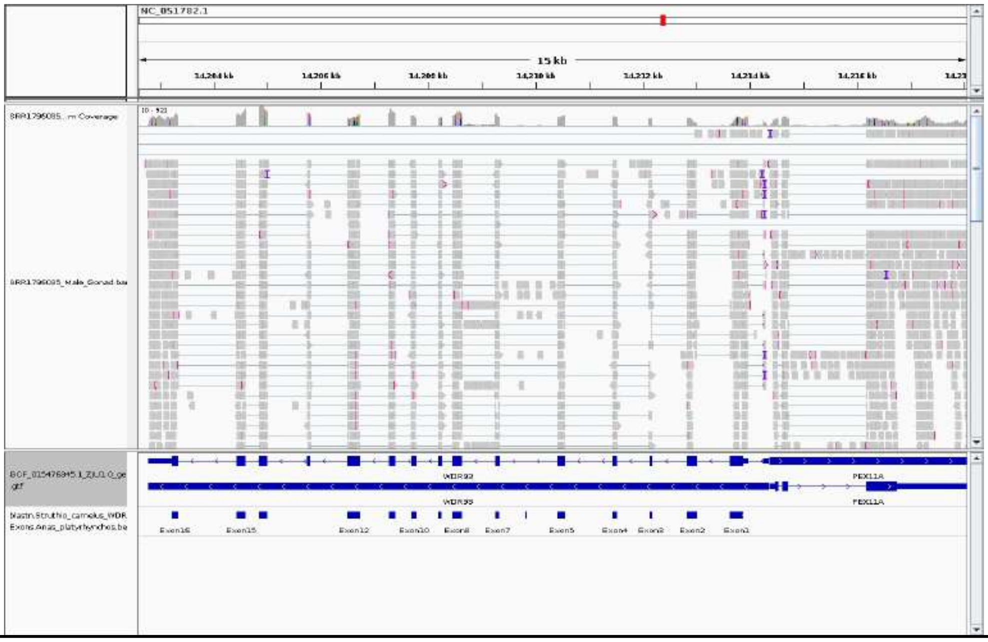


**Supplementary Figure 14: The *WDR93* gene is expressed in the testis of mallard (*Anas platyrhynchos*).** IGV screenshot for *WDR93* expression for male gonad (SRR1796035) mapped to GCF_015476345.1 genome assembly of mallard (*Anas platyrhynchos*), using STAR mapper.

**Figure S15: Mallard (*Anas platyrhynchos*) *WDR93* Expressed: Sashimi plot**

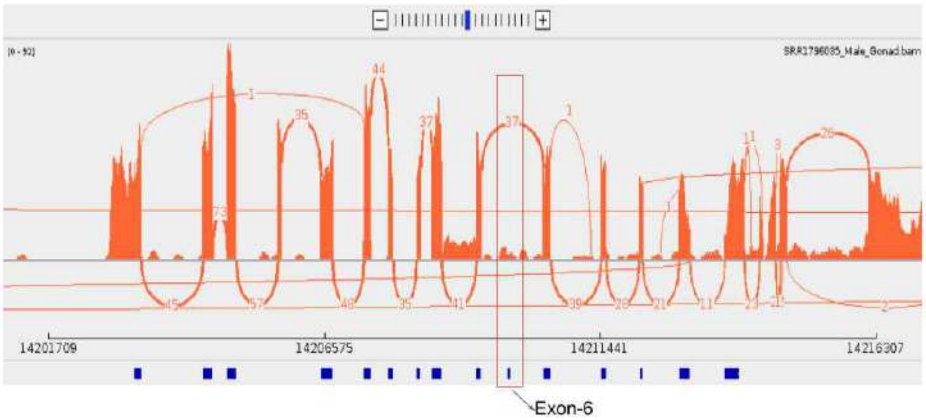


**Supplementary Figure 15:** The sashimi plot showing the splice junction at the Exon boundaries of mallard (*Anas platyrhynchos*). Only fifteen exons are expressed, and Exon6 is not expressed, shown in a red rectangular box.

**Figure S16: Muscovy duck (*Cairina moschata*) *WDR93* Expressed: IGV screenshot**

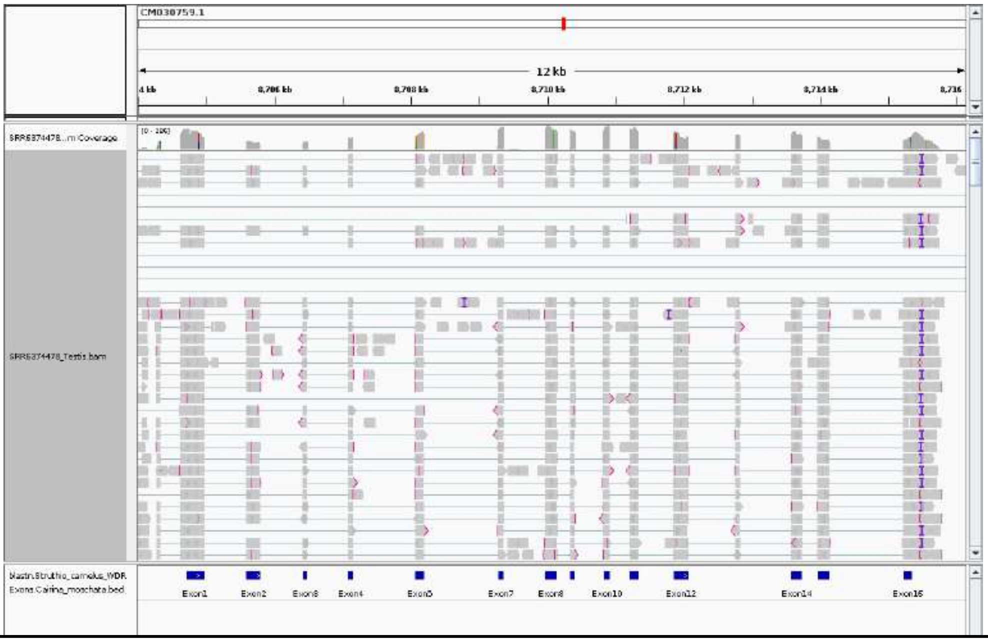


**Supplementary Figure 16: The *WDR93* gene is expressed in the testis of muscovy duck (*Cairina moschata*).** IGV screenshot for *WDR93* expression for male gonad (SRR6374478) mapped to GCA_018104995.1 genome assembly of muscovy duck (*Cairina moschata*), using STAR mapper.

**Figure S17: Muscovy duck (*Cairina moschata*) *WDR93* Expressed: Sashimi plot**

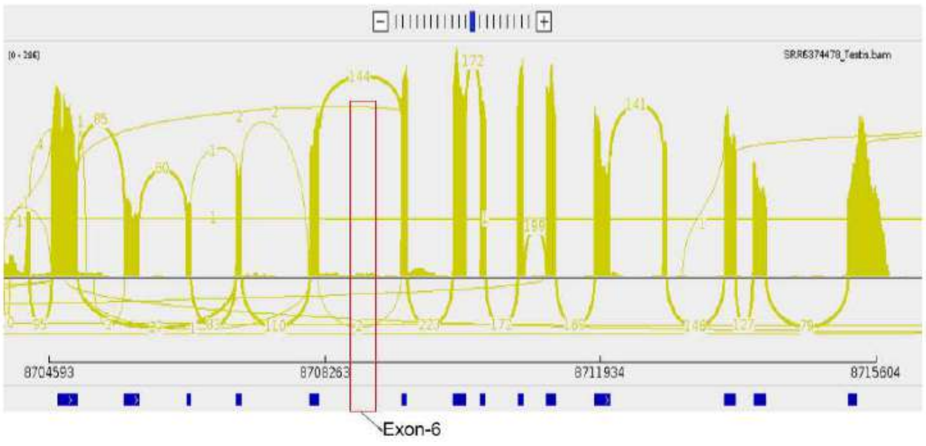


**Supplementary Figure 17:** The sashimi plot showing the splice junction at Exon boundaries of mallard (*Anas platyrhynchos*). Only fifteen exons are expressed, and Exon6 is not expressed, shown in a red rectangular box.

**Figure S18: Great tit (*Parus major*) *WDR93* Expressed: IGV screenshot**

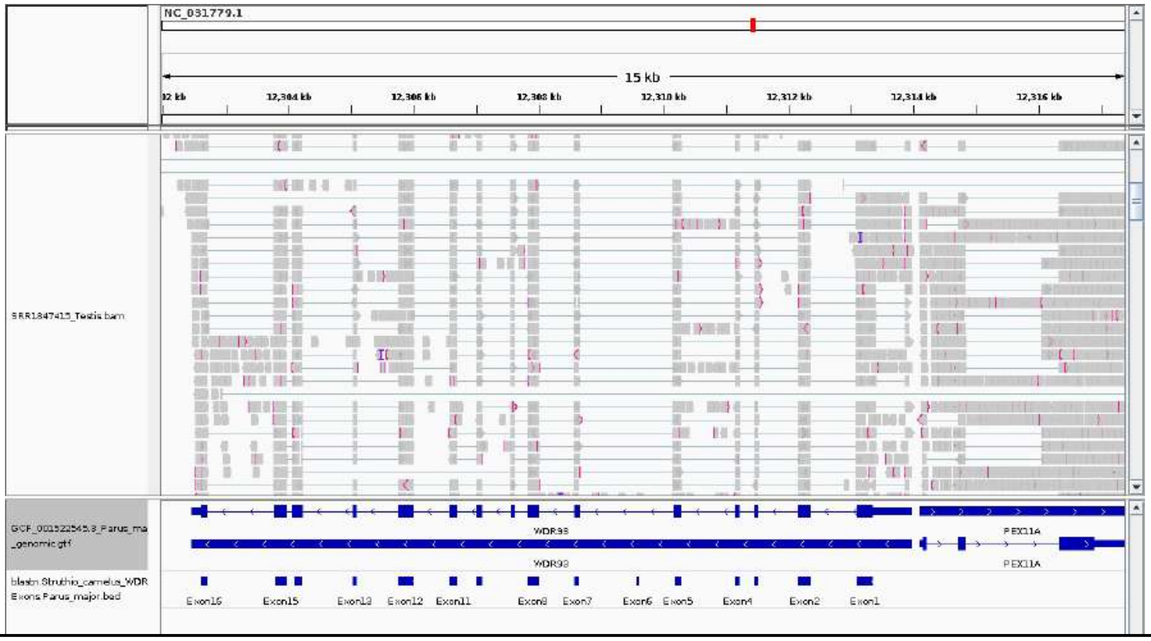


**Supplementary Figure 18: The *WDR93* gene is expressed in the testis of the great tit (*Parus major*).** IGV screenshot for *WDR93* expression for male gonad (SRR1847415) mapped to GCF_001522545.3 genome assembly of great tit (*Parus major*), using STAR mapper.

**Figure S19: Great tit (*Parus major*) *WDR93* Expressed: Sashimi plot**

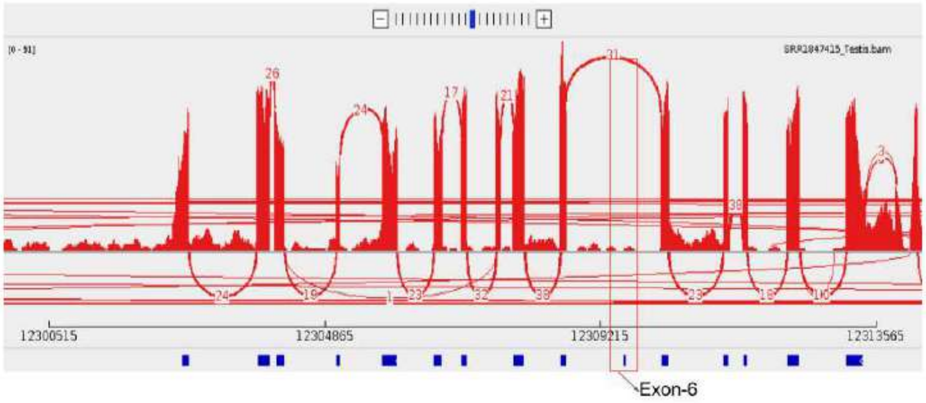


**Supplementary Figure 19:** The sashimi plot showing the splice junction at Exon boundaries of great tit (*Parus major*). Only fifteen exons are expressed, and Exon6 is not expressed, shown in a red rectangular box.

**Figure S20: Zebra finch (*Taeniopygia guttata*) *WDR93* Expressed: IGV screenshot**

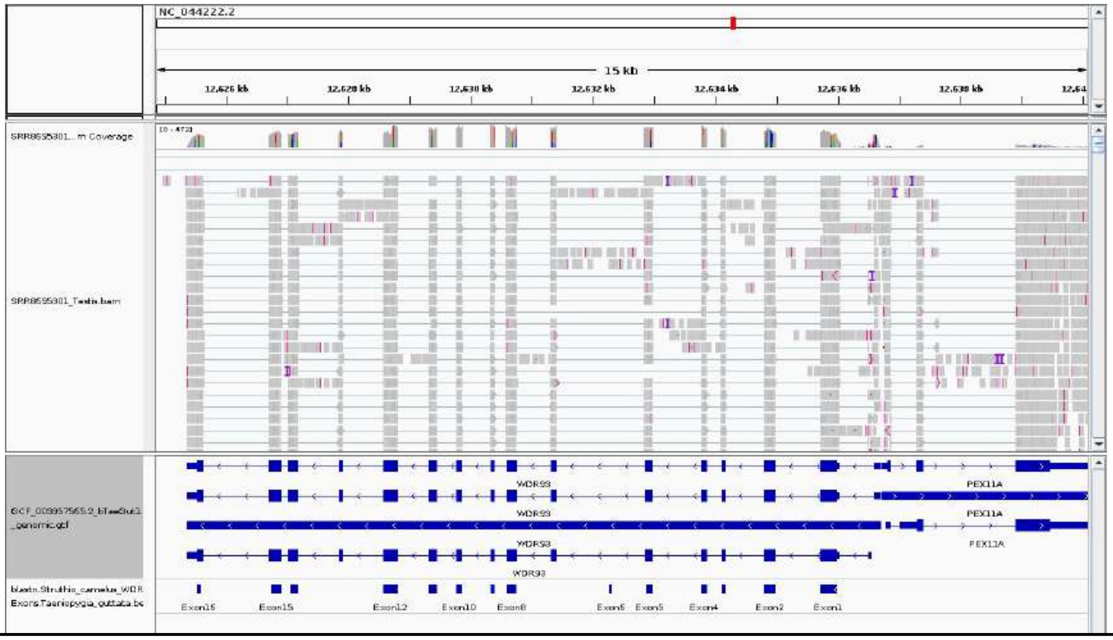


**Supplementary Figure 20: The *WDR93* gene is expressed in the testis of zebra finch (*Taeniopygia guttata*).** IGV screenshot for *WDR93* expression for male gonad (SRR8695301) mapped to GCF_003957565.2 genome assembly of the zebra finch (*Taeniopygia guttata*), using STAR mapper.

**Figure S21: Zebra finch (*Taeniopygia guttata*) *WDR93* Expressed: Sashimi plot**

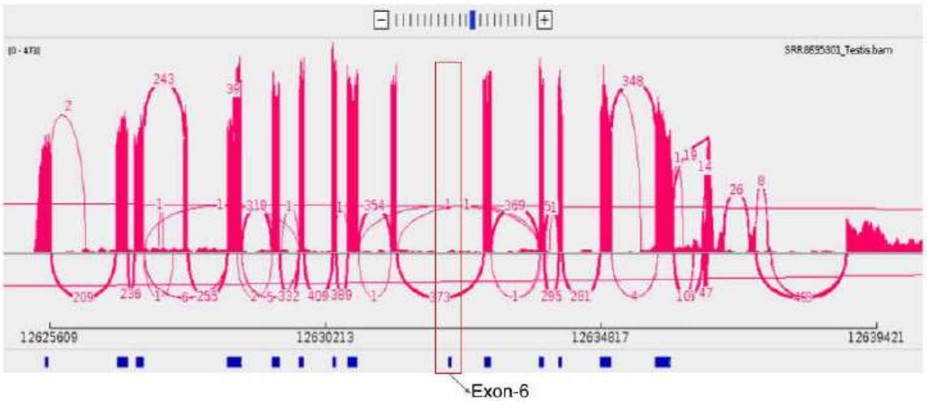


**Supplementary Figure 21:** The sashimi plot showing the splice junction at Exon boundaries of zebra finch (*Taeniopygia guttata*). Only fifteen exons are expressed, and Exon6 is not expressed, shown in a red rectangular box.

**Figure S22: *WDR93* expressed in the gonads & cerebellum, swan goose (*Anser cygnoides*) shows the noisy expression.**

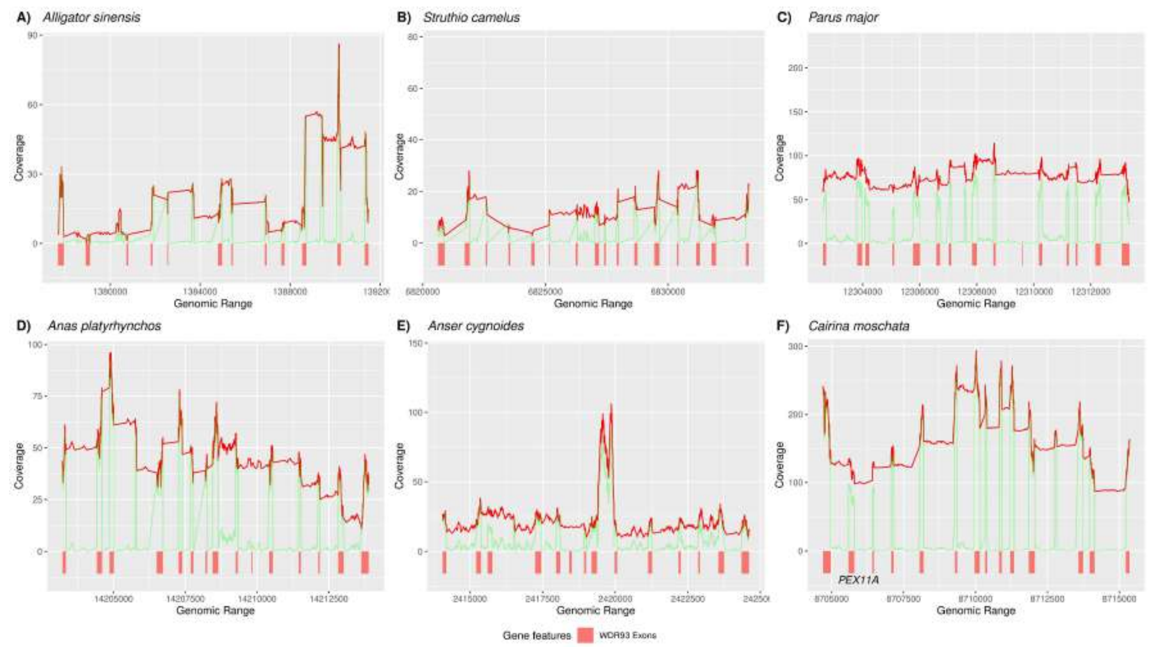


**Supplementary Figure 22: Expression status of *WDR93* at Exonic level. A)**Chinese alligator (*Alligator sinensis*), **B)** ostrich (*Struthio camelus*), **C)** great tit (*Parus major*), **D)** mallard (*Anas platyrhynchos*), **E)** swan goose (*Anser cygnoides*) and **F)** Muscovy duck (*Cairina moschata*). Overall WDR93 expression is shown by the red line, whereas the green lines represent the expression for each exon. The gonads and cerebellum tissues are used to check the expression. In Swan Goose, expression is noisy and unclear. The *WDR93* exons are shown by brick red color rectangular boxes at the bottom. The coverage is shown on the y-axis and the x-axis shows the Genomic positions.

**Figure S23: Chicken (*Gallus gallus*), turkey (*Meleagris gallopava*), and common pheasant (*Phasianus colchicus*) lack *WDR93* expression**

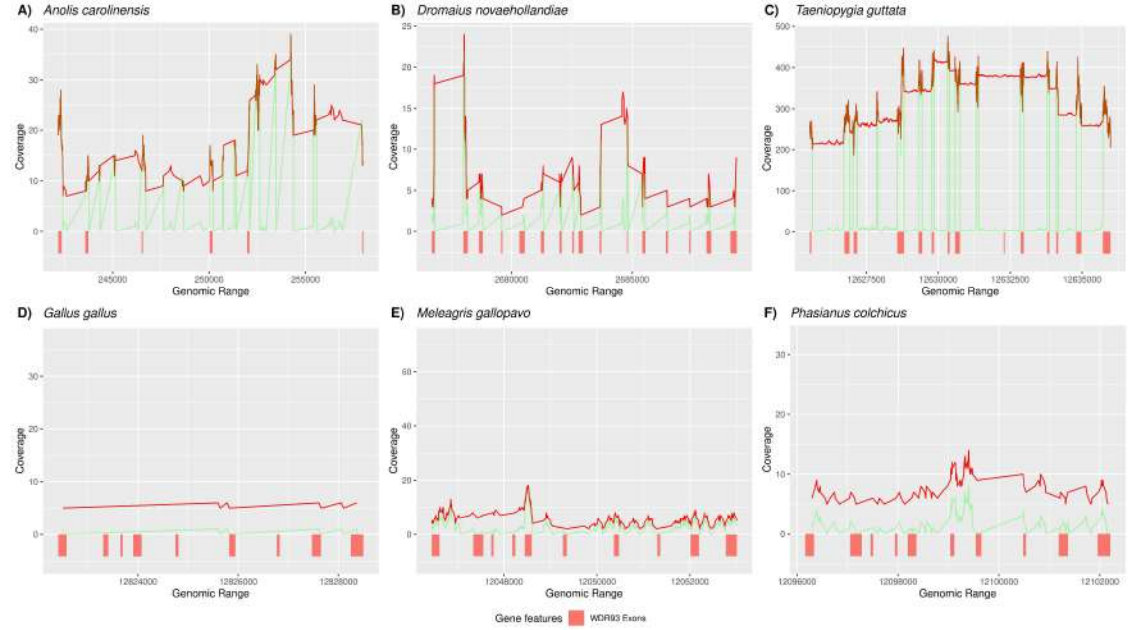


**Supplementary Figure 23: Galliformes lack the *WDR93* expression at its syntenic locus**. **A)** Green anole (*Anolis carolinensis*), **B)** Emu (*Dromaius novaehollandiae*), and **C)** Zebra finch (*Taeniopygia guttata*) with clear exon-level expression (shown in green line) and also gene level (shown in red line). However, Galliformes species such as **D)** chicken (*Gallus gallus*), E) turkey (*Meleagris gallopava*), and F) common pheasant (*Phasianus colchicus*) lack WDR93 expression. Brick-red rectangular boxes show the *WDR93* exons at the bottom. The coverage is shown on the y-axis, and the x-axis shows the Genomic positions.

**Figure S24: Helmeted guineafowl (*Numida meleagris*) lacks *WDR93* Expression: IGV screenshot**

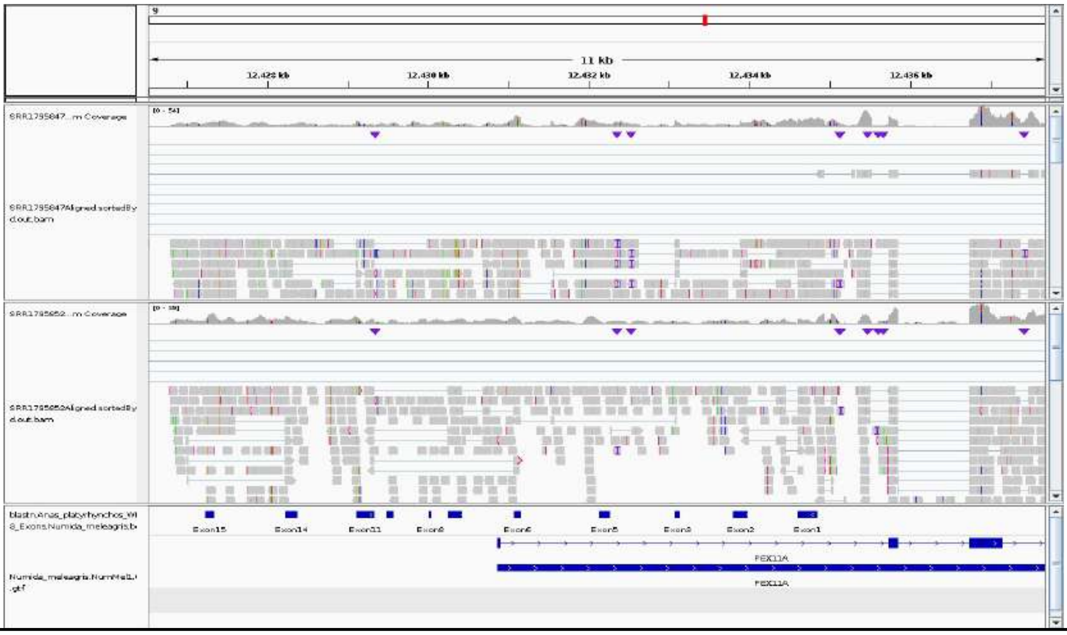


**Supplementary Figure 24:** IGV Screenshot of RNA-seq reads aligned to *WDR93* exons 1-3, 5-9, 11, 14-15 (found based on the comparison with the *WDR93* gene of Mallard) along with a part of its adjoining gene *PEX11A* in Helmeted guineafowl (*Numida meleagris*). Gray boxes indicate the reads aligned to the genome, and thin lines connecting them represent the spliced read alignments. Datasets from male gonad (SRR1795847 and SRR1795852) have been mapped to the Numida_meleagris.NumMel1.0.dna_sm.toplevel.fa genome. The bed record in the bottom row shows the locations of each of the exons (shown by blue boxes).

**Figure S25: Indian peafowl (*Pavo cristatus*) lacks *WDR93* Expression : IGV screenshot**

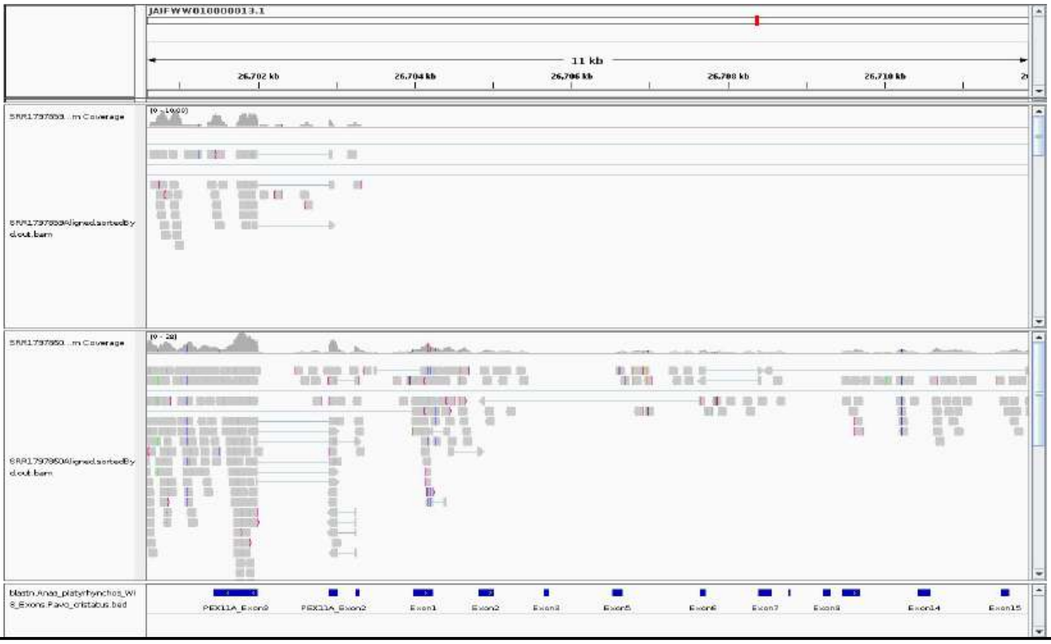


**Supplementary Figure 25:** IGV screenshot of RNA-seq reads aligned to *WDR93* exons 1-3, 5-9, 11, 14-15 (found based on the comparison with the *WDR93* gene of Mallard) along with a part of its adjoining gene *PEX11A* in Indian peafowl (*Pavo cristatus*). Gray boxes indicate the reads aligned to the genome, and thin lines connecting them represent the spliced read alignments. Datasets from the female spleen (SRR1797859) and male gonad(SRR1797860) have been mapped to the GCA_021513735.1_PavCris_1.0_genomic.fna genome. The bed record in the bottom row shows the locations of each of the exons (shown by blue boxes).

**Figure S26: Japanese quail (*Coturnix japonica*) lacks *WDR93* Expression: IGV screenshot**

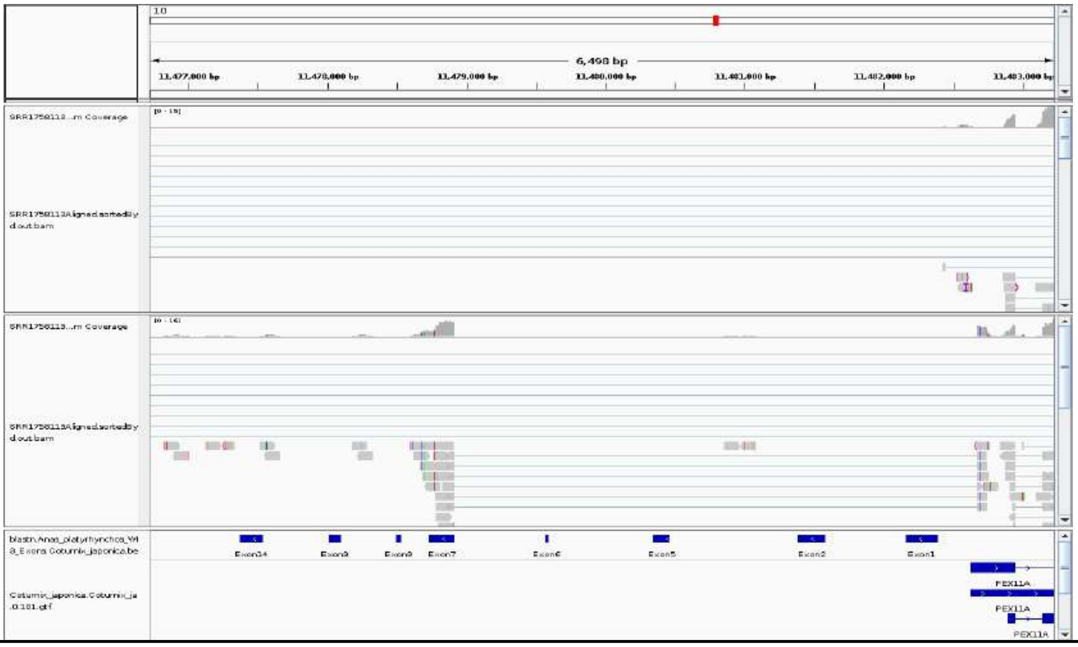


**Supplementary Figure 26:** IGV Screenshot of RNA-seq reads aligned to *WDR93* exons 1-2, 5-9, 14(found based on the comparison with the *WDR93* gene of mallard) along with a part of its adjoining gene *PEX11A* in *Coturnix japonica*. Gray boxes indicate the reads aligned to the genome, and thin lines connecting them represent the spliced read alignments. Datasets from the brain (SRR1758113) and testis (SRR1758119) have been mapped to the Coturnix_japonica.Coturnix_japonica_2.0.dna_sm.toplevel.fa genome. The bed record in the bottom row shows the locations of each of the exons (shown by blue boxes).

**Figure S27: Rifleman (*Acanthisitta chloris*) assembly verification at *WDR93* gene locus**

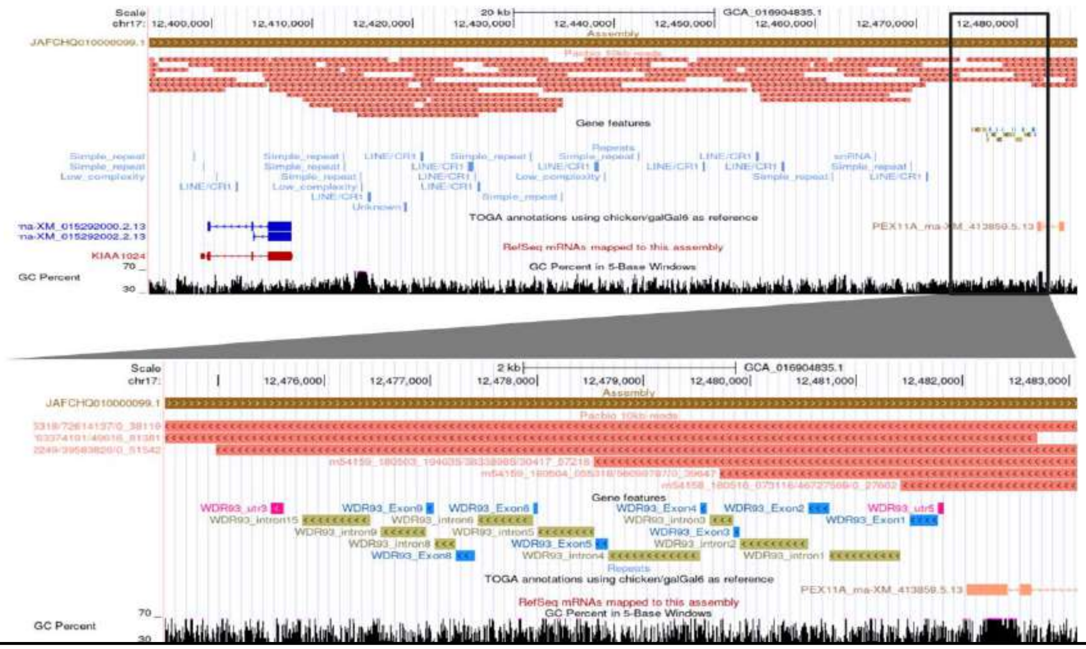


**Supplementary Figure 27: Verification of the rifleman (*Acanthisitta chloris*) genome assembly using long reads.** The UCSC genome browser image showing the alignment of ≥10 kb long reads generated using PacBio sequencing (salmon colored), aligned to the rifleman (GCA_016904835.1) at the syntenic location of the *WDR93* gene remnants. The image shown at the bottom is the zoomed-in view of the region highlighted (focusing on the *WDR93* remnants). Blue-colored boxes represent the *WDR93* exon remnants, olive-green-colored boxes show the *WDR93* introns, and the pink box represents the 5’ and 3’ UTR (shown in the genomic features track). The RepeatMasker BED track shows the repeats in that region. The accession IDs of the reads are specified on the left side of the reads beside them. Overlapping reads are found to span the entire region at the syntenic location of the *WDR93* gene, including the flanking genes (*KIAA1024/MINAR1* and *PEX11A*).

**Figure S28: Anna's hummingbird (Calypte anna) polymorphic frameshifting insertion in Exon-2**

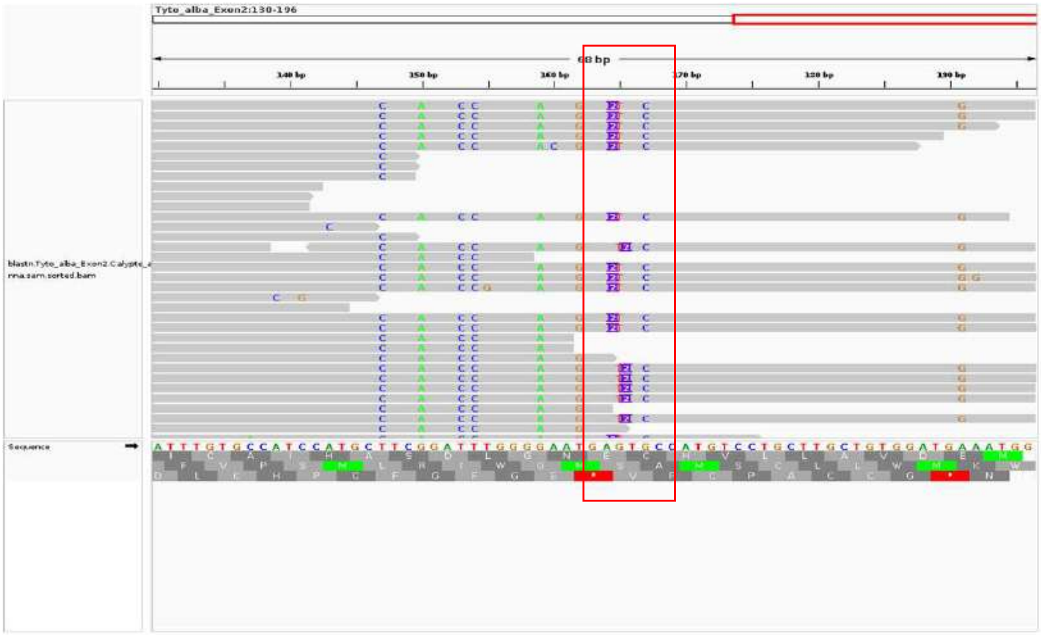


**Supplementary Figure 28: Polymorphic insertion.** The polymorphic site of two base insertions in Exon-2 of Anna's hummingbird (*Calypte anna*), compared to Ostrich (*Struthio camelus*). The grey rectangular boxes are blastn reads aligned (in bam format) to ostrich Exon2 (shown at the bottom as sequence track).

**Figure S29: Anna's hummingbird (*Calypte anna*) assembly verification at *WDR93* gene locus**

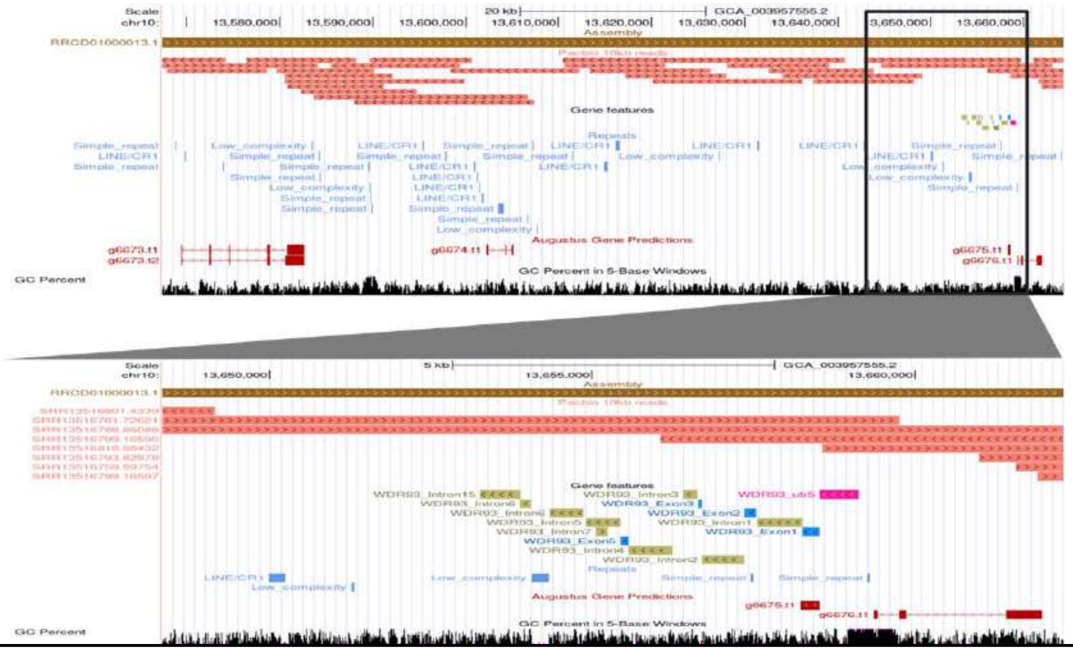


**Supplementary Figure 29: Verification of Anna's hummingbird (*Calypte anna*) genome assembly using long reads.** The UCSC genome browser image showing the alignment of ≥10 kb long reads generated using PacBio sequencing technology (salmon colored), aligned to Anna's hummingbird (GCA_003957555.2) at the syntenic location of the *WDR93* gene remnants. The image shown at the bottom is the zoomed-in view of the region highlighted (focusing on the *WDR93* remnants). Blue-colored boxes represent the *WDR93* exon remnants, olive green-colored boxes show the *WDR93* introns and the pink box represents the 5' UTR (shown in the genomic features track). The RepeatMasker BED track shows the repeats in that region. The accession IDs of the reads are specified on the left side of the reads beside them. Overlapping reads are found to span the entire region at the syntenic location of the *WDR93* gene, including the flanking genes (*KIAA1024/MINAR1* and *PEX11A*).

**Figure S30: Mallard (*Anas platyrhynchos*) assembly verification at *CFAP46* gene locus**

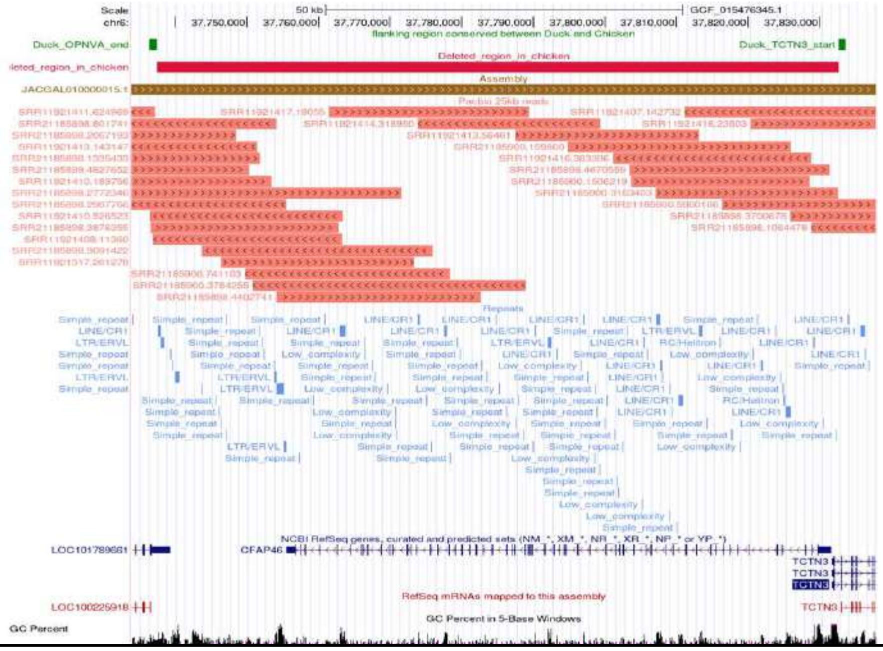


**Supplementary Figure 30: Verification of the mallard (*Anas platyrhynchos*) genome assembly at CFAP46 using long reads.** The UCSC genome browser image shows the alignment of ≥25 kb long reads generated using PacBio sequencing technology (salmon colored), aligned to the mallard (GCF_015476345.1) at the syntenic location of the *CFAP46* gene. The brick-red box labeled with the "deleted region in chicken" shows the approximate deleted region, whereas the green boxes at the top show the end of *OPNVA* and *TCTN3* start. The RepeatMasker BED track shows the repeats in that region(indicated by blue boxes). The accession IDs of the reads are specified on the left side of the reads beside them. Overlapping reads span the entire region at the syntenic location of the *CFAP46* gene, including the flanking genes (*OPNVA* and *TCTN3*).

**Figure S31: Chicken (*Gallus gallus*) assembly verification at *CFAP46* gene locus**

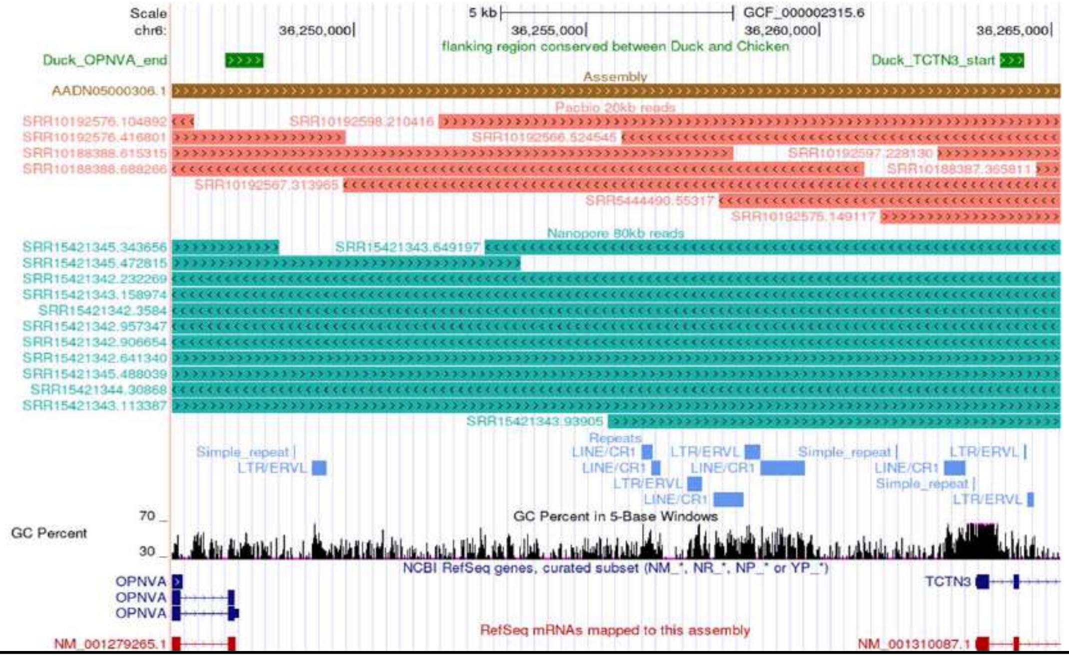


**Supplementary Figure 31: Verification of the Chicken (*Gallus gallus*) genome assembly using long reads.** The UCSC genome browser image shows the alignment of ≥20 kb long reads generated using PacBio sequencing (salmon colored) and cyan blue showing the ≥80 kb long reads generated using Nanopore sequencing, aligned to the chicken (GRCg6a) at the syntenic location, which *OPNVA* and *TCTN3* flank. The RepeatMasker BED track shows the repeats in that region (blue boxes). The accession IDs of the reads are specified on the left side of the reads beside them. Overlapping reads are found to span the entire region of flanking genes (*OPNVA* and *TCTN3*).

**Figure S32: Deletion size in comparison to mallard (*Anas platyrhynchos*)-with merge**

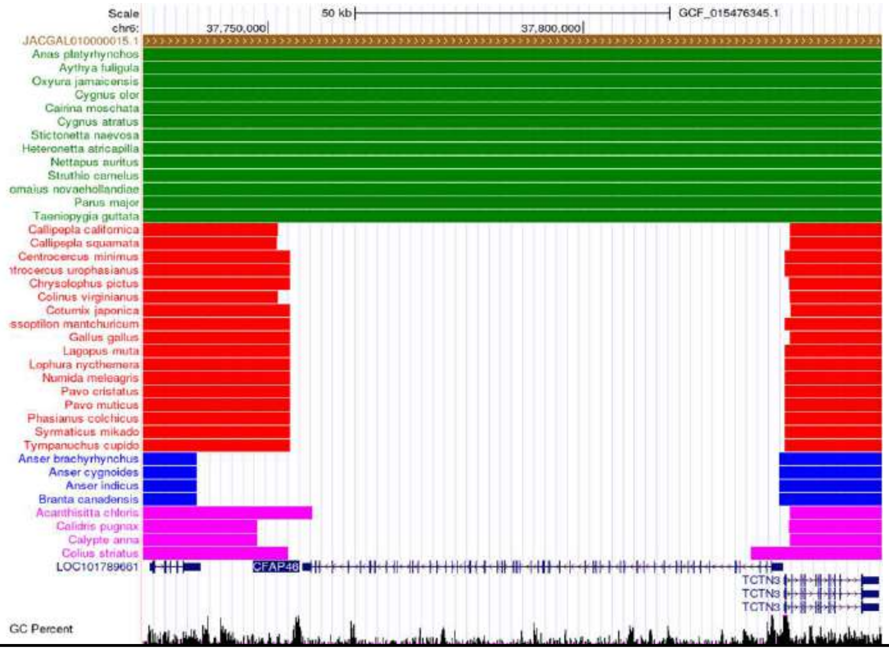


**Supplementary Figure 32: Deletion size comparison of *CFAP46* gene-containing segment across the six phylogenetically distant independent species with mallard (*Anas platyrhynchos*).** The species with green boxes have an intact *CFAP46* gene and do not show the deletion. In comparison, the species with red color are Galliformes with *CFAP46* deletion. The blue colored boxes show the deletion in geese species. The pink-colored boxes show deletion in rifleman (*Acanthisitta chloris*), ruff (*Calidris pugnax*), Anna's hummingbird (*Calypte anna*), and speckled mousebird (*Colius striatus*). For deletion size estimation, the blastn of shown species was performed with mallard repeat masked chromosome and blast results merged with bedtools merge with –d 30000 (the deleted region is longer than 30000 bp as identified by pairwise alignment of chicken and duck genomes).

**Figure S33: Deletion size in comparison to mallard (*Anas platyrhynchos*)-without merge**

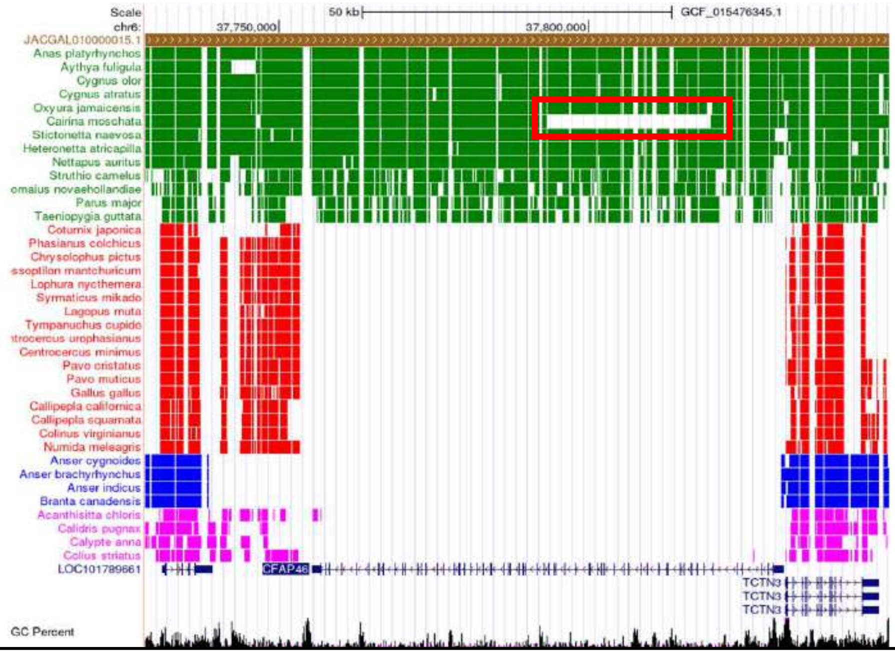


**Supplementary Figure 33: Deletion size comparison of *CFAP46* gene containing segment across the six phylogenetically distant independent species with mallard (*Anas platyrhynchos*) without merging blastn**. The species with green boxes have an intact *CFAP46* gene and do not show the deletion. In comparison, the species with red color are Galliformes which has *CFAP46* deletion. The blue colored boxes show the deletion in geese species. The pink-colored boxes show deletion in rifleman (*Acanthisitta chloris*), ruff (*Calidris pugnax*), Anna's hummingbird (*Calypte anna*), and speckled mousebird (*Colius striatus*). The large white space (shown in a red rectangular box) in the Muscovy duck (*Cairina moschata*) is due to the presence of a gap (N's) in the genomic sequence and is not a deletion.

**Figure S34: Swan goose (*Anser cygnoides*) assembly verification at *CFAP46* gene locus**

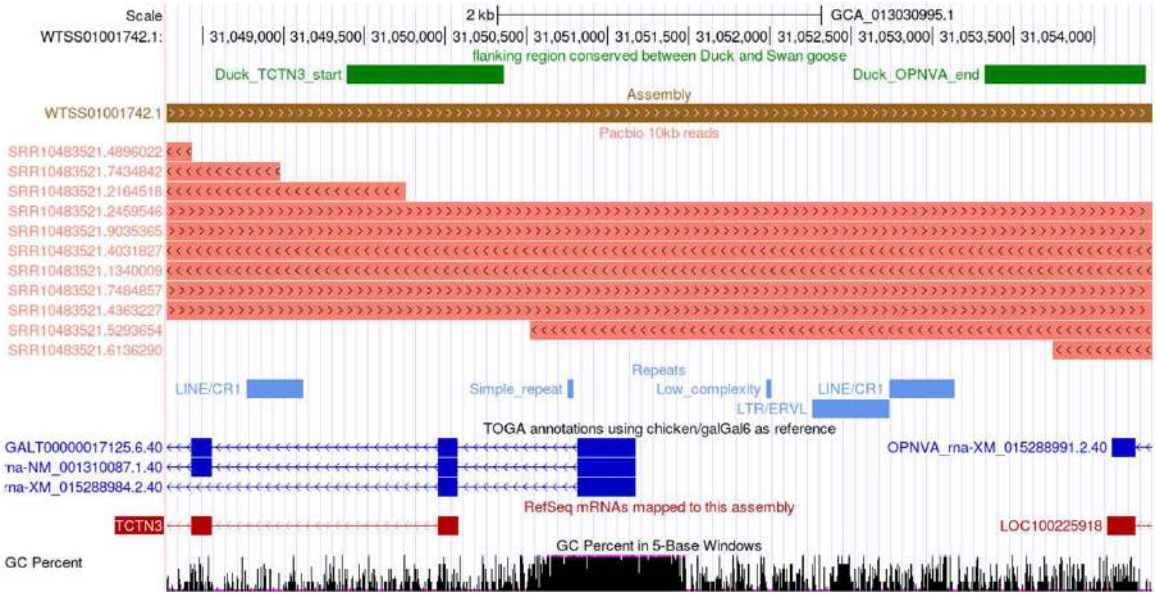


**Supplementary Figure 34: Verification of the swan goose (*Anser cygnoides*) genome assembly at *CFAP46* using long reads.** The UCSC genome browser image showing the alignment of ≥10 kb long reads generated using PacBio sequencing technology (salmon coloured), aligned to the swan goose (GCA_013030995.1) at the syntenic location which is flanked by *OPNVA* and *TCTN3* . The RepeatMasker BED track shows the repeats in that region (blue boxes). The accession IDs of the reads are specified on the left side of the reads beside them. Overlapping reads are found to span the entire region of flanking genes (*OPNVA* and *TCTN3*; shown at bottom and start-end sequence positions by green with track name "flanking region conserved between Duck and swan goose").

**Figure S35: Rifleman (*Acanthisitta chloris*) assembly verification at *CFAP46* gene locus**

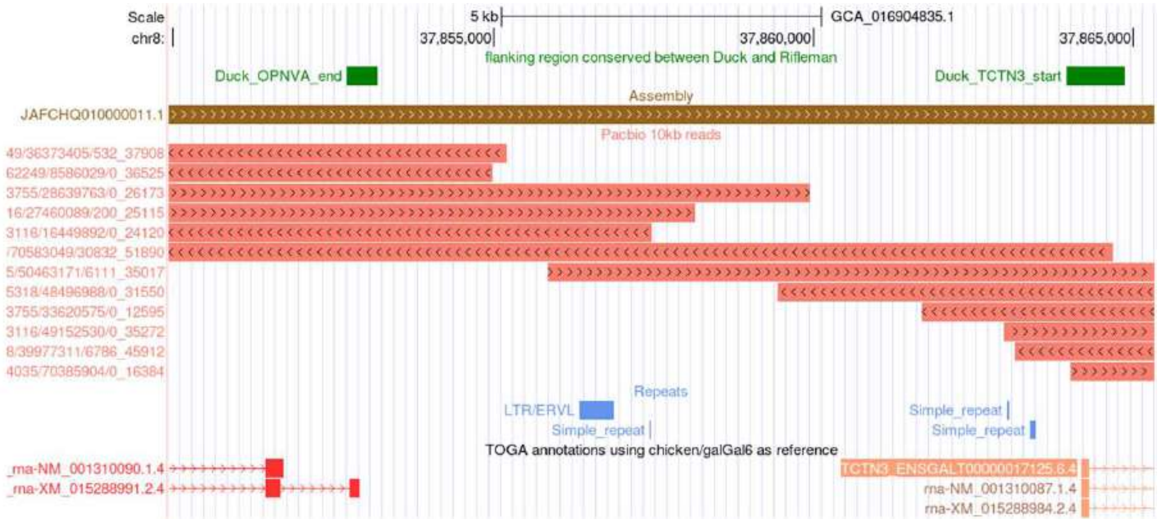


**Supplementary Figure 35: Verification of the Rifleman (*Acanthisitta chloris*) genome assembly at *CFAP46* using long reads.** The UCSC genome browser image showing the alignment of ≥10 kb long reads generated using PacBio sequencing technology (salmon coloured), aligned to the rifleman (GCA_016904835.1) at the syntenic location which is flanked by *OPNVA* and *TCTN3* . The RepeatMasker BED track shows the repeats in that region (blue boxes). The accession IDs of the reads are specified on the left side of the reads beside them. Overlapping reads are found to span the entire region of flanking genes (*OPNVA* and *TCTN3*; shown at bottom and start-end sequence positions by green with track name “flanking region conserved between Duck and Rifleman”).

**Figure S36: Anna's hummingbird (*Calypte anna*) assembly verification at *CFAP46* gene locus**

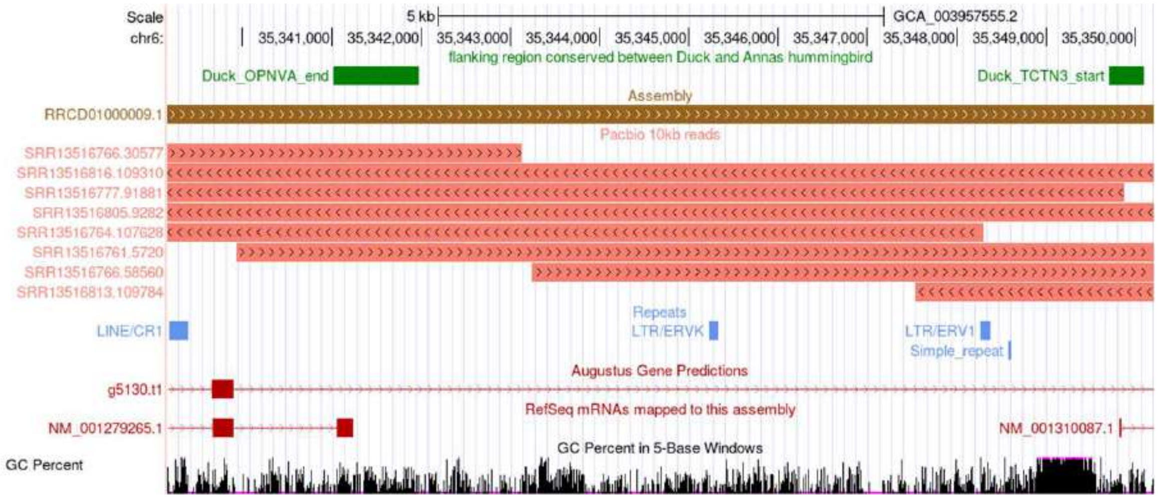


**Supplementary Figure 36: Verification of Anna's hummingbird (*Calypte anna*) genome assembly at *CFAP46* using long reads.** The UCSC genome browser image showing the alignment of ≥10 kb long reads generated using PacBio sequencing (salmon colored), aligned to Anna's hummingbirds (GCA_003957555.2) at the *CFAP46* syntenic location, which is flanked by OPNVA and TCTN3. The RepeatMasker BED track shows the repeats in that region (blue boxes). The accession IDs of the reads are specified on the left side of the reads beside them. Overlapping reads are found to span the entire region of flanking genes (*OPNVA* and *TCTN3*; shown at bottom and start-end sequence positions by green with track name "flanking region conserved between Duck and Anna's hummingbird").

**Figure S37: Graphical abstract**

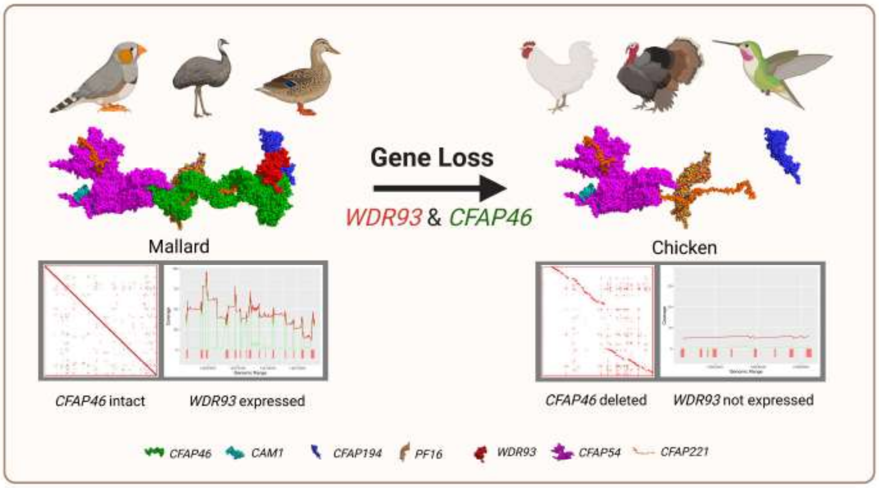


**Supplementary Figure 37: Graphical abstract summarizing the main findings.** At the top, the species with intact *WDR93* and *CFAP46* are shown on the left, whereas those with loss are on the right. In the middle, the cartoon representation of C1d projection is made from PDB 7N6G (*Chlamydomonas reinhardtii*) by keeping (on the left) and hiding (on the right) the lost genes in the pymol. The third row compares *CFAP46* deletion and expression status of *WDR93* in mallard and chicken. The last row shows the protein components of the C1d projection of *Chlamydomonas reinhardtii*.
